# Applying standardised methods to assess the potential risks of alien freshwater crayfish introductions to South Africa

**DOI:** 10.1101/2025.11.12.687953

**Authors:** Lee-Anne Botha, Christian T. Chimimba, Tsungai A. Zengeya

## Abstract

Alien freshwater crayfish have been widely introduced worldwide for aquaculture and the pet trade. Despite providing societal benefits, crayfish introductions are also known to cause adverse impacts in areas of introduction. This study used the recently developed Risk Analysis for Alien Taxa (RAAT) framework to assess the risk associated with 14 alien freshwater crayfish introductions in South Africa. Thirteen of the 14 species were assessed as high risk because they are likely to be introduced to South Africa and have the potential to cause major environmental impacts. Eight of these species are listed under the South African National Environmental Management: Biodiversity Act (NEM: BA) Alien and Invasive Species (A&IS) Regulations, which implies there is an obligation to manage them. This notion was supported by the recommendations from the risk analyses for *Cherax cainii*, *C. tenuimanus*, and *C. quadricarinatus*. The two marron species (*C. cainii* and *C. tenuimanus*) have no known naturalised populations and are likely confined to aquaculture facilities. *Cherax quadricarinatus* is already widespread and control methods should focus on minimising spread, as eradication is no longer feasible. The recommendations from some of the risk analyses do not agree with the current listing under the A&IS Regulations. *Procambarus clarkii* is already known to be invasive but is not listed and there is anecdotal evidence that *C. destructor*, *Faxonius limosus*, *F. rusticus*, *Pacifastacus leniusculus*, and *Pontastacus leptodactylus* are present in the pet trade and their presence needs to be verified. Crayfish can become conflict-generating alien species because they have both major negative impacts and socio-economic benefits. Therefore, the recommended management options aim to preserve benefits while limiting negative impacts by restricting the importation of high-risk species for new introductions and implementing interventions to prevent spread and minimise impacts for established introductions.

## Introduction

Biological invasions are a significant global problem that seems to be growing despite some concerted efforts to develop and implement measures to either halt or minimise their impacts (Essl et al. 2020; Pyšek et al. 2020; IPBES 2023). The primary reason for the alien species introductions has been to meet societal needs such as the provision for food, raw materials such as timber, ornamental horticulture species and the pet trade (Hulme et al. 2008; Bernery et al. 2022). Some of these introduced species have become invasive and have been implicated in causing adverse effects on biodiversity, ecosystem functioning, and human livelihood and health (IPBES 2023). This has necessitated the need for risk analysis protocols that restrict the importation of high-risk alien species while allowing for the introduction and utilisation of low-risk species (e.g., Roy et al. 2018). This is especially pertinent as the number of established alien species has increased worldwide during the past two centuries, mainly because of increased connectivity through increased travel and trade (Seebens et al. 2017).

Freshwater crayfish are a diverse group of decapods with over 600 species that naturally occur on all continents except continental Africa and Antarctica (Crandall & Budhay 2008). Despite their global distribution, crayfish are among some of the most widely introduced freshwater invertebrates around the world, mainly through aquaculture and the aquarium trade (Crandall & Buhay 2008; Chucholl 2013; Patoka et al. 2014; Faulkes 2015; FAO 2022; Olden & Carvalho 2024). While aquaculture and aquarium trade provide positive impacts, crayfish introductions are also known to have negative environmental and non-environmental impacts (Lodge et al. 2012; Souty-Grosset et al. 2016; Madzivanzira et al. 2020). Crayfish invasions have been implicated in causing adverse environmental impacts such as hybridisation with native species (e.g., Perry et al. 2001), extirpation of native species through competition and predation (e.g., Dunn et al. 2009), and/or disease transmission (e.g., Longshaw 2011; Chucholl & Schimpf 2016), modifying aquatic-terrestrial linkages (e.g., Dana et al. 2010), altering food-web structure (Jackson et al. 2016; Zengeya et al. 2022), and disrupting nutrient cycles (Eby et al. 2006). They are also known to cause non-environmental impacts that affect human livelihoods and well-being (Kouba et al. 2022). For example, crayfish invasions have been implicated in disrupting recreational activities (Keller et al. 2008), their feeding and burrowing activities impact aquaculture and crop production (Souty-Grosset et al. 2016), and damage fishing equipment (Gherardi et al. 2011b; Madzivanzira et al. 2023).

Eight crayfish species are presumed to be present in South Africa based on historical introduction records (Figure 1). However, there is uncertainty on the occurrence and identity of some of the species (De Moor 2002; Nunes et al. 2017a; Wilson & Kumschick 2024). There is evidence that redclaw crayfish (*Cherax quadricarinatus*) and red swamp crayfish (*Procambarus clarkii*) are present in South Africa, as they have established populations in river systems across several provinces (De Moor 2002; Du Preez & Smit 2013; Nunes et al. 2017b; Nunes et al. 2017c; Barkhuizen et al. 2022; CapeNature unpublished data).

**Figure 1.**
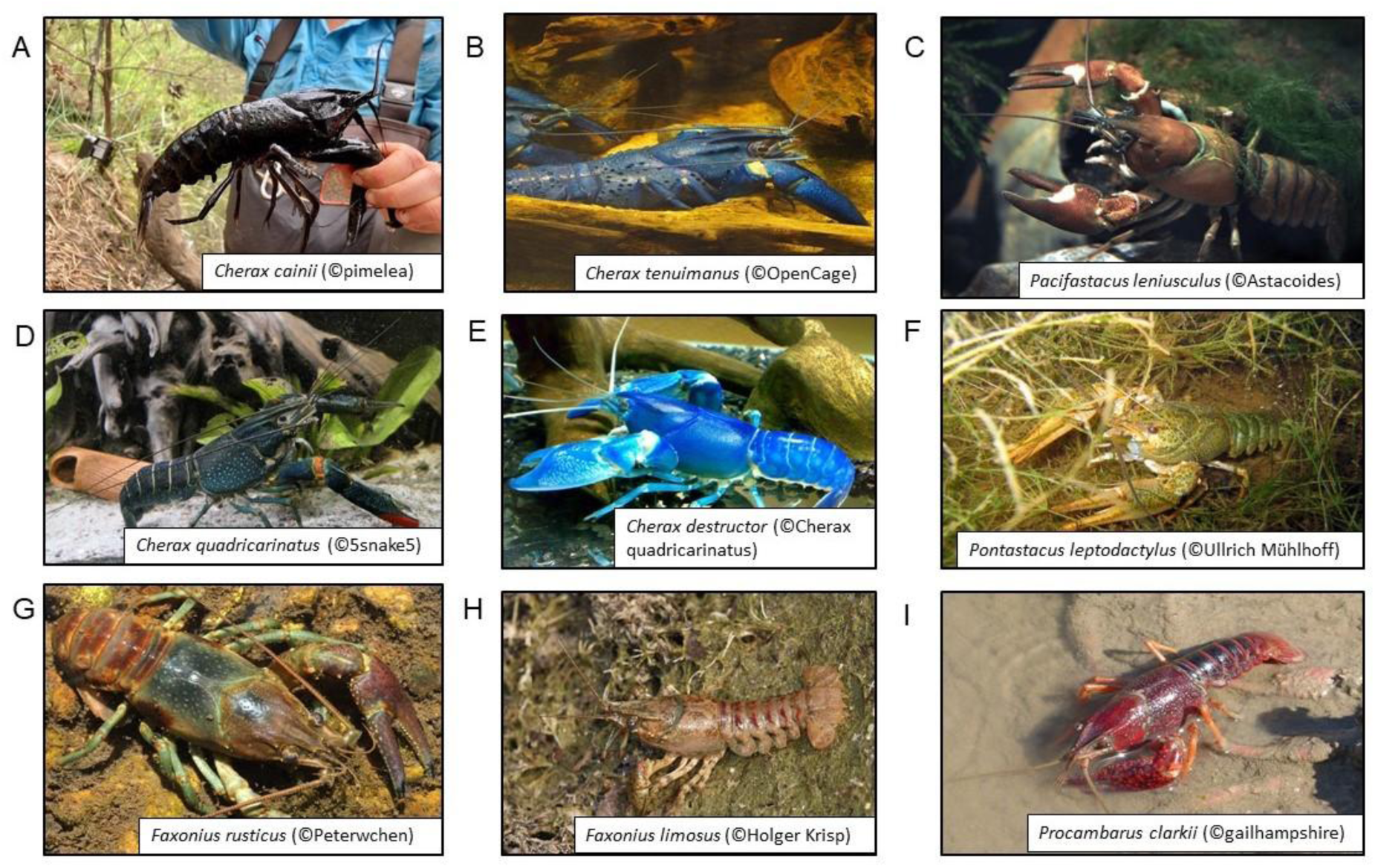
Nine crayfish species that are presumed to be present in South Africa. All of the images were obtained from https://commons.wikimedia.org/wiki/Main_Page.

There is, however, equivocal evidence that the other six species are present in South Africa. Two marron species (*Cherax cainii* and C. *tenuimanus*) were introduced for aquaculture but it is uncertain if both species are present in South Africa (Wilson & Kumschick 2024). A 2002 taxonomic revision of *C. tenuimanus* indicated that the species was not homogenous but instead consisted of two genetically distinct species, *C. tenuimanus* and *C. cainii* (Austin & Ryan 2002; Duffy et al. 2014). As a result, introduction records of marron in South Africa prior to 2002 refer to *C. tenuimanus* but recent import permit records indicate that both species are likely present in the country (Wilson & Kumschick 2024). *Cherax destructor* was introduced into South Africa in 1988 as a possible candidate species for aquaculture, but it is uncertain if it is still present in cultivation or in the wild (Nunes et al. 2017a). There is also anecdotal evidence that several crayfish species might be present in South Africa through the pet trade. These include the spiny-cheek crayfish (*Faxonius limosus*), rusty crayfish (*Faxonius rusticus*), narrow-clawed crayfish (*Pontastacus leptodactylus*) and signal crayfish (*Pacifastacus leniusculus*).

In South Africa, biological invasions are managed through the Alien and Invasive Species Regulations (RSA 2020) of its National Environmental Management: Biodiversity Act (NEM:BA) (Act no. 10 of 2004) (RSA 2004). The rationale behind the A&IS Regulations is to restrict the importation of high-risk alien species, reduce the number of alien species becoming invasive, and limit the extent and impact of well-established invaders, while allowing for the utilisation of low-risk alien species (Wilson & Kumschick 2024). The implementation of the A&IS Regulations is underpinned by a South African-developed risk analysis framework, termed Risk Analysis for Alien Taxa (RAAT) that outlines a normative process to assess an alien taxon’s likelihood of invasion, realised and potential impacts, and options for management (Kumschick et al. 2020a). The RAAT framework is an objective, transparent and consultative process that ensures that it is clear what the risks posed by the regulated species are and what can be done to mitigate or prevent impacts (Kumschick et al. 2020b). This paper applied the RAAT framework to evaluate the risk posed by the introduction of freshwater crayfish in South Africa. It provides an update on the status of alien crayfish species that are known to occur in South Africa, and recommendations on appropriate management interventions. It also identified alien freshwater crayfish that have a global invasion history and have the potential to become invasive if introduced into South Africa, and provides policy decision-makers with information both on the risks posed and on what can be done to mitigate or prevent impacts.

## Methods

### Species selection

A database with all crayfish species was compiled from the primary literature, the International Union for Conservation of Nature (IUCN) Red List (www.iucnredlist.org/), and the Global Biodiversity Information Facility (GBIF) (www.gbif.org/en/). The invasion status of the crayfish species was quantified based on data from the primary literature, the Centre for Agriculture and Bioscience International (CABI) invasive species compendium (www.cabi.org/isc), the Global Invasive Species Database (GISD) (www.iucngisd.org/gisd/), and the Non-Indigenous Aquatic Species (NAS) dataset (www.nas.er.usgs.gov/). Invasion status was defined according to the different stages of the unified framework (see Blackburn et al. 2011), and species were grouped in four broad categories: 1) not introduced = species that have no record of introduction to areas outside their native range; 2) introduced = species that are introduced to a country but are not naturalised in the wild; 3) established = species that have established in the wild but are not yet invasive; and 4) invasive = species with self-sustaining populations that have spread from initial sites of introduction.

### Impact assessments

Environmental impacts were assessed using the Environmental Impact Classification for Alien Taxa (EICAT; Blackburn et al. 2014; Hawkins et al. 2015) and non-environmental impact using the Socio-Economic Classification for Alien Taxa (SEICAT; Bacher et al. 2017). For both the EICAT and SEICAT assessments an extensive literature review of impact studies was undertaken using Google Scholar (https://scholar.google.co.za/) and Web of Science (http://apps.webofknowledge.com) search engines. The search thread *\*invasive*crayfish\** or *\*impacts*species name\** were used. The relevant literature was compiled, and recorded impacts were then assessed and classified using procedures outlined for EICAT and SEICAT. The EICAT assessment classified environmental impacts across 12 different mechanisms and assigned a magnitude score across five impact levels [Minimal Concern (MC), Minor (MN), Moderate (MO), Major (MR) and Massive (MV)] (Blackburn et al. 2014; Hawkins et al. 2015). The SEICAT was used to assess the impact alien freshwater crayfish have on human well-being using four impact categories (safety, material or immaterial assets, health, and social, spiritual or cultural) and the magnitude of the impacts were assessed across five levels that are similar to the EICAT assessment (Bacher et al. 2017).

### Risk analysis

The risks associated with crayfish introduction were assessed using the RAAT framework that consists of the following four components: 1) risk identification; 2) risk assessment; 3) risk management; and 4) risk communication (Kumschick et al. 2020a).

### Risk identification

Biological invasions present various risks that can be broadly grouped in terms of species, pathways and areas, and in this study, the risks associated with biological invasions from alien crayfish were identified in terms of species (Kumschick et al. 2020b).

### Risk assessment

This step evaluated the likelihood of a particular crayfish species being introduced, establishing and spreading in South Africa, and the consequences (negative impacts) thereof (Kumschick et al. 2020a). Information on environmental and non-environmental impacts were derived from the EICAT and SEICAT assessments, and in cases where a species had no documented impacts, it was classified as Data Deficient (DD); potential impacts were inferred from a closely related species (Kumschick et al. 2020a). Non-environmental impact assessments focus on human well-being and how the alien species affects related issues such as livelihoods, farming practices and recreational activities (e.g., Westman et al. 2002; Laverty et al. 2015). The risk score was calculated using the outcomes of the assessment of: 1) likelihood of introduction; 2) establishment and spread; and 3) potential to cause negative impacts (consequences) (Kumschick et al. 2020a).

### Risk management

This step included the evaluation of the best management options for the freshwater crayfish species that are known to be present in South Africa to mitigate spread and impacts, while allowing utilisation (Kumschick et al. 2020a). In South Africa, alien taxa are managed under A&IS Regulations, which comprise of lists (i.e., notices) for regulated species and the management and control option for each listed species (Van Wilgen & Wilson 2018). The management options are grouped into four categories: 1) Category 1a – species that should be eradicated; 2) Category 1b *–* species that should be controlled as part of national programmes, and cannot be traded or allowed to spread; 3) Category 2 – species that have the same restrictions as Category 1b species but a permit can be issued to allow utilisation under specific conditions that aim to prevent spread and minimise impacts; and 4) Category 3 – species that can be utilised without a permit but they cannot be traded or further propagated and should be controlled when they occur in biodiversity-sensitive areas such as protected areas or riparian zones (Kumschick et al. 2020b). These regulation categories apply only to species that are already present in the country. Permits are required for new introductions into the country, and these are only considered if supported by a risk analysis (Kumschick et al. 2020b). Possible management interventions were then evaluated and allocated a score (low, medium and high) based on ease of management score (Kumschick et al. 2020a).

### Risk communication and recommendations

This included the collation and summary of the complete background information of the RAAT framework process to make recommendations for management, regulations and engagement with relevant stakeholders (Kumschick et al. 2020a). The impact assessments were done at a global scale, whereas for the risk analysis, the area of interest was South Africa.

The risk analysis reports were prepared for submission to the South African Alien Species Risk Analysis Review Panel (ASRARP), a committee that is tasked with reviewing risk analyses attached to import applications and listing of species under national legislation to ensure they are scientifically robust and consider the best available evidence (Kumschick et al. 2020b). ASRARP is an independent body coordinated by the South African Biodiversity Institute (SANBI) as the secretariat, and its members consist of scientists and taxon experts working on various issues in biological invasions. Risk analysis reports submitted to ASRARP are peer-reviewed before approval, following a process like that used by peer-reviewed journals (Figure 2). The ASRARP committee then provides recommendations to the Risk Analysis Review Committee (RARC), an interdepartmental panel set up by the South African Department of Forestry, Fisheries, and the Environment (DFFE) that is tasked with granting import permits and approving changes to regulations on biological invasions. Seven risk analysis reports included in this paper have already been submitted and approved by ASRARP. These include reports for *Cherax cainii*, *C. tenuimanus, C. destructor*, *Faxonius rusticus*, *Procambarus clarkii, Pacifastacus leniusculus* and *Pontastacus leptodactylus*. These species were prioritised because they are either listed under the A&IS Regulations and or are now invasive in the country. Risk analysis reports for *Faxonius immunis, F. juvenilis, F. limosus, F. virilis, Procambarus acutus* and *P. virginalis* are in the process of being submitted to ASRARP.

**Figure 2.**
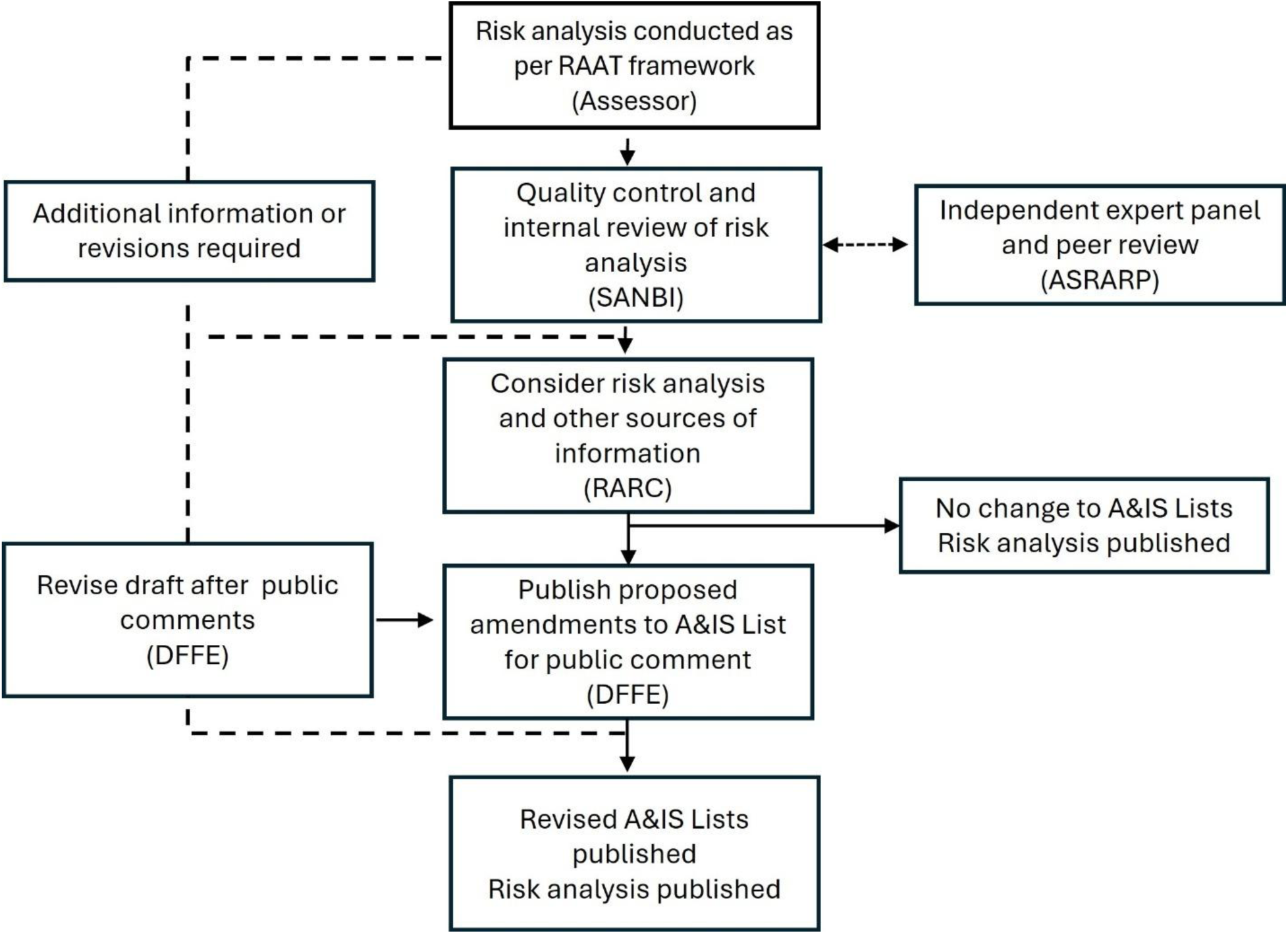
The process of how risk analysis reports are prepared, reviewed and used to inform the listing of alien species on the Alien and Invasive Species Regulations in South Africa. SANBI = South African National Biodiversity Institute; ASRARP = Alien Species Risk Analysis Review Panel; RARC = Risk Analysis Review Committee; DFFE = South African Department of Forestry, Fisheries, and the Environment. Figure adapted from Wilson and Kumschick (2024).

## Results

### Species selection

The introduction status of 658 crayfish species was assessed and 14 crayfish species that have been introduced to areas outside their natural range, where they have either become established and or invasive, were then selected for the impact assessments and risk analyses (Table 1). These included five *Faxonius* species: calico crayfish (*F. immunis*), Kentucky River crayfish (*F. juvenilis*), spiny-cheek crayfish (*F. limosus*), rusty crayfish (*F. rusticus*) and virile crayfish (*F. virilis*); two *Procambarus* species: White River crayfish (*P. acutus*) and red swamp crayfish (*P. clarkii*); and signal crayfish (*Pacifastacus leniusculus*) that are native to North America; narrow clawed crayfish (*Pontastacus leptodactylus*), which is native to Europe; and four *Cherax* species: smooth marron (*C. cainii*), hairy marron (*C. tenuimanus*), yabby crayfish (*C. destructor*) and redclaw crayfish (*C. quadricarinatus*) that are native to Australia. The marmorkrebs crayfish (*Procambarus virginalis*) has an unknown native distribution.

**Table 1.** A summary of the risk analysis results for 14 alien crayfish species that are known to be invasive or have been introduced in areas outside their native range. Invasion status was assessed for a species’ global introduced range based on the unified framework (Blackburn et al. 2011) and was grouped in three level descriptors: 1) introduced = species that are introduced to a country but are not naturalised in the wild; 2) established = species that have established in the wild but are not yet invasive; and 3) invasive = species with self-sustaining populations that have spread from initial sites of introduction. The current regulatory listing [based on the South African National Environmental Management: Biodiversity Act (NEM:BA) of 2020 Alien and Invasive Species (A&IS) Lists] and the listing recommended in the risk analyses (Supplement 1.1–1.14) are also included to indicate where change of listing has been proposed.

| Species | Native region | Global invasion status | Likelihood | Consequence | Risk | Ease of management | Current listing | Recommended listing |
| --- | --- | --- | --- | --- | --- | --- | --- | --- |
| Spiny-cheek crayfish (*Faxonius limosus*) | North America | Invasive | Very unlikely | Major | High | Medium | 1a | 1a<sup>#</sup> |
| Rusty crayfish (*Faxonius rusticus*) | North America | Invasive | Fairly probable | Major | High | Medium | 1a | 1a<sup>#</sup> |
| Yabby (*Cherax destructor*) | Australia | Established | Fairly probable | Major | High | Difficult | 1a | 1a<sup>#</sup> |
| Narrow-clawed crayfish (*Pontastacus leptodactylus*) | Europe | Invasive | Very unlikely | Major | High | Medium | 1a | 1a<sup>#</sup> |
| Signal crayfish (*Pacifastacus leniusculus*) | North America | Invasive | Probable | Major | High | Difficult | 1a | Prohibited list<sup>#</sup> |
| Redclaw crayfish (*Cherax quadricarinatus*) | Australia | Invasive | Probable | Major | High | Difficult | 1b | 1b |
| Red swamp crayfish (*Procambarus clarkii*) | North America | Invasive | Probable | Major | High | Difficult | Not listed | 1b |
| Smooth marron (*Cherax cainii*) | Australia | Introduced | Fairly probable | Major | High | Medium | 2 | 2 |
| Hairy marron (*Cherax tenuimanus*) | Australia | Introduced | Fairly probable | Major | High | Medium | 2 | 2<sup>#</sup> |
| Calico crayfish (*Faxonius immunis*) | North America | Introduced | Unlikely | Major | High | Medium | Not listed | Prohibited list<sup>#</sup> |
| Kentucky River crayfish (*Faxonius juvenilis*) | North America | Established | Very unlikely | Major | High | Medium | Not listed | Prohibited list<sup>#</sup> |
| Virile crayfish (*Faxonius virilis*) | North America | Invasive | Fairly probable | Major | High | Difficult | Not listed | Prohibited list<sup>#</sup> |
| White River crayfish (*Procambarus acutus*) | North America | Introduced | Very unlikely | Major | Medium | Medium | Not listed | Prohibited list<sup>#</sup> |
| Marmorkrebs (*Procambarus virginalis*) | Unknown | Established | Fairly probable | Major | High | Difficult | Not listed | Prohibited list<sup>#</sup> |

<sup>#</sup>See Table 4 for notes, exemptions and recommendations

### Likelihood of entry

The likelihood of entry into South Africa for most of the assessed crayfish species (64%) varied from fairly probable to probable, because there is some evidence that the species are present in the country in the pet trade, aquaculture facilities and/or neighbouring countries (Nunes et al. 2017a; Madzivanzira et al. 2020). However, the level of confidence in some of the evidence is low and requires verification through follow up studies. For example, there is reliable evidence that *Cherax quadricarinatus* and *Procambarus clarkii* are present in South Africa, but it is unclear if both marron crayfish species (*Cherax cainii* and *C. tenuimanus*) are present. Three species (*C. quadricarinatus*, *Pacifastacus leniusculus* and *Procambarus clarkii*) were assigned a score of probable for likelihood of entry into South Africa (Table 2). Two of the three species (*C. quadricarinatus* and *P. clarkii*) have been formally documented to be present in the country and there is anecdotal evidence that *P. leniusculus* is likely present in the country through the pet trade. *Procambarus clarkii* is known to occur in South Africa at several sites that are located across three provinces (Free State, Mpumalanga and Western Cape) (Barkhuizen et al. 2022; Nunes et al. 2017c; CapeNature unpublished data). Six species (*Cherax cainii*, *C. destructor*, *C. tenuimanus*, *Faxonius rusticus*, *F. virilis* and *Procambarus virginalis*) were assigned a score of fairly probable because of their availability in the global pet trade industry. Recent import permit records indicate that *C. tenuimanus* may be present in the country; however, this still needs to be confirmed because of the taxonomic uncertainty of whether the species imported was *C. cainii* or *C. tenuimanus* (Table 1). The likelihood of entry for the remainder of the assessed species was scored as unlikely for *Faxonius immunis* and very unlikely for *F. juvenilis*, *F. limosus*, *Pacifastacus leptodactylus*, and *Procambarus acutus* because there are no known records of the species in South Africa or in neighbouring countries (Table 1).

**Table 2.**
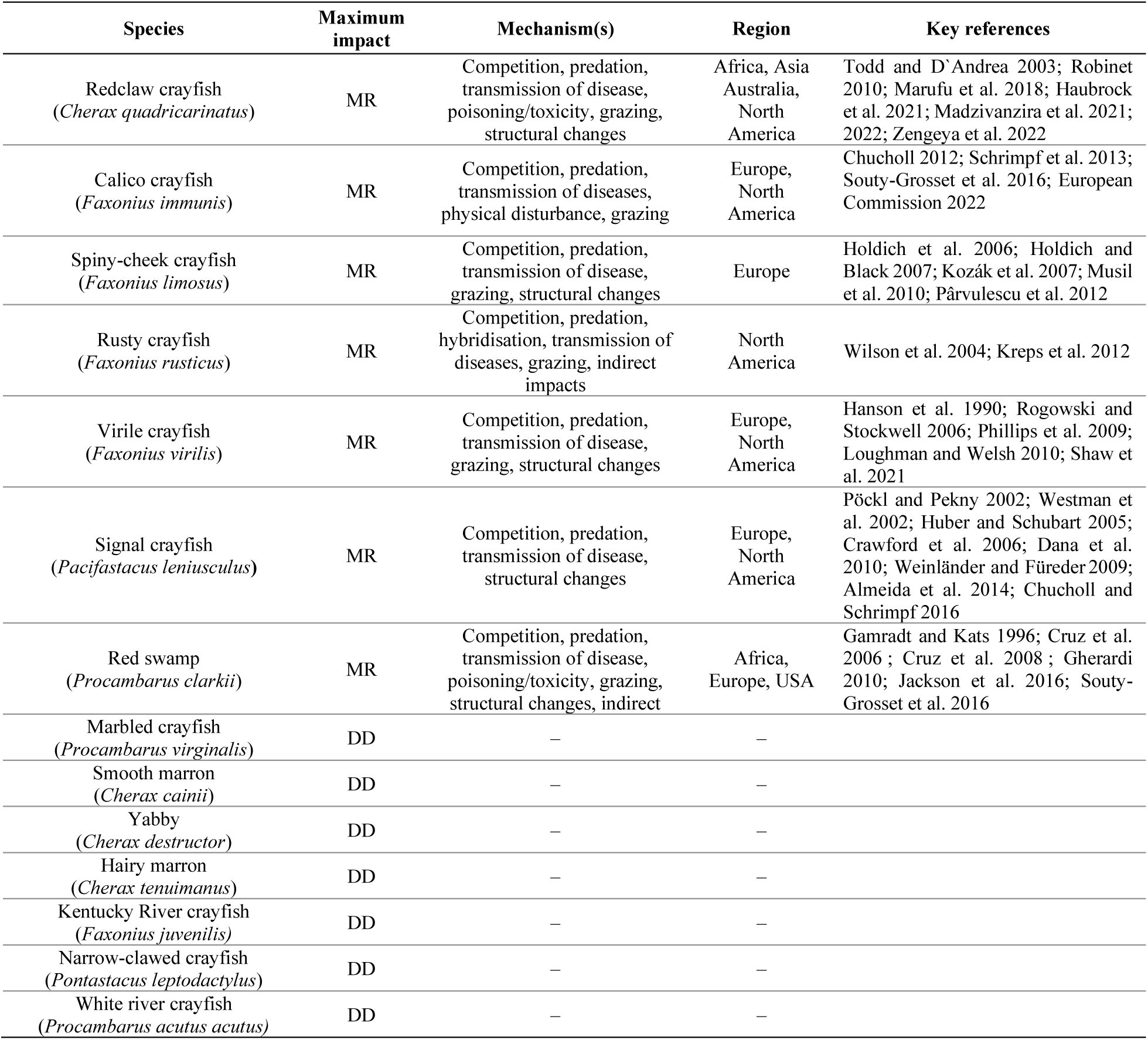
Environmental Impact Classification for Alien Taxa (EICAT) assessments of 14 alien freshwater crayfish that have a global invasion history. Key references provided but detailed information on EICAT assessments and literature cited are provided in Supplement 2.1. Impact mechanisms: MR = local extinctions of at least one species; DD = Data Deficient.

### Consequences

Of the 14 alien freshwater crayfish species assessed, seven species (*Cherax quadricarinatus*, *Faxonius immunis*, *F. limosus, F. rusticus*, *F. virilis, Pacifastacus leniusculus*, and *Procambarus clarkii*) had recorded environmental impacts in their introduced range and the remainder were all classified as Data Deficient (DD) (Table 2). The environmental impacts were mainly associated with predation, competition, transmission of diseases, and the magnitude of the impacts varied mainly from minor (24%), moderate (34%) to major (32%) (Figure 3). Most of the impact studies were recorded from Europe and North America, with a few case studies from Africa, Asia, Australia, and South America (Figure 3).

**Figure 3.**
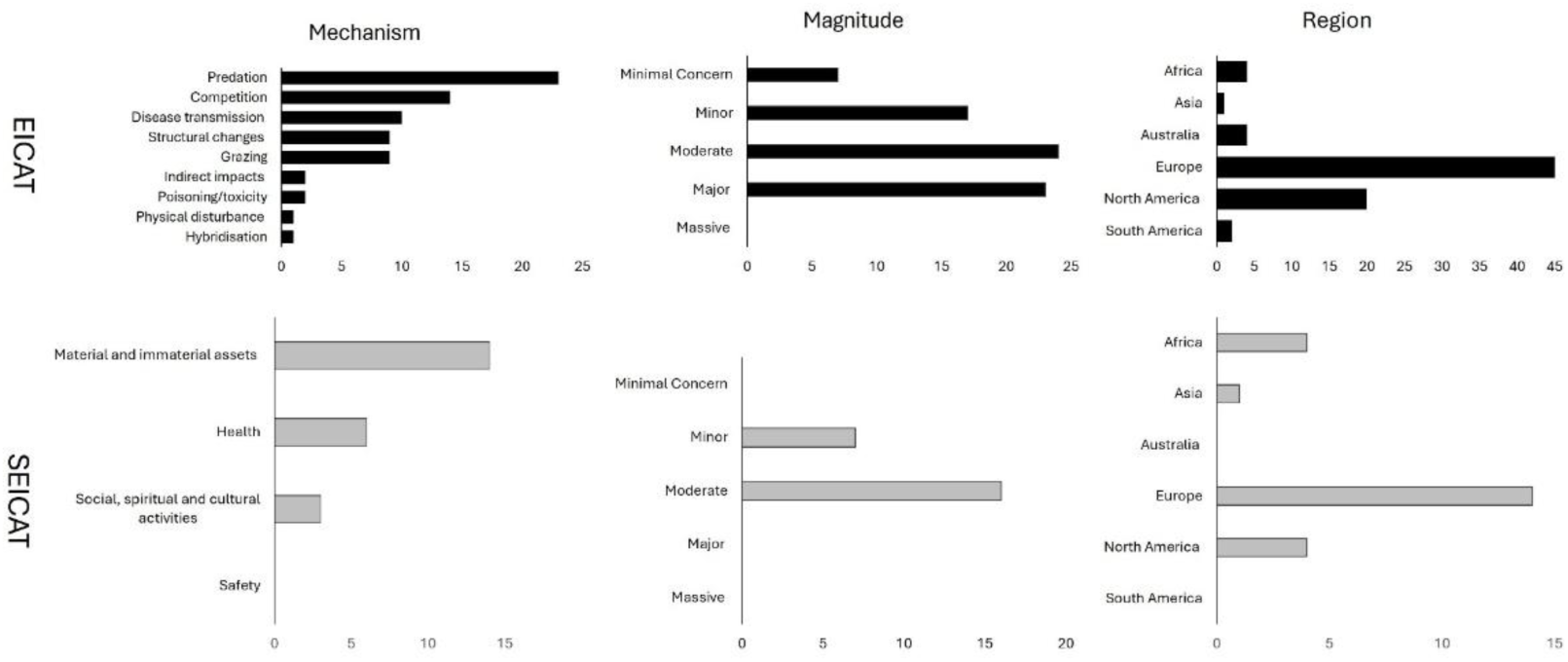
The mechanism, magnitude and region associated with environmental and non-environmental impacts of alien crayfish in their global invasive range. EICAT = Environmental Impact Classification for Alien Taxa; SEICAT = Socio-Economic Impact Classification for Alien.

Similar to the EICAT, seven species (*Cherax quadricarinatus*, *Faxonius immunis*, *F. limosus, F. rusticus*, *F. virilis, Pacifastacus leniusculus,* and *Procambarus clarkii*, had recorded non-environmental impacts in their alien range and the remainder were all classified as Data Deficient (DD) (Table 3). The non-environmental impacts were mainly on material and immaterial assets, human health and social, spiritual and cultural activities, and the magnitude of the impacts was either minor (30%) or moderate (70%) (Figure 2). Most of the impacts (61%) were recorded from Europe and only a few studies were from Africa, Asia and North America (Figure 3).

**Table 3.** Socio-Economic Impact Classification for Alien Taxa (SEICAT) assessments of 14 alien freshwater crayfish that are currently known to have a global invasion history. Detailed information on SEICAT assessments and the relevant literature cited are provided in Supplement 2.2. Impact mechanisms: MN = well-being of individual people is reduced; MO = change to human activity sizes; MR = local extinctions of at least one species; and DD = Data Deficient.

| Species | Maximum impact | Mechanism(s) | Region(s) where impacts were recorded | Key references |
| --- | --- | --- | --- | --- |
| Redclaw crayfish<br>(*Cherax quadricarinatus*) | MO | Material and immaterial assets | Africa, North America | Vega-Villasante et al. 2015; Douthwaite et al. 2018; Haubrock et al. 2021; Madzivanzira et al. 2022; Chakandinakira et al. 2023; Madzivanzira et al. 2023 |
| Calico crayfish<br>(*Faxonius immunis*) | MN | Material assets | Europe | Souty-Grosset et al. 2016 |
| Spiny-cheek crayfish<br>(*Faxonius limosus*) | MO | Material and immaterial assets, social, spiritual and cultural relations | Europe | Souty-Grosset et al. 2006; Aldridge 2011 |
| Rusty crayfish<br>(*Faxonius rusticus*) | MO | Material and immaterial assets | North America | Keller et al. 2008 |
| Virile crayfish<br>(*Faxonius virilis*) | MN | Material and immaterial assets | North America | GISD 2023; NEMESIS 2023 |
| Signal crayfish<br>(*Pacifastacus leniusculus*) | MO | Material and immaterial assets, social, spiritual and cultural activities | Europe | Holdich et al. 2009; Gherardi et al. 2011b; Kouba et al. 2022 |
| Red swamp<br>(*Procambarus clarkii*) | MO | Material assets, health, social, spiritual and cultural relations | Africa, Europe, North America | Lane et al. 2009; Anda et al. 2011; Gherardi et al. 2011b; Souty-Grosset et al. 2016; Putra et al. 2018; Sara and El Moutaouakil 2019 |
| Smooth marron<br>(*Cherax cainii*) | DD | — | — |  |
| Yabby<br>(*Cherax destructor*) | DD | — | — |  |
| Hairy marron<br>(*Cherax tenuimanus*) | DD | — | — |  |
| Kentucky River crayfish<br>(*Faxonius juvenilis*) | DD | — | — |  |
| Narrow-clawed crayfish<br>(*Pontastacus leptodactylus*) | DD | — | — |  |
| White river crayfish<br>(*Procambarus acutus*) | DD | — | — |  |
| Marbled crayfish<br>(*Procambarus virginalis*) | DD | — | — |  |

There are several management options and recommendations for management of 14 alien crayfish species in South Africa (Table 4). Four species (*Cherax cainii*, *C. tenuimanus*, *C. quadricarinatus* and *Procambarus clarkii*) are known to occur in South Africa and are currently listed under the A&IS Regulations. Management options should focus on genetically-verifying species identity (*Cherax cainii* and *C. tenuimanus*) and containing further spread of *C. quadricarinatus* and *Procambarus clarkii*. Five species (*Cherax destructor*, *Faxonius limosus*, *F. rusticus*, *Pacifastacus leniusculus* and *Pontastacus leptodactylus*) are listed under the A&IS Regulations but there is no evidence that they are present in the country, and management options should focus on confirming presence through dedicated assessment of the ornamental trade. The remainder (*Faxonius immunis*, *F. juvenilis*, *F. virilis*, *Procambarus acutus* and *P. virginalis*) have no documented evidence that they are present in South Africa and are not listed on the A&IS Regulations. Management options should therefore focus on preventing introduction.

**Table 4.**
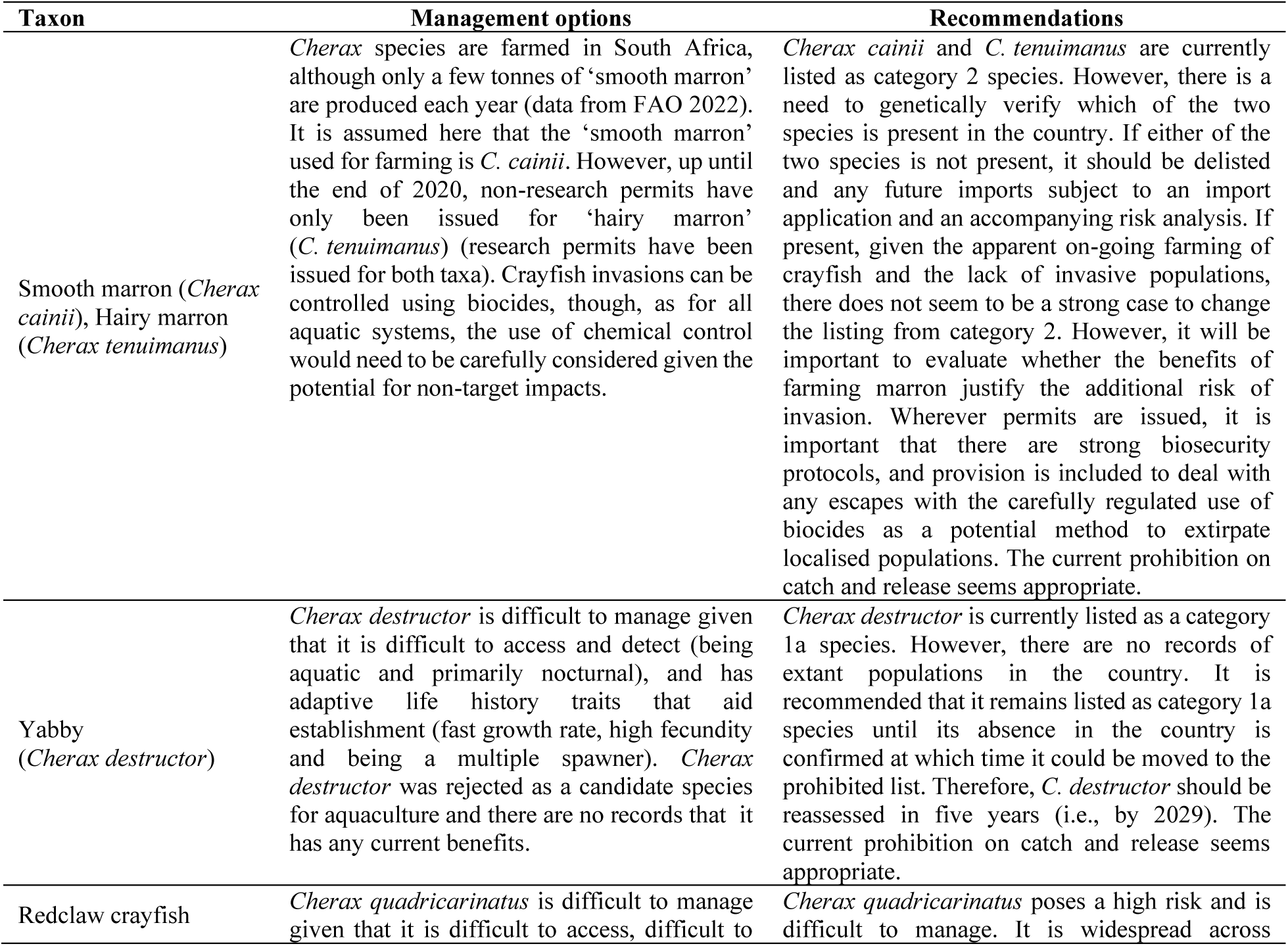
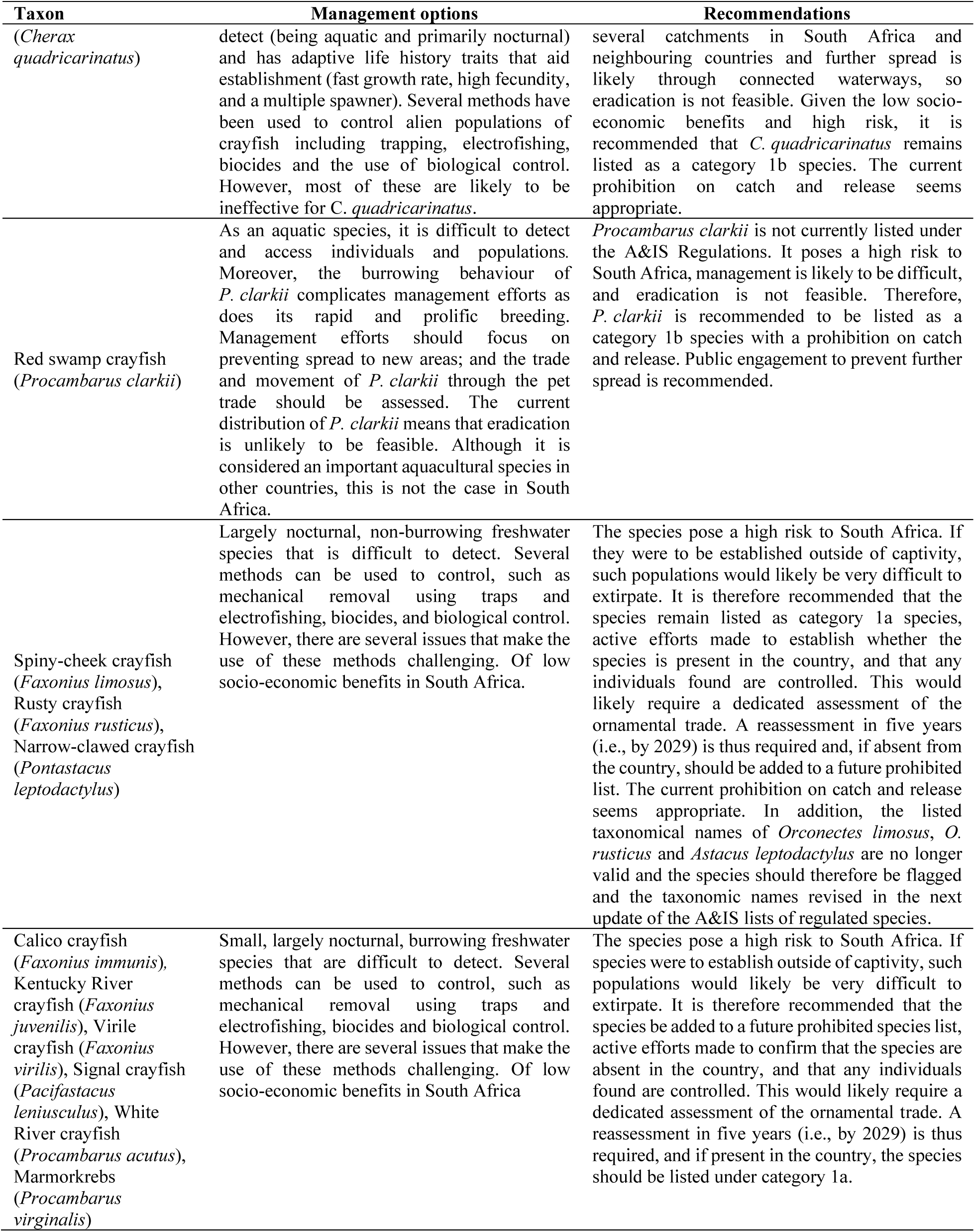
Management options and recommendations for management of 14 alien crayfish species in South Africa. Four species are known to be present in the country and the proposed management actions include genetically verifying species identity and containing further spread of widespread species. Five species are listed under the Alien and Invasive Species (A&IS) Regulations but there is no evidence that they are present in the country, and management options should focus on confirming presence through dedicated assessment of the ornamental trade. The remaining five species have no documented evidence that they are present in South Africa and are not listed on the A&IS Regulations. Management options should therefore focus on preventing introduction.

## Discussion

The risk analyses in this study identified 14 crayfish species that have a global invasion history of which 13 species were classified as a high risk for South Africa because they have a potential to cause major negative impacts. Alien freshwater crayfish have been widely introduced worldwide for aquaculture and the pet trade (Faulkes 2015; Olden & Carvalho 2024). For example, freshwater crayfish account for 5% of global aquaculture production annually by quantity (2.5 million tonnes) and 13% by value (USD 21 billion) per annum (FAO 2022). About 91% of the global crayfish production is provided by alien crayfish species, mainly from the production of *Procambarus clarkii* (Lodge et al. 2012; FAO 2022). There is also extensive trade of crayfish species as pets in aquariums globally and over 120 crayfish species are available for sale as pets (Chucholl 2013; Faulkes 2015; Olden & Carvalho 2024). The socio-economic benefits derived from alien crayfish in South Africa is however low. Aquaculture of crayfish is restricted to marron (*Cherax cainii* and C. *tenuimanus*) in a few small-scale aquaculture farms in Eastern and Western Cape provinces (Nunes et al. 2017a; Madzivanzira et al. 2020). The benefits derived from alien crayfish in the pet trade in South Africa is unknown and it is likely going to be difficult to quantify because trade data are not available. This lack of trade information in the pet trade is a common global problem and is likely due to several reasons (Bush et al. 2014). For example, traded animals and plants transit through many different entities and locations before they reach a consumer and trade data are not always systematically collected, curated and made readily accessible (Sinclair et al. 2021). Some of the pet trade activities are illegal and difficult to track because they try to circumvent regulations that protect endangered plants and animals from the threats of international trade (Olden & Carvalho 2024).

Despite providing societal benefits, crayfish introductions are also known to cause adverse impacts in areas of introduction. The environmental impacts of invasive crayfish are mainly associated with competition, predation and transmission of diseases. For example, crayfish species such as *Faxonius rusticus* and *Procambarus clarkii* are facultative omnivores and aggressive competitors that often displace native species through competition and predation (Jonas et al. 2005; Cruz et al. 2006; Roth et al. 2006; Bobeldyk & Lamberti 2008; Keller et al. 2008; Jackson et al. 2016). Alien crayfish can also harbour pathogens, parasites and diseases that can be transmitted to native congeneric species and cause major impacts in recipient ecological communities (Mastitsky et al. 2010; Longshaw 2011; Lodge et al. 2012). For example, several North American crayfish species such as *Pacifastacus leniusculus* and *Procambarus clarkii* have been implicated in the transmission of crayfish plague to native crayfish species in Europe leading to a decline in populations and in some cases the collapse of ecological communities (Holdich & Reeve 1991; Holdich et al. 2009; Chucholl & Schimpf 2016; Souty-Grosset et al. 2016).

Invasive crayfish can cause substantial economic losses, and they have been estimated to have caused up to USD 120 million in reported costs on material and immaterial assets, human health, and social and cultural activities (Kouba et al. 2022). For example, some crayfish species such as *P. clarkii* construct burrows in rice fields that can affect crop yields, leading to a loss in revenue (Souty-Grosset et al. 2016). Its burrows can also compromise bank morphology and accelerate soil erosion, making invaded areas susceptible to flooding (Haubrock et al. 2019). The presence of alien crayfish can also threaten human livelihoods by replacing native species with higher economic value (Marbuah et. al. 2014). Alien crayfish can also affect artisanal fisheries by damaging nets and spoiling catches leading to economic losses for fishers (Keller et al. 2008; Gherardi et al. 2011b; Chakandinakira et al. 2023; Madzivanzira et al. 2023). Alien crayfish can also cause human health issues, for example when poorly-cooked crayfish are consumed (Lane et al. 2009) or when handling infected crayfish (Anda et al. 2001).

Most of the impacts from crayfish invasions have been documented for Europe and North America and a few from countries in the global south (Lodge et al 2012). This highlights the need for more studies on impacts, especially in Africa, where there are no native freshwater crayfish (except for Madagascar) and invasive crayfish are phenotypically novel and can cause major impacts as they can act as novel predators, competitors and vectors of pathogens (Lodge et al. 2012; Twardochleb et al. 2013; Madzivanzira et al. 2020). For example, crayfish invasions might accelerate ecosystem functions such as shredding and decomposition rates of plant material leading to altered ecosystem functions (e.g., Jackson et al. 2016). In South Africa, there is a growing body of research around invasive crayfish (Zengeya & Wilson 2020; Van Wilgen et al. 2022). These include reviews of status and trends of crayfish invasions (Nunes et al. 2017a, c; Madzivanzira et al. 2020; Weyl et al. 2020), the extent and distribution of crayfish introductions (Nunes et al. 2017b), aspects of ecological community invisibility (e.g., South et al. 2020), and evaluations of potential impacts (e.g., Du Preez & Smith 2013; Tavakol et al. 2021; Madzivanzira et al. 2021, 2022; Zengeya et al. 2022). There is also a growing number of studies on crayfish invasion in neighbouring countries, especially in the upper and middle Zambezi River (e.g., Douthwaite et al. 2018; Marufu et al. 2018; Madzivanzira et al. 2021; Chakandinakira et al. 2023; Nawa et al. 2024).

There are several crayfish species that have a global invasion history but are not present in South Africa. This signifies a significant invasion debt to South Africa and management efforts should focus on preventing introductions, especially illegal introductions through the pet trade. For example, *Procambarus virginalis* was first discovered in the pet trade in Germany and is now also known to occur in several countries in Europe (Faulkes 2010; Gutekunst et al. 2021; Olden & Carvalho 2024) and Madagascar (Jones et al. 2009). This species poses a significant threat in areas of introduction as it reproduces through parthenogenesis and there is limited information on potential impacts in invaded areas (Jones et al. 2009). The pet trade industry is a cause for concern, particularly in South Africa, because the movement of crayfish has not been evaluated (De Moor 2002; Nunes et al. 2017a; Madzivanzira et al. 2020). Other countries such as Australia and New Zealand have used risk analysis protocols to identify and prohibit imports of potential harmful species through prominent pathways such as the pet trade (Lodge et al. 2016). South Africa is adopting a similar approach, where risk analyses will form the basis of evidence to inform management and develop policy of alien species (Kumschick et al. 2020b). The importation of alien freshwater species for aquaculture appears to be regulated better under a functioning permitting system (Wilson & Kumschick 2024). However, the situation is not clear in the pet trade and live bait industries (e.g., Douthwaite et al. 2018; Vezi et al. 2024). Various studies have reported that these industries are generally not well-regulated (Distefano et al. 2009) and where regulations have been developed, they are either not enforced adequately or are difficult to enforce (Kilian et al. 2012; Patoka et al. 2018). In South Africa, several alien crayfish species are available to buy online and in pet shops, although there is no legal documentation of their introduction (De Moor 2002; Nunes et al. 2017a; Madzivanzira et al. 2020). From the information available, there is no indication whether the movement of crayfish species through the pet trade and aquarium industries has been evaluated to fully anticipate the level of risk they pose as pathways of introduction (Nunes et al. 2017a; Madzivanzira et al. 2020).

*Cherax quadricarinatus* and *Procambarus clarkii* are now widespread across several catchments in South Africa and neighbouring countries and further spread is likely through connected waterways, so eradication is not feasible. *Procambarus clarkii* is not currently listed as under the A&IS Regulations, but it is now widespread across several sites in South Africa (Nunes et al. 2017b; Barkhuizen et al. 2022; CapeNature unpublished data). It was previously listed as a prohibited species (i.e., a species that is not present in South Africa and import is not permitted) but was removed when the A&IS Regulations were updated in 2021 (Wilson & Kumschick 2024). The rationale for removing the prohibited list was that there were no risk analyses done to inform why a species should be prohibited and therefore the lists might have been subject to disapproval. Therefore, for any species that is not present in South Africa, a risk analysis is required to inform whether the introduction should be allowed and if so, what are the appropriate management interventions required to minimise risk of spread and impacts. The utilisation and spread of *P. clarkii* were therefore illegal and this was likely confounded by the uncertainty around its listing, and unfortunately likely lead to its spread across the country. The primary pathway of introduction and spread are likely through the pet trade and intentional release as bait for angling (Nunes et al. 2017b). *Procambarus clarkii* is popular in the global pet trade (Faulkes 2015; Patoka et al. 2018) and as bait for angling (Olden et al. 2009; Larson & Olden 2011; Kilian et al. 2012; Kerr 2014). Previous attempts to eradicate *P. clarkii* in South Africa using mechanical methods at two sites in the Free State and Mpumalanga provinces were unsuccessful (Nunes et al. 2017c; Barkhuizen et al. 2022). There are several reasons why the methods (partial dewatering and physical removal) were not effective. The control methods were implemented over a short period of time with no long-term commitment to follow up to ensure complete eradication (e.g., Loureiro et al. 2018). *Procambarus clarkii* can construct burrows that can impede control methods (e.g., Gherardi 2006; Gherardi 2011a; Nunes et al. 2017c; Haubrock et al. 2019), and mechanical removal methods can be size-selective leading to community compensatory mechanisms that counteract removal efforts (Manfrin et al. 2019; Chadwick et al. 2020).

## Conclusion

Thirteen crayfish were assessed as high risk because they are likely to be introduced into South Africa and have the potential to cause major impacts. The environmental impacts of invasive crayfish are mainly associated with competition, predation and transmission of diseases, while non-environmental impacts are associated with material and immaterial assets, human health, and social and cultural activities. Most of these impacts were documented in Europe and North America and a few from countries in the global south. This highlights the need for more studies on impacts, especially in Africa, where there are no native freshwater crayfish (except for Madagascar) and invasive crayfish are phenotypically novel and can cause major impacts. This study provides several recommendations for management of alien crayfish species in South Africa. There is evidence that four crayfish species are present in the country and the proposed management actions include genetically verifying the identity of marron species (*Cherax cainii* and *C. tenuimanus*) and containing further spread of widespread species such as *Cherax quadricarinatus* and *Procambarus clarkii*. There is anecdotal evidence that *Faxonius limosus*, *F. rusticus*, *Cherax destructor*, *Pontastacus leptodactylus* and *Pacifastacus leniusculus* are present through the pet trade but this needs to be verified. The pet trade is a significant pathway for the introduction of crayfish globally, and it is concerning that the movement of crayfish into South Africa through this pathway has not been evaluated. There are several other crayfish species (*Faxonius immunis*, *F. juvenilis*, *F. virilis*, *Procambarus acutus* and *P. virginalis*) that have a global invasion history but are not present in South Africa and this signifies a significant invasion debt for the country and beyond. Management options should therefore focus on preventing introduction.

This paper demonstrated the use of a formal science-based risk analysis framework and how it can be used to support policy decision-makers on the risks posed by alien crayfish introductions in South Africa and provided recommendations on potential mitigation measures that can prevent the introduction of harmful species and minimise impacts. It is envisaged that the risk analyses could inform the implementation appropriate control methods, and policies that improve the management of alien freshwater crayfish in South Africa and beyond.

## Acknowledgements

This project was funded by the Department of Forestry, Fisheries and the Environment (DFFE) through the South African National Biodiversity Institute (SANBI) and the Centre for Invasion Biology (CIB). DFFE is thanked for funding, noting that this publication does not necessarily represent the views or opinions of DFFE or its employees. We also thank the anonymous reviewers who provided feedback that improved the risk analyses and the manuscript.

## Authors’ contributions

L.B., C.T.C., T.A.Z. designed and conceptualised the study, L.B. led the writing and analysis (as part of her MSc dissertation). All authors read, edited and agreed to the published version of the manuscript.

## References

Aldridge, D., 2011. ‘Spinycheek Crayfish, Orconectes limosus’, Sand Hutton, UK, GB Non-native Species secretariat, https://www.nonnativespecies.org/non-native-species/information-portal/view/2441#.

Almeida, D., Ellis, A., England, J. & Copp, G.H., 2014, ‘Time-series analysis of native and non-native crayfish dynamics in the Thames River Basin (south-eastern England)’, Aquatic Conservation: Marine and Freshwater Ecosystems 24, 192–202, 10.1002/aqc.2366.

Anda, P., Del Pozo, J.S., García, J.MD., Escudero, R., Peña, F.J.G, Velasco, M.C, Sellek, R.E., Chillarón, M.R.J., Serrano, L.P.S. & Navarro, J.F.M., 2001, ‘Waterborne outbreak of tularemia associated with crayfish fishing’, Emerging Infectious Diseases, 7, 575–582, 10.3201/eid0707.010740.

Austin, C.M. & Ryan, S.G., 2002, ‘Allozyme evidence for a new species of freshwater crayfish of genus *Cherax* Erichson (Decapoda: Parastacidae) from the south-west of Western Australia’, Invertebrate Systematics 16, 357–367, 10.1071/it01010.

Bacher, S., Blackburn, T.M., Essl, F., Genovesi, P., Heikkilä, J., Jeschke, J.M., Jones, G., Keller, R., Kenis, M., Kueffer, C., Martinou, A.F., Nentwig, W., Pergl, J., Pyšek, P., Rabitsch, W., Richardson, D.M., Roy, H.E., Saul, W.C., Scalera, R. Vilà, M., Wilson, J.R.U. & Kumschick, S., 2017, ‘Socio-economic impact classification of alien taxa (SEICAT)’, Methods in Ecology and Evolution, 159–168, 10.1111/2041-210X.12844.

Barkhuizen, L.M., Madzivanzira, T.C. & South, J., 2022, ‘Population ecology of a wild population of red swamp crayfish *Procambarus clarkii* (Girard, 1852) in the Free State Province, South Africa and implications for eradication efforts’, BioInvasions Record, 11(1), 181–191, 10.3391/bir.2022.11.1.18.

Bernery, C., Bellard, C., Courchamp, F., Brosse, S., Gozlan, R.E., Jarić, I., Teletchea, F. & Leroy, B., 2022, ‘Freshwater fish invasions: A comprehensive review’, Annual Review of Ecology, Evolution, and Systematics 53, 427–456, 10.1146/annurev-ecolsys-032522-015551.

Blackburn, T.M., Essl, F., Evans, T., Hulme, P.E., Jeschke, J.M., Kühn, I., Kumschick, S., Marková, Z., Mrugała, A., Nentwig, W., Pergl, J., Pyšek, P., Rabitsch, W., Ricciardi, A., Richardson, D.M., Sendek, A., Vilá, M., Wilson, J.R.U., Winter, M., Genovesi, P. & Bacher, S., 2014, ‘A unified classification of alien species based on the magnitude of their environmental impacts’, PLoS Biology, 12(5), e1001850, 10.1371/journal.pbio.1001850.

Blackburn, T.M., Pyšek, P., Bacher, S., Carlton, J.T., Duncan, R.P., Jarošík, V., Wilson, J.R.U. & Richardson, D.M., 2011, ‘A proposed unified framework for biological invasions’, Trends in Ecology and Evolution, 26(7), 333–339, 10.1016/j.tree.2011.03.023.

Bobeldyk, A.M. & Lamberti, G.A., 2008, ‘A decade after invasion: Evaluating the continuing effects of rusty crayfish on a Michigan River’, Journal of Great Lakes Research, 34, 265–275, 10.3394/0380-1330(2008)34[265:ADAIET]2.0.CO;2.

Bush, E.R., Baker, S.E. & MacDonald, D.W., 2014, ‘Global trade in exotic pets 2006–2012’, Conservation Biology, 28(3), 663–676, 10.1111/cobi.12240.

Chadwick, D.D.A, Pritchard, E.G, Bradley, P., Sayer, C.D., Chadwick, M.A., Eagle, L.J.B. & Axmacher, J.C., 2020, ‘A novel “triple down” method highlights deficiencies in invasive alien crayfish survey and control techniques’, Journal of Applied Ecology, 58(2), 316–326, 10.1111/1365-2664.13758.

Chakandinakira, A.T., Madzivanzira, T.C., Mashonga, S., Muzvondiwa, J.V., Ndlovu, N. & South, J. 2023, ‘Socioeconomic impacts of Australian redclaw crayfish *Cherax quadricarinatus* in Lake Kariba’, Biological Invasions 25, 2801–2812, 10.1007/s10530-023-03074-8.

Chucholl, C., 2012, ‘Understanding invasion success: life-history traits and feeding habits of the alien crayfish *Orconectes immunis* (Decapoda, Astacida, Cambaridae)’, Knowledge and Management of Aquatic Ecosystems, 404, 04, 10.1051/kmae/2011082.

Chucholl, C., 2013, ‘Invaders for sale: trade and determinants of introduction of ornamental freshwater crayfish’, Biological Invasions, 15, 125–141, 10.1007/s10530-012-0273-2.

Chucholl, C. & Schimpf, A., 2016, ‘The decline of endangered stone crayfish (*Austropotamobius torrentium*) in southern Germany is related to the spread of invasive alien species and land-use change’, Aquatic Conservation: Marine and Freshwater Ecosystems, 26, 44–56, 10.1002/aqc.2568.

Crandall, K.A. & Buhay, J.E., 2008, ‘Global diversity of crayfish (Astacidae, Cambaridae, and Parastacidae – Decapoda) in freshwater’, Hydrobiologia, 595, 295–301, 10.1007/s10750-007-9120-3.

Crawford, L., Yeomans, W.E. & Adams, C.E., 2006, ‘The impact of introduced signal crayfish *Pacifastacus leniusculus* on stream invertebrate communities’, Aquatic Conservation: Marine and Freshwater Ecosystems, 16, 611–621, 10.1002/aqc.761.

Cruz, M.J., Rebelo, R. & Crespo, E.G., 2006, ‘Effects of an introduced crayfish, *Procambarus clarkii*, on the distribution of south-western Iberian amphibians in their breeding habitats’, Ecography, 29, 329–338, 10.1111/j.2006.0906-7590.04333.x.

Cruz, M.J., Segurado, P., Sousa, P.M. & Rebelo, R., 2008, ‘Collapse of the amphibian community of the Paul do Boquilobo Natural Reserve (central Portugal) after the arrival of the exotic American crayfish *Procambarus clarkii*’, Herpetological Journal 18, 197–204.

Dana, E.D., López-Santiago, J., García-De-Lomas, J., García-Ocaña, D.M. & Ortega, F., 2010, ‘Long-term management of the invasive *Pacifastacus leniusculus* (Dana, 1852) in a small mountain stream’, Aquatic Invasions, 5, 317–322, 10.3391/ai.2010.5.3.10.

De Moor, I., 2002, ‘Potential impacts of alien freshwater crayfish in South Africa’, African Journal of Aquatic Science 27, 125–139, 10.2989/16085914.2002.9626584.

DiStefano, R.J., Litvan, L.E. & Horner, P.T., 2009, ‘The bait industry as a potential vector for alien crayfish introductions: Problem recognition by fisheries agencies and a Missouri evaluation’, Fisheries, 12, 586–597, 10.1577/1548-8446-34.12.586.

Douthwaite, R.J., Jones, E.W., Tyser, A.B. & Vrdoljak, S.M., 2018, ‘The introduction, spread and ecology of redclaw crayfish *Cherax quadricarinatus* in the Zambezi catchment’, African Journal of Aquatic Science, 43(4), 353–366, 10.2989/16085914.2018.1517080.

Du Preez, L.H. & Smit. N.J., 2013, ‘Double blow: Alien crayfish infected with invasive temnocephalan in South African waters’, South African Journal of Science 109(10/9), 4, 10.1590/sajs.2013/20130109.

Duffy, R., Ledger, J., Dias, J. & Snow, M., 2014, ‘The critically endangered harry marron, *Cherax tenuimanus* Smith 1912: A review of current knowledge and actions required to prevent extinction of a species’, Journal of Royal Society of Western Australia, 97, 297–306.

Dunn, J.C., McClymont, H.E., Christmas, M. & Dunn, A.M., 2009, ‘Competition and parasitism in the native White Clawed Crayfish *Austropotamobius pallipes* and the invasive Signal Crayfish *Pacifastacus leniusculus* in the UK’, Biological Invasions, 11, 315–324, 10.1007/s10530-008-9249-7.

Eby, L.A., Roach, W.J., Crowder, L.B. & Stanford, J.A., 2006, ‘Effects of stocking-up freshwater food webs’, Trends in Ecology & Evolution, 21(10), 576–584, 10.1016/j.tree.2006.06.016.

Essl, F., Latombe, G., Lenzner, B., Pagad, S., Seebens, H., Smith, K., Wilson, J.R.U. & Genovesi, P., 2020, ‘The Convention on Biological Diversity (CBD)’s Post-2020 target on invasive alien species – what should it include and how should it be monitored?’, in J.R. Wilson, S. Bacher, C.C. Daehler, Q.J. Groom, S. Kumschick, J.L. Lockwood, T.B. Robinson, T.A. Zengeya, & D.M. Richardson (eds), Frameworks used in Invasion Science, NeoBiota, 62, 99–121, 10.3897/neobiota.62.53972.

European Commission 2022, Directorate-General for Environment, Study on invasive alien species – Development of risk assessments to tackle priority species and enhance prevention – Final report (and annexes), Publications Office of the European Union, 2022, https://data.europa.eu/doi/10.2779/302048.

FAO, 2022, ‘Top 10 species groups in global, regional and national aquaculture 2020. Supplementary materials to the factsheet on Top 10 species groups in global aquaculture 2020’, World Aquaculture Performance Indicators (WAPI) factsheet, www.fao.org/3/cc0681en/cc0681en.pdf.

Faulkes, Z., 2010, ‘The spread of parthogenetic marbled crayfish, Marmorkebs (*Procambarus sp*.) in the North American pet trade’, Aquatic Invasions 5, 447–450, 10.3391/ai.2010.5.4.16.

Faulkes, Z., 2015, ‘The global trade in crayfish as pets’, Crustacean Research 44, 75–92, 10.18353/crustacea.44.0_75.

Gamradt, S.C. & Kats, L.B., 1996, ‘Effect of introduced crayfish and mosquitofish on California newts’, Conservation Biology 10(4), 1155–1162, 10.1046/j.1523-1739.1996.10041155.x.

Gherardi, F., 2006, ‘Crayfish invading Europe: the case study of *Procambarus clarkii*’, Marine and Freshwater Behaviour and Physiology 39, 175–191, 10.1080/10236240600869702.

Gherardi, F., 2010, ‘Invasive crayfish and freshwater fishes of the world’, Scientific and Technical Review of the Office International des Epizooties, 29, 241–254, 10.20506/rst.29.2.1973.

Gherardi, F., Aquiloni, L., Diéguez-Uribeondo, J. & Tricarico E., 2011a, ‘Managing invasive crayfish: is there a hope?’, Aquatic Sciences, 73, 185–200, 10.1007/s00027-011-0181-z.

Gherardi, F., Britton, J.R., Mavuti, K.M., Pacini, N., Grey, J., Tricarico, E. & Harper, D.M., 2011b, ‘A review of allodiversity in Lake Naivasha, Kenya: Developing conservation actions to protect East African lakes from the negative impacts of alien species’, Biological Conservation, 144, 2585–2596, 10.1016/j.biocon.2011.07.020.

GISD, 2023. Species profile: *Orconectes virilis*. Global Invasive Species Database, http://www.iucngisd.org/gisd/species.php?sc=218.

Gutekunst, J., Maiakovska, O., Hanna, K., Provataris, P., Horn, H., Wolf, S., Skelton, C.E., Dorn, N.J. & Lyko, F., 2021, ‘Phylogeographic reconstruction of the marbled crayfish origin’, Communications Biology, 4(1), 1096, 10.1038/s42003-021-02609-w.

Hanson, J.M., Chambers, P.A. & Prepas, E.E., 1990, ‘Selective foraging by the crayfish *Orconectes virilis* and its impact on macroinvertebrates’, Freshwater Biology, 24, 69–80, 10.1111/j.1365-2427.1990.tb00308.x.

Haubrock, P.J., Inghilesi, A.F., Mazza, G., Bendoni, M., Solari, L. & Ttricarico, E., 2019, ‘Burrowing activity of *Procambarus clarkii* on levees: analysing behaviour and burrow structure’, Wetlands Ecology Management, 27, 497–511, 10.1007/s11273-019-09674-3.

Haubrock, P.J., Oficialdegui, F.J., Zeng, Y., Patoka, J., Yeo, D.C.J. & Kouba, A., 2021, ‘The redclaw crayfish: A prominent aquaculture species with invasive potential in tropical and subtropical biodiversity hotspots’, Reviews in Aquaculture 13, 1488–1530, 10.1111/raq.12531.

Hawkins, C.L., Bacher, S., Essl, F., Hulme, P.E., Jeschke, J.M., Kühn, I., Kumschick, S., Nentwig, W., Pergl, J., Pyšek, P., Rabitsch, W., Richardson, D.M., Vilá, M., Wilson, J.R.U., Genovesi, P. & Blackburn, T.M., 2015, ‘Framework and guidelines for implementing the proposed IUCN Environmental Impact Classification for Alien Taxa (EICAT)’, Diversity and Distributions, 21, 1360–1363, 10.1111/ddi.12379.

Holdich, D. & Black, J., 2007, ‘The spiny-cheek crayfish, *Orconectes limosus* (Rafinesque, 1817) [Crustacea: Decapoda: Cambaridae], digs into the UK’, Aquatic Invasions, 2(1), 1–16, 10.3391/ai.2007.2.1.1.

Holdich, D.M., Haffner, P. & Noël, P., 2006, ‘Species files’, in C. Souty-Grosset, D.M. Holdich, P.Y. Noël, J.D. Reynolds & P. Haffner (eds), Atlas of Crayfish in Europe, Muséum national d’Histoire naturelle, Paris, Patrimoines naturels, 64, 50–129.

Holdich, D.M. & Reeve, I.D., 1991, ‘Distribution of freshwater crayfish in the British Isles, with particular reference to crayfish plague, alien introductions and water quality’, Aquatic Conservation: Marine and Freshwater Ecosystems 1, 139–158, 10.1002/aqc.3270010204.

Holdich, D.M., Reynolds, J.D. & Sibley, P.J., 2009, ‘A review of the ever-increasing threat to European crayfish from non-indigenous crayfish species’, Knowledge and Management of Aquatic Ecosystems, 394–395, 11, 10.1051/kmae/2009025.

Huber, M.G. & Schubart, C.D., 2005, ‘Distribution and reproductive biology of *Austropotamobius torrentium* in Bavaria and documentation of a contact zone with the alien crayfish *Pacifastacus leniusculus**’,* Bulletin Français de la Pêche et de la Pisciculture 376, 759–776, 10.1051/kmae:2005031.

Hulme, E., Bacher, S., Kenis, M., Klotz, S., Kühn, I., Minchin, D., Nentwig, W., Oleni, S., Panov, V., Pergl, J., Pyšek, P., Roques, A., Sol, D., Solarz, W. & Vilà, M., 2008, ‘Grasping at the routes of biological invasions: a framework for integrating pathways into policy’, Journal of Applied Ecology, 45, 403–414, 10.1111/j.1365-2664.2007.01442.x.

IPBES, 2023, Summary for Policymakers of the Thematic Assessment Report on Invasive Alien Species and their Control of the Intergovernmental Science-Policy Platform on Biodiversity and Ecosystem Services, Roy, H.E., Pauchard, A., Stoett, P., Renard Truong, T., Bacher, S., Galil, B.S., Hulme, P.E., Ikeda, T., Sankaran, K.V., McGeoch, M.A., Meyerson, L.A., Nuñez, M.A., Ordonez, A., Rahlao, S.J., Schwindt, E., Seebens, H., Sheppard, A.W., & Vandvik, V. (eds), IPBES secretariat, Bonn, Germany, 10.5281/zenodo.7430692.

Jackson, M.C., Grey, J., Miller, K., Britton, J.R. & Donohue, I., 2016, ‘Dietary niche constriction when invaders meet natives: evidence from freshwater decapods’, Journal of Animal Ecology, 85, 1098–1107, 10.1111/1365-2656.12533.

Jonas, J.L., Claramunt, R.M., Fitzsimons, J.D., Marsden, J.E. & Ellrott, B.J., 2005, ‘Estimates of egg deposition and effects of lake trout (*Salvelinus namaycush*) egg predators in three regions of the Great Lakes. Canadian Journal of Fisheries and Aquatic Sciences 62: 2254–2264, 10.1139/f05-141.

Jones, J.P.G., Rasamy, J.R., Harvey, A., Toon, A., Oidtmann, B., Randrianarison, M.H., Raminosoa, N. & Ravoahangimalala, O.R., 2009, ‘The perfect invader: a parthenogenetic crayfish poses a new threat to Madagascar’s freshwater biodiversity’, Biological Invasions, 11, 1475–1482, 10.1007/s10530-008-9334-y.

Keller, R.P., Frang, K. & Lodge, D.M., 2008, ‘Preventing the spread of invasive species: economic benefits of intervention guided by ecological predictions’, Conservation Biology, 22, 80–88, 10.1111/j.1523-1739.2007.00811.x.

Kerr, S.J., 2014, ‘The introduction and spread of aquatic invasive species through the recreational use of bait: a literature review’, report prepared for Biodiversity Branch, Ontario Ministry of Natural Resources, Peterborough, Ontario, Canada.

Kilian, J.V., Klauda, R.J., Widman, S., Kashiwagi, M., Bourquin, R., Weglein, S. & Schuster, J., 2012, ‘An assessment of a bait industry and angler behavior as a vector of invasive species’, Biological Invasions, 14(7), 1469–1481, 10.1007/s10530-012-0173-5.

Kouba, A., Oficialdegui, F.J., Cuthbert, R.N., Kourantidou, M., South, J., Tricarico, E., Gozlan, R.E., Courchamp, F. & Haubrock, P.J., 2022, ‘Identifying economic costs and knowledge gaps of invasive aquatic crustaceans’, Science of the Total Environment, 813, 152325, 10.1016/j.scitotenv.2021.152325.

Kozák, P., Buřič, M., Policar, T., Hamáčková, J. & Lepičová, A., 2007, ‘The effect of inter-and intra-specific competition on survival and growth rate of native juvenile noble crayfish *Astacus astacus* and alien spiny-cheek crayfish *Orconectes limosus*’, Hydrobiologia, 590, 85–94, 10.1007/s10750-007-0760-0.

Kreps, T.A., Baldridge, A.K. & Lodge, D.M., 2012, ‘The impact of an invasive predator (*Orconectes rusticus*) on freshwater snail communities: insights on habitat-specific effects from a multilake long-term study’, Canadian Journal of Fisheries and Aquatic Sciences, 69, 1164–1173, 10.1139/f2012-052.

Kumschick, S., Foxcroft, L. & Wilson, J.R.U., 2020a, ‘A framework to support alien species regulation: the Risk Analysis for Alien Taxa (RAAT)’, in J.R. Wilson, , S. Bacher, C.C. Daehler, Q.J. Groom, S. Kumschick, J.L. Lockwood, T.B. Robinson, T.A. Zengeya, & D.M. Richardson (eds), Frameworks used in Invasion Science, NeoBiota, 62, 213–239, 10.3897/neobiota.62.51031.

Kumschick, S., Foxcroft, L. & Wilson, J.R.U., 2020b, ‘Analysing the risks posed by biological invasions to South Africa’, in B.W. Van Wilgen, J. Measey, D.M. Richardson, J.R. Wilson, & T.A. Zengeya (eds), Biological Invasions in South Africa, Springer, Berlin, pp. 573–595, 10.1007/978-3-030-32394-3_20.

Lane, M.A., Barsanti, M.C., Santos, C.A., Yeung, M., Lubner, S.J. & Weil, G.J., 2009, ‘Human paragonimiasis in North America following ingestion of raw crayfish’, Clinical Infectious Diseases, 49, 55–61, 10.1086/605534.

Larson, E.R. & Olden, J.D., 2011, ‘The state of crayfish in the Pacific Northwest’, Fisheries, 36(2), 60–73, 10.1577/03632415.2011.10389069.

Laverty, C., Nentwig, W., Dick, J.T.A. & Lucy, F.E., 2015, ‘Alien aquatics in Europe: assessing the relative environmental and socioeconomic impacts of invasive aquatic macroinvertebrates and other taxa’, Management of Biological Invasions, 6, 341–350, 10.3391/mbi.2015.6.4.03.

Lodge, D.M., Deines, A., Gherardi, F., Yeo, D.C.J., Arcella, T., Baldridge, A.K, Barnes, M.A., Chadderton, W.L., Feder, J.L., Gantz, C.A., Howard, G.W., Jerde, C.L., Peters, B.W, Peters, J.A., Sargent, L.W., Turner, C.R., Wittmann, M.E. & Zeng, Y., 2012, ‘Global introductions of crayfishes: evaluating the impact of species invasions on ecosystem services’, Annual Review of Ecology, Evolution, and Systematics, 43, 449–472, 10.1146/annurev-ecolsys-111511-103919.

Lodge, D.M., Simonin, P.W., Burgiel, S.W., Keller, R.P., Bossenbroek, J.M., Jerde, C.L., Kramer, A.M., Rutherford, E.S., Barnes, M.A., Wittmann, M.E., Chadderton, W.L., Apriesnig, J.l., Beletsky, D., Cooke, R.M., Drake, J.M., Egan, S.P., Finnoff, D.C., Gantz, C.A., Grey, E.K., Hoff, M.H., Howeth, J.G., Jensen, R.A., Larson, E.R., Mandrak, N.E., Mason, D.M., Martinez, F.A., Newcomb, T.J., Rothlisberger, J.D., Tucker, A.J. & Warziniack, T.W., 2016, ‘Risk analysis and bioeconomics of invasive species to inform policy and management’, Annual Review of Environment and Resources 41, 453–488, 10.1146/annurev-environ-110615-085532.

Longshaw, M., 2011, ‘Diseases of crayfish: a review’, Journal of Invertebrate Pathology 106, 54–70, 10.1016/j.jip.2010.09.013.

Loughman, Z.J. & Welsh, S.A., 2010, ‘Distribution and conservation standing of West Virginia crayfishes’, Southeastern Naturalist, 9(3), 63–78, 10.1656/058.009.s304.

Loureiro, T.G, Anastácio, P.M, De Siqueira Bueno, S.L. & Araujo, P.B., 2018, ‘Management of invasive populations of the freshwater crayfish *Procambarus clarkii* (Decapoda, Cambaridae): test of a population-control method and proposal of a standard monitoring approach’, Environmental Monitoring Assessment, 190, 559, 10.1007/s10661-018-6942-6.

Madzivanzira, T.C., Chakandinakira, A.T., Mungenge, C.P., O’Brien, G., Dalu, T. & South, J., 2023, ‘Get it before it gets to my catch: misdirection traps to mitigate against socioeconomic impacts associated with crayfish invasion’, Management of Biological Invasions 14(2), 335–346, 10.3391/mbi.2023.14.2.10.

Madzivanzira, T.C., South, J. & Weyl, O.L.F., 2021, ‘Invasive crayfish outperform Potamonautid crabs at higher temperatures’, Freshwater Biology 66, 978–991, 10.1111/fwb.13691.

Madzivanzira, T.C, South, J., Wood, L.E., Nunes, A.L. & Weyl, O.L.F., 2020, ‘A review of freshwater crayfish introductions in Africa’, Reviews in Fisheries Science & Aquaculture, 29, 218–241, 10.1080/23308249.2020.1802405.

Madzivanzira, T.C., Weyl, O.L.F. & South, J., 2022, ‘Ecological and potential socioeconomic impacts of two globally invasive crayfish’, NeoBiota 72, 25–43, 10.3897/neobiota.72.71868.

Manfrin, C., Souty-Grosset, C., Anastacio, P.M., Reynolds, J. & Giulianini, P.G., 2019, ‘Detection and control of invasive freshwater crayfish: from traditional to innovative methods. Diversity, 11, 1–16, 10.3390/d11010005.

Marbuah, G., Gen, I.M. & McKie, B., 2014, ‘Economics of harmful invasive species: a review’. Diversity 6, 500–523, 10.3390/d6030500.

Marufu, L., Dalu, T., Barson, M., Simango, R., Utete, B. & Nhiwatiwa, T., 2018, ‘The diet of an invasive crayfish, *Cherax quadricarinatus* (Von Martens, 1868), in Lake Kariba, inferred using stomach content and stable isotope analyses’, BioInvasions Records, 7, 121–132, 10.3391/bir.2018.7.2.03.

Mastitsky, S.E., Karatayev, A.Y., Burlakova, L.E. & Molloy, D.P., 2010, ‘Parasites of exotic species in invaded areas: does lower diversity mean lower epizootic impact?’ Diversity and Distributions 16, 798–803, 10.1111/j.1472-4642.2010.00693.x.

Musil, M., Buřič, M., Policar, T., Kouba, A. & Kozák, P., 2010, ‘Comparison of diurnal and nocturnal activity between noble crayfish (*Astacus astacus*) and spinycheek crayfish (*Orconectes limosus*)’, Freshwater Crayfish, 17, 189–193, 10.5869/fc.2010.v17.189.

Nawa, N., South, J., Ellender, B.R., Pegg, J., Madzivanzira, T.C. & Wasserman, R.J., 2024, ‘Complex selection processes on invasive crayfish phenotype at the invasion front of the Zambezi floodplains ecoregion’, Freshwater Biology, 69, 1322–1337, 10.1111/fwb.14308,

NEMESIS, 2023, ‘Faxonius virilis, Crustaceans-Crayfish: virile crayfish, Edgewater, MD: National Estuarine and Marine Exotic Species Information System (NEMESIS), Smithsonian Environmental Research Center, https://invasions.si.edu/nemesis/species_summary/97425.

Nunes, A.L., Hoffman, C.A., Zengeya, T.A., Measey, G.J. & Weyl, O.L.F., 2017c, ‘Red swamp crayfish, *Procambarus clarkii*, found in South Africa 22 years after attempted eradication’, Aquatic Conservation: Marine Freshwater Ecosystems, 27, 1334–1340, 10.1002/aqc.2741.

Nunes, A.L, Zengeya, T.A, Measey, G.J. & Weyl, O.L.F., 2017a, ‘Freshwater crayfish invasions in South Africa: past, present and potential future’, African Journal of Aquatic Science, 42, 309–323, 10.2989/16085914.2017.1405788.

Nunes, A.L., Zengeya, T.A., Hoffman, A.C., Measey, G.J. & Weyl, O.L.F., 2017b, ‘Distribution and establishment of the alien Australian redclaw crayfish, *Cherax quadricarinatus*, in South Africa and Swaziland’, PeerJ, 5, 1–21, 10.7717/peerj.3135.

Olden, J.D., Adams, J.W. & Larson, E.R., 2009, ‘First record of *Orconectes rusticus* (Girard, 1852) (Decapoda, Cambaridae) west of the great continental divide in North America’, Crustaceana 82, 1347–1351, 10.1163/156854009X448934.

Olden, J.D. & Carvalho, F.A.C., 2024, ‘Global invasion and biosecurity risk from the online trade in ornamental crayfish’, Conservation Biology, 38, e14359, 10.1111/cobi.14359.

Pârvulescu, L., Schrimpf, A., Kozubíková, E., Resino, S. C., Vrålstad, T., Petrusek, A. & Schulz, R., 2012, ‘Invasive crayfish and crayfish plague on the move: first detection of the plague agent *Aphanomyces astaci* in the Romanian Danube’, Diseases of Aquatic Organisms, 98(1), 85–94, 10.3354/dao02432.

Patoka, J., Kalous, L. & Kopecký O., 2014, ‘Risk assessment of the crayfish pet trade based on data from the Czech Republic’, Biological Invasions, 16, 2489–2494, 10.1007/s10530-014-0682-5.

Patoka, J., Magalhães, A.L.B., Kouba, A., Faulkes, Z., Jerikho, R. & Vitule, J.R.S., 2018, ‘Invasive aquatic pets: failed policies increase risks of harmful invasions,’ Biodiversity and Conservation, 27, 3037–3046, 10.1007/s10531-018-1581-3.

Perry, W.L., Feder, J.L., Dwyer, G. & Lodge, D.M., 2001, ‘Hybrid zone dynamics and species replacement between *Orconectes* crayfishes in a northern Wisconsin Lake’, Evolution 55, 1153–1166, 10.1111/j.0014-3820.2001.tb00635.x.

Phillips, I.D., Vinebrooke, R.D. & Turner, M.A., 2009, ‘Ecosystem consequences of potential range expansions of *Orconectes virilis* and *Orvconectes rusticus* crayfish in Canada – a review’, Environmental Reviews, 17, 235–248, 10.1139/A09-011.

Pöckl, M. & Pekny, R., 2002. ‘Interaction between native and alien species of crayfish in Austria: case studies’, Bulletin Français de la Pêche et de la Pisciculture 367, 763–776, 10.1051/kmae:2002064.

Putra, M.D., Bláha, M., Wardiatno, Y., Krisanti, M., Yonvitner, Jerikho, R., Kamal, M.M., Mojžišová, M., Bystřický, P.K., Kouba, A. & Kalous, L., 2018, ‘*Procambarus clarkii* (Girard, 1852) and crayfish plague as new threats for biodiversity in Indonesia’, Aquatic Conservation: Marine and Freshwater Ecosystems, 28, 1434–1440, 10.1002/aqc.2970.

Pyšek, P., Hulme, P.E, Simberloff, D., Bacher, S., Blackburn, T.M., Carlton, J.T, Dawson, W., Essl, F., Foxcroft, L.C., Genovesi, P., Jeschke, J.M., Liebhold, A.M., Mandrak, N.E., Meyerson, L.A., Pauchard, A., Pergl, J., Roy, H.E., Seebens, H., Van Kleunen, M., Vilà, M., Wingfield, M.J. & Richardson, D.M., 2020, ‘Scientists’ warning on invasive alien species’, Biological Reviews, 95, 1511–1534, 10.1111/brv.12627.

Republic of South Africa (RSA), 2004, National Environmental Management: Biodiversity Act 10 of 2004, Proc. R47/Government Gazette No. 26887/20041008.

Republic of South Africa (RSA), 2020, National Environmental Management: Biodiversity Act (Act No. 10 of 2004) Alien and Invasive Species Lists, 2020. Notice 1003 of 2020, Government Gazette No. 43726

Robinet, O., 2010, ‘Stratégie de lutte contre les espèces invasives à la Réunion’ Parc national Ile de La Réunion, France.

Rogowski, D.L. & Stockwell, C.A., 2006, ‘Assessment of potential impacts of exotic species on populations of a threatened species, White Sands Pupfish, *Cyprinodon tularosa*’, Biological Invasions, 8(1), 79–87, 10.1007/s10530-005-0238-9.

Roth, B.M., Hein, C.L. & Vander Zanden, M.J., 2006, ‘Using bioenergetics and stable isotopes to assess the trophic role of rusty crayfish (*Orconectes rusticus*) in lake littoral zones’, Canadian Journal of Fisheries and Aquatic Sciences 63, 335–344, 10.1139/f05-217.

Roy, H.E., Rabitsch, W., Scalera, R., Stewart, A., Gallardo, B., Genovesi, P., Essl, F., Adriaens, T., Bacher, S., Booy, O. & Branquart, E., 2018, ‘Developing a framework of minimum standards for the risk assessment of alien species’, Journal of Applied Ecology 55(2), 526–538, 10.1111/1365-2664.13025.

Sara, S. & El Moutaouakil, M.E., 2019, ‘Study on the spread of *Procambarus clarkii* at Gharb (Morocco) and its impact on rice growing’, *Journal of Agriculture*, Science and Technology, 9, 81–92, https://www.davidpublisher.com/index.php/Home/Article/index?id=40647.html.

Schrimpf, A., Chucholl, C., Schmidt, T., & Schulz, R., 2013, ‘Crayfish plague agent detected in populations of the invasive North American crayfish *Orconectes immunis* (Hagen, 1870) in the Rhine River, Germany’, Aquatic Invasions, 8(1), 103–109, 10.3391/ai.2013.8.1.12.

Seebens, H., Blackburn, T.M., Dyer, E., Genovesi, P., Hulme, P.E., Jeschke, J.M., Pagad, S., Pyšek, P., Winter, M., Arianoutsou, M., Dawson, W., Dullinger, S., Fuentes, N., Jager, H., Kartesz, J., Kenis, M., Kreft, H., Kuhn, I., Lezner, B., Liebhold, A., Mosena, A., Moser, D., Nishino, M., Pearman, D., Pergl, J., Rabitsch, W., Rojas-Sandoval, J., Roques, A., Rorke, S., Rossinelli, S., Roy, H.E., Scalera, R., Schindler, S., Stajerova, K., Tokarska-Gozik, B., Van Kleunen, M., Walker, K., Weigelt, P., Yamanaka, T. & Essl, F., 2017, ‘No saturation in the accumulation of alien species worldwide’, Nature Communications 8, 14435, 10.1038/ncomms14435.

Shaw, P.C., Larson, E.R. & Billman, E.J., 2021, ‘Invasion of Virile Crayfish *Faxonius virilis* (Hagen 1870) in the Lower Henrys Fork Drainage, Idaho’, Northwest Science 95(1), 106–113, 10.3955/046.095.0107.

Sinclair, J.S., Stringham, O.C., Udell, B., Mandrak, N.E., Leung, B., Romagosa, C.M. & Lockwood, J.L., 2021, ‘The international vertebrate pet trade network and insights from US imports of exotic pets’, Bioscience, 71(9), 977–990, 10.1093/biosci/biab056.

South, J., Madzivanzira, T.C., Tshali, N., Measey, J. & Weyl, O.L.F., 2020, ‘In a Pinch: Mechanisms behind potential biotic resistance toward two invasive crayfish by native Africa freshwater crabs’, Frontiers in Ecology and Evolution 8, 72, 10.3389/fevo.2020.00072.

Souty-Grosset, C, Manuel, P., Aquiloni, L., Banha, F., Choquer, J., Chucholl, C. & Tricarico, E., 2016, ‘The red swamp crayfish *Procambarus clarkii* in Europe: impacts on aquatic ecosystems and human well-being’, Limnologica 58, 78–93, 10.1016/j.limno.2016.03.003.

Souty-Grosset, C., Holdich, D.M., Noel, P.Y., Reynolds, J.D. & Haffner, P. (eds), 2006. Atlas of crayfish in Europe, Paris, France: Muséum national d’Histoire naturelle, 187 pp.

Tavakol, S., Blair, D., Morgan, J.A.T., Halajian, A. & Luus-Powell, W.J., 2021, ‘Molecular characterization of two Australian temnocephalans (Temnocephalida, Platyhelminthes) introduced with alien crayfish (Parastacidae, Decapoda) into South Africa’, Aquaculture Research, 52, 4613–4618, 10.1111/are.15294.

Todd, S. & D’Andrea, M., 2003, ‘Alien crayfish invasion of Jamaican rivers’, Crayfish News 25, 17–18.

Twardochleb, L.A, Olden, J.D. & Larson, E.R., 2013, ‘A global meta-analysis of the ecological impacts of nonnative crayfish’, Freshwater Science 32, 1367–1382, 10.1899/12-203.1

Van Wilgen, B.W. & Wilson, J.R. (eds), 2018, ‘The status of biological invasions and their management in South Africa in 2017’, South African National Biodiversity Institute, Kirstenbosch and DST-NRF Centre of Excellence for Invasion Biology, Stellenbosch.

Van Wilgen, B.W., Zengeya, T.A. & Richardson, D.M., 2022, ‘A review of the impacts of invasive alien species in South Africa’, Biological Invasions, 24, 27–50, 10.1007/s10530-021-02623-3.

Vega-Villasante, F., Ávalos-Aguilar, J.J., Nolasco-Soria, H., Vargas-Ceballos, M.A., Bortolini-Rosales, J.L., Chong-Carrillo, O., Ruiz-Núñez, M.F. & Morales-Hernández, J.C., 2015, ‘Wild populations of the invasive Australian red claw crayfish *Cherax quadricarinatus* (Crustacea, Decapoda) near the northern coast of Jalisco, Mexico: a new fishing and profitable resource’, Latin American Journal of Aquatic Research 43, 781–785, 10.3856/vol43-issue4-fulltext-17.

Vezi, M.S., Downs, C.T. & Zengeya, T.A., 2024, ‘Ornamental fish in the South African pet shop trade: potential risk to natural aquatic ecosystems’, Biological Invasions, 26, 3031–3047, 10.1007/s10530-024-03349-8.

Weinländer, M. & Füreder, L., 2009, ‘The continuing spread of *Pacifastacus leniusculus* in Carinthia (Austria)’, Knowledge and Management of Aquatic Ecosystems 17, 394–395, 10.1051/kmae/20010011.

Westman, K., Savolainen, R. & Julkunen, M., 2002, ‘Replacement of the native crayfish *Astacus astacus* by the introduced species *Pacifastacus leniusculus* in a small, enclosed Finnish lake: a 30-year study’, Ecography 25, 53–73, 10.1034/j.1600-0587.2002.250107.x.

Weyl, O.L.F., Ellender, B., Wassermann, R.J., Truter, M., Dalu, T., Zengeya T.A. & Smit, N.J., 2020, ‘Alien freshwater fauna in South Africa’, in B. Van Wilgen, J. Measey, D. Richardson, J. Wilson, T. Zengeya. (eds), Biological Invasions in South Africa, Vol. 14, Springer, Cham, 10.1007/978-3-030-32394-3_6.

Wilson, J.R., & Kumschick, S., 2024, ‘The regulation of alien species in South Africa’, South African Journal of Science, 120(5/6), 10.17159/sajs.2024/17002.

Wilson, K.A., Magnuson, J.J., Lodge, D.M., Hill, A.M., Kratz, T.K., Perry, W.L. & Willis, T.V., 2004, ‘A long-term rusty crayfish (*Orconectes rusticus*) invasion: dispersal patterns and community change in a north temperate lake’, Canadian Journal of Fisheries and Aquatic Sciences 61, 2255–2266, 10.1139/F04-170.

Zengeya, T.A. & Wilson, J.R. (eds), 2020, The status of biological invasions and their management in South Africa in 2019, South African National Biodiversity Institute, Kirstenbosch and DSI-NRF Centre of Excellence for Invasion Biology, Stellenbosch, 10.5281/zenodo.3947613.

Zengeya, T.A., Lombard, R.J.H., Nelwamondo, V.E., Nunes, A.L., Measey, J.G. & Weyl, O.L.F., 2022, ‘Trophic niche of an invasive generalist consumer: Australian redclaw crayfish, *Cherax quadricarinatus*, in the Inkomati River Basin, South Africa’, Austral Ecology, 47, 1480–1494, 10.1111/aec.13230.

