## Supplement 1 for "Applying standardised methods to assess the potential risks of alien freshwater crayfish introductions to South Africa"

#### 1.1 Risk analysis report for smooth marron (*Cherax cainii*)

| | |
|---|---|
| **Taxon:** *Cherax cainii* Austin and Ryan, 2002 | **Area*:*** South Africa |
| **Picture of *Taxon***<br>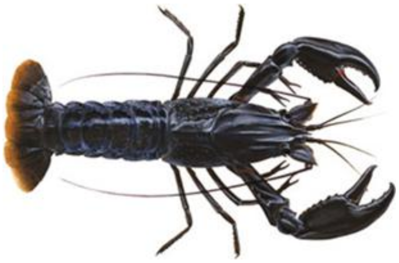<br>Photo: http://www.fish.wa.gov.au/Species/<br>Marron/Pages/default.aspx | **Alien distribution map**<br>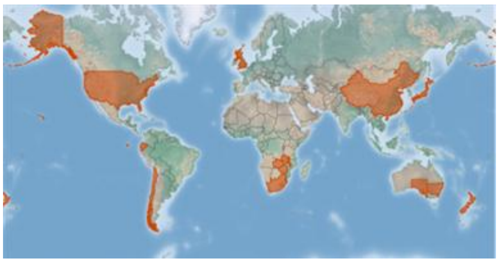<br>Reference: CABI, 2021. *Cherax cainii* www.cabi.org/isc. |
| **Risk Assessment summary:**<br>*Cherax cainii* was described in 2002 after it was distinguished genetically from *C. tenuimanus*. Records of introductions to South Africa prior to 2002 therefore only refer to *C. tenuimanus*. However, given that *C. cainii* is more widespread globally and is more commonly used in aquaculture, it is widely assumed that all imports were of *C. cainii*. There are no known naturalised populations of *Cherax* in South Africa though there are historical reports of *Cherax* outside of captivity. In the absence of evidence of impact by invasions of *Cherax* species, potential impacts of *C. cainii* were inferred from *Procambarus clarkii*—a crayfish with similar traits. Invasions of *P. clarkii* have caused major impacts on native communities through competition and predation leading to decreased abundance and local extirpation of native species. *Procambarus clarkii* has also caused habitat loss and modification through intensive grazing and stalk cutting of aquatic macrophytes that provide food, refuge and spawning sites for fish and other aquatic fauna. | **Risk score:**<br>High |
| **Management options summary:**<br>*Cherax* are farmed in South Africa, although only a few tonnes of 'smooth marron' are produced each year (data from FAO). It is assumed here that the 'smooth marron' used for farming is *C. cainii*. However, up until the end of 2020, non-research permits have only been issued for *C. tenuimanus* (research permits have been issued for both taxa). Crayfish invasions can be controlled using biocides, though, as for all aquatic systems, the use of chemical control would need to be carefully considered given the potential for non-target impacts. | **Ease of management:**<br>Medium |
| **Recommendations:**<br>First, there is a need to genetically verify that *C. cainii* is indeed present in the country. If *C. cainii* is not present, it should be delisted and any future imports | **Listing under NEM:BA A&IS Lists of 2020:** |

| | |
|---|---|
| subject to an import application and an accompanying risk analysis. If *C. cainii* is present, given the apparent on-going farming of crayfish and the lack of invasive populations, there does not seem to be a strong case to change the listing from category 2. However, it will be important to evaluate whether the benefits of farming *C. cainii* justify the additional the risk of invasion. Wherever permits are issued, it is important that strong biosecurity protocols are in place, and provision is included to deal with any escapes with the carefully regulated use of biocides as a potential method to extirpate localised populations. The current prohibition on catch and release seems appropriate. | 2 as *Cherax cainii* (Austin & Ryan, 2002), with catch and release prohibited |
| | **Recommended listing category:** 2 as *Cherax cainii* Austin & Ryan, 2002, with catch and release prohibited |

#### 1. Background

| **BAC1 Name of assessor(s)** | |
|---|---|
| Name of lead assessor | Lee-Anne Botha |
| Additional assessor (1) | Tsungai Zengeya |
| Additional assessor (2) | Christian Chimimba |
| Lead assessor | Organisational affiliation: Department of Zoology and Entomology, University of Pretoria and The South African National Biodiversity Institute (SANBI). |
| | |
| Additional assessor (1) | Organisational affiliation: South African National Biodiversity Institute (SANBI) |
| | |
| Additional assessor (2) | Organisational affiliation: Department of Zoology and Entomology, University of Pretoria and The South African National Biodiversity Institute (SANBI). |
| | |
| Comments: none | |

| **BAC4 Scientific name of *Taxon* under assessment** | |
|---|---|
| Taxon name: *Cherax cainii* | Authority: Austin and Ryan, 2002 |
| Comments: *Cherax cainii* was originally described as *Cherax tenuimanus* by Smith in 1912, from populations in the Margaret River in Western Australia (Austin & Ryan 2002). Examination of marron populations from different river systems in Australia, however, revealed two genetically distinct marron species, *C. tenuimanus* that is restricted to the Margaret River, Western Australia, and *C. cainii* which is widespread within south-west of Western Australia (Austin & Ryan 2002). Notably, *C. cainii* has been extensively introduced outside of its native range within Australia for aquaculture in South Australia and Victoria (Morrissy 1976; Austin & Ryan 2002). | |
| References:<br>Austin, C.M. & Ryan, S.G., 2002, 'Allozyme evidence for a new species of freshwater crayfish of genus *Cherax* Erichson (Decapoda: Parastacidae) form the south-west of Western Australia', *Invertebrate Systematics* 16, 357–367, https://doi.org/10.1071/IT01010. | |

| **BAC5 Synonym(s) considered** |
|---|
| Synonyms: None |
| Comments: *Cherax cainii* and *C. tenuimanus* were initially classified as one species, thus literature records prior to 2002 could be referring to either one of the two species (Austin & Ryan 2002). For this risk analysis, information on both species was used. |
| References:<br>Austin. C.M. & Ryan, S.G., 2002, 'Allozyme evidence for a new species of freshwater crayfish of genus *Cherax* Erichson (Decapoda: Parastacidae) form the south-west of Western Australia', *Invertebrate Systematics* 16, 357–367, https://doi.org/10.1071/IT01010. |

| **BAC6 Common name(s) considered** |
|---|
| Common names: smooth marron/marron |
| Comments: *Cherax cainii* has two common names: smooth marron and marron (Austin & Ryan 2002; Beatty et al. 2004). |
| References:<br>Austin, C.M. & Ryan, S.G., 2002, 'Allozyme evidence for a new species of freshwater crayfish of genus *Cherax* Erichson (Decapoda: Parastacidae) form the south-west of Western Australia', *Invertebrate Systematics* 16, 357–367, https://doi.org/10.1071/IT01010.<br>Beatty, S.J., Morgan, D.L. & Gill, H.S., 2004, 'Biology of a translocated population of the large freshwater crayfish, *Cherax cainii* Austin & Ryan, 2002 in a Western Australian River', *Crustaceana* 77,1329–1351, http://www.jstor.org/stable/20107445. |

<table>
<tr><td colspan="2">BAC7 What is the native range of the Taxon?</td></tr>
<tr><td>Response: South-west of Western Australia</td><td>Confidence: High</td></tr>
<tr><td colspan="2">Comments: Cherax cainii is native to the South-west of Western Australia (Austin & Ryan 2002; Beatty et al. 2004).</td></tr>
<tr><td colspan="2">References:<br>Austin, C.M. & Ryan, S.G., 2002, ‘Allozyme evidence for a new species of freshwater crayfish of genus Cherax Erichson (Decapoda: Parastacidae) form the south-west of Western Australia’, Invertebrate Systematics 16, 357–367, https://doi.org/10.1071/IT01010.<br>Beatty, S.J., Morgan, D. L. & Gill, H. S., 2004, ‘Biology of a translocated population of the large freshwater crayfish, Cherax cainii Austin & Ryan, 2002 in a Western Australian River’, Crustaceana 77, 1329–1351, http://www.jstor.org/stable/20107445.</td></tr>
<tr><td colspan="2">BAC8 What is the global alien range of the Taxon?</td></tr>
<tr><td>Response: Australia, Chile, China, Japan, New Zealand, South Africa, the U.K., the U.S.A, Zambia, and Zimbabwe (Nunes et al. 2017), see Appendix BAC8a, b.</td><td>Confidence: Medium</td></tr>
<tr><td colspan="2">Comments: Mostly due to commercial aquaculture and recreational fishing industries, C. cainii has been translocated to different parts of Australia (Western Australia, South Australia, Queensland, Victoria and New South Wales (Bryant & Papas 2007)). Globally, C. cainii has been introduced for aquaculture in several countries, but there is no evidence of naturalised populations in any of the areas of introduction (Lawrence & Jones 2002; Beatty et al. 2004; CABI 2019).</td></tr>
<tr><td colspan="2">References:<br>Beatty, S.J., Morgan, D.L. & Gill, H.S., 2004, ‘Biology of a translocated population of the large freshwater crayfish, Cherax cainii Austin & Ryan, 2002 in a Western Australian River’, Crustaceana 77: 1329–1351, http://www.jstor.org/stable/20107445.<br>Lawrence, C & Jones, C., 2002, ‘Cherax’, in D.M. Holdich (ed.), Biology of freshwater crayfish, Blackwell Science, U.K<br>Bryant, D. & Papas, P., 2007, ‘Marron Cherax cainii (Austin) in Victoria – a literature review’, Arthur Rylah Institute for Environmental Research Technical Report Series No. 167, Department of Sustainability and Environment, Heidelberg.<br>CABI, 2019, ‘Invasive species compendium’, from https://www.cabi.org/isc/datasheet/89136.</td></tr>
<tr><td colspan="2">BAC9 Geographic scope = the Area under consideration</td></tr>
<tr><td colspan="2">Area of assessment: South Africa</td></tr>
<tr><td colspan="2">Comments: Geographic scope of assessment is limited to South Africa.</td></tr>
<tr><td colspan="2">BAC10 Is the Taxon present in the Area?</td></tr>
<tr><td>Response: Yes</td><td>Confidence: Medium</td></tr>
<tr><td colspan="2">Comments: Cherax cainii is likely to be present in South Africa, because i) this species is widely utilised for aquaculture and more widespread globally, whereas C. tenuimanus seems to be largely restricted to the Margaret River, Australia (Austin & Ryan 2002); and ii) there are import permits records that indicate that marron was introduced for aquaculture in the country starting in the 1970s (Nunes et al. 2017). Over the years it has been introduced to aquaculture farms in KwaZulu Natal, Free State, North West Province and the Eastern and Western Cape (Nunes et al, 2017; Madzivanzira et al, 2020) although more recently it has been reported that the only active facilities are now restricted to the Eastern Cape (Hinrichsen, 2019). However, there is still uncertainty as to whether the species being farmed are C. cainii or C. tenuimanus. This needs to be genetically verified (Austin & Ryan 2002; de Moor 2002; Nunes et al. 2017). Specimens of Cherax have previously been found outside of captivity in South Africa, but currently there are no known naturalised populations (Nunes et al. 2017; Zengeya & Wilson 2020).</td></tr>
<tr><td colspan="2">References:<br>Austin, C.M. & Ryan, S.G., 2002, ‘Allozyme evidence for a new species of freshwater crayfish of genus Cherax Erichson (Decapoda: Parastacidae) form the south-west of Western Australia’, Invertebrate Systematics 16, 357–367, https://doi.org/10.1071/IT01010.</td></tr>
</table>

de Moor, I., 2002, 'Potential impacts of alien freshwater crayfish in South Africa', *African Journal of Aquatic Science* 27, 125–139, https://doi.org/10.2989/16085914.2002.9626584.

Hinrichsen, E. 2019. Biodiversity risk and benefit assessment for Smooth Marron (*Cherax cainii*) and Hairy Marron (*Cherax tenuimanus*) in South Africa. Report prepared for the Department of Environment, Forestry and Fisheries. Appendix C – 1.5 BRBA. 79pp.

Madzivanzira, T.C., South, J., Wood, L.E., Nunes, A.L. & Weyl, O.L.F., 2020, 'A review of freshwater crayfish introductions in Africa', *Reviews in Fisheries Science & Aquaculture*, 1-21 https://doi.org/10.1080/23308249.2020.1802405.

Nunes, A.L., Zengeya, T.A., Measey, G.J. & Weyl, O.L.F., 2017, 'Freshwater crayfish invasions in South Africa: past, present and potential future', *African Journal of Aquatic Science* 42, 309-323, https://doi.org/10.2989/16085914.2017.1405788.

Zengeya, T.A. & Wilson, J.R., 2020, 'The status of biological invasions and their management in South Africa in 2019', South African National Biodiversity Institute, Kirstenbosch and DSI-NRF Centre of Excellence for Invasion Biology, Stellenbosch.

**BAC11 Availability of physical specimen**

| | |
|---|---|
| Response: No | Confidence in ID: |

Herbarium or museum accession number:

References:

**BAC12 Is the *Taxon* native to the *Area* or part of the *Area*?**

| | | |
|---|---|---|
| The *Taxon* is native to (part of) the *Area*. | No | Confidence: High |
| The *Taxon* is alien in (part of) the *Area*. | Yes | Confidence: High |

Comments: *Cherax cainii* is native to Australia and there are no freshwater crayfish species native to South Africa (de Moor 2002; Nunes et al. 2017).

References:

de Moor, I., 2002, 'Potential impacts of alien freshwater crayfish in South Africa', *African Journal of Aquatic Science* 27, 125–139, , https://doi.org/10.2989/16085914.2002.9626584.

Nunes, A.L., Zengeya, T.A., Measey, G.J. & Weyl, O.L.F., 2017, 'Freshwater crayfish invasions in South Africa: past, present and potential future', *African Journal of Aquatic Science* 42, 309–323, https://doi.org/10.2989/16085914.2017.1405788.

**BAC13 What is the *Taxon*'s introduction status in the *Area*?**

| | | |
|---|---|---|
| The *Taxon* is in cultivation/containment. | Yes | Confidence: Medium |
| The *Taxon* is present outside of cultivation/containment. | No | Confidence: Medium |
| The *Taxon* has established/naturalised. | No | Confidence: Medium |
| The *Taxon* is invasive. | No | Confidence: |

Comments: *Cherax cainii* is likely present in the country in a few aquaculture farms in the Eastern and Western Cape (Nunes et al. 2017). Specimens of *Cherax* spp. have previously been found outside of captivity in South Africa; however, there are currently no known naturalised populations (de Moor 2002; Nunes et al. 2017; Zengeya & Wilson 2020).

References:

de Moor, I., 2002, 'Potential impacts of alien freshwater crayfish in South Africa', *African Journal of Aquatic Science* 27, 125–139, , https://doi.org/10.2989/16085914.2002.9626584.

Nunes, A.L., Zengeya, T.A., Measey, G.J. & Weyl, O.L.F., 2017, 'Freshwater crayfish invasions in South Africa: past, present and potential future', *African Journal of Aquatic Science* 42, 309–323, https://doi.org/10.2989/16085914.2017.1405788.

Zengeya, T.A. & Wilson, J.R., 2020, 'The status of biological invasions and their management in South Africa in 2019', South African National Biodiversity Institute, Kirstenbosch and DSI-NRF Centre of Excellence for Invasion Biology, Stellenbosch.

**BAC14 Primary (introduction) pathways**

| | | |
|---|---|---|
| Release | None | Confidence: |

| | | |
|---|---|---|
| Escape | aquaculture | Confidence: Medium |
| Contaminant | None | Confidence: |
| Stowaway | None | Confidence: |
| Corridor | None | Confidence: |
| Unaided | None | Confidence: |
| Comments: A permit was issued to import marron in 2017 (most likely *C. cainii* though listed as *C. tenuimanus*) presumably for aquaculture in South Africa (Zengeya and Wilson 2020). *Cherax cainii* appears not to be popular in the pet trade industry (Faulkes 2015). | | |
| References:<br>de Moor, I., 2002, 'Potential impacts of alien freshwater crayfish in South Africa', *African Journal of Aquatic Science* 27: 125–139, https://doi.org/10.2989/16085914.2002.9626584.<br>Faulkes, Z., 2015, 'The global trade in crayfish as pets', *Crustacean Research* 44, 75–92.<br>Nunes, A.L., Zengeya, T.A., Measey, G.J. & Weyl, O.L.F., 2017, 'Freshwater crayfish invasions in South Africa: past, present and potential future', *African Journal of Aquatic Science* 42, 309–323, https://doi.org/10.2989/16085914.2017.1405788.<br>Zengeya, T.A. & Wilson, J.R., 2020, 'The status of biological invasions and their management in South Africa in 2019', South African National Biodiversity Institute, Kirstenbosch and DSI-NRF Centre of Excellence for Invasion Biology, Stellenbosch. | | |

**2. Likelihood**

| **LIK1 Likelihood of entry via unaided primary pathways** | |
|---|---|
| Response: Present in the country (p=1) | Confidence: Medium |
| Rationale: *Cherax cainii* is believed to be currently present in the country (de Moor 2002; Nunes et al. 2017). In terms of the likelihood of future introductions, *C. cainii* is not known to be present in neighbouring countries, but as an aquatic species capable of dispersal overland, if *C. cainii* were present, it would be difficult to stop natural dispersal (Nunes et al. 2017). | |
| References:<br><br>de Moor, I., 2002, 'Potential impacts of alien freshwater crayfish in South Africa', *African Journal of Aquatic Science* 27, 125–139, https://doi.org/10.2989/16085914.2002.9626584.<br>Nunes, A.L., Zengeya, T.A., Measey, G.J. & Weyl, O.L.F., 2017, 'Freshwater crayfish invasions in South Africa: past, present and potential future', *African Journal of Aquatic Science* 42, 309–323, https://doi.org/10.2989/16085914.2017.1405788. | |

| **LIK2 Likelihood of entry via human aided primary pathways** | |
|---|---|
| Response: Present in the country (p=1) | Confidence: Medium |
| Rationale: *Cherax cainii* is believed to be currently present in the country (de Moor 2002; Nunes et al. 2017). In terms of the likelihood of future introductions, *C. cainii* has been introduced primarily for aquaculture purposes (de Moor 2002; Nunes et al. 2017) and there might be demand for future introductions. These would be subject to regulation, though illegal introductions are possible. | |
| References:<br>de Moor, I., 2002, 'Potential impacts of alien freshwater crayfish in South Africa', *African Journal of Aquatic Science* 27, 125–139, https://doi.org/10.2989/16085914.2002.9626584.<br>Nunes, A.L., Zengeya, T.A., Measey, G.J. & Weyl, O.L.F., 2017, 'Freshwater crayfish invasions in South Africa: past, present and potential future', *African Journal of Aquatic Science* 42, 309–323, https://doi.org/10.2989/16085914.2017.1405788. | |

| **LIK3 Habitat suitability** | |
|---|---|
| Response: Fairly probable (p = 0.5) | Confidence: Medium |
| Rationale: *Cherax cainii* occurs in deep perennial rivers and prefers sandy areas in rivers, particularly where organic matter accumulates (Beatty et al. 2004). Areas susceptible to *C. cainii* in South Africa are cool permanent streams in the Highveld and in the southern and south-western Cape (de Moor 2002; Nunes et al. 2017). | |
| References:<br><br>Beatty, S.J., Morgan, D.L. & Gill, H.S., 2004, 'Biology of a translocated population of the large freshwater crayfish, *Cherax cainii* Austin & Ryan, 2002 in a Western Australian River', *Crustaceana* 77, 1329–1351, http://www.jstor.org/stable/20107445.<br>de Moor, I., 2002, 'Potential impacts of alien freshwater crayfish in South Africa', *African Journal of Aquatic Science* 27, 125–139, https://doi.org/10.2989/16085914.2002.9626584.<br>Nunes, A.L., Zengeya, T.A., Measey, G.J. & Weyl, O.L.F., 2017, 'Freshwater crayfish invasions in South Africa: past, present and potential future', *African Journal of Aquatic Science* 42, 309–323, https://doi.org/10.2989/16085914.2017.1405788. | |

| **LIK4 Climate suitability** | |
|---|---|
| Response: Fairly probable (p = 0.5) | Confidence: High |
| Rationale: In aquaculture conditions, *Cherax cainii* has a thermal tolerance range of 8-29°C, with an optimal range of 17-25°C (Bryant & Papas 2007). Growth ceases when temperature is <12.5 °C (Bryant & Papas 2007). The projected areas that are climatically suitable for *C. cainii* in South Africa are located mainly in the eastern part of the country and a few areas in the Western Cape (Nunes et al. 2017). The suitable areas were mainly restricted to upland areas of the Greater Berg, Kromme, Great Kei, Mzimvubu, uMngeni, Phongolo, Crocodile and Limpopo catchment areas (Nunes et al. 2017). | |
| References:<br>Bryant, D. & Papas, P., 2007, 'Marron *Cherax cainii* (Austin) in Victoria - a Literature review', Arthur Rylah Institute for Environmental Research, Technical Report Series 167.<br>Nunes, A.L., Zengeya, T.A., Measey, G.J. & Weyl, O.L.F., 2017, 'Freshwater crayfish invasions in South Africa: past, present and potential future', *African Journal of Aquatic Science* 42, 309–323, https://doi.org/10.2989/16085914.2017.1405788.<br>Zengeya, T.A., 'Risk assessment of *Cherax* sp in South Africa (Unpublished data)'. | |

| **LIK5 Unaided secondary (dispersal) pathways** | |
|---|---|
| Response: Unlikely (p = 0.027) | Confidence: Medium |
| Rationale: There are no naturalised populations of the species in South Africa, or in neighbouring countries, that could act as sources for secondary dispersal (de Moor 2002; Nunes et al. 2017). | |
| References:<br>de Moor, I., 2002, 'Potential impacts of alien freshwater crayfish in South Africa', *African Journal of Aquatic Science* 27, 125–139, https://doi.org/10.2989/16085914.2002.9626584.<br>Nunes, A.L., Zengeya, T.A., Measey, G.J. & Weyl, O.L.F., 2017, 'Freshwater crayfish invasions in South Africa: past, present and potential future', *African Journal of Aquatic Science* 42, 309–323, https://doi.org/10.2989/16085914.2017.1405788. | |

| **LIK6 Human aided secondary (dispersal) pathways** |
|---|

| Response: Probable (p = 1) | Confidence: Medium |
|---|---|
| Rationale: *Cherax cainii* is used for aquaculture and the potential for intentional releases, or unintentional escapes, is high (de Moor 2002; Bryant & Papas 2007; Nunes et al. 2017). | |
| References:<br><br>Bryant, D., Papas, P., 2007, 'Marron *Cherax cainii* (Austin) in Victoria – a literature review', Arthur Rylah Institute for Environmental Research Technical Report Series No. 167, Department of Sustainability and Environment, Heidelberg.<br>de Moor, I., 2002, 'Potential impacts of alien freshwater crayfish in South Africa', *African Journal of Aquatic Science* 27, 125–139, https://doi.org/10.2989/16085914.2002.9626584.<br>Nunes, A.L., Zengeya, T.A., Measey, G.J. & Weyl, O.L.F., 2017, 'Freshwater crayfish invasions in South Africa: past, present and potential future', *African Journal of Aquatic Science* 42, 309–323, https://doi.org/10.2989/16085914.2017.1405788. | |

**3. Consequences**

| **CON1 Environmental impact** | |
|---|---|
| **CON1a: Competition** | |
| Response: Data Deficient (DD) | Confidence: Medium |
| Rationale: No documented information available to assess the level of impact. | |
| References: | |
| **CON1b: Predation** | |
| Response: Data Deficient (DD) | Confidence: Medium |
| Rationale: No documented information available to assess the level of impact. | |
| References: | |
| **CON1c: Hybridisation** | |
| Response: Data Deficient (DD) | Confidence: Medium |
| Rationale: There are no native *Cherax* species in South Africa. | |
| References: | |
| **CON1d: Transmission of disease** | |
| Response: Data Deficient (DD) | Confidence: Medium |
| Rationale: No documented information available to assess the level of impact. | |
| References: | |
| **CON1e: Parasitism** | |
| Response: Data Deficient (DD) | Confidence: Medium |

| | |
|---|---|
| Rationale: No documented information available to assess the level of impact. | |
| References: | |
| **CON1f: Poisoning/toxicity** | |
| Response: Data Deficient (DD) | Confidence: Medium |
| Rationale: No documented information available to assess the level of impact. | |
| References: | |
| **CON1g: Bio-fouling or other direct physical disturbance** | |
| Response: Data Deficient (DD) | Confidence: Medium |
| Rationale: No documented information available to assess the level of impact. | |
| References: | |
| **CON1h: Grazing/herbivory/browsing** | |
| Response: Data Deficient (DD) | Confidence: Medium |
| Rationale: No documented information available to assess the level of impact. | |
| References: | |
| **CON1i: Chemical, physical or structural impact on ecosystem** | |
| Response: Data Deficient (DD) | Confidence: Medium |
| Rationale: No documented information available to assess the level of impact. | |
| References: | |
| **CON1k: Indirect impacts through interactions with other species** | |
| Response: Data Deficient (DD) | Confidence: Medium |
| Rationale: No documented information available to assess the level of impact. | |
| References: | |
| **CON1 Maximum environmental impact** | |
| Response: Data Deficient (DD) | Confidence: Medium |
| Rationale: | |
| References: | |

| |
|---|
| **CON2 Socio-economic impact** |
| **CON2a: Safety** |

| | |
|---|---|
| Response: Data Deficient (DD) | Confidence: Medium |
| Rationale: No documented information available to assess the level of impact. | |
| References: | |
| **CON2b: Material and immaterial assets** | |
| Response: Data Deficient (DD) | Confidence: Medium |
| Rationale: No documented information available to assess the level of impact. | |
| References: | |
| **CON2c: Health** | |
| Response: Data Deficient (DD) | Confidence: Medium |
| Rationale: No documented information available to assess the level of impact. | |
| References: | |
| **CON2d: Social, spiritual and cultural relations** | |
| Response: Data Deficient (DD) | Confidence: Medium |
| Rationale: No documented information available to assess the level of impact. | |
| References: | |
| **CON2 Maximum socio-economic impact** | |
| Response: Data Deficient (DD) | Confidence: Medium |
| Rationale: No documented information available to assess the level of impact. | |
| References: | |

| | |
|---|---|
| **CON3 Closely related species' environmental impact** | |
| Response: Major (MR) | Confidence: High |
| Rationale: In the absence of evidence of impact of *Cherax* species in its global introduced range, potential impacts of *C. cainii* were inferred from *Procambarus clarkii* (a crayfish with similar traits and with documented evidence of impact in its introduction range).<br><br>Direct predation and competition for food, shelter, and spawning sites have led to local extinctions and a decrease in the abundance of native amphibians and crayfish species (Cruz et al. 2006; Gherardi & Acquistapace 2007; Jackson et al. 2016). Impacts have been recorded in the Iberian Peninsula, Sweden, Italy (Gherardi & Acquistapace 2007), Japan, and U.S.A. (California) (Holdich et al. 2009; Lodge et al. 2012). The most significant impacts caused by *P. clarkii* are highlighted below.<br><br><u>Competition; Major (MR); Confidence: High</u><br>*Procambarus clarkii* is very aggressive and usually out-competes native species for shelter and spawning sites, and these often lead to reproductive interference (Cruz et al. 2006; Lodge et al. 2012). For example, in areas were *P. clarkii* has been introduced , some amphibian species (e.g., *Bufo bufo*, *B. calamita*, *Rana* sp., *Taricha torosa* and *Triturus vulgaris*) have been excluded or displaced from their natural habitats, resulting in local | |

extinctions through reproductive failure (Cruz et al. 2006; Lodge et al. 2012). Furthermore, direct competition for food has caused dietary niche constriction and declines in populations of native crabs in some areas invaded by *P. clarkii* (Jackson et al. 2016).

Predation; Major (MR); Confidence: High

In its invasive range, there is evidence that predation by *P. clarkii* can result in the local or population extinctions of at least one native species (Lodge et al. 2012; Souty-Grosset et al. 2016). For example, *P. clarkii* has been implicated in causing population declines of several species of fish and amphibians by reducing their breeding success through predation on eggs and larval amphibians (Cruz et al. 2006; Ficetola et al. 2011). Indirect effects of predation can also cause trophic cascades that can lead to changes in ecosystem structure and function. For example, predation on invertebrates can release algae from grazing pressure and lead to changes in the abundance and dominance of species in algal communities (Gherardi & Barbaresi 2008).

Transmission of disease; Major (MR); Confidence: High

*Procambarus clarkii* can harbour many pathogens, parasites, and diseases that can be transmitted to other congeneric species (Longshaw 2011; Lodge et al. 2012). There is evidence that transmission of diseases and parasites by *P. clarkii* to native species has caused local or population extinctions of at least one native species, leading to changes in community composition (Lodge et al. 2012). For example, *P. clarkii* is a vector of crayfish plague, a disease caused by the parasitic oomycete, *Aphanomyces astaci* has proven to be fatal when transferred to other crayfish species (Lodge et al. 2012). However, in South Africa, there are no native crayfish species and it is not known if transmission will occur to native decapods such as shrimps and crabs. *Procambarus clarkii* can also harbour white spot syndrome disease – a viral infections of crustaceans, and fungal pathogens such as *Batrachochytrium dendrobatids* that causes chytridiomycosis – a lethal skin infection in amphibians (Longshaw 2011; McMahon et al. 2013).

Grazing/herbivory/browsing; Moderate (MO); Confidence: Medium

Grazing by *P. clarkii* can lead to habitat loss and modification through the removal of macrophytes. Habitat loss can lead to a decline in populations of species that utilise the macrophyte stands as a food source, nesting sites, and as refugia from predation (Rosenthal et al. 2005). *Procambarus clarkii* can also cause changes to community composition through trophic cascades (Gherardi & Aquistapace 2007; Souty-Grosset et al. 2016). For example, in Lake Chozas (northwest Spain), grazing by *P. clarkii* caused a reduction in macrophyte communities and this impact cascaded up the food chain with declines in invertebrates, amphibians, and waterfowl (Rodriguez et al 2003). Grazing can also cause changes to community composition and structure, such as changing ecosystems from macrophyte-dominated areas with clear water, to turbid phytoplankton-dominated areas (Matsizaki et al. 2009). Excessive grazing can also lead to accelerated rates of important processes such as litter breakdown and decomposition (Gherardi & Aquistapace 2007; Souty-Grosset et al. 2016).

References:

Cruz, M.J., Rebelo, R. & Crespo, E.G., 2006, ‘Effects of an introduced crayfish, *Procambarus clarkii*, on the distribution of south-western Iberian amphibians in their breeding habitats’, *Ecography* 29, 329-338, https://doi.org/10.1111/j.2006.0906-7590.04333.x.

Ficetola, G.F., Siesa, M.E., Manenti, R., Bottoni, L., De Bernardi, F. & Padoa-Schioppa, E., 2011, ‘Early assessment of the impact of alien species: Differential consequences of an invasive crayfish on adult and larval amphibians’, *Diversity and Distributions* 17,1141–1151, https://doi.org/10.1111/j.1472-4642.2011.00797.x

Gherardi. F. & Acquistapace, P., 2007, ‘Invasive crayfish in Europe: The impact of *Procambarus clarkii* on the littoral community of a Mediterranean lake’, *Freshwater Biology* 52, 1249-59, https://doi.org/10.1111/j.1365-2427.2007.01760.x.

Holdich, D.M., Reynolds, J.D., Sibley, P.J., 2009, ‘A review of the ever increasing threat to European crayfish from non-indigenous crayfish species’, *Knowledge and Management of Aquatic Ecosystems* 11, 394–395, https://doi.org/10.1051/kmae/2009025.

Jackson, M.C., Grey, J., Miller, K., Britton, J.R. & Donohue, I., 2016, 'Dietary niche constriction when invaders meet natives: Evidence from freshwater decapods', *Journal of Animal Ecology*, **85**, 1098–1107, https://doi.org/10.1051/kmae:2005031.

Lodge, D.M., Deines, A., Gherardi, F., Yeo, D.C.J., Arcella, T., Baldridge, A.K., Branes, M.A., Chadderton, W.L., Feder, J.L., Grantz, C.A., Howard, G.W., Jerdde, C.L., Peters, J.A., Sargent, L.W., Turner, C.R., Wittmann, M.E. & Zeng, Y., 2012, 'Global introductions of crayfishes: Evaluating the impact of species invasions on ecosystem services', *Annual Review of Ecology, Evolution and Systematics* 43, 449 –472, https://doi.org/10.1146/annurev-ecolsys-111511-103919.

Longshaw, M., 2011, 'Diseases of crayfish: A review', *Journal of Invertebrate Pathology* 106, 54–70, https://doi.org/10.1016/j.jip.2010.09.013.

Matsuzaki, S.S., Usio, N., Takamura, N. & Washitani, I., 2009, 'Contrasting impacts of invasive engineers on freshwater ecosystems: An experiment and meta-analysis', *Oecologia* 158, 673–686, https://doi.org/10.1007/s00442-008-1180-1.

McMahon, T.A., Brannelly, L.A., Chatfield, M.W., Johnson, P.T., Joseph, M.B., McKenzie, V.J., Richards-Zawacki, C.L., Venesky, M.D. & Rohr, J.R., 2013, 'Chytrid fungus *Batrachochytrium dendrobatidis* has non-amphibian hosts and releases chemicals that cause pathology in the absence of infection', *Proceedings of the National Academy of Sciences of the United States of America* 110, 210–215, https://doi.org/10.1073/pnas.1200592110.

Rodríguez, C.F., Bécares, E. & Fernández-Aláez, M., 2003, 'Shift from clear to turbid phase in Lake Chozas (NW Spain) due to the introduction of American red swamp crayfish (*Procambarus clarkii*)', *Hydrobiologia* 506, 421–26, https://doi.org/10.1023/B:HYDR.0000008626.07042.87

Rosenthal, S.K., Lodge, D.M., Mavuti, K.M., Muohi, W., Ochieng, P., Mungai, B.N. & Mkoji, G.M., 2005, 'Comparing macrophyte herbivory by introduced Louisiana crayfish (*Procambarus clarkii*) (Crustacea: Cambaridea) and native Dytiscid beetles (*Cybister tripunctatus*) in Kenya, *African Journal of Aquatic Science*, 30, 157–162, https://doi.org/10.2989/16085910509503850

Souty-Grosset, C., Anastácio, P.M., Aquiloni, L., Banha, F., Choquer, J., Chucholl, C. & Tricarico, E., 2016, 'The red swamp crayfish *Procambarus clarkii* in Europe: Impacts on aquatic ecosystems and human well-being', *Limnologica* 58, 78–93, https://doi.org/10.1016/j.limno.2016.03.003.

| **CON4 Closely related species' socio-economic impact** | |
|---|---|
| Response: Moderate (MO) | Confidence: Medium |

Rationale: In the absence of evidence of impact of *Cherax* species in its global introduced range, potential impacts of *C. cainii* were inferred from *Procambarus clarkii* (a crayfish with similar traits and with documented evidence of impact in its introduction range).

In its invaded range *P.clarkii* had caused harmful impacts in agricultural fields (Anastácio et al. 2005) and has disrupted some recreational activities leading to economic loss (Gherardi et al. 2011). The most significant impacts caused by *P. clarkii* are highlighted below.

Material and immaterial assets; Moderate (MO); Confidence: Medium

*Procambarus clarkii* often inhabits agricultural fields and their burrowing activities can cause damage to infrastructure such as irrigation canals and dam walls (Lodge et al. 2012; Arce & Diéguez-Uribeondo 2015). Burrowing activities can also alter soil hydrology leading to water loss. Grazing causes crop damage and reduces yield (Anastácio et al. 2005; Arce & Diéguez-Uribeondo 2015). In Europe for example, *P. clarkii* affects rice production through field water loss, damage to rice field banks and ditches, direct consumption of rice seed and plants, and clogging of pipes ( Anastácio et al. 2005; Souty-Grosset et al. 2016).

Health; Minor (MN); Confidence: Medium

*Procambarus clarkii* can bio-accumulate toxins and metals from the environment (Gherardi et al. 2011; Alcorlo et al. 2016; Souty-Grosset et al. 2016). These pollutants are often harmful and can be transferred to higher food web levels through the consumption of affected crayfish by humans and predators such as otters, birds, and fish (Anda et al. 2001; Gherardi et al. 2011; Lodge et al. 2012).

| |
|---|
| *Procambarus clarkii* serves as vector for several parasites and diseases some of which are zoonotic, for example the parasitic fluke flatworms (*Paragonimus* spp.) that causes lung fluke disease in humans, tularemia-causing bacterium *Francisella tularensis*, rat lungworm *Angiostrongylus cantonensis* that causes meningitis, and the nematode *Gnathostoma spinigerum* that causes human gnathostomiasis (Edgerton et al. 2002; Lane et al 2009; Souty-Grosset et al. 2016).<br><br>Social, spiritual and cultural relations; Moderate (MO); Confidence: Medium<br>Rationale: In the absence of evidence of impact of *Cherax* species in its global introduced range, potential impacts of *C. tenuimanus* were inferred from *Procambarus clarkii* (a crayfish with similar traits and with documented evidence of impact in its introduction range).<br>In its invaded range *P.clarkii* had caused harmful impacts in agricultural fields (Anastácio et al. 2005) and has disrupted some recreational activities leading to economic loss (Gherardi et al. 2011). |
| References:<br>Alcorlo, P., Otero, M., Crehuet, M., Baltanás, A. & Montes, C., 2006, 'The use of the red swamp crayfish (*Procambarus clarkii* Girard) as indicator of the bioavailability of heavy metals in environmental monitoring in the River Guadiamar (SW Spain)', *Science of the Total Environment* 366, 380–390, https://doi.org/10.1016/j.scitotenv.2006.02.023.<br>Anastácio, P.M., Correia, A.M. & Menino, J.P., 2005, 'Processes and patterns of plant destruction by crayfish: effects of crayfish size and developmental stages of rice', *Archiv für Hydrobiologie* 162, 37-51, https://doi.org/10.1127/0003-9136/2005/0162-0037.<br>Anda, P., Segura del Pozo, J., Díaz García, J.M., Escudero, R.,Garcia Pena, F.J.,Lopez Velasco, M.C., Sellek, R.E., Jimenez Chillaron, M.R., Sanchez Serrano, L.P. & Martinez Navarro, J.F., 2001, 'Waterborne outbreak of tularemia associated with crayfish fishing. *Emerging Infectious Diseases* **7**, 575–82, https://doi.org/10.3201/eid0707.010740.<br>Arce, J.A. & Diéguez-Uribeondo, J., 2015, 'Structural damage caused by the invasive crayfish *Procambarus clarkii* (Girard, 1852) in rice fields of the Iberian Peninsula: A study case', *Fundamental and Applied Limnology* 186, 259-269, https://doi.org/10.1127/fal/2015/0715.<br>Edgerton, B.F., Evans, L.H., Stephens, F.J. & Overstreet, R.M., 2002, 'Synopsis of freshwater crayfish diseases and commensal organisms', *Aquaculture* 206, 57–135, https://doi.org/10.1016/S0044-8486(01)00865-1<br>Gherardi, F., Barbaresi, S., Vaselli, O. & Bencini, A., 2002, 'A comparison of trace metal accumulation in indigenous and alien freshwater macro-decapods', *Marine and Freshwater Behaviour and Physiology* 35, 179–88, https://doi.org/10.1080/10236240210000014761.<br>Lane, M.A., Barsanti, M.C., Santos, C.A., Yeung, M., Lubner, S.J. & Weil, G.J., 2009, 'Human paragonimiasis in North America following ingestion of raw crayfish', *Clinical and Infectious Diseases* 49, 55–61, https://doi.org/10.1086/605534.<br>Lodge, D.M., Deines, A., Gherardi, F., Yeo, D.C.J., Arcella, T., Baldridge, A.K., Branes, M.A., Chadderton, W.L., Feder, J.L., Grantz, C.A., Howard, G.W., Jerdde, C.L., Peters, J.A., Sargent, L.W., Turner, C.R., Wittmann, M.E. & Zeng, Y., 2012, 'Global introductions of crayfishes: Evaluating the impact of species invasions on ecosystem services'. *Annual Review of Ecology, Evolution and Systematics* 43, 449-472, https://doi.org/10.1146/annurev-ecolsys-111511-103919.<br>Souty-Grosset, C., Anastácio, P.M., Aquiloni, L., Banha, F., Choquer, J., Chucholl, C. & Tricarico, E., 2016, 'The red swamp crayfish *Procambarus clarkii* in Europe: Impacts on aquatic ecosystems and human well-being', *Limnologica* 58, 78-93, https://doi.org/10.1016/j.limno.2016.03.003. |

| **CON5 Potential impact** | |
|---|---|
| Response: Moderate (MO) | Confidence: Low |
| Rationale: In South Africa, there are no native freshwater crayfish species, but other closely related decapods such as crabs are likely to have broad habitat and trophic overlaps (de Moor 2002; Jackson et al. 2016). The outcome of such resource overlap between *C. cainii* and indigenous fauna is unknown, but given that *P. clarkii* has caused adverse impact in other areas of introduction there is a cause of concern for possible impacts of *C.* | |

*cainii* in South African river systems (de Moor 2002; Lodge et al. 2012). *Cherax cainii* is a functional omnivore and may have an impact on macroinvertebrates and macrophytes communities (de Moor 2002). Another major concern is the transmission of parasites to native decapods and other freshwater fauna because species in the *Cherax* genus are known to harbour temnocephelan worms that can be transferred to other decapods, however impacts still need to be recorded (Avenant-Oldewage 2013; Tavakol et al. 2016). Known populations of *C. cainii* in South Africa are restricted to aquaculture facilities and there are no known naturalised populations. In the event that *C. cainii* manages to escape from such facilities, it is very unlikely that it will spread rapidly due to its low tolerance to a wide range of environmental conditions (Byrant & Pappas; Nunes et al. 2017).

References:

Avenant-Oldewage, A., 1993, 'Occurrence of *Temnocephala chaeropsis* on *Cherax tenuimanus* imported into South Africa, and notes on its infestation of an indigenous crab', *South African Journal of Science* 89, 427–428.

Bryant, D. & Papas, P., 2007, 'Marron *Cherax cainii* (Austin) in Victoria – a literature review', Arthur Rylah Institute for Environmental Research Technical Report Series No. 167, Department of Sustainability and Environment, Heidelberg.

de Moor, I., 2002, 'Potential impacts of alien freshwater crayfish in South Africa', *African Journal of Aquatic Science* 27, 125–139, https://doi.org/10.2989/16085914.2002.9626584.

Jackson, M.C., Grey, J., Miller. K., Britton, J.R. & Donohue, I., 2016, 'Dietary niche constriction when invaders meet natives: Evidence from freshwater decapods', *Journal of Animal Ecology* 85, 1098–1107, https://doi.org/10.1051/kmae:2005031.

Lodge, D.M., Deines, A., Gherardi, F., Yeo, D.C.J., Arcella, T., Baldridge, A.K., Branes, M.A., Chadderton, W.L., Feder, J.L., Grantz, C.A., Howard, G.W., Jerdde, C.L., Peters, J.A., Sargent, L.W., Turner, C.R., Wittmann, M.E. & Zeng, Y., 2012, 'Global introductions of crayfishes: evaluating the impact of species invasions on ecosystem services', *Annual Review of Ecology Evolution Systematics* 43, 449–472, , https://doi.org/10.1146/annurev-ecolsys-111511-103919

Nunes, A.L., Zengeya, T.A., Measey, G.J., Weyl, O.L.F., 2017, 'Freshwater crayfish invasions in South Africa: past, present and potential future', *African Journal of Aquatic Science* 42, 309–323, https://doi.org/10.2989/16085914.2017.1405788

Tavakol, S., Luus-Powell, W.J., Smit, W.J., Baker, C., Hoffman, A. & AlHalajian, A., 2016, 'First introduction of two Australian Temnocephalan species into Africa with an alien host: Double Trouble', *Journal of Parasitology* 102, 653–658.

#### 4. Management

| **MAN1 What is the feasibility to stop future immigration?** | |
|---|---|
| Response: High | Confidence: Medium |
| Rationale: *Cherax cainii* is mainly used for aquaculture and there are no known populations in neighbouring countries (de Moor 2002; Nunes et al. 2017). However, if it were in neighbouring countries, it would be difficult to stop natural dispersal (Nunes et al. 2017). | |
| References:<br>de Moor, I., 2002, 'Potential impacts of alien freshwater crayfish in South Africa', *African Journal of Aquatic Science* 27, 125–139, https://doi.org/10.2989/16085914.2002.9626584.<br>Nunes, A.L., Zengeya, T.A., Measey, G.J. & Weyl, O.L.F., 2017, 'Freshwater crayfish invasions in South Africa: past, present and potential future', *African Journal of Aquatic Science* 42, 309–323, https://doi.org/10.2989/16085914.2017.1405788. | |

| **MAN2 Benefits of the Taxon** | |
|---|---|
| **MAN2a Socio-economic benefits of the *Taxon*** | |
| Response: High | Confidence: Low |
| Rationale: Production of crayfish in South Africa has had mixed success because of several challenges that have been encountered (Nunes et al. 2017). As a result, marron aquaculture is now restricted to a few small scale aquaculture farms in the Eastern Cape (Hinrichsen, 2019). Levels of production are small. Tonnes live weight produced of 'smooth marron' (presumably *C. cainii*): 4 in 2018; 7 in 2017; 4 in 2016; and 4 in 2015 (FAO, 2022). Nine research permits have been issued for work on *C. cainii* in South Africa as of December 2020, and two research and twenty other permits have been issued for *C. tenuimanus* (SANBI, unpublished). | |
| References:<br>FAO, 2022, 'Global Aquaculture Production Quantity (1950–2019)'. Fisheries and Aquaculture Division [online]. Rome. [Cited Tuesday, March 22nd 2022]. https://www.fao.org/fishery/en/collection/aquaculture?lang=en<br>Hinrichsen, E. 2019. Biodiversity risk and benefit assessment for Smooth Marron (*Cherax cainii*) and Hairy Marron (*Cherax tenuimanus*) in South Africa. Report prepared for the Department of Environment, Forestry and Fisheries. Appendix C – 1.5 BRBA. 79pp.<br>Madzivanzira, T.C., South, J., Wood, L.E., Nunes, A.L. & Weyl, O.L.F., 2020, 'A review of freshwater crayfish introductions in Africa', *Reviews in Fisheries Science & Aquaculture*, 1-21, https://doi.org/10.1080/23308249.2020.1802405<br>Nunes, A.L., Zengeya, T.A., Measey, G.J. & Weyl, O.L.F., 2017, 'Freshwater crayfish invasions In South Africa: past, present and potential future', *African Journal of Aquatic Science* 42, 309-323, https://doi.org/10.2989/16085914.2017.1405788<br>SANBI (unpublished) Permits issued in respect of South Africa's National Environmental Management: Biodiversity Act's Alien and Invasive Species Regulations over the period October 2014 – December 2020. doi: 10.5281/zenodo.5101758 | |
| **MAN2b Environmental benefits of the *Taxon*** | |
| Response: None | Confidence: Low |
| Rationale: No documented information available. | |
| References: | |

| MAN3 Ease of management | |
|---|---|
| **MAN3a How accessible are populations?** | |
| Response: Easy Access (0) | Confidence: Low |
| Rationale: Although there are records of permit applications at CapeNature to import *C. cainii* for aquaculture in the Western Cape (Nunes et al. 2017) marron aquaculture is apparently now restricted to a few small-scale aquaculture farms in the Eastern Cape (Hinrichsen, 2019). As such current known populations are easy to access, though if there were naturalised populations, accessibility might become more difficult. | |
| References:<br>Hinrichsen, E. 2019. Biodiversity risk and benefit assessment for Smooth Marron (*Cherax cainii*) and Hairy Marron (*Cherax tenuimanus*) in South Africa. Report prepared for the Department of Environment, Forestry and Fisheries. Appendix C – 1.5 BRBA. 79pp.<br>Nunes, A.L., Zengeya, T.A., Measey, G.J. & Weyl, O.L.F., 2017, 'Freshwater crayfish invasions in South Africa: past, present and potential future', *African Journal of Aquatic Science* 42, 309–323, https://doi.org/10.2989/16085914.2017.1405788 | |
| **MAN3b Is detectability critically time-dependent?** | |
| Response: Yes (2) | Confidence: Low |
| Rationale: *Cherax cainii*, as a freshwater species, is generally difficult to detect, particularly given the species seem to be more active at night (Bryrant & Papas 2007). | |
| References:<br>Bryant, D. & Papas, P., 2007, 'Marron *Cherax cainii* (Austin) in Victoria – a literature review', Arthur Rylah Institute for Environmental Research Technical Report Series No. 167, Department of Sustainability and Environment, Heidelberg. | |
| **MAN3c Time to reproduction** | |
| Response: 1-3 years (1) | Confidence: Low |
| Rationale: *Cherax cainii* reaches sexual maturity when two to three years (Beatty et al. 2004). | |
| References:<br>Beatty, S.J., Morgan, D.L. & Gill, H.S., 2004, 'Biology of a translocated population of the large freshwater crayfish, *Cherax cainii* Austin & Ryan, 2002 in a Western Australian River', *Crustaceana* 77, 1329–1351, https://www.jstor.org/stable/20107445. | |
| **MAN3d Propagule persistence** | |
| Response: 0 | Confidence: High |
| Rationale: Marron crayfish usually spawns in spring and can produce between 200-400 eggs per spawning event. The eggs can take up to half a year to develop inside the females. Once the eggs have been fertilised, the female carries the eggs until they hatch. Hatchlings hang under the female's tale for many weeks until they can feed on their own. | |
| References:<br><br>Department of Fisheries, Government of Australia. 2011. *Fisheries Factsheet Marron* https://www.fish.wa.gov.au/Documents/recreational_fishing/fact_sheets/fact_sheet_marron.pdf | |
| **MAN3 Ease of management** | |
| Response: Medium | Confidence: Low |

| |
|---|
| Rationale: As a fairly small, largely nocturnal, freshwater species, *C. cainii* are difficult to detect. However current populations in the Area are under permit and so finding and accessing populations is not currently difficult. |
| References:<br>de Moor, I., 2002, 'Potential impacts of alien freshwater crayfish in South Africa', *African Journal of Aquatic Science* 27, 125-139, https://doi.org/10.2989/16085914.2002.9626584.<br>Madzivanzira, T.C., South, J., Wood, L.E., Nunes, A.L. & Weyl, O.L.F., 2020, 'A review of freshwater crayfish introductions in Africa', *Reviews in Fisheries Science & Aquaculture*, 1-21, https://doi.org/10.1080/23308249.2020.1802405<br>Nunes, A.L., Zengeya, T.A., Measey, G.J. & Weyl, O.L.F., 2017, 'Freshwater crayfish invasions in South Africa: past, present and potential future', *African Journal of Aquatic Science* 42, 309-323, https://doi.org/10.2989/16085914.2017.1405788 |

| **MAN4 Has the feasibility of eradication been evaluated?** | |
|---|---|
| Response: No | Confidence: High |
| Rationale: Currently there are no known populations outside of captivity in the country (de Moor 2002; Nunes et al. 2017), and so there has been no need to look at eradication to date. | |
| References:<br>de Moor, I., 2002, 'Potential impacts of alien freshwater crayfish in South Africa', *African Journal of Aquatic Science* 27, 125-139, https://doi.org/10.2989/16085914.2002.9626584.<br>Nunes, A.L., Zengeya, T.A., Measey, G.J. & Weyl, O.L.F., 2017, 'Freshwater crayfish invasions in South Africa: past, present and potential future', *African Journal of Aquatic Science* 42, 309-323, https://doi.org/10.2989/16085914.2017.1405788. | |

| MAN5 Control options and monitoring approaches available for the *Taxon* |
|---|
| Response: In New Zealand biocides have been used to control and eradicate closely related *Cherax tenuimanus* hairy marron crayfish (Gould 2005), noting that such an approach would need to be carefully regulated given the potential for non-target impacts. |
| References:<br><br>Gould B, 2005, 'Marron Interagency collaboration follows surprise catch'. *Biosecurity* 60: 10-11 |

| MAN6 Any other management considerations to highlight? | |
|---|---|
| Response | No |

**5. Calculations**

**Likelihood = Fairly probable**

| Parameter | Likelihood | Stages | Final assessment |
|---|---|---|---|
| LIK1 | 1 | P(entry) = 1 | P (invasion) = 0.5 |
| LIK2 | 1 | | |
| LIK3 | 0.5 | P(establishment) = 0.5 | |
| LIK4 | 0.5 | | |
| LIK5 | 0.027 | P (spread) = 1 | |
| LIK6 | 1 | | |

**Consequence = MR**

| Parameter | Mechanism/sector | Response |
|---|---|---|
| CON1a | Competition | DD |
| CON1b | Predation | DD |
| CON1c | Hybridisation | DD |
| CON1d | Disease transmission | DD |
| CON1e | Parasitism | DD |
| CON1f | Poisoning/toxicity | DD |
| CON1g | Bio-fouling or other direct physical disturbance | DD |
| CON1h | Grazing/herbivory/browsing | DD |

| | | |
|---|---|---|
| CON1i | Chemical, physical, structural impact | DD |
| CON1k | Indirect impacts through interactions with other species | DD |
| CON1 | Maximum environmental impact | DD |
| CON2a | Safety | DD |
| CON2b | Material and immaterial assets | DD |
| CON2c | Health | DD |
| CON2d | Social, spiritual and cultural relations | DD |
| CON2 | Maximum socio-economic impact | DD |
| CON3 | Environmental impact of closely related taxa | MR |
| CON4 | Socio-economic impact of closely related taxa | MO |
| CON5 | Potential impact based on traits, experiments, or models | MO |

**Risk score = High**

| | | **Consequences** | | | | |
|---|---|---|---|---|---|---|
| | | MC | MN | MO | MR | MV |
| **Likelihood** | Extremely unlikely | low | low | low | medium | medium |
| | Very unlikely | low | low | low | medium | high |
| | Unlikely | low | low | medium | high | high |
| | Fairly probable | medium | medium | high | high | high |
| | Probable | medium | high | high | high | high |

**Ease of management = Medium**

| **Parameter** | **Question** | **Response** |
|---|---|---|
| MAN3a | How accessible are populations? | 0 |
| MAN3b | Is detectability critically time-dependent? | 2 |
| MAN3c | Time to reproduction | 1 |
| MAN3d | Propagule persistence | 0 |
| **MAN3** | **SUM** | 3 |

**Appendix BAC7:** Map of the records of *Cherax cainii* from Western Australia (taken here to be the native range). From GBIF (2021)—*Cherax cainii* Austin & Ryan, 2002 in GBIF Secretariat (2021). GBIF Backbone Taxonomy. Checklist dataset https://doi.org/10.15468/39omei accessed via GBIF.org on 2022-03-18.

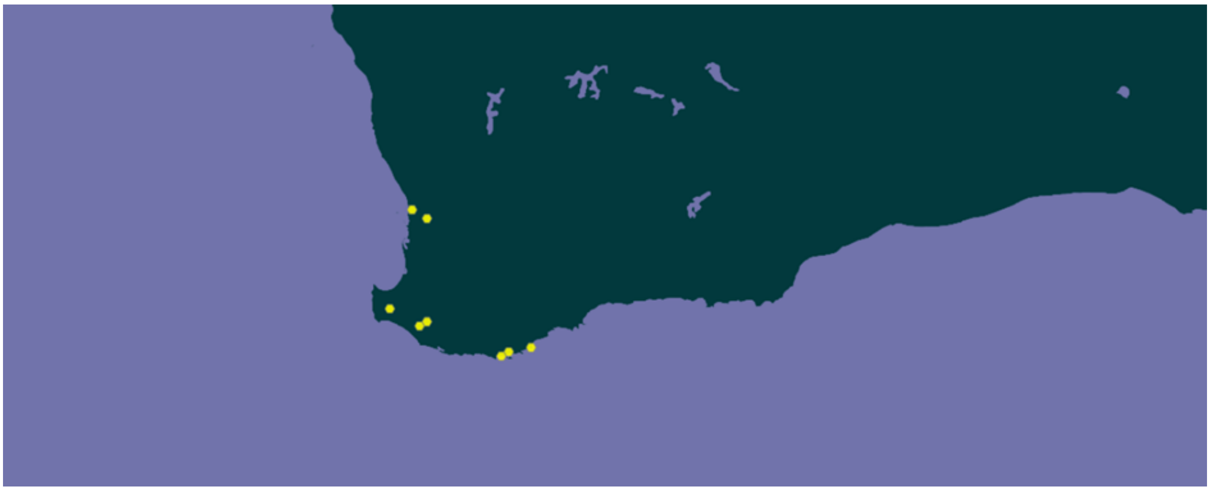

**Appendix BAC8(a)**: Map of the global alien range of *Cherax cainii* at a state level for Australia and a country level for elsewhere. Source: CABI, 2021. *Cherax cainii* [original text by Uma Sabapathy Allen]. In: Invasive Species Compendium. Wallingford, UK: CAB International. www.cabi.org/isc.

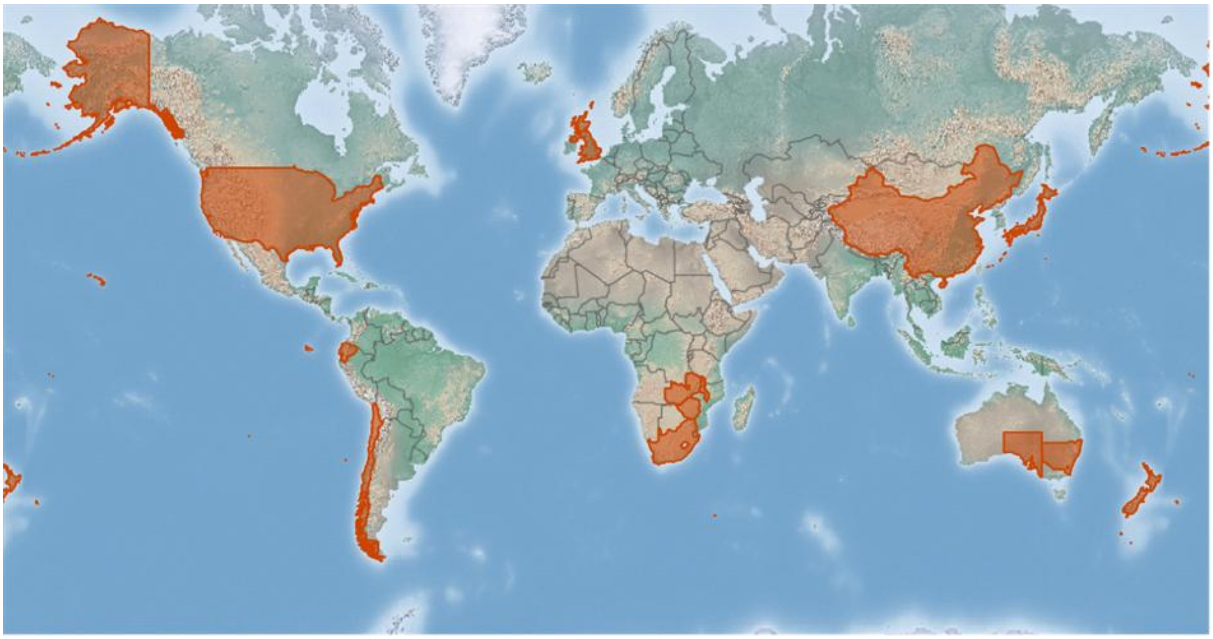

#### 1.2 Risk analysis report for hairy marron (*Cherax tenuimanus*)

| | |
|---|---|
| **Taxon:** *Cherax tenuimanus* Smith, 1912 | **Area:** South Africa |
| **Compiled by:** Lee-Anne Botha | **Conditionally Approved by:** [1] |
| **Picture of *Taxon***<br>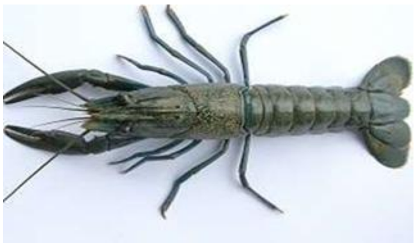<br>Photo: https://phys.org/news/2014-01-captive-hairy-marron-populace-bolstered.html | **Global distribution map**<br>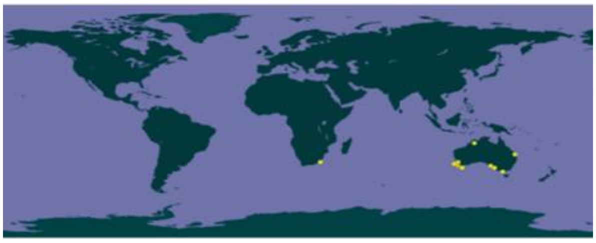<br>Reference GBIF.org (06 August 2019) https://doi.org/10.15468/dl.tvqoxp |
| **Risk Assessment summary:**<br><br>*Cherax tenuimanus* was originally described in 1912, but in 2002 the taxon was, based on a genetic analysis, split into two species: *C. tenuimanus* (that is restricted to the Margaret River in Western Australia) and *C. cainii* (that is more widespread in SW Australia). It is widely assumed that all imports to South Africa were of *C. cainii* as this species is more common around the world (including in aquaculture). There are no known naturalised populations of *Cherax* spp. in South Africa though there are historical reports of *Cherax* spp. outside of captivity. In the absence of evidence of impact by invasions of *Cherax* species, potential impacts of *C. tenuimanus* were inferred from *Procambarus clarkii*—a crayfish with similar traits. Invasions of *P. clarkii* have caused major impacts on native communities through competition and predation leading to decreased abundance and local extirpation of native species. *Procambarus clarkii* has also caused habitat loss and modification through intensive grazing and stalk cutting of aquatic macrophytes that provide food, refuge and spawning sites for fish and other aquatic fauna. | **Risk score:**<br><br>High |
| **Management options summary:**<br><br>*Cherax* are farmed in South Africa, although only a few tonnes of 'smooth marron' are produced each year (data from FAO). It is assumed here that the 'smooth marron' used for farming is *C. cainii*. However, up until the end of 2020, non-research permits have only been issued for *C. tenuimanus* (although research permits have been issued for both taxa). Crayfish invasions can be controlled using biocides, though, as for all aquatic systems, the use of chemical control would need to be carefully considered given the potential for non-target impacts. | **Ease of management:**<br><br>Medium |
| **Recommendations:**<br><br>First, there is a need to verify that *C. tenuimanus* is present in the country. If *C. tenuimanus* is not present, it should be delisted, and future import permit requests accompanied by a risk analysis. If *C. tenuimanus* is present, given the apparent | **Listing under NEM:BA A&IS lists of 2021:**<br><br>2 as *Cherax tenuimanus* (Smith, 1912), with |

[1] Within the next five years, i.e. by 2027, the presence of *C. tenuimanus* in the country should be reassessed, and an evaluation made comparing the demand for and benefits of farming to the risks of invasions. The listing might then need to be updated.

| | |
|---|---|
| on-going farming of crayfish and the lack of invasive populations, there does not seem to be a strong case to change the listing from category 2. However, it will be important to evaluate whether the benefits of farming *C. tenuimanus* justify the additional the risk of invasion. Wherever permits are issued, it is important that strong biosecurity protocols are in place, and provision is included to deal with any escapes with the carefully regulated use of biocides as a potential method to extirpate localised populations. The current prohibition on catch and release seems appropriate. | catch and release prohibited<br>**Recommended listing category:**<br>2 *Cherax tenuimanus* Smith, 1912, with catch and release prohibited |

#### 1. Background

| **BAC1 Name of assessor(s)** | |
|---|---|
| Name of lead assessor | Lee-Anne Botha |
| Additional assessor (1) | Tsungai Zengeya |
| Additional assessor (2) | Christian Chimimba |
| **BAC2 Contact details of assessor (s)** | |
| Lead assessor | Organisational affiliation: Department of Zoology and Entomology, University of Pretoria and The South African National Biodiversity Institute (SANBI). |
| | |
| Additional assessor (1) | Organisational affiliation: South African National Biodiversity Institute (SANBI) |
| | |
| Additional assessor (2) | Organisational affiliation: Department of Zoology and Entomology, University of Pretoria and The South African National Biodiversity Institute (SANBI). |
| | |
| Comments: none | |

| **BAC4 Scientific name of *Taxon* under assessment** | |
|---|---|
| *Taxon* name: *Cherax tenuimanus* | Authority: Smith, 1912 |

Comments: *Cherax tenuimanus* was originally described by Smith in 1912 from the Margaret River in Western Australia (Austin & Ryan 2002). Examination of marron populations from different river systems in Australia, however, revealed two genetically distinct marron species, *C. tenuimanus* that is restricted to the Margaret River, Western Australia, and *C. cainii* which is widespread within south-west of Western Australia (Austin & Ryan 2002). Notably, *C. cainii* has been extensively introduced outside of its native range within Australia for aquaculture in South Australia and Victoria (Morrissy 1976; Austin & Ryan 2002).

References:
Austin, C.M. & Ryan, S.G., 2002, 'Allozyme evidence for a new species of freshwater crayfish of genus *Cherax* Erichson (Decapoda: Parastacidae) from the south-west of Western Australia', *Invertebrate Systematics* 16, 357-367, https://doi.org/10.1071/IT01010.
Morrissy, N.M., 1976, 'Aquaculture of Marron, *Cherax tenuimanus* (Smith) Part 2: Breeding and Early Rearing', *Fisheries Research Bulletin of Western Australia* 17, 1-32.

**BAC5 Synonym(s) considered**

Synonyms: None

Comments: *Cherax cainii* and *C. tenuimanus* were initially classified as one species, thus literature records prior to 2002 could be referring to either one of the two species (Austin & Ryan 2002). For this risk analysis, information on both species was used.

References:
Austin, C.M. & Ryan, S.G., 2002, 'Allozyme evidence for a new species of freshwater crayfish of genus *Cherax* Erichson (Decapoda: Parastacidae) from the south-west of Western Australia', *Invertebrate Systematics* 16, 357-367, https://doi.org/10.1071/IT01010.

**BAC6 Common name(s) considered**

Common names: hairy marron/ marron

Comments: *Cherax tenuimanus* has two common names: hairy marron and marron (Austin & Ryan 2002; Beatty et al. 2004).

References:
Austin, C.M. & Ryan, S.G., 2002, 'Allozyme evidence for a new species of freshwater crayfish of genus *Cherax* Erichson (Decapoda: Parastacidae) from the south-west of Western Australia', *Invertebrate Systematics* 16, 357-367, https://doi.org/10.1071/IT01010.
Beatty, S.J., Morgan, D.L. & Gill, H.S., 2004, 'Biology of a translocated population of the large freshwater crayfish, *Cherax cainii* Austin & Ryan, 2002 in a Western Australian River', *Crustaceana* 77,1329-1351, https://www.jstor.org/stable/20107445.

**BAC7 What is the native range of the *Taxon*?**

| Response: South-west of Western Australia (the Margaret River) | Confidence: High |
|---|---|

Comments: *Cherax tenuimanus* is native to the Margaret River that is located in South-west of Western Australia (Austin & Ryan 2002; Beatty et al. 2004).

References:
Austin, C.M. & Ryan, S.G., 2002, 'Allozyme evidence for a new species of freshwater crayfish of genus *Cherax* Erichson (Decapoda: Parastacidae) form the south-west of Western Australia', *Invertebrate Systematics* 16, 357-367, https://doi.org/10.1071/IT01010.
Beatty, S.J., Morgan, D.L. & Gill, H.S., 2004, 'Biology of a translocated population of the large freshwater crayfish, *Cherax cainii* Austin & Ryan, 2002 in a Western Australian River', *Crustaceana* 77, 1329-1351, https://www.jstor.org/stable/20107445.
CABI n.d., from https://www.cabi.org/isc/datasheet/89136

**BAC8 What is the global alien range of the *Taxon*?**

| Response: Australia | Confidence: Medium |
|---|---|

Comments: *Cherax tenuimanus* is largely restricted to the upper reaches of the Margaret River, but it has been translocated to river systems within this region and other states in Australia (Austin & Ryan 2002; Lawrence & Jones 2002; Beatty et al. 2004). Prior to 2002, introduction records of marron to South Africa and several other countries outside Australia refer to *C. tenuimanus*, but these introductions need to be genetically verified as they might actually refer to *C. cainii* (Zengeya & Wilson 2020; US and Wildlife Services, 2017).

References:
Austin, C.M. & Ryan, S.G., 2002, 'Allozyme evidence for a new species of freshwater crayfish of genus *Cherax* Erichson (Decapoda: Parastacidae) form the south-west of Western Australia', *Invertebrate Systematics* 16, 357-367, https://doi.org/10.1071/IT01010.
Beatty, S.J., Morgan, D.L. & Gill, H.S., 2004, 'Biology of a translocated population of the large freshwater crayfish, *Cherax cainii* Austin & Ryan, 2002 in a Western Australian River', *Crustaceana* 77, 1329-1351, https://www.jstor.org/stable/20107445.
CABI n.d., from https://www.cabi.org/isc/datasheet/89136
Lawrence, C. & Jones, C., 2002, '*Cherax*', in D.M. Holdich (ed.), *Biology of freshwater crayfish. Blackwell Science*, U.K.
US and Wildlife Services, 2017, 'Ecological Risk Screening Summary', from https://www.fws.gov/fisheries/ans/erss/uncertainrisk/ERSS-Cherax-tenuimanus_Final.pdf.
Zengeya, T.A, & Wilson, J.R., 2020, 'The status of biological invasions and their management in South Africa in 2019', South African National Biodiversity Institute, Kirstenbosch and DSI-NRF Centre of Excellence for Invasion Biology, Stellenbosch

**BAC9 Geographic scope = the *Area* under consideration**

*Area* of assessment: South Africa

Comments: Geographic scope of assessment is limited to South Africa.

**BAC10 Is the *Taxon* present in the *Area*?**

| Response: Yes | Confidence: Medium |
|---|---|

Comments: *Cherax tenuimanus* is likely to be present in South Africa, because there are import permits records that indicate that marron was introduced for aquaculture in the country starting in the 1970s (Nunes et a. 2017). However, there is uncertainty whether the introduced species was *C. cainii* or *C. tenuimanus*.

Specimens of marron have previously been found outside of captivity in South Africa, but currently there are no known naturalised populations of *C. tenuimanus* (Nunes et al. 2017; Zengeya & Wilson 2020). A few aquaculture farms in the Eastern and Western Cape are reportedly rearing marron but the identity of the farmed species needs to be genetically verified to ascertain if it is *C. cainii* and/or *C. tenuimanus* (Austin & Ryan 2002; de Moor 2002; Nunes et al. 2017). It is assumed to be *C. cainii* because this species is widely utilised for aquaculture and more widespread globally, whereas *C. tenuimanus* seems to be largely restricted to the Margaret River, Australia (Austin & Ryan 2002).

References:

Austin, C.M. & Ryan, S.G., 2002, 'Allozyme evidence for a new species of freshwater crayfish of genus *Cherax* Erichson (Decapoda: Parastacidae) form the south-west of Western Australia', *Invertebrate Systematics* 16, 357-367, https://doi.org/10.1071/IT01010.

Beatty, S.J., Morgan, D.L. & Gill, H.S., 2004, 'Biology of a translocated population of the large freshwater crayfish, *Cherax cainii* Austin & Ryan, 2002 in a Western Australian River', *Crustaceana* 77, 1329-1351, https://www.jstor.org/stable/20107445.

de Moor, I., 2002, 'Potential impacts of alien freshwater crayfish in South Africa', *African Journal of Aquatic Science* 27, 125-139, https://doi.org/10.2989/16085914.2002.9626584.

Nunes, A.L., Zengeya, T.A., Measey, G.J. & Weyl, O.L.F., 2017, 'Freshwater crayfish invasions in South Africa: past, present and potential future', *African Journal of Aquatic Science* 42, 309-323, https://doi.org/10.2989/16085914.2017.1405788.

Zengeya, T.A., Wilson, J.R., 2020, 'The status of biological invasions and their management in South Africa in 2019', South African National Biodiversity Institute, Kirstenbosch and DSI-NRF Centre of Excellence for Invasion Biology, Stellenbosch.

**BAC11 Availability of physical specimen**

| | |
|---|---|
| Response: Yes | Confidence in ID: Low |

Herbarium or museum accession number: GEN 833A. Species was collected in South Africa, Arigata Fish Farm near Marina Beach (-30.9300000000, 30.27000000) and recorded as *Cherax tenuimanus.*

References:

Albany Museum Grahamstown, South Africa

GBIF n.d., from https://www.gbif.org/occurrence/1299981578

**BAC12 Is the *Taxon* native to the *Area* or part of the *Area*?**

| | | |
|---|---|---|
| The *Taxon* is native to (part of) the *Area*. | No | Confidence: High |
| The *Taxon* is alien in (part of) the *Area*. | Yes | Confidence: High |

Comments: *Cherax tenuimanus* is native to Australia and there are no freshwater crayfish native to South Africa (Austin & Ryan 2002, de Moor 2002, Nunes et al. 2017).

References:

Austin, C.M. & Ryan, S.G., 2002, 'Allozyme evidence for a new species of freshwater crayfish of genus *Cherax* Erichson (Decapoda: Parastacidae) form the south-west of Western Australia', *Invertebrate Systematics* 16, 357-367, https://doi.org/10.1071/IT01010.

de Moor, I., 2002, 'Potential impacts of alien freshwater crayfish in South Africa', *African Journal of Aquatic Science* 27, 125-139, https://doi.org/10.2989/16085914.2002.962658.

Nunes, A.L., Zengeya, T.A., Measey, G.J. & Weyl, O.L.F., 2017, 'Freshwater crayfish invasions in South Africa: past, present and potential future', *African Journal of Aquatic Science* 42, 309-323, https://doi.org/10.2989/16085914.2017.1405788

**BAC13 What is the *Taxon*'s introduction status in the *Area*?**

| | | |
|---|---|---|
| The *Taxon* is in cultivation/containment. | Yes | Confidence: Medium |
| The *Taxon* is present outside of cultivation/containment. | No | Confidence: Medium |
| The *Taxon* has established/naturalised. | No | Confidence: Medium |
| The *Taxon* is invasive. | No | Confidence: Low |

Comments: The presence of *C. tenuimanus* in the country needs to be verified. While there are import permit records for the introduction of *C. tenuimanus* for aquaculture purposes and crayfish are in cultivation, it is likely these are largely *C. cainii*. Specimens of *Cherax* spp. have previously been found outside of captivity in South Africa; however, there are currently no known naturalised populations (de Moor 2002; Nunes et al. 2017; Zengeya & Wilson 2020).

References:

de Moor, I., 2002, 'Potential impacts of alien freshwater crayfish in South Africa', *African Journal of Aquatic Science* 27, 125-139, https://doi.org/10.2989/16085914.2002.962658.

Nunes, A.L., Zengeya, T.A., Measey, G.J. & Weyl, O.L.F., 2017, 'Freshwater crayfish invasions in South Africa: past, present and potential future', *African Journal of Aquatic Science* 42, 309-323, https://doi.org/10.2989/16085914.2017.1405788.

Zengeya, T.A. & Wilson, J.R., 2020, 'The status of biological invasions and their management in South Africa in 2019', South African National Biodiversity Institute, Kirstenbosch and DSI-NRF Centre of Excellence for Invasion Biology, Stellenbosch.

**BAC14 Primary (introduction) pathways**

| | | |
|---|---|---|
| Release | None | Confidence: |
| Escape | aquaculture | Confidence: High |
| Contaminant | None | Confidence: |
| Stowaway | None | Confidence: |
| Corridor | None | Confidence: |
| Unaided | None | Confidence: |

Comments: A permit was issued to import marron in 2017 (most likely *C. cainii* though listed as *C. tenuimanus*) presumably for aquaculture in South Africa (Zengeya and Wilson 2020). *Cherax cainii* appears not to be popular in the pet trade industry (Faulkes 2015).

References:

de Moor, I., 2002, 'Potential impacts of alien freshwater crayfish in South Africa', *African Journal of Aquatic Science* 27, 125-139, https://doi.org/10.2989/16085914.2002.962658.

Faulkes, Z., 2015, 'The global trade in crayfish as pets', *Crustacean Research* 44, 75–92. https://doi.org/10.18353/crustacea.44.0_75

Nunes, A.L., Zengeya, T.A., Measey, G.J. & Weyl, O.L.F., 2017, 'Freshwater crayfish invasions in South Africa: past, present and potential future', *African Journal of Aquatic Science* 42, 309-323, https://doi.org/10.2989/16085914.2017.1405788

Zengeya, T.A. & Wilson, J.R., 2020, 'The status of biological invasions and their management in South Africa in 2019', South African National Biodiversity Institute, Kirstenbosch and DSI-NRF Centre of Excellence for Invasion Biology, Stellenbosch.

#### 2. Likelihood

**LIK1 Likelihood of entry via unaided primary pathways**

| | |
|---|---|
| Response: Present in the country (p=1) | Confidence: Medium |

Rationale: *Cherax tenuimanus* is probably currently present in the country (de Moor 2002; Nunes et al. 2017), but this needs to be verified. In terms of the likelihood of future introductions, *C. tenuimanus* is not known to be present in neighbouring countries, but as an aquatic species capable of dispersal overland, if *C. tenuimanus* were present, it would be difficult to stop natural dispersal (Nunes et al. 2017).

References:

CABI n.d., from https://www.cabi.org/isc/datasheet/89136.

de Moor, I., 2002, 'Potential impacts of alien freshwater crayfish in South Africa', *African Journal of Aquatic Science* 27, 125-139, https://doi.org/10.2989/16085914.2002.962658.

| |
|---|
| Zengeya, T.A. & Wilson, J.R., 2020, 'The status of biological invasions and their management in South Africa in 2019', South African National Biodiversity Institute, Kirstenbosch and DSI-NRF Centre of Excellence for Invasion Biology, Stellenbosch. |

| **LIK2 Likelihood of entry via human aided primary pathways** | |
|---|---|
| Response: Present in the country (p=1) | Confidence: Medium |
| Rationale: *Cherax tenuimanus* is probably currently present in the country (de Moor 2002; Nunes et al. 2017), but this needs to be verified. In terms of the likelihood of future introductions, *C. tenuimanus* has been introduced primarily for aquaculture purposes (de Moor 2002; Nunes et al. 2017) and there might be demand for future introductions. These would be subject to regulation, though illegal introductions are possible. | |
| References:<br><br>de Moor, I., 2002, 'Potential impacts of alien freshwater crayfish in South Africa', *African Journal of Aquatic Science* 27, 125-139, https://doi.org/10.2989/16085914.2002.962658.<br>Nunes, A.L., Zengeya, T.A., Measey, G.J. & Weyl, O.L.F., 2017, 'Freshwater crayfish invasions in South Africa: past, present and potential future', *African Journal of Aquatic Science* 42, 309-323, https://doi.org/10.2989/16085914.2017.1405788<br>Zengeya, T.A. & Wilson, J.R., 2020, 'The status of biological invasions and their management in South Africa in 2019', South African National Biodiversity Institute, Kirstenbosch and DSI-NRF Centre of Excellence for Invasion Biology, Stellenbosch. | |

| **LIK3 Habitat suitability** | |
|---|---|
| Response: Fairly probable (p = 0.5) | Confidence: Low |
| Rationale: Little is known of how the ecology and habitat requirements differ between the hairy and smooth marron species (Department of the Environment 2013). Marron occurs in deep perennial rivers and prefers sandy areas in rivers particularly where organic matter accumulates (Beatty et al. 2004). *Cherax tenuimanus* requires structural diversity for shelter and refuge (de Moor 2002) and relatively good water quality, and may struggle to persist in disturbed habitats (Department of the Environment 2013). Areas in South Africa susceptible to *C. cainii* are cool permanent streams in the Highveld and in the southern and south-western Cape (Nunes et al. 2017). It is assumed here that *C. tenuimanus* would have similar habitat requirements, but the confidence is placed at low. | |
| References:<br><br>Department of the Environment, 2013, *Approved Conservation Advice for Cherax tenuimanus (hairy marron)*, Canberra, Department of the Environment, from http://www.environment.gov.au/biodiversity/threatened/species/pubs/78931-conservation-advice.pdf<br>Beatty, S.J., Morgan, D.L. & Gill, H.S., 2004, 'Biology of a translocated population of the large freshwater crayfish, *Cherax cainii* Austin & Ryan, 2002 in a Western Australian River', *Crustaceana* 77, 1329-1351, https://www.jstor.org/stable/20107445.<br>de Moor, I., 2002, 'Potential impacts of alien freshwater crayfish in South Africa', *African Journal of Aquatic Science* 27, 125-139, https://doi.org/10.2989/16085914.2002.9626584.<br>Nunes, A.L., Zengeya, T.A. & Measey, G.J. & Weyl, O.L.F., 2017, 'Freshwater crayfish invasions in South Africa: past, present and potential future', *African Journal of Aquatic Science* 42, 309-323, https://doi.org/10.2989/16085914.2017.1405788. | |

| **LIK4 Climate suitability** | |
|---|---|
| Response: Fairly probable (p = 0.5) | Confidence: Medium |

| |
|---|
| Rationale: In aquaculture conditions, *Cherax tenuimanus* has a thermal tolerance range of 8-29°C with an optimal range of 17-25°C (Department of Agriculture, Landform and Rural Development, 2018). Growth ceases when temperature is <12.5 °C (Department of Agriculture, Landform and Rural Development, 2018). The projected areas that are climatically suitable for marron in South Africa (predicted using *C. cainii*) are located mainly in the eastern part of the country and a few areas in the Western Cape (Nunes et al. 2017). The suitable areas were mainly restricted to upland areas of the Greater Berg, Kromme, Great Kei, Mzimvubu, uMngeni, Phongolo, Crocodile and Limpopo catchment areas (Nunes et al. 2017). |
| References:<br>Department of Agriculture, Landform and Rural Development, viewed 10 August 2018, from https://www.nda.agric.za/doadev/sidemenu/fisheries/03_areasofwork/Aquaculture/BIODIVERSITY/C%20%20tenuimanus%20final%20BRBA.pdf<br>Nunes, A.L., Zengeya, T.A. & Measey, G.J. & Weyl, O.L.F., 2017, 'Freshwater crayfish invasions in South Africa: past, present and potential future', *African Journal of Aquatic Science* 42, 309-323, https://doi.org/10.2989/16085914.2017.1405788. |

| **LIK5 Unaided secondary (dispersal) pathways** | |
|---|---|
| Response: Unlikely (p = 0.027) | Confidence: Medium |
| Rationale: There are no naturalised populations in South Africa or neighbouring countries that could act as sources for secondary dispersal (de Moor 2002; Nunes et al. 2017). | |
| References:<br>de Moor, I., 2002, 'Potential impacts of alien freshwater crayfish in South Africa', *African Journal of Aquatic Science* 27, 125-139, https://doi.org/10.2989/16085914.2002.9626584.<br>Nunes, A.L., Zengeya, T.A. & Measey, G.J. & Weyl, O.L.F., 2017, 'Freshwater crayfish invasions in South Africa: past, present and potential future', *African Journal of Aquatic Science* 42, 309-323, https://doi.org/10.2989/16085914.2017.1405788. | |

| **LIK6 Human aided secondary (dispersal) pathways** | |
|---|---|
| Response: Probable (p = 1) | Confidence: Medium |
| Rationale: *Cherax tenuimanus* is only used for aquaculture in South Africa and the potential for intentional release, or unintentional escape, is high (de Moor 2002; Nunes et al. 2017). | |
| References:<br>Bryant, D. & Papas, P., 2007, 'Marron *Cherax cainii* (Austin) in Victoria – a literature review', Arthur Rylah Institute for Environmental Research Technical Report Series No. 167, Department of Sustainability and Environment, Heidelberg.<br>de Moor, I., 2002, 'Potential impacts of alien freshwater crayfish in South Africa', *African Journal of Aquatic Science* 27, 125-139, https://doi.org/10.2989/16085914.2002.9626584.<br>Nunes, A.L., Zengeya, T.A. & Measey, G.J. & Weyl, O.L.F., 2017, 'Freshwater crayfish invasions in South Africa: past, present and potential future', *African Journal of Aquatic Science* 42, 309-323, https://doi.org/10.2989/16085914.2017.1405788. | |

**3. Consequences**

| **CON1 Environmental impact** | |
|---|---|
| **CON1a: Competition** | |
| Response: Data Deficient (DD) | Confidence: Medium |
| Rationale: No documented information available to assess the level of impact. | |
| References: | |
| **CON1b: Predation** | |
| Response: Data Deficient (DD) | Confidence: Medium |
| Rationale: No documented information available to assess the level of impact. | |
| References: | |
| **CON1c: Hybridisation** | |
| Response: Data Deficient (DD) | Confidence: Medium |
| Rationale: There are no native *Cherax* species in South Africa. | |
| References: | |
| **CON1d: Transmission of disease** | |
| Response: Data Deficient (DD) | Confidence: Medium |
| Rationale: No documented information available to assess the level of impact. | |
| References: | |
| **CON1e: Parasitism** | |
| Response: Data Deficient (DD) | Confidence: Medium |
| Rationale: No documented information available to assess the level of impact. | |
| References: | |
| **CON1f: Poisoning/toxicity** | |
| Response: Data Deficient (DD) | Confidence: Medium |
| Rationale: No documented information available to assess the level of impact. | |
| References: | |
| **CON1g: Bio-fouling or other direct physical disturbance** | |
| Response: Data Deficient (DD) | Confidence: Medium |
| Rationale: No documented information available to assess the level of impact. | |

| | |
|---|---|
| References: | |
| **CON1h: Grazing/herbivory/browsing** | |
| Response: Data Deficient (DD) | Confidence: Medium |
| Rationale: No documented information available to assess the level of impact. | |
| References: | |
| **CON1i: Chemical, physical or structural impact on ecosystem** | |
| Response: Data Deficient (DD) | Confidence: Medium |
| Rationale: No documented information available to assess the level of impact. | |
| References: | |
| **CON1k: Indirect impacts through interactions with other species** | |
| Response: Data Deficient (DD) | Confidence: Medium |
| Rationale: No documented information available to assess the level of impact. | |
| References: | |
| **CON1 Maximum environmental impact** | |
| Response: Data Deficient (DD) | Confidence: Medium |
| Rationale: No documented information available to assess the level of impact | |
| References: | |

| | |
|---|---|
| **CON2 Socio-economic impact** | |
| **CON2a: Safety** | |
| Response: Data Deficient (DD) | Confidence: Medium |
| Rationale: No documented information available to assess the level of impact. | |
| References: | |
| **CON2b: Material and immaterial assets** | |
| Response: Data Deficient (DD) | Confidence: Medium |
| Rationale: No documented information available to assess the level of impact. | |
| References: | |
| **CON2c: Health** | |
| Response: Data Deficient (DD) | Confidence: Medium |

| | |
|---|---|
| Rationale: No documented information available to assess the level of impact. | |
| References: | |
| **CON2d: Social, spiritual and cultural relations** | |
| Response: Data Deficient (DD) | Confidence: Medium |
| Rationale: | |
| References: | |
| **CON2 Maximum socio-economic impact** | |
| Response: Data Deficient (DD) | Confidence: Medium |
| Rationale: No documented information available to assess the level of impact. | |
| References: | |

| | |
|---|---|
| **CON3 Closely related species' environmental impact** | |
| Response: Major (MR) | Confidence: High |

Rationale: In the absence of evidence of impact of *Cherax* species in its global introduced range, potential impacts of *C. tenuimanus* were inferred from *Procambarus clarkii* (a crayfish with similar traits and with documented evidence of impact in its introduction range).

Direct predation and competition for food, shelter, and spawning sites have led to local extinctions and a decrease in the abundance of native amphibians and crayfish species (Cruz et al. 2006; Gherardi & Acquistapace 2007; Jackson et al. 2016). Impacts have been recorded in the Iberian Peninsula, Sweden, Italy (Gherardi & Acquistapace 2007), Japan, and U.S.A. (California) (Holdich et al. 2009; Lodge et al. 2012). The most significant impacts caused by *P. clarkii* are highlighted below.

Competition; Major (MR); Confidence: High
*Procambarus clarkii* is very aggressive and usually out-competes native species for shelter and spawning sites, and these often lead to reproductive interference (Cruz et al. 2006; Lodge et al. 2012). For example, in areas were *P. clarkii* has been introduced , some amphibian species (e.g., *Bufo bufo*, *B. calamita*, *Rana* sp., *Taricha torosa* and *Triturus vulgaris*) have been excluded or displaced from their natural habitats, resulting in local extinctions through reproductive failure (Cruz et al. 2006; Lodge et al. 2012). Furthermore, direct competition for food has caused dietary niche constriction and declines in populations of native crabs in some areas invaded by *P. clarkii* (Jackson et al. 2016).

Predation; Major (MR); Confidence: High
In its invasive range, there is evidence that predation by *P. clarkii* can result in the local or population extinctions of at least one native species (Lodge et al. 2012; Souty-Grosset et al. 2016). For example, *P. clarkii* has been implicated in causing population declines of several species of fish and amphibians by reducing their breeding success through predation on eggs and larval amphibians (Cruz et al. 2006; Ficetola et al. 2011). Indirect effects of predation can also cause trophic cascades that can lead to changes in ecosystem structure and function. For example, predation on invertebrates can release algae from grazing pressure and lead to changes in the abundance and dominance of species in algal communities (Gherardi & Barbaresi 2008).

Transmission of disease; Major (MR); Confidence: High
*Procambarus clarkii* can harbour many pathogens, parasites, and diseases that can be transmitted to other congeneric species (Longshaw 2011; Lodge et al. 2012). There is evidence that transmission of diseases and

parasites by *P. clarkii* to native species has caused local or population extinctions of at least one native species, leading to changes in community composition (Lodge et al. 2012). For example, *P. clarkii* is a vector of crayfish plague, a disease caused by the parasitic oomycete, *Aphanomyces astaci* has proven to be fatal when transferred to other crayfish species (Lodge et al. 2012). However, in South Africa, there are no native crayfish species and it is not known if transmission will occur to native decapods such as shrimps and crabs. *Procambarus clarkii* can also harbour white spot syndrome disease – a viral infections of crustaceans, and fungal pathogens such as *Batrachochytrium dendrobatids* that causes chytridiomycosis – a lethal skin infection in amphibians (Longshaw 2011; McMahon et al. 2013).

Grazing/herbivory/browsing; Moderate (MO); Confidence: Medium

Grazing by *P. clarkii* can lead to habitat loss and modification through the removal of macrophytes. Habitat loss can lead to a decline in populations of species that utilise the macrophyte stands as a food source, nesting sites, and as refugia from predation (Rosenthal et al. 2005). *Procambarus clarkii* can also cause changes to community composition through trophic cascades (Gherardi & Aquistapace 2007; Souty-Grosset et al. 2016). For example, in Lake Chozas (northwest Spain), grazing by *P. clarkii* caused a reduction in macrophyte communities and this impact cascaded up the food chain with declines in invertebrates, amphibians, and waterfowl (Rodriguez et al 2003). Grazing can also cause changes to community composition and structure, such as changing ecosystems from macrophyte-dominated areas with clear water, to turbid phytoplankton-dominated areas (Matsizaki et al. 2009). Excessive grazing can also lead to accelerated rates of important processes such as litter breakdown and decomposition (Gherardi & Aquistapace 2007; Souty-Grosset et al. 2016).

References:

Cruz, M.J., Rebelo, R. & Crespo, E.G., 2006, 'Effects of an introduced crayfish, *Procambarus clarkii*, on the distribution of south-western Iberian amphibians in their breeding habitats', *Ecography* 29, 329-338, https://doi.org/10.1111/j.2006.0906-7590.04333.x

Ficetola, G.F., Siesa, M.E., Manenti, R., Bottoni, L., De Bernardi, F. & Padoa-Schioppa, E., 2011, 'Early assessment of the impact of alien species: Differential consequences of an invasive crayfish on adult and larval amphibians', *Diversity and Distributions* 17,1141–1151, https://doi.org/10.1111/j.1472-4642.2011.00797.x

Gherardi. F. & Acquistapace, P., 2007, 'Invasive crayfish in Europe: The impact of *Procambarus clarkii* on the littoral community of a Mediterranean lake', *Freshwater Biology* 52, 1249-59, https://doi.org/10.1111/j.1365-2427.2007.01760.x.

Holdich, D.M., Reynolds, J.D., Sibley, P.J., 2009, 'A review of the ever increasing threat to European crayfish from non-indigenous crayfish species', *Knowledge and Management of Aquatic Ecosystems* 11, 394–395, https://doi.org/10.1051/kmae/2009025.

Jackson, M.C., Grey, J., Miller, K., Britton, J.R. & Donohue, I., 2016, 'Dietary niche constriction when invaders meet natives: Evidence from freshwater decapods', *Journal of Animal Ecology*, **85**, 1098–1107, https://doi.org/10.1051/kmae:2005031.

Lodge, D.M., Deines, A., Gherardi, F., Yeo, D.C.J., Arcella, T., Baldridge, A.K., Branes, M.A., Chadderton, W.L., Feder, J.L., Grantz, C.A., Howard, G.W., Jerdde, C.L., Peters, J.A., Sargent, L.W., Turner, C.R., Wittmann, M.E. & Zeng, Y., 2012, 'Global introductions of crayfishes: Evaluating the impact of species invasions on ecosystem services', *Annual Review of Ecology, Evolution and Systematics* 43, 449 –472, https://doi.org/10.1146/annurev-ecolsys-111511-103919.

Longshaw, M., 2011, 'Diseases of crayfish: A review', *Journal of Invertebrate Pathology* 106, 54–70, https://doi.org/10.1016/j.jip.2010.09.013.

Matsuzaki, S.S., Usio, N., Takamura, N. & Washitani, I., 2009, 'Contrasting impacts of invasive engineers on freshwater ecosystems: An experiment and meta-analysis', *Oecologia* 158, 673–686, https://doi.org/10.1007/s00442-008-1180-1

McMahon, T.A., Brannelly, L.A., Chatfield, M.W., Johnson, P.T., Joseph, M.B., McKenzie, V.J., Richards-Zawacki, C.L., Venesky, M.D. & Rohr, J.R., 2013, 'Chytrid fungus *Batrachochytrium dendrobatidis* has non-amphibian hosts and releases chemicals that cause pathology in the absence of infection',

*Proceedings of the National Academy of Sciences of the United States of America* 110, 210–215, https://doi.org/10.1073/pnas.1200592110

Rodríguez, C.F., Bécares, E. & Fernández-Aláez, M., 2003, 'Shift from clear to turbid phase in Lake Chozas (NW Spain) due to the introduction of American red swamp crayfish (*Procambarus clarkii*)', *Hydrobiologia* 506, 421–26, https://doi.org/10.1023/B:HYDR.0000008626.07042.87.

Rosenthal, S.K., Lodge, D.M., Mavuti, K.M., Muohi, W., Ochieng, P., Mungai, B.N. & Mkoji, G.M., 2005, 'Comparing macrophyte herbivory by introduced Louisiana crayfish (*Procambarus clarkii*) (Crustacea: Cambaridea) and native Dytiscid beetles (*Cybister tripunctatus*) in Kenya, *African Journal of Aquatic Science*, 30, 157–162, https://doi.org/10.2989/16085910509503850.

Souty-Grosset, C., Anastácio, P.M., Aquiloni, L., Banha, F., Choquer, J., Chucholl, C. & Tricarico, E., 2016, 'The red swamp crayfish *Procambarus clarkii* in Europe: Impacts on aquatic ecosystems and human well-being', *Limnologica* 58, 78–93, https://doi.org/10.1016/j.limno.2016.03.003.

**CON4 Closely related species' socio-economic impact**

| Response: Moderate (MO) | Confidence: Medium |
|---|---|

Rationale: In the absence of evidence of impact of *Cherax* species in its global introduced range, potential impacts of *C. tenuimanus* were inferred from *Procambarus clarkii* (a crayfish with similar traits and with documented evidence of impact in its introduction range). In its invaded range *P.clarkii* had caused harmful impacts in agricultural fields (Anastácio et al. 2005) and has disrupted some recreational activities leading to economic loss (Gherardi et al. 2011). The most significant impacts caused by *P. clarkii* are highlighted below.

Material and immaterial assets; Moderate (MO); Confidence: Medium

*Procambarus clarkii* often inhabits agricultural fields and their burrowing activities can cause damage to infrastructure such as irrigation canals and dam walls (Lodge et al. 2012; Arce & Diéguez-Uribeondo 2015). Burrowing activities can also alter soil hydrology leading to water loss. Grazing causes crop damage and reduces yield (Anastácio et al. 2005; Arce & Diéguez-Uribeondo 2015). In Europe for example, *P. clarkii* affects rice production through field water loss, damage to rice field banks and ditches, direct consumption of rice seed and plants, and clogging of pipes ( Anastácio et al. 2005; Souty-Grosset et al. 2016).

Health; Minor (MN); Confidence: Medium

*Procambarus clarkii* can bio-accumulate toxins and metals from the environment (Gherardi et al. 2011; Alcorlo et al. 2016; Souty-Grosset et al. 2016). These pollutants are often harmful and can be transferred to higher food web levels through the consumption of affected crayfish by humans and predators such as otters, birds, and fish (Anda et al. 2001; Gherardi et al. 2011; Lodge et al. 2012).

*Procambarus clarkii* serves as vector for several parasites and diseases some of which are zoonotic, for example the parasitic fluke flatworms (*Paragonimus* spp.) that causes lung fluke disease in humans, tularemia-causing bacterium *Francisella tularensis*, rat lungworm *Angiostrongylus cantonensis* that causes meningitis, and the nematode *Gnathostoma spinigerum* that causes human gnathostomiasis (Edgerton et al. 2002; Lane et al 2009; Souty-Grosset et al. 2016).

Social, spiritual and cultural relations; Moderate (MO); Confidence: Medium

Rationale: In the absence of evidence of impact of *Cherax* species in its global introduced range, potential impacts of *C. tenuimanus* were inferred from *Procambarus clarkii* (a crayfish with similar traits and with documented evidence of impact in its introduction range).

In its invaded range *P.clarkii* had caused harmful impacts in agricultural fields (Anastácio et al. 2005) and has disrupted some recreational activities leading to economic loss (Gherardi et al. 2011).

References:

Alcorlo, P., Otero, M., Crehuet, M., Baltanás, A. & Montes, C., 2006, 'The use of the red swamp crayfish (*Procambarus clarkii* Girard) as indicator of the bioavailability of heavy metals in environmental monitoring in the River Guadiamar (SW Spain)', *Science of the Total Environment* 366, 380–390, https://doi.org/10.1016/j.scitotenv.2006.02.023.

Anastácio, P.M., Correia, A.M. & Menino, J.P., 2005, 'Processes and patterns of plant destruction by crayfish: effects of crayfish size and developmental stages of rice', *Archiv für Hydrobiologie* 162, 37-51, https://doi.org/10.1127/0003-9136/2005/0162-0037.

Anda, P., Segura del Pozo, J., Díaz García, J.M., Escudero, R.,Garcia Pena, F.J.,Lopez Velasco, M.C., Sellek, R.E., Jimenez Chillaron, M.R., Sanchez Serrano, L.P. & Martinez Navarro, J.F., 2001, 'Waterborne outbreak of tularemia associated with crayfish fishing. *Emerging Infectious Diseases* **7**, 575–82, https://doi.org/10.3201/eid0707.010740.

Arce, J.A. & Diéguez-Uribeondo, J., 2015, 'Structural damage caused by the invasive crayfish *Procambarus clarkii* (Girard, 1852) in rice fields of the Iberian Peninsula: A study case', *Fundamental and Applied Limnology* 186, 259-269, https://doi.org/10.1127/fal/2015/0715.

Edgerton, B.F., Evans, L.H., Stephens, F.J. & Overstreet, R.M., 2002, 'Synopsis of freshwater crayfish diseases and commensal organisms', *Aquaculture* 206, 57–135, https://doi.org/10.1016/S0044-8486(01)00865-1.

Gherardi, F., Barbaresi, S., Vaselli, O. & Bencini, A., 2002, 'A comparison of trace metal accumulation in indigenous and alien freshwater macro-decapods', *Marine and Freshwater Behaviour and Physiology* 35, 179–88, https://doi.org/10.1080/1023624021000014761.

Lane, M.A., Barsanti, M.C., Santos, C.A., Yeung, M., Lubner, S.J. & Weil, G.J., 2009, 'Human paragonimiasis in North America following ingestion of raw crayfish', *Clinical and Infectious Diseases* 49, 55–61, https://doi.org/10.1086/605534.

Lodge, D.M., Deines, A., Gherardi, F., Yeo, D.C.J., Arcella, T., Baldridge, A.K., Branes, M.A., Chadderton, W.L., Feder, J.L., Grantz, C.A., Howard, G.W., Jerdde, C.L., Peters, J.A., Sargent, L.W., Turner, C.R., Wittmann, M.E. & Zeng, Y., 2012, 'Global introductions of crayfishes: Evaluating the impact of species invasions on ecosystem services'. *Annual Review of Ecology, Evolution and Systematics* 43, 449-472, https://doi.org/10.1146/annurev-ecolsys-111511-103919.

Souty-Grosset, C., Anastácio, P.M., Aquiloni, L., Banha, F., Choquer, J., Chucholl, C. & Tricarico, E., 2016, 'The red swamp crayfish *Procambarus clarkii* in Europe: Impacts on aquatic ecosystems and human well-being', *Limnologica* 58, 78-93, https://doi.org/10.1016/j.limno.2016.03.003

| **CON5 Potential impact** | |
|---|---|
| Response: Moderate (MO) | Confidence: Low |

Rationale: In South Africa, there are no freshwater crayfish species but other closely related decapods such as crabs are likely to have broad habitat and trophic overlaps (Jackson et al. 2016). The outcome of such resource overlap between *C. tenuimanus* and indigenous fauna is unknown, but given that *P. clarkii* has caused adverse impact in other areas of introduction there is a cause of concern for possible impacts of *C. tenuimanus* in South African river systems (Lodge et al. 2012). Another major concern is the transmission of parasites for example, temnocephala worms to native decapods and other freshwater fauna (Avenant-Oldewage 1993, Tavakol et al. 2016). The potential for hybridisation in South Africa is very unlikely as there are no native freshwater crayfish (de Moor 2002). There are no known naturalised populations, although, should *C. tenuimanus* be present in South Africa, it is most likely confined to aquaculture facilities (Zengeya & Wilson 2019). In the event that *C. tenuimanus* manages to escape from such facilities, it's very unlikely that it will spread rapidly due to its low tolerance to a wide range of environmental conditions (Bryant & Pappas 2007).

References:

Avenant-Oldewage, A., 1993, 'Occurrence of *Temnocephala chaeropsis* on *Cherax tenuimanus* imported into South Africa, and notes on its infestation of an indigenous crab', *South African Journal of Science* 89, 427–428.

Bryant, D. & Papas, P., 2007, 'Marron *Cherax cainii* (Austin) in Victoria – a literature review', Arthur Rylah Institute for Environmental Research Technical Report Series No. 167, Department of Sustainability and Environment, Heidelberg.

de Moor, I., 2002, ‘Potential impacts of alien freshwater crayfish in South Africa’, African Journal of Aquatic Science 27, 125-139, https://doi.org/10.2989/16085914.2002.9626584.
Jackson, M.C., Grey, J., Miller, K., Britton, J.R. & Donohue, I., 2016, ‘Dietary niche constriction when invaders meet natives: Evidence from freshwater decapods’, *Journal of Animal Ecology* 85, 1098–1107, https://doi.org/10.1051/kmae:2005031.
Lodge, D.M., Deines, A., Gherardi, F., Yeo, D.C.J., Arcella, T., Baldridge, A.K., Branes, M.A., Chadderton, W.L., Feder, J.L., Grantz, C.A., Howard, G.W., Jerdde, C.L., Peters, J.A., Sargent, L.W., Turner, C.R., Wittmann, M.E. & Zeng, Y., 2012, ‘Global introductions of crayfishes: evaluating the impact of species invasions on ecosystem services’, *Annual Review of Ecology Evolution Systematics* 43, 449-72, , https://doi.org/10.1146/annurev-ecolsys-111511-103919
Tavakol, S., Luus-Powell, W.J., Smit, W.J., Baker, C., Hoffman, A. & AlHalajian, A., 2016, ‘First introduction of two Australian Temnocephalan species into Africa with an alien host: Double Trouble’, *Journal of Parasitology* 102, 653–658.
Zengeya, T.A. & Wilson, J.R., 2020, ‘The status of biological invasions and their management in South Africa in 2019’, South African National Biodiversity Institute, Kirstenbosch and DSI-NRF Centre of Excellence for Invasion Biology, Stellenbosch.

**4. Management**

| **MAN1 What is the feasibility to stop future immigration?** | |
|---|---|
| Response: High | Confidence: Medium |
| Rationale: *Cherax tenuimanus* was purportedly introduced for aquaculture in South Africa but the identity of the introduced populations still needs to be verified. There are no known populations in neighbouring countries (Burgess 2007; de Moor 2002; Nunes et al. 2017). However, if *C. tenuimanus* were in neighbouring countries it would be difficult to stop natural dispersal (Nunes et al. 2017). | |
| References:<br><br>Burgess, M., 2007, ‘Pioneers of SA marron production’, Farmer's Weekly Magazine, from http://www.farmersweekly.co.za/article.aspx?id=520&h=Pioneers-of-SA-marron-production<br>de Moor, I., 2002, ‘Potential impacts of alien freshwater crayfish in South Africa’, *African Journal of Aquatic Science* 27, 125-139, , https://doi.org/10.2989/16085914.2002.9626584.<br>Nunes, A.L., Zengeya, T.A., Measey, G.J. & Weyl, O.L.F., 2017, ‘Freshwater crayfish invasions in South Africa: past, present and potential future’, *African Journal of Aquatic Science* 42, 309-323, https://doi.org/10.2989/16085914.2017.1405788 | |

| **MAN2 Benefits of the Taxon** | |
|---|---|
| **MAN2a Socio-economic benefits of the *Taxon*** | |
| Response: High | Confidence: Low |
| Rationale: Production of crayfish in South Africa has had mixed success because of several challenges that have been encountered (Nunes et al. 2017). As a result, marron aquaculture is now restricted to a few small scale aquaculture farms in the Eastern Cape (Hinrichsen, 2019). Levels of production are small. Tonnes live weight produced of ‘smooth marron’ (presumably *C. cainii*): 4 in 2018; 7 in 2017; 4 in 2016; and 4 in 2015 (FAO, 2022). Nine research permits have been issued for work on *C. cainii* in South Africa as of December 2020, and two research and twenty other permits have been issued for *C. tenuimanus* (SANBI, unpublished). | |
| References: | |

FAO, 2022, 'Global Aquaculture Production Quantity (1950–2019)'. Fisheries and Aquaculture Division [online]. Rome. [Cited Tuesday, March 22nd 2022]. https://www.fao.org/fishery/en/collection/aquaculture?lang=en

Hinrichsen, E. 2019. Biodiversity risk and benefit assessment for Smooth Marron (*Cherax cainii*) and Hairy Marron (*Cherax tenuimanus*) in South Africa. Report prepared for the Department of Environment, Forestry and Fisheries. Appendix C – 1.5 BRBA. 79pp.

Madzivanzira, T.C., South, J., Wood, L.E., Nunes, A.L. & Weyl, O.L.F., 2020, 'A review of freshwater crayfish introductions in Africa', *Reviews in Fisheries Science & Aquaculture*, 1-21 https://doi.org/10.1080/23308249.2020.1802405

Nunes, A.L., Zengeya, T.A., Measey, G.J. & Weyl, O.L.F., 2017, 'Freshwater crayfish invasions In South Africa: past, present and potential future', *African Journal of Aquatic Science* 42, 309-323, https://doi.org/10.2989/16085914.2017.1405788

SANBI (unpublished) Permits issued in respect of South Africa's National Environmental Management: Biodiversity Act's Alien and Invasive Species Regulations over the period October 2014 – December 2020. doi: 10.5281/zenodo.5101758

| **MAN2b Environmental benefits of the *Taxon*** | |
|---|---|
| Response: None | Confidence: Medium |
| Rationale: No documented information available. | |
| References: | |

| **MAN3 Ease of management** | |
|---|---|
| **MAN3a How accessible are populations?** | |
| Response: Easy Access (0) | Confidence: Low |
| Rationale: There are records of permit applications at CapeNature to import *C. cainii* for aquaculture in the Western Cape (Nunes et al. 2017). A follow up study needs to be undertaken to check if farms are still in production and to ascertain the identity of the species utilised (de Moor 2002; Burgess 2007; Nunes et al. 2017). As such current known populations are easy to access, though if there were naturalised populations, accessibility might become more difficult. | |
| References:<br>Burgess, M., 2007, 'Pioneers of SA marron production', Farmer's Weekly Magazine, from https://www.farmersweekly.co.za/animals/aquaculture/pioneers-of-sa-marron-production/<br>de Moor, I., 2002, 'Potential impacts of alien freshwater crayfish in South Africa', *African Journal of Aquatic Science* 27, 125-139, , https://doi.org/10.2989/16085914.2002.9626584.<br>Nunes, A.L., Zengeya, T.A., Measey, G.J. & Weyl, O.L.F., 2017, 'Freshwater crayfish invasions in South Africa: past, present and potential future', *African Journal of Aquatic Science* 42, 309-323, https://doi.org/10.2989/16085914.2017.1405788. | |
| **MAN3b Is detectability critically time-dependent?** | |
| Response: Yes (2) | Confidence: Low |
| Rationale: *Cherax cainii*, as a freshwater species, is generally difficult to detect, particularly given the species seem to be more active at night (Bryrant & Papas 2007), the same would be expected for *C. tenuimanus*. | |
| References: | |

Bryant, D. & Papas, P., 2007, ‘Marron *Cherax cainii* (Austin) in Victoria – a literature review’, Arthur Rylah Institute for Environmental Research Technical Report Series No. 167, Department of Sustainability and Environment, Heidelberg.

**MAN3c Time to reproduction**

| Response: 1-3 years (1) | Confidence: Low |
|---|---|

Rationale: Marron crayfish reaches sexual maturity when two to three years (Beatty et al. 2004).

References:

Beatty, S.J., Morgan, D.L. & Gill, H.S., 2004, ‘Biology of a translocated population of the large freshwater crayfish, *Cherax cainii* Austin & Ryan, 2002 in a Western Australian River’, *Crustaceana* 77, 1329–1351 https://www.jstor.org/stable/20107445.

**MAN3d Propagule persistence**

| Response: 0 | Confidence: High |
|---|---|

Rationale: Marron crayfish usually spawns in spring and can produce between 200-400 eggs per spawning event. The eggs can take up to half a year to develop inside the females. Once the eggs have been fertilised, the female carries the eggs until they hatch. Hatchlings hang under the female's tale for many weeks until they can feed on their own.

References:

Department of Fisheries, Government of Australia. 2011. *Fisheries Factsheet Marron* https://www.fish.wa.gov.au/Documents/recreational_fishing/fact_sheets/fact_sheet_marron.pdf

**MAN3 Ease of management**

| Response: Medium | Confidence: Low |
|---|---|

Rationale: As a fairly small, largely nocturnal, freshwater species, *C. tenuimanus* are difficult to detect. However current populations in the Area are under permit and so finding and accessing populations is not currently difficult.

References:

de Moor, I., 2002, ‘Potential impacts of alien freshwater crayfish in South Africa’, *African Journal of Aquatic Science* 27, 125-139, , https://doi.org/10.2989/16085914.2002.9626584.

Madzivanzira, T.C., South, J., Wood, L.E., Nunes, A.L. & Weyl, O.L.F., 2020, ‘A review of freshwater crayfish introductions in Africa’, *Reviews in Fisheries Science & Aquaculture*, 1-21, https://doi.org/10.1080/23308249.2020.1802405

Nunes, A.L., Zengeya, T.A., Measey, G.J. & Weyl, O.L.F., 2017, ‘Freshwater crayfish invasions in South Africa: past, present and potential future’, *African Journal of Aquatic Science* 42, 309-323, https://doi.org/10.2989/16085914.2017.1405788.

**MAN4 Has the feasibility of eradication been evaluated?**

| Response: No | Confidence: Low |
|---|---|

Rationale: Currently there are no known populations outside of captivity in the country (de Moor 2002; Nunes et al. 2017), and so there has been no need to look at eradication to date.

| |
|---|
| References:<br><br>de Moor, I., 2002, 'Potential impacts of alien freshwater crayfish in South Africa', *African Journal of Aquatic Science* 27, 125-139, , https://doi.org/10.2989/16085914.2002.9626584.<br>Nunes, A.L., Zengeya, T.A., Measey, G.J. & Weyl, O.L.F., 2017, 'Freshwater crayfish invasions in South Africa: past, present and potential future', *African Journal of Aquatic Science* 42, 309-323, https://doi.org/10.2989/16085914.2017.1405788. |

| **MAN5 Control options and monitoring approaches available for the *Taxon*** |
|---|
| Response: Yes |
| Rationale: In New Zealand biocides have been used to control and eradicate *Cherax tenuimanus* (Gould 2005), noting that such an approach would need to be carefully regulated given the potential for non-target impacts. |
| References:<br><br>Gould, B., 2005, 'Marron. Interagency collaboration follows surprise catch', *Biosecurity* 60, 10-11. |

| **MAN6 Any other management considerations to highlight?** | |
|---|---|
| Response | No |

#### 5. Calculations

**Likelihood = Fairly probable**

| Parameter | Likelihood | Stages | Final assessment |
|---|---|---|---|
| LIK1 | 1 | P(entry) = 1 | P (invasion) = 0.5 |
| LIK2 | 1 | | |
| LIK3 | 0.5 | P(establishment) = 0.5 | |
| LIK4 | 0.5 | | |
| LIK5 | 0.027 | P (spread) = 1 | |
| LIK6 | 1 | | |

**Consequence = MR**

| Parameter | Mechanism/sector | Response |
|---|---|---|
| CON1a | Competition | DD |
| CON1b | Predation | DD |
| CON1c | Hybridisation | DD |
| CON1d | Disease transmission | DD |
| CON1e | Parasitism | DD |
| CON1f | Poisoning/toxicity | DD |
| CON1g | Bio-fouling or other direct physical disturbance | DD |
| CON1h | Grazing/herbivory/browsing | DD |
| CON1i | Chemical, physical, structural impact | DD |
| CON1k | Indirect impacts through interactions with other species | DD |
| CON1 | Maximum environmental impact | DD |
| CON2a | Safety | DD |
| CON2b | Material and immaterial assets | DD |
| CON2c | Health | DD |
| CON2d | Social, spiritual and cultural relations | DD |
| CON2 | Maximum socio-economic impact | DD |
| CON3 | Environmental impact of closely related taxa | MR |

| | | |
|---|---|---|
| CON4 | Socio-economic impact of closely related taxa | MO |
| CON5 | Potential impact based on traits, experiments, or models | MO |

**Risk score = High**

| | | Consequences | | | | |
|---|---|---|---|---|---|---|
| | | MC | MN | MO | MR | MV |
| Likelihood | Extremely unlikely | low | low | low | medium | medium |
| | Very unlikely | low | low | low | medium | high |
| | Unlikely | low | low | medium | high | high |
| | Fairly probable | medium | medium | high | high | high |
| | Probable | medium | high | high | high | high |

**Ease of management = Medium**

| Parameter | Question | Response |
|---|---|---|
| MAN3a | How accessible are populations? | 0 |
| MAN3b | Is detectability critically time-dependent? | 2 |
| MAN3c | Time to reproduction | 1 |
| MAN3d | Propagule persistence | 0 |
| **MAN3** | **SUM** | 3 |

**Appendix BAC7:** Map of the native range of *Cherax tenuimanus*. **Source**: Allen, M.G., Beatty, S.J., Morgan, D.M. (2012). Stategic action plan for protecting aquatic biodiversity in the Cape to Cape region. Freshwater Fish Group and Fish Health Unit (Murdoch University) report to the Cape to Cape Research Group Department of Evironment and Conservation, Government of Western Australia

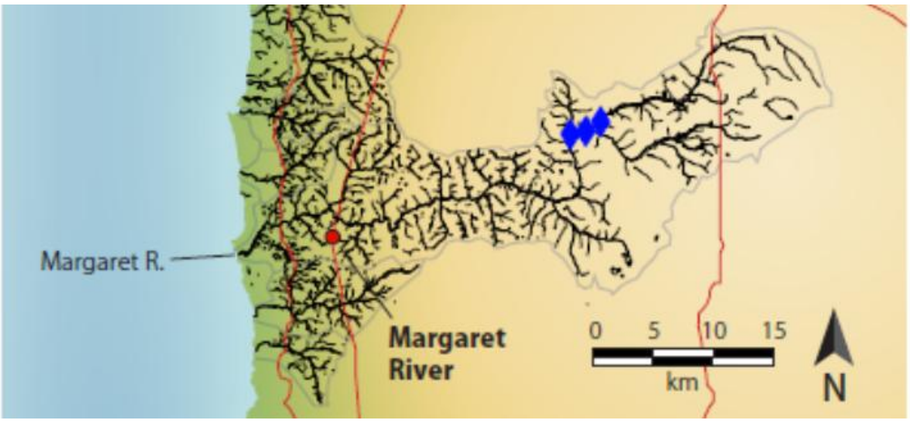

**Appendix BAC8(a)**: Map of the global range of *Cherax tenuimanus.* Source*:* GBIF.org (06 August 2019) GBIF Occurrence Download https://doi.org/10.15468/dl.tvqoxp. The presence of *C. tenuimanus* in South Africa needs to be verified as these records may have been of *C. cainii*.

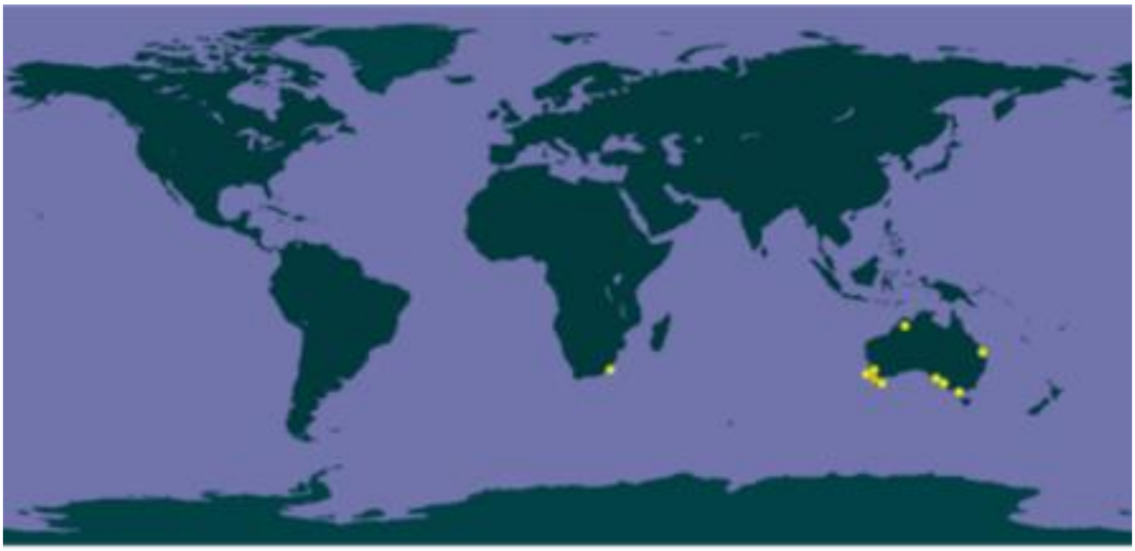

#### 1.3 Risk analysis report for yabby (*Cherax destructor*)

| ***Taxon:*** *Cherax destructor* Clark 1936 | ***Area:*** South Africa |
|---|---|
| **Picture of *Taxon***<br>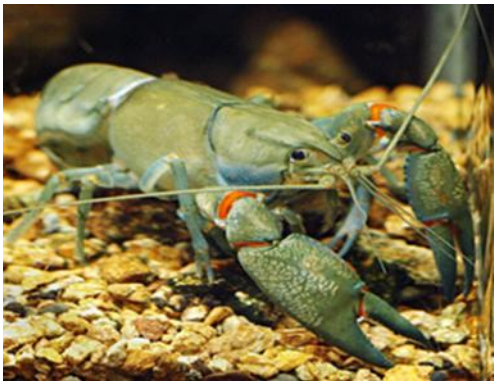<br>Photo: CABI, 2021. *Cherax destructor* [original text by Uma Sabapathy Allen]. In: Invasive Species Compendium. Wallingford, UK: CAB International. www.cabi.org/isc. | **Alien distribution map**<br>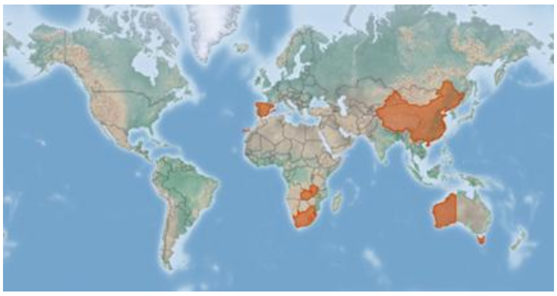<br>Reference: CABI, 2021. *Cherax destructor* [original text by Uma Sabapathy Allen]. In: Invasive Species Compendium. Wallingford, UK: CAB International. www.cabi.org/isc. |
| **Risk Assessment summary:**<br><br>*Cherax destructor* is popular in aquaculture and the pet trade and was introduced to South Africa in 1988 for experimental research to assess its suitability for aquaculture. However, it was found to be unsuitable for aquaculture and although international databases still indicate it is present in South Africa, there are currently no locality records for the country. *Cherax destructor* has a high environmental tolerance, occupying a variety of habitats. Suitable climates are present in a large part of South Africa. *Cherax destructor* is mobile, and dispersal is not limited to interconnected waterways. There are no recorded environmental and socio-economic impacts for *C. destructor*. *Cherax quadricarinatus,* a closely related species, is known to have adverse impacts on invaded ecosystems through competition, predation, and transmission of diseases, and impact local fisheries by replacing high-value native species, damaging nets, and spoiling fish catches. Similar impacts are expected if *C. destructor* were to become invasive. | **Risk score:**<br><br>High |
| **Management summary:**<br><br>*Cherax destructor* is difficult to manage given that it is difficult to access, difficult to detect (being aquatic and primarily nocturnal), and has adaptive life history traits that aid establishment (fast growth rate, high fecundity, and being a multiple spawner). *Cherax destructor* was rejected as a candidate species for aquaculture and there are no records that *C. destructor* has any current benefits. | **Ease of management:**<br><br>Difficult |
| **Recommendations:**<br><br>*Cherax destructor* is currently listed as category 1a. However, there are no records of extant populations in the country. It is recommended that *C. destructor* remain listed as | **Listing under NEM:BA A&IS lists of 2020:** |

| | |
|---|---|
| category 1a until its absence in the country is confirmed at which time it could be moved to the prohibited list. Therefore, this taxon should be reassessed in five years. | 1a[1] |
| | **Recommended listing category:**<br><br>1a |

[1] The presence of *C. destructor* in South Africa needs to be confirmed and the taxon should be reassessed in five years (i.e., by 2029).

#### 1. Background

| **BAC1 Name of assessor(s)** | |
|---|---|
| Name of lead assessor | Lee-Anne Botha |
| Additional assessor (1) | Tsungai Zengeya |
| Additional assessor (2) | Christian Chimimba |
| **BAC2 Contact details of assessor (s)** | |
| Lead assessor | Organisational affiliation: University of Pretoria, Department of Zoology and Entomology/South African National Biodiversity Institute (SANBI) |
| | |
| Additional assessor (1) | Organisational affiliation: University of Pretoria, Department of Zoology and Entomology/South African National Biodiversity Institute (SANBI) |
| | |
| Additional assessor (2) | Organisational affiliation: Department of Zoology and Entomology, University of Pretoria and The South African National Biodiversity Institute (SANBI). |
| | |
| **BAC3 Name(s) and contact details of expert(s) consulted** | |
| Expert (1) | Name: |
| | email: |
| | Phone: |
| Expert (2) | Name: |
| | email: |
| | Phone: |
| Comments: none | |
| **BAC4 Scientific name of *Taxon* under assessment** | |
| *Taxon* name: *Cherax destructor* | Authority: Clarke, 1939 |

Comments: The taxonomic status of *C. destructor* has been subject to several revisions (Austin et al. 2003; Campbell, Geddes & Adams 1994; Munasinghe, Burridge & Austin 2004; Riek 1951, 1956, 1967, 1969; Sokol 1988). The current consensus is that there is one species of *C. destructor*, with two subspecies *C. d. albidus,* and *C. d. destructor* (Austin et al. 2003; Munasinghe et al. 2004), both of which were considered in this risk analysis.

References:

Austin, C.M., Nguyen, T.T.T., Meewan, M.M. & Jerry, D.R., 2003, 'The taxonomy and phylogeny of the '*Cherax destructor*' complex (Decapoda: Parastacidae) examined using mitochondrial 16S sequences', *Australian Journal of Zoology* 51, 99–110, https://doi.org/10.1071/ZO02054.

Campbell, N.J.H., Geddes, M.C. & Adams, M., 1994, 'Genetic variation in yabbies', *Cherax destructor* and *C. albidus* (Crustacea: Decapoda: Parastacidae), indicates the presence of a single, highly sub-structured species', *Australian Journal of Zoology* 42, 1–16, https://doi.org/10.1071/ZO9940745.

Munasinghe, D.H.N., Burridge, C.P. & Austin, C.M., 2004, 'Molecular phylogeny and zoogeography of the freshwater crayfish genus *Cherax* Erichson (Decapoda: Parastacidae) in Australia', *Biological Journal of the Linnean Society* 81, 553–563, https://doi.org/10.1111/j.1095-8312.2003.00299.x.

Riek, E.F., 1951, 'The freshwater crayfish (Family: Parastacidae) of Queensland. With an appendix describing other Australian species', *Records of the Australian Museum* 22,368–388. https://doi.org/10.3853/j.0067-1975.22.1951.615.

Riek, E.F., 1956, 'Addition to the Australian freshwater crayfish', *Records of the Australian Museum* 24, 1–6. https://doi.org/10.3853/j.0067-1975.24.1956.639.

Riek, E.F., 1967, 'The freshwater crayfish of Western Australia (Decapoda: Parastacidae)', *Australian Journal of Zoology* 14, 103–121. https://doi.org/10.1071/ZO9670103.

Riek, E.F., 1969, 'The Australian freshwater crayfish (Crustacea: Decapoda: Parastacidae), with the description of new species', *Australian Journal of Zoology* 17, 855-918. https://doi.org/10.1071/ZO9690855.

| |
|---|
| Sokol, A., 1988, 'Morphological variation in relation to the taxonomy of the destructor group of the genus *Cherax*', *Invertebrate Taxonomy* 2, 55–79, https://doi.org/10.1071/IT9880055. |

| **BAC5 Synonym(s) considered** |
|---|
| Synonyms: *Cherax davis* Clark, 1941 \| *Cherax esculus* Riek, 1956 |
| Comments: The abovementioned synonyms (*Cherax davis* and *Cherax esculus*) were considered in the literature search for this assessment (Crandall & de Grave (2017) |
| References:<br>Crandall, K.A., & De Grave, S., 2017, 'An updated classification of the freshwater crayfishes (Decapoda: Astacidea) of the world, with a complete species list', *The Journal of Crustacean Biology 37*(5), 615-653. https://doi.org/10.1093/jcbiol/rux070. |

| **BAC6 Common name(s) considered** |
|---|
| Common names: Yabby \| common yabby |
| Comments: *Cherax destructor* is commonly known as the yabby or common yabby (Beatty, Morgan & Gill 2005), both these common names were included in the literature search. |
| References:<br>Beatty, S., Morgan, D. & Gill, H., 2005, 'Role of life history strategy in the colonisation of Western Australian aquatic systems by the introduced crayfish *Cherax destructor* Clark, 1936', *Hydrobiologia* 549, 219–237. https://doi.org/10.1007/s10750-005-5443-0. |

| **BAC7 What is the native range of the *Taxon*?** | |
|---|---|
| Response: Australia | Confidence: High |
| Comments: *Cherax destructor* is native to central and eastern Australia (McCormack 2014; Riek 1967; Sokol 1988; Wingfield 2002), see Appendix BAC7. | |
| References:<br>McCormack, R.B., 2014,' New records and review of the translocation of the yabby *Cherax destructor* into eastern drainages of New South Wales, Australia', *Australian Zoologist* 34, 85–94. https://doi.org/10.7882/AZ.2014.006.<br>Riek, E.F., 1967, 'The freshwater crayfish of Western Australia (Decapoda: Parastacidae)', *Australian Journal of Zoology* 14, 103–121. https://doi.org/10.1071/ZO9670103.<br>Sokol, A., 1988, 'Morphological variation in relation to the taxonomy of the destructor group of the genus *Cherax'*, *Invertebrate Taxonomy* 2, 55–79. https://doi.org/10.1071/IT9880055.<br>Wingfield, M., 2002, 'An overview of the Australian freshwater crayfish farming industry', *Freshwater Crayfish* 13(1), 177-184. | |

| **BAC8 What is the global alien range of the *Taxon*?** | |
|---|---|
| Response: Italy \| Spain \| Switzerland \| Ireland \| China \| Western Australia \| South Africa \| Zambia | Confidence: Medium |
| Comments: *Cherax destructor* is reported as introduced to China, western Australian, Switzerland, Ireland, South Africa, and Zambia (CABI 2021), with naturalised populations in Spain and Italy (Scalici et al. 2009). The population in Italy might have been eradicated by the crayfish plague (Mazza et al. 2018). Although *C. destructor* was introduced to South Africa and Zambia, there are no recent records of presence (Nunes et al. 2017). | |
| References:<br>CABI, 2021. *Cherax destructor* [original text by Uma Sabapathy Allen]. In: Invasive Species Compendium. Wallingford, UK: CAB International. www.cabi.org/isc.<br>Scalici, M., Chiesa, S., Gherardi, F., Ruffini, M., Gibertini, G. & Marzano, F.N., 2009, 'The new threat to Italian inland waters from the alien crayfish "gang": the Australian *Cherax destructor* Clark, 1936', *Hydrobiologia* 632, 341–345. https://doi.org/10.1007/s10750-009-9839-0.<br>Mazza, G., Scalici, M., Inghilesi, A.F., Aquiloni, L., Pretto, T., Monaco, A., et al., 2018, 'The red alien vs. the blue destructor: The eradication of *Cherax destructor* by *Procambarus clarkii* in Latium (Central Italy)', *Diversity* 10(4), 126. https://doi.org/10.3390/d10040126 | |

Nunes, A.L., Zengeya, T.A., Measey, G.J. & Weyl, O.L.F., 2017, 'Freshwater crayfish invasions in South Africa: Past, present and potential future', *African Journal of Aquatic Science* 42, 309–323. https://doi.org/10.2989/16085914.2017.1405788

**BAC9 Geographic scope = the *Area* under consideration**

*Area* of assessment: South Africa

Comments: The geographic scope of this assessment is limited to South Africa.

**BAC10 Is the *Taxon* present in the *Area*?**

| | |
|---|---|
| Response: Yes | Confidence: Low |

Comments: *Cherax destructor* was imported to South Africa in 1988 for experimental research to assess its suitability for aquaculture and there is a possibility that escapees and illegal introductions might have established populations outside of captivity (de Moor 2002). However, there is no recent evidence that *C. destructor* is still present and so the confidence is scored as low (Nunes et al. 2017).

References:

de Moor, I., 2002, 'Potential impacts of alien freshwater crayfish in South Africa', *African Journal of Aquatic Science* 27, 125–139. https://doi.org/10.2989/16085914.2002.9626584.

Nunes, A.L., Zengeya, T.A., Measey, G.J. & Weyl, O.L.F., 2017, 'Freshwater crayfish invasions in South Africa: Past, present and potential future', *African Journal of Aquatic Science* 42, 309–323. https://doi.org/10.2989/16085914.2017.1405788.

**BAC11 Availability of physical specimen**

| | |
|---|---|
| Response: No physical specimen is available | Confidence in ID: |

Herbarium or museum accession number:

References:

**BAC12 Is the *Taxon* native to the *Area* or part of the *Area*?**

| | | |
|---|---|---|
| The *Taxon* is native to (part of) the *Area*. | No | Confidence: High |
| The *Taxon* is alien in (part of) the *Area*. | Yes | Confidence: High |

Comments: *Cherax destructor* is native to central and eastern parts of Australia (McCormack 2014; Riek 1967; Sokol 1988; Wingfield 2002).

References:

McCormack, R.B., 2014,' New records and review of the translocation of the yabby *Cherax destructor* into eastern drainages of New South Wales, Australia', *Australian Zoologist* 34, 85–94. http://dx.doi.org/10.7882/AZ.2014.006.

Riek, E.F., 1967, 'The freshwater crayfish of Western Australia (Decapoda: Parastacidae)', *Australian Journal of Zoology* 14, 103–121. https://doi.org/10.1071/ZO9670103.

Sokol, A., 1988, 'Morphological variation in relation to the taxonomy of the destructor group of the genus *Cherax'*, *Invertebrate Taxonomy* 2, 55–79. https://doi.org/10.1071/IT9880055.

Wingfield, M., 2002, 'An overview of the Australian freshwater crayfish farming industry', *Freshwater Crayfish* 13(1), 177-184.

**BAC13 What is the *Taxon*'s introduction status in the *Area*?**

| | | |
|---|---|---|
| The *Taxon* is in cultivation/containment. | No | Confidence: Low |
| The *Taxon* is present outside of cultivation/containment. | No | Confidence: Low |
| The *Taxon* has established/naturalised. | No | Confidence: High |
| The *Taxon* is invasive. | No | Confidence: High |

Comments: *Cherax destructor* was introduced in 1988 for research to see if was suitable for aquaculture, but there is no evidence that indicates that the species succeeded in establishing populations. There are anecdotal records of *C. destructor* introductions by fishers into several South African dams. These records, gathered by fishing and aquarium enthusiasts, need to be verified (Nunes et al. 2017).

References:

Nunes, A.L., Zengeya, T.A., Measey, G.J. & Weyl, O.L.F., 2017, 'Freshwater crayfish invasions in South Africa: Past, present and potential future', *African Journal of Aquatic Science* 42, 309–323. https://doi.org/10.2989/16085914.2017.1405788.

| **BAC14 Primary (introduction) pathways** | | |
|---|---|---|
| Release | none | Confidence: Medium |
| Escape | aquaculture/mariculture | Confidence: High |
| | live food and live bait | Confidence: High |
| Contaminant | pet/aquarium/terrarium species (including live food for such species) | Confidence: High |
| Stowaway | none | Confidence: Medium |
| Corridor | none | Confidence: Medium |
| Unaided | none | Confidence: Medium |
| Comments: *Cherax destructor* is common in the global pet trade (Chucholl 2013; Faulkes 2015) and sold as live bait to fishers (McCormack 2014). Furthermore, *C. destructor* has also been introduced for aquaculture purposes (Nunes et al. 2017). | | |
| References: Chucholl, C., 2013, 'Invaders for sale: trade and determinants of introduction of ornamental freshwater crayfish', *Biological Invasions* 15, 125–141. https://doi.org/10.1007/s10530-012-0273-2 Faulkes, Z., 2015, 'The global trade in crayfish as pets', *Crustacean Research* 44, 75–92. https://doi.org/10.18353/crustacea.44.0_75. McCormack, R.B., 2014,' New records and review of the translocation of the yabby *Cherax destructor* into eastern drainages of New South Wales, Australia', *Australian Zoologist* 34, 85–94. https://doi.org/10.7882/AZ.2014.006. Nunes, A.L., Zengeya, T.A., Measey, G.J. & Weyl, O.L.F., 2017, 'Freshwater crayfish invasions in South Africa: Past, present and potential future', *African Journal of Aquatic Science* 42, 309–323. https://doi.org/10.2989/16085914.2017.1405788. | | |

#### 2. Likelihood

| **LIK1 Likelihood of entry via unaided primary pathways** | |
|---|---|
| Response: Very unlikely (p = 0.0027) | Confidence: Low |
| Rationale: There are no known populations in South Africa or neighbouring countries that could act as a source of introduction (Madzivanzira et al. 2020). Although *C. destructor* was introduced in Zambia there are no current records of them in captivity or in natural environments (Madzivanzira et al. 2020). Populations outside of captivity are only known from Spain (Scalici et al 2009). | |
| References: Madzivanzira, T.C., South, J., Wood, L.E., Nunes, A.L. & Weyl, O.L.F., 2020, 'A review of freshwater crayfish introductions in Africa', *Reviews in Fisheries Science & Aquaculture* 1-21. https://doi.org/10.1080/23308249.2020.1802405 Scalici, M., Chiesa, S., Gherardi, F., Ruffini, M., Gibertini, G. & Marzano, F.N., 2009, 'The new threat to Italian inland waters from the alien crayfish "gang": the Australian *Cherax destructor* Clark, 1936', *Hydrobiologia* 632, 341–345. https://doi.org/10.1007/s10750-009-9839-0. | |

| **LIK2 Likelihood of entry via human aided primary pathways** | |
|---|---|
| Response: Fairly probable (p = 0.5) | Confidence: Medium |
| Rationale: *Cherax destructor* was introduced into South Africa for experimental research in 1988 but there are no confirmed populations in captivity or in natural environments (Nunes et al. 2017). It could potentially be introduced in the future for aquaculture, through the pet trade or as bait for fishing (Chucholl 2013; Faulkes 2015; Nguyen 2005). | |
| References: | |

Chucholl, C., 2013, 'Invaders for sale: trade and determinants of introduction of ornamental freshwater crayfish', *Biological Invasions* 15, 125–141. https://doi.org/10.1007/s10530-012-0273-2
Faulkes, Z., 2015, 'The global trade in crayfish as pets', *Crustacean Research* 44, 75–92. https://doi.org/10.18353/crustacea.44.0_75.
Nguyen, T.T.T., 2005, 'A genetic investigation on translocation of Australian commercial freshwater crayfish, *Cherax destructor*', *Aquatic Living Resources* 18, 319–323. https://doi.org/10.1051/alr:2005035.
Nunes, A.L., Zengeya, T.A., Measey, G.J. & Weyl, O.L.F., 2017, 'Freshwater crayfish invasions in South Africa: Past, present and potential future', *African Journal of Aquatic Science* 42, 309–323. https://doi.org/10.2989/16085914.2017.1405788.

**LIK3 Habitat suitability**

| Response: Probable (p = 1) | Confidence: High |
|---|---|

Rationale: *Cherax destructor* occurs in a wide range of habitats and is known to have a high tolerance to salinity (15000ppm), low oxygen conditions (1ppm) and poor water quality (Beatty 2005). *Cherax destructor* occurs in desert mound springs, alpine streams, subtropical creeks, rivers, ephemeral lakes, swamps, farms dams, and irrigation canals (de Moor 2002). *Cherax destructor* is a burrowing species that can excavate shafts that are 0.2–5 m deep, which are often used as 'refugia' from desiccation in the dry season (Beatty et al. 2005; Kouba et al. 2016; Withnall 2000). Suitable habitat is therefore widely available in South Africa (Nunes et al. 2017).

References:

Beatty, S., 2005, 'Translocations of freshwater crayfish: contributions from life histories, trophic relations and diseases of three species in Western Australia', Doctoral dissertation, Murdoch University, Australia.
de Moor, I., 2002, 'Potential impacts of alien freshwater crayfish in South Africa', *African Journal of Aquatic Science* 27, 125–139. https://doi.org/10.2989/16085914.2002.9626584.
Kouba, A., Tíkal, J., Císař, P., Veselý, L., Fořt, M., Příborský, J., et al., 2016, 'The significance of droughts for hyporheic dwellers: evidence from freshwater crayfish', *Scientific reports* 6(1), 26569. https://doi.org/10.1038/srep26569.
Nunes, A.L., Zengeya, T.A., Measey, G.J. & Weyl, O.L.F., 2017, 'Freshwater crayfish invasions in South Africa: Past, present and potential future', *African Journal of Aquatic Science* 42, 309–323. https://doi.org/10.2989/16085914.2017.1405788.
Withnall, F., 2000, 'Biology of yabbies (*Cherax destructor*)', *Aquaculture notes of the department of Natural Resources and Environment*, Victoria, Australia.

**LIK4 Climate suitability**

| Response: Probable (p = 1) | Confidence: High |
|---|---|

Rationale: *Cherax destructor* has a wide thermal tolerance range (1-35°C), with an optimal range of 22-28°C with growth ceasing at <15 °C and >34°C (McCormack 2014). A substantial part of South Africa was projected to be climatically suitable for *C. destructor* (>20% of the country) including ecoregions in the Soutpansberg, North Eastern Highlands, Eastern Bankenveld, Northern Escarpment Mountains, the northern half of the North Eastern Uplands, South Eastern Uplands and sections of the Southern Fold Mountains and Southern Coastal Belt (de Moor 2002; Nunes et al. 2017).

| |
|---|
| References:<br><br>de Moor, I., 2002, 'Potential impacts of alien freshwater crayfish in South Africa', *African Journal of Aquatic Science* 27, 125–139, https://doi.org/10.2989/16085914.2002.9626584.<br>McCormack, R.B., 2014,' New records and review of the translocation of the yabby *Cherax destructor* into eastern drainages of New South Wales, Australia', *Australian Zoologist* 34, 85–94. https://doi.org/10.7882/AZ.2014.006.<br>Nunes, A.L., Zengeya, T.A., Measey, G.J. & Weyl, O.L.F., 2017, 'Freshwater crayfish invasions in South Africa: Past, present and potential future', *African Journal of Aquatic Science* 42, 309–323. https://doi.org/10.2989/16085914.2017.1405788. |

| **LIK5 Unaided secondary (dispersal) pathways** | |
|---|---|
| Response: Fairly probable (p = 0.5) | Confidence: High |
| Rationale: *Cherax destructor*, like other crayfish species, is highly mobile and can migrate overland (Beatty 2005; Withnall 2000), and is thus capable of unaided secondary dispersal. | |
| References:<br><br>Beatty, S., 2005, 'Translocations of freshwater crayfish: contributions from life histories, trophic relations and diseases of three species in Western Australia', Doctoral dissertation, Murdoch University, Australia.<br>Withnall, F., 2000, 'Biology of yabbies (*Cherax destructor*)', *Aquaculture notes of the department of Natural Resources and Environment*, State of Victoria, Australia. | |

| **LIK6 Human aided secondary (dispersal) pathways** | |
|---|---|
| Response: Fairly probable (p = 0.5) | Confidence: Low |
| Rationale: *Cherax destructor* is a popular species in the global pet trade and could be intentionally introduced into the wild in new areas as unwanted pets (Chucholl 2013; Faulkes 2015; Nunes et al 2017). | |
| References:<br><br>Chucholl, C., 2013, 'Invaders for sale: trade and determinants of introduction of ornamental freshwater crayfish', *Biological Invasions* 15, 125–141. https://doi.org/10.1007/s10530-012-0273-2<br>Faulkes, Z., 2015, 'The global trade in crayfish as pets', *Crustacean Research* 44, 75–92. https://doi.org/10.18353/crustacea.44.0_75<br>Nunes, A.L., Zengeya, T.A., Measey, G.J. & Weyl, O.L.F., 2017, 'Freshwater crayfish invasions in South Africa: Past, present and potential future', *African Journal of Aquatic Science* 42, 309–323. https://doi.org/10.2989/16085914.2017.1405788. | |

**3. Consequences**

| **CON1 Environmental impact** | |
|---|---|
| **CON1a: Competition** | |
| Response: Data Deficient (DD) | Confidence: |
| Rationale: | |
| References: | |
| **CON1b: Predation** | |
| Response: Data Deficient (DD) | Confidence: |
| Rationale: | |
| References: | |
| **CON1c: Hybridisation** | |
| Response: Data Deficient (DD) | Confidence: |
| Rationale: | |
| References: | |
| **CON1d: Transmission of disease** | |
| Response: Data Deficient (DD) | Confidence: |
| Rationale: | |
| References: | |
| **CON1e: Parasitism** | |
| Response: Data Deficient (DD) | Confidence: |
| Rationale: | |
| References: | |
| **CON1f: Poisoning/toxicity** | |
| Response: Data Deficient (DD) | Confidence: |
| Rationale: | |
| References: | |
| **CON1g: Bio-fouling or other direct physical disturbance** | |
| Response: Data Deficient (DD) | Confidence: |
| Rationale: | |

| | |
|---|---|
| References: | |
| **CON1h: Grazing/herbivory/browsing** | |
| Response: Data Deficient (DD) | Confidence: |
| Rationale: | |
| References: | |
| **CON1i: Chemical, physical or structural impact on ecosystem** | |
| Response: Data Deficient (DD) | Confidence: |
| Rationale: | |
| References: | |
| **CON1k: Indirect impacts through interactions with other species** | |
| Response: Data Deficient (DD) | Confidence: |
| Rationale: | |
| References: | |
| **CON1 Maximum environmental impact** | |
| Response: Data Deficient (DD) | Confidence: |
| Rationale: | |
| References: | |

| | |
|---|---|
| **CON2 Socio-economic impact** | |
| **CON2a: Safety** | |
| Response: Data Deficient (DD) | Confidence: |
| Rationale: | |
| References: | |
| **CON2b: Material and immaterial assets** | |
| Response: Data Deficient (DD) | Confidence: |
| Rationale: | |
| References: | |
| **CON2c: Health** | |
| Response: Data Deficient (DD) | Confidence: |

| |
|---|
| Rationale: |
| References: |

| | |
|---|---|
| **CON2d: Social, spiritual and cultural relations** | |
| Response: Data Deficient (DD) | Confidence: |
| Rationale: | |
| References: | |
| **CON2 Maximum socio-economic impact** | |
| Response: Data Deficient (DD) | Confidence: |
| Rationale: | |
| References: | |

| | |
|---|---|
| **CON3 Closely related species' environmental impact** | |
| Response: Major (MR) | Confidence: Medium |
| Rationale: There is limited information on the impacts of *C. destructor* (e.g., Coghran & Daly 2012). Therefore, the closely related *Cherax quadricarinatus* has been used to estimate the impacts that may arise from an introduction of *C. destructor*. *Cherax quadricarinatus* invasions have been implicated in the decline of native species of macrophytes, macroinvertebrates, and fish (Douthwaite et al. 2018; Marufu et al. 2018; Pinder et al. 2019). In South Africa, there have been laboratory studies to quantify the impacts of *C. quadricarinatus* on various aquatic and terrestrial resources ubiquitous to South African ecosystems. *Cherax quadricarinatus* exhibited deleterious impacts on various ecosystem resources that include fish, macroinvertebrates, and aquatic plants which are all important for ecosystem structure and functioning (Madzivanzira et al. 2021, 2022). | |
| References: Belle, C.C., Wong, J.Q.H., Yeo, D.C.J., Tan, S.H., Tan, H.H., Clews, E., et al., 2011, 'Ornamental trade as a pathway for Australian redclaw crayfish introduction and establishment', *Aquatic Biology* 12, 69-79. https://doi.org/10.3354/ab00324.<br>Coghran, J., & Daly, G., 2012, 'Potential threats posed by a translocated crayfish: the case of *Cherax destructor* in coastal drainages of New South Wales, Australia', *Crustacean Research* (7), 5-13. https://doi.org/10.18353/crustacea.Special2012.7_5.<br>Douthwaite, R., Jones, E., Tyser, A. & Vrdoljak, S., 2018, 'The introduction, spread and ecology of redclaw crayfish *Cherax quadricarinatus* in the Zambezi catchment', *African Journal of Aquatic Science* 43, 353–366. https://doi.org/10.2989/16085914.2018.1517080.<br>Madzivanzira, T.C., South, J. & Weyl, O.L.F., 2021,' Invasive crayfish outperform Potamonautid crabs at higher temperatures', *Freshwater Biology* 66(5), 978–991. https://doi.org/10.1111/fwb.13691.<br>Madzivanzira, T.C., Weyl, O.L.F. & South, J., 2022, 'Ecological and potential socioeconomic impacts of two globally invasive crayfish', *NeoBiota* 72, 25–43. https://doi.org/10.3897/neobiota.72.71868.<br>Marufu, L.T., Dalu, T., Phiri, C., Barson, M., Simango, R., Utete, B. et al., 2018, 'The diet of an invasive crayfish, *Cherax quadricarinatus* (Von Martens, 1868), in Lake Kariba, inferred using stomach content and stable isotope analyses', *BioInvasions Records* 7, 121–132. https://doi.org/10.3391/bir.2018.7.2.03.<br>Pinder, A., Harman, A., Bird, C., Quinlan, K., Angel, F., Cowan, M. et al., 2019,' Spread of the non-native redclaw crayfish *Cherax quadricarinatus* (von Martens, 1868) into natural waters of the Pilbara region | |

of Western Australia, with observations on potential adverse ecological effects', *BioInvasions Records* 8, 882–897. https://doi.org/10.3391/bir.2019.8.4.17.

Yeo, D.C.J., 2010, 'Introduced decapod crustaceans in Singapore's reservoirs', *Cosmos* 6, 83-88.

| **CON4 Closely related species' socio-economic impact** | |
|---|---|
| Response: Moderate (MO) | Confidence: Low |

Rationale:

There are no recorded socio-economic impacts of *C. destructor*. Therefore, the closely related *Cherax quadricarinatus* has been used to estimate the impacts that may arise from the introduction of *C. destructor*. The presence of *C. quadricarinatus* in local fisheries can threaten the livelihoods of fishers by replacing native species that are of higher economic value, damaging fishing gear and spoiling fish catches (Chakandinakira et al. 2023; Douthwaite et al. 2018; Haubrock et al. 2023; Madzivanzira et al. 2022; Vega-Villasante et al. 2015).

References:

Chakandinakira, A.T., Madzivanzira, T.C., Mashonga, S., Muzvondiwa, J.V., Ndlovu, N. & South, J. 2023, 'Socioeconomic impacts of Australian redclaw crayfish *Cherax quadricarinatus* in Lake Kariba', *Biological Invasions* 25, 2801-2812. https://doi.org/10.1007/s10530-023-03074-8

Douthwaite, R., Jones, E., Tyser, A. & Vrdoljak, S., 2018, 'The introduction, spread and ecology of redclaw crayfish *Cherax quadricarinatus* in the Zambezi catchment', *African Journal of Aquatic Science* 43, 353–366. https://doi.org/10.2989/16085914.2018.1517080.

Haubrock, P.J., Oficialdegui, F.J., Zeng, Y., Patoka, J., Yeo, D.C., & Kouba, A., 2021, 'The redclaw crayfish: A prominent aquaculture species with invasive potential in tropical and subtropical biodiversity hotspots', *Reviews in Aquaculture* 13(3), 1488-1530, https://doi.org/10.1111/raq.12531.

Madzivanzira, T.C., Weyl, O.L.F. & South, J., 2022, 'Ecological and potential socioeconomic impacts of two globally invasive crayfish', *NeoBiota* 72, 25–43. https://doi.org/10.3897/neobiota.72.71868.

Vega-Villasante, F., Avalos-Aguilar, J.J., Nolasco-Soria, H., Vargas-Ceballos, M.A., Bortolini-Rosales, J.L., Chong-Carrillo O. et al., 2015, 'Wild populations of the invasive Australian red claw crayfish *Cherax quadricarinatus* (Crustacea, Decapoda) near the northern coast of Jalisco, Mexico: a new fishing and profitable resource', *Latin American Journal of Aquatic Research* 43, 781–785. https://doi.org/10.3856/vol43-issue4-fulltext-17.

| **CON5 Potential impact** | |
|---|---|
| Response: Major (MR) | Confidence: Medium |

Rationale: *Cherax destructor* can withstand variety of environmental conditions that help to facilitate its establishment. *Cherax destructor* can also switch from a diet of fish in summer to a predominantly herbaceous/detrital diet in winter (Beaty 2005). Therefore, it may compete for food resources with closely related native decapods (de Moor 2002; Nunes et al. 2017). *Cherax destructor* is a possible vector of parasites and diseases as it is known to harbour recognised pathogens such protozoans, bacteria, and viruses that can be transmitted to native species in invaded ecosystems (Edgerton et al. 2002; Longshaw 2011).

Furthermore, the burrowing behaviour of *C. destructor* is a cause for concern (Withnall 2000), with very deep burrows of 50 cm to 2 m (Withnall 2005). Burrows are connected by access shafts to the water. In the event of the water drying up, they can survive over summer in the burrows. This behaviour can destroy the integrity of dam walls by increasing soil erosion (de Moor 2002).

Particular life history traits (high fecundity, fast growth, large size, omnivorous diet, and wide environmental tolerances) have been inferred to predispose the closely related *Cherax quadricarinatus* to have major impacts on ecosystems through predation, competition, and transfer of diseases (e.g., Ahyong & Yeo 2007; Bortolini et al. 2007; Lynas et al. 2007; Madzivanzira et al. 2021, 2022; Marufu et al. 2018; Yeo 2010; Zeng et al. 2019; Zengeya et al. 2022).

References:

Ahyong, S.T. & Yeo, D.C.J., 2007, 'Feral populations of the Australian redclaw crayfish (*Cherax quadricarinatus* von Martens) in water supply catchments of Singapore', *Biological Invasions* 9, 943-946, https://doi.org/10.1007/s10530-007-9094-0.

Beatty, S., 2005, 'Translocations of freshwater crayfish: contributions from life histories, trophic relations and diseases of three species in Western Australia', Doctoral dissertation, Murdoch University, Australia. https://doi.org/10.1007/s10750-005-5443-0

Bortolini, J.L., Alvarez, F. & Rodríguez-Almaraz, G., 2007, 'On the presence of the Australian redclaw crayfish *Cherax quadricarinatus* in Mexico', *Biological Invasions* 9, 615-620. https://doi.org/10.1007/s10530-006-9054-0.

de Moor, I., 2002, 'Potential impacts of alien freshwater crayfish in South Africa', *African Journal of Aquatic Science* 27, 125–139. https://doi.org/10.2989/16085914.2002.9626584.

Du Preez, L. & Smit, N., 2013, 'Double blow: Alien crayfish infected with invasive temnocephalan in South African waters', *South African Journal of Science* 109**,** 01–04. https://hdl.handle.net/10520/EJC142057

Edgerton, B.F., Evans L.H., Stephens, F.J. & Overstreet, R.M., 2002, 'Synopsis of freshwater crayfish diseases and commensal organisms', *Aquaculture* 206, 57–135. https://doi.org/10.1016/S0044-8486(01)00865-1

Longshaw, M., 2011, 'Diseases of crayfish: a review', *Journal of Invertebrate Pathology* 106, 54–70. https://doi.org/10.1016/j.jip.2010.09.013.

Lynas, J., Storey, A. & Knott, B., 2007, 'Introduction and spread of crayfish (Parastacidae) in Western Australia and their potential to displace indigenous species', In: Gherardi F (ed.) *Biological invaders in inland waters: profiles, distribution and threats*, pp. 577-596, Springer, Dordrecht, Netherlands.

Madzivanzira, T.C., South, J. & Weyl, O.L.F., 2021,' Invasive crayfish outperform Potamonautid crabs at higher temperatures', *Freshwater Biology* 66(5), 978–991. https://doi.org/10.1111/fwb.13691.

Madzivanzira, T.C., Weyl, O.L.F. & South, J., 2022, 'Ecological and potential socioeconomic impacts of two globally invasive crayfish', *NeoBiota* 72, 25–43. https://doi.org/10.3897/neobiota.72.71868.

Marufu, L.T., Dalu, T., Phiri, C., Barson, M., Simango, R., Utete, B. et al., 2018, 'The diet of an invasive crayfish, *Cherax quadricarinatus* (Von Martens, 1868), in Lake Kariba, inferred using stomach content and stable isotope analyses', *BioInvasions Records* 7, 121–132. https://doi.org/10.3391/bir.2018.7.2.03.

Nunes, A.L., Zengeya, T.A., Measey, G.J. & Weyl, O.L.F., 2017, 'Freshwater crayfish invasions in South Africa: Past, present and potential future', *African Journal of Aquatic Science* 42, 309–323. https://doi.org/10.2989/16085914.2017.1405788.

Withnall, F., 2000, 'Biology of yabbies (*Cherax destructor*)', *Aquaculture notes of the department of Natural Resources and Environment*, Victoria, Australia.

Zeng, Y., Shakir, K.K. & Yeo, D.C.J. 2019, 'Competition between a native freshwater crab and an invasive crayfish in tropical Southeast Asia', *Biological Invasions* 21, 2653–2663. https://doi.org/10.1007/s10530-019-02009-6.

Zengeya, T.A, Lombard, R.J.H., Nelwamondo, V.E., Nunes, A.L., Measey, J.G. & Weyl, O.L.F., 2022, 'Trophic niche of an invasive generalist consumer: Australian redclaw crayfish, *Cherax quadricarinatus*, in the Inkomati River Basin, South Africa', *Austral Ecology* 47, 1480-1494. https://doi.org/10.1111/aec.13230.

#### 4. Management

| MAN1 What is the feasibility to stop future immigration? | |
|---|---|
| Response: High | Confidence: Low |
| Rationale: There are no known populations in neighbouring countries, thus the probability of species entering via unaided primary pathways is very low (de Moor 2002; Nunes et al. 2017). Intentional introductions can be prevented through the regulation of imports (Zengeya & Wilson 2020). However, illegal imports can still be problematic (Faulkes 2015; Nunes et al. 2017). *Cherax destructor* is present in the global pet trade industry (Faulkes 2015) but its potential presence in the pet trade in South Africa is still to be assessed (Nunes et al. 2017). | |
| References:<br>de Moor, I., 2002, 'Potential impacts of alien freshwater crayfish in South Africa', *African Journal of Aquatic Science* 27, 125–139, https://doi.org/10.2989/16085914.2002.9626584.<br>Faulkes, Z., 2015, 'The global trade in crayfish as pets', *Crustacean Research* 44, 75–92. https://doi.org/10.18353/crustacea.44.0_75.<br>Nunes, A.L., Zengeya, T.A., Measey, G.J. & Weyl, O.L.F., 2017, 'Freshwater crayfish invasions in South Africa: Past, present and potential future', *African Journal of Aquatic Science* 42, 309–323. https://doi.org/10.2989/16085914.2017.1405788.<br>Zengeya, T.A. & Wilson, J.R., 2020, *The status of biological invasions and their management in South Africa in 2019*, South African National Biodiversity Institute, Kirstenbosch and DSI-NRF Centre of Excellence for Invasion Biology, Stellenbosch. http://dx.doi.org/10.5281/zenodo.3947613. | |

| MAN2 Benefits of the Taxon | |
|---|---|
| **MAN2a Socio-economic benefits of the *Taxon*** | |
| Response: Low | Confidence: Low |
| Rationale: Production of crayfish in general in South Africa has had mixed success. Experimental research for the use of *C. destructor* in aquaculture was unsuccessful (Nunes et al. 2017). As a result, aquaculture of crayfish has been restricted to *Cherax cainii* in a few small scale aquaculture farms in Eastern Cape and Western Cape (Madzivanzira et al. 2020). | |
| References:<br>Madzivanzira, T.C., South, J., Wood, L.E., Nunes, A.L. & Weyl, O.L.F., 2020, 'A review of freshwater crayfish introductions in Africa', *Reviews in Fisheries Science & Aquaculture* 29(2), 1-21. https://doi.org/10.1080/23308249.2020.1802405.<br>Nunes, A.L., Zengeya, T.A., Measey, G.J. & Weyl, O.L.F., 2017, 'Freshwater crayfish invasions in South Africa: Past, present and potential future', *African Journal of Aquatic Science* 42, 309–323. https://doi.org/10.2989/16085914.2017.1405788. | |
| **MAN2b Environmental benefits of the *Taxon*** | |
| Response: Low | Confidence: Low |
| Rationale: There are no recorded environmental benefits associated with *C. destructor*. | |
| References: | |

| MAN3 Ease of management |
|---|

| **MAN3a How accessible are populations?** | |
|---|---|
| Response: 2 (difficult to access) | Confidence: Low |
| Rationale: *Cherax destructor* is a small nocturnal aquatic species that is difficult to access (Larson et al. 2016). | |
| References:<br><br>Larson, E.R. & Olden, J.D., 2016, 'Field sampling techniques for crayfish', In: Longshaw, M. & Stebbing, P. (eds), *Biology and Ecology of Crayfish 1st edn*., pp. 287- 323, CRC Press, Boca Raton. | |
| **MAN3b Is detectability critically time-dependent?** | |
| Response: 2 (yes) | Confidence: High |
| Rationale: *Cherax destructor* is a small nocturnal aquatic species that is difficult to detect (Larson et al. 2016). Moreover, *C. destructor* can burrow which further impedes detection (McCormack 2014). Detectability is not particularly seasonal however (de Moor 2002; Nunes et al. 2017). | |
| References:<br><br>de Moor, I., 2002, 'Potential impacts of alien freshwater crayfish in South Africa', *African Journal of Aquatic Science* 27, 125–139. https://doi.org/10.2989/16085914.2002.9626584.<br>Nunes, A.L., Zengeya, T.A., Measey, G.J. & Weyl, O.L.F., 2017, 'Freshwater crayfish invasions in South Africa: Past, present and potential future', *African Journal of Aquatic Science* 42, 309–323. https://doi.org/10.2989/16085914.2017.1405788.<br>Larson, E.R. & Olden, J.D., 2016, 'Field sampling techniques for crayfish', In: Longshaw, M. & Stebbing, P. (eds), *Biology and Ecology of Crayfish 1st edn*., pp. 287- 323, CRC Press, Boca Raton.<br>McCormack, R.B., 2014, 'New records and review of the translocation of the yabby *Cherax destructor* into eastern drainages of New South Wales, Australia', *Australian Zoologist* 34, 85–94. https://doi.org/10.7882/AZ.2014.006. | |
| **MAN3c Time to reproduction** | |
| Response: 2 (<1 year) | Confidence: High |
| Rationale: *Cherax destructor* can reach sexual maturity within six months and has a short gestation period (six to eight weeks) where the female protects eggs until they hatched, and juveniles are ready to leave the female (Beatty et al. 2005; McCormack 2014; Mills 1983; Withnall 2000). | |
| References:<br>Beatty, S.J., Morgan, D.L. & Gill, H.S., 2005, 'Role of life history strategy in the colonisation of Western Australian aquatic systems by the introduced crayfish *Cherax destructor* Clark, 1936', *Hydrobiologia* 549, 219-237. https://doi.org/10.1007/s10750-005-5443-0.<br>McCormack, R.B., 2014, 'New records and review of the translocation of the yabby *Cherax destructor* into eastern drainages of New South Wales, Australia', *Australian Zoologist* 34, 85–94. https://doi.org/10.7882/AZ.2014.006.<br>Mills, B.J., 1983, 'Aquaculture of yabbies', In: *Proceedings of the First Freshwater Aquaculture Workshop*, Department of Agriculture, New South Wales. Australia.<br>Withnall, F., 2000, 'Biology of yabbies (*Cherax destructor*)', *Aquaculture notes of the department of Natural Resources and Environment*. State of Victoria, Australia. | |
| **MAN3d Propagule persistence** | |
| Response: 0 (<1 year) | Confidence: High |

| |
|---|
| Rationale: *Cherax destructor* can have multiple spawning events within one year where one brood can produce 300 eggs on average (Beatty et al. 2005; Mills 1983; McCormack 2014; Withnall 2000) |
| References:<br><br>Beatty, S.J., Morgan, D.L. & Gill, H.S., 2005, 'Role of life history strategy in the colonisation of Western Australian aquatic systems by the introduced crayfish *Cherax destructor* Clark, 1936', *Hydrobiologia* 549, 219-237https://doi.org/10.1007/s10750-005-5443-0.<br>McCormack, R.B., 2014, 'New records and review of the translocation of the yabby *Cherax destructor* into eastern drainages of New South Wales, Australia', *Australian Zoologist* 34, 85–94. https://doi.org/10.7882/AZ.2014.006.<br>Mills, B.J., 1983, 'Aquaculture of yabbies', In: *Proceedings of the First Freshwater Aquaculture Workshop*, Department of Agriculture, New South Wales. Australia.<br>Withnall, F., 2000, 'Biology of yabbies (*Cherax destructor*)', *Aquaculture notes of the department of Natural Resources and Environment*. State of Victoria, Australia. |

| **MAN3 Ease of management** | |
|---|---|
| Response: Difficult | Confidence: High |
| Rationale: *Cherax destructor* is difficult to manage given that it is difficult to access, difficult to detect (being aquatic and primarily nocturnal) and has adaptive life history traits that aid establishment (fast growth rate, high fecundity, and a multiple spawner) (Beatty et al. 2005; Larson & Olden 2016; McCormack 2014; Mills 1983; Nunes et al. 2017; Withnall 2000; van Kuijk et al. 2021). | |
| References:<br><br>Beatty, S.J., Morgan, D.L. & Gill, H.S., 2005, 'Role of life history strategy in the colonisation of Western Australian aquatic systems by the introduced crayfish *Cherax destructor* Clark, 1936', *Hydrobiologia* 549, 219-237. https://doi.org/10.1007/s10750-005-5443-0.<br>Larson, E.R. & Olden, J.D., 2016, 'Field sampling techniques for crayfish', In: Longshaw, M. & Stebbing, P. (eds), *Biology and Ecology of Crayfish 1st edn*., pp. 287- 323, CRC Press, Boca Raton.<br>McCormack, R.B., 2014, 'New records and review of the translocation of the yabby *Cherax destructor* into eastern drainages of New South Wales, Australia', *Australian Zoologist* 34, 85–94. https://doi.org/10.7882/AZ.2014.006.<br>Mills, B.J., 1983, 'Aquaculture of yabbies', In: *Proceedings of the First Freshwater Aquaculture Workshop*, Department of Agriculture, New South Wales. Australia.<br>Withnall, F., 2000, 'Biology of yabbies (*Cherax destructor*)', *Aquaculture notes of the department of Natural Resources and Environment*. State of Victoria, Australia.<br>Van Kuijk T., Biesmeijer, J.C., van der Hoorn, B.B.& Verdonschot, P.M.F., 2021,' Functional traits explain crayfish invasive success in the Netherlands', *Scientific Reports* 11, 2772. https://doi.org/10.1038/s41598-021-82302-4. | |

| **MAN4 Has the feasibility of eradication been evaluated?** | |
|---|---|
| Response: No | Confidence: High |
| Rationale: There has been no eradication feasibility study conducted for *C. destructor* in South Africa as wild populations have not been detected (Nunes et al. 2017). | |
| References:<br><br>Nunes, A.L., Zengeya, T.A., Measey, G.J. & Weyl, O.L.F., 2017, 'Freshwater crayfish invasions in South Africa: Past, present and potential future', *African Journal of Aquatic Science* 42, 309–323. https://doi.org/10.2989/16085914.2017.1405788. | |

| **MAN5 Control options and monitoring approaches available for the *Taxon*** |
|---|
| Response: Several methods can be used to monitor and control invasive crayfish populations including mechanical trapping (e.g., Barkhuizen et al. 2022; Madzivanzira et al. 2021), electrofishing (Nunes et al. 2017), use of biocides, and biological control (Aquiloni et al 2010; Stebbing et al. 2014). However, there are several issues that make management of crayfish invasions challenging (Gherardi et al. 2011). For example, the use of biocides is often effective for localised populations in dams and ponds (Peay et al. 2019) but they are non-specific and need to be applied with caution (Gherardi et al. 2011). Mechanical control methods are also effective in small discrete water bodies but are expensive and require commitment to follow on to complete eradication (Loureiro et al. 2018). In addition, alien crayfish species are regarded as potential conflict-generating species in South Africa and potential management interventions should consider the concerns of stakeholders to avoid conflict (Zengeya et al. 2017). |
| References:<br>Aquiloni, L., Brusconi, S., Cecchinelli, E., Tricarico, E., Mazza, G., Paglianti, A., et al., 2010, 'Biological control of invasive populations of crayfish: the European eel (*Anguilla anguilla*) as a predator of *Procambarus clarkii*', *Biological Invasions*, 12, 3817-3824. https://doi.org/ 10.1007/s10530-010-9774-z.<br>Barkhuizen, L.M., Madzivanzira, T.C. & South, J. 2022, 'Population ecology of a wild population of red swamp crayfish *Procambarus clarkii* (Girard, 1852) in the Free State Province, South Africa and implications for eradication efforts', *BioInvasions Records* 11, 181–191https://doi.org/10.3391/bir.2022.11.1.18<br>Gherardi, F., Aquiloni, L., Diéguez-Uribeondo, J. & Tricarico E., 2011, 'Managing invasive crayfish: is there a hope?', *Aquatic Sciences* 73, 185–200. https://doi.org/10.1007/s00027-011-0181-z<br>Loureiro, T.G., Anastácio, P.M., de Siqueira Bueno, S.L. & Araujo, P.B., 2018, 'Management of invasive populations of the freshwater crayfish *Procambarus clarkii* (Decapoda, Cambaridae): test of a population-control method and proposal of a standard monitoring approach', *Environmental Monitoring Assessment* 190, 559, https://doi.org/10.1007/s10661-018-6942-6.<br>Madzivanzira, T.C, South, J., Nhiwatiwa, T. & Weyl, O.L.F., 2021, 'Standardisation of Australian redclaw crayfish *Cherax quadricarinatus* sampling gear in southern Africa', *Water SA* 47, 380–384. https://hdl.handle.net/10520/ejc-waters-v47-n3-a10.<br>Nunes, A.L., Zengeya, T.A., Hoffman, A.C., Measey, G.J. & Weyl, O.L., 2017, 'Distribution and establishment of the alien Australian redclaw crayfish, *Cherax quadricarinatus*, in South Africa and Swaziland', *PeerJ* 5, 1-21. https://doi.org/10.7717/peerj.3135.<br>Peay, S., Johnsen, S.I, Bean, S.W., Dunn, A.M., Sandodden, R. & Edsman, L, 2019, 'Biocide treatment of invasive signal crayfish: Successes, failures and lessons learned', *Diversity* 11, 29. https://doi.org/10.3390/d11030029<br>Stebbing, P., Longshaw, M. & Scott, A., 2014, 'Review of methods for the management of nonindigenous crayfish, with particular reference to Great Britain', *Ethology Ecology & Evolution* 26, 204-231. https://doi.org/10.1080/03949370.2014.908326.<br>Zengeya, T., Ivey, P., Woodford, D.J., Weyl, O, Novoa, A., Shackleton, R., et al., 2017, 'Managing conflict-generating invasive species in South Africa: Challenges and trade-offs', *Bothalia* 47, a2160. https://hdl.handle.net/10520/EJC-727738f3d |

| **MAN6 Any other management considerations to highlight?** | |
|---|---|
| Response | No |

#### 5. Calculations

**Likelihood = Fairly probable**

| Parameter | Likelihood | Stages | Final assessment |
|---|---|---|---|
| LIK1 | 0.0027 | P(entry) = 0.5 | P (invasion) = 0.25 |
| LIK2 | 0.5 | | |
| LIK3 | 1 | P(establishment) = 1 | |
| LIK4 | 1 | | |
| LIK5 | 0.5 | P (spread) =0.5 | |
| LIK6 | 0.5 | | |

**Consequence = Major (MR)**

| Parameter | Mechanism/sector | Response |
|---|---|---|
| CON1a | Competition | DD |
| CON1b | Predation | DD |
| CON1c | Hybridisation | DD |
| CON1d | Disease transmission | DD |
| CON1e | Parasitism | DD |
| CON1f | Poisoning/toxicity | DD |
| CON1g | Bio-fouling or other direct physical disturbance | DD |
| CON1h | Grazing/herbivory/browsing | DD |
| CON1i | Chemical, physical, structural impact | DD |
| CON1k | Indirect impacts through interactions with other species | DD |
| CON1 | Maximum environmental impact | DD |
| CON2a | Safety | DD |
| CON2b | Material and immaterial assets | DD |
| CON2c | Health | DD |
| CON2d | Social, spiritual and cultural relations | DD |
| CON2 | Maximum socio-economic impact | DD |
| CON3 | Environmental impact of closely related taxa | MR |
| CON4 | Socio-economic impact of closely related taxa | MO |
| CON5 | Potential impact based on traits, experiments, or models | MR |

**Risk score = High**

| | | Consequences | | | | |
|---|---|---|---|---|---|---|
| | | MC | MN | MO | MR | MV |
| Likelihood | Extremely unlikely | low | low | low | medium | medium |
| | Very unlikely | low | low | low | medium | high |
| | Unlikely | low | low | medium | high | high |
| | Fairly probable | medium | medium | high | high | high |
| | Probable | medium | high | high | high | high |

**Ease of management = Difficult**

| Parameter | Question | Response |
|---|---|---|
| MAN3a | How accessible are populations? | 2 |
| MAN3b | Is detectability critically time-dependent? | 2 |
| MAN3c | Time to reproduction | 2 |
| MAN3d | Propagule persistence | 0 |
| **MAN3** | **SUM** | 6 |

**Appendix BAC7**: Native range of *Cherax* species in Australia. Source: Wingfield, M., 2002, 'An overview of the Australian freshwater crayfish farming industry', *Freshwater Crayfish* 13(1), 177-184.

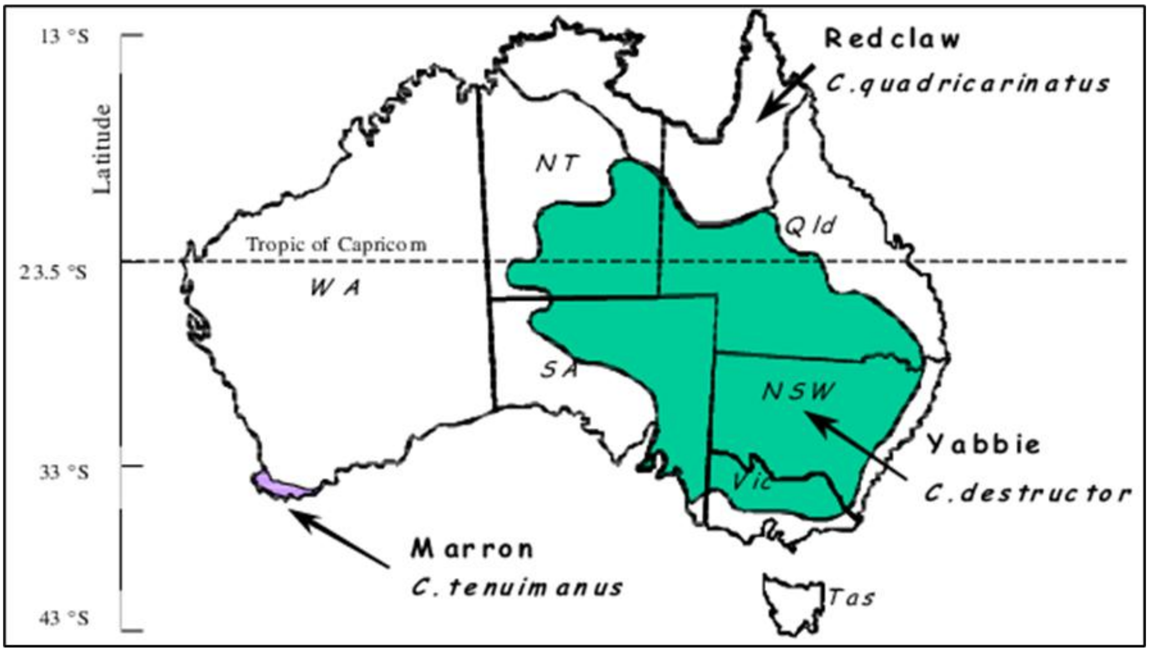

**Appendix BAC8(a)**: Global alien range of *Cherax destructor.* Source: CABI, 2021. *Cherax destructor* [original text by Uma Sabapathy Allen]. In: Invasive Species Compendium. Wallingford, UK: CAB International. www.cabi.org/isc. It is noted that recent field survey results did not show *C. destructor* being present in South Africa (Nunes et al. 2017).

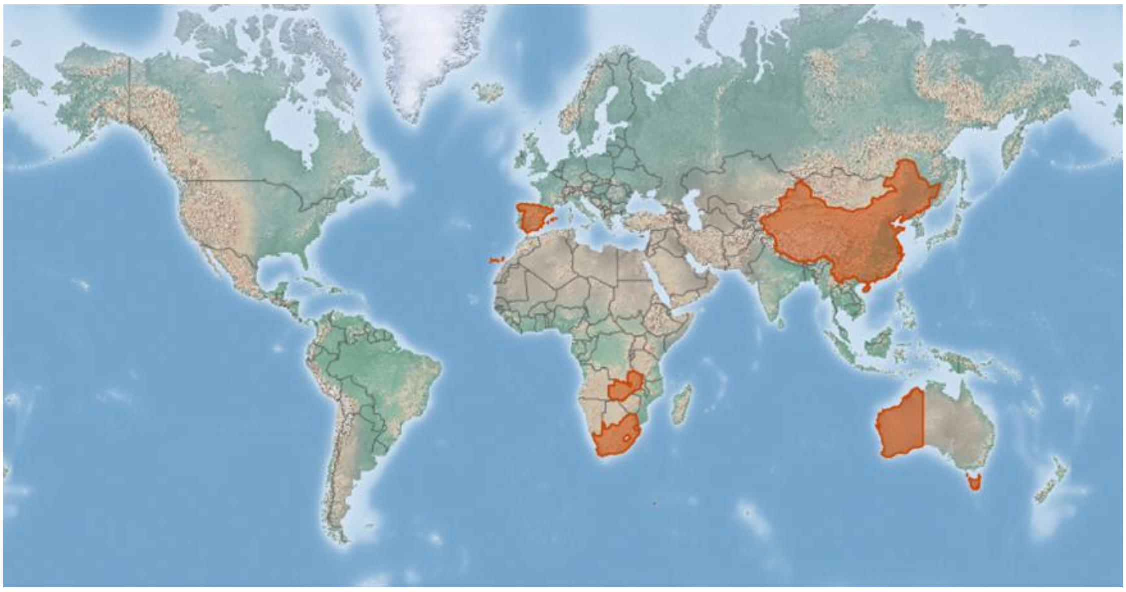

Appendix BAC8(b) Historic records for freshwater crayfish introductions in South Africa (including *Cherax destructor* in blue.) It is noted that recent field survey results did not show *C. destructor* being present in South Africa (Source: Nunes, A.L., Zengeya, T.A., Measey, G.J. & Weyl, O.L.F., 2017, 'Freshwater crayfish invasions in South Africa: Past, present and potential future', *African Journal of Aquatic Science* 42, 309–323. https://doi.org/10.2989/16085914.2017.1405788).

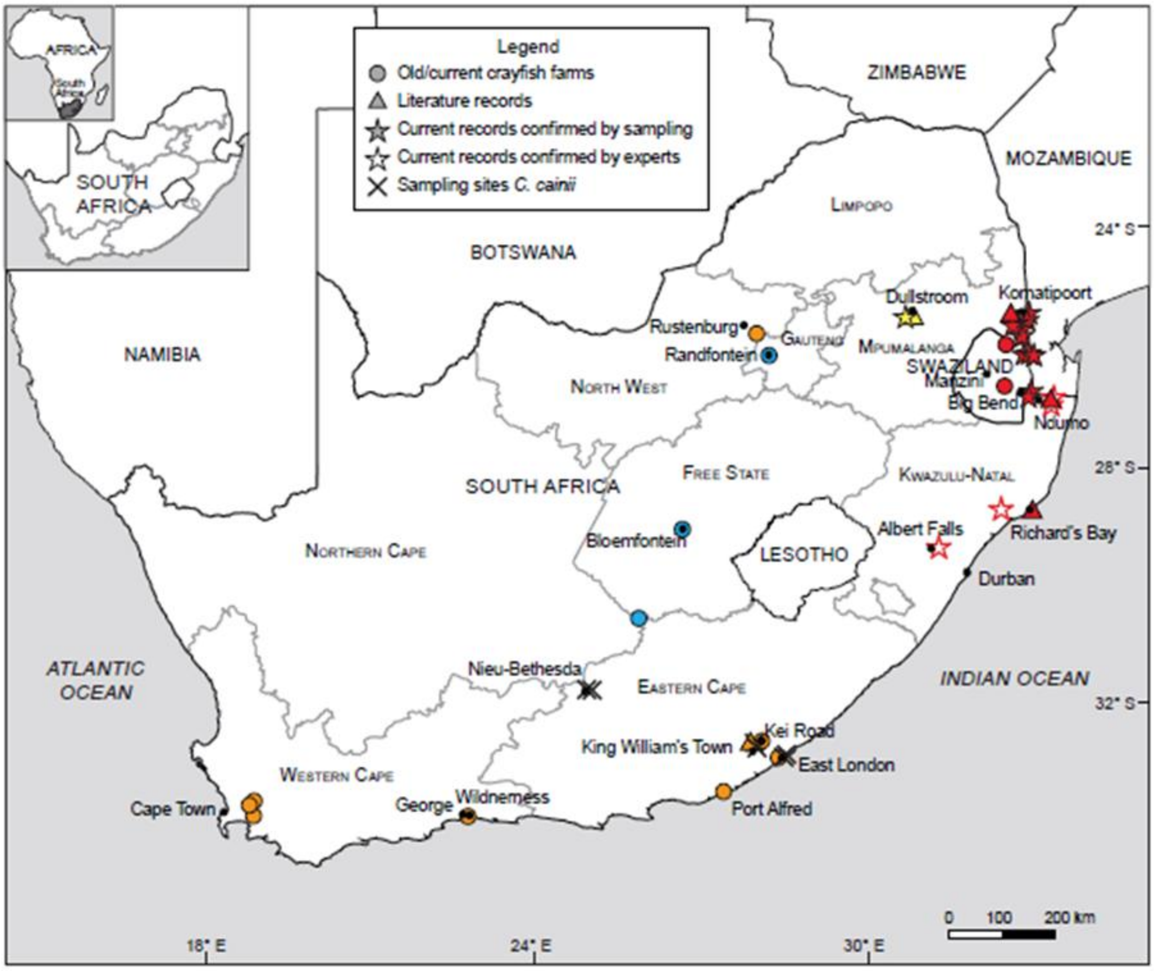

### 1.4 Risk analysis report for redclaw crayfish (*Cherax quadricarinatus*)

| | |
|---|---|
| ***Taxon:*** *Cherax quadricarinatus* (von Martens,1868) | ***Area:*** South Africa |
| **Compiled by:** Tsungai Zengeya | **Approved by:** |
| **Picture of *Taxon***<br>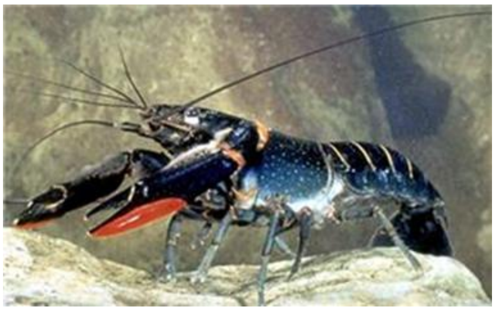<br>CABI, 2021. *Cherax quadricarinatus* [original text by Clive Jones]. In: Invasive Species Compendium. Wallingford, UK: CAB International. www.cabi.org/isc. | **Alien distribution map**<br>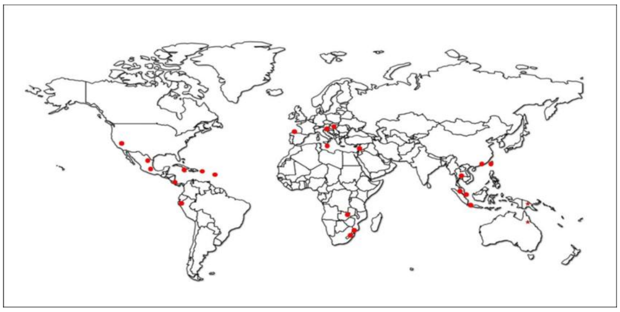<br>Reference: Sallehuddin, A.S., Kamarudin, A.S & Ismail, N., 2021, 'Review on the global distribution of wild population of Australian Redclaw Crayfish, *Cherax quadricarinatus* (von Martens, 1868)', *Bioscience Research* 18, 194–207. |

| | |
|---|---|
| **Risk Assessment summary:**<br><br>*Cherax quadricarinatus* (redclaw crayfish) has been introduced for aquaculture worldwide and is popular in the aquarium and pet trade. *Cherax quadricarinatus* was introduced into South Africa in 1988 for research purposes and is now established in South Africa and neighbouring countries. *Cherax quadricarinatus* has a high environmental tolerance, occupying a wide variety of habitats. *Cherax quadricarinatus* has negative impacts on invaded ecosystems through competition, predation, and transmission of diseases, and also on local fisheries by replacing high-value native species, damaging nets, and spoiling fish catches. | **Risk score:**<br><br>High |
| **Management summary:**<br><br>*Cherax quadricarinatus* is difficult to manage given that it is difficult to access, difficult to detect (being aquatic and primarily nocturnal) and has adaptive life history traits that aid establishment (fast growth rate, high fecundity, and a multiple spawner). Several methods have been used to control alien populations of crayfish including trapping, electrofishing, biocides, and the use of biological control. However, most of these are likely to be ineffective for C. *quadricarinatus*. | **Ease of management:**<br><br>Difficult |
| **Recommendations:**<br><br>*Cherax quadricarinatus* poses a high risk and is difficult to manage. *Cherax quadricarinatus* is widespread across several catchments in South Africa and neighbouring countries and further spread is likely through connected waterways so eradication is infeasible. Given the low socio-economic | **Listing under NEM:BA A&IS lists of 2020:**<br><br>1b<br><br>**Recommended listing category:** |

| | |
|---|---|
| benefits, and high risk, it is recommended that *C. quadricarinatus* remains listed as category 1b. | 1b |

#### 1. Background

| **BAC1 Name of assessor(s)** | |
|---|---|
| Name of lead assessor | Lee-Anne Botha |
| Additional assessor (1) | Tsungai Zengeya |
| Additional assessor (2) | Christian Chimimba |
| **BAC2 Contact details of assessor (s)** | |
| Lead assessor | Organisational affiliation: University of Pretoria, Department of Zoology and Entomology/ South African National Biodiversity Institute (SANBI) |
| | |
| Additional assessor (1) | Organisational affiliation: South African National Biodiversity Institute (SANBI) |
| | |
| Additional assessor (2) | Organisational affiliation: Department of Zoology and Entomology, University of Pretoria and The South African National Biodiversity Institute (SANBI). |
| | |
| **BAC3 Name(s) and contact details of expert(s) consulted** | |
| Expert (1) | Name: |
| | email: |
| | Phone: |
| Expert (2) | Name: |
| | email: |
| | Phone: |
| Comments: none | |

| **BAC4 Scientific name of *Taxon* under assessment** | |
|---|---|
| *Taxon* name: *Cherax quadricarinatus* | Authority: (von Martens,1868) |
| Comments: The scientific name of the taxon under assessment is accepted (Crandall & De Grave 2017). | |
| References:<br>Crandall, K.A. & De Grave, S., 2017, 'An updated classification of the freshwater crayfishes (Decapoda: Astacidea) of the world, with a complete species list', *Journal of Crustacean Biology* 27, 615–653. https://doi.org/10.1093/jcbiol/rux070. | |
| **BAC5 Synonym(s) considered** | |
| Synonyms: *Astacus quadricarinatus* von Martens, 1868 | |
| Comments: The abovementioned synonym (*Astacus quadricarinatus*) was considered in the literature search for this assessment (GBIF 2023). | |
| References:<br>*Cherax quadricarinatus* (von Martens, 1868) in GBIF Secretariat. GBIF Backbone Taxonomy. Checklist dataset https://doi.org/10.15468/39omei accessed via GBIF.org on 2023-09-11. | |
| **BAC6 Common name(s) considered** | |
| Common names: Australian redclaw crayfish \| Queensland red claw \| red claw \| tropical blue crayfish \| blue lobster \| redclaw crayfish | |
| Comments: The abovementioned common names were considered in the literature search for this assessment (GBIF 2023). | |
| References:<br>*Cherax quadricarinatus* (von Martens, 1868) in GBIF Secretariat. GBIF Backbone Taxonomy. Checklist dataset https://doi.org/10.15468/39omei accessed via GBIF.org on 2023-09-11. | |
| **BAC7 What is the native range of the *Taxon*?** | |
| Response: Australia \| New Guinea | Confidence: High |
| Comments: *Cherax quadricarinatus* is native to north-eastern Queensland and the northern and eastern parts of the Northern Territory in Australia, and to the southern part of New Guinea (Riek 1969) | |

| |
|---|
| References:<br>Riek, E.F., 1969, 'The Australian freshwater crayfish (Crustacea: Decapoda: Parastacidae), with descriptions of new species', *Australian Journal of Zoology*, 17, 855-918. https://doi.org/10.1071/ZO9690855. |
| **BAC8 What is the global alien range of the *Taxon*?** |

| | |
|---|---|
| Response: Asia \| Europe \| America \| Africa | Confidence: High |

| |
|---|
| Comments: *Cherax quadricarinatus* has been introduced to Malaysia, Indonesia, Singapore, Thailand, Taiwan, Hong Kong, Italy, Hungary, Spain, Slovenia, Puerto Rico, Jamaica, United States (Lake Balboa), Mexico, Martinique Island, Costa Rica, Morocco, Zambia, Zimbabwe, Mozambique, Eswatini, and South Africa (Haubrock et al. 2021; Lodge et al. 2012; Nunes et al. 2017; Madzivanzira et al. 2021; Sallehuddin et al. 2021). |
| References:<br>Haubrock, P.J., Oficialdegui, F.J., Zeng, Y., Patoka, J., Yeo, D.C.J. & Kouba, A. 2021, 'The redclaw crayfish: A prominent aquaculture species with invasive potential in tropical and subtropical biodiversity hotspots', *Reviews in Aquaculture* 13, 1488-1530https://doi.org/10.1111/raq.12531.<br>Lodge, D.M., Deines, A., Gherardi, F., Yeo, D.C.J., Arcella, T., Baldridge, A.K., et al., 2012, 'Global introductions of crayfishes: Evaluating the impact of species invasions on ecosystem services', *Annual Review of Ecology, Evolution and Systematics*, 43, 449-472. https://doi.org/10.1146/annurev-ecolsys-111511-103919.<br>Madzivanzira, T.C., South, J,, Ellender. B.R., Chalmers, R., Chisule, G., Coppinger, C.R., et al., 2021, 'Distribution and establishment of the alien Australian redclaw crayfish, *Cherax quadricarinatus*, in the Zambezi Basin', *Aquatic conservation: Marine and Freshwater Ecosystems*, 31(11), 3156-3168. https://doi.org/10.1002/aqc.3703.<br>Nunes, A.L., Zengeya, T.A., Hoffman, A.C., Measey, G.J. & Weyl, O.L.F., 2017, 'Distribution and establishment of the alien Australian redclaw crayfish, *Cherax quadricarinatus*, in South Africa and Swaziland', *PeerJ*, 5, e3135., http://dx.doi.org/10.7717/peerj.3135<br>Sallehuddin, A.S., Kamarudin, A.S & Ismail, N., 2021, 'Review on the global distribution of wild population of Australian Redclaw Crayfish, *Cherax quadricarinatus* (von Martens, 1868)', *Bioscience Research* 18, 194–207. |
| **BAC9 Geographic scope = the *Area* under consideration** |
| *Area* of assessment: South Africa |
| Comments: The geographic scope of this assessment is limited to South Africa. |
| **BAC10 Is the *Taxon* present in the *Area*?** |

| | |
|---|---|
| Response: Yes | Confidence: High |

| |
|---|
| Comments: *Cherax quadricarinatus* is widespread in the Inkomati and adjacent river systems in South Africa and neighbouring countries such as Mozambique, Eswatini, Zambia, and Zimbabwe (de Moor 2002; Madzivanzira et al. 2021; Nunes et al. 2017a, 2017b). |
| References:<br>de Moor, I., 2002, 'Potential impacts of alien freshwater crayfish in South Africa', *African Journal of Aquatic Science* 27, 125–139. https://doi.org/10.2989/16085914.2002.9626584.<br>Madzivanzira, T.C., South, J,, Ellender. B.R., Chalmers, R., Chisule, G., Coppinger, C.R., et al., 2021, 'Distribution and establishment of the alien Australian redclaw crayfish, *Cherax quadricarinatus*, in the Zambezi Basin', *Aquatic conservation: Marine and Freshwater Ecosystems*, 31(11), 3156-3168. https://doi.org/10.1002/aqc.3703.<br>Nunes, A.L., Zengeya, T.A., Hoffman, A.C., Measey, G.J. & Weyl, O.L., 2017a, 'Distribution and establishment of the alien Australian redclaw crayfish, *Cherax quadricarinatus*, in South Africa and Swaziland', *PeerJ* 5, 1-21. http://dx.doi.org/10.7717/peerj.3135.<br>Nunes, A.L., Zengeya, T.A., Measey, G.J. & Weyl, O.L.F., 2017b, 'Freshwater crayfish invasions in South Africa: Past, present and potential future', *African Journal of Aquatic Science* 42, 309-323. https://doi.org/10.2989/16085914.2017.1405788. |

| **BAC11 Availability of physical specimen** | | |
|---|---|---|
| Response: Albany Museum, Grahamstown | | Confidence in ID: Medium |
| Herbarium or museum accession number: GEN 1565A | | |
| References:<br>Albany Museum, Grahamstown [personal communication] | | |
| **BAC12 Is the *Taxon* native to the *Area* or part of the *Area*?** | | |
| The *Taxon* is native to (part of) the *Area*. | No | Confidence: High |
| The *Taxon* is alien in (part of) the *Area*. | Yes | Confidence: High |
| Comments: There are no native freshwater crayfish species in South Africa (de Moor 2002; Nunes et al. 2017) | | |
| References:<br>de Moor, I., 2002, 'Potential impacts of alien freshwater crayfish in South Africa', African *Journal of Aquatic Science* 27, 125–139. https://doi.org/10.2989/16085914.2002.9626584.<br>Nunes, A.L., Zengeya, T.A., Hoffman, A.C., Measey, G.J. & Weyl, O.L., 2017, 'Distribution and establishment of the alien Australian redclaw crayfish, *Cherax quadricarinatus*, in South Africa and Swaziland', *PeerJ* 5, 1-21. http://dx.doi.org/10.7717/peerj.3135. | | |
| **BAC13 What is the *Taxon*'s introduction status in the *Area*?** | | |
| The *Taxon* is in cultivation/containment. | Yes | Confidence: High |
| The *Taxon* is present outside of cultivation/containment. | Yes | Confidence: High |
| The *Taxon* has established/naturalised. | Yes | Confidence: High |
| The *Taxon* is invasive. | Yes | Confidence: High |
| Comments: *Cherax quadricarinatus* was introduced in 1988 for research (Nunes et al. 2017) and there are still several valid permits for possession and research (plus one for breeding) (SANBI 2023). *Cherax quadricarinatus* is invasive and widespread in the Inkomati River Catchment and adjacent rivers in South Africa and neighbouring countries such as Mozambique, Eswatini, Zambia, and Zimbabwe (Nunes et al. 2017; Douthwaite et al. 2018; Marufu et al. 2018; Madzivanzira et al. 2021). | | |
| References:<br>Douthwaite, R.J., Jones, E.W., Tyser, A.B. & Vrdoljak, S.M., 2018, 'The introduction, spread and ecology of redclaw crayfish *Cherax quadricarinatus* in the Zambezi catchment', *African Journal of Aquatic Science* 43, 353–366 https://doi.org/10.2989/16085914.2018.1517080<br>Madzivanzira, T.C., South, J,, Ellender. B.R., Chalmers, R., Chisule, G., Coppinger, C.R., et al., 2021, 'Distribution and establishment of the alien Australian redclaw crayfish, *Cherax quadricarinatus*, in the Zambezi Basin', *Aquatic conservation: Marine and Freshwater Ecosystems*, 31(11), 3156-3168. https://doi.org/10.1002/aqc.3703.<br>Marufu, L., Barson, M., Chifamba, P., Tiki, M. & Nhiwatiwa, T., 2018, 'The population dynamics of a recently introduced crayfish, *Cherax quadricarinatus* (von Martens, 1868), in the Sanyati Basin of Lake Kariba, Zimbabwe', *African Zoology* 53, 17–22. https://doi.org/10.1080/15627020.2018.1448719.<br>Nunes, A.L., Zengeya, T.A., Hoffman, A.C., Measey, G.J. & Weyl, O.L., 2017, 'Distribution and establishment of the alien Australian redclaw crayfish, *Cherax quadricarinatus*, in South Africa and Swaziland', *PeerJ* 5, 1-21, http://dx.doi.org/10.7717/peerj.3135<br>SANBI (2023) Permits issued in respect of South Africa's National Environmental Management: Biodiversity Act's Alien and Invasive Species Regulations over the period October 2014 – December 2022. version 20230809. Zenodo. https://dx.doi.org/10.5281/zenodo.8229321 | | |
| **BAC14 Primary (introduction) pathways** | | |
| Release | none | Confidence: High |
| Escape | aquaculture/mariculture | Confidence: High |

| | | |
|---|---|---|
| | pet/aquarium/terrarium species (including live food for such species) | Confidence: High |
| | live food and live bait | Confidence: High |
| Contaminant | none | Confidence: High |
| Stowaway | none | Confidence: High |
| Corridor | none | Confidence: High |
| Unaided | natural dispersal across borders of invasive alien organism | Confidence: High |
| Comments: *Cherax quadricarinatus* is introduced for aquaculture purposes (FAO 2023; Haubrock et al. 2021), is sold as pets in various countries, and used as live bait by anglers (Akmal et al. 2023). | | |

References:

Akmal, S.G., Yonvitner, Yulianda, F., Adrianto, L. & Patoka, J., 2023, 'Potential spread of invasive crayfish used as life bait by Indonesian anglers', *Human Dimensions of Wildlife*, 1-13. https://doi.org/10.1080/10871209.2023.2234394.

FAO 2023. PARASTACIDAE. Cultured Aquatic Species Information Programme. Text by Jones, C. Fisheries and Aquaculture Division [online]. Rome. Updated 2011-10-10 [Accessed Thursday, September 7th 2023]. https://www.fao.org/fishery/en/culturedspecies/cherax_quadricarinatus/en

Haubrock, P.J., Oficialdegui, F.J., Zeng, Y., Patoka, J., Yeo, D.C.J. & Kouba, A. 2021, 'The redclaw crayfish: A prominent aquaculture species with invasive potential in tropical and subtropical biodiversity hotspots', *Reviews in Aquaculture* 13, 1488-1530, https://doi.org/10.1111/raq.12531.

**2. Likelihood**

| **LIK1 Likelihood of entry via unaided primary pathways** | |
|---|---|
| Response: Present in the country (p = 1) | Confidence: High |
| Rationale: *Cherax quadricarinatus* is present in the country and has established populations in natural environments (Nunes et al. 2017; Petersen et al. 2017). In terms of the likelihood of future introductions, there are populations outside of captivity in several countries in southern Africa (Douthwaite et al. 2018; Madzivanzira et al. 2021; Marufu et al. 2018). *Cherax quadricarinatus* is capable of spreading naturally via interconnected waterways (Nunes et al. 2017), therefore it is probable (p = 1) that *C. quadricarinatus* will re-enter the country via unaided pathways. | |

References:

Douthwaite, R.J., Jones, E.W., Tyser, A.B. & Vrdoljak, S.M., 2018, 'The introduction, spread and ecology of redclaw crayfish *Cherax quadricarinatus* in the Zambezi catchment', *African Journal of Aquatic Science* 43, 353–366. https://doi.org/10.2989/16085914.2018.1517080.

Madzivanzira, T.C., South, J,, Ellender. B.R., Chalmers, R., Chisule, G., Coppinger, C.R., et al., 2021, 'Distribution and establishment of the alien Australian redclaw crayfish, *Cherax quadricarinatus*, in the Zambezi Basin', *Aquatic conservation: Marine and Freshwater Ecosystems*, 31(11), 3156-3168. . https://doi.org/10.1002/aqc.3703.

Marufu, L., Barson, M., Chifamba, P., Tiki, M. & Nhiwatiwa, T., 2018, 'The population dynamics of a recently introduced crayfish, *Cherax quadricarinatus* (von Martens, 1868), in the Sanyati Basin of Lake Kariba, Zimbabwe', *African Zoology* 53, 17–22. https://doi.org/10.1080/15627020.2018.1448719.

Nunes, A.L., Zengeya, T.A., Hoffman, A.C., Measey, G.J. & Weyl, O.L., 2017, 'Distribution and establishment of the alien Australian redclaw crayfish, *Cherax quadricarinatus*, in South Africa and Swaziland', *PeerJ* 5, 1-21. http://dx.doi.org/10.7717/peerj.3135.

Petersen, R.M., Hoffman, A.C., Kotze, P. & Marr, S.M., 2017, 'First record of the invasive Australian redclaw crayfish *Cherax quadricarinatus* (von Martens, 1868) in the Crocodile River, Kruger National Park, South Africa', *Koedoe* 59, 1–3. https://hdl.handle.net/10520/EJC-731116ccd.

| LIK2 Likelihood of entry via human aided primary pathways | |
|---|---|
| Response: Present in the country (p = 1) | Confidence: High |
| Rationale: *Cherax quadricarinatus* is currently present in the country (Nunes et al. 2017; Petersen et al. 2017a). In terms of the likelihood of future introductions into South Africa, *C. quadricarinatus* is likely to be introduced for aquaculture and the pet trade, from which escapees are likely to occur (Nunes et al. 2017; Petersen et al. 2017). In addition, crayfish are used as bait for fishing in neighbouring countries (Douthwaite et al. 2018). | |
| References:<br>Douthwaite, R.J., Jones, E.W., Tyser, A.B. & Vrdoljak, S.M., 2018, 'The introduction, spread and ecology of redclaw crayfish *Cherax quadricarinatus* in the Zambezi catchment', *African Journal of Aquatic Science* 43, 353–366. https://doi.org/10.2989/16085914.2018.1517080.<br>Nunes, A.L., Zengeya, T.A., Hoffman, A.C., Measey, G.J. & Weyl, O.L., 2017, 'Distribution and establishment of the alien Australian redclaw crayfish, *Cherax quadricarinatus*, in South Africa and Swaziland', *PeerJ* 5, 1-21. https://doi.org/10.7717/peerj.3135.<br>Petersen, R.M., Hoffman, A.C., Kotze, P. & Marr, S.M., 2017, 'First record of the invasive Australian redclaw crayfish *Cherax quadricarinatus* (von Martens, 1868) in the Crocodile River, Kruger National Park, South Africa', *Koedoe* 59, 1–3. https://hdl.handle.net/10520/EJC-731116ccd | |

| LIK3 Habitat suitability | |
|---|---|
| Response: Probable (p = 1) | Confidence: High |
| Rationale: *Cherax quadricarinatus* can occupy a wide range of aquatic systems including ponds and rivers (Sallehuddin et al. 2021). *Cherax quadricarinatus* prefers rocky habitats with plenty of crevices for shelter and foraging, especially during the moulting phase (de Moor 2002). These are present across the country (Nunes et al. 2017). | |
| References:<br>de Moor, I., 2002, 'Potential impacts of alien freshwater crayfish in South Africa', *African Journal of Aquatic Science* 27, 125–139. https://doi.org/10.2989/16085914.2002.9626584.<br>Nunes, A.L., Zengeya, T.A., Hoffman, A.C., Measey, G.J. & Weyl, O.L., 2017, 'Distribution and establishment of the alien Australian redclaw crayfish, *Cherax quadricarinatus*, in South Africa and Swaziland', *PeerJ* 5, 1-21. https://doi.org/10.7717/peerj.3135.<br>Sallehuddin, A.S., Kamarudin, A.S & Ismail, N., 2021, 'Review on the global distribution of wild population of Australian Redclaw Crayfish, *Cherax quadricarinatus* (von Martens, 1868)', *Bioscience Research* 18, 194–207. | |

| LIK4 Climate suitability | |
|---|---|
| Response: Probable (p = 1) | Confidence: High |
| Rationale: The thermal tolerance for *C. quadricarinatus* is between 10°C and 34°C, but the optimal thermal range is between 22°C and 32°C (FAO 2023). Based on the niche models produced by Nunes et al. (2017b), climatically suitable regions for *C. quadricarinatus* were largely restricted to the north and eastern parts of South Africa, including the Soutpansberg, North Eastern Highlands, Eastern Bankenveld, Highveld, Northern Coastal Belt and South Eastern Uplands (Nunes et al. 2017b). However, there is evidence that *C. quadricarinatus* is adaptable to new temperature regimes if moved [e.g., in Tasmania feral populations of *C. quadricarinatus* have been reported to survive in water temperatures as low as 2°C to 4°C (de Moor 2002)]. | |
| References: | |

de Moor, I., 2002, 'Potential impacts of alien freshwater crayfish in South Africa', *African Journal of Aquatic Science* 27, 125–139, https://doi.org/10.2989/16085914.2002.9626584

FAO 2023. PARASTACIDAE. Cultured Aquatic Species Information Programme. Text by Jones, C. Fisheries and Aquaculture Division [online]. Rome. Updated 2011-10-10 [Accessed Thursday, September 7th 2023]. https://www.fao.org/fishery/en/culturedspecies/cherax_quadricarinatus/en

Nunes, A.L., Zengeya, T.A., Measey, G.J. & Weyl, O.L.F., 2017b, 'Freshwater crayfish invasions in South Africa: Past, present and potential future', *African Journal of Aquatic Science* 42, 309-323. https://doi.org/10.2989/16085914.2017.1405788.

| **LIK5 Unaided secondary (dispersal) pathways** | |
|---|---|
| Response: Probable (p = 1) | Confidence: High |
| Rationale: Similar to other crayfish species, *C. quadricarinatus* is highly mobile and can disperse via connected waterways and overland (Lodge et al. 2012). | |
| References:<br><br>Lodge, D.M., Deines, A., Gherardi, F., Yeo, D.C., Arcella, T., Baldridge, A.K., et al., 2012, 'Global introductions of crayfishes: evaluating the impact of species invasions on ecosystem services', *Annual Review of Ecology, Evolution, and Systematics* 43**,** 449–472. https://doi.org/10.1146/annurev-ecolsys-111511-103919. | |

| **LIK6 Human aided secondary (dispersal) pathways** | |
|---|---|
| Response: Probable (p = 1) | Confidence: High |
| Rationale: *Cherax quadricarinatus* may be introduced to new locations for purposes of aquaculture, via the aquarium and pet trade, and as bait by anglers (Madzivanzira et al. 2020)—there are reports that *C. quadricarinatus* has been released by anglers into dams (Nunes et al. 2017a). *Cherax quadricarinatus* is present in the pet trade industry in several countries globally, including South Africa and adjacent countries, leading to the intentional release of unwanted pets (Faulkes 2015). | |
| References:<br>Faulkes, Z., 2015, 'The global trade in crayfish as pets', *Crustacean Research* 44, 75–92. https://doi.org/10.18353/crustacea.44.0_75.<br>Madzivanzira, T.C., South, J., Wood, L.E., Nunes, A.L. & Weyl, O.L.F., 2020, 'A review of freshwater crayfish introductions in Africa', *Reviews in Fisheries Science & Aquaculture* 1-21. https://doi.org/10.1080/23308249.2020.1802405.<br>Nunes, A.L., Zengeya, T.A., Hoffman, A.C., Measey, G.J. & Weyl, O.L., 2017a, 'Distribution and establishment of the alien Australian redclaw crayfish, *Cherax quadricarinatus*, in South Africa and Swaziland', *PeerJ* 5, 1-21. https://doi.org/10.7717/peerj.3135. | |

**3. Consequences**

| **CON1 Environmental impact** | |
|---|---|
| **CON1a: Competition** | |
| Response: Minimal Concern (MC) | Confidence: Medium |
| Rationale: There is limited direct evidence for negative environmental impacts for *C. quadricarinatus* in its global invasive range and most of the cited impacts are based on potential rather than observed impacts (Haubrouck et al. 2021; Madzivanzira et al. 2021, 2022; Zengeya et al. 2022). | |

| | |
|---|---|
| References:<br>Haubrock, P.J., Oficialdegui, F.J., Zeng, Y., Patoka, J., Yeo, D.C.J. & Kouba, A. 2021, 'The redclaw crayfish: A prominent aquaculture species with invasive potential in tropical and subtropical biodiversity hotspots', *Reviews in Aquaculture* 13, 1488-1530. https://doi.org/10.1111/raq.12531.<br>Madzivanzira, T.C., South, J. & Weyl, O.L.F., 2021, 'Invasive crayfish outperform Potamonautid crabs at higher temperatures', *Freshwater Biology* 66(5), 978–991. https://doi.org/10.1111/ fwb.13691.<br>Madzivanzira, T.C., Weyl, O.L.F. & South, J., 2022, 'Ecological and potential socioeconomic impacts of two globally invasive crayfish', *NeoBiota* 72, 25–43. https://doi.org/10.3897/neobiota.72.71868<br>Zengeya, T.A., Lombard, R.J.H., Nelwamondo, V.E., Nunes, A.L., Measey, J.G. & Weyl, O.L.F., 2022, 'Trophic niche of an invasive generalist consumer: Australian redclaw crayfish, *Cherax quadricarinatus*, in the Inkomati River Basin, South Africa', *Austral Ecology* 47, 1480-1494. https://doi.org/10.1111/aec.13230. | |
| **CON1b: Predation** | |
| Response: Minimal Concern (MC) | Confidence: Medium |
| Rationale: *Cherax quadricarinatus* is omnivorous and feeds on plant and animal material and is likely to cause major impacts on invaded ecosystems through predation on invertebrates and fish (Haubrock et al. 2021, Madzivanzira et al. 2021, 2022; Zengeya et al. 2022) | |
| References:<br>Haubrock, P.J., Oficialdegui, F.J., Zeng, Y., Patoka, J., Yeo, D.C.J. & Kouba, A. 2021, 'The redclaw crayfish: A prominent aquaculture species with invasive potential in tropical and subtropical biodiversity hotspots', *Reviews in Aquaculture* 13, 1488-1530. https://doi.org/10.1111/raq.12531.<br>Madzivanzira, T.C., South, J. & Weyl, O.L.F., 2021, 'Invasive crayfish outperform Potamonautid crabs at higher temperatures', *Freshwater Biology* 66(5), 978–991. https://doi.org/10.1111/ fwb.13691.<br>Madzivanzira, T.C., Weyl, O.L.F. & South, J., 2022, 'Ecological and potential socioeconomic impacts of two globally invasive crayfish', *NeoBiota* 72, 25–43. https://doi.org/10.3897/neobiota.72.71868<br>Zengeya, T.A., Lombard, R.J.H., Nelwamondo, V.E., Nunes, A.L., Measey, J.G. & Weyl, O.L.F., 2022, 'Trophic niche of an invasive generalist consumer: Australian redclaw crayfish, *Cherax quadricarinatus*, in the Inkomati River Basin, South Africa', *Austral Ecology* 47, 1480-1494. https://doi.org/10.1111/aec.13230. | |
| **CON1c: Hybridisation** | |
| Response: Data Deficient (DD) | Confidence: High |
| Rationale: There are no native freshwater crayfish in South Africa, therefore hybridisation is not possible (de Moor 2002). | |
| References:<br>de Moor, I., 2002, 'Potential impacts of alien freshwater crayfish in South Africa', African *Journal of Aquatic Science* 27, 125–139. https://doi.org/10.2989/16085914.2002.9626584. | |
| **CON1d: Transmission of disease** | |
| Response: Major (MR) | Confidence: Medium |
| Rationale: *Cherax quadricarinatus* has been shown to cause major impacts to areas of introduction through transmission of disease (Haubrock et al. 2021). For example, *C. quadricarinatus* infected by the Decapod Iridescent Virus 1 (DIV1) have been implicated in the decline of crayfish and shrimp species in southern China (Qiu et al. 2019). Similarly, introductions of *C. quadricarinatus* infected with the *Cherax quadricarinatus bacilliform virus* (CqBV) have caused declines in aquaculture production (Edgerton & Owens 1997). | |

| |
|---|
| References:<br>Edgerton, B. & Owens, L., 1997,' Age at first infection of *Cherax quadricarinatus* by *Cherax quadricarinatus bacilliform* virus and *Cherax giardiavirus*-like virus, and production of putative virus-free crayfish', *Aquaculture* 152, 1–12. https://doi.org/10.1016/S0044-8486(97)00006-9.<br>Haubrock, P.J., Oficialdegui, F.J., Zeng, Y., Patoka, J., Yeo, D.C.J. & Kouba, A. 2021, 'The redclaw crayfish: A prominent aquaculture species with invasive potential in tropical and subtropical biodiversity hotspots', *Reviews in Aquaculture* 13, 1488-1530. https://doi.org/10.1111/raq.12531.<br>Longshaw, M., 2011, 'Diseases of crayfish: a review', *Journal of Invertebrate Pathology* 106, 54–70. https://doi.org/10.1016/j.jip.2010.09.013.<br>Saoud, I.P., Ghanawi, J., Thompson, K.R. & Webster, C.D., 2013, 'A review of culture and diseases of redclaw crayfish *Cherax quadricarinatus* (Von Martens 1868)', *Journal of the World Aquaculture Society* 44, 1-29. https://doi.org/10.1111/jwas.12011.<br>Qiu, L., Chen, X., Zhao, R.-H., Li, C., Gao, W., Zhang, Q.L., et al., 2019, 'Description of a natural Infection with Decapod Iridescent Virus 1 in farmed giant freshwater prawn *Macrobrachium rosenbergii*', *Viruses* 11, 354. https://doi.org/10.3390/v11040354. |

| **CON1e: Parasitism** | |
|---|---|
| Response: Data Deficient (DD) | Confidence: High |
| Rationale: *Cherax quadricarinatus* is not a parasitic taxon (Haubrock et al. 2021) | |
| References:<br>Haubrock, P.J., Oficialdegui, F.J., Zeng, Y., Patoka, J., Yeo, D.C.J. & Kouba, A. 2021, 'The redclaw crayfish: A prominent aquaculture species with invasive potential in tropical and subtropical biodiversity hotspots', *Reviews in Aquaculture* 13, 1488-1530. https://doi.org/10.1111/raq.12531. | |

| **CON1f: Poisoning/toxicity** | |
|---|---|
| Response: Minimal Concern (MC) | Confidence: High |
| Rationale: *Cherax quadricarinatus* can bio–accumulate toxins (e.g. cyanobacteria) in its tissues and organs (Saker & Eaglesham 1999) but there have been no recorded adverse impacts as a result of poisoning or toxicity from infected crayfish (Haubrock et al. 2021). | |
| References:<br>Saker, M.L. & Eaglesham, G,K., 1999, 'The accumulation of cylindrospermopsin from the cyanobacteria *Cylindrospermopsis raciborskii* in tissues of redclaw crayfish *Cherax quadricarinatus*', *Toxicon* 37, 1065-1077. https://doi.org/10.1016/S0041-0101(98)00240-2.<br>Haubrock, P.J., Oficialdegui, F.J., Zeng, Y., Patoka, J., Yeo, D.C.J. & Kouba, A. 2021, 'The redclaw crayfish: A prominent aquaculture species with invasive potential in tropical and subtropical biodiversity hotspots', *Reviews in Aquaculture* 13, 1488-1530. https://doi.org/10.1111/raq.12531. | |

| **CON1g: Bio-fouling or other direct physical disturbance** | |
|---|---|
| Response: Data Deficient (DD) | Confidence: High |
| Rationale: There are no recorded impacts by *C. quadricarinatus* through bio-fouling/physical disturbance (Haubrock et al. 2021) | |
| References:<br>Haubrock, P.J., Oficialdegui, F.J., Zeng, Y., Patoka, J., Yeo, D.C.J. & Kouba, A. 2021, 'The redclaw crayfish: A prominent aquaculture species with invasive potential in tropical and subtropical biodiversity hotspots', *Reviews in Aquaculture* 13, 1488-1530. https://doi.org/10.1111/raq.12531. | |

| CON1h: Grazing/herbivory/browsing | |
|---|---|
| Response: Minimal Concern (MC) | Confidence: Medium |
| Rationale: *Cherax quadricarinatus* is omnivorous and feeds on plant and animal material and is likely to cause moderate impacts on invaded ecosystems through grazing on macrophytes (Haubrock et al. 2021, Madzivanzira et al. 2022, Zengeya et al. 2022) | |
| References:<br><br>Haubrock, P.J., Oficialdegui, F.J., Zeng, Y., Patoka, J., Yeo, D.C.J. & Kouba, A. 2021, 'The redclaw crayfish: A prominent aquaculture species with invasive potential in tropical and subtropical biodiversity hotspots', *Reviews in Aquaculture* 13, 1488-1530. https://doi.org/10.1111/raq.12531.<br>Madzivanzira, T.C., Weyl, O.L.F. & South, J., 2022, 'Ecological and potential socioeconomic impacts of two globally invasive crayfish', *NeoBiota* 72, 25-43. https://doi.org/10.3897/neobiota.72.71868<br>Zengeya, T.A., Lombard, R.J.H., Nelwamondo, V.E., Nunes, A.L., Measey, J.G. & Weyl, O.L.F., 2022, 'Trophic niche of an invasive generalist consumer: Australian redclaw crayfish, *Cherax quadricarinatus*, in the Inkomati River Basin, South Africa', *Austral Ecology* 47, 1480-1494. https://doi.org/10.1111/aec.13230. | |
| **CON1i: Chemical, physical or structural impact on ecosystem** | |
| Response: Minimal Concern (MC) | Confidence: Medium |
| Rationale: Similar to other invasive crayfish, *C. quadricarinatus* is considered to be an ecosystem engineer due to its omnivorous feeding strategy that affects multiple trophic levels. For example, excessive grazing by *C. quadricarinatus* may cause habitat modification leading to cascading ecosystem impacts such as habitat loss for invertebrates and fish (Marufu et al. 2018; Weyl et al. 2017) and changes to nutrient cycling (Zengeya et al. 2022). Although *C. quadricarinatus* is not considered a burrowing species, it has been observed to construct burrows in Jamaica and La Réunion (Robinet 2010; Todd & D'Andrea 2003) that could potentially lead to adverse chemical, physical or structural impact on ecosystems as observed for other invasive crayfish (e.g. Souty-Grosset et al. 2016). However, Haubrouck et al. (2021) scored impacts of *C. quadricarinatus* through chemical, physical or structural impact on ecosystems as "Minimal Concern" because there is limited evidence of recorded impacts as most literature cites potential rather than observed impacts. This might change as more evidence becomes available. | |
| References:<br><br>Haubrock, P.J., Oficialdegui, F.J., Zeng, Y., Patoka, J., Yeo, D.C.J. & Kouba, A. 2021, 'The redclaw crayfish: A prominent aquaculture species with invasive potential in tropical and subtropical biodiversity hotspots', *Reviews in Aquaculture* 13, 1488-1530. https://doi.org/10.1111/raq.12531.<br>Marufu, L.T., Dalu, T., Phiri, C., Barson, M., Simango, R., Utete, B. et al., 2018, 'The diet of an invasive crayfish, *Cherax quadricarinatus* (Von Martens, 1868), in Lake Kariba, inferred using stomach content and stable isotope analyses', *BioInvasions Records* 7, 121–132. https://doi.org/10.3391/bir.2018.7.2.03.<br>Robinet, O., 2010, 'Stratégie de lutte contre les espèces invasives à la Réunion. Parc national Ile de La Réunion., France<br>Souty-Grosset, C., Anastacio, P.M., Aquiloni, L., Banha, F., Choquer, J., Chucholl, C. & Tricarico, E., 2016, 'The red swamp crayfish *Procambarus clarkii* in Europe: Impacts on aquatic ecosystems and human well–being', *Limnologica* 58, 78–93. , https://doi.org/10.1016/j.limno.2016.03.003.<br>Todd, S. & D'Andrea, M., 2003, 'Alien crayfish invasion of Jamaican rivers', *Crayfish News* 25, 17–18.<br>Weyl, O., Nunes, A., Ellender, B., Weyl, P., Chilala, A., Jacobs, F. et al., 2017, 'Why suggesting Australian redclaw crayfish *Cherax quadricarinatus* as biological control agents for snails is a bad idea', *African Journal of Aquatic Science* 42, 325–327. https://doi.org/10.2989/16085914.2017.1414685. | |

Zengeya, T,A., Lombard, R.J.H., Nelwamondo, V.E., Nunes, A.L., Measey, J.G., Weyl, O.L.F., 2022, 'Trophic niche of an invasive generalist consumer: Australian redclaw crayfish, *Cherax quadricarinatus*, in the Inkomati River Basin, South Africa', *Austral Ecology* 47, 1480-1494. https://doi.org/10.1111/aec.13230.

**CON1k: Indirect impacts through interactions with other species**

Response: Data Deficient (DD) | Confidence: Medium

Rationale:

References:

**CON1 Maximum environmental impact**

Response: Major (MR) | Confidence: Medium

Rationale: There is limited direct evidence of adverse impacts of *C. quadricarinatus* in its global invasive range – including South Africa - with most cited literature revealing circumstantial evidence for likely impacts on ecosystems through competition and predation (Haubrock et al. 2021). There is evidence however to show that *C. quadricarinatus* can cause major impacts to areas of introduction through transmission of disease (Haubrock et al. 2021).

References:

Haubrock, P.J., Oficialdegui, F.J., Zeng, Y., Patoka, J., Yeo, D.C.J. & Kouba, A. 2021, 'The redclaw crayfish: A prominent aquaculture species with invasive potential in tropical and subtropical biodiversity hotspots', *Reviews in Aquaculture* 13, 1488-1530. https://doi.org/10.1111/raq.12531.

**CON2 Socio-economic impact**

**CON2a: Safety**

Response: Data Deficient (DD) | Confidence: Medium

Rationale: There are no recorded impacts by *C. quadricarinatus* on human safety (Haubrock et al. 2021)

References:

Haubrock, P.J., Oficialdegui, F.J., Zeng, Y., Patoka, J., Yeo, D.C.J. & Kouba, A. 2021, 'The redclaw crayfish: A prominent aquaculture species with invasive potential in tropical and subtropical biodiversity hotspots', *Reviews in Aquaculture* 13, 1488-1530. https://doi.org/10.1111/raq.12531.

**CON2b: Material and immaterial assets**

Response: Moderate (MO) | Confidence: Medium

Rationale: Invasive crayfish can cause shifts in artisanal fisheries that can adversely affect the livelihood of fishers (Gherardi et al. 2011; Haubrock et al. 2021). For example, *C. quadricarinatus* has adversely affected local fisheries in Mexico (Vega-Villasante et al. 2015), Zambia, and Zimbabwe (Douthwaite et al. 2018) by replacing native species which have a higher economic value. In addition, *C. quadricarinatus* can also cause damage to gillnets and spoil fish catches resulting in some economic loss (Chakandinakira et al. 2023; Madzivanzira et al. 2022, 2023).

References:

Chakandinakira, A.T., Madzivanzira, T.C., Mashonga, S., Muzvondiwa, J.V., Ndlovu, N., & South, J. 2023, 'Socioeconomic impacts of Australian redclaw crayfish *Cherax quadricarinatus* in Lake Kariba', *Biological Invasions* 25, 2801-2812. https://doi.org/10.1007/s10530-023-03074-8.

Douthwaite, R., Jones, E., Tyser, A. & Vrdoljak, S., 2018, 'The introduction, spread and ecology of redclaw crayfish *Cherax quadricarinatus* in the Zambezi catchment', *African Journal of Aquatic Science* 43, 353–366. https://doi.org/10.2989/16085914.2018.1517080.

Gherardi, F., Britton, J.R., Mavuti, K.M., Pacini, N., Grey, J., Tricarico, E., et al., 2011, 'A review of allodiversity in Lake Naivasha, Kenya: Developing conservation actions to protect East African lakes from the negative impacts of alien species', *Biological* Conservation 144, 2585–2596. https://doi.org/10.1016/j.biocon.2011.07.020.

Haubrock, P.J., Oficialdegui, F.J., Zeng, Y., Patoka, J., Yeo, D.C.J. & Kouba, A. 2021, 'The redclaw crayfish: A prominent aquaculture species with invasive potential in tropical and subtropical biodiversity hotspots', *Reviews in Aquaculture* 13, 1488-1530. https://doi.org/10.1111/raq.12531.

Madzivanzira, T.C., Weyl, O.L.F. & South, J., 2022, 'Ecological and potential socioeconomic impacts of two globally invasive crayfish', *NeoBiota* 72, 25–43, https://doi.org/10.3897/neobiota.72.71868.

Madzivanzira, T.C., Chakandinakira, A.T., Mungenge, C.P., O'Brien, G., Dalu, T. & South, J., 2023, 'Get it before it gets to my catch: misdirection traps to mitigate against socioeconomic impacts associated with crayfish invasion', *Management of Biological Invasions* 14(2), 335–346. https://doi.org/10.3391/mbi.2023.14.2.10.

**CON2c: Health**

| Response: Data Deficient (DD) | Confidence: Medium |
|---|---|

Rationale: There are no recorded impacts of *C. quadricarinatus* on human health (Haubrock et al. 2021).

References:

Haubrock, P.J., Oficialdegui, F.J., Zeng, Y., Patoka, J., Yeo, D.C.J. & Kouba, A. 2021, 'The redclaw crayfish: A prominent aquaculture species with invasive potential in tropical and subtropical biodiversity hotspots', *Reviews in Aquaculture* 13, 1488-1530. https://doi.org/10.1111/raq.12531.

**CON2d: Social, spiritual and cultural relations**

| Response: Data Deficient (DD) | Confidence: |
|---|---|

Rationale: There are no recorded impact of *C. quadricarinatus* on human social, spiritual and cultural relations (Haubrock et al. 2021).

References:

Haubrock, P.J., Oficialdegui, F.J., Zeng, Y., Patoka, J., Yeo, D.C.J. & Kouba, A. 2021, 'The redclaw crayfish: A prominent aquaculture species with invasive potential in tropical and subtropical biodiversity hotspots', *Reviews in Aquaculture* 13, 1488-1530. https://doi.org/10.1111/raq.12531.

**CON2 Maximum socio-economic impact**

| Response: Moderate (MO) | Confidence: Medium |
|---|---|

Rationale: The presence of *C. quadricarinatus* in local fisheries can threaten the livelihoods of fishers by replacing native species which have higher economic value, damaging fishing gear and spoiling fish catches (Chakandinakira et al. 2023; Douthwaite et al. 2018; Madzivanzira et al. 2022; Vega-Villasante et al. 2015).

References:

Chakandinakira, A.T., Madzivanzira, T.C., Mashonga, S., Muzvondiwa, J.V., Ndlovu, N. & South, J. 2023, 'Socioeconomic impacts of Australian redclaw crayfish *Cherax quadricarinatus* in Lake Kariba', *Biological Invasions* 25, 2801-2812. https://doi.org/10.1007/s10530-023-03074-8

Douthwaite, R., Jones, E., Tyser, A. & Vrdoljak, S., 2018, 'The introduction, spread and ecology of redclaw crayfish *Cherax quadricarinatus* in the Zambezi catchment', *African Journal of Aquatic Science* 43, 353–366. https://doi.org/10.2989/16085914.2018.1517080.
Haubrock, P.J., Oficialdegui, F.J., Zeng, Y., Patoka, J., Yeo, D.C., & Kouba, A., 2021, 'The redclaw crayfish: A prominent aquaculture species with invasive potential in tropical and subtropical biodiversity hotspots', *Reviews in Aquaculture* 13(3), 1488-1530, https://doi.org/10.1111/raq.12531.
Madzivanzira, T.C., Weyl, O.L.F. & South, J., 2022, 'Ecological and potential socioeconomic impacts of two globally invasive crayfish', *NeoBiota* 72, 25–43. https://doi.org/10.3897/neobiota.72.71868.
Vega-Villasante, F., Avalos-Aguilar, J.J., Nolasco-Soria, H., Vargas-Ceballos, M.A., Bortolini-Rosales, J.L., Chong-Carrillo O. et al., 2015, 'Wild populations of the invasive Australian red claw crayfish *Cherax quadricarinatus* (Crustacea, Decapoda) near the northern coast of Jalisco, Mexico: a new fishing and profitable resource', *Latin American Journal of Aquatic Research* 43, 781–785. https://doi.org/10.3856/vol43-issue4-fulltext-17.

| **CON3 Closely related species' environmental impact** | |
| --- | --- |
| Response: Not scored as CON1 scored | Confidence: |
| Rationale: There is information to score CON1, so CON3 does not need to be scored as per the framework | |
| References: | |

| **CON4 Closely related species' socio-economic impact** | |
| --- | --- |
| Response: Not scored as CON2 scored | Confidence: |
| Rationale: There is information to score CON2, so CON4 does not need to be scored as per the framework. | |
| References: | |

| **CON5 Potential impact** | |
| --- | --- |
| Response: Major (MR) | Confidence: Medium |

Rationale:

Life history traits (high fecundity, fast growth, large size, omnivorous diet, and wide environmental tolerances) have been suggested to predispose *C. quadricarinatus* to have Major impacts on ecosystems through predation, competition, and transfer of diseases (e.g., Ahyong & Yeo 2007; Bortolini et al. 2007; Lynas et al. 2007; Yeo 2010; Madzivanzira et al. 2020, 2021; Marufu et al. 2018; Zeng et al. 2019; Zengeya et al. 2022). For example, *C. quadricarinatus* invasions have been implicated in the decline of native species of macrophytes, macroinvertebrates and fish (Douthwaite et al. 2018; Marufu et al. 2018; Pinder et al. 2019). In South Africa, there have been laboratory studies to quantify the impacts of *C. quadricarinatus* on various aquatic and terrestrial resources ubiquitous to South African ecosystems. In these studies, *C. quadricarinatus* exhibited deleterious impacts on various ecosystem resources including fish, macroinvertebrates, and aquatic plants, all of which are important for ecosystem structure and functioning (Madzivanzira et al. 2021, 2022).

*Cherax quadricarinatus* is a functional omnivore that can outcompete native species for shared resources, predate on macroinvertebrates and fish and modify habitats through intensive grazing (de Moor 2002; Madzivanzira et al. 2021, 2022). Crayfish species in general are known hosts of parasites such as commensal worms (*temnocephalidans* species) (Tavakol et al. 2016). These parasites can be harmful when transferred to native decapods such as crabs and shrimp resulting in decline of populations or affect their overall performance. *Cherax quadricarinatus* is a possible vector of parasites and diseases as it known to harbour recognised pathogens such protozoans, bacteria and viruses (e.g., Edgerton et al. 2002; du Preez & Smit 2013; Saoud & Ghanawi 2013).

References:

Ahyong, S.T. & Yeo, D.C.J., 2007, 'Feral populations of the Australian redclaw crayfish (*Cherax quadricarinatus* von Martens) in water supply catchments of Singapore', *Biological Invasions* 9, 943-946, https://doi.org/10.1007/s10530-007-9094-0.

Bortolini, J.L., Alvarez, F. & Rodríguez-Almaraz, G., 2007, 'On the presence of the Australian redclaw crayfish *Cherax quadricarinatus* in Mexico,, *Biological Invasions* 9, 615-620. https://doi.org/10.1007/s10530-006-9054-0.

de Moor, I., 2002, 'Potential impacts of alien freshwater crayfish in South Africa', African *Journal of Aquatic Science* 27, 125–139. https://doi.org/10.2989/16085914.2002.9626584

Douthwaite, R., Jones, E., Tyser, A. & Vrdoljak, S., 2018, 'The introduction, spread and ecology of redclaw crayfish *Cherax quadricarinatus* in the Zambezi catchment', *African Journal of Aquatic Science* 43, 353–366. https://doi.org/10.2989/16085914.2018.1517080.

Du Preez, L. & Smit, N., 2013, 'Double blow: Alien crayfish infected with invasive temnocephalan in South Africa waters', *African Journal of Science* 109, 1-4 https://hdl.handle.net/10520/EJC142057.

Edgerton, B.F., Evans, L.H., Stephens, F.J. & Overstreet, R.M., 2002, 'Synopsis of freshwater crayfish diseases and commensal organisms', *Aquaculture* 206, 57–135. https://doi.org/10.1016/S0044-8486(01)00865-1.

Haubrock, P.J., Oficialdegui, F.J., Zeng, Y., Patoka, J., Yeo, D.C.J. & Kouba, A. 2021, 'The redclaw crayfish: A prominent aquaculture species with invasive potential in tropical and subtropical biodiversity hotspots', *Reviews in Aquaculture* 13, 1488-1530. https://doi.org/10.1111/raq.12531.

Lynas, J., Storey, A. & Knott, B., 2007, 'Introduction and spread of crayfish (Parastacidae) in Western Australia and their potential to displace indigenous species', In: Gherardi F (ed.) *Biological invaders in inland waters: profiles, distribution and threats*, pp. 577-596, Springer, Dordrecht, Netherlands.

Madzivanzira, T.C., South, J. & Weyl, O.L.F., 2021, 'Invasive crayfish outperform Potamonautid crabs at higher temperatures', *Freshwater Biology* 66(5), 978–991. https://doi.org/10.1111/fwb.13691.

Madzivanzira, T.C., Weyl, O.L.F. & South, J., 2022, 'Ecological and potential socioeconomic impacts of two globally invasive crayfish', *NeoBiota* 72, 25–43. https://doi.org/10.3897/neobiota.72.71868

Marufu, L.T., Dalu, T., Phiri, C., Barson, M., Simango, R., Utete, B. et al., 2018, 'The diet of an invasive crayfish, *Cherax quadricarinatus* (Von Martens, 1868), in Lake Kariba, inferred using stomach content and stable isotope analyses', *BioInvasions Records* 7, 121–132. https://doi.org/10.3391/bir.2018.7.2.03.

Nunes, A.L., Zengeya, T.A., Hoffman, A.C., Measey, G.J. & Weyl, O.L., 2017, 'Distribution and establishment of the alien Australian redclaw crayfish, *Cherax quadricarinatus*, in South Africa and Swaziland', *PeerJ* 5, 1–21. https://doi.org/10.7717/peerj.3135.

Pinder, A., Harman, A., Bird, C., Quinlan, K., Angel, F., Cowan, M. et al., 2019, 'Spread of the non-native redclaw crayfish *Cherax quadricarinatus* (von Martens, 1868) into natural waters of the Pilbara region of Western Australia, with observations on potential adverse ecological effects', *BioInvasions Records* 8, 882–897. https://doi.org/10.3391/bir.2019.8.4.17.

Tavakol, S., Luus-Powell, W.J., Smit, W.J., Baker, C., Hoffman, A. & Halajian, A., 2016, 'First introduction of two Australian Temnocephalan species into Africa with an alien host: double trouble', *Journal of Parasitology* 102, 653–658.

Yeo, D.C.J., 2010, 'Introduced decapod crustaceans in Singapore's reservoirs', *Cosmos* 6, 83-88. https://doi.org/10.1142/S0219607710000498

Zeng, Y., Shakir, K.K. & Yeo, D.C.J., 2019, 'Competition between a native freshwater crab and an invasive crayfish in tropical Southeast Asia', *Biological Invasions* 21, 2653–2663. https://doi.org/10.1007/s10530-019-02009-6.
Zengeya, T.A., Lombard, R.J.H., Nelwamondo, V.E., Nunes, A.L., Measey, J.G. & Weyl, O.L.F., 2022, 'Trophic niche of an invasive generalist consumer: Australian redclaw crayfish, *Cherax quadricarinatus*, in the Inkomati River Basin, South Africa,' *Austral Ecology* 47, 1480-1494 https://doi.org/10.1111/aec.13230.

**4. Management**

**MAN1 What is the feasibility to stop future immigration?**

| Response: Low | Confidence: Medium |
|---|---|

Rationale: The feasibility of preventing unaided immigration of *C. quadricarinatus* into South Africa is low. *Cherax quadricarinatus* is present in neighbouring countries (i.e., Eswatini, Mozambique, and Zimbabwe) and is capable of moving overland through unaided dispersal (Nunes et al. 2017a; Douthwaite et al. 2018; Marufu et al. 2018). Intentional introductions can be prevented through the regulation of imports (Zengeya & Wilson 2020). However, illegal imports can still be problematic (Faulkes 2015; Nunes et al. 2017b). *Cherax quadricarinatus* is present in the pet trade industry in various countries (Faulkes 2015). The significance of *C. quadricarinatus* in the pet trade industry in South Africa is still to be assessed.

References:

Douthwaite, R.J., Jones, E.W., Tyser, A.B. & Vrdoljak, S.M., 2018, 'The introduction, spread and ecology of redclaw crayfish *Cherax quadricarinatus* in the Zambezi catchment', *African Journal of Aquatic Science* 43, 353–366. https://doi.org/10.2989/16085914.2018.1517080.
Faulkes, Z., 2015, 'The global trade in crayfish as pets', *Crustacean Research* 44, 75–92. https://doi.org/10.18353/crustacea.44.0_75.
Marufu, L., Barson, M., Chifamba, P., Tiki, M. & Nhiwatiwa, 2018, 'The population dynamics of a recently introduced crayfish, *Cherax quadricarinatus* (von Martens, 1868), in the Sanyati Basin of Lake Kariba, Zimbabwe', *African Zoology* 53, 17–22. https://doi.org/10.3391/bir.2018.7.2.03.
Nunes, A.L., Zengeya, T.A., Hoffman, A.C., Measey, G.J. & Weyl, O.L.F., 2017a, 'Distribution and establishment of the alien Australian redclaw crayfish, *Cherax quadricarinatus*, in South Africa and Swaziland', *PeerJ* 5**,** 1–21. https://doi.org/10.7717/peerj.3135.
Nunes, A.L., Zengeya, T.A., Measey, G.J. & Weyl, O.L.F., 2017b, 'Freshwater crayfish invasions in South Africa: Past, present and potential future', *African Journal of Aquatic Science* 42, 309–323. https://doi.org/10.2989/16085914.2017.1405788.
Zengeya, T.A. & Wilson, J.R., 2020, *The status of biological invasions and their management in South Africa in 2019*, South African National Biodiversity Institute, Kirstenbosch and DSI-NRF Centre of Excellence for Invasion Biology, Stellenbosch. http://dx.doi.org/10.5281/zenodo.3947613.

**MAN2 Benefits of the Taxon**

**MAN2a Socio-economic benefits of the *Taxon***

| Response: Low | Confidence: Medium |
|---|---|

Rationale: There is no information on the production of *C. quadricarinatus* in South Africa (FAO 2023), but other crayfish species (*C. tenuimanus*) have been farmed on very small scales (Burgess 2007). Therefore, the response has been scored as low.

References:

| | |
|---|---|
| Burgess, M., 2007, 'Pioneers of SA marron production', *farmer's weekly*, viewed 11 August 2023, from https://www.farmersweekly.co.za/animals/aquaculture/pioneers-of-sa-marron-production/.<br>FAO 2023. PARASTACIDAE. Cultured Aquatic Species Information Programme. Text by Jones, C. Fisheries and Aquaculture Division [online]. Rome. Updated 2011-10-10 [Cited Thursday, September 7th 2023]. https://www.fao.org/fishery/en/culturedspecies/cherax_quadricarinatus/en. | |
| **MAN2b Environmental benefits of the *Taxon*** | |
| Response: Low | Confidence: High |
| Rationale: There are no recorded environmental benefits associated with *C. quadricarinatus*. | |
| References: | |

| | |
|---|---|
| **MAN3 Ease of management** | |
| **MAN3a How accessible are populations?** | |
| Response: 2 (difficult to access) | Confidence: High |
| Rationale: *Cherax quadricarinatus* is a small nocturnal aquatic species that is difficult to sample (Larson et al. 2016). In South Africa, *C. quadricarinatus* is widespread in the Inkomati and adjacent river systems (Nunes et al. 2017). Large sections of these river systems flow through private land; hence gaining access to populations will be dependent on the permission of landowners. Furthermore, there are safety risks associated with gaining access to these populations as there are dangerous animals present in these river systems (i.e., crocodiles and hippopotamuses). | |
| References:<br><br>Nunes, A.L., Zengeya, T.A., Hoffman, A.C., Measey, G.J. & Weyl, O.L.F., 2017, 'Distribution and establishment of the alien Australian redclaw crayfish, *Cherax quadricarinatus*, in South Africa and Swaziland', *PeerJ* 5**,** 1–21. https://doi.org/10.7717/peerj.3135.<br>Larson, E.R. & Olden, J.D., 2016, 'Field sampling techniques for crayfish', In: Longshaw, M., Stebbing, P. (eds), *Biology and Ecology of Crayfish 1 st edn*, pp. 287- 323, CRC Press, Boca Raton. | |
| **MAN3b Is detectability critically time-dependent?** | |
| Response: 2 (yes) | Confidence: High |
| Rationale: *Cherax quadricarinatus* can be detected throughout the year; but it is a small nocturnal aquatic species that is difficult to detect during the day (de Moor 2002; Larson et al. 2016; Nunes et al. 2017). | |
| References:<br>de Moor, I., 2002, 'Potential impacts of alien freshwater crayfish in South Africa', African *Journal of Aquatic Science* 27, 125–139. https://doi.org/10.2989/16085914.2002.9626584<br>Nunes, A.L., Zengeya, T.A., Measey, G.J. & Weyl, O.L.F., 2017b, 'Freshwater crayfish invasions in South Africa: Past, present and potential future', *African Journal of Aquatic Science* 42, 309–323. https://doi.org/10.2989/16085914.2017.1405788.<br>Larson, E.R. & Olden, J.D., 2016, 'Field sampling techniques for crayfish', In: Longshaw, M., Stebbing, P. (eds), *Biology and Ecology of Crayfish 1 st edn*, pp. 287- 323, CRC Press, Boca Raton. | |
| **MAN3c Time to reproduction** | |
| Response: 2 (<1 year) | Confidence: High |

| |
|---|
| Rationale: *Cherax quadricarinatus* reaches sexual maturity within the first year and can have multiple spawning events per year when environmental conditions are optimal (de Moor 2002). Clutch size can reach up to 1000 eggs (van Kuijk et al. 2021) however the average is about 500 eggs (de Moor 2002) |
| References:<br>de Moor, I., 2002, 'Potential impacts of alien freshwater crayfish in South Africa', African *Journal of Aquatic Science* 27, 125–139. https://doi.org/10.2989/16085914.2002.9626584<br>van Kuijk, T., Biesmeijer, J.C., van der Hoorn, B.B.& Verdonschot, P.M.F., 2021, 'Functional traits explain crayfish invasive success in the Netherlands', *Scientific Reports* 11, 2772. https://doi.org/10.1038/s41598-021-82302-4 |

| **MAN3d Propagule persistence** | |
|---|---|
| Response: 0 (<1 year) | Confidence: High |
| Rationale: Depending on environmental conditions, *C. quadricarinatus* can have multiple spawning events per year producing up to 1000 eggs (van Kuijk et al. 2021). | |
| References:<br><br>van Kuijk T., Biesmeijer, J.C., van der Hoorn, B.B.& Verdonschot, P.M.F., 2021, 'Functional traits explain crayfish invasive success in the Netherlands', *Scientific Reports* 11, 2772. https://doi.org/10.1038/s41598-021-82302-4. | |

| **MAN3 Ease of management** | |
|---|---|
| Response: Difficult | Confidence: High |
| Rationale: *Cherax quadricarinatus* is difficult to manage given that it is difficult to access, difficult to detect (being aquatic and primarily nocturnal) and has adaptive life history traits that aid establishment (parental care, fast growth rate, high fecundity, and a multiple spawner) (Nunes et al. 2017; van Kuijk et al. 2021). | |
| References:<br><br>Nunes, A.L., Zengeya, T.A., Hoffman, A.C., Measey, G.J. & Weyl, O.L.F., 2017a, 'Distribution and establishment of the alien Australian redclaw crayfish, *Cherax quadricarinatus*, in South Africa and Swaziland', *PeerJ* 5**,** 1–21. https://doi.org/10.7717/peerj.3135.<br>Van Kuijk T., Biesmeijer, J.C., van der Hoorn, B.B.& Verdonschot, P.M.F., 2021, 'Functional traits explain crayfish invasive success in the Netherlands', *Scientific Reports* 11, 2772. https://doi.org/10.1038/s41598-021-82302-4. | |

| **MAN4 Has the feasibility of eradication been evaluated?** | |
|---|---|
| Response: No | Confidence: |
| Rationale: There has been no evaluation of eradication feasibility for *C. quadricarinatus* in South Africa. However, *C. quadricarinatus* is already present in several river systems and so the species should be managed as part of a national programme to prevent future intentional introductions and to minimise natural spread (de Moor 2002; Nunes et al. 2017) rather than attempting eradication. | |
| References:<br>de Moor, I., 2002, 'Potential impacts of alien freshwater crayfish in South Africa', African *Journal of Aquatic Science* 27, 125–139. https://doi.org/10.2989/16085914.2002.9626584<br>Nunes, A.L., Zengeya, T.A., Hoffman, A.C., Measey, G.J. & Weyl, O.L., 2017, 'Distribution and establishment of the alien Australian redclaw crayfish, *Cherax quadricarinatus*, in South Africa and Swaziland', *PeerJ* 5, 1-21. https://doi.org/10.7717/peerj.3135. | |

| **MAN5 Control options and monitoring approaches available for the *Taxon*** |
| --- |
| Response:<br><br>Several methods can be used to monitor and control invasive crayfish populations, and these include mechanical trapping (e.g., Barkhuizen et al. 2022; Madzivanzira et al. 2021), electrofishing (Nunes et al. 2017), and use of biocides, and biological control (Aquiloni et al. 2010; Stebbing et al. 2014). -However, there are several issues that make management of crayfish invasions challenging (Gherardi et al. 2011). For example, the use of biocides is often effective for localised populations in dams and ponds (Peay et al. 2019) but these are non-specific and need to be applied with caution (Gherardi et al. 2011). Mechanical controls are also effective in small discrete water bodies but are expensive and require commitment to follow on to complete eradication (Loureiro et al. 2018). In addition, alien crayfish species are regarded as potential conflict-generating species in South Africa and potential management interventions should consider the concerns of stakeholders to avoid conflict (Zengeya et al. 2017). |
| References:<br><br>Aquiloni, L., Brusconi, S., Cecchinelli, E., Tricarico, E., Mazza, G., Paglianti, A. & Gherardi, F. 2010, 'Biological control of invasive populations of crayfish: the European eel (*Anguilla anguilla*) as a predator of *Procambarus clarkii*', *Biological Invasions* 12 3817-3824. https://doi.org/10.1007/s10530-007-9094-0.<br>Barkhuizen, L.M, Madzivanzira, T.C., South, J., 2022, 'Population ecology of a wild population of red swamp crayfish *Procambarus clarkii* (Girard, 1852) in the Free State Province, South Africa and implications for eradication efforts', *BioInvasions Records* 11, 181–191. https://doi.org/10.3391/bir.2022.<br>Gherardi, F., Aquiloni, L., Diéguez-Uribeondo, J. & Tricarico E., 2011, 'Managing invasive crayfish: is there a hope? *Aquatic Sciences* 73, 185–200. https://doi.org/10.1007/s00027-011-0181-z.<br>Loureiro, T.G, Anastácio, P.M, de Siqueira Bueno, S.L. & Araujo, P.B., 2018, 'Management of invasive populations of the freshwater crayfish *Procambarus clarkii* (Decapoda, Cambaridae): test of a population-control method and proposal of a standard monitoring approach', *Environmental Monitoring Assessment* 190, 559, https://doi.org/10.1007/s10661-018-6942-6.<br>Madzivanzira, T.C., South, J., Nhiwatiwa, T. & Weyl, O.L.F., 2021, 'Standardisation of Australian redclaw crayfish *Cherax quadricarinatus* sampling gear in southern Africa', *Water SA* 47, 380–384. https://hdl.handle.net/10520/ejc-waters-v47-n3-a10.<br>Nunes, A.L., Zengeya, T.A., Hoffman, A.C., Measey, G.J. & Weyl, O.L., 2017, 'Distribution and establishment of the alien Australian redclaw crayfish, *Cherax quadricarinatus*, in South Africa and Swaziland', *PeerJ* 5, 1-21. https://doi.org/10.7717/peerj.3135.<br>Peay, S., Johnsen, S.I, Bean, S.W., Dunn, A.M., Sandodden, R. & Edsman, L, 2019, 'Biocide treatment of invasive signal crayfish: Successes, failures and lessons learned', *Diversity* 11, 29. https://doi.org/10.3390/d11030029<br>Stebbing, P., Longshaw, M., Scott, A., 2014, 'Review of methods for the management of nonindigenous crayfish, with particular reference to Great Britain', *Ethology Ecology & Evolution* 26, 204-231. https://doi.org/10.1080/03949370.2014.908326.<br>Woodford, D.J., Ivey, P., Novoa, A., Shackleton, R., Richardson, D., Weyl, O., Van Wilgen, B. and Zengeya, T., 2017, 'Managing conflict-generating invasive species in South Africa: Challenges and trade-offs', *Bothalia* 47, a2160. https://doi. org/10.4102/abc.v47i2.2160. |

| **MAN6 Any other management considerations to highlight?** | |
| --- | --- |
| Response | No |

**5. Calculations**

**Likelihood = Probable**

| Parameter | Likelihood | Stages | Final assessment |
|---|---|---|---|
| LIK1 | 1 | P(entry) = 1 | P (invasion) = 1 |
| LIK2 | 1 | | |
| LIK3 | 1 | P(establishment) = 1 | |
| LIK4 | 1 | | |
| LIK5 | 1 | P (spread) =1 | |
| LIK6 | 1 | | |

**Consequence = Major (MR)**

| Parameter | Mechanism/sector | Response |
|---|---|---|
| CON1a | Competition | DD |
| CON1b | Predation | DD |
| CON1c | Hybridisation | DD |
| CON1d | Disease transmission | MR |
| CON1e | Parasitism | DD |
| CON1f | Poisoning/toxicity | MC |
| CON1g | Bio-fouling or other direct physical disturbance | DD |
| CON1h | Grazing/herbivory/browsing | DD |
| CON1i | Chemical, physical, structural impact | MC |
| CON1k | Indirect impacts through interactions with other species | DD |
| CON1 | Maximum environmental impact | MR |
| CON2a | Safety | DD |
| CON2b | Material and immaterial assets | MO |
| CON2c | Health | DD |
| CON2d | Social, spiritual and cultural relations | DD |
| CON2 | Maximum socio-economic impact | MO |
| CON3 | Environmental impact of closely related taxa | Not scored as CON1 scored |
| CON4 | Socio-economic impact of closely related taxa | Not scored as CON2 scored |

| | | |
|---|---|---|
| CON5 | Potential impact based on traits, experiments, or models | MR |

**Risk score = High**

| | | Consequences | | | | |
|---|---|---|---|---|---|---|
| | | MC | MN | MO | MR | MV |
| Likelihood | Extremely unlikely | low | low | low | medium | medium |
| | Very unlikely | low | low | low | medium | high |
| | Unlikely | low | low | medium | high | high |
| | Fairly probable | medium | medium | high | high | high |
| | Probable | medium | high | high | high | high |

**Ease of management = Difficult**

| Parameter | Question | Response |
|---|---|---|
| MAN3a | How accessible are populations? | 2 |
| MAN3b | Is detectability critically time-dependent? | 2 |
| MAN3c | Time to reproduction | 2 |
| MAN3d | Propagule persistence | 0 |
| **MAN3** | **SUM** | 6 |

**Appendix BAC7**: The native distribution of *Cherax quadricarinatus*. Source: Bláha, M., Patoka, J., Kozák, P., & Kouba, A. (2016). Unrecognized diversity in New Guinean crayfish species (Decapoda, Parastacidae): The evidence from molecular data. Integrative Zoology, 11(6), 457-468. https://doi.org/10.1111/1749-4877.12211.

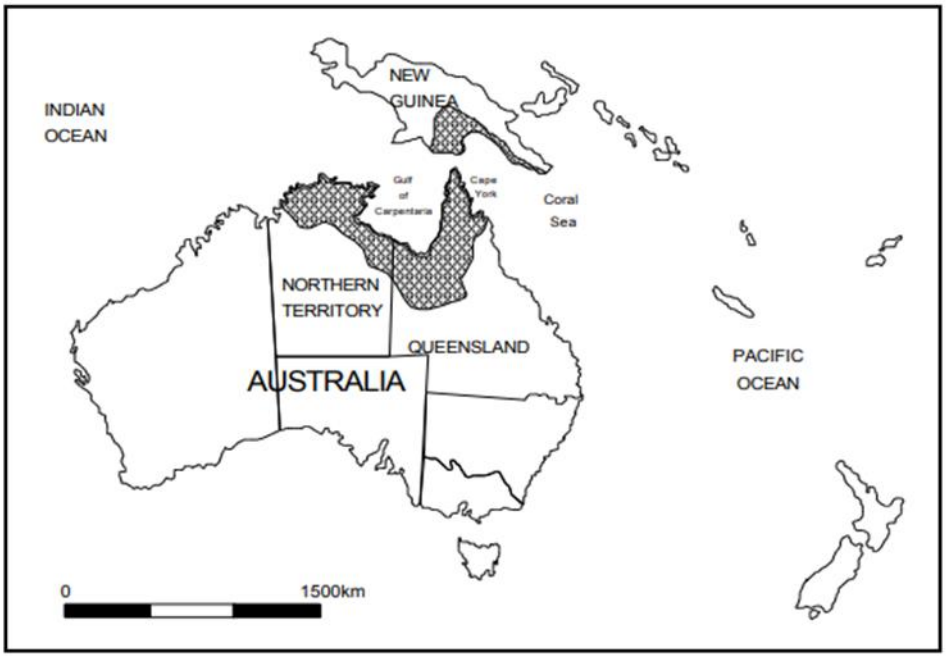

**Appendix BAC8(a)**: The global alien distribution of *Cherax quadricarinatus*. Source: Sallehuddin, A.S., Kamarudin, A.S & Ismail, N., 2021, 'Review on the global distribution of wild population of Australian Redclaw Crayfish, *Cherax quadricarinatus* (von Martens, 1868)', *Bioscience Research* 18, 194–207.

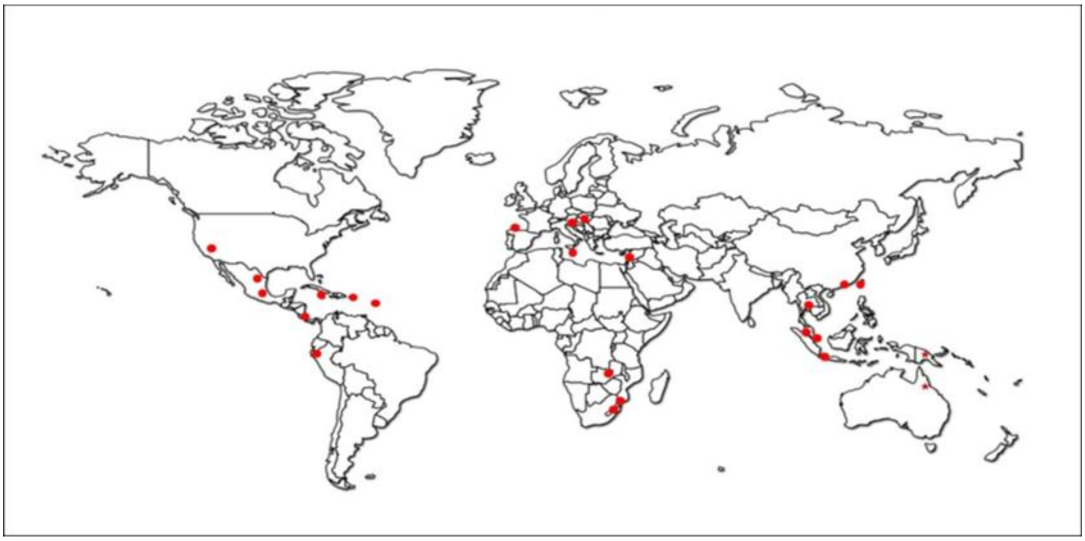

**Appendix BAC8(b)**: Distribution of *Cherax quadricarinatus* in South Africa. Locality records sourced from GBIF.org (11 September 2023) GBIF Occurrence Download https://doi.org/10.15468/dl.gaa77h.

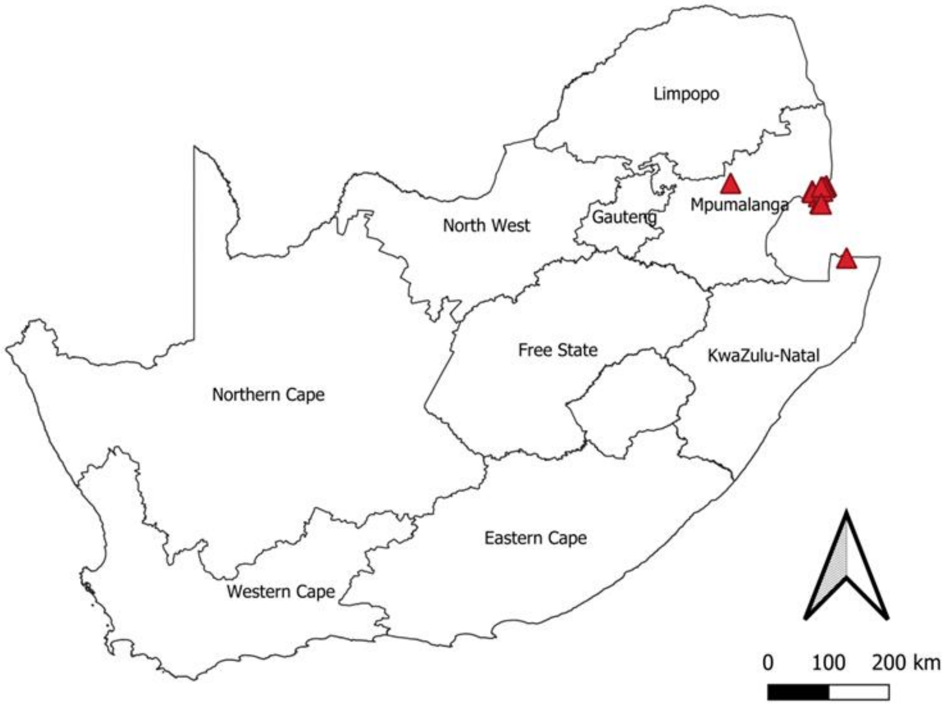

#### 1.5 Risk analysis report for Calico crayfish (*Faxonius immunis*)

<table>
<tr><td colspan="2">Taxon: Faxonius immunis (Hagen, 1870)</td><td>Area: South Africa</td></tr>
<tr><td colspan="2">Picture of Taxon<br>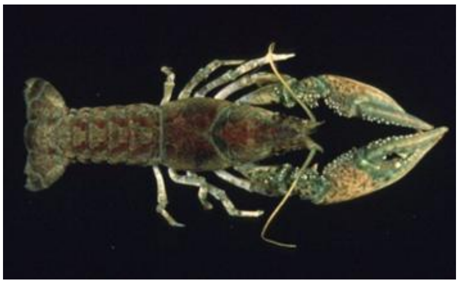<br>https://alchetron.com/Orconectes-immunis</td><td>Alien distribution map<br>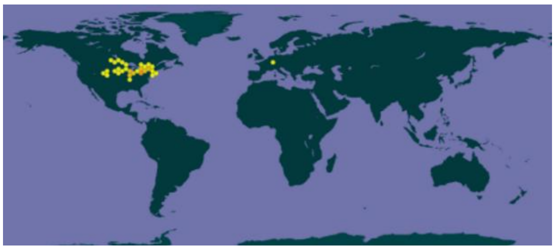<br>https://www.gbif.org/species/2227004</td></tr>
<tr><td colspan="2">Risk Assessment summary:<br>Calico crayfish (Faxonius immunis) is native to eastern parts of North America but has been introduced to western parts North America and to Europe. There are no know populations of F. immunis in South Africa but there is anecdotal evidence that it is present in the ornamental trade and this needs to be verified. There are no known populations in neighbouring countries, therefore via unaided pathways is unlikely. The illegal pet trade industry remains a concern because F. immunis may be moved around by humans and there is a high risk of release of unwanted pets. Faxinous immunis is omnivorous species, it is likely to cause a decrease in macrophyte cover, macroinvertebrates abundance and diversity, changing trophic interaction and community composition. It is also regarded as an agricultural pest due to its grazing and burrowing activities.</td><td>Risk score:<br>High</td></tr>
<tr><td colspan="2">Management options summary:<br>Faxonius immunis is a small, largely nocturnal, burrowing freshwater species that is difficult to detect. Several methods can be used to control F. immunis such as mechanical removal using traps and electrofishing, biocides, and biological control. However, there are several issues that make the use of these methods challenging. The benefits of F. immunis were scored as low.</td><td>Ease of management:<br>Medium</td></tr>
<tr><td colspan="2" rowspan="2">Recommendations:<br><br>Faxonius immunis poses a high risk to South Africa. If F. immunis were to establish outside of captivity, such populations would likely be very difficult to extirpate.<br><br>Faxonius immunis is recommended to be added to a future prohibited species list, active efforts made to confirm that the species is absent in the country, and that any individuals found are controlled. This would likely require a dedicated assessment of the ornamental trade. A reassessment in five years is thus required and, if present in the country, F. immunis should be listed under category 1a.</td><td>Listing under NEM:BA A&IS lists of 2020:<br><br>Not listed</td></tr>
<tr><td>Recommended listing category:<br><br>Prohibited list[1]</td></tr>
</table>

[1] The absence of *F. immunis* in South Africa should be confirmed and the taxon should be reassessed in five years (i.e., by 2029).

#### 1. Background

| **BAC1 Name of assessor(s)** | |
|---|---|
| Name of lead assessor | Lee-Anne Botha |
| Additional assessor (1) | Tsungai Zengeya |
| Additional assessor (2) | Christian Chimimba |
| **BAC2 Contact details of assessor (s)** | |
| Lead assessor | Organisational affiliation: Department of Zoology and Entomology, University of Pretoria and The South African National Biodiversity Institute (SANBI). |
| | |
| Additional assessor (1) | Organisational affiliation: Department of Zoology and Entomology, University of Pretoria and The South African National Biodiversity Institute (SANBI). |
| | |
| Additional assessor (2) | Organisational affiliation: Department of Zoology and Entomology, University of Pretoria and The South African National Biodiversity Institute (SANBI). |
| | |
| **BAC3 Name(s) and contact details of expert(s) consulted** | |
| Expert (1) | Name: |
| | email: |
| | Phone: |
| Expert (2) | Name: |
| | email: |
| | Phone: |
| Comments: none | |
| **BAC4 Scientific name of *Taxon* under assessment** | |
| *Taxon* name: *Faxonius immunis* | Authority: (Hagen, 1870) |
| Comments: The group of surface-dwelling crayfish in the *Orconectes* genus was moved to *Faxonius* (Crandall & De Grave 2017). The taxonomic status of *Faxonius immunis* has been verified (GBIF 2024). | |
| References:<br>Crandall, K.A. & De Grave, S., 2017, 'An updated classification of the freshwater crayfishes (Decapoda: Astacidea) of the world, with a complete species list', *Journal of Crustacean Biology*, 27, 615–653. https://doi.org/10.1093/jcbiol/rux070<br>*Faxonius immunis* (Hagen, 1870) in GBIF Secretariat (2023). GBIF Backbone Taxonomy. Checklist dataset https://doi.org/10.15468/39omei accessed via GBIF.org on 2024-06-05. | |
| **BAC5 Synonym(s) considered** | |
| Synonyms: *Cambarus immunis* Hagen, 1870 | |
| Comments: Cave inhabiting crayfish and the surface-dwelling crayfish were split into two groups (Crandall & De Grave 2017). The non-cave dwelling crayfish in the *Orconectes* genus was moved to *Faxonius* (Crandall & De Grave 2017). Therefore, the abovementioned synonym was used in the literature search of this risk analysis. | |
| References:<br>Crandall, K.A. & De Grave, S., 2017, 'An updated classification of the freshwater crayfishes (Decapoda: Astacidea) of the world, with a complete species list', *Journal of Crustacean Biology*, 27, 615–653. https://doi.org/10.1093/jcbiol/rux070 | |
| **BAC6 Common name(s) considered** | |
| Common names: Calico crayfish | |

| | |
|---|---|
| Comments: Calico crayfish is listed as the common name for *F. immunis* on the GBIF and the IUCN Red List (Adams et al. 2010; GBIF 2024). This was the only common name used for literature search in this risk analysis. | |
| References:<br>Adams, S., Schuster, G.A. & Taylor, C.A., 2010, *Orconectes immunis. The IUCN Red List of ThreatenedSpecies2010*:e.T153925A4564415. http://www.iucnredlist.org/details/153925/0. Accessed on 01/12/2018<br>*Faxonius immunis* (Hagen, 1870) in GBIF Secretariat (2023). GBIF Backbone Taxonomy. Checklist dataset https://doi.org/10.15468/39omei accessed via GBIF.org on 2024-06-05. | |
| **BAC7 What is the native range of the *Taxon*?** (add map in Appendix BAC7) | |
| Response: Canada \| United States | Confidence: High |
| Comments: *Faxonius immunis* is native to eastern North America. Its range extends as far east as Maine and Connecticut, and as far west as eastern Colorado and Wyoming. Its northernmost range includes southern Manitoba, Ontario and Quebec; Its southernmost range extends to Kentucky (Capelli and Magnuson 1983; Adams et al. 2010). | |
| References:<br>Adams, S., G.A., Schuster, & Taylor, C.A., 2010, *Orconectes immunis*. The IUCN Red List of Threatened Species. http://www.iucnredlist.org/details/153925/0. Accessed on 01/12/2018.<br>Capelli, G.M. & Magnuson, J.J., 1983, 'Morphoedaphic and biogeographic analysis of crayfish distribution in northern Wisconsin', *Journal of Crustacean Biology*, 3, 548–564. https://doi.org/10.1163/193724083X00210. | |
| **BAC8 What is the global alien range of the *Taxon*?** (add map in Appendix BAC8) | |
| Response: western North America \| Germany \| France | Confidence: High |
| Comments: *Faxonius immunis* is native to eastern North America but has being introduced and has become established in western North America (Newkirk et al. 2023) and Europe (Holdrich 2009; Chucholl 2012; Chucholl and Chucholl 2021; Herrmann et al. 2022) | |
| References:<br>Chucholl, C., 2012, 'Understanding invasion success: life-history traits and feeding habits of the alien crayfish *Orconectes immunis* (Decapoda, Astacida, Cambaridae)', *Knowledge and Management of Aquatic Ecosystems*, 404: 04. https://doi.org/10.1051/kmae/2011082.<br>Chucholl, F. & Chucholl, C., 2021, 'Differences in the functional response of four invasive and one native crayfish species suggest invader-specific ecological impacts', *Freshwater Biology*, 66, 2051–2063. https://doi.org/10.1111/fwb.13813<br>Herrmann, A., Grabow, K. & Martens, A., 2022, 'The invasive *Faxonius immunis* causes the collapse of macroinvertebrate communities in Central European ponds', *Aquatic Ecology*, 56, 741–750. https://doi.org/10.1007/s10452-021-09935-5.<br>Holdich, D.M., Reynolds, J.D., & Sibley, P.J., 2009, 'A review of the ever increasing threat to European crayfish from non-indigenous crayfish species', *Knowledge and Management of Aquatic Ecosystems* 11, 394–395. https://doi.org/10.1051/kmae/2009025 .<br>Newkirk, B.M., Larson, E.R., Walker, A.D. & Walters, A.D., 2023, 'Winners and losers over a ½ century of change in crayfish assemblages of Wyoming, USA', *Freshwater Science,* 42, 146–160. | |
| **BAC9 Geographic scope = the *Area* under consideration** | |
| *Area* of assessment: South Africa | |
| Comments: The geographic scope of this assessment is limited to South Africa. | |
| **BAC10 Is the *Taxon* present in the *Area*?** | |
| Response: No | Confidence: Low |
| Comments: There are no records of *Faxonius immunis* being in the country (Nunes et al. 2017). *Faxonius immunis* might be present through the pet trade, although this needs to be assessed (Chucholl 2012; Faulkes 2015). | |

| |
|---|
| References:<br>Chucholl, C., 2013, ‘Invaders for sale: trade and determinants of introduction of ornamental freshwater crayfish’, *Biological Invasions* 15, 125–141. https://doi.org/10.1007/s10530-012-0273-2.<br>Faulkes, Z., 2015, ‘The global trade in crayfish as pets’, *Crustacean Research* 44, 75–92. https://doi.org/10.18353/crustacea.44.0_75<br>Nunes, A.L., Zengeya, T.A., Measey, G.J. & Weyl, O.L.F., 2017b, ‘Freshwater crayfish invasions in South Africa: past, present and potential future’, *African Journal of Aquatic Science* 42, 309–323. https://doi.org/10.2989/16085914.2017.1405788 |

| **BAC11 Availability of physical specimen** | |
|---|---|
| Response: No | Confidence in ID: |
| Herbarium or museum accession number: no physical specimen is lodged as present in South Africa (GBIF 2024) | |
| References:<br>*Faxonius immunis* (Hagen, 1870) in GBIF Secretariat (2023). GBIF Backbone Taxonomy. Checklist dataset https://doi.org/10.15468/39omei accessed via GBIF.org on 2024-06-05. | |

| **BAC12 Is the *Taxon* native to the *Area* or part of the *Area*?** | | |
|---|---|---|
| The *Taxon* is native to (part of) the *Area*. | No | Confidence: High |
| The *Taxon* is alien in (part of) the *Area*. | Yes | Confidence: High |
| Comments: South Africa has no indigenous freshwater crayfish (de Moor 2002, Nunes et al. 2017). | | |
| References:<br>de Moor, I., 2002, ‘Potential impacts of alien freshwater crayfish in South Africa’, *African Journal of Aquatic Science* 27, 125–139. https://doi.org/10.2989/26085914.2002.9626584.<br>Nunes, A.L., Zengeya, T.A., Measey, G.J. & Weyl, O.L.F., 2017b, ‘Freshwater crayfish invasions in South Africa: past, present and potential future’, *African Journal of Aquatic Science* 42, 309–323. https://doi.org/10.2989/16085914.2017.1405788. | | |

| **BAC13 What is the *Taxon*’s introduction status in the *Area*?** | | |
|---|---|---|
| The *Taxon* is in cultivation/containment. | Don’t know | Confidence: Low |
| The *Taxon* is present in the wild. | None | Confidence: Low |
| The *Taxon* has established/naturalised. | None | Confidence: Low |
| The *Taxon* is invasive. | None | Confidence: Low |
| Comments: There are no records of *F. immunis* being present in South Africa, and, there is furthermore no evidence available that indicates it present and been traded in the pet trade industry (de Moor 2002; Nunes et al. 2017). | | |
| References:<br>de Moor, I., 2002, ‘Potential impacts of alien freshwater crayfish in South Africa’, *African Journal of Aquatic Science*, 27, 125–139. https://doi.org/10.2989/26085914.2002.9626584.<br>Nunes, A.L., Zengeya, T.A., Measey, G.J. & Weyl, O.L.F., 2017, ‘Freshwater crayfish invasions in South Africa: past, present and potential future’, African *Journal of Aquatic Science* 42, 309–323. https://doi.org/10.2989/16085914.2017.1405788 | | |

| **BAC14 Primary (introduction) pathways** | | |
|---|---|---|
| Release | None | Confidence: |
| Escape | Aquarium trade \| Live bait | Confidence: Low |
| Contaminant | None | Confidence: |
| Stowaway | None | Confidence: |
| Corridor | None | Confidence: |
| Unaided | None | Confidence: |
| Comments: Introduced as bait in areas outside its native range in North America and the introduction pathway is unclear outside the USA but likely aquarium trade and or fishing bait (Dehus et al. 1999; Chocholl 2013, Schrimpf et al. 2013). | | |

| |
|---|
| References:<br>Chucholl, C., 2013, 'Invaders for sale: trade and determinants of introduction of ornamental freshwater crayfish', *Biological Invasions* 15, 125–141. https://doi.org/10.1007/s10530-012-0273-2.<br>Dehus, P., Dussling, U. & Hoffmann, C., 1999, 'Notes on the occurrence of the calico crayfish (*Orconectes immunis*) in Germany', *Freshwater Crayfish*, 12, 786–790.<br>Schrimpf, A., Chucholl, C., Schmidt, T., & Schulz, R., 2013, 'Crayfish plague agent detected in populations of the invasive North American crayfish *Orconectes immunis* (Hagen, 1870) in the Rhine River, Germany', *Aquatic Invasions*, 8(1). http://dx.doi.org/10.3391/ai.2013.8.1.12 |

**2. Likelihood**

| **LIK1 Likelihood of entry via unaided primary pathways** | |
|---|---|
| Response: Very unlikely (p = 0.0027) | Confidence: Low |
| Rationale: There is no evidence that *F. immunis* is present in South Africa or has established in the wild in neighbouring countries (Nunes et al. 2017; Madzivanzira et al. 2020) therefore the probability of *F. immunis* entering South Africa through unaided pathways (connected waterways) is very unlikely (p = 0.0027). | |
| References:<br>Madzivanzira, T.C., South, J., Wood, L.E., Nunes, A.L. & Weyl, O.L.F., 2020, 'A review of freshwater crayfish introductions in Africa', *Reviews in Fisheries Science & Aquaculture* 1-21. https://doi.org/10.1080/23308249.2020.1802405<br>Nunes, A.L., Zengeya, T.A., Measey, G.J. & Weyl, O.L.F., 2017b, 'Freshwater crayfish invasions in South Africa: past, present and potential future', *African Journal of Aquatic Science* 42, 309–323. https://doi.org/10.2989/16085914.2017.1405788. | |

| **LIK2 Likelihood of entry via human aided primary pathways** | |
|---|---|
| Response: Unlikely (p = 0.027) | Confidence: Low |
| Rationale: *Faxonius immunis* is present in the global pet trade, however it is considered rare and has only been found in the Germany and the United States of America (Chucholl 2013; Faulkes 2015). There is no evidence to show that *F. immunis* is currently traded in the South Africa pet trade industry, but this needs to be assessed (Nunes et al. 2017). | |
| References:<br>Chucholl, C., 2013, 'Invaders for sale: trade and determinants of introduction of ornamental freshwater crayfish', *Biological Invasions* 15, 125–141. https://doi.org/10.1007/s10530-012-0273-2<br>Faulkes, Z. 2015, 'The global trade in crayfish as pets', *Crustacean Research* 44, 75–92. https://doi.org/10.18353/crustacea.44.0_75<br>Nunes, A.L., Zengeya, T.A., Measey, G.J. & Weyl, O.L.F., 2017b, 'Freshwater crayfish invasions in South Africa: past, present and potential future', *African Journal of Aquatic Science* 42, 309–323. https://doi.org/10.2989/16085914.2017.1405788. | |

| **LIK3 Habitat suitability** | |
|---|---|
| Response: Fairly probable (p = 0.5) | Confidence: Low |
| Rationale: *Faxonius immunis* is a generalist species can occupy a variety of freshwater habitats such as ponds, floodplains, drainage ditches and small sluggish streams (Adams et al 2010). It can also occur in higher gradient gravel-bedded streams (Chucholl 2009; Soes & Koese 2010). The substrate in the habitats of this species is generally soft mud or clay with abundant aquatic vegetation. *Faxonius immunis* can travel across dry ground, especially in wet weather enabling the species to colonise new areas (Chucholl 2009; Soes & Koese 2010). Despite the broad ecological niche of *Faxonius immunis*, it is unable to colonise fast flowing streams, restricting its distribution (Chucholl 2009; Soes & Koese 2010). *Faxonius immunis* are known to | |

| |
|---|
| dig deep burrows, which allow it to occupy shallow and temporary water bodies (Chucholl 2009; Soes & Koese 2010). The habitats are present in most aquatic systems across South Africa. |
| References:<br><br>Adams, S., Schuster, G.A., Taylor, C.A., 2010, *Orconectes immunis. The IUCN Red List of Threatened Species 2010*: e.T153925A4564415.<br>Chucholl, C., 2009. 'The 'newcomer' *Orconectes immunis* keeps spreading in the upper Rhine plain', *Crayfish News: IAA Newsletter*, 31: 4–5.<br>Soes, M. & Koese, B., 2010, Invasive freshwater crayfish in the Netherlands: a preliminary risk analysis. Invasive freshwater crayfish in the Netherlands: a preliminary risk analysis |

| **LIK4 Climate suitability** | |
|---|---|
| Response: Fairly probable (p = 0.5) | Confidence: Low |
| Rationale: *Faxonius immunis* is native to North America, ranging from southern Quebec and New England westward across the upper Midwest to Wyoming and eastern Colorado and south to extreme northwestern Tennessee (Hobbs 1989; Hosabettu & Daniel 2020). Its native range covers different climatic zones (following the classification of Köppen: oceanic; warm summer/humid continental; hot summer/humid continental; cold semi-arid; warm summer/mediterranean continental) (EU 2022). A large portion (>20%) of South Africa is climatical suitable for the establishment of *F. immunis* | |
| References:<br>Global Invasive Species Database (GISD) 2015. Species profile *Orconectes rusticus*.Available from: http://www.iucngisd.org/gisd/species.php?sc=217 [Accessed 03 July2018]<br>European Commission 2022, Directorate-General for Environment, Study on invasive alien species – Development of risk assessments to tackle priority species and enhance prevention – Final report (and annexes), Publications Office of the European Union, 2022, https://data.europa.eu/doi/10.2779/302048<br>Hobbs, H.H. Jr., 1989, 'An illustrated checklist of the American crayfishes (Decapoda: Astacidae, Cambaridae, and Parastacidae)', *Smithsonian Contributions to Zoology*, 480, 1–236.<br>Hosabettu, M., Daniel, W.M., 2020, *Faxonius immunis* (Hagen, 1870): U.S. Geological Survey, Nonindigenous Aquatic Species Database, Gainesville, FL,, Revision Date: 4/1/2020, Access Date: 5/29/2020 | |

| **LIK5 Unaided secondary (dispersal) pathways** | |
|---|---|
| Response: Unlikely (p = 0.027) | Confidence: Low |
| Rationale: *Faxinous immunis* can disperse naturally in aquatic environments and is known to disperse overland (Chucholl 2012; Herrmann et al. 2018). There are no wild populations in South Africa or neighbouring countries, thus unaided secondary dispersal through connected waterways is very unlikely (Madzivanzira et al. 2020). | |
| References:<br><br>Chucholl, C., 2012, 'Understanding invasion success: life-history traits and feeding habits of the alien crayfish *Orconectes immunis* (Decapoda, Astacida, Cambaridae)', *Knowledge and Management of Aquatic Ecosystems*, 404: 04. https://doi.org/10.1051/kmae/2011082.<br>Herrmann, A., Schnabler, A. & Martens, A., 2018, 'Phenology of overland dispersal in the invasive crayfish *Faxonius immunis* (Hagen) at the Upper Rhine River area', *Knowledge & Management of Aquatic Ecosystems* 419: 30. https://doi.org/10.1007/s10452-021-09935-5<br>Madzivanzira, T.C., South, J., Wood, L.E., Nunes, A.L. & Weyl, O.L.F., 2020, 'A review of freshwater crayfish introductions in Africa', *Reviews in Fisheries Science & Aquaculture* 1–21. https://doi.org/10.1080/23308249.2020.1802405 | |

| **LIK6 Human aided secondary (dispersal) pathways** | |
|---|---|
| Response: Fairly probable (p = 0.5) | Confidence: Low |
| Rationale: Intentional release into the wild through the pet/ aquarium species and live bait pathways remains a concern. In areas of introduction, *Faxonius immunis* has been released into the wild by humans as unwanted pets and bucket release by anglers into waterways where they are used as bait, dispersing into new areas (Dehus et al. 1999; Chucholl 2013, Schrimpf et al. 2013). | |
| References:<br><br>Chucholl, C., 2013, 'Invaders for sale: trade and determinants of introduction of ornamental freshwater crayfish', *Biological Invasions* 15, 125–141. https://doi.org/10.1007/s10530-012-0273-2.<br>Dehus, P., Dussling, U. & Hoffmann, C., 1999, 'Notes on the occurrence of the calico crayfish (*Orconectes immunis*) in Germany', *Freshwater Crayfish*, 12, 786–790.<br>Schrimpf, A., Chucholl, C., Schmidt, T., & Schulz, R., 2013, 'Crayfish plague agent detected in populations of the invasive North American crayfish *Orconectes immunis* (Hagen, 1870) in the Rhine River, Germany', *Aquatic Invasions*, *8*(1). https://dx.doi.org/10.3391/ai.2013.8.1.12. | |

**3. Consequences**

| **CON1 Environmental impact** | |
|---|---|
| **CON1a: Competition** | |
| Response: Moderate (MO) | Confidence: Low |
| Rationale: There is limited direct evidence for negative environmental impacts for *F. immunius* in its global invasive range and most of the cited impacts are based on potential rather than observed impacts (e.g., Jansen et al. 2009; EU 2022). For example, *Faxinous immunis* is an omnivore and in its alien range in the USA it is considered as been more aggressive than native congeneric crayfish species such *F. limosus* which it is likely to displace through coemption for resources (food and shelter) (Jansen et al. 2009) | |
| References:<br><br>European Commission 2022, Directorate-General for Environment, Study on invasive alien species – Development of risk assessments to tackle priority species and enhance prevention – Final report (and annexes), Publications Office of the European Union, 2022, https://data.europa.eu/doi/10.2779/302048<br>Jansen, W., Geard, N., Mosindy, T., Olson, G. & Turner, M., 2009, 'Relative abundance and habitat association of three crayfish (*Orconectes virilis*, *O. rusticus*, and *O. immunis*) near an invasion front of *O. rusticus*, and long-term changes in their distribution in Lake of the Woods, Canada', *Aquatic Invasions*, 4, 627-649. https://doi.org/10.3391/ai.2009.4.4.9. | |
| **CON1b: Predation** | |
| Response: Major (MR) | Confidence: Medium |
| Rationale: *Faxinous immunis* is a facultative omnivore that consumes detritus, macrophytes and invertebrates (Chucholl 2012). In areas of introduction, direct predation by *F. immunis* on native fauna has resulted in a decline in population size of at least one native fish species. For example, in Germany, predation by *F. immunius* has been implicated in the decline of native populations of invertebrates such as dragonflies, Coleoptera, Odonata, Mollusca and Trichoptera ((EU 2022). | |
| References: | |

European Commission 2022, Directorate-General for Environment, Study on invasive alien species – Development of risk assessments to tackle priority species and enhance prevention – Final report (and annexes), Publications Office of the European Union, 2022, https://data.europa.eu/doi/10.2779/302048

Chucholl, C., 2012, 'Understanding invasion success: life-history traits and feeding habits of the alien crayfish *Orconectes immunis* (Decapoda, Astacida, Cambaridae)', *Knowledge and Management of Aquatic Ecosystems*, 404: 04. https://doi.org/10.1007/s10530-012-0273-2.

**CON1c: Hybridisation**

| Response: Data Deficient (DD) | Confidence: High |
|---|---|

Rationale: There are no native freshwater crayfish in South Africa (de Moor 2002; Crandall & de Grave 2017), so there is no possibility of hybridisation with native species.

References:

de Moor, I., 2002, 'Potential impacts of alien freshwater crayfish in South Africa', African *Journal of Aquatic Science* 27, 125–139. https://doi.org/10.2989/16085914.2002.9626584.

Crandall, K.A. & De Grave, S., 2017, 'An updated classification of the freshwater crayfishes (Decapoda: Astacidea) of the world, with a complete species list', *Journal of Crustacean Biology*, 37(5), 615–653. https://doi.org/10.1093/jcbiol/rux070.

**CON1d: Transmission of disease**

| Response: Minor (MN) | Confidence: Low |
|---|---|

Rationale: *Faxinous immunis* is a known vector of pathogen such as the crayfish plague agent, *Aphanomyces astaci* (Schrimpf et al. 2013)

References:

Schrimpf, A., Chucholl, C., Schmidt, T., & Schulz, R., 2013, 'Crayfish plague agent detected in populations of the invasive North American crayfish *Orconectes immunis* (Hagen, 1870) in the Rhine River, Germany', *Aquatic Invasions*, 8(1). https://dx.doi.org/10.3391/ai.2013.8.1.12.

**CON1e: Parasitism**

| Response: Data Deficient (DD) | Confidence: High |
|---|---|

Rationale: *Faxinous immunis* not a parasitic taxon (Lodge et al. 2012)

References:

Lodge, D.M., Deines, A., Gherardi, F., Yeo, D.C., Arcella, T., Baldridge, A.K., et al., 2012, 'Global introductions of crayfishes: evaluating the impact of species invasions on ecosystem services', *Annual Review of Ecology, Evolution, and Systematics* 43**,** 449–472. https://doi.org/10.1146/annurev-ecolsys-111511-103919

**CON1f: Poisoning/toxicity**

| Response: Data Deficient (DD) | Confidence: |
|---|---|

Rationale: No information is available to assess the level of impact.

References:

| **CON1g: Bio-fouling or other direct physical disturbance** | |
|---|---|
| Response: Minor (MN) | Confidence: Low |
| Rationale: *Faxinious immunis* is a burrowing species and is likely to cause damages to riverbanks and impacts water quality through increased turbidity as observed from other burrowing alien crayfish species in Europe (Souty-Grosset et al. 2016) | |
| References:<br><br>Souty-Grosset, C., Anastácio, P., Aquiloni, L., Banha, F., Choquer, J., Chucholl, C. & Tricarico, E., 2016, 'Impacts of the red swamp crayfish *Procambarus clarkii* on European aquatic ecosystems and human well-being', *Limnologica*, 58: 78-93. https://doi.org/10.1016/j.limno.2016.03.003. | |
| **CON1h: Grazing/herbivory/browsing** | |
| Response: Minor (MN) | Confidence: Low |
| Rationale: *Faxonius immunis* is an omnivorous species, it can cause a decrease in macrophyte cover, macroinvertebrates abundance and diversity, changing trophic interaction and community composition (EU 2022) | |
| References:<br><br>European Commission 2022, Directorate-General for Environment, Study on invasive alien species – Development of risk assessments to tackle priority species and enhance prevention – Final report (and annexes), Publications Office of the European Union, 2022, https://data.europa.eu/doi/10.2779/302048 | |
| **CON1i: Chemical, physical or structural impact on ecosystem** | |
| Response: Data Deficient (DD) | Confidence: |
| Rationale: No information is available to assess the level of impact | |
| References: | |
| **CON1k: Indirect impacts through interactions with other species** | |
| Response: Data Deficient (DD) | Confidence: |
| Rationale: No information is available to assess the level of impact | |
| References: | |
| **CON1 Maximum environmental impact** | |
| Response: Major (MR) | Confidence: Medium |
| Rationale: *Faxinous immunis is* omnivorous species, it is likely to cause a decrease in macrophyte cover, macroinvertebrates abundance and diversity, changing trophic interaction and community composition (EU 2022) | |
| References: | |

European Commission 2022, Directorate-General for Environment, Study on invasive alien species – Development of risk assessments to tackle priority species and enhance prevention – Final report (and annexes), Publications Office of the European Union, 2022. https://data.europa.eu/doi/10.2779/302048.

| CON2 Socio-economic impact | |
|---|---|
| **CON2a: Safety** | |
| Response: Data Deficient (DD) | Confidence: |
| Rationale: No information is available to assess the level of impact | |
| References: | |
| **CON2b: Material and immaterial assets** | |
| Response: Minor (MN) | Confidence: Medium |
| Rationale: Regarded as agricultural pest due to its burrowing and grazing activities (Souty-Grosset et al. 2006) | |
| References:<br><br>Souty Grosset, C., Holdich, D.M., Noel, P.Y., Reynolds, J.D. & Haffner, P., (eds), 2006, 'Atlas of crayfish in Europe', Museum national d'Histoire naturelle, Paris, 187p (Patrimoines naturels 64). | |
| **CON2c: Health** | |
| Response: Data Deficient (DD) | Confidence: |
| Rationale: | |
| References: | |
| **CON2d: Social, spiritual and cultural relations** | |
| Response: Data Deficient (DD) | Confidence: |
| Rationale: | |
| References: | |
| **CON2 Maximum socio-economic impact** | |
| Response: Minor (MN) | Confidence: |
| Rationale: *Faxonius immunis* is regarded as agricultural pest due to its grazing and burrowing activities (Chucholl 2012; EU 2022). | |
| References:<br><br>European Commission 2022, Directorate-General for Environment, Study on invasive alien species – Development of risk assessments to tackle priority species and enhance prevention – Final report (and annexes), Publications Office of the European Union, 2022, https://data.europa.eu/doi/10.2779/302048 | |

| |
|---|
| Chucholl, C., 2012, 'Understanding invasion success: life-history traits and feeding habits of the alien crayfish *Orconectes immunis* (Decapoda, Astacida, Cambaridae)', *Knowledge and Management of Aquatic Ecosystems*, 404: 04. https://doi.org/10.1051/kmae/2011082. |

| **CON3 Closely related species' environmental impact** | |
|---|---|
| Response: Not scored as CON1 scored | Confidence: |
| Rationale: There is information to score CON1, so CON3 does not need to be scored as per the framework. | |
| References: | |

| **CON4 Closely related species' socio-economic impact** | |
|---|---|
| Response: Not scored as CON1 scored | Confidence: |
| Rationale: There is information to score CON1, so CON3 does not need to be scored as per the framework. | |
| References: | |

| **CON5 Potential impact** | |
|---|---|
| Response: MR | Confidence: Medium |
| Rationale: *Faxonius immunis* is a functional omnivore and may also have detrimental impacts on macroinvertebrates and macrophyte communities in areas where invaded (Klocker and Strayer 2004; Roth et al. 2006; Holdich et al. 2009; Kreps et al. 2012; EU 2022). *Faxonius immunis* can also influence the occurrence and species composition of fish communities (Keller et al. 2008). In addition, being a vector for the crayfish plague is another cause of concern because this disease may be transferable and could be detrimental for native freshwater crustaceans in South Africa (Lodge et al. 2012). *Faxonius immunis* also digs deep burrows, which allow it to occupy shallow and temporary water bodies. Burrowing activities by invasive crayfish species may destabilise riverbanks, causing erosion (Chucholl 2009; Soes & Koese 2010). | |
| References:<br>Chucholl, C., 2009, 'The 'newcomer' *Orconectes immunis* keeps spreading in the upper Rhine plain', *Crayfish News*, *IAA Newsletter*, 31, 4–5.<br>European Commission 2022, Directorate-General for Environment, Study on invasive alien species – Development of risk assessments to tackle priority species and enhance prevention – Final report (and annexes), Publications Office of the European Union, 2022, https://data.europa.eu/doi/10.2779/302048<br>Keller, R.P., Frang, K. & Lodge, D.M., 2008, 'Preventing the spread of invasive species: Economic benefits of intervention guided by ecological predictions', *Conservation Biology*, 22, 80–88. https://doi.org/10.1111/j.1523-1739.2007.00811.x<br>Klocker, C.A. & Strayer, D.L., 2004, 'Interactions among an invasive crayfish (*Orconectes rusticus*), a native crayfish (*Orconectes limosus*), and native bivalves (Sphaeriidae and Unionidae)', *Northeastern Naturalist*, 11, 167–178. https://doi.org/10.1656/1092-6194(2004)011[0167:IAAICO]2.0.CO;2.<br>Kreps, T.A., Baldridge, A.K. & Lodge, D.M., 2012, 'The impact of an invasive predator (*Orconectes rusticus*) on freshwater snail communities: Insights on habitat-specific effects from a multilake long-term study', *Canadian Journal of Fisheries and Aquatic Sciences*, 69, 1164–1173. https://doi.org/10.1139/f2012-052 | |

Lodge, D.M., Deines, A., Gherardi, F., Yeo, D.C.J, Arcella, T., Baldridge, A.K., Branes, M.A., Chadderton, W.L., Feder, J.L., Grantz, C.A., Howard, G.W., Jerdde, C.L., Peters, J.A., Sargent, L.W., Turner, C.R., Wittmann, M.E. & Zeng Y, 2012, 'Global introductions of crayfishes: evaluating the impact of species invasions on ecosystem services', *Annual Review of Ecology, Evolution and Systematics*, 43, 449–72. https://doi.org/10.1146/annurev-ecolsys-111511-103919.

Roth, B.M., Hein, C.L. & Vander Zanden, M.J., 2006, 'Using bioenergetics and stable isotopes to assess the trophic role of rusty crayfish (*Orconectes rusticus*) in lake littoral zones', *Canadian Journal of Fisheries and Aquatic Sciences*, 63, 335–344. https://doi.org/10.1139/f05-217.

Soes, M. & Koese, B., 2010, Invasive freshwater crayfish in the Netherlands: a preliminary risk analysis. Invasive freshwater crayfish in the Netherlands: a preliminary risk analysis.

**4. Management**

**MAN1 What is the feasibility to stop future immigration?**

| Response: High | Confidence: Low |
|---|---|

Rationale: *Faxonius immunis* is known to be present in the pet trade industry in Germany and the USA (Chucholl 2013; Faulkes 2015). It is also a popular bait species among anglers (Kerr 2014). There are no records of *Faxonius immunis* in South Africa but its potential presence in the pet trade needs to be checked (Madzivanzira et al. 2021; Nunes et al. 2017). The species is absent in neighbouring countries, so unaided immigration is unlikely. Moreover, it should be possible to regulate intentional introductions legislative regulations and having an effective border surveillance and monitoring strategy (Zengeya & Wilson 2020; Wilson & Kumschick 2024).

References:

Chucholl, C., 2013, 'Invaders for sale: trade and determinants of introduction of ornamental freshwater crayfish', *Biological Invasions* 15, 125–141. https://doi.org/10.1007/s10530-012-0273-2.

Faulkes, Z., 2015, 'The global trade in crayfish as pets', *Crustacean Research* 44: 75–92.

Madzivanzira, T.C., South, J., Wood, L.E., Nunes, A.L. & Weyl, O.L.F., 2020, 'A review of freshwater crayfish introductions in Africa', *Reviews in Fisheries Science & Aquaculture* 1-21. https://doi.org/10.1080/23308249.2020.1802405

Nunes, A.L., Zengeya, T.A., Measey, G.J. & Weyl, O.L.F., 2017b, 'Freshwater crayfish invasions in South Africa: past, present and potential future', *African Journal of Aquatic Science* 42, 309–323. https://doi.org/10.2989/16085914.2017.1405788.

Wilson, J. R., & Kumschick, S., 2024, 'The regulation of alien species in South Africa', *South African Journal of Science*, 120(5/6). https://doi.org/10.17159/sajs.2024/17002.

Zengeya, T.A. & Wilson, J.R., (eds.) 2020, *The status of biological invasions and their management in South Africa in 2019*, pp.71, South African National Biodiversity Institute, Kirstenbosch and DSI-NRF Centre of Excellence for Invasion Biology, Stellenbosch. http://dx.doi.org/10.5281/zenodo.3947613.

**MAN2 Benefits of the Taxon**

**MAN2a Socio-economic benefits of the *Taxon***

| Response: Low | Confidence: Medium |
|---|---|

Rationale: *Faxonius immunis* has been introduced as bait by anglers for recreational fishing is present in the pet trade industry in Germany and the USA (Dehus 1999; Chucholl 2013; Schrimpf et al. 2013). No socio-economic benefits have been recorded to date in South Africa and even if *F. immunis* is present the benefits are likely to be low (Madzivanzira et al. 2021).

| References: |  |
|---|---|
| Chucholl, C., 2013, 'Invaders for sale: trade and determinants of introduction of ornamental freshwater crayfish', *Biological Invasions* 15, 125–141. https://doi.org/10.1007/s10530-012-0273-2.<br>Dehus, P., Dussling. U. & Hoffmann, C., 1999, 'Notes on the occurrence of the calico crayfish (*Orconectes immunis*) in Germany', *Freshwater Crayfish*, 12, 786–790.<br>Schrimpf, A., Chucholl, C., Schmidt, T., & Schulz, R. 2013', 'Crayfish plague agent detected in populations of the invasive North American crayfish *Orconectes immunis* (Hagen, 1870) in the Rhine River, Germany', *Aquatic Invasions*, 8(1). http://dx.doi.org/10.3391/ai.2013.8.1.12.<br>Madzivanzira, T.C., South, J., Wood, L.E., Nunes, A.L. & Weyl, O.L.F., 2020, 'A review of freshwater crayfish introductions in Africa', *Reviews in Fisheries Science & Aquaculture* 1-21. https://doi.org/10.1080/23308249.2020.1802405 |  |
| **MAN2b Environmental benefits of the *Taxon*** |  |
| Response: Low | Confidence: High |
| Rationale: There are no recorded environmental benefits of *Faxonius immunis.* |  |
| References: |  |

| **MAN3 Ease of management** |  |
|---|---|
| **MAN3a How accessible are populations?** |  |
| Response: 2 (difficult to access) | Confidence: High |
| Rationale: There are no locality records for *Faxonius immunis* in South Africa (Madzivanzira et al. 2021; Nunes et al. 2017). However, given that it is an aquatic species, accessibility is likely to be difficult. |  |
| References:<br>Madzivanzira, T.C., South, J., Weyl, O.L.F., 2021, 'Invasive crayfish outperform potamonautid crabs at higher temperatures', *Freshwater Biology* 66, 978–991. https://doi.org/10.1111/fwb.13691<br>Nunes, A.L., Zengeya, T.A., Measey, G.J. & Weyl, O.L.F., 2017, 'Freshwater crayfish invasions in South Africa: past, present and potential future', *African Journal of Aquatic Science* 42, 309–323, https://doi.org/10.2989/16085914.2017.1405788 |  |
| **MAN3b Is detectability critically time-dependent?** |  |
| Response: 2 (yes) | Confidence: |
| Rationale: *Faxonius immunis* is a small, largely nocturnal freshwater species that is difficult to detect (Chucholl 2012). |  |
| References:<br>Chucholl, C., 2012, 'Understanding invasion success: life-history traits and feeding habits of the alien crayfish *Orconectes immunis* (Decapoda, Astacida, Cambaridae)', *Knowledge and Management of Aquatic Ecosystems* 404: 04. https://doi.org/10.1051/kmae/2011082. |  |
| **MAN3c Time to reproduction** |  |
| Response: 1 (1-3 years) | Confidence: High |
| Rationale: *Faxonius immunis* reach sexual maturity in one-two years (Chucholl 2012). |  |

| |
|---|
| References:<br><br>Chucholl, C., 2012, 'Understanding invasion success: life-history traits and feeding habits of the alien crayfish *Orconectes immunis* (Decapoda, Astacida, Cambaridae)', *Knowledge and Management of Aquatic Ecosystems*, (404), 04. https://doi.org/10.1051/kmae/2011082. |

| **MAN3d Propagule persistence** | |
|---|---|
| Response: 0 (<1 year) | Confidence: High |
| Rationale: *Faxonius immunis* are highly fecund, a female produces up to 261 eggs, with limited parental care (Chucholl 2012). | |
| References:<br><br>Chucholl, C., 2012, 'Understanding invasion success: life-history traits and feeding habits of the alien crayfish *Orconectes immunis* (Decapoda, Astacida, Cambaridae)', *Knowledge and Management of Aquatic Ecosystems*, (404), 04. https://doi.org/10.1051/kmae/2011082. | |
| **MAN3 Ease of management** (Table 4) | |
| Response: Medium | Confidence: High |
| Rationale: *Faxonius immunis* is a small, largely nocturnal freshwater species that is difficult to detect, reaches sexual maturity in one-two years and is highly fecund (up to 261 eggs) (Chucholl 2012). | |
| References:<br><br>Chucholl, C., 2012, 'Understanding invasion success: life-history traits and feeding habits of the alien crayfish *Orconectes immunis* (Decapoda, Astacida, Cambaridae)', *Knowledge and Management of Aquatic Ecosystems*, (404), 04. https://doi.org/10.1051/kmae/2011082. | |

| **MAN4 Has the feasibility of eradication been evaluated?** | |
|---|---|
| Response: No | Confidence: Medium |
| Rationale: There are no known population of *F. immunis* in South Africa, therefore eradication feasibility has not been no evaluated (Nunes et al. 2017) but see MAN5 for summary of control efforts used to manage invasive crayfish populations. | |
| References:<br><br>Nunes, A.L., Zengeya, T.A., Measey, G.J. & Weyl, O.L.F., 2017, 'Freshwater crayfish invasions in South Africa: past, present and potential future', *African Journal of Aquatic Science* 42, 309–323, https://doi.org/10.2989/16085914.2017.1405788 | |

| **MAN5 Control options and monitoring approaches available for the *Taxon*** |
|---|
| Response: Several methods can be used to monitor and control invasive crayfish populations including mechanical trapping (e.g., Barkhuizen et al. 2022; Madzivanzira et al. 2021), electrofishing (Nunes et al. 2017), use of biocides, and biological control (Aquiloni et al 2010; Stebbing et al. 2014). However, there are several issues that make management of crayfish invasions challenging (Gherardi et al. 2011). For example, the use of biocides is often effective for localised populations in dams and ponds (Peay et al. 2019) but they are non-specific and need to be applied with caution (Gherardi et al. 2011). Mechanical control methods are also |

effective in small discrete water bodies but are expensive and require commitment to follow on to complete eradication (Loureiro et al. 2018). In addition, alien crayfish species are regarded as potential conflict-generating species in South Africa and potential management interventions should consider the concerns of stakeholders to avoid conflict (Zengeya et al. 2017).

References:

Aquiloni, L., Brusconi, S., Cecchinelli, E., Tricarico, E., Mazza, G., Paglianti, A. & Gherardi, F. 2010, 'Biological control of invasive populations of crayfish: the European eel (*Anguilla anguilla*) as a predator of *Procambarus clarkii*', *Biological Invasions* 12 3817-3824. https://doi.org/10.1007/s10530-007-9094-0.

Barkhuizen, L.M, Madzivanzira, T.C., South, J., 2022, 'Population ecology of a wild population of red swamp crayfish *Procambarus clarkii* (Girard, 1852) in the Free State Province, South Africa and implications for eradication efforts', *BioInvasions Records* 11, 181–191. https://doi.org/10.3391/bir.2022.

Gherardi, F., Aquiloni, L., Diéguez-Uribeondo, J. & Tricarico E., 2011, 'Managing invasive crayfish: is there a hope? *Aquatic Sciences* 73, 185–200. https://doi.org/10.1007/s00027-011-0181-z.

Loureiro, T.G, Anastácio, P.M, de Siqueira Bueno, S.L. & Araujo, P.B., 2018, 'Management of invasive populations of the freshwater crayfish *Procambarus clarkii* (Decapoda, Cambaridae): test of a population-control method and proposal of a standard monitoring approach', *Environmental Monitoring Assessment* 190, 559, https://doi.org/10.1007/s10661-018-6942-6.

Madzivanzira, T.C., South, J., Nhiwatiwa, T. & Weyl, O.L.F., 2021, 'Standardisation of Australian redclaw crayfish *Cherax quadricarinatus* sampling gear in southern Africa', *Water SA* 47, 380–384. https://hdl.handle.net/10520/ejc-waters-v47-n3-a10.

Nunes, A.L., Zengeya, T.A., Hoffman, A.C., Measey, G.J. & Weyl, O.L., 2017, 'Distribution and establishment of the alien Australian redclaw crayfish, *Cherax quadricarinatus*, in South Africa and Swaziland', *PeerJ* 5, 1-21. https://doi.org/10.7717/peerj.3135.

Peay, S., Johnsen, S.I, Bean, S.W., Dunn, A.M., Sandodden, R. & Edsman, L, 2019, 'Biocide treatment of invasive signal crayfish: Successes, failures and lessons learned', *Diversity* 11, 29. https://doi.org/10.3390/d11030029

Stebbing, P., Longshaw, M., Scott, A., 2014, 'Review of methods for the management of nonindigenous crayfish, with particular reference to Great Britain', *Ethology Ecology & Evolution* 26, 204-231. https://doi.org/10.1080/03949370.2014.908326.

Woodford, D.J., Ivey, P., Novoa, A., Shackleton, R., Richardson, D., Weyl, O., Van Wilgen, B. and Zengeya, T., 2017, 'Managing conflict-generating invasive species in South Africa: Challenges and trade-offs', *Bothalia* 47, a2160. https://doi. org/10.4102/abc.v47i2.2160

| **MAN6 Any other management considerations to highlight?** (if yes, fill in Appendix MAN6) | |
|---|---|
| Response | No |

**5. Calculations**

**Likelihood = Unlikely**

| Parameter | Likelihood | Stages | Final assessment |
|---|---|---|---|
| LIK1 | Very unlikely | P(entry) = 0.027 | P (invasion) = 0.003 |
| LIK2 | Unlikely | | |
| LIK3 | Fairly Probable | P(establishment) = 0.5 | |
| LIK4 | Fairly Probable | | |

| | | |
|---|---|---|
| LIK5 | Unlikely | P (spread) = 0.5 |
| LIK6 | Fairly probable | |

**Consequence = Major (MR)**

| Parameter | Mechanism/sector | Response |
|---|---|---|
| CON1a | Competition | MO |
| CON1b | Predation | MR |
| CON1c | Hybridisation | DD |
| CON1d | Disease transmission | MN |
| CON1e | Parasitism | DD |
| CON1f | Poisoning/toxicity | DD |
| CON1g | Bio-fouling or other direct physical disturbance | MN |
| CON1h | Grazing/herbivory/browsing | MN |
| CON1i | Chemical, physical, structural impact | DD |
| CON1k | Indirect impacts through interactions with other species | DD |
| **CON1** | **Maximum environmental impact** | **MR** |
| CON2a | Safety | DD |
| CON2b | Material and immaterial assets | DD |
| CON2c | Health | DD |
| CON2d | Social, spiritual and cultural relations | MN |
| **CON2** | **Maximum socio-economic impact** | **MN** |
| CON3 | Maximum environmental impact (closely related taxa) | Not scored as CON1 scored |
| CON4 | Maximum socio-economic impact (closely related taxa) | Not scored as CON2 scored |
| CON5 | Potential impact based on traits, experiments, or models | MR |

**Risk score: High**

| | | **Consequences** | | | | |
|---|---|---|---|---|---|---|
| | | MC | MN | MO | MR | MV |
| **Likelihood** | Extremely unlikely | low | low | low | medium | medium |
| | Very unlikely | low | low | low | medium | high |
| | Unlikely | low | low | medium | high | high |
| | Fairly probable | medium | medium | high | high | high |
| | Probable | medium | high | high | high | high |

**Ease of management: Medium**

| Parameter | Question | Response |
|---|---|---|
| MAN3a | How accessible are populations? | 2 |
| MAN3b | Is detectability critically time-dependent? | 2 |
| MAN3c | Time to reproduction | 1 |
| MAN3d | Propagule persistence | 0 |
| **MAN3** | **SUM** | 5 |

**Appendix BAC7**: Global alien range for *Orconectes immunis*
Map from GBIF: https://www.gbif.org/species/2227004

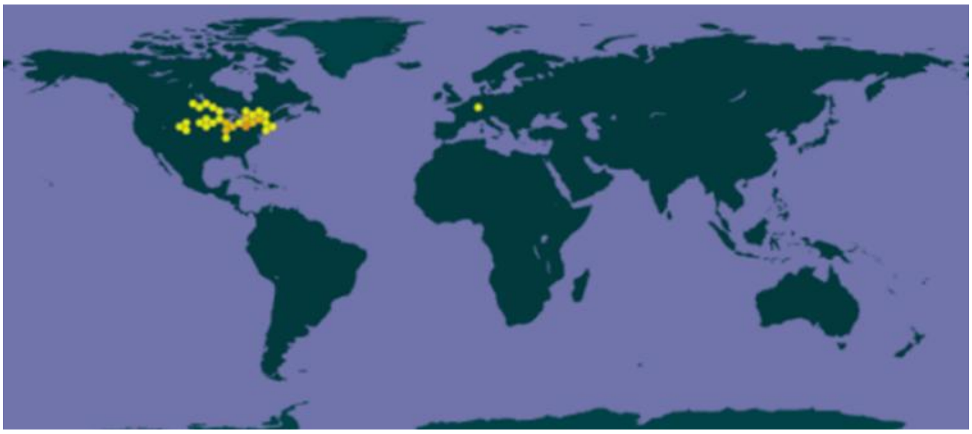

#### 1.6 Risk analysis report for Kentucky River crayfish (*Faxonius juvenilis*)

| **Taxon**: *Faxonius juvenilis* (Hagen, 1870) | **Area**: South Africa |
|---|---|
| **Picture of *Taxon***<br>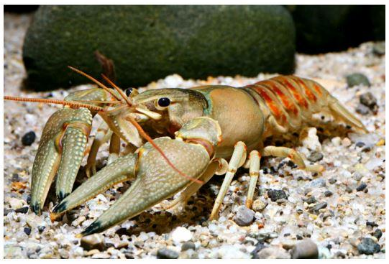<br>Chucholl & Daudey 2008. | **Alien distribution map**<br>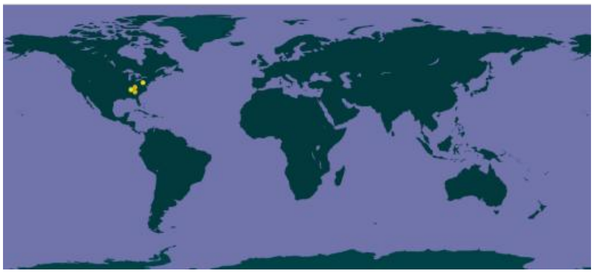<br>**https://www.gbif.org/species/5789975** |
| **Risk Assessment summary:**<br><br>Kentucky River crayfish (*Faxonius juvenilis*) are native to the Ohio River basin, USA but have been introduced to several states outside it natural range. *Faxonius juvenilis* have also been introduced to France. There are no know populations of *F. juvenilis* in South Africa and in neighbouring countries, therefore via unaided pathways is unlikely. The illegal pet trade industry remains a concern because *F. juvenilis* may be moved around by humans and there is a high risk of release of unwanted pets. There is no information on impacts caused by *F. juvenilis* and therefore impacts were inferred from *F. rusticus*, a closely related species, that has been implicated in causing detrimental impacts in its invaded range. *Faxonius rusticus* is an aggressive competitor and in its introduced range it has been implicated in causing the decline of native freshwater crayfish and fish species through competition and predation. In addition, it is an omnivore and intensive grazing have contributed to a decline in freshwater invertebrate populations and macrophyte communities. *Faxonius juvenilis* is likely to have similar impacts in areas of introduction. | **Risk score:**<br><br>High |
| **Management options summary:**<br><br>*Faxonius juvenilis* is a small, largely nocturnal freshwater species that is difficult to detect. Several methods can be used to control *F. juvenilis* such as mechanical removal using traps and electrofishing, biocides, and biological control. However, there are several issues that make the use of these methods challenging. The benefits of *F. juvenilis* were scored as low. | **Ease of management:**<br><br>Medium |
| **Recommendations:**<br><br>*Faxonius juvenilis* poses a high risk to South Africa. If *F. juvenilis* were to establish outside of captivity, such populations would likely be very difficult to extirpate.<br><br>*Faxonius juvenilis* is recommended to be added to a future prohibited species list, active efforts made to confirm that the species is absent in the country, and that any individuals found are controlled. This would likely require a dedicated assessment of the ornamental trade. A reassessment in five years is thus required and, if present in the country, *F. juvenilis* should be listed under category 1a. | **Listing under NEM:BA A&IS lists of 2020:**<br><br>Not listed<br><br>**Recommended listing category:**<br><br>Prohibited list[1] |

[1] The absence of *F. juvenilis* in South Africa should be confirmed and the taxon should be reassessed in five years (i.e., by 2029).

#### 1. Background

| | | |
|---|---|---|
| **BAC1 Name of assessor(s)** | | |
| Name of lead assessor | Lee-Anne Botha | |
| Additional assessor (1) | Tsungai Zengeya | |
| Additional assessor (2) | Christian Chimimba | |
| **BAC2 Contact details of assessor (s)** | | |
| Lead assessor | Organisational affiliation: Department of Zoology and Entomology, University of Pretoria and The South African National Biodiversity Institute (SANBI). | |
| | | |
| Additional assessor (1) | Organisational affiliation: Department of Zoology and Entomology, University of Pretoria and The South African National Biodiversity Institute (SANBI). | |
| | | |
| Additional assessor (2) | Organisational affiliation: Department of Zoology and Entomology, University of Pretoria and The South African National Biodiversity Institute (SANBI). | |
| | | |
| **BAC3 Name(s) and contact details of expert(s) consulted** | | |
| Expert (1) | Name: | |
| | email: | |
| | Phone: | |
| Expert (2) | Name: | |
| | email: | |
| | Phone: | |
| Comments: none | | |
| **BAC4 Scientific name of *Taxon* under assessment** | | |
| *Taxon* name: *Faxonius juvenilis* | | Authority: (Hagen,1870) |
| Comments: The group of surface-dwelling crayfish in the *Orconectes* genus was moved to *Faxonius* (Crandall & De Grave 2017). The taxonomic status of *Faxonius juvenilis* has been verified (GBIF 2024). | | |
| References:<br>Crandall, K.A. & De Grave, S., 2017, 'An updated classification of the freshwater crayfishes (Decapoda: Astacidea) of the world, with a complete species list', *Journal of Crustacean Biology*, 27, 615–653. https://doi.org/10.1093/jcbiol/rux070.<br>*Faxonius juvenilis* (Hagen, 1870) in GBIF Secretariat (2023). GBIF Backbone Taxonomy. Checklist dataset https://doi.org/10.15468/39omei accessed via GBIF.org on 2024-07-09. | | |
| **BAC5 Synonym(s) considered** | | |
| Synonyms: *Cambarus juvenilis* Hagen, 1870 | | |
| Comments:<br>The genus *Orconectes* previously included cave-dwelling and surface-dwelling crayfish (Crandall & De Grave 2017). The non-cave dwelling crayfish in the *Orconectes* genus were moved to *Faxonius* (Crandall & De Grave 2017). Therefore, the abovementioned synonym was used in the literature search of this risk analysis. | | |
| References:<br>Crandall, K.A. & De Grave, S., 2017, 'An updated classification of the freshwater crayfishes (Decapoda: Astacidea) of the world, with a complete species list', *Journal of Crustacean Biology*, 27, 615–653. https://doi.org/10.1093/jcbiol/rux070<br>*Faxonius juvenilis* (Hagen, 1870) in GBIF Secretariat (2023). GBIF Backbone Taxonomy. Checklist dataset https://doi.org/10.15468/39omei accessed via GBIF.org on 2024-07-09. | | |

| BAC6 Common name(s) considered | |
|---|---|
| Common names: Kentucky River crayfish | |
| Comments: Literature indicates that this the common name for *F. juvenilis* and therefore it was used in the literature search of this risk analysis (Adams et al. 2010; GBIF 2023) | |
| References:<br>Adams, S., Schuster, G.A. & Taylor, C.A. 2010, '*Orconectes juvenilis*'. *The IUCN Red List of Threatened Species* 2010: https://www.iucnredlist.org/species/153954/4568495.<br>*Faxonius juvenilis* (Hagen, 1870) in GBIF Secretariat (2023). GBIF Backbone Taxonomy. Checklist dataset https://doi.org/10.15468/39omei accessed via GBIF.org on 2024-07-09. | |
| **BAC7 What is the native range of the *Taxon*?** (add map in Appendix BAC7) | |
| Response: United States of America (Indiana, Kentucky) | Confidence: low |
| Comments: The native range of *F. juvenilis* comprises the Ohio river basin, spanning tributaries in northcentral Kentucky and southeastern Indiana (Taylor 2000; Adams et al. 2010, Daniel 2024). | |
| References:<br>Adams, S., Schuster, G.A. & Taylor, C.A. 2010, '*Orconectes juvenilis*'. *The IUCN Red List of Threatened Species* 2010: https://www.iucnredlist.org/species/153954/4568495.<br>Daniel, W.M., 2024, *Faxonius juvenilis* (Hagen, 1870): U.S. Geological Survey, Nonindigenous Aquatic Species Database, Gainesville, FL, https://nas.er.usgs.gov/queries/FactSheet.aspx?SpeciesID=2904 , Revision Date: 6/14/2017, Access Date: 7/9/2024<br>Taylor, C.A., 2000, 'Systematic studies of the *Orconectes juvenilis* complex (Decapoda: Cambaridae), with descriptions of two new species', *Journal of Crustacean Biology*, 20(1), 132-152. | |
| **BAC8 What is the global alien range of the *Taxon*?** (add map in Appendix BAC8) | |
| Response: tranlocated to several states in the USA and introduced France | Confidence: Medium |
| Comments: *Faxonius juvenilis* has been translocated outside its native range to several states in the USA (Daniels 2024) and there are known introductions in France (Chucholl & Daude 2008). | |
| References:<br>Daniel, W.M., 2024, *Faxonius juvenilis* (Hagen, 1870): U.S. Geological Survey, Nonindigenous Aquatic Species Database, Gainesville, FL, https://nas.er.usgs.gov/queries/FactSheet.aspx?SpeciesID=2904 , Revision Date: 6/14/2017, Access Date: 7/9/2024<br>Chucholl, C. & Daude, T., 2008, 'First record of *Orconectes juvenilis* (Hagen, 1870) in eastern France: update to the species identity of a recently introduced orconectid crayfish (Crustacea: Astacida)', *Aquatic Invasions*, 3, 105–107. https://doi.org/10.3391/ai.2008.3.1.19. | |
| **BAC9 Geographic scope = the *Area* under consideration** | |
| *Area* of assessment: South Africa | |
| Comments: The geographic scope of this assessment is limited to South Africa. | |
| **BAC10 Is the *Taxon* present in the *Area*?** | |
| Response: No | Confidence: Low |
| Comments: There are no records of species being in the country (Nunes et al. 2017). *Faxonius juvenilis* is present in pet trade globally (Chucholl 2013; Faulkes 2015), but there are no verified records of its presence in South Africa. | |
| References:<br>Chucholl, C., 2013, 'Invaders for sale: trade and determinants of introduction of ornamental freshwater crayfish', *Biological Invasions* 15, 125–141. https://doi.org/10.1007/s10530-012-0273-2<br>Faulkes, Z., 2015, 'The global trade in crayfish as pets', *Crustacean Research*, 44, 75–92. https://doi.org/10.18353/crustacea.44.0_75<br>Nunes, A.L., Zengeya, T.A., Measey, G.J. & Weyl, O.L.F. 2017, 'Freshwater crayfish invasions in South Africa: past, present and potential future', *African Journal of Aquatic Science*, 42, 309–323. https://doi.org/10.2989/16085914.2017.1405788. | |
| **BAC11 Availability of physical specimen** | |
| Response: No | Confidence in ID: |

Herbarium or museum accession number: No *Faxonius juvenilis* specimens have been found in South Africa and it is assumed the species is not present in South Africa, although it requires further assessment, especially linked to the pet trade industry (Nunes et al. 2017).

References:

Nunes, A.L., Zengeya, T.A., Measey, G.J. & Weyl, O.L.F. 2017, 'Freshwater crayfish invasions in South Africa: past, present and potential future', *African Journal of Aquatic Science*, 42, 309–323. https://doi.org/10.2989/16085914.2017.1405788.

**BAC12 Is the *Taxon* native to the *Area* or part of the *Area*?**

| | | |
|---|---|---|
| The *Taxon* is native to (part of) the *Area*. | No | Confidence: High |
| The *Taxon* is alien in (part of) the *Area*. | No | Confidence: High |

Comments: There are no native freshwater crayfish species native in South Africa (de Moor 2002; Nunes et al. 2017). *Faxonius juvenilis* is native to the Ohio river basin in the USA (Taylor et al. 2000) and has been introduced globally through the pet trade (Chucholl & Daude 2008, Chucholl 2013). However, there are no verified records of its presence in South Africa (Nunes et al. 2017).

References:

de Moor, I., 2002, 'Potential impacts of alien freshwater crayfish in South Africa', *African Journal of Aquatic Science* 27, 125–139. https://doi.org/10.2989/16085914.2002.9626584

Chucholl, C. & Daude, T., 2008, 'First record of *Orconectes juvenilis* (Hagen, 1870) in eastern France: update to the species identity of a recently introduced orconectid crayfish (Crustacea: Astacida)', *Aquatic Invasions*, 3,105–107. https://doi.org/10.3391/ai.2008.3.1.19.

Chucholl, C., 2013, 'Invaders for sale: trade and determinants of introduction of ornamental freshwater crayfish', *Biological Invasions* 15, 125–141. https://doi.org/10.1007/s10530-012-0273-2.

Nunes, A.L., Zengeya, T.A., Measey, G.J. & Weyl, O.L.F., 2017b, 'Freshwater crayfish invasions in South Africa: past, present and potential future', *African Journal of Aquatic Science* 42, 309–323. https://doi.org/10.2989/16085914.2017.1405788.

Taylor, C.A. 2000, 'Systematic studies of the *Orconectes juvenilis* complex (Decapoda: Cambaridae), with descriptions of two new species', *Journal of Crustacean Biology*, 20(1), 132-152. https://doi.org/10.1163/20021975-99990023.

**BAC13 What is the *Taxon*'s introduction status in the *Area*?**

| | | |
|---|---|---|
| The *Taxon* is in cultivation/containment. | Don't know | Confidence: Low |
| The *Taxon* is present in the wild. | No | Confidence: Low |
| The *Taxon* has established/naturalised. | No | Confidence: Low |
| The *Taxon* is invasive. | No | Confidence: Low |

Comments: *Faxonius juvenilis* is present in the pet trade globally (Chucholl & Daude 2008, Chucholl 2013) but there are no verified records of its presence in South Africa (Nunes et al. 2017).

References:

Chucholl, C. & Daude, T., 2008, 'First record of *Orconectes juvenilis* (Hagen, 1870) in eastern France: update to the species identity of a recently introduced orconectid crayfish (Crustacea: Astacida)', *Aquatic Invasions*, 3,105–107. https://doi.org/10.3391/ai.2008.3.1.19.

Chucholl, C., 2013, 'Invaders for sale: trade and determinants of introduction of ornamental freshwater crayfish', *Biological Invasions* 15, 125–141. https://doi.org/10.1007/s10530-012-0273-2.

Nunes, A.L., Zengeya, T.A., Measey, G.J. & Weyl, O.L.F., 2017b, 'Freshwater crayfish invasions in South Africa: past, present and potential future', *African Journal of Aquatic Science* 42, 309–323. https://doi.org/10.2989/16085914.2017.1405788

**BAC14 Primary (introduction) pathways**

| | | |
|---|---|---|
| Release | None | Confidence: |
| Escape | Pet/aquarium/terrarium species (including live food for such species) | Confidence: Medium |
| Contaminant | Live food and bait | Confidence: Medium |
| Stowaway | None | Confidence: Low |
| Corridor | None | Confidence: Low |
| Unaided | None | Confidence: Low |

| |
|---|
| Comments: *Faxonius juvenilis* is known to be present in the pet trade industry globally (Chucholl 2013; Faulkes 2015). It has also been introduced through accidental release of live individual kept for food and as bait (Holdich et al. 2009; Chucholl & Daude 2008, Daniels 2024). For example, in France it is suspected that individuals escaped from a breeding population in ponds at a restaurant advertising crayfish as a delicacy into the Dessoubre river in eastern France where it was discovered in 2005 and has since become established (Chucholl & Daude 2008). |
| References:<br>Chucholl, C. & Daude, T., 2008, 'First record of *Orconectes juvenilis* (Hagen, 1870) in eastern France: update to the species identity of a recently introduced orconectid crayfish (Crustacea: Astacida)', *Aquatic Invasions*, 3,105–107. https://doi.org/10.3391/ai.2008.3.1.19.<br>Chucholl, C., 2013, 'Invaders for sale: trade and determinants of introduction of ornamental freshwater crayfish', *Biological Invasions* 15, 125–141. https://doi.org/10.1007/s10530-012-0273-2<br>Daniel, W.M., 2024, *Faxonius juvenilis* (Hagen, 1870): U.S. Geological Survey, Nonindigenous Aquatic Species Database, Gainesville, FL, https://nas.er.usgs.gov/queries/FactSheet.aspx?SpeciesID=2904 , Revision Date: 6/14/2017, Access Date: 7/9/2024<br>Faulkes, Z., 2015, 'The global trade in crayfish as pets', *Crustacean Research*, 44, 75–92. https://doi.org/10.18353/crustacea.44.0_75.<br>Holdich, D.M., Reynolds, J.D., & Sibley, P.J., 2009, 'A review of the ever increasing threat to European crayfish from non-indigenous crayfish species', *Knowledge and Management of Aquatic Ecosystems* 11, 394–395. https://doi.org/10.1051/kmae/2009025. |

**2. Likelihood**

| **LIK1 Likelihood of entry via unaided primary pathways** | |
|---|---|
| Response: Very unlikely (p = 0.0027) | Confidence: Medium |
| Rationale: There is no evidence that *F. juvenilis* is present in South Africa or has established in the wild in neighbouring countries in the SADC Region (Lodge et al. 2012; Nunes et al. 2017; Madzivanzira et al. 2020) therefore the probability of *F. juvenilis* entering South Africa through unaided pathways (connected waterways) is very unlikely (p = 0.0027). | |
| References:<br>Lodge, D.M., Deines, A., Gherardi, F., Yeo, D.C.J., Arcella, T., Baldridge, A.K., Branes, M.A., Chadderton, W.L., Feder, J.L., Grantz, C.A., Howard, G.W., Jerdde, C.L., Peters, J.A., Sargent, L.W., Turner, C.R., Wittmann, M.E. & Zeng, Y., 2012, 'Global introductions of crayfishes: evaluating the impact of species invasions on ecosystem services', *Annual Review of Ecology Evolution Systematics* 43, 449–472, , https://doi.org/10.1146/annurev-ecolsys-111511-103919.<br>Madzivanzira, T.C., South, J., Wood, L.E., Nunes, A.L. & Weyl, O.L.F., 2020, 'A review of freshwater crayfish introductions in Africa', *Reviews in Fisheries Science & Aquaculture* 1-21. https://doi.org/10.1080/23308249.2020.1802405.<br>Nunes, A.L., Zengeya, T.A., Measey, G.J. & Weyl, O.L.F., 2017b, 'Freshwater crayfish invasions in South Africa: past, present and potential future', *African Journal of Aquatic Science* 42, 309–323. https://doi.org/10.2989/16085914.2017.1405788. | |

| **LIK2 Likelihood of entry via human aided primary pathways** | |
|---|---|
| Response: Fairly probable (p = 0.5) | Confidence: Medium |
| Rationale: *Faxonius juvenilis* is present in the global trade (Holdich et al. 2009; Chucholl 2013), it is likely that it could be introduced in the pet trade in South Africa. | |

| |
|---|
| References:<br><br>Chucholl, C., 2013, 'Invaders for sale: trade and determinants of introduction of ornamental freshwater crayfish', *Biological Invasions* 15, 125–141. https://doi.org/10.1007/s10530-012-0273-2.<br>Holdich, D.M., Reynolds, J.D., & Sibley, P.J., 2009, 'A review of the ever increasing threat to European crayfish from non-indigenous crayfish species', *Knowledge and Management of Aquatic Ecosystems* 11, 394–395. https://doi.org/10.1051/kmae/2009025. |

| **LIK3 Habitat suitability** | |
|---|---|
| Response: Fairly probable (p = 0.5) | Confidence: Medium |
| Rationale: *Faxonius juvenilis* is a habitat generalist that inhabits small streams to rivers with a range of substrates such as mud, gravel and bedrock substrates (Daniels 2024). *Faxonius juvenilis* prefers areas that consist of rocks, logs or undercut banks that provide shelter (Taylor 2000). These habitats are present in most aquatic systems across South Africa. | |
| References:<br><br>Daniel, W.M., 2024, *Faxonius juvenilis* (Hagen, 1870): U.S. Geological Survey, Nonindigenous Aquatic Species Database, Gainesville, FL, https://nas.er.usgs.gov/queries/FactSheet.aspx?SpeciesID=2904 , Revision Date: 6/14/2017, Access Date: 7/9/2024<br>Taylor, C.A., 2000, 'Systematic studies of the *Orconectes juvenilis* complex (Decapoda: Cambaridae), with descriptions of two new species', *Journal of Crustacean Biology*, 20(1), 132-152. https://doi.org/10.1163/20021975-99990023 | |

| **LIK4 Climate suitability** | |
|---|---|
| Response: Fairly probable (p = 0.5) | Confidence: Medium |
| Rationale: Suitable climates are present in South Africa for *F. juvenilis* to establish. There is no study that has assessed the climate suitability of South Africa for *F. juvenilis* however its ecological traits are assumed to be similar to *F. rusticus* which is known to prefer well oxygenated water and an optimal temperature range of 20-25°C and but can withstand seasonal water temperatures of 0-39°C within its native range (GISD 2015). In temperatures over 30°C, adults have been observed digging burrows to escape the heat (GISD 2015). | |
| References:<br><br>Global Invasive Species Database (GISD) 2015. Species profile *Orconectes rusticus*. Available from: http://www.iucngisd.org/gisd/species.php?sc=217 [Accessed 03 July2018]. | |

| **LIK5 Unaided secondary (dispersal) pathways** | |
|---|---|
| Response: Very unlikely (p = 0.0027) | Confidence: Low |
| Rationale: Currently this is very unlikely due to no feral populations present in neighbouring countries that could disperse naturally through connected waterways (Nunes et al. 2019; Madzivanzira et al. 2020). | |
| References:<br>Madzivanzira, T.C., South, J., Wood, L.E., Nunes, A.L. & Weyl, O.L.F., 2020, 'A review of freshwater crayfish introductions in Africa', *Reviews in Fisheries Science & Aquaculture* 1-21. https://doi.org/10.1080/23308249.2020.1802405 | |

| |
|---|
| Nunes, A.L., Zengeya, T.A., Measey, G.J. & Weyl, O.L.F., 2017b, 'Freshwater crayfish invasions in South Africa: past, present and potential future', *African Journal of Aquatic Science* 42, 309–323. https://doi.org/10.2989/16085914.2017.1405788 |

| **LIK6 Human aided secondary (dispersal) pathways** | |
|---|---|
| Response: Fairly probable (p = 0.5) | Confidence: Low |
| Rationale: Intentional release into the wild through the pet/ aquarium/ terrarium species and live bait pathways remains a concern. In areas of introduction, *Faxonius juvenilis* has been released into the wild through deliberate stocking for human consumption and bucket release by anglers into waterways where they are used as bait, dispersing into new areas (Taylor 2000; Chuchol & Daudely 2008). | |
| References:<br><br>Chucholl, C. & Daude, T., 2008, 'First record of *Orconectes juvenilis* (Hagen, 1870) in eastern France: update to the species identity of a recently introduced orconectid crayfish (Crustacea: Astacida)', *Aquatic Invasions*, 3, 105–107. https://doi.org/10.3391/ai.2008.3.1.19<br>Taylor, C.A., 2000, 'Systematic studies of the *Orconectes juvenilis* complex (Decapoda: Cambaridae), with descriptions of two new species', *Journal of Crustacean Biology*, 20(1), 132-152. https://doi.org/10.1163/20021975-99990023 | |

**3. Consequences**

| **CON1 Environmental impact** | |
|---|---|
| **CON1a: Competition** | |
| Response: Data Deficient (DD) | Confidence: |
| Rationale: No documented information available to assess the level of impact. | |
| References: | |
| **CON1b: Predation** | |
| Response: Data Deficient (DD) | Confidence: |
| Rationale: No documented information available to assess the level of impact. | |
| References: | |
| **CON1c: Hybridisation** | |
| Response: Data Deficient (DD) | Confidence: |
| Rationale: No documented information available to assess the level of impact. | |
| References: | |
| **CON1d: Transmission of disease** | |
| Response: Data Deficient (DD) | Confidence: |
| Rationale: No documented information available to assess the level of impact. | |

| | |
|---|---|
| References: | |
| **CON1e: Parasitism** | |
| Response: Data Deficient (DD) | Confidence: |
| Rationale: No documented information available to assess the level of impact. | |
| References: | |
| **CON1f: Poisoning/toxicity** | |
| Response: Data Deficient (DD) | Confidence: |
| Rationale: No documented information available to assess the level of impact. | |
| References: | |
| **CON1g: Bio-fouling or other direct physical disturbance** | |
| Response: Data Deficient (DD) | Confidence: |
| Rationale: No documented information available to assess the level of impact. | |
| References: | |
| **CON1h: Grazing/herbivory/browsing** | |
| Response: Data Deficient (DD) | Confidence: |
| Rationale: No documented information available to assess the level of impact. | |
| References: | |
| **CON1i: Chemical, physical or structural impact on ecosystem** | |
| Response: Data Deficient (DD) | Confidence: |
| Rationale: No documented information available to assess the level of impact. | |
| References: | |
| **CON1k: Indirect impacts through interactions with other species** | |
| Response: Data Deficient (DD) | Confidence: |
| Rationale: No documented information available to assess the level of impact. | |
| References: | |
| **CON1 Maximum environmental impact** | |
| Response: Data Deficient (DD) | Confidence: |
| Rationale: No documented information available to assess the level of impact. | |
| References: | |

| CON2 Socio-economic impact | |
|---|---|
| **CON2a: Safety** | |
| Response: Data Deficient (DD) | Confidence: |
| Rationale: No documented information available to assess the level of impact. | |
| References: | |
| **CON2b: Material and immaterial assets** | |
| Response: Data Deficient (DD) | Confidence: |
| Rationale: No documented information available to assess the level of impact. | |
| References: | |
| **CON2c: Health** | |
| Response: Data Deficient (DD) | Confidence: |
| Rationale: No documented information available to assess the level of impact. | |
| References: | |
| **CON2d: Social, spiritual and cultural relations** | |
| Response: Data Deficient (DD) | Confidence: |
| Rationale: No documented information available to assess the level of impact. | |
| References: | |
| **CON2 Maximum socio-economic impact** | |
| Response: Data Deficient (DD) | Confidence: |
| Rationale: No documented information available to assess the level of impact. | |
| References: | |

| CON3 Closely related species' environmental impact | |
|---|---|
| Response: Major (MR) | Confidence: High |
| Rationale: There are no recorded environmental impacts for *F. juvenilis*, therefore impacts were inferred from a closely related taxon, *Faxonius rusticus.* In introduced areas, direct predation on native invertebrates, competition for food, shelter and spawning sites have led to local extinctions and a decrease in the abundance of native crayfish and fish species (e.g., Garvey & Stein 1993; Klocker & Strayer 2004; Lodge et al. 2012). *Faxonius rusticus* feeds on macrophytes leading to changes in structural communities (e.g., Hobb et al. 1989, Lodge et al. 1994; Wilson et al. 2004, Roth et al. 2006; Rosenthal et al. 2006; Baldridge & Lodge 2014). *Faxinous rusticus* is a known vector of pathogen such as the crayfish plague agent, Aphanomyces astaci (Panteleit et al. 2019; Butler et al. 2020) and microsporidia (Cambaraspora sp.) (Stratton et al. 2023). *Faxonius* | |

| |
|---|
| *rusticus* invasions have caused disruption in recreational activities by reducing sport fish populations though egg predation and/or competition with juveniles (Keller et al. 2008). |

References:

Baldridge, A. K., & Lodge, D. M., 2014, 'Long-term studies of crayfish invaded lakes reveal limited potential for macrophyte recovery from the seed bank', *Freshwater Science*, 33, 788–797. https://doi.org/10.1086/677070.

Butler, E., Crigler, P., Robbins, G. & Blair, J.E., 2020, 'Preliminary survey of *Aphanomyces* sp. associated with native and invasive crayfish in the lower Susquehanna watershed of south central Pennsylvania', *Journal of Freshwater Ecology*, 35(1), 223-233. https://doi.org/10.1080/02705060.2020.1779141.

Garvey, J.E. & Stein, R.A, 1993, 'Evaluating how chela size influences the invasion potential of an introduced crayfish (*Orconectes rusticus*)', *American Midland Naturalist* 129, 172–181. https://doi.org/10.2307/2426446.

Hobbs, H.H., Jass, J.P. & Huner, J.V., 1989, 'A review of global crayfish introductions with particular emphasis on two North American species (Decapoda, Cambaridae)', *Crustaceana* 56(3), 299-316. https://www.jstor.org/stable/20104461.

Klocker, C.A. & Strayer, D.L., 2004, 'Interactions among an invasive crayfish (*Orconectes rusticus*), a native crayfish (*Orconectes limosus*), and native bivalves (Sphaeriidae and Unionidae)', *Northeastern Naturalist* 11, 167–178. https://doi.org/10.1656/1092-6194(2004)011[0167:IAAICO]2.0.CO;2.

Keller, R.P., Frang, K. & Lodge, D.M., 2008, 'Preventing the spread of invasive species: Economic benefits of intervention guided by ecological predictions' *Conservation Biology* 22, 80–88. https://doi.org/10.1111/j.1523-1739.2007.00811.

Lodge, D.M., Deines, A., Gherardi, F., Yeo, D.C.J., Arcella. T, Baldridge, A.K., et al., 2012, 'Global introductions of crayfishes: Evaluating the impact of species invasions on ecosystem services', *Annual Review of Ecology Evolution Systematics* 43, 449–472. https://doi.org/10.1146/annurev-ecolsys-111511-103919.

Lodge, D.M., Kershner, M.W., Aloi, J.E. & Covich, A.P., 1994, 'Effects of an omnivorous crayfish (*Orconectes rusticus*) on a freshwater littoral food web', *Ecology*, 75, 1265-1281. https://doi.org/10.2307/1937452

Panteleit, J., Horvath, T., Jussila, J., Makkonen, J., Perry, W., Schulz, R., Theissinger, K. & Schrimpf, A., 2019. Invasive rusty crayfish (*Faxonius rusticus*) populations in North America are infected with the crayfish plague disease agent (*Aphanomyces astaci*)', *Freshwater Science*, 38(2), 425-433. https://doi.org/10.1086/703417

Perry, W.L., Feder, J.L., Dwyer, G. & Lodge, D.M., 2001, 'Hybrid zone dynamics and species replacement between *Orconectes* crayfishes in a northern Wisconsin lake', *Evolution* 55, 1153–1166. https://doi.org/10.1111/j.0014-3820.2001.tb00635.x.

Rosenthal, S. K., Stevens, S. S., & Lodge, D. M., 2006, 'Whole-lake effects of invasive crayfish (*Orconectes* spp.) and the potential for restoration', *Canadian Journal of Fisheries and Aquatic Sciences*, 63, 1276–1285. https://doi.org/10.1139/f06-037.

Roth, B.M, Hein, C.L. & Vander Zanden, M.J., 2006, 'Using bioenergetics and stable isotopes to assess the trophic role of rusty crayfish (*Orconectes rusticus*) in lake littoral zones,' *Canadian Journal of Fisheries and Aquatic Sciences* 63, 335–344. https://doi.org/10.1139/f05-217

Stratton, C. E., Kabalan, B. A., Bolds, S. A., Reisinger, L. S., Behringer, D. C. & Bojko, J., 2023, '*Cambaraspora faxoni* n. sp. (Microsporidia: Glugeida) from native and invasive crayfish in the USA and a novel host of *Cambaraspora floridanus', Journal of Invertebrate Pathology* 199:107949. https://doi.org/10.1016/j.jip.2023.107949.

Wilson, K.A., Magnuson, J.J., Lodge, D.M., Hill, A.M., Kratz, T.K., Perry, W.L., et al., 2004, 'A long-term rusty crayfish (*Orconectes rusticus*) invasion: Dispersal patterns and community change in a north temperate lake', *Canadian Journal of Fisheries and Aquatic Sciences* 61, 2255–2266. https://doi.org/10.1139/F04-170.

| **CON4 Closely related species' socio-economic impact** | |
|---|---|
| Response: Moderate (MO) | Confidence: Low |
| Rationale: There are no recorded socio-economic impacts for *F. juvenilis*, therefore impacts were inferred from a closely related taxon, *Faxonius rusticus. Faxonius rusticus* invasions have caused disruption in recreational activities by reducing sport fish populations though egg predation and/or competition with juveniles (Keller et al. 2008). | |
| References:<br><br>Keller, R.P., Frang, K. & Lodge, D.M., 2008, 'Preventing the spread of invasive species: Economic benefits of intervention guided by ecological predictions' *Conservation Biology* 22, 80–88. https://doi.org/10.1111/j.1523-1739.2007.00811.x. | |

| **CON5 Potential impact** | |
|---|---|
| Response: MR | Confidence: High |
| Rationale: There are no recorded environmental impacts for *F. juvenilis*, therefore potential impacts were inferred from a closely related taxon, *Faxonius rusticus* that is known to cause impacts in introduced areas, mainly through competition, predation and excessive grazing (Lodge et al. 2012). *Faxonius rusticus* is highly fecund and an aggressive competitor (Garvey & Stein 1993). *Faxonius rusticus* may out-compete native freshwater crab species in South Africa (Jackson et al. 2016; Twardochleb et al. 2018). Being a functional omnivore, *F. rusticus* may also have detrimental impacts on macroinvertebrates and macrophyte communities when occurring in high densities (Wilson et al. 2004; Roth et al. 2006). *Faxonius rusticus* can reduce invertebrate communities, changing their composition in invaded areas (Klocker & Strayer 2004; Johnson et al. 2009). Furthermore, being a vector for the crayfish plague is another cause of concern, as it may be transferable and could be detrimental to native freshwater crustaceans in South Africa (Twardochleb et al. 2013; Panteleit et al. 2019; Butler et al. 2020)). Although *F. rusticus* has not been implicated as an active vector, all north American crayfish are immune to the disease and can easily infect other species. | |
| References:<br>Butler, E., Crigler, P., Robbins, G. & Blair, J. E., 2020, 'Preliminary survey of *Aphanomyces* sp. associated with native and invasive crayfish in the lower Susquehanna watershed of south central Pennsylvania', *Journal of Freshwater Ecology*, 35(1), 223-233. https://doi.org/10.1080/02705060.2020.1779141.<br>Garvey, J.E. & Stein, R.A., 1993, 'Evaluating how chela size influences the invasion potential of an introduced crayfish (*Orconectes rusticus*)', *American Midland Naturalist 129,* 172–181. https://doi.org/10.2307/2426446.<br>Holdich, D.M. & Reeve, I.D., 1991, 'Distribution of freshwater crayfish in the British Isles, with particular reference to crayfish plague, alien introductions and water quality', *Aquatic Conservation: Marine and Freshwater Ecosystems* 1, 139–158. https://doi.org/10.1002/aqc.3270010204.<br>Johnson, P.T., Olden, J.D., Solomon, C.T. & Vander Zanden, M.J., 2009, 'Interactions among Invaders: Community and ecosystem effects of multiple invasive species in an experimental aquatic system', *Oecologia* 159, 161–170. https://doi.org/10.1007/s00442-008-1176-x.<br>Klocker, C.A. & Strayer, D.L., 2004, 'Interactions among an invasive crayfish (*Orconectes rusticus*), a native crayfish (*Orconectes limosus*), and native bivalves (Sphaeriidae and Unionidae)', *Northeastern Naturalist* 11, 167–178. https://doi.org/10.1656/1092-6194(2004)011[0167:IAAICO]2.0.CO;2.<br>Lodge, D.M., Kershner, M.W., Aloi, J.E. & Covich, A.P., 1994, Effects of an omnivorous crayfish (*Orconectes rusticus*) on a freshwater littoral food web. *Ecology*, 75, 1265-1281. https://doi.org/10.2307/1937452<br>Panteleit, J., Horvath, T., Jussila, J., Makkonen, J., Perry, W., Schulz, R., Theissinger, K. & Schrimpf, A., 2019, 'Invasive rusty crayfish (*Faxonius rusticus*) populations in North America are infected with the | |

crayfish plague disease agent (*Aphanomyces astaci*)', *Freshwater Science*, 38(2), 425-433. https://doi.org/10.1086/703417

Roth, B.M, Hein, C.L. & Vander Zanden, M.J., 2006, 'Using bioenergetics and stable isotopes to assess the trophic role of rusty crayfish (*Orconectes rusticus*) in lake littoral zones,' *Canadian Journal of Fisheries and Aquatic Sciences* 63, 335–344. https://doi.org/10.1139/F05-217.

Twardochleb, L.A, Olden, J.D. & Larson, E.R., 2013, 'A global meta-analysis of the ecological impacts of nonnative crayfish', *Freshwater Science* 32, 1367–1382.

Wilson, K.A., Magnuson, J.J., Lodge, D.M., Hill, A.M., Kratz, T.K., Perry, W.L., et al., 2004, 'A long-term rusty crayfish (*Orconectes rusticus*) invasion: Dispersal patterns and community change in a north temperate lake', *Canadian Journal of Fisheries and Aquatic Sciences* 61, 2255–2266. https://doi.org/10.1139/f04-170.

**4. Management**

| **MAN1 What is the feasibility to stop future immigration?** | |
|---|---|
| Response: High | Confidence: low |
| Rationale: *Faxonius juvenilis* is absent in neighbouring countries, so unaided immigration is unlikely. Moreover, it should be possible to regulate intentional introductions through legislative regulations and having an effective border surveillance and monitoring strategy (Zengeya & Wilson 2020, Wilson and Kumschick 2024). | |
| References:<br>Zengeya, T.A. & Wilson, J.R., (eds.) 2020, *The status of biological invasions and their management in South Africa in 2019*, pp.71, South African National Biodiversity Institute, Kirstenbosch and DSI-NRF Centre of Excellence for Invasion Biology, Stellenbosch. http://dx.doi.org/10.5281/zenodo.3947613.<br>Wilson, J. R., & Kumschick, S., 2024, 'The regulation of alien species in South Africa', *South African Journal of Science*, 120(5/6). https://doi.org/10.17159/sajs.2024/17002. | |

| **MAN2 Benefits of the Taxon** | |
|---|---|
| **MAN2a Socio-economic benefits of the *Taxon*** | |
| Response: Low | Confidence: Low |
| Rationale: *Faxonius juvenilis* is used as a bait species among anglers in the USA and it was intentional introduced for human consumption in France (Taylor 2000; Chucholl & Daude 2008). No socio-economic benefits have been recorded to date in South Africa and even if *F.juvenilis* is present the benefits are likely to be low (Madzivanzira et al. 2021). | |
| References:<br><br>Chucholl, C. & Daude, T., 2008, 'First record of *Orconectes juvenilis* (Hagen, 1870) in eastern France: update to the species identity of a recently introduced orconectid crayfish (Crustacea: Astacida)', *Aquatic Invasions* 3:105–107, https://doi.org/10.3391/ai.2008.3.1.19<br>Taylor, C.A., 2000, 'Systematic studies of the *Orconectes juvenilis* complex (Decapoda: Cambaridae), with descriptions of two new species', *Journal of Crustacean Biology*, 20(1), 132-152. https://doi.org/10.1163/20021975-99990023.<br>Madzivanzira, T.C., South, J., Wood, L.E., Nunes, A.L. & Weyl, O.L.F., 2020, 'A review of freshwater crayfish introductions in Africa', *Reviews in Fisheries Science & Aquaculture*, 1-21. https://doi.org/10.1080/23308249.2020.1802405. | |
| **MAN2b Environmental benefits of the *Taxon*** | |
| Response: Low | Confidence: High |
| Rationale: There are no recorded environmental benefits of *Faxonius juvenilis.* | |
| References: | |

| **MAN3 Ease of management** | |
|---|---|
| **MAN3a How accessible are populations?** | |
| Response: 2 (difficult to access) | Confidence: Low |

| |
|---|
| Rationale: There are no locality records for *Faxonius juvenilis* in South Africa (Nunes et al. 2017). However, given that it is a small and largely nocturnal aquatic species, accessibility is likely to be difficult (Taylor et al. 2000). |
| References:<br>Nunes, A.L., Zengeya, T.A., Measey, G.J. & Weyl, O.L.F., 2017, 'Freshwater crayfish invasions in South Africa: past, present and potential future', *African Journal of Aquatic Science* 42, 309–323, https://doi.org/10.2989/16085914.2017.1405788. |

| **MAN3b Is detectability critically time-dependent?** | |
|---|---|
| Response: 2 (yes) | Confidence: High |
| Rationale: *Faxonius juvenilis* is a small, largely nocturnal freshwater species that is difficult to detect (Taylor & Schuster 2004). | |
| References:<br>Taylor, C.A. & Schuster, G.A. 2004, 'Crayfishes of Kentucky', Illinois Natural History Survey, Special Publication 28. | |

| **MAN3c Time to reproduction** | |
|---|---|
| Response: 1 (1-3 years) | Confidence: High |
| Rationale: *Faxonius juvenilis* reach sexual maturity in one-two years (Taylor & Schuster 2004). | |
| References:<br>Taylor, C.A. & Schuster, G.A. 2004, 'Crayfishes of Kentucky', Illinois Natural History Survey, Special Publication 28. | |

| **MAN3d Propagule persistence** | |
|---|---|
| Response: 0 (< 1 year) | Confidence: High |
| Rationale: *Faxonius juvenilis* are highly fecund, a female produces up to 141 eggs, and with some parental care of eggs and hatchlings (Taylor & Schuster 2004). | |
| References:<br>Taylor, C.A. & Schuster, G.A. 2004, 'Crayfishes of Kentucky', Illinois Natural History Survey, Special Publication 28. | |

| **MAN3 Ease of management** (Table 4) | |
|---|---|
| Response: Medium | Confidence: High |
| Rationale: *Faxonius juvenilis* is a small, largely nocturnal freshwater species that is difficult to detect, it reaches sexual maturity in one-two years and is highly fecund (up to 141 eggs) (Taylor & Schuster 2004). | |
| References:<br>Taylor, C.A. & Schuster, G.A. 2004, 'Crayfishes of Kentucky', Illinois Natural History Survey, Special Publication 28. | |

| MAN4 Has the feasibility of eradication been evaluated? | |
|---|---|
| Response: No | Confidence: Low |
| Rationale: There are no known population of *F. juvenilis* in South Africa, therefore eradication feasibility has not been no evaluated (Nunes et al. 2017) but see MAN5 for summary of control efforts used to manage invasive crayfish populations. | |
| References:<br><br>Nunes, A.L., Zengeya, T.A., Measey, G.J. & Weyl, O.L.F., 2017, 'Freshwater crayfish invasions in South Africa: past, present and potential future', *African Journal of Aquatic Science* 42, 309–323, https://doi.org/10.2989/16085914.2017.1405788. | |

| MAN5 Control options and monitoring approaches available for the *Taxon* |
|---|
| Response: Several methods can be used to monitor and control invasive crayfish populations including mechanical trapping (e.g., Barkhuizen et al. 2022; Madzivanzira et al. 2021), electrofishing (Nunes et al. 2017), use of biocides, and biological control (Aquiloni et al 2010; Stebbing et al. 2014). However, there are several issues that make management of crayfish invasions challenging (Gherardi et al. 2011). For example, the use of biocides is often effective for localised populations in dams and ponds (Peay et al. 2019) but they are non-specific and need to be applied with caution (Gherardi et al. 2011). Mechanical control methods are also effective in small discrete water bodies but are expensive and require commitment to follow on to complete eradication (Loureiro et al. 2018). In addition, alien crayfish species are regarded as potential conflict-generating species in South Africa and potential management interventions should consider the concerns of stakeholders to avoid conflict (Zengeya et al. 2017). |
| References:<br><br>Aquiloni, L., Brusconi, S., Cecchinelli, E., Tricarico, E., Mazza, G., Paglianti, A. & Gherardi, F. 2010, 'Biological control of invasive populations of crayfish: the European eel (*Anguilla anguilla*) as a predator of *Procambarus clarkii*', *Biological Invasions* 12 3817-3824. https://doi.org/10.1007/s10530-007-9094-0.<br>Barkhuizen, L.M, Madzivanzira, T.C., South, J., 2022, 'Population ecology of a wild population of red swamp crayfish *Procambarus clarkii* (Girard, 1852) in the Free State Province, South Africa and implications for eradication efforts', *BioInvasions Records* 11, 181–191. https://doi.org/10.3391/bir.2022.<br>Gherardi, F., Aquiloni, L., Diéguez-Uribeondo, J. & Tricarico E., 2011, 'Managing invasive crayfish: is there a hope? *Aquatic Sciences* 73, 185–200. https://doi.org/10.1007/s00027-011-0181-z.<br>Loureiro, T.G, Anastácio, P.M, de Siqueira Bueno, S.L. & Araujo, P.B., 2018, 'Management of invasive populations of the freshwater crayfish *Procambarus clarkii* (Decapoda, Cambaridae): test of a population-control method and proposal of a standard monitoring approach', *Environmental Monitoring Assessment* 190, 559, https://doi.org/10.1007/s10661-018-6942-6.<br>Madzivanzira, T.C., South, J., Nhiwatiwa, T. & Weyl, O.L.F., 2021, 'Standardisation of Australian redclaw crayfish *Cherax quadricarinatus* sampling gear in southern Africa', *Water SA* 47, 380–384. https://hdl.handle.net/10520/ejc-waters-v47-n3-a10.<br>Nunes, A.L., Zengeya, T.A., Hoffman, A.C., Measey, G.J. & Weyl, O.L., 2017, 'Distribution and establishment of the alien Australian redclaw crayfish, *Cherax quadricarinatus*, in South Africa and Swaziland', *PeerJ* 5, 1-21. https://doi.org/10.7717/peerj.3135.<br>Peay, S., Johnsen, S.I, Bean, S.W., Dunn, A.M., Sandodden, R. & Edsman, L, 2019, 'Biocide treatment of invasive signal crayfish: Successes, failures and lessons learned', *Diversity* 11, 29. https://doi.org/10.3390/d11030029<br>Stebbing, P., Longshaw, M., Scott, A., 2014, 'Review of methods for the management of nonindigenous crayfish, with particular reference to Great Britain', *Ethology Ecology & Evolution* 26, 204-231. https://doi.org/10.1080/03949370.2014.908326. |

| |
|---|
| Woodford, D.J., Ivey, P., Novoa, A., Shackleton, R., Richardson, D., Weyl, O., Van Wilgen, B. and Zengeya, T., 2017, ‘Managing conflict-generating invasive species in South Africa: Challenges and trade-offs’, *Bothalia* 47, a2160. https://doi. org/10.4102/abc.v47i2.2160. |

| **MAN6 Any other management considerations to highlight?** (if yes, fill in Appendix MAN6) | |
|---|---|
| Response | No |

**5. Calculations**

**Likelihood = Very unlikely**

| Parameter | Likelihood | Stages | Final assessment |
|---|---|---|---|
| LIK1 | 0.0027 | P(entry) = 0.5 | P (invasion) = 0.125 |
| LIK2 | 0.5 | | |
| LIK3 | 0.5 | P(establishment) = 0.5 | |
| LIK4 | 0.5 | | |
| LIK5 | 0.0027 | P (spread) = 0.5 | |
| LIK6 | 0.5 | | |

**Consequence = Major (MR)**

| Parameter | Mechanism/sector | Response |
|---|---|---|
| CON1 | Maximum environmental impact | DD |
| CON2 | Maximum socio-economic impact | DD |
| CON3a | Competition | DD |
| CON3b | Predation | DD |
| CON3c | Hybridisation | DD |
| CON3d | Disease transmission | DD |
| CON3e | Parasitism | DD |
| CON3f | Poisoning/toxicity | DD |
| CON3g | Bio-fouling or other direct physical disturbance | DD |
| CON3h | Grazing/herbivory/browsing | DD |
| CON3i | Chemical, physical, structural impact | DD |
| CON3k | Indirect impacts through interactions with other species | DD |

| | | |
|---|---|---|
| CON3 | Maximum environmental impact (closely related taxa) | MR |
| CON4a | Safety | DD |
| CON4b | Material and immaterial assets | DD |
| CON4c | Health | DD |
| CON4d | Social, spiritual and cultural relations | DD |
| CON4 | Maximum socio-economic impact (closely related taxa) | MO |
| CON5 | Potential impact based on traits, experiments, or models | MR |

**Risk = High**

| | | Consequences | | | | |
|---|---|---|---|---|---|---|
| | | MC | MN | MO | MR | MV |
| Likelihood | Extremely unlikely | low | low | low | medium | medium |
| | Very unlikely | low | low | low | medium | high |
| | Unlikely | low | low | medium | high | high |
| | Fairly probable | medium | medium | high | high | high |
| | Probable | medium | high | high | high | high |

**Ease of management: Medium**

| Parameter | Question | Response |
|---|---|---|
| MAN3a | How accessible are populations? | 2 |
| MAN3b | Is detectability critically time-dependent? | 2 |
| MAN3c | Time to reproduction | 1 |
| MAN3d | Propagule persistence | 0 |
| **MAN3** | **SUM** | 5 |

**Appendix BAC7**: Global alien range of *Orconectes juvenilis*
Map from GBIF: https://www.gbif.org/species/5789975

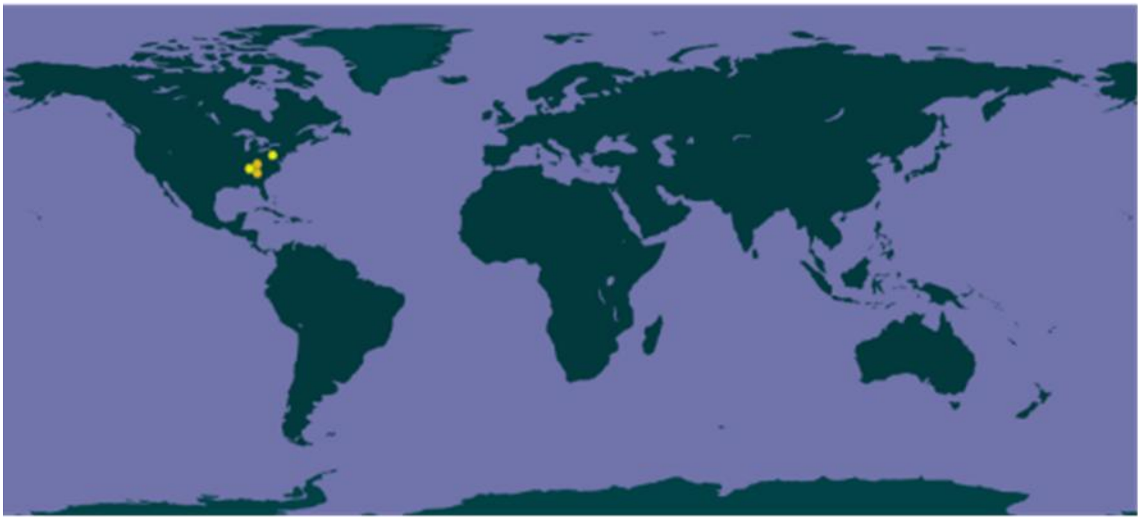

#### 1.7 Risk analysis report for Spinycheek crayfish (*Faxonius limosus*)

***Taxon:*** *Faxonius limosus* (Rafinesque,1817)

***Area:*** South Africa

**Picture of *Taxon***

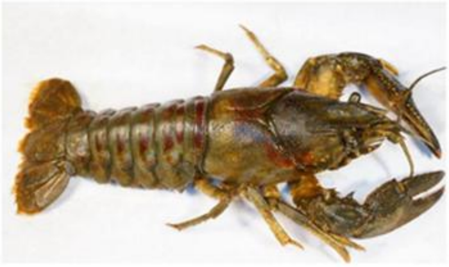

Photo: Alekhnovich A, Buřič M. 2017: NOBANIS – Invasive Alien Species Fact Sheet – *Orconectes limosus*. – From : Online Database of the European Network on Invasive Alien Species – NOBANIS www.nobanis.org

**Alien distribution map**

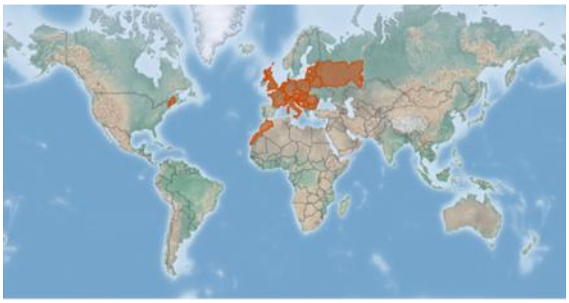

Reference: CABI, 2021. *Faxonius limosus* [original text by Elena Tricarico]. In: Invasive Species Compendium. Wallingford, UK: CAB International. www.cabi.org/isc.

**Risk Assessment summary:**

Spiny-cheek crayfish (*Faxonius limosus*) are native to North America but have been widely distributed in Europe. There are no know populations of *F. limosus* in South Africa but there is anecdotal evidence that it is present in the ornamental trade and this needs to be verified. There are no known populations in neighbouring countries, therefore via unaided pathways is unlikely. The illegal pet trade industry remains a concern because *F. limosus* may be moved around by humans and there is a high risk of release of unwanted pets. *Faxonius limosus* has displaced native crayfish in Europe through the competition for resources and transmission of crayfish plague. The decrease in native population of crayfish in Europe because of *F. limosus* invasions also led to loss of livelihoods for communities' dependent harvesting native crayfish.

**Risk score:**

High

**Management options summary:**

*Faxonius limosus* is a small, largely nocturnal, burrowing freshwater species that is difficult to detect. Several methods can be used to control *F. limosus* such as mechanical removal using traps and electrofishing, biocides, and biological control. However, there are several issues that make the use of these methods challenging. The benefits of *F. limosus* were scored as low.

**Ease of management:**

Medium

**Recommendations:**

*Faxonius limosus* poses a high risk to South Africa. If *F. limosus* were to establish outside of captivity, such populations would likely be very difficult to extirpate.

**Listing under NEM:BA A&IS lists of 2020:**

1a[1]

[1] The current listing is "*Orconectes limosus* (Girard, 1852)", the recommended listing is "*Faxonius limosus* (Girard, 1852)", i.e., it is proposed that the regulatory name be updated to the accepted scientific name of the taxon under assessment.

| | |
|---|---|
| As such it is recommended that *F. limosus* remains listed as category 1a, active efforts made to establish whether the species is present in the country, and that any individuals found are controlled. This would likely require a dedicated assessment of the ornamental trade. A reassessment in five years is thus required and, if absent from the country, *F. limosus s*hould be added to a future prohibited list. | **Recommended listing category:**<br><br>1a[1] |

[1] The presence of *F. limosus* in South Africa should be confirmed and the taxon should be reassessed in five years (i.e., by 2029).

**1. Background**

<table>
<tr><td colspan="3">BAC1 Name of assessor(s)</td></tr>
<tr><td>Name of lead assessor</td><td colspan="2">Lee-Anne Botha</td></tr>
<tr><td>Additional assessor (1)</td><td colspan="2">Tsungai Zengeya</td></tr>
<tr><td>Additional assessor (2)</td><td colspan="2">Christian Chimimba</td></tr>
<tr><td colspan="3">BAC2 Contact details of assessor (s)</td></tr>
<tr><td rowspan="2">Lead assessor</td><td colspan="2">Organisational affiliation: Department of Zoology and Entomology, University of Pretoria and The South African National Biodiversity Institute (SANBI).</td></tr>
<tr><td colspan="2"></td></tr>
<tr><td rowspan="2">Additional assessor (1)</td><td colspan="2">Organisational affiliation: Department of Zoology and Entomology, University of Pretoria and The South African National Biodiversity Institute (SANBI).</td></tr>
<tr><td colspan="2"></td></tr>
<tr><td rowspan="2">Additional assessor (2)</td><td colspan="2">Organisational affiliation: Department of Zoology and Entomology, University of Pretoria and The South African National Biodiversity Institute (SANBI).</td></tr>
<tr><td colspan="2"></td></tr>
<tr><td colspan="3">BAC3 Name(s) and contact details of expert(s) consulted</td></tr>
<tr><td rowspan="3">Expert (1)</td><td colspan="2">Name:</td></tr>
<tr><td colspan="2">email:</td></tr>
<tr><td colspan="2">Phone:</td></tr>
<tr><td rowspan="3">Expert (2)</td><td colspan="2">Name:</td></tr>
<tr><td colspan="2">email:</td></tr>
<tr><td colspan="2">Phone:</td></tr>
<tr><td colspan="3">Comments: none</td></tr>
<tr><td colspan="3">BAC4 Scientific name of Taxon under assessment</td></tr>
<tr><td colspan="2">Taxon name: Faxonius limosus</td><td>Authority: (Rafinesque,1817)</td></tr>
<tr><td colspan="3">Comments: The genus Orconectes previously included cave-dwelling and surface-dwelling crayfish (Crandall & De Grave 2017). The non-cave dwelling crayfish in the Orconectes genus were moved to Faxonius (Crandall & De Grave 2017). The taxonomic status of F. rusticus has been verified (GBIF 2023).</td></tr>
<tr><td colspan="3">References:<br>Crandall, K.A., De Grave, S., 2017, ‘An updated classification of the freshwater crayfishes (Decapoda: Astacidea) of the world, with a complete species list’, Journal of Crustacean Biology, 27, 615–653.<br>Faxonius limosus (Rafinesque, 1817) in GBIF Secretariat (2023). GBIF Backbone Taxonomy. Checklist dataset https://doi.org/10.15468/39omei accessed via GBIF.org on 2024-07-10.</td></tr>
<tr><td colspan="3">BAC5 Synonym(s) considered</td></tr>
<tr><td colspan="3">Synonyms: Cambarus affinis | Cambarus limosus | Orconectes limosus | Astacus limosus</td></tr>
<tr><td colspan="3">Comments: The abovementioned synonyms are known for F. limosus (Crandall and de Grave 2017, GBIF, 2023) and were used in the literature search of this risk analysis.</td></tr>
<tr><td colspan="3">References:<br>Crandall, K.A., De Grave, S., 2017, ‘An updated classification of the freshwater crayfishes (Decapoda: Astacidea) of the world, with a complete species list’, Journal of Crustacean Biology, 27, 615–653. https://doi.org/10.1093/jcbiol/rux070.<br>Faxonius limosus (Rafinesque, 1817) in GBIF Secretariat (2023). GBIF Backbone Taxonomy. Checklist dataset https://doi.org/10.15468/39omei accessed via GBIF.org on 2024-07-10.</td></tr>
<tr><td colspan="3">BAC6 Common name(s) considered</td></tr>
<tr><td colspan="3">Common names: spiny-cheek crayfish</td></tr>
<tr><td colspan="3">Comments: The abovementioned common name is known for F. limosus (GBIF 2023) and were used in the literature search of this risk analysis</td></tr>
</table>

| | |
|---|---|
| References:<br>*Faxonius limosus* (Rafinesque, 1817) in GBIF Secretariat (2023). GBIF Backbone Taxonomy. Checklist dataset https://doi.org/10.15468/39omei accessed via GBIF.org on 2024-07-10. | |
| **BAC7 What is the native range of the *Taxon*?** | |
| Response: **Canada**: New Brunswick \| Quebec<br>**United States**: Connecticut \| Delaware \| District of Columbia \| Maine \| Maryland \| Massachusetts \| New Hampshire \| New Jersey \| New York \| Pennsylvania \| Rhode Island \| Vermont \| Virginia \| West Virginia | Confidence: High |
| Comments: Native to Canada and several states in the USA (Pilotto et al. 2008; Adams et al. 2010) | |
| References:<br>Adams, S., Schuster, G.A., Taylor, C.A., 2010, *Orconectes limosus. The IUCN Red List of Threatened Species 2010*: https://www.iucnredlist.org/species/153764/4541724.<br>Pilotto, F., Free, G., Crosa, G., Sena, F., Ghiani, M. & Cardoso, A.C., 2008, 'The invasive crayfish *Orconectes limosus* in Lake Varese: estimating abundance and population size structure in the context of habitat and methodological constraints', *Journal of Crustacean Biology*, 28, 633–640. https://doi.org/10.1651/07-2967.1 | |
| **BAC8 What is the global alien range of the *Taxon*?** | |
| Response: Austria; Belgium; Czech Republic; France; Germany; Hungary; Italy; Lithuania; Luxembourg; Montenegro; Morocco; Netherlands; Poland; Russian Federation (Kaliningrad); Switzerland; United Kingdom (Great Britain) | Confidence: High |
| Comments: Widely introduced to several countries in Europe (Adams et al. 2010; CABI 2024) | |
| References:<br>Adams, S., Schuster, G.A. & Taylor, C.A., 2010, *Orconectes limosus. The IUCN Red List of Threatened Species 2010*. https://www.iucnredlist.org/species/153764/4541724.<br>CABI, 2024. *Faxonius limosus* (Spiny-cheek crayfish). Invasive Species Compendium, CABI Publishing, Wallingford, UK. https://www. cabi.org/isc/datasheet/72033 (accessed 10 July 2024) | |
| **BAC9 Geographic scope = the *Area* under consideration** | |
| *Area* of assessment: South Africa | |
| Comments: The geographic scope of this assessment is limited to South Africa | |
| **BAC10 Is the *Taxon* present in the *Area*?** | |
| Response: No | Confidence: Low |
| Comments: *Faxonius limosus* is present in the pet trade globally (Chucholl 2013; Faulkes 2015), but there are no records that it is present in South Africa (Nunes et al. 2017). | |
| References:<br>Chucholl, C., 2013, 'Invaders for sale: trade and determinants of introduction of ornamental freshwater crayfish', *Biological Invasions* 15, 125–141. https://doi.org/10.1007/s10530-012-0273-2<br>Faulkes, Z., 2015, 'The global trade in crayfish as pets', *Crustacean Research*, 44, 75–92. https://doi.org/10.18353/crustacea.44.0_75<br>Nunes, A.L., Zengeya, T.A., Measey, G.J. & Weyl, O.L.F., 2017b, 'Freshwater crayfish invasions in South Africa: past, present and potential future', *African Journal of Aquatic Science* 42, 309–323. https://doi.org/10.2989/16085914.2017.1405788 | |
| **BAC11 Availability of physical specimen** | |
| Response: No | Confidence in ID: |
| Herbarium or museum accession number: No *Faxonius limosus* specimens have been found in South Africa and it is assumed the species is not present in South Africa, although it requires further assessment, especially linked to the pet trade industry (Nunes et al. 2017). | |
| References:<br>Nunes, A.L., Zengeya, T.A., Measey, G.J. & Weyl, O.L.F., 2017b, 'Freshwater crayfish invasions in South Africa: past, present and potential future', *African Journal of Aquatic Science* 42, 309–323. https://doi.org/10.2989/16085914.2017.1405788 | |

**BAC12 Is the *Taxon* native to the *Area* or part of the *Area*?**

| | | |
|---|---|---|
| The *Taxon* is native to (part of) the *Area*. | No | Confidence: High |
| The *Taxon* is alien in (part of) the *Area*. | Yes | Confidence: High |

Comments: There are no native freshwater crayfish species native in South Africa (de Moor 2002; Nunes et al. 2017).

References:

de Moor, I., 2002, 'Potential impacts of alien freshwater crayfish in South Africa', *African Journal of Aquatic Science* 27, 125–139. , https://doi.org/10.2989/16085914.2002.9626584.

Nunes AL, Zengeya TA, Measey GJ, Weyl OLF. 2017. Freshwater crayfish invasions in South Africa: past, present and potential future. *African Journal of Aquatic Science* 42: 309–323. https://doi.org/10.2989/16085914.2017.1405788.

**BAC13 What is the *Taxon*'s introduction status in the *Area*?**

| | | |
|---|---|---|
| The *Taxon* is in cultivation/containment. | Don't know | Confidence: Low |
| The *Taxon* is present outside of cultivation/containment. | None | Confidence: Low |
| The *Taxon* has established/naturalised. | None | Confidence: Low |
| The *Taxon* is invasive. | None | Confidence: Low |

Comments: There are no records of *F. limosus* being present in South Africa (de Moor 2002; Nunes et al. 2017). It is present in the pet trade in several countries in Europe (Chucholl 2013; Faulkes 2015) and could likely be traded in South Africa but this needs to be assessed.

References:

Chucholl, C., 2013, 'Invaders for sale: trade and determinants of introduction of ornamental freshwater crayfish', *Biological Invasions* 15, 125–141. https://doi.org/10.1007/s10530-012-0273-2.

de Moor, I., 2002, 'Potential impacts of alien freshwater crayfish in South Africa', *African Journal of Aquatic Science* 27, 125–139, https://doi.org/10.2989/16085914.2002.9626584.

Faulkes, Z., 2015, 'The global trade in crayfish as pets', *Crustacean Research*, 44, 75–92. https://doi.org/10.18353/crustacea.44.0_75.

Nunes, A.L., Zengeya, T.A., Measey, G.J. & Weyl, O.L.F., 2017b, 'Freshwater crayfish invasions in South Africa: past, present and potential future', *African Journal of Aquatic Science* 42, 309–323. https://doi.org/10.2989/16085914.2017.1405788.

**BAC14 Primary (introduction) pathways**

| | | |
|---|---|---|
| Release | None | Confidence: |
| Escape | Aquaculture/bait/Pet/aquarium/terrarium species (including live food for such species) | Confidence: High |
| Contaminant | Live food and live bait | Confidence: High |
| Stowaway | None | Confidence: |
| Corridor | None | Confidence: |
| Unaided | None | Confidence: |

Comments: *Faxonius limosus* has been introduced as live bait, unwanted aquarium species, and as contaminants of fish stocks (Holdich et al. 2006; Holdich & Black 2007; Aquiloni et al. 2010).

References:

Aquiloni, L., Tricarico, E. & Gherardi, F., 2010, 'Crayfish in Italy: distribution, threats and management', *International Aquatic Research*, 2(1) 1-14.

Holdich, D. & Black, J., 2007, 'The spiny-cheek crayfish, *Orconectes limosus* (Rafinesque, 1817) [Crustacea: Decapoda: Cambaridae], digs into the UK', *Aquatic Invasions*, 2(1), 1-16. http://www.aquaticinvasions.ru/2007/AI_2007_2_1_Holdich_Black.pdf

Holdich, D.M., Haffner, P. & Noël, P., 2006, Species files. In: *Atlas of Crayfish in Europe*, [ed. by Souty-Grosset, C., Holdich, D.M., Noël. P.Y., Reynolds. J.D. & Haffner, P.]. Paris, France: Museum national d'Histoire naturelle. 50-129.

#### 2. Likelihood

| **LIK1 Likelihood of entry via unaided primary pathways** | |
|---|---|
| Response: Very unlikely (p =0.0027) | Confidence: Medium |
| Rationale: There is no evidence that *F. limosus* is present in South Africa or has established in the wild in neighbouring countries in the SADC Region (Nunes et al. 2017; Madzivanzira et al. 2020) therefore the probability of *F. limosus* entering South Africa through unaided pathways (connected waterways) is very unlikely (p = 0.0027). | |
| References:<br><br>Madzivanzira, T.C., South, J., Wood, L.E., Nunes, A.L. & Weyl, O.L.F., 2020, 'A review of freshwater crayfish introductions in Africa', *Reviews in Fisheries Science & Aquaculture* 1-21. https://doi.org/10.1080/23308249.2020.1802405.<br>Nunes, A.L., Zengeya, T.A., Measey, G.J. & Weyl, O.L.F., 2017b, 'Freshwater crayfish invasions in South Africa: past, present and potential future', *African Journal of Aquatic Science* 42, 309–323. https://doi.org/10.2989/16085914.2017.1405788 | |

| **LIK2 Likelihood of entry via human aided primary pathways** | |
|---|---|
| Response: Fairly probable (p = 0.5) | Confidence: Low |
| Rationale: *Faxonius limosus* has been introduced as live bait, unwanted aquarium species, and as contaminants of fish stocks (Holdich & Black 2007, Aquiloni et al. 2010). It is likely that it could be present in the pet trade in South Africa, but this needs to be assessed. | |
| References:<br>Aquiloni, L., Tricarico, E. & Gherardi, F., 2010, 'Crayfish in Italy: distribution, threats and management. *International Aquatic Research*, 2(1), 1-14.<br>Holdich, D. & Black, J., 2007, 'The spiny-cheek crayfish, *Orconectes limosus* (Rafinesque, 1817) [Crustacea: Decapoda: Cambaridae], digs into the UK', *Aquatic Invasions*, 2(1), 1-16. http://www.aquaticinvasions.ru/2007/AI_2007_2_1_Holdich_Black.pdf | |

| **LIK3 Habitat suitability** | |
|---|---|
| Response: Fairly probable (p = 0.5) | Confidence: Low |
| Rationale: *Faxonius limosus* is a habitat generalist that inhabits permanent streams and lakes with a range of substrates such as clay, silt, sand and gravel (Hamr 2002; Adams et al. 2010, Kouba et al. 2016). Individuals are often found in lakes, shallow depressions in pools and have rarely been captured where silt is absent from the substrate (Holdich et al. 2009; Adams et al. 2010). Generally, it is very tolerant to a wide range of environmental conditions and is able to cope with polluted canals and organically enriched lakes and ponds (Holdich et al 2009). | |
| References:<br><br>Adams, S., Schuster, G.A. & Taylor, C.A. 2010, *Orconectes limosus. The IUCN Red List of Threatened Species 2010*: https://www.iucnredlist.org/species/153764/4541724.<br>Hamr, P., 2002. *Orconectes*. In: *Biology of freshwater crayfish*, [ed. by Holdich, D. M.]. Oxford, UK: Blackwell Science. 585-608.<br>Holdich, D.M., Reynolds, J.D., & Sibley, P.J., 2009, 'A review of the ever-increasing threat to European crayfish from non-indigenous crayfish species', *Knowledge and Management of Aquatic Ecosystems* 11, 394–395. https://doi.org/10.1051/kmae/2009025. | |

| |
|---|
| Kouba, A., Tíkal, J., Císar, P., Veselý. L., Fort, M., Príborský, J., Patoka, J., Buřič, M., 2016, 'The significance of droughts for hyporheic dwellers: evidence from freshwater crayfish', *Scientific Reports*, 6265-269. https://doi.org/10.1038/srep26569. |

| **LIK4 Climate suitability** | |
|---|---|
| Response: Fairly probable (p = 0.5) | Confidence: Low |
| Rationale: *Faxonius limosus* is known to prefer a temperate mesothermal climate were the average temperature of coldest month > 0°C and < 18°C and the mean warmest mouth is > 10°C. it also prefers a continental microthermal climate where the average temperatures in the coldest month are <0°C and in the mean warmest month are >10°C (CABI 2024). These climatic conditions are available in South Africa. | |
| References:<br><br>CABI, 2024. *Faxonius limosus* (Spiny-cheek crayfish). Invasive Species Compendium, CABI Publishing, Wallingford, UK. https://www. cabi.org/isc/datasheet/72033 (accessed 10 July 2024) | |

| **LIK5 Unaided secondary (dispersal) pathways** | |
|---|---|
| Response: Very unlikely (p = 0.0027) | Confidence: Low |
| Rationale: *Faxinous limosus* can disperse naturally in aquatic environments but has limited ability to disperse overland (Holdich et al 2009; Soes & Koese 2010). There are no known populations of *F. limosus* present in countries neighbouring South Africa that could lead to secondary dispersal events through connected waterways (Nunes et al. 2017; Madzivanzira et al. 2020). | |
| References:<br>Holdich, D.M., Reynolds, J.D., & Sibley, P.J., 2009, 'A review of the ever-increasing threat to European crayfish from non-indigenous crayfish species', *Knowledge and Management of Aquatic Ecosystems* 11, 394–395. https://doi.org/10.1051/kmae/2009025.<br>Madzivanzira, T.C., South, J., Wood, L.E., Nunes, A.L. & Weyl, O.L.F., 2020, 'A review of freshwater crayfish introductions in Africa', *Reviews in Fisheries Science & Aquaculture* 1-21. https://doi.org/10.1080/23308249.2020.1802405<br>Nunes, A.L., Zengeya, T.A., Measey, G.J. & Weyl, O.L.F., 2017, 'Freshwater crayfish invasions in South Africa: past, present and potential future', *African Journal of Aquatic Science* 42, 309–323. https://doi.org/10.2989/16085914.2017.1405788.<br>Soes, D.M. & Koese, B., 2010, 'Invasive crayfish in the Netherlands: A preliminary risk analysis. *Interim report, Bureau Waardenburg bv, Stichting EIS-Nederland'*, Invasive Alien Species Team, Waardenburg | |

| **LIK6 Human aided secondary (dispersal) pathways** | |
|---|---|
| Response: Fairly probable (p = 0.5) | Confidence: Low |
| Rationale:<br><br>Intentional release into the wild through the pet/ aquarium/ terrarium species and live bait pathways remains a concern. In areas of introduction, *Faxonius limosus* has been released into the wild by humans as live bait, unwanted aquarium species, and as contaminants of fish stocks (Holdich & Black 2007, Aquiloni et al. 2010). It is likely that it could be present in the pet trade in South Africa, but this needs to be assessed. | |
| References: | |

Aquiloni, L., Tricarico, E. & Gherardi, F., 2010, 'Crayfish in Italy: distribution, threats and management', *International Aquatic Research*, 2(1), 1-14.
Holdich, D. & Black, J., 2007, 'The spiny-cheek crayfish, *Orconectes limosus* (Rafinesque, 1817) [Crustacea: Decapoda: Cambaridae], digs into the UK', *Aquatic Invasions*, 2(1), 1-16. http://www.aquaticinvasions.ru/2007/AI_2007_2_1_Holdich_Black.pdf

**3. Consequences**

| **CON1 Environmental impact** | |
|---|---|
| **CON1a: Competition** | |
| Response: Major (MR) | Confidence: Medium |
| Rationale: *Faxonius limosus* has displaced native crayfish in Europe through the competition for resources (Holdich et al., 2006; Holdich and Black, 2007; Kozák et al., 2007; Musil et al., 2010; Pârvulescu et al., 2012). For example, in Hungary *F. limosus* is now the most abundant crayfish after supplanting the native crayfish *Astacus astacus* (Holdich et al., 2006). | |
| References:<br>Holdich, D.M., Haffner. P. & Noël, P., 2006, Species files. In: *Atlas of Crayfish in Europe*, [ed. by Souty-Grosset, C., Holdich, D.M., Noël, P.Y., Reynolds, J.D. & Haffner, P.]. Paris, France: Museum national d'Histoire naturelle. 50-129.<br>Holdich, D. & Black, J., 2007, 'The spiny-cheek crayfish, *Orconectes limosus* (Rafinesque, 1817) [Crustacea: Decapoda: Cambaridae], digs into the UK', *Aquatic Invasions*, 2(1), 1-16. http://www.aquaticinvasions.ru/2007/AI_2007_2_1_Holdich_Black.pdf<br>Kozák, P., Buřič, M., Policar, T., Hamáčková, J. & Lepičová, A., 2007, 'The effect of inter- and intra-specific competition on survival and growth rate of native juvenile noble crayfish *Astacus astacus* and alien spiny-cheek crayfish *Orconectes limosus*', *Hydrobiologia*, 59085-94. https://doi.org/10.1007/s10750-007-0760-0.<br>Musil, M., Buřič, M., Policar, T., Kouba, A. & Kozák, P., 2010, 'Comparison of diurnal and nocturnal activity between noble crayfish (*Astacus astacus*) and spinycheek crayfish (*Orconectes limosus*)', *Freshwater Crayfish*, 17189-193.<br>Pârvulescu, L., Schrimpf, A., Kozubíková, E., Resino, S. C., Vrålstad, T., Petrusek, A. & Schulz, R., 2012, Invasive crayfish and crayfish plague on the move: first detection of the plague agent *Aphanomyces astaci* in the Romanian Danube', *Diseases of Aquatic Organisms*, 98(1), 85-94. https://doi.org/10.3354/dao02432. | |
| **CON1b: Predation** | |
| Response: Moderate (MO) | Confidence: Low |
| Rationale: *Faxinous limosus* is omnivorous species, it is likely to cause a decrease in macrophyte cover, macroinvertebrates abundance and diversity, changing trophic interaction and community composition (Vojkovská et al. 2014; Šidagytė et al. 2017). | |
| References:<br>Šidagytė, E., Razlutskij, V., Alekhnovich, A., Rybakovas, A., Moroz, M., Šniaukštaitė, V., Gintaitas, V. & Arbačiauskas, K., 2017, 'Predatory diet and potential effects of *Orconectes limosus* on river macroinvertebrate assemblages of the southeastern Baltic Sea basin: implications for ecological assessment', *Aquatic Invasions*, 12(4). https://doi.org/10.3391/ai.2017.12.4.09.<br>Vojkovská, R., Horká, I. & Ďuriš, Z., 2014, 'The diet of the spiny-cheek crayfish *Orconectes limosus* in the Czech Republic', *Central European Journal of Biology*, 9(1), 58-69. http://rd.springer.com/article/10.2478/s11535-013-0189-y | |

| **CON1c: Hybridisation** | |
|---|---|
| Response: Data Deficient (DD) | Confidence: High |
| Rationale: Hybridisation is unlikely in South Africa because there are no native freshwater crayfish species (Nunes et al. 2017) | |
| References:<br><br>Nunes, A.L., Zengeya, T.A., Measey, G.J. & Weyl, O.L.F., 2017, 'Freshwater crayfish invasions in South Africa: past, present and potential future', *African Journal of Aquatic Science* 42, 309–323. https://doi.org/10.2989/16085914.2017.1405788. | |
| **CON1d: Transmission of disease** | |
| Response: Major (MR) | Confidence: Medium |
| Rationale: *Faxonius limosus* has displaced native crayfish in Europe through the competition for resources and transmission of crayfish plague (Holdich et al., 2006; Holdich and Black, 2007; Kozák et al., 2007; Musil et al., 2010; Pârvulescu et al., 2012). There are no native crayfish species in South Africa, but crayfish plague pathogen has been shown to infect other freshwater decapods such as native potamonautid crabs (Svoboda et al. 2014) | |
| References:<br>Holdich, D.M., Haffner, P. & Noël, P., 2006, Species files. In: *Atlas of Crayfish in Europe*, [ed. by Souty-Grosset, C., Holdich, D.M., Noël, P.Y., Reynolds, J.D. & Haffner, P.]. Paris, France: Museum national d'Histoire naturelle. 50-129.<br>Holdich, D. & Black, J., 2007, 'The spiny-cheek crayfish, *Orconectes limosus* (Rafinesque, 1817) [Crustacea: Decapoda: Cambaridae], digs into the UK', *Aquatic Invasions*, 2(1), 1-16. http://www.aquaticinvasions.ru/2007/AI_2007_2_1_Holdich_Black.pdf<br>Kozák, P., Buřič, M., Policar, T., Hamáčková, J., Lepičová, A., 2007, 'The effect of inter- and intra-specific competition on survival and growth rate of native juvenile noble crayfish *Astacus astacus* and alien spiny-cheek crayfish *Orconectes limosus*', *Hydrobiologia*, 59085-94. https://doi.org/10.1007/s10750-007-0760-0.<br>Musil, M., Buřič, M., Policar, T., Kouba, A. & Kozák. P., 2010, 'Comparison of diurnal and nocturnal activity between noble crayfish (*Astacus astacus*) and spinycheek crayfish (*Orconectes limosus*)', *Freshwater Crayfish*, 17189-193.<br>Pârvulescu, L., Schrimpf, A., Kozubíková, E., Resino, S. C., Vrålstad, T., Petrusek, A. & Schulz, R., 2012, Invasive crayfish and crayfish plague on the move: first detection of the plague agent *Aphanomyces astaci* in the Romanian Danube', *Diseases of Aquatic Organisms*, 98(1), 85-94. https://doi.org/10.3354/dao02432.<br>Svoboda, J., Strand, D.A., Vrålstad, T., Grandjean, F., Edsman, L., Kozák, P., et al., 2014, 'The crayfish plague pathogen can infect freshwater–inhabiting crabs', *Freshwater Biology* 59, 918–929. https://doi–org.uplib.idm.oclc.org/10.1111/fwb.12315. | |
| **CON1e: Parasitism** | |
| Response: Data Deficient (DD) | Confidence: High |
| Rationale: *Faxonius limosus* is not a parasitic taxon (Hamr 2002) | |
| References:<br><br>Hamr, P., 2002. *Orconectes*. In: *Biology of freshwater crayfish*, [ed. by Holdich, D. M.]. Oxford, UK: Blackwell Science. 585-608. | |

| **CON1f: Poisoning/toxicity** | |
|---|---|
| Response: Data Deficient (DD) | Confidence: |
| Rationale: No documented information available to assess the level of impact. | |
| References: | |
| **CON1g: Bio-fouling or other direct physical disturbance** | |
| Response: Data Deficient (DD) | Confidence: |
| Rationale: No documented information available to assess the level of impact. | |
| References: | |
| **CON1h: Grazing/herbivory/browsing** | |
| Response: Minor (MN) | Confidence: Medium |
| Rationale: *Faxinous limosus is* omnivorous species, it is likely to cause a decrease in macrophyte cover, macroinvertebrates abundance and diversity, changing trophic interaction and community composition (Vojkovská et al. 2014; Šidagytė et al. 2017) | |
| References:<br>Šidagytė, E., Razlutskij, V., Alekhnovich, A., Rybakovas, A., Moroz, M., Šniaukštaitė, V., Gintaitas, V. & Arbačiauskas, K., 2017, 'Predatory diet and potential effects of *Orconectes limosus* on river macroinvertebrate assemblages of the southeastern Baltic Sea basin: implications for ecological assessment', *Aquatic Invasions*, 12(4).<br>Vojkovská, R., Horká, I. & Ďuriš, Z., 2014, 'The diet of the spiny-cheek crayfish *Orconectes limosus* in the Czech Republic', *Central European Journal of Biology*, 9(1) 58-69. http://rd.springer.com/article/10.2478/s11535-013-0189-y | |
| **CON1i: Chemical, physical or structural impact on ecosystem** | |
| Response: Minor (MN) | Confidence: Low |
| Rationale: Bioturbation by *F. limosus* has been shown to change the structure of bottom substrate and this can lead to changes in the abundance and structure of benthic communities (Statzner et al. 2000). Burrowing behaviour may destabilize riverbanks, increase erosion and damage nearby infrastructure (Aldridge 2011) | |
| References:<br>Statzner, B., Fièvet, E., Champagne, J. Y., Morel, R. & Herouin, E., 2000, 'Crayfish as geomorphic agents and ecosystem engineers: biological behavior affects sand and gravel erosion in experimental streams', *Limnology and Oceanography*, 45(5), 1030-1040. https://doi.org/10.4319/lo.2000.45.5.1030<br>Aldridge, D., 2011. Spinycheek Crayfish, *Orconectes limosus*. Sand Hutton, UK: GB Non-native Species Secretariat. http://www.nonnativespecies.org/factsheet/factsheet.cfm?speciesId=2441 | |
| **CON1k: Indirect impacts through interactions with other species** | |
| Response: Data Deficient (DD) | Confidence: |
| Rationale: No documented information available to assess the level of impact. | |
| References: | |
| **CON1 Maximum environmental impact** | |

| Response: Major (MR) | Confidence: Medium |
|---|---|
| Rationale: *Faxonius limosus* has displaced native crayfish in Europe through the competition for resources and transmission of crayfish plague (Holdich et al., 2006; Holdich & Black, 2007; Kozák et al., 2007; Musil et al., 2010; Pârvulescu et al., 2012). | |

References:

Holdich, D.M., Haffner, P. & Noël, P., 2006, Species files. In: *Atlas of Crayfish in Europe*, [ed. by Souty-Grosset, C., Holdich. D.M., Noël, P.Y., Reynolds, J.D. & Haffner, P]. Paris, France: Museum national d'Histoire naturelle. 50-129.

Holdich, D. & Black, J., 2007, 'The spiny-cheek crayfish, *Orconectes limosus* (Rafinesque, 1817) [Crustacea: Decapoda: Cambaridae], digs into the UK', *Aquatic Invasions*, 2(1), 1-16. http://www.aquaticinvasions.ru/2007/AI_2007_2_1_Holdich_Black.pdf

Kozák, P., Buřič, M., Policar, T., Hamáčková, J., Lepičová, A., 2007, 'The effect of inter- and intra-specific competition on survival and growth rate of native juvenile noble crayfish *Astacus astacus* and alien spiny-cheek crayfish *Orconectes limosus*', *Hydrobiologia*, 59085-94. https://doi.org/10.1007/s10750-007-0760-0.

Musil, M., Buřič, M., Policar, T., Kouba, A. & Kozák, P., 2010, 'Comparison of diurnal and nocturnal activity between noble crayfish (*Astacus astacus*) and spinycheek crayfish (*Orconectes limosus*)', *Freshwater Crayfish*, 17189-193.

Pârvulescu, L., Schrimpf, A., Kozubíková, E., Resino, S. C., Vrålstad, T., Petrusek, A., Schulz, R., 2012, 'Invasive crayfish and crayfish plague on the move: first detection of the plague agent *Aphanomyces astaci* in the Romanian Danube', *Diseases of Aquatic Organisms*, 98(1), 85-94. https://doi.org/10.3354/dao02432.

| **CON2 Socio-economic impact** | |
|---|---|
| **CON2a: Safety** | |
| Response: Data Deficient (DD) | Confidence: |
| Rationale: No documented information available to assess the level of impact. | |
| References: | |
| **CON2b: Material and immaterial assets** | |
| Response: Moderate (MO) | Confidence: Medium |
| Rationale: The decrease in native population of crayfish in Europe because of *F. limosus* invasions led to economic losses to the crayfish industry dependent on the native crayfish (Laurent 1998). In addition, the burrowing activities of *F. limosus* can cause structural damage to riverbanks, increase bank erosion and damage nearby infrastructure (Aldridge 2011). *Faxonius limosus* can also cause disruptions to recreational activities by reducing sport fish populations though egg predation and/or competition with juveniles (Souty-Grosset et al. 2006; Aldridge 2011). | |

References:

Laurent, P.J., 1988, '*Austropotamobius pallipes* and *A. torrentium*, with observations on their interactions with other species in Europe. In: *Biology of freshwater crayfish: biology, management and exploitation*, [ed. by Holdich, D.M., Lowery, R.S]. London, UK: Croom Helm. 341-364.

Aldridge D., 2011. Spinycheek Crayfish, *Orconectes limosus*. Sand Hutton, UK: GB Non-native Species Secretariat. http://www.nonnativespecies.org/factsheet/factsheet.cfm?speciesId=2441

Souty-Grosset, C., Holdich, D. M., Noel, P. Y., Reynolds, J. D. & Haffner, P. (ed), 2006. *Atlas of crayfish in Europe*, Paris, France: Muséum national d'Histoire naturelle. 187 pp.

| CON2c: Health | |
|---|---|
| Response: Data Deficient (DD) | Confidence: Medium |
| Rationale: No documented information available to assess the level of impact. | |
| References: | |
| **CON2d: Social, spiritual and cultural relations** | |
| Response: Minor (MN) | Confidence: Low |
| Rationale: The decrease in native population of crayfish in Europe because of *F. limosus* invasions led to loss of livelihoods for communities dependent harvesting native crayfish (Laurent 1988; Souty-Grosset et al. 2006; Aldridge 2011). | |
| References:<br>Laurent, P.J., 1988, '*Austropotamobius pallipes* and *A. torrentium*, with observations on their interactions with other species in Europe. In: *Biology of freshwater crayfish: biology, management and exploitation*, [ed. by Holdich, D.M., Lowery, R.S]. London, UK: Croom Helm. 341-364.<br>Aldridge D., 2011. Spinycheek Crayfish, *Orconectes limosus*. Sand Hutton, UK: GB Non-native Species Secretariat. http://www.nonnativespecies.org/factsheet/factsheet.cfm?speciesId=2441.<br>Souty-Grosset, C., Holdich, D. M., Noel, P. Y., Reynolds, J. D. & Haffner, P. (ed), 2006. *Atlas of crayfish in Europe*, Paris, France: Muséum national d'Histoire naturelle. 187 pp. | |
| **CON2 Maximum socio-economic impact** | |
| Response: Moderate (MO) | Confidence: Medium |
| Rationale: *Faxonius limosus* has caused a decrease in native population of crayfish in Europe leading to economic losses to the crayfish industry (Laurent 1998). | |
| References:<br><br>Laurent, P.J., 1988, '*Austropotamobius pallipes* and *A. torrentium*, with observations on their interactions with other species in Europe', In: *Biology of freshwater crayfish: biology, management and exploitation*, [ed. by Holdich, D.M. & Lowery, R.S]. London, UK: Croom Helm. 341-364. | |

| CON3 Closely related species' environmental impact | |
|---|---|
| Response: Not scored as CON1 scored | Confidence: |
| Rationale: There is information to score CON1, so CON3 does not need to be scored as per the framework. | |
| References: | |

| CON4 Closely related species' socio-economic impact | |
|---|---|
| Response: Not scored as CON2 scored | Confidence: |
| Rationale: There is information to score CON2, so CON4 does not need to be scored as per the framework. | |
| References: | |

| **CON5 Potential impact** | |
|---|---|
| Response: MR | Confidence: High |
| Rationale: *Faxonius limosus* may outcompete native freshwater crab species in South Africa, displacing it in freshwater ecosystems (Jackson et al. 2016). Being a functional omnivore, this species may also have detrimental impacts on macroinvertebrates and macrophyte communities in areas where invaded (Klocker and Strayer 2004; Roth et al. 2006; Kreps et al 2012). They can also influence occurrence and species composition of fish communities (Keller et al. 2008). In addition, being a vector for the crayfish plague is another cause of concern (Lodge et al. 2012). This disease may be transferable and could be detrimental to native freshwater crustaceans in South Africa. | |
| References:<br><br>Keller, R.P., Frang, K. & Lodge, D.M., 2008, 'Preventing the spread of invasive species: Economic benefits of intervention guided by ecological predictions', *Conservation Biology*, 22, 80–88. https://doi.org/10.1111/j.1523-1739.2007.00811.x<br>Klocker, C.A. & Strayer, D.L., 2004, 'Interactions among an invasive crayfish (*Orconectes rusticus*), a native crayfish (*Orconectes limosus*), and native bivalves (Sphaeriidae and Unionidae)', *Northeastern Naturalist*, 11, 167–178. https://doi.org/10.1656/1092-6194(2004)011[0167:IAAICO]2.0.CO;2<br>Kreps, T.A., Baldridge, A.K. & Lodge, D.M., 2012, 'The impact of an invasive predator (*Orconectes rusticus*) on freshwater snail communities: Insights on habitat-specific effects from a multilake long-term study', *Canadian Journal of Fisheries and Aquatic Sciences*, 69, 1164–1173. https://doi.org/10.1139/f2012-052<br>Lodge, D.M., Deines, A., Gherardi, F., Yeo, D.C.J., Arcella, T., Baldridge, A.K., Branes, M.A., Chadderton, W.L., Feder, J.L., Grantz, C.A., Howard, G.W., Jerdde, C.L., Peters, J.A., Sargent, L.W., Turner, C.R., Wittmann, M.E. & Zeng, Y. 2012, 'Global introductions of crayfishes: evaluating the impact of species invasions on ecosystem services', *Annual Review of Ecology, Evolution and Systematics*, 43, 449–72. https://doi.org/10.1146/annurev-ecolsys-111511-103919.<br>Roth, B.M., Hein, C.L. & Vander Zanden, M.J., 2006, 'Using bioenergetics and stable isotopes to assess the trophic role of rusty crayfish (*Orconectes rusticus*) in lake littoral zones', *Canadian Journal of Fisheries and Aquatic Sciences*, 63, 335–344. https://doi.org/10.1139/f05-217 | |

#### 4. Management

| **MAN1 What is the feasibility to stop future immigration?** | |
|---|---|
| Response: High | Confidence: low |
| Rationale: *Faxonius limosus* is absent in neighbouring countries, so unaided immigration is unlikely. Moreover, it should be possible to regulate intentional introductions through legislative regulations and having an effective border surveillance and monitoring strategy (Zengeya & Wilson 2020; Wilson & Kumschick 2024). | |
| References:<br><br>Zengeya, T.A. & Wilson, J.R., (eds.) 2020, *The status of biological invasions and their management in South Africa in 2019*, pp.71, South African National Biodiversity Institute, Kirstenbosch and DSI-NRF Centre of Excellence for Invasion Biology, Stellenbosch. http://dx.doi.org/10.5281/zenodo.3947613.<br>Wilson, J. R., & Kumschick, S., 2024, 'The regulation of alien species in South Africa', *South African Journal of Science*, 120(5/6). https://doi.org/10.17159/sajs.2024/17002. | |

| **MAN2 Benefits of the Taxon** |
|---|
| **MAN2a Socio-economic benefits of the *Taxon*** |

| | |
|---|---|
| Response: Low | Confidence: Low |

Rationale: *Faxonius limosus* has been used as live bait and the aquarium trade (Holdich et al. 2006; Holdich & Black 2007; Aquiloni et al. 2010). No socio-economic benefits have been recorded to date in South Africa and even if *F. limosus* is present the benefits are likely to be low (Madzivanzira et al. 2021).

References:

Aquiloni, L., Tricarico, E. & Gherardi, F., 2010, 'Crayfish in Italy: distribution, threats and management, *International Aquatic Research*, 2(1), 1-14. http://www.intelaquares.com/doc/1b.pdf

Holdich, D. & Black, J., 2007, 'The spiny-cheek crayfish, *Orconectes limosus* (Rafinesque, 1817) [Crustacea: Decapoda: Cambaridae], digs into the UK', *Aquatic Invasions*, 2(1), 1-16. http://www.aquaticinvasions.ru/2007/AI_2007_2_1_Holdich_Black.pdf

Holdich, D.M., Haffner, P. & Noël, P., 2006, Species files. In: *Atlas of Crayfish in Europe*, [ed. by Souty-Grosset, C., Holdich, D.M., Noël, P.Y., Reynolds, J.D., Haffner, P]. Paris, France: Museum national d'Histoire naturelle. 50-129.

Madzivanzira, T.C., South, J., Wood, L.E., Nunes, A.L. & Weyl, O.L.F., 2020, 'A review of freshwater crayfish introductions in Africa', *Reviews in Fisheries Science & Aquaculture*, 1-21. https://doi.org/10.1080/23308249.2020.1802405.

**MAN2b Environmental benefits of the *Taxon***

| | |
|---|---|
| Response: Low | Confidence: High |

Rationale: There are no recorded environmental benefits of *Faxonius limosus.*

References:

**MAN3 Ease of management**

**MAN3a How accessible are populations?**

| | |
|---|---|
| Response: 2 (difficult to access) | Confidence: Low |

Rationale: There are no locality records for *Faxonius limosus* in South Africa (Madzivanzira et al. 2020; Nunes et al. 2017). However, given that it is an aquatic species, accessibility is likely to be difficult.

References:

Madzivanzira, T.C., South, J., Wood, L.E., Nunes, A.L. & Weyl, O.L.F., 2020, 'A review of freshwater crayfish introductions in Africa', *Reviews in Fisheries Science & Aquaculture*, 1-21. https://doi.org/10.1080/23308249.2020.1802405.

Nunes, A.L., Zengeya, T.A., Measey, G.J. & Weyl, O.L.F., 2017, 'Freshwater crayfish invasions in South Africa: past, present and potential future', *African Journal of Aquatic Science* 42, 309–323. https://doi.org/10.2989/16085914.2017.1405788.

**MAN3b Is detectability critically time-dependent?**

| | |
|---|---|
| Response: 2 (yes) | Confidence: High |

Rationale: *Faxonius limosus* is a small, largely nocturnal freshwater species that is difficult to detect (Hamr 2002).

References:

| |
|---|
| Hamr, P., 2002. *Orconectes*. In: *Biology of freshwater crayfish*, [ed. by Holdich, D.M.]. Oxford, UK: Blackwell Science. 585-608. |

| | |
|---|---|
| **MAN3c Time to reproduction** | |
| Response: 1 (1-3 years) | Confidence: High |
| Rationale: *Faxonius limosus* reach sexual maturity in one-two years (Hamr 2002; Alekhnovich & Buřič 2017). | |
| References:<br>Alekhnovich, A. & Buřič, M., 2017, NOBANIS – Invasive Alien Species Fact Sheet: *Orconectes limosus*. European Network on Invasive Alien Species (NOBANIS). https://www.nobanis.org/globalassets/speciesinfo/o/orconectes-limosus/orconectes-limosus.pdf<br>Hamr, P., 2002. *Orconectes*. In: *Biology of freshwater crayfish*, [ed. by Holdich, D. M.]. Oxford, UK: Blackwell Science. 585-608. | |
| **MAN3d Propagule persistence** | |
| Response: 0 (< 1 year) | Confidence: High |
| Rationale: *Faxonius limosus* are highly fecund, a female produces up to 440 eggs, and with some parental care of eggs and hatchlings (Hamr 2002; Alekhnovich & Buřič 2017). Females can reproduce by facultative parthenogenesis and are able store sperm (Buřič et al., 2011, 2013). | |
| References:<br>Alekhnovich, A., Buřič, M., 2017, NOBANIS – Invasive Alien Species Fact Sheet: *Orconectes limosus*. European Network on Invasive Alien Species (NOBANIS). https://www.nobanis.org/globalassets/speciesinfo/o/orconectes-limosus/orconectes-limosus.pdf<br>Buřič, M., Hulák, M., Kouba, A., Petrusek, A., & Kozák, P., 2011, 'A successful crayfish invader is capable of facultative parthenogenesis: a novel reproductive mode in decapod crustaceans', *PLoS ONE*, 6e20281. https://doi.org/10.1371/journal.pone.0077597<br>Hamr, P., 2002. *Orconectes*. In: *Biology of freshwater crayfish*, [ed. by Holdich, D. M.]. Oxford, UK: Blackwell Science. 585-608. | |
| **MAN3 Ease of management** | |
| Response: Medium | Confidence: High |
| Rationale: *Faxonius limosus* is a small, largely nocturnal freshwater species that is difficult to detect, it reaches sexual maturity in one-two years, is highly fecund (up to 440 eggs) and can reproduce parthenogenetically (Hamr 2002). | |
| References:<br>Hamr, P., 2002. *Orconectes*. In: *Biology of freshwater crayfish*, [ed. by Holdich, D.M.]. Oxford, UK: Blackwell Science. 585-608. | |

| | |
|---|---|
| **MAN4 Has the feasibility of eradication been evaluated?** | |
| Response: No | Confidence: Low |

| |
|---|
| Rationale: There are no known population of *F. limosus* in South Africa, therefore eradication feasibility has not been no evaluated (Nunes et al. 2017) but see MAN5 for summary of control efforts used to manage invasive crayfish populations. |
| References:<br><br>Nunes, A.L., Zengeya, T.A., Measey, G.J. & Weyl, O.L.F., 2017, 'Freshwater crayfish invasions in South Africa: past, present and potential future', *African Journal of Aquatic Science* 42, 309–323, https://doi.org/10.2989/16085914.2017.1405788. |

| **MAN5 Control options and monitoring approaches available for the *Taxon*** |
|---|
| Response: Several methods can be used to monitor and control invasive crayfish populations including mechanical trapping (e.g., Barkhuizen et al. 2022; Madzivanzira et al. 2021), electrofishing (Nunes et al. 2017), use of biocides, and biological control (Aquiloni et al 2010; Stebbing et al. 2014). However, there are several issues that make management of crayfish invasions challenging (Gherardi et al. 2011). For example, the use of biocides is often effective for localised populations in dams and ponds (Peay et al. 2019) but they are non-specific and need to be applied with caution (Gherardi et al. 2011). Mechanical control methods are also effective in small discrete water bodies but are expensive and require commitment to follow on to complete eradication (Loureiro et al. 2018). In addition, alien crayfish species are regarded as potential conflict-generating species in South Africa and potential management interventions should consider the concerns of stakeholders to avoid conflict (Zengeya et al. 2017). |
| References:<br>Aquiloni, L., Brusconi, S., Cecchinelli, E., Tricarico, E., Mazza, G., Paglianti, A. & Gherardi, F. 2010, 'Biological control of invasive populations of crayfish: the European eel (*Anguilla anguilla*) as a predator of *Procambarus clarkii*', *Biological Invasions* 12 3817-3824. https://doi.org/10.1007/s10530-007-9094-0.<br>Barkhuizen, L.M, Madzivanzira, T.C., South, J., 2022, 'Population ecology of a wild population of red swamp crayfish *Procambarus clarkii* (Girard, 1852) in the Free State Province, South Africa and implications for eradication efforts', *BioInvasions Records* 11, 181–191. https://doi.org/10.3391/bir.2022.<br>Gherardi, F., Aquiloni, L., Diéguez-Uribeondo, J. & Tricarico E., 2011, 'Managing invasive crayfish: is there a hope? *Aquatic Sciences* 73, 185–200. https://doi.org/10.1007/s00027-011-0181-z.<br>Loureiro, T.G, Anastácio, P.M, de Siqueira Bueno, S.L. & Araujo, P.B., 2018, 'Management of invasive populations of the freshwater crayfish *Procambarus clarkii* (Decapoda, Cambaridae): test of a population-control method and proposal of a standard monitoring approach', *Environmental Monitoring Assessment* 190, 559, https://doi.org/10.1007/s10661-018-6942-6.<br>Madzivanzira, T.C., South, J., Nhiwatiwa, T. & Weyl, O.L.F., 2021, 'Standardisation of Australian redclaw crayfish *Cherax quadricarinatus* sampling gear in southern Africa', *Water SA* 47, 380–384. https://hdl.handle.net/10520/ejc-waters-v47-n3-a10.<br>Nunes, A.L., Zengeya, T.A., Hoffman, A.C., Measey, G.J. & Weyl, O.L., 2017, 'Distribution and establishment of the alien Australian redclaw crayfish, *Cherax quadricarinatus*, in South Africa and Swaziland', *PeerJ* 5, 1-21. https://doi.org/10.7717/peerj.3135.<br>Peay, S., Johnsen, S.I, Bean, S.W., Dunn, A.M., Sandodden, R. & Edsman, L, 2019, 'Biocide treatment of invasive signal crayfish: Successes, failures and lessons learned', *Diversity* 11, 29. https://doi.org/10.3390/d11030029<br>Stebbing, P., Longshaw, M., Scott, A., 2014, 'Review of methods for the management of nonindigenous crayfish, with particular reference to Great Britain', *Ethology Ecology & Evolution* 26, 204-231. https://doi.org/10.1080/03949370.2014.908326. |

Woodford, D.J., Ivey, P., Novoa, A., Shackleton, R., Richardson, D., Weyl, O., Van Wilgen, B. and Zengeya, T., 2017, ‘Managing conflict-generating invasive species in South Africa: Challenges and trade-offs’, *Bothalia* 47, a2160. https://doi. org/10.4102/abc.v47i2.2160.

| **MAN6 Any other management considerations to highlight?** | |
|---|---|
| Response | No |

#### 5. Calculations

**Likelihood = Very unlikely**

| **Parameter** | **Likelihood** | **Stages** | **Final assessment** |
|---|---|---|---|
| LIK1 | 0.0027 | P(entry) = 0.5 | P (invasion) = 0.125 |
| LIK2 | 0.5 | | |
| LIK3 | 0.5 | P(establishment) = 0.5 | |
| LIK4 | 0.5 | | |
| LIK5 | 0.0027 | P (spread) = 0.5 | |
| LIK6 | 0.5 | | |

**Consequence = Major (MR)**

| **Parameter** | Mechanism/sector | **Response** |
|---|---|---|
| CON1a | Competition | MR |
| CON1b | Predation | MO |
| CON1c | Hybridisation | DD |
| CON1d | Disease transmission | MR |
| CON1e | Parasitism | DD |
| CON1f | Poisoning/toxicity | DD |
| CON1g | Bio-fouling or other direct physical disturbance | DD |
| CON1h | Grazing/herbivory/browsing | MN |
| CON1i | Chemical, physical, structural impact | MN |
| CON1k | Indirect impacts through interactions with other species | DD |
| **CON1** | Maximum environmental impact | MR |
| CON2a | Safety | DD |
| CON2b | Material and immaterial assets | MO |
| CON2c | Health | DD |
| CON2d | Social, spiritual and cultural relations | MN |
| **CON2** | Maximum socio-economic impact | MO |

| | | |
|---|---|---|
| CON3 | Maximum environmental impact of closely related species | Not scored as CON1 scored |
| CON4 | Maximum socio-economic impact of closely related species | Not scored as CON2 scored |
| **CON5** | Potential impact based on traits, experiments, or models | MR |

**Risk score = High**

| | | Consequences | | | | |
|---|---|---|---|---|---|---|
| | | MC | MN | MO | MR | MV |
| **Likelihood** | Extremely unlikely | low | low | low | medium | medium |
| | Very unlikely | low | low | low | medium | high |
| | Unlikely | low | low | medium | high | high |
| | Fairly probable | medium | medium | high | high | high |
| | Probable | medium | high | high | high | high |

**Ease of management = Medium**

| Parameter | Question | Response |
|---|---|---|
| MAN3a | How accessible are populations? | 2 |
| MAN3b | Is detectability critically time-dependent? | 2 |
| MAN3c | Time to reproduction | 1 |
| MAN3d | Propagule persistence | 0 |
| **MAN3** | **SUM** | 5 |

**Appendix BAC7**:

**Appendix BAC8(a)**: Global alien distribution of *Faxonius limosus*. Source: CABI, 2021. *Faxonius limosus* [original text by Elena Tricarico]. In: Invasive Species Compendium. Wallingford, UK: CAB International. www.cabi.org/isc.

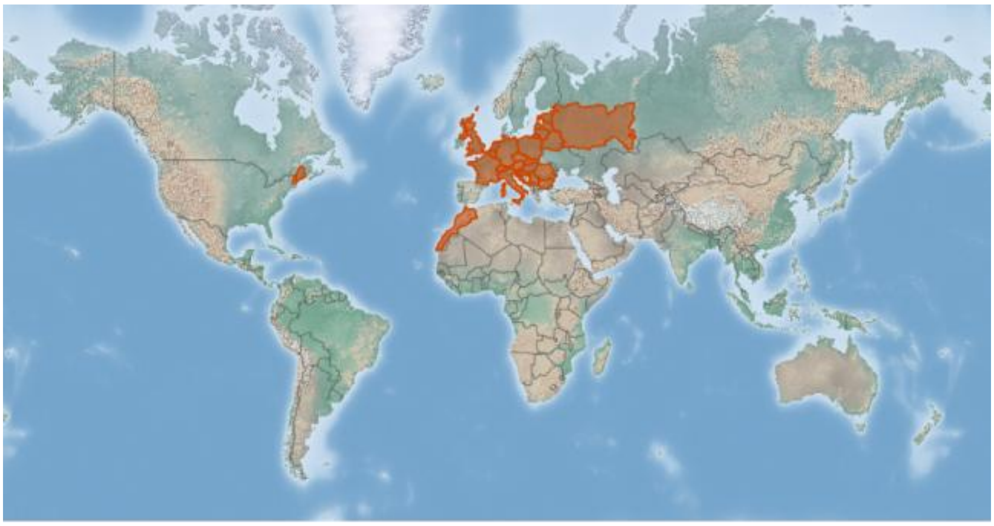

**Appendix BAC8(b)**:

#### 1.8 Risk analysis report for rusty crayfish (*Faxonius rusticus*)

***Risk Analysis Report***

| | |
|---|---|
| ***Taxon:*** *Faxonius rusticus* (Girard, 1852) | ***Area:*** South Africa |
| **Compiled by:** Lee-Anne Botha | **Approved by:** |
| **Picture of *Taxon***<br>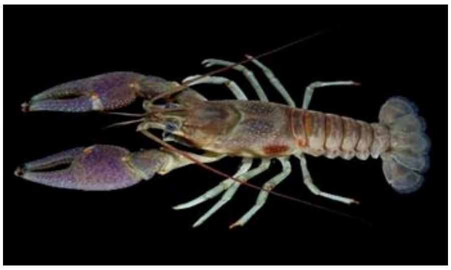<br>Photo: Tennessee Wildlife Resources Agency, NAS - Nonindigenous Aquatic Species. https://nas.er.usgs.gov/queries/FactSheet.aspx?speciesID=214 | **Alien distribution map**<br>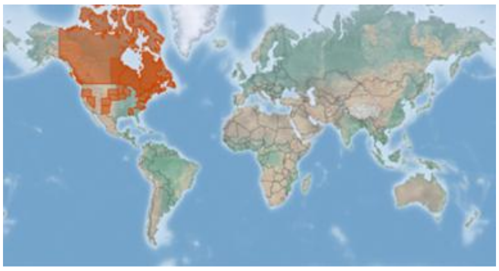<br>Reference: CABI, 2021. *Faxonius rusticus* [original text by Elena Tricarico]. In: Invasive Species Compendium. Wallingford, UK: CAB International. www.cabi.org/isc. |
| **Risk Assessment summary:**<br><br>Rusty crayfish (*Faxonius rusticus*) has been widely introduced outside its native range in the USA and Canada. It has also been established in France and is present in the ornamental trade in Europe. There are no know populations of *F. rusticus* in South Africa but there is anecdotal evidence that it is present in the ornamental trade and this needs to be verified. There are no known populations in neighbouring countries, therefore via unaided pathways is unlikely. The illegal pet trade industry remains a concern because *F. rusticus* may be moved around by humans and there is a high risk of release of unwanted pets. *Faxonius rusticus* is an aggressive competitor and in its introduced range it has been implicated in causing the decline of native freshwater crayfish and fish species through competition and predation. In addition, it is an omnivore and intensive grazing have contributed to a decline in freshwater invertebrate populations and macrophyte communities. *Faxonius rusticus* has also been implicated in the decline of native fish populations that has adversely affecting recreational fishing. | **Risk score:**<br>High |
| **Management options summary:**<br><br>*Faxonius rusticus* is a small, largely nocturnal, non-burrowing freshwater species that is difficult to detect. Several methods have been used to control alien populations of *F. rusticus* such as intensive trapping and fish predation. However, there are several issues that make the use of these methods challenging. The benefits of *F. rusticus* were scored as low. | **Ease of management:**<br><br>Medium |
| **Recommendations:** | **Listing under NEM:BA A&IS lists of 2020:** |

| | |
|---|---|
| *Faxonius rusticus* poses a high risk to South Africa. If *F. rusticus* were to establish outside of captivity, such populations would likely be very difficult to extirpate.<br><br>As such it is recommended that *F. rusticus* remains listed as category 1a, active efforts made to establish whether the species is preset in the country, and that any individuals found are controlled. This would likely require a dedicated assessment of the ornamental trade. A reassessment in five years is thus required and, if absent from the country, *F. rusticus s*hould be added to a future prohibited list. | 1a[1]<br><br>**Recommended listing category:**<br><br>1a[2] |

[1] The current listing is "*Orconectes rusticus* (Girard, 1852)", the recommended listing is "*Faxonius rusticus* (Girard, 1852)", i.e., it is proposed that the regulatory name be updated to the accepted scientific name of the taxon under assessment.
[2] The presence of *F. rusticus* in South Africa should be confirmed and the taxon should be reassessed in five years (i.e., by 2029).

**1. Background**

| | |
|---|---|
| **BAC1 Name of assessor(s)** | |
| Name of lead assessor | Lee-Anne Botha |
| Additional assessor (1) | Tsungai Zengeya |
| Additional assessor (2) | Christian Chimimba |
| **BAC2 Contact details of assessor (s)** | |
| Lead assessor | Organisational affiliation: Department of Zoology and Entomology, University of Pretoria/South African National Biodiversity Institute (SANBI). |
| | |
| Additional assessor (1) | Organisational affiliation: Department of Zoology and Entomology, University of Pretoria/South African National Biodiversity Institute (SANBI). |
| | |
| Additional assessor (2) | Organisational affiliation: Department of Zoology and Entomology, University of Pretoria |
| | |
| | Phone: |
| **BAC3 Name(s) and contact details of expert(s) consulted** | |
| Expert (1) | Name |
| | email: |
| | Phone: |
| Expert (2) | Name: |
| | email: |
| | Phone: |
| Comments: none. | |
| **BAC4 Scientific name of *Taxon* under assessment** | |
| *Taxon* name: *Faxonius rusticus* | Authority: (Girard, 1852) |
| Comments: The genus *Orconectes* previously included cave-dwelling and surface-dwelling crayfish (Crandall & De Grave 2017). The non-cave dwelling crayfish in the *Orconectes* genus were moved to *Faxonius* (Centre for Agriculture and Bioscience International 2019; Crandall & De Grave 2017). The taxonomic status of *F. rusticus* has been verified (Integrated Taxonomic Information System 2022). | |
| References:<br>Centre for Agriculture and Bioscience International (CABI), 2021, '*Faxonius rusticus*', In: Invasive Species Compendium. Wallingford, UK: CAB International. www.cabi.org/isc.<br>Crandall, K.A. & De Grave, S., 2017, 'An updated classification of the freshwater crayfishes (Decapoda: Astacidea) of the world, with a complete species list', *Journal of Crustacean Biology* 27, 615–653. https://doi.org/10.1093/jcbiol/rux070.<br>Integrated Taxonomic Information System (ITIS), 2022, '*Faxonius rusticus* (Girard, 1852)', viewed 13 October 2022, from: https://www.itis.gov/servlet/SingleRpt/SingleRpt?search_topic=TSN&search_value=1133773#null | |
| **BAC5 Synonym(s) considered** | |
| Synonyms: *Orconectes rusticus* | |
| Comments: The genus *Orconectes* previously included cave-dwelling and surface-dwelling crayfish (Crandall & De Grave 2017). The non-cave dwelling crayfish in the *Orconectes* genus were moved to *Faxonius* (Crandall & De Grave 2017). Therefore, the abovementioned synonym was used in the literature search of this risk analysis. | |
| References:<br>Crandall, K.A. & De Grave, S., 2017, 'An updated classification of the freshwater crayfishes (Decapoda: Astacidea) of the world, with a complete species list', *Journal of Crustacean Biology* 27, 615–653. https://doi.org/10.1093/jcbiol/rux070. | |
| **BAC6 Common name(s) considered** | |

| | |
|---|---|
| Common names: Rusty crayfish | |
| Comments: Literature indicates that this is the only common name for *F. rusticus* that was used in the literature search of this risk analysis. | |
| References:<br>Centre for Agriculture and Bioscience International (CABI), 2021, '*Faxonius rusticus*', In: Invasive Species Compendium. Wallingford, UK: CAB International. www.cabi.org/isc. | |
| **BAC7 What is the native range of the *Taxon*?** | |
| Response: United States of America | Confidence: High |
| Comments: The native range of *F. rusticus* comprises the Ohio river basin, spanning tributaries in Western Ohio, Indiana, Kentucky, and Northern Tennessee (Olden et al. 2006; GISD 2015). | |
| References:<br>Global Invasive Species Database (GISD) 2015. Species profile *Orconectes rusticus*. Available from: http://www.iucngisd.org/gisd/species.php?sc=217 [Accessed 03 July2018]<br>Olden, J.D., McCarthy, J.M., Maxted, J.T., Fetzer, W.W. & Van der Zanden, M.J., 2006, 'The rapid spread of rusty crayfish (*Orconectes rusticus*) with observations on native crayfish declines in Wisconsin (USA.) over the past 130 years', *Biological Invasions* 8, 1621–1628. https://doi.org/10.1007/s10530-005-7854-2. | |
| **BAC8 What is the global alien range of the *Taxon*?** | |
| Response: Canada and widespread across several states in the USA | Confidence: High |
| Comments: *Faxonius rusticus* has been translocated outside its native range to 20 states in the USA and there are known introductions in Canada and France (GISD 2015; Lodge et al. 2012; Laffitte et al. 2023). | |
| References:<br>Global Invasive Species Database (GISD) 2015. Species profile *Orconectes rusticus*. Available from: http://www.iucngisd.org/gisd/species.php?sc=217 [Accessed 03 July2018]<br>Lodge, D.M., Deines, A., Gherardi, F., Yeo, D.C.J., Arcella. T, Baldridge, A.K., et al., 2012, 'Global introductions of crayfishes: Evaluating the impact of species invasions on ecosystem services', *Annual Review of Ecology Evolution Systematics* 43, 449–472. https://doi.org/10.1146/annurev-ecolsys-111511-103919<br>Laffitte, M., Baudry, T., Guilmet, M., Andrieu, T., Poulet, N., Duperray, T., Carine, D., Collas, M., Moumen, B. & Grandjean F., 2023,'A new invader in freshwater ecosystems in France: the rusty crayfish *Faxonius rusticus* (Girard, 1852)', *BioInvasions Records,* 12(2), 457–468. https://doi.org/10.3391/bir.2023.12.2.10.<br>https://nas.er.usgs.gov/queries/SpeciesAnimatedMap.aspx?SpeciesID=214 (accessed 22 Sept 2023) | |
| **BAC9 Geographic scope = the *Area* under consideration** | |
| *Area* of assessment: South Africa | |
| Comments: The geographic scope of this assessment is limited to South Africa. | |
| **BAC10 Is the *Taxon* present in the *Area*?** | |
| Response: No | Confidence: Low |
| Comments: There are no records of *F. rusticus* species being in South Africa ; Nunes et al. 2017). *Faxonius rusticus* is present in the pet trade globally ((Lodge et al. 2012; Faulkes 2015), and it might be present in industry in South Africa but this needs to be assessed (Nunes et al. 2017). | |
| References:<br>Faulkes, Z, 2015, 'The global trade in crayfish as pets', *Crustacean Research* 44, 75–92. https://doi.org/10.18353/crustacea.44.0_75.<br>Lodge, D.M., Deines, A., Gherardi, F., Yeo, D.C.J., Arcella. T, Baldridge, A.K., et al., 2012, 'Global introductions of crayfishes: Evaluating the impact of species invasions on ecosystem services', *Annual Review of Ecology Evolution Systematics* 43, 449–472. https://doi.org/10.1146/annurev-ecolsys-111511-103919<br>Nunes, A.L., Zengeya, T.A., Measey, G.J. & Weyl, O.L.F., 2017, 'Freshwater crayfish invasions in South Africa: past, present and potential future', *African Journal of Aquatic Science* 42, 309–323. https://doi.org/10.2989/16085914.2017.1405788. | |
| **BAC11 Availability of physical specimen** | |

| Response: No | Confidence in ID: |
|---|---|

Herbarium or museum accession number: No *Faxonius rusticus* specimens have been found in South Africa and it is assumed the species is not present in South Africa, although it requires further assessment, especially linked to the pet trade industry (Nunes et al. 2017).

References:

Nunes AL, Zengeya TA, Measey GJ, Weyl OLF. 2017. Freshwater crayfish invasions in South Africa: past, present and potential future. *African Journal of Aquatic Science* 42: 309–323, https://doi.org/10.2989/16085914.2017.1405788

**BAC12 Is the *Taxon* native to the *Area* or part of the *Area*?**

| | | |
|---|---|---|
| The *Taxon* is native to (part of) the *Area*. | No | Confidence: High |
| The *Taxon* is alien in (part of) the *Area*. | Yes | Confidence: High |

Comments: There are no native freshwater crayfish species native in South Africa (de Moor 2002; Nunes et al. 2017).

References:

de Moor, I., 2002, 'Potential impacts of alien freshwater crayfish in South Africa', *African Journal of Aquatic Science* 27, 125–139. https://doi.org/10.2989/16085914.2002.9626584.

Nunes, A.L., Zengeya, T.A., Measey, G.J. & Weyl, O.L.F., 2017, 'Freshwater crayfish invasions in South Africa: past, present and potential future', *African Journal of Aquatic Science* 42, 309–323. https://doi.org/10.2989/16085914.2017.1405788.

**BAC13 What is the *Taxon*'s introduction status in the *Area*?**

| | | |
|---|---|---|
| The *Taxon* is in cultivation/containment. | Don't know | Confidence: Low |
| The *Taxon* is present outside of cultivation/containment. | None | Confidence: Low |
| The *Taxon* has established/naturalised. | None | Confidence: Low |
| The *Taxon* is invasive. | None | Confidence: Low |

Comments: There are no records of *F. rusticus* being present in South Africa. It is present in the pet trade in Germany and the USA , and could likely be traded in South Africa but this needs to be assessed (de Moor 2002; Nunes et al. 2017).

References:

de Moor, I., 2002, 'Potential impacts of alien freshwater crayfish in South Africa', *African Journal of Aquatic Science* 27, 125–139. https://doi.org/10.2989/16085914.2002.9626584.

Nunes, A.L., Zengeya, T.A., Measey, G.J. & Weyl, O.L.F., 2017, 'Freshwater crayfish invasions in South Africa: past, present and potential future', *African Journal of Aquatic Science* 42, 309–323. https://doi.org/10.2989/16085914.2017.1405788.

**BAC14 Primary (introduction) pathways**

| | | |
|---|---|---|
| Release | None | Confidence: Medium |
| Escape | Pet/aquarium/terrarium species (including live food for such species) | Confidence: Medium |
| Contaminant | Live food and live bait | Confidence: Medium |
| Stowaway | None | Confidence: Low |
| Corridor | None | Confidence: Low |
| Unaided | None | Confidence: Low |

Comments: *Faxonius rusticus* is known to be present in the pet trade industry in Germany and the USA (Chucholl 2013; Faulkes 2015). In these areas, crayfish have established populations after accidental or intentional release by owners as unwanted pets (Faulkes 2015). *Faxonius rusticus* is furthermore a popular bait species among anglers (Kerr 2014) and is stocked by fishermen that could also lead to the unintentional release of *F. rusticus*.

| |
|---|
| References:<br>Chucholl, C., 2013, 'Invaders for sale: Trade and determinants of introduction of ornamental freshwater crayfish', *Biological Invasions* 15, 125–141. https://doi.org/10.1007/s10530-012-0273-2<br>Faulkes, Z, 2015, 'The global trade in crayfish as pets', *Crustacean Research* 44, 75–92. https://doi.org/10.18353/crustacea.44.0_75.<br>Kerr, S.J., 2014, 'The introduction and spread of aquatic invasive species through the recreational use of bait: A literature review", Report prepared for Biodiversity Branch. Ontario Ministry of Natural Resources. Peterborough', Ontario, Canada. |

**2. Likelihood**

| **LIK1 Likelihood of entry via unaided primary pathways** | |
|---|---|
| Response: Very unlikely (p = 0.0027) | Confidence: Medium |
| Rationale: There is no evidence that *F. rusticus* is present in South Africa or has established in the wild in neighbouring countries in the SADC Region (Lodge et al. 2012; Madzivanzira et al. 2020) therefore the probability of *F. rusticus* entering South Africa through unaided pathways (connected waterways) is very unlikely (p = 0.0027). | |
| References:<br><br>Madzivanzira, T.C., South, J., Wood, L.E., Nunes, A.L. & Weyl, O.L.F., 2020, 'A review of freshwater crayfish introductions in Africa', *Reviews in Fisheries Science & Aquaculture*, 1-21. https://doi.org/10.1080/23308249.2020.1802405.<br>Lodge, D.M., Deines, A., Gherardi, F., Yeo, D.C.J., Arcella. T, Baldridge, A.K., et al., 2012, 'Global introductions of crayfishes: Evaluating the impact of species invasions on ecosystem services', *Annual Review of Ecology Evolution Systematics* 43, 449–472. https://doi.org/10.1146/annurev-ecolsys-111511-103919 | |

| **LIK2 Likelihood of entry via human aided primary pathways** | |
|---|---|
| Response: Fairly probable (p = 0.5) | Confidence: Medium |
| Rationale: *Faxonius rusticus* is popular as a live bait used by anglers for recreational fishing (Olden et al. 2009; Kerr 2014). *Faxonius rusticus* is present in the pet trade industry in Germany and the USA (Faulkes 2015). It is likely that it could be present in the pet trade in South Africa but this needs to be assessed. | |
| References:<br><br>Faulkes, Z, 2015, 'The global trade in crayfish as pets', *Crustacean Research* 44, 75–92. https://doi.org/10.18353/crustacea.44.0_75.<br>Kerr, S.J., 2014, 'The introduction and spread of aquatic invasive species through the recreational use of bait: A literature review", Report prepared for Biodiversity Branch. Ontario Ministry of Natural Resources. Peterborough', Ontario, Canada.<br>Olden, J.D., Adams, J.W. & Larson, E.R., 2009, 'First record of *Orconectes rusticus* (Girard, 1852) (Decapoda, Cambaridae) west of the great continental divide in North America', *Crustaceana* 82, 1347–1351. https://doi.org/10.1163/156854009X448934. | |

| **LIK3 Habitat suitability** | |
|---|---|
| Response: Fairy probable (p = 0.5) | Confidence: Medium |

| |
|---|
| Rationale: *Faxonius rusticus* is a habitat generalist that inhabits permanent streams and lakes with a range of substrates such as clay, silt, sand and gravel (GISD 2015). *Faxonius rusticus* prefers areas that consist of rocks, logs or other debris used to construct shallow excavations underneath (GISD 2015). *Faxonius rusticus* lives in open water during most life stages and burrows only under extreme adverse conditions (GISD 2015). The habitats are present in most aquatic systems across South Africa. |
| References:<br><br>Global Invasive Species Database (GISD) 2015. Species profile *Orconectes rusticus*. Available from: http://www.iucngisd.org/gisd/species.php?sc=217 [Accessed 03 July2018]. |

| **LIK4 Climate suitability** | |
|---|---|
| Response: Fairy probable (p = 0.5) | Confidence: Medium |
| Rationale: Based on the Koppen-Geiger climate classification map, *F. rusticus* predominantly occurs in cold D and arid B regions (Peel et al. 2007). South Africa's main climate type is arid B, therefore, *F. rusticus* is likely to survive in these areas (Peel et al. 2007). *Faxonius rusticus* prefers well-oxygenated water and optimal water temperature is 20-25°C, but can endure a wide range of temperatures (0–39°C) within the native range (GISD 2015). When temperatures exceed 30° C, adults have been observed digging burrows to escape the heat (GISD 2015). | |
| References:<br><br>Global Invasive Species Database (GISD) 2015. Species profile *Orconectes rusticus*. Available from: http://www.iucngisd.org/gisd/species.php?sc=217 [Accessed 03 July 2018]<br>Peel, M.C., Finlayson, B.L. & McMahon, T.A., 2007, 'Updated world map of the Koppen-Geiger climate classification', *Hydrology and Earth System Sciences* 11, 1633–1644. https://doi.org/10.5194/hess-11-1633-2007. | |

| **LIK5 Unaided secondary (dispersal) pathways** | |
|---|---|
| Response: Fairly probable (p = 0.5) | Confidence: Medium |
| Rationale: *Faxinous rusticus* can disperse naturally in aquatic environments but has limited ability to disperse overland (Claussen et al. 2000; Byron & Wilson 2001; Hamr 2002). There are no known populations of *F. rusticus* present in countries neighbouring South Africa that could lead to secondary dispersal events through connected waterways (Madzivanzira et al. 2020). | |
| References:<br>Byron, C.J. & Wilson, K.A., 2001, 'Rusty crayfish (*Orconectes rusticus*) movement within and between habitats in Trout Lake, Vilas County, Wisconsin', *Journal of the North American Benthological Society* 20, 606–614.<br>Hamr, P., 2002. Orconectes. In: *Biology of freshwater crayfish*, [ed. by Holdich, D. M.]. Oxford, UK: Blackwell Science. 585-608.<br>Claussen DL, Hopper RA, Sanker AM, 2000. The effects of temperature, body size, and hydration state on the terrestrial locomotion of the crayfish *Orconectes rusticus*. *Journal of Crustacean Biology*, 20218-223<br>Madzivanzira, T.C., South, J., Wood, L.E., Nunes, A.L. & Weyl, O.L.F., 2020, 'A review of freshwater crayfish introductions in Africa', *Reviews in Fisheries Science & Aquaculture*, 1-21. https://doi.org/10.1080/23308249.2020.1802405. | |

| **LIK6 Human aided secondary (dispersal) pathways** |
|---|

| Response: Fairly probable (p = 0.5) | Confidence: Medium |
|---|---|
| Rationale: Intentional release into the wild through the pet/ aquarium/ terrarium species and live bait pathways remains a concern. In areas of introduction, *Faxonius rusticus* has been released into the wild by humans as unwanted pets and bucket release by anglers into waterways where they are used as bait, dispersing into new areas (Faulkes 2015; Olden et al. 2009). | |
| References:<br>Faulkes, Z., 2015, 'The global trade in crayfish as pets', *Crustacean Research* 44, 75–92. https://doi.org/10.18353/crustacea.44.0_75.<br>Olden, J.D., Adams, J.W. & Larson, E.R., 2009, 'First record of *Orconectes rusticus* (Girard, 1852) (Decapoda, Cambaridae) west of the great continental divide in North America', *Crustaceana* 82, 1347–1351. https://doi.org/10.1163/156854009X448934. | |

**3. Consequences**

| **CON1 Environmental impact** | |
|---|---|
| **CON1a: Competition** | |
| Response: Major (MR) | Confidence: High |
| Rationale: *Faxonius rusticus* invasions have been implicated in the decline and or disappearance of native crayfish mainly through competition for resources. For example, *F. rusticus* has displaced native crayfish (*F. propinquus, F. sanborni* and *F. virilis*) in the invaded range in the USA through direct competition for food and shelter (Lodge et al. 1995). This competitive displacement of congeners is facilitated by adaptive life history traits such as earlier recruitment, higher fecundity, faster growth rate, relatively larger body sizes, interference reproduction, and wider habitat preferences (Butler and Stein 1985; Hobbs et al. 1989; Hill and Lodge 1994; Garvey & Stein 1993; Klocker & Strayer 2004; Roth & Kitchell 2005; Lodge et al. 2012). In addition, the displaced native species become more susceptible to predation because of a lack of shelter (Byron & Wilson 2001, Garvey & Stein 1993, Lodge et al. 2012). | |
| References:<br>Butler, M.J. & Stein, R.A., 1985, 'An analysis of the mechanisms governing species replacements in crayfish', *Oecologia* 66, 168-177. https://doi.org/10.1007/BF00379851.<br>Byron, C.J. & Wilson, K.A., 2001, 'Rusty crayfish (*Orconectes rusticus*) movement within and between habitats in Trout Lake, Vilas County, Wisconsin', *Journal of the North American Benthological Society* 20, 606–614. https://doi.org/10.2307/1468091.<br>Garvey, J.E. & Stein, R.A, 1993, 'Evaluating how chela size influences the invasion potential of an introduced crayfish (*Orconectes rusticus*)', *American Midland Naturalist 129,* 172–181. https://doi.org/10.2307/2426446 .<br>Hill, A.M. & Lodge, D.M., 1994, 'Diel changes in resource demand: Competition and predation in species replacement among crayfish, *Ecology* 75(7), 2118-2126. https://doi.org/10.2307/1941615.<br>Hobbs, H.H., Jass, J.P., Huner, J.V., 1989, 'A review of global crayfish introductions with particular emphasis on two North American species (Decapoda, Cambaridae), *Crustaceana* 56(3), 299-316. https://www.jstor.org/stable/20104461.<br>Klocker, C.A. & Strayer, D.L., 2004, 'Interactions among an invasive crayfish (*Orconectes rusticus*), a native crayfish (*Orconectes limosus*), and native bivalves (Sphaeriidae and Unionidae)', *Northeastern Naturalist* 11, 167–178. https://doi.org/10.1656/1092-6194(2004)011[0167:IAAICO]2.0.CO;2.<br>Lodge, D.M., Deines, A., Gherardi, F., Yeo, D.C.J., Arcella. T, Baldridge, A.K., et al., 2012, 'Global introductions of crayfishes: Evaluating the impact of species invasions on ecosystem services', *Annual* | |

*Review of Ecology Evolution Systematics* 43, 449–472. https://doi.org/10.1146/annurev-ecolsys-111511-103919

Lodge, D.M., Kershner, M.W. & Aloi, J.E., 1995, 'Effects of an omnivorous crayfish (*Orconectes rusticus*) on a freshwater littoral food web', *Ecology* 75, 1265–1281. https://doi.org/10.2307/1937452.

Roth, B.M. & Kitchell, J.F., 2005, 'The role of size-selective predation in the displacement of *Orconectes* crayfishes following rusty crayfish invasion', *Crustaceana* 78, 297-310. https://doi.org/10.1139/f05-217.

Wilson, K.A., Magnuson, J.J., Lodge, D.M., Hill, A.M., Kratz, T.K., Perry, W.L., et al., 2004, 'A long-term rusty crayfish (*Orconectes rusticus*) invasion: Dispersal patterns and community change in a north temperate lake', *Canadian Journal of Fisheries and Aquatic Sciences* 61, 2255–2266. https://doi.org/10.1139/f04-170.

**CON1b: Predation**

| Response: Major (MR) | Confidence: High |
|---|---|

Rationale: *Faxinous rasticus* is a facultative omnivore that consumes detritus, macrophytes and invertebrates (Lodge et al. 1995, Wilson et al. 2004). In areas of introduction, direct predation by *F. rusticus* on native fauna has resulted in a decline in population size of at least one native fish species (Willis & Magnuson 2006; Lodge et al. 2012). For example, in Wisconsin, USA, predation by *F. rusticus* has been implicated in the decline of native fish populations of bluegill sunfish (*Lepomis macrochirus*) (Wilson et al. 2004), and lake trout (*Salvelinus namaycush*) (Jonas et al. 2005). *Faxonius rusticus* is also known to feed on invertebrates, and in Trout Lake, Wisconsin, USA, direct predation on invertebrate communities has led to a decrease in the mean abundance of several invertebrates (Klocker & Strayer 2004, Kreps et al. 2012, Lodge et al. 1995; Wilson et al. 2004).

References:

Johnson, P.T., Olden, J.D., Solomon, C.T. & Vander Zanden, M.J., 2009, 'Interactions among invaders: community and ecosystem effects of multiple invasive species in an experimental aquatic system', *Oecologia* 159, 161–170. https://doi.org/10.1007/s00442-008-1176-x.

Jonas, J.L., Claramunt, R.M., Fitzsimons, J.D., Marsden, J.E. & Ellrott, B.J., 2005, 'Estimates of egg deposition and effects of lake trout (*Salvelinus namaycush*) egg predators in three regions of the Great Lakes', *Canadian Journal of Fisheries and Aquatic Sciences* 62, 2254–2264. https://doi.org/10.1139/f05-141.

Kreps, T.A., Baldridge, A.K. & Lodge, D.M., 2012, 'The impact of an invasive predator (*Orconectes rusticus*) on freshwater snail communities: Insights on habitat-specific effects from a multilake long-term study', *Canadian Journal of Fisheries and Aquatic Sciences* 69, 1164–1173. https://doi.org/10.1139/f2012-052.

Klocker, C.A. & Strayer, D.L., 2004, 'Interactions among an invasive crayfish (*Orconectes rusticus*), a native crayfish (*Orconectes limosus*), and native bivalves (Sphaeriidae and Unionidae)', *Northeastern Naturalist* 11, 167–178. https://doi.org/10.1656/1092-6194(2004)011[0167:IAAICO]2.0.CO;2.

Lodge, D.M., Kershner, M.W. & Aloi, J.E., 1995, 'Effects of an omnivorous crayfish (*Orconectes rusticus*) on a freshwater littoral food web', *Ecology* 75, 1265–1281. https://doi.org/10.2307/1937452.

Lodge, D.M., Deines, A., Gherardi, F., Yeo, D.C., Arcella, T., Baldridge, A.K., et al., 2012, 'Global introductions of crayfishes: evaluating the impact of species invasions on ecosystem services', *Annual Review of Ecology, Evolution, and Systematics* 43, 449–472. https://doi.org/10.1146/annurev-ecolsys-111511-103919.

Wilson, K.A., Magnuson, J.J., Lodge, D.M., Hill, A.M., Kratz, T.K., Perry, W.L., et al., 2004, 'A long-term rusty crayfish (*Orconectes rusticus*) invasion: Dispersal patterns and community change in a north temperate lake', *Canadian Journal of Fisheries and Aquatic Sciences* 61, 2255–2266. https://doi.org/10.1139/f04-170.

Willis, T.V. & Magnuson, J.J., 2006, 'Responses of fish communities in five north temperate lakes to exotic species and climate', *Limnology and Oceanography*, 51, https://doi.org/10.4319/lo.2006.51.6.2808.

**CON1c: Hybridisation**

| Response: Data Deficient (DD) | Confidence: High |
|---|---|
| Rationale: There are no native freshwater crayfish in South Africa (de Moor 2002; Crandall & de Grave 2017), so there is no possibility of hybridisation with native species. | |
| References:<br><br>de Moor, I., 2002, 'Potential impacts of alien freshwater crayfish in South Africa', African *Journal of Aquatic Science* 27, 125–139. https://doi.org/10.2989/16085914.2002.9626584.<br>Crandall, K.A. & De Grave, S., 2017, 'An updated classification of the freshwater crayfishes (Decapoda: Astacidea) of the world, with a complete species list', *Journal of Crustacean Biology*, 37(5), 615–653. https://doi.org/10.1093/jcbiol/rux070. | |
| **CON1d: Transmission of disease** | |
| Response: Minor (MN) | Confidence: Low |
| Rationale: *Faxinous rusticus* is a known vector of pathogen such as the crayfish plague agent, *Aphanomyces astaci* (Panteleit et al. 2019; Butler et al. 2020) and microsporidia (*Cambaraspora* sp.) (Stratton et al. 2023). There are no native crayfish species in South Africa but crayfish plague pathogen has been shown to infect other freshwater decapods such as native potamonautid crabs (Svoboda et al. 2014) | |
| References:<br>Butler, E., P. Crigler, G. Robbins, and J. E. Blair. 2020. Preliminary survey of *Aphanomyces* sp. associated with native and invasive crayfish in the lower Susquehanna watershed of south central Pennsylvania. *Journal of Freshwater Ecology* 35(1), 223-233. https://doi.org/10.1080/02705060.2020.1779141.<br>Panteleit, J., T. Horvath, J. Jussila, J. Makkonen, W. Perry, R. Schulz, K. Theissinger, and A. Schrimpf. 2019. Invasive rusty crayfish (*Faxonius rusticus*) populations in North America are infected with the crayfish plague disease agent (*Aphanomyces astaci*). *Freshwater Science*, 38(2), 425-433. https://doi.org/10.1086/703417<br>Stratton, C. E., B. A. Kabalan, S. A. Bolds, L. S. Reisinger, D. C. Behringer, and J. Bojko. 2023. *Cambaraspora faxoni* n. sp. (Microsporidia: Glugeida) from native and invasive crayfish in the USA and a novel host of *Cambaraspora floridanus*. *Journal of Invertebrate Pathology* 199:107949. https://doi.org/10.1016/j.jip.2023.107949.<br>Svoboda, J., Strand, D.A., Vrålstad, T., Grandjean, F., Edsman, L., Kozák, P., Kouba, A., Fristad, R.F., Bahadir Koca, S. and Petrusek, A. 2014, 'The crayfish plague pathogen can infect freshwater-inhabiting crabs', *Freshwater Biology*, 59, 918-929, https://doi.org/10.1111/fwb.12315 | |
| **CON1e: Parasitism** | |
| Response: Data Deficient (DD) | Confidence: Medium |
| Rationale: *Faxinous rusticus*is not a parasitic taxon (Lodge et al. 2012). | |
| References:<br><br>Lodge, D.M., Deines, A., Gherardi, F., Yeo, D.C., Arcella, T., Baldridge, A.K., et al., 2012, 'Global introductions of crayfishes: evaluating the impact of species invasions on ecosystem services', *Annual Review of Ecology, Evolution, and Systematics* 43, 449–472. https://doi.org/10.1146/annurev-ecolsys-111511-103919 | |
| **CON1f: Poisoning/toxicity** | |
| Response: Data Deficient (DD) | Confidence: Medium |
| Rationale: No information is available to assess the level of impact. | |

| References: | |
|---|---|
| **CON1g: Bio-fouling or other direct physical disturbance** | |
| Response: Data Deficient (DD) | Confidence: Medium |
| Rationale: No information is available to assess the level of impact. | |
| References: | |
| **CON1h: Grazing/herbivory/browsing** | |
| Response: Major (MR) | Confidence: Medium |
| Rationale: *Faxonius rusticus* is a functional omnivore and feeds on plant and animal material and is known to cause major impacts on invaded ecosystems through grazing on macrophytes (e.g., Hobb et al. 1989, Lodge et al. 1994; Wilson et al. 2004, Roth et al. 2006; Rosenthal et al. 2006; Baldridge & Lodge 2014). | |
| References:<br>Baldridge, A. K., & Lodge, D. M., 2014, 'Long-term studies of crayfish invaded lakes reveal limited potential for macrophyte recovery from the seed bank', *Freshwater Science*, 33, 788–797. https://doi.org/10.1086/677070<br>Hobbs, H.H., Jass, J.P. & Huner, J.V., 1989, 'A review of global crayfish introductions with particular emphasis on two North American species (Decapoda, Cambaridae)', *Crustaceana* 56(3), 299-316. https://www.jstor.org/stable/20104461.<br>Lodge, D.M., Kershner, M.W., Aloi, J.E. & Covich, A.P., 1995, 'Effects of an omnivorous crayfish (*Orconectes rusticus*) on a freshwater littoral food web', *Ecology*, 75: 1265-1281. https://doi.org/10.2307/1937452<br>Rosenthal, S. K., Stevens, S. S., & Lodge, D. M., 2006, 'Whole-lake effects of invasive crayfish (*Orconectes* spp.) and the potential for restoration', *Canadian Journal of Fisheries and Aquatic Sciences*, 63, 1276–1285. https://doi.org/10.1139/f06-037<br>Roth, B.M, Hein, C.L. & Vander Zanden, M.J., 2006, 'Using bioenergetics and stable isotopes to assess the trophic role of rusty crayfish (*Orconectes rusticus*) in lake littoral zones', *Canadian Journal of Fisheries and Aquatic Sciences* 63, 335–344. https://doi.org/10.1139/f05-217.<br>Wilson, K.A., Magnuson, J.J., Lodge, D.M., Hill, A.M., Kratz, T.K., Perry, W.L., et al., 2004, 'A long-term rusty crayfish (*Orconectes rusticus*) invasion: Dispersal patterns and community change in a north temperate lake', *Canadian Journal of Fisheries and Aquatic Sciences* 61, 2255–2266. https://doi.org/10.1139/f04-170 | |
| **CON1i: Chemical, physical or structural impact on ecosystem** | |
| Response: Data Deficient (DD) | Confidence: Low |
| Rationale: *Faxonius rusticus* lives in open water during most life stages and burrows only under extreme adverse conditions (GISD 2015). There is no information on this burrowing behaviour having a chemical, physical and or structural impact on ecosystems. | |
| References:<br>Global Invasive Species Database (GISD) 2015. Species profile *Orconectes rusticus*. Available from: http://www.iucngisd.org/gisd/species.php?sc=217 [Accessed 03 July2018]. | |
| **CON1k: Indirect impacts through interactions with other species** | |
| Response: Moderate (MO) | Confidence: Medium |

Rationale: In its invasive range, *Faxonius rusticus* may occur with other alien species, and interactions with other these species could facilitate impacts that ultimately lead to a decline in population size of native fauna (Jonson et al. 2009; Lodge et al 2012). In USA, *F. rusticus* occurs with the invasive snail *Bellamya chinesis* that has a thick shell that prevents predation by *F. rusticus* (Johnson et al. 2009). However, competition and predation pressure from both invasive species have reduced native snail biomass (Johnson et al. 2009).

References:

Johnson, P.T., Olden, J.D., Solomon, C.T. & Vander Zanden, M.J., 2009, 'Interactions among invaders: community and ecosystem effects of multiple invasive species in an experimental aquatic system', *Oecologia* 159, 161–170. https://doi.org/10.1007/s00442-008-1176-x.

**CON1 Maximum environmental impact**

| Response: Major (MR) | Confidence: High |
| --- | --- |

Rationale: In introduced areas, direct predation on native invertebrates, competition for food, shelter and spawning sites have led to local extinctions and a decrease in the abundance of native crayfish and fish species (Garvey & Stein 1993; Klocker & Strayer 2004; Lodge et al. 2012). *Faxonius rusticus* feeds on macrophytes leading to changes in structural communities. There are also records of hybridisation in the alien range, threatening indigenous crayfish populations (Perry et al. 2001).

References:

Garvey, J.E. & Stein, R.A, 1993, 'Evaluating how chela size influences the invasion potential of an introduced crayfish (*Orconectes rusticus*)', *American Midland Naturalist 129,* 172–181. , https://doi.org/10.1046/j.1523-1739.1996.10041155.x.

Klocker, C.A. & Strayer, D.L., 2004, 'Interactions among an invasive crayfish (*Orconectes rusticus*), a native crayfish (*Orconectes limosus*), and native bivalves (Sphaeriidae and Unionidae)', *Northeastern Naturalist* 11**,**167–178https://doi.org/10.1656/1092-6194(2004)011[0167:IAAICO]2.0.CO;2.

Lodge, D.M., Deines, A., Gherardi, F., Yeo, D.C.J., Arcella. T, Baldridge, A.K., et al., 2012, 'Global introductions of crayfishes: Evaluating the impact of species invasions on ecosystem services', *Annual Review of Ecology Evolution Systematics* 43, 449–472. https://doi.org/10.1146/annurev-ecolsys-111511-103919

Perry, W.L., Feder, J.L., Dwyer, G. & Lodge, D.M., 2001, 'Hybrid zone dynamics and species replacement between *Orconectes* crayfishes in a northern Wisconsin lake', *Evolution* 55, 1153–1166. https://doi.org/10.1111/j.0014-3820.2001.tb00635.x.

**CON2 Socio-economic impact**

**CON2a: Safety**

| Response: Data Deficient (DD) | Confidence: Medium |
| --- | --- |

Rationale: No information is available to assess the level of impact.

References:

**CON2b: Material and immaterial assets**

| Response: Moderate (MO) | Confidence: Medium |
| --- | --- |

Rationale: *Faxonius rusticus* invasions disrupt recreational activities in the invaded range leading to lost income from recreational angling (Keller et al. 2008). For example, in Vilas County (USA), *F. rusticus* has

| | |
|---|---|
| reduced sport fish populations through egg predation and/or competition with juvenile fish. Consequently, resulting in an estimated annual loss of 1.5 million US dollars. | |
| References:<br><br>Keller, R.P., Frang, K. & Lodge, D.M., 2008, 'Preventing the spread of invasive species: Economic benefits of intervention guided by ecological predictions' *Conservation Biology* 22, 80–88. https://doi.org/10.1111/j.1523-1739.2007.00811.x | |
| **CON2c: Health** | |
| Response: Data Deficient (DD) | Confidence: Medium |
| Rationale: No information is available to assess the level of impact. | |
| References: | |
| **CON2d: Social, spiritual and cultural relations** | |
| Response: Data Deficient (DD) | Confidence: Medium |
| Rationale: No information is available to assess the level of impact. | |
| References: | |
| **CON2 Maximum socio-economic impact** | |
| Response: Moderate (MO) | Confidence: Low |
| Rationale: *Faxonius rusticus* invasions have caused disruption in recreational activities by reducing sport fish populations though egg predation and/or competition with juveniles (Keller et al. 2008). | |
| References:<br><br>Keller, R.P., Frang, K. & Lodge, D.M., 2008, 'Preventing the spread of invasive species: Economic benefits of intervention guided by ecological predictions' *Conservation Biology* 22, 80–88. https://doi.org/10.1111/j.1523-1739.2007.00811.x. | |

| | |
|---|---|
| **CON3 Closely related species' environmental impact** | |
| Response: Not scored as CON1 scored | Confidence: |
| Rationale: There is information to score CON1 so CON3 does not need to be scored as per the framework. | |
| References: | |

| | |
|---|---|
| **CON4 Closely related species' socio-economic impact** | |
| Response: Not scored as CON2 scored | Confidence: |
| Rationale: There is information to score CON2 so CON4 does not need to be scored as per the framework. | |
| References: | |

| **CON5 Potential impact** | |
|---|---|
| Response: Major (MR) | Confidence: High |
| Rationale: *Faxonius rusticus* is known to cause impacts in introduced areas, mainly through competition, predation and excessive grazing (Lodge et al. 2012). *Faxonius rusticus* is highly fecund and an aggressive competitor (Garvey & Stein 1993). *Faxonius rusticus* may out-compete native freshwater crab species in South Africa (Jackson et al. 2016; Twardochleb et al. 2018). Being a functional omnivore, *F. rusticus* may also have detrimental impacts on macroinvertebrates and macrophyte communities when occurring in high densities (Wilson et al. 2004; Roth et al. 2006). *Faxonius rusticus* can reduce invertebrate communities, changing their composition in invaded areas (Klocker & Strayer 2004; Johnson et al. 2009). Furthermore, being a vector for the crayfish plague is another cause of concern, as it may be transferable and could be detrimental to native freshwater crustaceans in South Africa (Twardochleb et al. 2013; Panteleit et al. 2019; Butler et al. 2020). Although *F. rusticus* has not been implicated as an active vector, all north American crayfish are immune to the disease and can easily infect other species. | |

References:

Butler, E., P. Crigler, G. Robbins, & Blair, B.E., 2020, 'Preliminary survey of *Aphanomyces* sp. associated with native and invasive crayfish in the lower Susquehanna watershed of south central Pennsylvania', *Journal of Freshwater Ecology* 35(1), 223-233. https://doi.org/10.1080/02705060.2020.1779141.

Garvey, J.E. & Stein, R.A, 1993, 'Evaluating how chela size influences the invasion potential of an introduced crayfish (*Orconectes rusticus*)', *American Midland Naturalist 129,* 172–181. https://doi.org/10.1046/j.1523-1739.1996.10041155.x.

Holdich, D.M. & Reeve, I.D., 1991, 'Distribution of freshwater crayfish in the British Isles, with particular reference to crayfish plague, alien introductions and water quality', *Aquatic Conservation: Marine and Freshwater Ecosystems* 1, 139–158. https://doi.org/10.1002/aqc.3270010204.

Johnson, P.T., Olden, J.D., Solomon, C.T. & Vander Zanden, M.J., 2009, 'Interactions among Invaders: Community and ecosystem effects of multiple invasive species in an experimental aquatic system', *Oecologia* 159, 161–170. https://doi.org/10.1007/s00442-008-1176-x

Klocker, C.A. & Strayer, D.L., 2004, 'Interactions among an invasive crayfish (*Orconectes rusticus*), a native crayfish (*Orconectes limosus*), and native bivalves (Sphaeriidae and Unionidae)', *Northeastern Naturalist* 11**,** 167–178. https://doi.org/10.1656/1092-6194.

Lodge, D.M., Kershner, M.W., Aloi, J.E. & Covich, A.P., 1995, 'Effects of an omnivorous crayfish (*Orconectes rusticus*) on a freshwater littoral food web', *Ecology*, 75: 1265-1281. https://doi.org/10.2307/1937452

Panteleit, J., Horvath, T., Jussila, J., Makkonen, J., Perry, W., Schulz, R., Theissinger, K. & Schrimpf, A., 2019, 'Invasive rusty crayfish (Faxonius rusticus) populations in North America are infected with the crayfish plague disease agent (*Aphanomyces astaci*)', *Freshwater Science*, 38(2), 425-433. https://doi.org/10.1086/703417

Roth, B.M, Hein, C.L. & Vander Zanden, M.J., 2006, 'Using bioenergetics and stable isotopes to assess the trophic role of rusty crayfish (*Orconectes rusticus*) in lake littoral zones', *Canadian Journal of Fisheries and Aquatic Sciences* 63, 335–344. https://doi.org/10.1139/F05-217.

Twardochleb, L.A, Olden, J.D. & Larson, E.R., 2013, 'A global meta-analysis of the ecological impacts of nonnative crayfish', *Freshwater Science* 32, 1367–1382. http://dx.doi.org/10.1899/12-203.1

Wilson, K.A., Magnuson, J.J., Lodge, D.M., Hill, A.M., Kratz, T.K., Perry, W.L., et al., 2004, 'A long-term rusty crayfish (*Orconectes rusticus*) invasion: Dispersal patterns and community change in a north temperate lake', *Canadian Journal of Fisheries and Aquatic Sciences* 61, 2255–2266. https://doi.org/10.1139/f04-170.

#### 4. Management

| MAN1 What is the feasibility to stop future immigration? | |
|---|---|
| Response: High | Confidence: Low |
| Rationale: *Faxonius rusticus* is a popular bait species among anglers in the USA and is often stocked by fishermen (Kerr 2014). It also present in the pet trade industry in Germany and the USA (Chucholl 2013; Faulkes 2015). It is suspected that *Faxonius rusticus* is present in the pet trade in South Africa, but this needs to be confirmed (de Moore 2002; Nunes et al. 2017). The species is absent in neighbouring countries (Madzivanzira et al. 2021), so unaided immigration is unlikely. Moreover, it should be possible to regulate intentional introductions through legislative regulations and having an effective border surveillance and monitoring strategy (Zengeya & Wilson 2020; Wilson & Kumschick 2024). | |
| References:<br><br>de Moor, I., 2002, 'Potential impacts of alien freshwater crayfish in South Africa', *African Journal of Aquatic Science* 27, 125–139https://doi.org/10.2989/16085914.2002.9626584.<br>Chucholl, C., 2013, 'Invaders for sale: Trade and determinants of introduction of ornamental freshwater crayfish', *Biological Invasions* 15, 125–141. https://doi.org/10.1007/s10530-012-0273-2.<br>Faulkes, Z, 2015, 'The global trade in crayfish as pets', *Crustacean Research* 44, 75–92. https://doi.org/10.18353/crustacea.44.0_75.<br>Kerr, S.J., 2014, 'The Introduction and spread of aquatic invasive species through the recreational use of bait: A literature review. Report prepared for Biodiversity Branch. Ontario Ministry of Natural Resources. Peterborough', Ontario, Canada.<br>Madzivanzira, T.C., South, J., Wood, L.E., Nunes, A.L. & Weyl, O.L.F., 2020, 'A review of freshwater crayfish introductions in Africa', *Reviews in Fisheries Science & Aquaculture*, 1-21. https://doi.org/10.1080/23308249.2020.1802405.<br>Nunes, A.L., Zengeya, T.A., Measey, G.J. & Weyl, O.L.F., 2017, 'Freshwater crayfish invasions in South Africa: past, present and potential future', *African Journal of Aquatic Science* 42, 309–323. https://doi.org/10.2989/16085914.2017.1405788.<br>Wilson, J. R., & Kumschick, S., 2024, 'The regulation of alien species in South Africa', South *African Journal of Science*, *120*(5/6). https://doi.org/10.17159/sajs.2024/17002<br>Zengeya, T.A. & Wilson, J.R., (eds.) 2020, *The status of biological invasions and their management in South Africa in 2019*, pp.71, South African National Biodiversity Institute, Kirstenbosch and DSI-NRF Centre of Excellence for Invasion Biology, Stellenbosch. http://dx.doi.org/10.5281/zenodo.3947613. | |

| MAN2 Benefits of the Taxon | |
|---|---|
| **MAN2a Socio-economic benefits of the *Taxon*** | |
| Response: Low | Confidence: Low |
| Rationale: *Faxonius rusticus* is popularly used as a live bait by anglers for recreational fishing (Kerr 2014; Olden et al. 2009). *Faxonius rusticus* is present in the pet trade industry in Germany and the USA (Faulkes 2015). No socio-economic benefits have been recorded to date in South Africa and even if *F. rusticus* is present the benefits are likely to be low (Madzivanzira et al. 2021). | |
| References:<br><br>Faulkes, Z, 2015, 'The global trade in crayfish as pets', *Crustacean Research* 44, 75–92. https://doi.org/10.18353/crustacea.44.0_75. | |

| |
|---|
| Kerr, S.J., 2014, 'The introduction and spread of aquatic invasive species through the recreational use of bait: A literature review", Report prepared for Biodiversity Branch. Ontario Ministry of Natural Resources. Peterborough', Ontario, Canada.<br>Olden, J.D., Adams, J.W. & Larson, E.R., 2009, 'First record of *Orconectes rusticus* (Girard, 1852) (Decapoda, Cambaridae) west of the great continental divide in North America', *Crustaceana* 82, 1347–1351. https://doi.org/10.56854009X448934.<br>Madzivanzira, T.C., South, J., Wood, L.E., Nunes, A.L. & Weyl, O.L.F., 2020, 'A review of freshwater crayfish introductions in Africa', *Reviews in Fisheries Science & Aquaculture*, 1-21. https://doi.org/10.1080/23308249.2020.1802405 . |

| **MAN2b Environmental benefits of the *Taxon*** | |
|---|---|
| Response: Low | Confidence: High |
| Rationale: There are no recorded environmental benefits of *Faxonius rusticus.* | |
| References: | |

| **MAN3 Ease of management** | |
|---|---|
| **MAN3a How accessible are populations?** | |
| Response: 2 (difficult to access) | Confidence: Low |
| Rationale: There are no locality records for *Faxonius rusticus* in South Africa (Madzivanzira et al. 2021; Nunes et al. 2017). However, given that it is an aquatic species, accessibility is likely to be difficult. | |
| References:<br><br>Madzivanzira, T.C., South, J., Weyl, O.L.F., 2021, 'Invasive crayfish outperform potamonautid crabs at higher temperatures', *Freshwater Biology* 66, 978–991. https://doi.org/10.1111/fwb.13691<br>Nunes, A.L., Zengeya, T.A., Measey, G.J. & Weyl, O.L.F., 2017, 'Freshwater crayfish invasions in South Africa: past, present and potential future', *African Journal of Aquatic Science* 42, 309–323, https://doi.org/10.2989/16085914.2017.1405788. | |
| **MAN3b Is detectability critically time-dependent?** | |
| Response: 2 (yes) | Confidence: High |
| Rationale: *Faxonius rusticus* is a small, largely nocturnal freshwater species that is difficult to detect (Hein et al. 2006). | |
| References:<br><br>Hein, C.L., Roth, B.M., Ives, A.R. & Vander Zanden, M.J., 2006, 'Fish predation and trapping for rusty crayfish (*Orconectes rusticus*) control: A whole-lake experiment', *Canadian Journal Fisheries and Aquatic Sciences*. 63, 383–393. https://doi/org/10.1139/F05-229. | |
| **MAN3c Time to reproduction** | |
| Response: 1 (1-3 years) | Confidence: High |
| Rationale: *Faxonius rusticus* reach sexual maturity in one-two years (Phillips 2010). | |
| References: | |

| |
|---|
| Phillips, I.D. 2010. Biological synopsis of the Rusty Crayfish (*Orconectes rusticus*). Canadian Manuscript Report of Fisheries and Aquatic Science. 2923: iv + 17 p. |

| **MAN3d Propagule persistence** | |
|---|---|
| Response: 0 (<1 year) | Confidence: High |
| Rationale: *Faxonius rusticus* are highly fecund, a female produces up to 200 eggs, and hatchlings hang under the female's tail for many weeks until they can feed on their own (Lodge et al. 1985). | |
| References:<br><br>Lodge, D.M., Beckel, A.L. & Magnuson J.J., 1985, 'Lake-bottom tyrant', *Natural History* 94, 32-37. | |

| **MAN3 Ease of management** | |
|---|---|
| Response: Medium | Confidence: High |
| Rationale: *Faxonius rusticus* is a small, largely nocturnal freshwater species that is difficult to detect (Hein et al. 2006). It reaches sexual maturity in one-two years and is highly fecund (up to 200 eggs) (Lodge et al. 1985; Phillips 2010). | |
| References:<br><br>Hein, C.L., Roth, B.M., Ives, A.R. & Vander Zanden, M.J., 2006, 'Fish predation and trapping for rusty crayfish (*Orconectes rusticus*) control: A whole-lake experiment', *Canadian Journal Fisheries and Aquatic Sciences*. 63, 383–393. https://doi/org/10.1139/F05-229.<br>Lodge, D.M., Beckel, A.L. & Magnuson J.J., 1985, 'Lake-bottom tyrant', *Natural History* 94, 32-37.<br>Phillips, I.D. 2010. Biological synopsis of the Rusty Crayfish (*Orconectes rusticus*). Canadian Manuscript Report of Fisheries and Aquatic Science. 2923: iv + 17 p. | |

| **MAN4 Has the feasibility of eradication been evaluated?** | |
|---|---|
| Response: No | Confidence: Low |
| Rationale: There are no known population of *F. rusticus* in South Africa, therefore eradication feasibility has not been no evaluated. There is no documented eradication elsewhere in the invasive range of *F. rusticus* but see MAN5 for an example of control efforts using trapping and enhancement of predatory fishes (Hein et al. 2007; Perales et al. 2021) | |
| References:<br><br>Hein, C.L., Roth, B.M., Ives, A.R. & Vander Zanden, M.J., 2006, 'Fish predation and trapping for rusty crayfish (*Orconectes rusticus*) control: A whole-lake experiment', *Canadian Journal Fisheries and Aquatic Sciences*. 63, 383–393. https://doi/org/10.1139/F05-229.<br>Perales, K.M., Hansen, G.J.A., Hein, C.L., Mrnak, J.T., Roth, B.M., Walsh, J.R., & van der Zanden, M. J., 2021, 'Spatial and temporal patterns in native and invasive crayfishes during a 19-year whole-lake invasive crayfish removal experiment', *Freshwater Biology*, 66, 2105–2117. https://doi.org/10.1111/fwb.13818 | |

| **MAN5 Control options and monitoring approaches available for the *Taxon*** |
|---|
| Response: *Faxonius rusticus* invasions in the USA have been controlled using trapping and enhancement of predatory fishes (Hein et al. 2007). For example, a whole lake experiment (2001-2008) was carried out to control *F. rusticus* in Northern Wisconsin, USA and resulting in a ≥95% reduction in abundance of rusty crayfish and |

recovery of native species populations (Hein et al., 2007; Hansen et al., 2013; Perales et al. 2021). There are several other methods that have been used to control alien populations of invasive crayfish these include mechanical removal using traps and electrofishing, biocides, and sterile male release (Gherardi et al. 2011). However, there are several issues that make management of crayfish invasions challenging. For example, the use of biocides is often effective for localised populations in dams and ponds but it is not effective when a species is widespread because of the large doses of treatment required and higher potential of adverse effects on non-targeted species (Gherardi et al. 2011; Peay et al. 2019). Mechanical control can be effective in small discrete water bodies but is expensive, requires commitment to follow up to complete extirpation and can lead to unwanted community compensation mechanisms such as enhanced recruitment (Chadwick et al. 2020). *Faxonius rusticus* is present in the global pet trade industry and there are unconfirmed records that it might be present South Africa. The pet trade industry is generally difficult to regulate because trade restrictions imposed on species of conservation concern that are potentially popular and in high demand, often lead to illegal trading that is compounded by a lack of reliable record–keeping of the type and number of organisms presently in the pet trade (Gippet & Bertelsmeier 2021)

References:

Chadwick, D.D.A., Pritchard, E.G., Bradley, P., Sayer, C.D., Chadwick, M.A., Eagle, L.J.B., et al., 2020, 'A novel triple down method highlights deficiencies in invasive alien crayfish survey and control techniques', *Journal of Applied Ecology* 58(2), 316–326. https://doi.org/10.1111/1365–2664.13758

Gherardi, F., Aquiloni, L., Diéguez-Uribeondo, J. & Tricarico, E., 2011, 'Managing invasive crayfish: is there a hope?', *Aquatic Sciences* 73, 185-200. https://doi.org/10.1007/s00027-011-0181-z.

Gippet, J.M.& Bertelsmeier, C. 2021, 'Invasiveness is linked to greater commercial success in the global pet trade', *Proceedings of the National Academy of Sciences* 118, e2016337118. https://doi.org/10.1073/pnas.2016337118.

Hansen, G. J. A., Hein, C. L., Roth, B. M., Vander Zanden, M. J., Gaeta, J. W., Latzka, A. W. & Carpenter, S. R., 2013, 'Food web consequences of long-term invasive crayfish control', *Canadian Journal of Fisheries and Aquatic Sciences*, 70, 1109–1122. https://doi.org/10.1139/cjfas-2012-0460

Hein, C.L., Roth, B.M., Ives, A.R. & Vander Zanden, M.J., 2006, 'Fish predation and trapping for rusty crayfish (*Orconectes rusticus*) control: A whole-lake experiment', *Canadian Journal Fisheries and Aquatic Sciences*. 63, 383–393. https://doi/org/10.1139/F05-229.

Peay, S., Johnsen, S.I., Bean, C.W., Dunn, A.M., Sandodden, R. & Edsman, L. 2019, 'Biocide treatment of invasive signal crayfish: successes, failures and lessons learned', *Diversity, 11,* 29. https://doi.org/10.3390/d11030029.

Perales, K.M., Hansen, G.J.A., Hein, C.L., Mrnak, J.T., Roth, B.M., Walsh, J.R., & Vander Zanden, M. J., 2021, 'Spatial and temporal patterns in native and invasive crayfishes during a 19-year whole-lake invasive crayfish removal experiment', *Freshwater Biology*, 66, 2105–2117. https://doi.org/10.1111/fwb.13818

| **MAN6 Any other management considerations to highlight?** | |
|---|---|
| Response | No |

**5. Calculations**

**Likelihood = Fairly probable**

| Parameter | Likelihood | Stages | Final assessment |
|---|---|---|---|
| LIK1 | 0.0027 | P(entry) = 0.5 | P (invasion) = 0.125 |
| LIK2 | 0.5 | | |
| LIK3 | 0.5 | P(establishment) = 0.5 | |
| LIK4 | 0.5 | | |
| LIK5 | 0.5 | P (spread) = 0.5 | |
| LIK6 | 0.5 | | |

**Consequence = Major (MR)**

| Parameter | Mechanism/sector | Response |
|---|---|---|
| CON1a | Competition | MR |
| CON1b | Predation | MR |
| CON1c | Hybridisation | DD |
| CON1d | Disease transmission | MN |
| CON1e | Parasitism | DD |
| CON1f | Poisoning/toxicity | DD |
| CON1g | Bio-fouling or other direct physical disturbance | DD |
| CON1h | Grazing/herbivory/browsing | MR |
| CON1i | Chemical, physical, structural impact | DD |
| CON1k | Indirect impacts through interactions with other species | MO |
| CON1 | Maximum environmental impact | MR |
| CON2a | Safety | DD |
| CON2b | Material and immaterial assets | MO |
| CON2c | Health | DD |
| CON2d | Social, spiritual and cultural relations | DD |
| CON2 | Maximum socio-economic impact | MO |
| CON3 | Environmental impact of closely related taxa | Not scored as CON1 scored |

| | | |
| --- | --- | --- |
| CON4 | Socio-economic impact of closely related taxa | Not scored as CON2 scored |
| CON5 | Potential impact based on traits, experiments, or models | MR |

**Risk score = High**

| | | Consequences | | | | |
| --- | --- | --- | --- | --- | --- | --- |
| | | MC | MN | MO | MR | MV |
| Likelihood | Extremely unlikely | low | low | low | medium | medium |
| | Very unlikely | low | low | low | medium | high |
| | Unlikely | low | low | medium | high | high |
| | Fairly probable | medium | medium | high | high | high |
| | Probable | medium | high | high | high | high |

**Ease of management** = *Faxonius rusticus* is not present in South Africa. According to the framework, MAN3 does not need to be scored.

| Parameter | Question | Response |
| --- | --- | --- |
| MAN3a | How accessible are populations? | 2 |
| MAN3b | Is detectability critically time-dependent? | 2 |
| MAN3c | Time to reproduction | 1 |
| MAN3d | Propagule persistence | 0 |
| **MAN3** | **SUM** | 5 |

**Appendix BAC7**: The global native range of *Faxonius rusticus*. Source: CABI, 2022. *Faxonius rusticus* [original text by Elena Tricarico]. In: Invasive Species Compendium. Wallingford, UK: CAB International. www.cabi.org/isc.

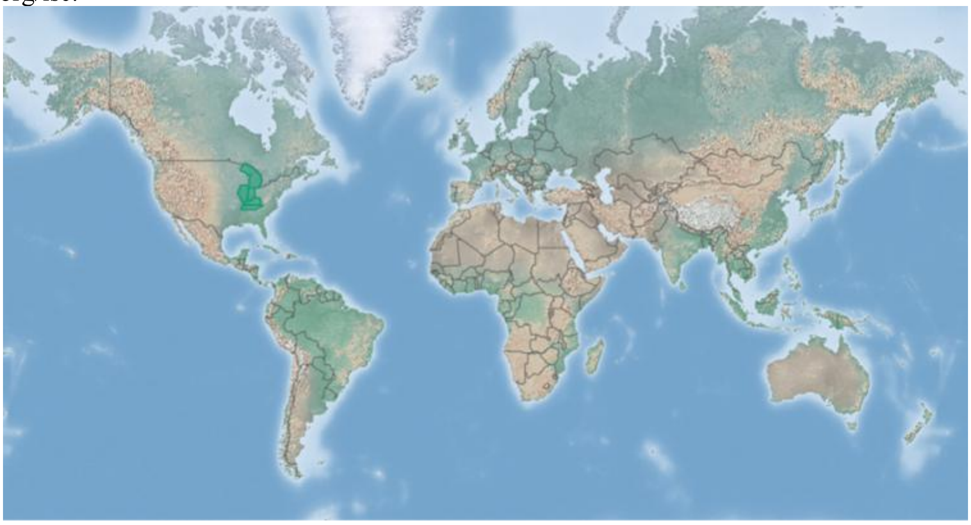

**Appendix BAC8(a)**: The global alien range of *Faxonius rusticus*. Source: CABI, 2021. *Faxonius rusticus* [original text by Elena Tricarico]. In: Invasive Species Compendium. Wallingford, UK: CAB International. www.cabi.org/isc.

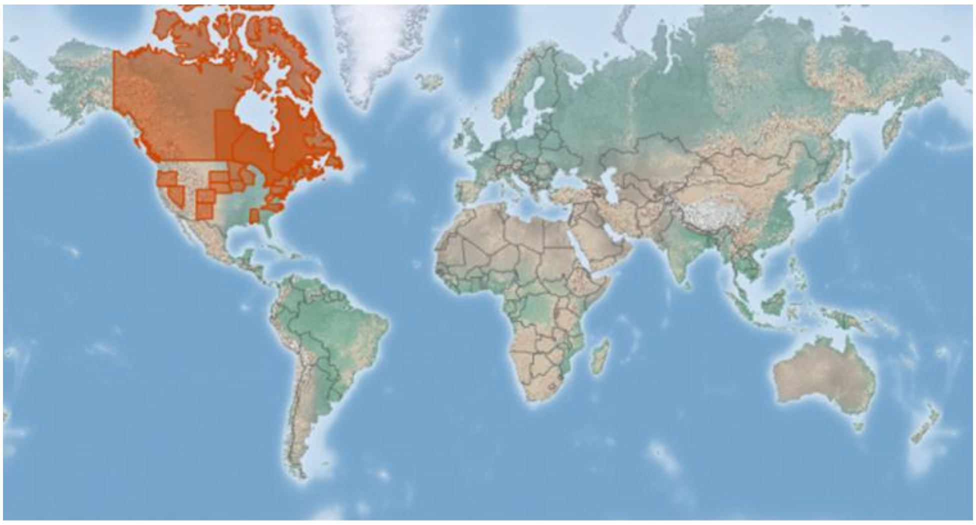

#### 1.9 Risk analysis report for virile crayfish (*Faxonius virilis*)

| | |
|---|---|
| ***Taxon***: *Faxonius virilis* (Hagen, 1870) | ***Area***: South Africa |
| **Picture of *Taxon***<br>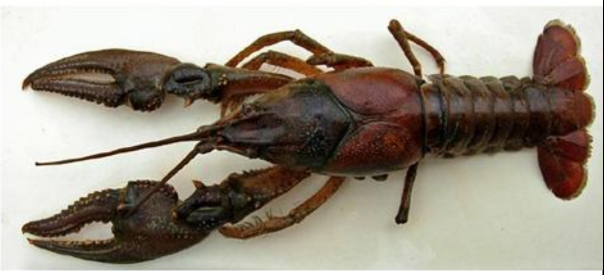<br>Ahern et al. 2008. | **Alien distribution map**<br>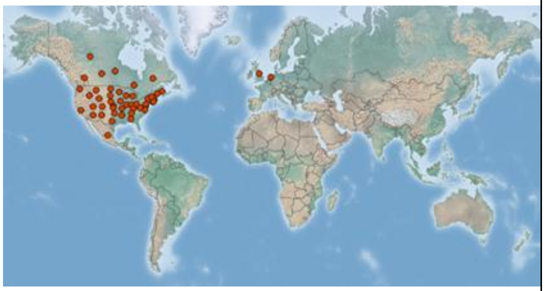<br>Sourced from CABI (2019): https://www.cabi.org/ISC/datasheet/72034 |
| **Risk Assessment summary:**<br><br>The Virile crayfish (*Faxonius virilis*) has been widely introduced outside its native in North America. It has also been introduced to several countries in Europe. There are no know populations of *F. virilis* in South Africa but there is anecdotal evidence that it is present in the ornamental trade and this needs to be verified. There are no known populations in neighbouring countries, therefore via unaided pathways is unlikely. The illegal pet trade industry remains a concern because *F. virilis* may be moved around by humans and there is a high risk of release of unwanted pets. *Faxonius virilis* has being implicated in the decline, displacement and local extirpation of native macroinvertebrates, crayfishes, snails, frogs and fish in its introduced range. *Faxonius virilis* is also a vector for the crayfish plague which may be transferred to native decapod species. It is known to burrow in its home ranges which may result in the destabilization of riverbanks, causing erosion. | **Risk score:**<br><br>High |
| **Management options summary:**<br><br>*Faxonius virilis* is a small, largely nocturnal, burrowing freshwater species that is difficult to detect. Several methods can be used to control *F. virilis* such as mechanical removal using traps and electrofishing, biocides, and biological control. However, there are several issues that make the use of these methods challenging. The benefits of *F. virilis* were scored as low. | **Ease of management:**<br><br>Difficult |
| **Recommendations:**<br><br>*Faxonius virilis* poses a high risk to South Africa. If *F. virilis* were to establish outside of captivity, such populations would likely be very difficult to extirpate. | **Listing under NEM:BA A&IS lists of 2020:**<br><br>Not listed |

| | |
|---|---|
| As such it is recommended that *F. virilis* is listed as category 1a, active efforts made to establish whether the species is preset in the country, and that any individuals found are controlled. This would likely require a dedicated assessment of the ornamental trade. A reassessment in five years is thus required and, if absent from the country, *F. virilis s*hould be added to a future prohibited list. | **Recommended listing category:**<br><br>1a[1] |

[1] The presence of *F. virilis* in South Africa should be confirmed and the taxon should be reassessed in five years (i.e., by 2029).

#### 1. Background

| BAC1 Name of assessor(s) | |
|---|---|
| Name of lead assessor | Lee-Anne Botha |
| Additional assessor (1) | Tsungai Zengeya |
| Additional assessor (2) | Christian Chimimba |
| **BAC2 Contact details of assessor (s)** | |
| Lead assessor | Organisational affiliation: Department of Zoology and Entomology, University of Pretoria and The South African National Biodiversity Institute (SANBI). |
| | |
| Additional assessor (1) | Organisational affiliation: Department of Zoology and Entomology, University of Pretoria and The South African National Biodiversity Institute (SANBI). |
| | |
| Additional assessor (2) | Organisational affiliation: Department of Zoology and Entomology, University of Pretoria |
| | |
| **BAC3 Name(s) and contact details of expert(s) consulted** | |
| Expert (1) | Name: |
| | email: |
| | Phone: |
| Expert (2) | Name: |
| | email: |
| | Phone: |
| Comments: none | |
| **BAC4 Scientific name of *Taxon* under assessment** | |
| *Taxon* name: *Faxonius virilis* | Authority: (Hagen, 1870) |
| Comments: The genus *Orconectes* previously included cave-dwelling and surface-dwelling crayfish (Crandall & De Grave 2017). The non-cave dwelling crayfish in the *Orconectes* genus were moved to *Faxonius* (Crandall & De Grave 2017). The taxonomic status of *F. virilis* has been verified (GBIF 2023). | |
| References:<br>Crandall, K.A., De Grave, S., 2017, 'An updated classification of the freshwater crayfishes (Decapoda: Astacidea) of the world, with a complete species list', *Journal of Crustacean Biology,* 27, 615–653. https://doi.org/10.1093/jcbiol/rux070.<br>*Faxonius virilis* (Hagen, 1870) in GBIF Secretariat (2023). GBIF Backbone Taxonomy. Checklist dataset https://doi.org/10.15468/39omei accessed via GBIF.org on 2024-07-11. | |
| **BAC5 Synonym(s) considered** | |
| Synonyms: *Cambarus virilis, Orconectes virilis, Cambarus couesi, Cambarus debilis, Cambarus wisconsinensis* | |
| Comments: There are several known synonyms of *F. varilis* (Crandall & De Grave 2017; GBIF 2023; CABI 2024) and the above listed synonyms were used for the literature search for this risk analysis. | |

References:

Crandall, K.A., De Grave, S., 2017, 'An updated classification of the freshwater crayfishes (Decapoda: Astacidea) of the world, with a complete species list', Journal of Crustacean Biology, 27, 615–653. https://doi.org/10.1093/jcbiol/rux070.

*Faxonius virilis* (Hagen, 1870) in GBIF Secretariat (2023). GBIF Backbone Taxonomy. Checklist dataset https://doi.org/10.15468/39omei accessed via GBIF.org on 2024-07-11.

CABI, 2024. *Faxonius virilis* (virile crayfish). Invasive Species Compendium, CABI Publishing, Wallingford, UK. https://www.cabidigitallibrary.org/doi/full/10.1079/cabicompendium.72034 (accessed 10 July 2024)

**BAC6 Common name(s) considered**

Common names: Virile crayfish, Northern crayfish, eastern crayfish, lake crayfish, common crayfish

Comments: There are several known synonyms of *F. varilis* (Crandall & De Grave 2017; GBIF 2023; CABI 2024) and the above listed synonyms were used for the literature search for this risk analysis.

References:

Crandall, K.A., De Grave, S., 2017, 'An updated classification of the freshwater crayfishes (Decapoda: Astacidea) of the world, with a complete species list', Journal of Crustacean Biology, 27, 615–653. https://doi.org/10.1093/jcbiol/rux070.

*Faxonius virilis* (Hagen, 1870) in GBIF Secretariat (2023). GBIF Backbone Taxonomy. Checklist dataset https://doi.org/10.15468/39omei accessed via GBIF.org on 2024-07-11.

CABI, 2024. *Faxonius virilis* (virile crayfish). Invasive Species Compendium, CABI Publishing, Wallingford, UK. https://www.cabidigitallibrary.org/doi/full/10.1079/cabicompendium.72034 (accessed 10 July 2024)

**BAC7 What is the native range of the *Taxon*?** (add map in Appendix BAC7)

| Response | Confidence |
|---|---|
| Response: USA (Missouri, Mississippi, Ohio, Montana, Missouri and Great Lakes) and Canada (from Alberta to Quebec) | Confidence: High |

Comments: Native to a large part of North America east of the continental divide (Adams et al. 2010; Larson and Olden 2011; USGS-NAS 2023).

References:

Adams S, Schuster GA, Taylor CA. 2010. *Orconectes virilis. The IUCN Red List of Threatened Species 2010*: https://www.iucnredlist.org/species/153831/4551026.

USGS-NAS, 2023. *Faxonius virilis* (Hagen, 1870). US Geological Survey, Nonindigenous Aquatic Species database. https://nas.er.usgs.gov/queries/FactSheet.aspx?speciesID=215

Larson, E.R., Olden, J.D., 2011. The state of crayfish in the Pacific Northwest. *Fisheries*, 36:60-73. https://doi.org/10.1577/03632415.2011.10389069

**BAC8 What is the global alien range of the *Taxon*?** (add map in Appendix BAC8)

| Response | Confidence |
|---|---|
| Response: **Europe**: France \| Netherlands \| Sweden \| United Kingdom **North America**: Wyoming \| West Virginia \| Vermont \| Utah \| Tennesse \| Rhode Island \| Pennsylvania \| New Mexico \| New Hampshire \| Montana \| Massachusetts \| Maryland \| Kansas \| Idaho \| Connecticut \| Colorado \| California \| Arizona \| Alabama \| Mexico | Confidence: Medium |

Comments: *Faxonius virilis* has been translocated within the United States of America outside of its native range. It has also been introduced to several countries Europe (Adams et al. 2010; USGS-NAS 2023; CABI 2024).

References:

Adams S, Schuster GA, Taylor CA. 2010. *Orconectes virilis. The IUCN Red List of Threatened Species 2010*: https://www.iucnredlist.org/species/153831/4551026 .

USGS-NAS, 2023. *Faxonius virilis* (Hagen, 1870). US Geological Survey, Nonindigenous Aquatic Species database. https://nas.er.usgs.gov/queries/FactSheet.aspx?speciesID=215

CABI, 2024. *Faxonius virilis* (virile crayfish). Invasive Species Compendium, CABI Publishing, Wallingford, UK. https://www.cabidigitallibrary.org/doi/full/10.1079/cabicompendium.72034 (accessed 10 July 2024)

**BAC9 Geographic scope = the *Area* under consideration**

| | | |
|---|---|---|
| *Area* of assessment: South Africa | | |
| Comments: The geographic scope of this assessment is limited to South Africa. | | |
| **BAC10 Is the *Taxon* present in the *Area*?** | | |
| Response: No | | Confidence: Low |
| Comments: There are no records of *F. virilis* in South Africa (Nunes et al. 2017). *Faxonius rusticus* is present in the pet trade globally (Ahern et al. 2008; Lodge et al. 2012; Faulkes 2015), and it might be present in industry in South Africa but this needs to be assessed (Nunes et al. 2017). | | |
| References:<br>Ahern, D., England, J. & Ellis, A., 2008, 'The virile crayfish, *Orconectes virilis* (Hagen, 1870) (Crustacea: Decapoda: Cambaridae), identified in the UK. Invasive species in inland waters of Europe and North America: Distribution and Impacts', *Aquatic Invasions*, 3(1), 102-104. https://www.reabic.net/aquaticinvasions/2008/AI_2008_3_1_Ahern_etal.pdf<br>Faulkes Z., 2015, 'The global trade in crayfish as pets', *Crustacean Research,* 44, 75–92. https://doi.org/10.18353/crustacea.44.0_75.<br>Lodge, D.M., Deines, A., Gherardi, F., Yeo, D.C.J., Arcella, T., Baldridge, A.K., Branes, M.A., Chadderton, W.L., Feder, J.L., Grantz, C.A., Howard, G.W., Jerdde, C.L., Peters, J.A., Sargent, L.W., Turner, C.R., Wittmann, M.E., Zeng, Y., 2012, 'Global introductions of crayfishes: evaluating the impact of species invasions on ecosystem services', *Annual Review of Ecology Evolution Systematics*, 43, 449–472. https://doi.org/10.1146/annurev-ecolsys-111511-103919.<br>Nunes, A.L., Zengeya, T.A., Measey, G.J. & Weyl, O.L.F., 2017b, 'Freshwater crayfish invasions in South Africa: past, present and potential future', *African Journal of Aquatic Science* 42, 309–323. https://doi.org/10.2989/16085914.2017.1405788. | | |
| **BAC11 Availability of physical specimen** | | |
| Response: No | | Confidence in ID: |
| Herbarium or museum accession number: No *Faxonius virilis* specimens have been found in South Africa and it is assumed the species is not present in South Africa, although it requires further assessment, especially linked to the pet trade industry (Nunes et al. 2017). | | |
| References:<br>Nunes, A.L., Zengeya, T.A., Measey, G.J. & Weyl, O.L.F., 2017b, 'Freshwater crayfish invasions in South Africa: past, present and potential future', *African Journal of Aquatic Science* 42, 309–323. https://doi.org/10.2989/16085914.2017.1405788. | | |
| **BAC12 Is the *Taxon* native to the *Area* or part of the *Area*?** | | |
| The *Taxon* is native to (part of) the *Area*. | No | Confidence: High |
| The *Taxon* is alien in (part of) the *Area*. | Yes | Confidence: High |
| Comments: There are no native freshwater crayfish species native in South Africa (de Moor 2002; Nunes et al. 2017). | | |
| References:<br>de Moor, I., 2002, 'Potential impacts of alien freshwater crayfish in South Africa', *African Journal of Aquatic Science* 27, 125–139, https://doi.org/10.2989/16085914.2002.9626584.<br>Nunes, A.L., Zengeya, T.A., Measey, G.J. & Weyl, O.L.F., 2017b, 'Freshwater crayfish invasions in South Africa: past, present and potential future', *African Journal of Aquatic Science* 42, 309–323. https://doi.org/10.2989/16085914.2017.1405788. | | |
| **BAC13 What is the *Taxon*'s introduction status in the *Area*?** | | |
| The *Taxon* is in cultivation/containment. | Don't known | Confidence: Low |
| The *Taxon* is present in the wild. | None | Confidence: Low |
| The *Taxon* has established/naturalised. | None | Confidence: Low |
| The *Taxon* is invasive. | None | Confidence: Low |
| Comments: There are no records of *F. virilis* being present in South Africa. It is present in the pet trade in Europe (Chucholl 2013; Faulkes 2015; CABI 2024), and could likely be traded in South Africa but this needs to be assessed (de Moor 2002; Nunes et al. 2017). | | |

References:

CABI, 2024. *Faxonius virilis* (virile crayfish). Invasive Species Compendium, CABI Publishing, Wallingford, UK. https://www.cabidigitallibrary.org/doi/full/10.1079/cabicompendium.72034 (accessed 10 July 2024)

Chucholl, C., 2013, 'Invaders for sale: trade and determinants of introduction of ornamental freshwater crayfish', *Biological Invasions* 15, 125–141. https://doi.org/10.1007/s10530-012-0273-2.

de Moor, I., 2002, 'Potential impacts of alien freshwater crayfish in South Africa', *African Journal of Aquatic Science* 27, 125–139. https://doi.org/10.2989/16085914.2002.9626584.

Faulkes, Z., 2015, 'The global trade in crayfish as pets', *Crustacean Research*, 44, 75–92. https://doi.org/10.18353/crustacea.44.0_75.

Nunes, A.L., Zengeya, T.A., Measey, G.J. & Weyl, O.L.F., 2017b, 'Freshwater crayfish invasions in South Africa: past, present and potential future', *African Journal of Aquatic Science* 42, 309–323. https://doi.org/10.2989/16085914.2017.1405788.

**BAC14 Primary (introduction) pathways**

| | | |
|---|---|---|
| Release | None | Confidence: |
| Escape | Pet/aquarium/bait | Confidence: High |
| Contaminant | None | Confidence: |
| Stowaway | None | Confidence: |
| Corridor | None | Confidence: |
| Unaided | None | Confidence: |

Comments: *Faxonius virilis* is known to be present in the pet trade industry in Netherlands and the United Kingdom (Chucholl 2013; Faulkes 2015). In these areas, crayfish have established populations after accidental or intentional release by owners as unwanted pets (Faulkes 2015). In the USA*, F. virilis* is a popular bait species among anglers and is also stocked for forage (Larson and Olden 2011, Kilian et al. 2012).

References:

Chucholl, C., 2013, 'Invaders for sale: trade and determinants of introduction of ornamental freshwater crayfish', *Biological Invasions* 15, 125–141. https://doi.org/10.1007/s10530-012-0273-2.

Faulkes, Z., 2015, 'The global trade in crayfish as pets', *Crustacean Research*, 44, 75–92. https://doi.org/10.18353/crustacea.44.0_75.

Kilian, J.V., Klauda, R.J., WIdman, S., Kashiwagi, M., Bourquin, R., Weglein, S. & Schuster, J., 2012, 'An assessment of a bait industry and angler behavior as a vector of invasive species', *Biological Invasions*, 14(7), 1469-1481. https://doi.org/10.1007/s10530-012-0173-5.

Larson, E.R. & Olden, J.D., 2011, 'The state of crayfish in the Pacific Northwest', *Fisheries*, 36(2), 60-73. https://doi.org/10.1577/03632415.2011.10389069

#### 2. Likelihood

**LIK1 Likelihood of entry via unaided primary pathways**

| | |
|---|---|
| Response: very unlikely (p = 0.0027) | Confidence: Medium |

Rationale: There is no evidence that *F. virilis* is present in South Africa or has established in the wild in neighbouring countries in the SADC Region (Lodge et al. 2012; Madzivanzira et al. 2020) therefore the probability of *F. virilis* entering South Africa through unaided pathways (connected waterways) is very unlikely (p = 0.0027).

References:

Madzivanzira, T.C., South, J., Wood, L.E., Nunes, A.L. & Weyl, O.L.F., 2020, 'A review of freshwater crayfish introductions in Africa', *Reviews in Fisheries Science & Aquaculture*, 1-21. https://doi.org/10.1080/23308249.2020.1802405.

Lodge, D.M., Deines, A., Gherardi, F., Yeo, D.C.J., Arcella, T., Baldridge, A.K., Branes, M.A., Chadderton, W.L., Feder, J.L., Grantz, C.A., Howard, G.W., Jerdde, C.L., Peters, J.A., Sargent, L.W., Turner, C.R.,

Wittmann, M.E., Zeng, Y., 2012, 'Global introductions of crayfishes: evaluating the impact of species invasions on ecosystem services', *Annual Review of Ecology Evolution Systematics*, 43, 449–472. https://doi.org/10.1146/annurev-ecolsys-111511-103919.

| **LIK2 Likelihood of entry via human aided primary pathways** | |
|---|---|
| Response: Fairly probable (p = 0.5) | Confidence: Medium |
| Rationale: *Faxonius virilis* used as bait and often stocked as forage in the USA (Larson & Olden 2011, Kilian et al. 2012). It is also known to be present in the pet trade industry in Netherlands and the United Kingdom (Chucholl 2013; Faulkes 2015). It is likely that it could be present in the pet trade in South Africa, but this needs to be assessed. | |
| References: | |

Chucholl, C., 2013, 'Invaders for sale: trade and determinants of introduction of ornamental freshwater crayfish', *Biological Invasions* 15, 125–141. https://doi.org/10.1007/s10530-012-0273-2.
Faulkes, Z., 2015, 'The global trade in crayfish as pets', *Crustacean Research*, 44, 75–92. https://doi.org/10.18353/crustacea.44.0_75.
Kilian, J.V., Klauda, R.J., WIdman, S., Kashiwagi, M., Bourquin, R., Weglein, S. & Schuster, J., 2012, 'An assessment of a bait industry and angler behavior as a vector of invasive species', *Biological Invasions*, 14(7), 1469-1481. https://doi.org/10.1007/s10530-012-0173-5.
Larson, E.R. & Olden, J.D., 2011, 'The state of crayfish in the Pacific Northwest', *Fisheries*, 36(2), 60-73. https://doi.org/10.1577/03632415.2011.10389069

| **LIK3 Habitat suitability** | |
|---|---|
| Response: Fairly probable (p = 0.5) | Confidence: Medium |
| Rationale: *Faxonius virilis* may inhabit rivers, streams, lakes, marshes and ponds that are permanent and well-oxygenated (GISD 2015). They prefer warm waters of moderate turbidity with cobble or rocky substrates and abundant logs, rocks, vegetation, and other debris to use as refuge (Soes & Koese 2010). These habitats are available is South Africa | |
| References: | |

Global Invasive Species Database (GISD) 2015. Species profile *Orconectes virilis*. Available from: http://www.iucngisd.org/gisd/species.php?sc=218 [Accessed 03 July 2018].
Soes, D.M. & Koese, B., 2010, 'Invasive crayfish in the Netherlands: A preliminary risk analysis. *Interim report, Bureau Waardenburg bv, Stichting EIS-Nederland'*, Invasive Alien Species Team, Waardenburg.

| **LIK4 Climate suitability** | |
|---|---|
| Response: Fairly probable (p = 0.5) | Confidence: Medium |
| Rationale: *Faxonius virilis* can survive a temperature range of 0-32°C and has a preferred temperature range of 24-25°C (GISD 2015). Its movement is halted at temperatures is below 10°C (GISD 2015). It known to occur in several climate types that include Tropical and dry savanna, mediterranean, humid subtropical that are known to occur in South Africa (CABI 2024). | |
| References: | |

CABI, 2024. *Faxonius virilis* (virile crayfish). Invasive Species Compendium, CABI Publishing, Wallingford, UK. https://www.cabidigitallibrary.org/doi/full/10.1079/cabicompendium.72034 (accessed 10 July 2024)
Global Invasive Species Database (GISD) 2015. Species profile *Orconectes virilis*. Available from: http://www.iucngisd.org/gisd/species.php?sc=218 [Accessed 03 July 2018]

| **LIK5 Unaided secondary (dispersal) pathways** | |
|---|---|
| Response: Very unlikely (p = 0.0027) | Confidence: Medium |
| Rationale: *Faxinous virilis* can disperse naturally in aquatic environments but has limited ability to disperse overland (Hamr 2002). There are no known populations of *F. virilis* present in countries neighbouring South Africa that could lead to secondary dispersal events through connected waterways (Nunes et al. 2017; Madzivanzira et al. 2020). | |

References:

Hamr, P., 2002, *Orconectes*. In: *Biology of freshwater crayfish*, [ed. by Holdich, D. M.]. Oxford, UK: Blackwell Science. 585-608.
Madzivanzira, T.C., South, J., Wood, L.E., Nunes, A.L. & Weyl, O.L.F., 2020, 'A review of freshwater crayfish introductions in Africa', *Reviews in Fisheries Science & Aquaculture* 1-21. https://doi.org/10.1080/23308249.2020.1802405
Nunes, A.L., Zengeya, T.A., Measey, G.J. & Weyl, O.L.F., 2017b, 'Freshwater crayfish invasions in South Africa: past, present and potential future', *African Journal of Aquatic Science* 42, 309–323. https://doi.org/10.2989/16085914.2017.1405788

| **LIK6 Human aided secondary (dispersal) pathways** | |
|---|---|
| Response: Fairly probable (p = 0.5) | Confidence: Medium |
| Rationale: Intentional release into the wild through the pet/ aquarium/ terrarium species and live bait pathways remains a concern. In areas of introduction, *Faxonius virilis* has been released into the wild by humans as unwanted pets and as bait (Larson and Olden 2011, Kilian et al. 2012; Chucholl 2013; Faulkes 2015) | |

References:

Chucholl, C., 2013, 'Invaders for sale: trade and determinants of introduction of ornamental freshwater crayfish', *Biological Invasions*, 15, 125–141. https://doi.org/10.1007/s10530-012-0273-2.
Faulkes, Z., 2015, 'The global trade in crayfish as pets', *Crustacean Research*, 44, 75–92. https://doi.org/10.18353/crustacea.44.0_75.
Kilian, J.V., Klauda, R.J., WIdman, S., Kashiwagi, M., Bourquin, R., Weglein, S. & Schuster, J., 2012, 'An assessment of a bait industry and angler behavior as a vector of invasive species', *Biological Invasions*, 14(7), 1469-1481. https://doi.org/10.1007/s10530-012-0173-5.
Larson, E.R. & Olden, J.D., 2011, 'The state of crayfish in the Pacific Northwest', *Fisheries*, 36(2), 60-73. https://doi.org/10.1577/03632415.2011.10389069.

**3. Consequences**

| **CON1 Environmental impact** | |
|---|---|
| **CON1a: Competition** | |
| Response: Major (MR) | Confidence: Medium |
| Rationale: *Faxonius virilis* has being implicated in the decline, displacement and local extirpation of native macroinvertebrates, crayfishes, snails, frogs and fish in the USA through competition and predation (Hanson | |

| |
|---|
| et al. 1990; Rogowski & Stockwell 2006, Philips et al. 2009; Loughman & Welsh 2010; Hale et al. 2016; Shaw et al. 2021; USGS-NAS 2023; GISD 2023). |
| References:<br><br>GISD, 2023. Species profile: *Orconectes virilis*. Global Invasive Species Database. http://www.iucngisd.org/gisd/species.php?sc=218<br>Hale, P., Wilson, J., Loughman, Z. & Henkanaththegedara, S., 2016, 'Potential impacts of invasive crayfish on native crayfish: insights from laboratory experiments', *Aquatic Invasions* 11(4), 451-458. http://dx.doi.org/10.3391/ai.2016.11.4.09.<br>Hanson, J.M., Chambers, P.A. & Prepas, E.E., 1990, 'Selective foraging by the crayfish *Orconectes virilis* and its impact on macroinvertebrates', *Freshwater Biology*, 24, 69-80. https://doi.org/10.1111/j.1365-2427.1990.tb00308.x<br>Loughman, Z. J. & Welsh, S.A., 2010, 'Distribution and conservation standing of West Virginia crayfishes', *Southeastern Naturalist*, 9(3), 63-78. https://doi.org/10.1656/058.009.s304.<br>Phillips, I.D., Vinebrooke, R.D. & Turner, M.A., 2009, 'Ecosystem consequences of potential range expansions of *Orconectes virilis* and *Orvconectes rusticus* crayfish in Canada- a review. *Environmental Review*, 17, 235-248. https://doi.org/10.1139/A09-011<br>Rogowski, D.L. & Stockwell, C.A., 2006, 'Assessment of potential impacts of exotic species on populations of a threatened species, white sands pupfish, *Cyprinodon Tularosa*', *Biological Invasions*, 8(1), 79-87.<br>Shaw, P.C., Larson, E.R. & Billman, E.J., 2021, 'Invasion of Virile Crayfish *Faxonius virilis* (Hagen 1870) in the Lower Henrys Fork Drainage, Idaho', *Northwest Science* 95(1), 106-113. https://doi.org/10.3955/046.095.0107.<br>USGS-NAS, 2023. *Faxonius virilis* (Hagen, 1870).US Geological Survey, Nonindigenous Aquatic Species database. https://nas.er.usgs.gov/queries/FactSheet.aspx?speciesID=215 |

| **CON1b: Predation** | |
|---|---|
| Response: Major (MR) | Confidence: Medium |
| Rationale: *Faxonius virilis* has being implicated in the decline, displacement and local extirpation of native macroinvertebrates, crayfishes, snails, frogs and fish in the USA through competition and predation (Hanson et al. 1990; Rogowski & Stockwell 2006, Philips et al. 2009; Loughman & Welsh 2010; Hale et al. 2016; Shaw et al. 2021; USGS-NAS 2023; GISD 2023). | |
| References:<br>GISD, 2023. Species profile: *Orconectes virilis*. Global Invasive Species Database. http://www.iucngisd.org/gisd/species.php?sc=218<br>Hale, P., Wilson, J., Loughman, Z. & Henkanaththegedara, S., 2016, 'Potential impacts of invasive crayfish on native crayfish: insights from laboratory experiments', *Aquatic Invasions* 11(4), 451-458. http://dx.doi.org/10.3391/ai.2016.11.4.09.<br>Hanson, J.M., Chambers, P.A. & Prepas, E.E., 1990, 'Selective foraging by the crayfish *Orconectes virilis* and its impact on macroinvertebrates', *Freshwater Biology*, 24, 69-80.<br>Loughman, Z. J. & Welsh, S.A., 2010, 'Distribution and conservation standing of West Virginia crayfishes', *Southeastern Naturalist*, 9(3), 63-78. https://doi.org/10.1656/058.009.s304.<br>Phillips, I.D., Vinebrooke, R.D. & Turner, M.A., 2009, 'Ecosystem consequences of potential range expansions of *Orconectes virilis* and *Orvconectes rusticus* crayfish in Canada- a review. *Environmental Review*, 17, 235-248. https://doi.org/10.1656/058.009.s304.<br>Rogowski, D.L. & Stockwell, C.A., 2006, 'Assessment of potential impacts of exotic species on populations of a threatened species, white sands pupfish, *Cyprinodon Tularosa*', *Biological Invasions*, 8(1), 79-87. https://doi.org/10.1007/s10530-005-0238-9. | |

| |
|---|
| Shaw, P.C., Larson, E.R. & Billman, E.J., 2021, 'Invasion of Virile Crayfish *Faxonius virilis* (Hagen 1870) in the Lower Henrys Fork Drainage, Idaho', *Northwest Science* 95(1), 106-113. https://doi.org/10.3955/046.095.0107.<br>USGS-NAS, 2023. *Faxonius virilis* (Hagen, 1870).US Geological Survey, Nonindigenous Aquatic Species database. https://nas.er.usgs.gov/queries/FactSheet.aspx?speciesID=215 |

| **CON1c: Hybridisation** | |
|---|---|
| Response: Data Deficient (DD) | Confidence: High |
| Rationale: Hybridisation is unlikely in South Africa because there are no native freshwater crayfish species (Nunes et al. 2017) | |
| References:<br><br>Nunes, A.L., Zengeya, T.A., Measey, G.J. & Weyl, O.L.F., 2017, 'Freshwater crayfish invasions in South Africa: past, present and potential future', *African Journal of Aquatic Science* 42, 309–323. https://doi.org/10.2989/16085914.2017.1405788. | |

| **CON1d: Transmission of disease** | |
|---|---|
| Response: Minor (MN) | Confidence: Medium |
| Rationale: *Faxinous virilis* is a known vector of pathogen such as the crayfish plague agent, *Aphanomyces astaci* (Ahern et al. 2008; Longshaw 2011). There are no native crayfish species in South Africa, but crayfish plague pathogen has been shown to infect other freshwater decapods such as native potamonautid crabs (Svoboda et al. 2014) | |
| References:<br>Ahern, D., England, J. & Ellis, A., 2008, 'The virile crayfish, *Orconectes virilis* (Hagen, 1870) (Crustacea: Decapoda: Cambaridae), identified in the UK. Invasive Species in Inland Waters of Europe and North America: Distribution and Impacts', *Aquatic Invasions*, 3(1), 102-104. https://doi.org/10.3391/ai.2008.3.1.18.<br>Longshaw, M., 2011, 'Diseases of crayfish: A review', *Journal of Invertebrate Pathology* 106, 54–70. https://doi.org/10.1016/j.jip.2010.09.013.<br>Svoboda, J., Strand, D.A., Vrålstad, T., Grandjean, F., Edsman, L., Kozák, P., et al., 2014, 'The crayfish plague pathogen can infect freshwater–inhabiting crabs', *Freshwater Biology* 59, 918–929, https://doi.org/10.1111/fwb.12315 | |

| **CON1e: Parasitism** | |
|---|---|
| Response: Data Deficient (DD) | Confidence: |
| Rationale: *Faxinous virilis* is not a parasitic taxon | |
| References: | |

| **CON1f: Poisoning/toxicity** | |
|---|---|
| Response: Data Deficient (DD) | Confidence: |
| Rationale: No documented information available to assess the level of impact. | |
| References: | |

| **CON1g: Bio-fouling or other direct physical disturbance** | |
|---|---|
| Response: Data Deficient (DD) | Confidence: |

| | |
|---|---|
| Rationale: No documented information available to assess the level of impact. | |
| References: | |
| **CON1h: Grazing/herbivory/browsing** | |
| Response: Moderate (MO) | Confidence: Low |
| Rationale: *Faxinous virilis is* omnivorous species, it is likely to cause a decrease in macrophyte cover, macroinvertebrates abundance and diversity, changing trophic interaction and community composition (Chambers et al. 1990; Roessink et al. 2017) | |
| References:<br><br>Chambers, P.A., Hanson, J.M. & Burke, J.M. 1990. The impact of the crayfish *Orconectes virilis* on aquatic macrophytes. *Freshwater Biology* 24:81-91. https://doi.org/10.1111/j.1365-2427.1990.tb00309.x.<br>Roessink, I., Gylstra, R., Heuts, P. G. M., Specken, B. & Ottburg, F., 2017, 'Impact of invasive crayfish on water quality and aquatic macrophytes in the Netherlands', *Aquatic Invasions* 12(3), 397-404. https://doi.org/10.3391/ai.2017.12.3.12. | |
| **CON1i: Chemical, physical or structural impact on ecosystem** | |
| Response: Minor (MN) | Confidence: Low |
| Rationale: Excessive grazing by *F. virilis* has led to habitat alteration through decreased macrophytes abundance and increased turbidity in invaded systems (Chambers et al. 1990; Roessink et al. 2017) | |
| References:<br><br>Chambers, P.A., Hanson, J.M., Burke, J.M., Prepas, E.E., 1990. The impact of the crayfish *Orconectes virilis* on aquatic macrophytes. *Freshwater Biology*, 24:81-91. https://doi.org/10.1111/j.1365-2427.1990.tb00309.x.<br>Roessink, I., Gylstra, R., Heuts, P. G. M., Specken, B., and F. Ottburg. 2017. Impact of invasive crayfish on water quality and aquatic macrophytes in the Netherlands. *Aquatic Invasions* 12(3): 397-404. https://doi.org/10.3391/ai.2017.12.3.12. | |
| **CON1k: Indirect impacts through interactions with other species** | |
| Response: Data Deficient (DD) | Confidence: |
| Rationale: No documented information available to assess the level of impact. | |
| References: | |
| **CON1 Maximum environmental impact** | |
| Response: Major (MR) | Confidence: Medium |
| Rationale: *Faxonius virilis* has being implicated in the decline, displacement and local extirpation of native macroinvertebrates, crayfishes, snails, frogs and fish in the USA through competition and predation (Hanson et al. 1990; Rogowski & Stockwell 2006, Philips et al. 2009; Loughman & Welsh 2010; Hale et al. 2016; Shaw et al. 2021; USGS-NAS 2023; GISD 2023). | |
| References:<br><br>GISD, 2023. Species profile: *Orconectes virilis*. Global Invasive Species Database. http://www.iucngisd.org/gisd/species.php?sc=218 | |

Hale, P., Wilson, J., Loughman, Z. & Henkanaththegedara, S., 2016, 'Potential impacts of invasive crayfish on native crayfish: insights from laboratory experiments', *Aquatic Invasions* 11(4), 451-458. https://doi.org/10.3391/ai.2016.11.4.09.

Hanson, J.M., Chambers, P.A. & Prepas, E.E., 1990, 'Selective foraging by the crayfish *Orconectes virilis* and its impact on macroinvertebrates', *Freshwater Biology*, 24, 69-80.

Loughman, Z. J. & Welsh, S.A., 2010, 'Distribution and conservation standing of West Virginia crayfishes', *Southeastern Naturalist*, 9(3), 63-78. https://doi.org/10.1656/058.009.s304.

Phillips, I.D., Vinebrooke, R.D. & Turner, M.A., 2009, 'Ecosystem consequences of potential range expansions of *Orconectes virilis* and *Orvconectes rusticus* crayfish in Canada- a review. *Environmental Review*, 17, 235-248. https://doi.org/10.1656/058.009.s304.

Rogowski, D.L. & Stockwell, C.A., 2006, 'Assessment of potential impacts of exotic species on populations of a threatened species, white sands pupfish, *Cyprinodon Tularosa*', *Biological Invasions*, 8(1), 79-87. https://doi.org/10.1007/s10530-005-0238-9.

Shaw, P.C., Larson, E.R. & Billman, E.J., 2021, 'Invasion of Virile Crayfish *Faxonius virilis* (Hagen 1870) in the Lower Henrys Fork Drainage, Idaho', *Northwest Science* 95(1), 106-113. https://doi.org/10.3955/046.095.0107.

USGS-NAS, 2023. *Faxonius virilis* (Hagen, 1870).US Geological Survey, Nonindigenous Aquatic Species database. https://nas.er.usgs.gov/queries/FactSheet.aspx?speciesID=215

| **CON2 Socio-economic impact** | |
|---|---|
| **CON2a: Safety** | |
| Response: Data Deficient (DD) | Confidence: |
| Rationale: No documented information available to assess the level of impact. | |
| References: | |
| **CON2b: Material and immaterial assets** | |
| Response: Minor (MN) | Confidence: Low |
| rationale: *Faxonius virilis* has been implicated in causing damage to levee and irrigation systems due to its burrowing activity ((GISD, 2023; Nemesis, 2023) | |
| References:<br><br>GISD, 2023. Species profile: *Orconectes virilis*. Global Invasive Species Database.http://www.iucngisd.org/gisd/species.php?sc=218<br>Nemesis, 2023. *Faxonius virilis*, Crustaceans-Crayfish: virile crayfish. Edgewater, MD: National Estuarine and Marine Exotic Species Information System (NEMESIS), Smithsonian Environmental Research Center. https://invasions.si.edu/nemesis/species_summary/97425 | |
| **CON2c: Health** | |
| Response: Data Deficient (DD) | Confidence: |
| Rationale: No documented information available to assess the level of impact. | |
| References: | |
| **CON2d: Social, spiritual and cultural relations** | |
| Response: Data Deficient (DD) | Confidence: |

| Rationale: |
|---|
| References: |

| CON2 Maximum socio-economic impact | |
|---|---|
| Response: Minor (MN) | Confidence: Low |
| rationale: *Faxonius virilis* has been implicated in causing damage to levee and irrigation systems due to its burrowing activity ((GISD, 2023; Nemesis, 2023) | |
| References:<br><br>GISD, 2023. Species profile: *Orconectes virilis*. Global Invasive Species Database.http://www.iucngisd.org/gisd/species.php?sc=218<br>Nemesis, 2023. *Faxonius virilis*, Crustaceans-Crayfish: virile crayfish. Edgewater, MD: National Estuarine and Marine Exotic Species Information System (NEMESIS), Smithsonian Environmental Research Center. https://invasions.si.edu/nemesis/species_summary/97425 | |

| CON3 Closely related species' environmental impact | |
|---|---|
| Response: Not scored as CON1 scored | Confidence: |
| Rationale: There is information to score CON1, so CON3 does not need to be scored as per the framework. | |
| References: | |

| CON4 Closely related species' socio-economic impact | |
|---|---|
| Response: Not scored as CON2 scored | Confidence: |
| Rationale: There is information to score CON2, so CON4 does not need to be scored as per the framework. | |
| References: | |

| CON5 Potential impact | |
|---|---|
| Response: Major (MR) | Confidence: High |
| Rationale: *Faxonius virilis* has being implicated in the decline, displacement and local extirpation of native macroinvertebrates, crayfishes, snails, frogs and fish in the USA through competition and predation (Hanson et al. 1990; Rogowski & Stockwell 2006, Philips et al. 2009; Loughman & Welsh 2010; Hale et al. 2016; Shaw et al. 2021; USGS-NAS 2023; GISD 2023). It is likely to cause similar impacts if introduced in South Africa. | |
| References:<br>GISD, 2023. Species profile: *Orconectes virilis*. Global Invasive Species Database. http://www.iucngisd.org/gisd/species.php?sc=218<br>Hale, P., Wilson, J., Loughman, Z. & Henkanaththegedara, S., 2016, 'Potential impacts of invasive crayfish on native crayfish: insights from laboratory experiments', *Aquatic Invasions* 11(4), 451-458. https://doi.org/10.3391/ai.2016.11.4.09.<br>Hanson, J.M., Chambers, P.A. & Prepas, E.E., 1990, 'Selective foraging by the crayfish *Orconectes virilis* and its impact on macroinvertebrates', *Freshwater Biology*, 24, 69-80. | |

Loughman, Z. J. & Welsh, S.A., 2010, ‘Distribution and conservation standing of West Virginia crayfishes’, *Southeastern Naturalist*, 9(3), 63-78. https://doi.org/10.1656/058.009.s304.
Phillips, I.D., Vinebrooke, R.D. & Turner, M.A., 2009, ‘Ecosystem consequences of potential range expansions of *Orconectes virilis* and *Orvconectes rusticus* crayfish in Canada- a review. *Environmental Review*, 17, 235-248. https://doi.org/10.1656/058.009.s304.
Rogowski, D.L. & Stockwell, C.A., 2006, ‘Assessment of potential impacts of exotic species on populations of a threatened species, white sands pupfish, *Cyprinodon Tularosa*’, *Biological Invasions*, 8(1), 79-87. https://doi.org/10.1007/s10530-005-0238-9.
Shaw, P.C., Larson, E.R. & Billman, E.J., 2021, ‘Invasion of Virile Crayfish *Faxonius virilis* (Hagen 1870) in the Lower Henrys Fork Drainage, Idaho’, *Northwest Science* 95(1), 106-113. https://doi.org/10.3955/046.095.0107.
USGS-NAS, 2023. *Faxonius virilis* (Hagen, 1870).US Geological Survey, Nonindigenous Aquatic Species database. https://nas.er.usgs.gov/queries/FactSheet.aspx?speciesID=215.

**4. Management**

| **MAN1 What is the feasibility to stop future immigration?** | |
|---|---|
| Response: High | Confidence: Medium |
| Rationale: *Faxonius virilis* is absent in neighbouring countries, so unaided immigration is unlikely. Moreover, it should be possible to regulate intentional introductions through legislative regulations and having an effective border surveillance and monitoring strategy (Zengeya & Wilson 2020; Wilson and Kumschick 2024). | |
| References:<br>Zengeya, T.A. & Wilson, J.R., (eds.) 2020, ‘*The status of biological invasions and their management in South Africa in 2019*’, pp.71, South African National Biodiversity Institute, Kirstenbosch and DSI-NRF Centre of Excellence for Invasion Biology, Stellenbosch. http://dx.doi.org/10.5281/zenodo.3947613.<br>Wilson, J. R., & Kumschick, S., 2024, ‘The regulation of alien species in South Africa’, *South African Journal of Science*, 120(5/6). https://doi.org/10.17159/sajs.2024/17002. | |

| **MAN2 Benefits of the Taxon** | |
|---|---|
| **MAN2a Socio-economic benefits of the *Taxon*** | |
| Response: Low | Confidence: Medium |
| Rationale: *Faxonius virilis* used as bait and often stocked as forage in the USA (Larson and Olden 2011, Kilian et al. 2012). It is also known to be present in the pet trade industry in Netherlands and the United Kingdom (Chucholl 2013; Faulkes 2015). No socio-economic benefits have been recorded to date in South Africa and even if *F. virilis* is present the benefits are likely to be low (Madzivanzira et al. 2021). | |
| References:<br><br>Chucholl, C., 2013, ‘Invaders for sale: trade and determinants of introduction of ornamental freshwater crayfish’, *Biological Invasions* 15, 125–141. https://doi.org/10.1007/s10530-012-0273-2.<br>Faulkes, Z., 2015, ‘The global trade in crayfish as pets’, *Crustacean Research*, 44, 75–92. https://doi.org/10.18353/crustacea.44.0_75.<br>Kilian, J.V., Klauda, R.J.; WIdman, S., Kashiwagi, M.,, Bourquin, R., Weglein, S. & Schuster, J., 2012, ‘An assessment of a bait industry and angler behavior as a vector of invasive species’, *Biological Invasions*, 14(7), 1469-1481. https://doi.org/10.1007/s10530-012-0173-5. | |

| | |
|---|---|
| Larson, E.R. & Olden, J.D., 2011, 'The state of crayfish in the Pacific Northwest', *Fisheries*, 36(2), 60-73. https://doi.org/10.1577/03632415.2011.10389069.<br>Madzivanzira, T.C., South, J., Wood, L.E., Nunes, A.L. & Weyl, O.L.F., 2020, 'A review of freshwater crayfish introductions in Africa', *Reviews in Fisheries Science & Aquaculture* 1-21. https://doi.org/10.1080/23308249.2020.1802405 | |
| **MAN2b Environmental benefits of the *Taxon*** | |
| Response: None | Confidence: |
| Rationale: No documented information available to assess the level of impact. | |
| References: | |

| | |
|---|---|
| **MAN3 Ease of management** | |
| **MAN3a How accessible are populations?** | |
| Response: 2 (difficulty to access) | Confidence: High |
| Rationale: There are no locality records for *Faxonius virilis* in South Africa (Madzivanzira et al. 2021; Nunes et al. 2017). However, given that it is an aquatic species, accessibility is likely to be difficult. | |
| References:<br>Madzivanzira, T.C., South, J. & Weyl, O.L.F., 2021, 'Invasive crayfish outperform Potamonautid crabs at higher temperatures', *Freshwater Biology* 66, 978–991. https://doi.org/10.1111/fwb.13691<br>Nunes, A.L., Zengeya, T.A., Measey, G.J. & Weyl, O.L.F., 2017, 'Freshwater crayfish invasions in South Africa: past, present and potential future', *African Journal of Aquatic Science* 42, 309–323, https://doi.org/10.2989/16085914.2017.1405788. | |
| **MAN3b Is detectability critically time-dependent?** | |
| Response: 2 (yes) | Confidence: High |
| Rationale: *Faxonius virilis* is a small, largely nocturnal freshwater species that is difficult to detect (Hamr 2002). | |
| References:<br><br>Hamr, P., 2002. *Orconectes*. In: *Biology of freshwater crayfish*, [ed. by Holdich, D. M.]. Oxford, UK: Blackwell Science. 585-608. | |
| **MAN3c Time to reproduction** | |
| Response: 1 (1-3 years) | Confidence: High |
| Rationale: *Faxonius virilis* reach sexual maturity in one-two years (Corey 1987; Hamr 2002). | |
| References:<br><br>Corey, S., 1987, 'Comparitive fecundity of four species of crayfish in southwestern Ontario, Canada (Decapoda, Astacidea)', *Crustaceana*, 52, 276-286. https://doi.org/10.1163/156854087X00510.<br>Hamr, P., 2002. *Orconectes*. In: *Biology of freshwater crayfish*, [ed. by Holdich, D. M.]. Oxford, UK: Blackwell Science. 585-608. | |
| **MAN3d Propagule persistence** | |
| Response: 0 (<1 year) | Confidence: High |
| Rationale: *Faxonius virilis* are highly fecund, a female produces up to 310 eggs, is able to store sperm and there is some parental care of eggs and hatchlings (Hamr 2002; Corey 1987). | |
| References:<br>Corey, S., 1987, 'Comparitive fecundity of four species of crayfish in southwestern Ontario, Canada (Decapoda, Astacidea)', *Crustaceana*, 52, 276-286. https://doi.org/10.1163/156854087X00510.<br>Hamr, P., 2002. *Orconectes*. In: *Biology of freshwater crayfish*, [ed. by Holdich, D. M.]. Oxford, UK: Blackwell Science. 585-608. | |
| **MAN3 Ease of management** | |
| Response: Medium | Confidence: High |

| |
|---|
| Rationale: *Faxonius virilis* is a small, largely nocturnal freshwater species that is difficult to detect, it reaches sexual maturity in one-two years, and is highly fecund (up to 310 eggs) (Hamr 2002). |
| References:<br><br>Hamr, P., 2002. *Orconectes*. In: *Biology of freshwater crayfish*, [ed. by Holdich, D. M.]. Oxford, UK: Blackwell Science. 585-608. |

| **MAN4 Has the feasibility of eradication been evaluated?** | |
|---|---|
| Response: No | Confidence: Low |
| Rationale: There are no known population of *F. virilis* in South Africa, therefore eradication feasibility has not been no evaluated (Nunes et al. 2017) but see MAN5 for summary of control efforts used to manage invasive crayfish populations. | |
| References:<br><br>Nunes, A.L., Zengeya, T.A., Measey, G.J. & Weyl, O.L.F., 2017, 'Freshwater crayfish invasions in South Africa: past, present and potential future', *African Journal of Aquatic Science* 42, 309–323, https://doi.org/10.2989/16085914.2017.1405788. | |

| **MAN5 Control options and monitoring approaches available for the *Taxon*** |
|---|
| Response: Several methods can be used to monitor and control invasive crayfish populations including mechanical trapping (e.g., Barkhuizen et al. 2022; Madzivanzira et al. 2021), electrofishing (Nunes et al. 2017), use of biocides, and biological control (Aquiloni et al 2010; Stebbing et al. 2014). However, there are several issues that make management of crayfish invasions challenging (Gherardi et al. 2011). For example, the use of biocides is often effective for localised populations in dams and ponds (Peay et al. 2019) but they are non-specific and need to be applied with caution (Gherardi et al. 2011). Mechanical control methods are also effective in small discrete water bodies but are expensive and require commitment to follow on to complete eradication (Loureiro et al. 2018). In addition, alien crayfish species are regarded as potential conflict-generating species in South Africa and potential management interventions should consider the concerns of stakeholders to avoid conflict (Zengeya et al. 2017). |
| References:<br><br>Aquiloni, L., Brusconi, S., Cecchinelli, E., Tricarico, E., Mazza, G., Paglianti, A. & Gherardi, F. 2010, 'Biological control of invasive populations of crayfish: the European eel (*Anguilla anguilla*) as a predator of *Procambarus clarkii*', *Biological Invasions* 12 3817-3824. https://doi.org/10.1007/s10530-007-9094-0.<br>Barkhuizen, L.M, Madzivanzira, T.C., South, J., 2022, 'Population ecology of a wild population of red swamp crayfish *Procambarus clarkii* (Girard, 1852) in the Free State Province, South Africa and implications for eradication efforts', *BioInvasions Records* 11, 181–191. https://doi.org/10.3391/bir.2022.<br>Gherardi, F., Aquiloni, L., Diéguez-Uribeondo, J. & Tricarico E., 2011, 'Managing invasive crayfish: is there a hope? *Aquatic Sciences* 73, 185–200. https://doi.org/10.1007/s00027-011-0181-z.<br>Loureiro, T.G, Anastácio, P.M, de Siqueira Bueno, S.L. & Araujo, P.B., 2018, 'Management of invasive populations of the freshwater crayfish *Procambarus clarkii* (Decapoda, Cambaridae): test of a population-control method and proposal of a standard monitoring approach', *Environmental Monitoring Assessment* 190, 559, https://doi.org/10.1007/s10661-018-6942-6.<br>Madzivanzira, T.C., South, J., Nhiwatiwa, T. & Weyl, O.L.F., 2021, 'Standardisation of Australian redclaw crayfish *Cherax quadricarinatus* sampling gear in southern Africa', *Water SA* 47, 380–384. https://hdl.handle.net/10520/ejc-waters-v47-n3-a10. |

Nunes, A.L., Zengeya, T.A., Hoffman, A.C., Measey, G.J. & Weyl, O.L., 2017, 'Distribution and establishment of the alien Australian redclaw crayfish, *Cherax quadricarinatus*, in South Africa and Swaziland', *PeerJ* 5, 1-21. https://doi.org/10.7717/peerj.3135.
Peay, S., Johnsen, S.I, Bean, S.W., Dunn, A.M., Sandodden, R. & Edsman, L, 2019, 'Biocide treatment of invasive signal crayfish: Successes, failures and lessons learned', *Diversity* 11, 29. https://doi.org/10.3390/d11030029
Stebbing, P., Longshaw, M., Scott, A., 2014, 'Review of methods for the management of nonindigenous crayfish, with particular reference to Great Britain', *Ethology Ecology & Evolution* 26, 204-231. https://doi.org/10.1080/03949370.2014.908326.
Woodford, D.J., Ivey, P., Novoa, A., Shackleton, R., Richardson, D., Weyl, O., Van Wilgen, B. and Zengeya, T., 2017, 'Managing conflict-generating invasive species in South Africa: Challenges and trade-offs', *Bothalia* 47, a2160. https://doi. org/10.4102/abc.v47i2.2160.

| **MAN6 Any other management considerations to highlight?** (if yes, fill in Appendix MAN6) | |
|---|---|
| Response | No |

**5. Calculations**

**Likelihood = Fairly probable**

| Parameter | Likelihood | Stages | Final assessment |
|---|---|---|---|
| LIK1 | 0.0027 | P(entry) = 0.5 | P (invasion) =0.125 |
| LIK2 | 0.5 | | |
| LIK3 | 0.5 | P(establishment) = 0.5 | |
| LIK4 | 0.5 | | |
| LIK5 | 0.0027 | P (spread) = 0.5 | |
| LIK6 | 0.5 | | |

**Consequence = Major (MR)**

| Parameter | Mechanism/sector | Response |
|---|---|---|
| CON3a | Competition | MR |
| CON3b | Predation | MR |
| CON3c | Hybridisation | DD |
| CON3d | Disease transmission | MN |
| CON3e | Parasitism | DD |
| CON3f | Poisoning/toxicity | DD |
| CON3g | Bio-fouling or other direct physical disturbance | DD |
| CON3h | Grazing/herbivory/browsing | MO |

| | | |
|---|---|---|
| CON3i | Chemical, physical, structural impact | MN |
| CON3k | Indirect impacts through interactions with other species | DD |
| **CON1** | Maximum environmental impact | MR |
| CON4a | Safety | DD |
| CON4b | Material and immaterial assets | MN |
| CON4c | Health | DD |
| CON4d | Social, spiritual and cultural relations | DD |
| **CON2** | Maximum socio-economic impact | MN |
| CON3 | Maximum environmental impact (closely related taxa) | Not scored as CON1 scored |
| CON4 | Maximum socio-economic impact (closely related taxa) | Not scored as CON2 scored |
| **CON5** | Potential impact based on traits, experiments, or models | MR |

**Risk score: High**

| | | Consequences | | | | |
|---|---|---|---|---|---|---|
| | | MC | MN | MO | MR | MV |
| Likelihood | Extremely unlikely | low | low | low | medium | medium |
| | Very unlikely | low | low | low | medium | high |
| | Unlikely | low | low | medium | high | high |
| | Fairly probable | medium | medium | high | high | high |
| | Probable | medium | high | high | high | high |

**Ease of management: Medium**

| Parameter | Question | Response |
|---|---|---|
| MAN3a | How accessible are populations? | 2 |
| MAN3b | Is detectability critically time-dependent? | 2 |
| MAN3c | Time to reproduction | 1 |
| MAN3d | Propagule persistence | 0 |
| **MAN3** | **SUM** | 5 |

**Supplementary to add to answer sheet**
**Appendix BAC7**: Global alien range of *Faxonius virilis.* Map form CABI:
https://www.cabi.org/ISC/datasheet/72034

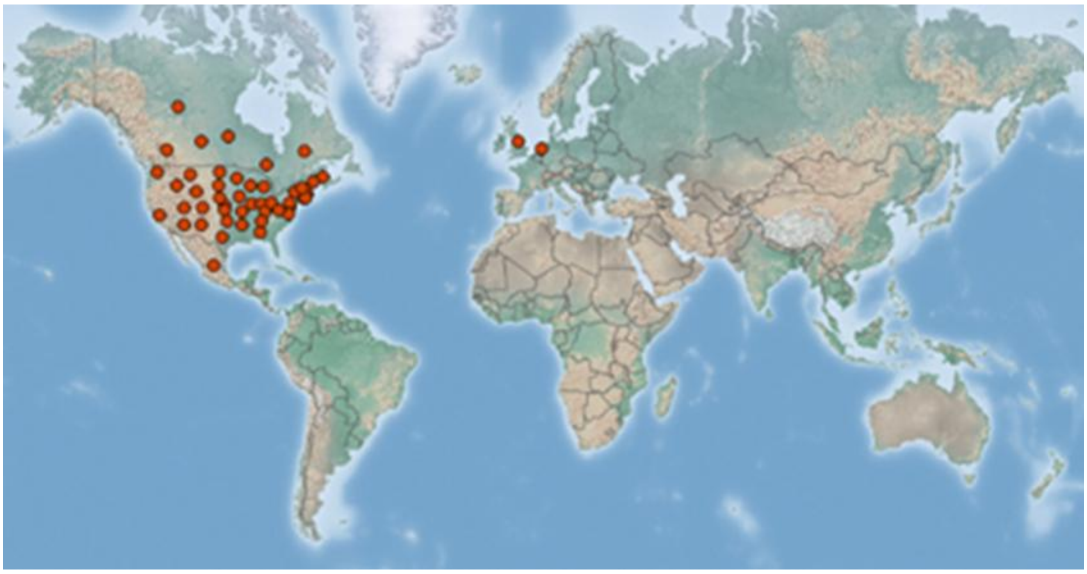

### 1.10 Risk analysis report for signal crayfish (*Pacifastacus leniusculus*)

| | |
|---|---|
| ***Taxon:*** *Pacifastacus leniusculus* (Dana, 1852) | ***Area:*** **South Africa** |
| **Compiled by:** Lee-Anne Botha | **Approved by:** |
| **Picture of *Taxon***<br>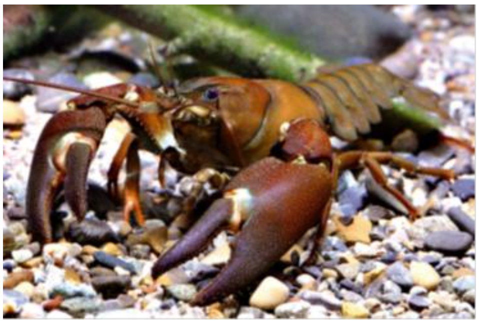<br>Photo: Holdich et al. 2009 | **Alien distribution map**<br>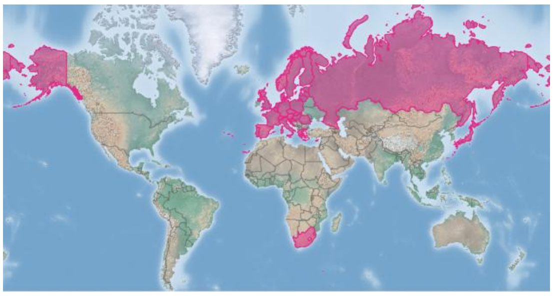<br>Reference: CABI (2022), https://doi/10.1079/cabicompendium.70581 |

| | |
|---|---|
| **Risk Assessment summary:**<br><br>Signal crayfish (*Pacifastacus leniusculus*) has been widely introduced for aquaculture and the global pet trade industry. There is anecdotal evidence that it might be present in South Africa but this needs to be verified. The are no known populations of *P. leniusculus* in neighbouring countries, therefore entry via unaided pathways is unlikely. The illegal pet trade industry remains a concern because *P. leniusculus* may be moved around by humans and there is a high risk of intentional release of unwanted pets into the wild. In the introduced range, *P. leniusculus* has been implicated in displacing native crayfish species through the transmission of crayfish plague and out-competing them for food and shelter. It has also been implicated in causing adverse impacts on macroinvertebrate communities through predation. Crayfish plague caused population crashes of native crayfish in Europe leading to a decline in economic and cultural benefits derived from the species. There are no native crayfish species in South Africa but the crayfish plague has been shown to infect other freshwater decapods and introduced crayfish have the potential to outcompete native decapods such as potamonautid crabs. | **Risk score:**<br>High |
| **Management summary:**<br><br>*Pacifastacus leniusculus* is a fairly small, largely nocturnal, burrowing freshwater species that is difficult to detect. Several methods have been used to control alien populations of *P. leniusculus* and these include mechanical removal using traps and electrofishing, biocides, and the release of sterile males. However, several issues make the management of crayfish invasions challenging. Biocides are effective for localized populations but can have non-target effects on native fauna. Mechanical controls are also effective in small discrete water bodies but are expensive, require a commitment to follow on to complete eradication, and can lead to unwanted community compensation mechanisms such as enhanced recruitment. *Pacifastacus leniusculus* is a popular species in the pet trade industry. The pet trade industry is generally difficult to regulate because trade restrictions imposed on species of conservation concern that are albeit popular and in high demand, often lead to illegal trading that is compounded | **Ease of management:**<br><br>Difficult |

| | |
|---|---|
| by a lack of reliable record-keeping of the type and number of organisms presently in the pet trade. | |
| **Recommendations:**<br><br>*Pacifastacus leniusculus* is recommended to be added to a future prohibited list because there are no known records in South Africa. The taxon should be included in a border surveillance and monitoring strategy to prevent entry into South Africa. In addition, surveys should be done to confirm which crayfish species are present in the pet trade in South Africa and if found to be present, it should be relisted as category 1a. | **Listing under NEM:BA A&IS lists of 2020:**<br><br>1a<br><br>**Recommended listing category:**<br><br>Future prohibited list |

#### 1. Background

| **BAC1 Name of assessor(s)** | |
|---|---|
| Name of lead assessor | Lee-Anne Botha |
| Additional assessor (1) | Tsungai Zengeya |
| Additional assessor (2) | Christian Chimimba |
| **BAC2 Contact details of assessor (s)** | |
| Lead assessor | Organisational affiliation: Department of Zoology and Entomology, University of Pretoria/South African National Biodiversity Institute (SANBI). |
| | |
| Additional assessor (1) | Organisational affiliation: Department of Zoology and Entomology, University of Pretoria and The South African National Biodiversity Institute (SANBI). |
| | |
| Additional assessor (2) | Organisational affiliation: Department of Zoology and Entomology, University of Pretoria |
| | |
| **BAC3 Name(s) and contact details of expert(s) consulted** | |
| Expert (1) | Name: |
| | email: |
| | Phone: |
| Expert (2) | Name: |
| | email: |
| | Phone: |
| Comments: none | |
| **BAC4 Scientific name of *Taxon* under assessment** | |
| *Taxon* name: *Pacifastacus leniusculus* | Authority: (Dana, 1852) |

Comments: *Pacifastacus leniusculus* is the valid name of the taxon under assessment (Integrated Taxonomic Information System 2023). The following subspecies have been listed for *P. leniusculus*: *Pacifastacus leniusculus klamathensis* (Stimpson, 1857), *Pacifastacus leniusculus leniusculus* (Dana, 1852) and *Pacifastacus leniusculus trowbridgii* (Stimpson, 1857) (Agerberg & Jansson 1995; Larson et al. 2012). However, ITIS (2023) lists these subspecies as invalid.

References:

Agerberg, A. & Jansson, H., 1995, 'Allozymic comparisons between three subspecies of the freshwater crayfish *Pacifastacus leniusculus* (Dana) and between populations introduced to Sweden', *Hereditas* 122. 33–39. https://doi.org/10.1111/j.1601-5223.1995.00033.x .

Integrated Taxonomic Information System (ITIS), 2023, '*Pacifastacus leniusculus*', viewed 27 July 2023, from https://www.itis.gov/servlet/SingleRpt/SingleRpt?search_topic=TSN&search_value=97326#null.

Larson, E.R., Abbott, C.L., Usio, N., Azuma, N., Wood, K.A., Herborg, L.M. & Olden, J.D., 2012, 'The signal crayfish is not a single species: Cryptic diversity and invasions in the Pacific Northwest range of *Pacifastacus leniusculus*', *Freshwater Biology* 57, 1823–1838. https://doi.org/10.1111/j.1365-2427.2012.02841.x.

**BAC5 Synonym(s) considered**

Synonyms: *Pacifastacus trowbridgi* (Stimpson, 1857), *Pacifastacus leniusculus leniusculus* (Dana, 1852), *Pacifastacus leniusculus klamathensis* (Stimpson, 1857), *Pacifastacus leniusculus trowbridgii* (Stimpson, 1857), *Astacus oreganus* Randall, 1840, *Astacus leniusculus* Dana, 1852, *Astacus klamathensis* Stimpson, 1857, and Astacus trowbridgii Stimpson, 1857.

Comments: The abovementioned synonyms of *Pacifastacus leniusculus* were considered for this assessment and are listed on the Integrated Taxonomy Information System (ITIS, 2023)

| | |
|---|---|
| References:<br>Integrated Taxonomic Information System (ITIS), 2023, ‘*Pacifastacus leniusculus*’, viewed 27 July 2023, from https://www.itis.gov/servlet/SingleRpt/SingleRpt?search_topic=TSN&search_value=97326#null. | |
| **BAC6 Common name(s) considered** | |
| Common names: signal crayfish \| Columbia river signal crayfish \| Klamath signal crayfish | |
| Comments: The abovementioned common names have been listed for *P. leniusculus* (Schuster, Taylor & Cardeiro 2010) and were considered for this assessment. | |
| References:<br>Schuster, G.A., Taylor, C.A. & Cordeiro, J., 2010, *Pacifastacus leniusculus. The IUCN Red List of Threatened Species 2010*: e.T153648A4526314. http://dx.doi.org/10.2305/IUCN.UK.2010-3.RLTS.T153648A4526314.en [Accessed 31 January 2019] | |
| **BAC7 What is the native range of the *Taxon*?** | |
| Response: North-western U.S.A. \| south-western Canada. | Confidence: High |
| Comments: *Pacifastacus leniusculus* is native to the north-western region of the USA and south-western Canada (GISD 2015; Lodge et al. 2012). | |
| References:<br>Global Invasive Species Database (GISD) 2015, Species profile *Pacifastacus leniusculus*. Available from: http://www.iucngisd.org/gisd/species.php?sc=725 [Accessed 02 July 2018]<br>Lodge, D.M., Deines, A., Gherardi, F., Yeo, D.C.J., Arcella, T., Baldridge, A.K., et al., 2012, ‘Global introductions of crayfishes: Evaluating the impact of species invasions on ecosystem services’, *Annual Review of Ecology, Evolution and Systematics* 43, 449-472. https://doi.org/10.1146/annurev-ecolsys-111511-103919. | |
| **BAC8 What is the global alien range of the *Taxon*?** | |
| Response: Europe \| Asia | Confidence: High |
| Comments: *Pacifastacus leniusculus* has been introduced into the following countries: Austria; Belgium; Cyprus; Croatia, Denmark; Finland; France; Germany; Greece; Hungary, Italy; Japan; Latvia; Lithuania; Luxembourg; Netherlands; Poland; Portugal; Russian Federation; Serbia, Spain; South Africa, Sweden; Switzerland; United Kingdom (Schuster et al. 2010, Kouba et al. 2014, Dragičević et al. 2020., Horvatovic et al. 2022., Ludanyi et al. 2022). | |

| |
|---|
| References:<br>Dragičević, P., Faller, M., Kutlesa, P. & Hudina, S. 2020."Update on the signal crayfish, *Pacifastacus leniusculus* (Dana, 1852) range expansion in Croatia: a 10-year report." *BioInvasions Records* 9(4): 793-807. https://doi.org/10.3391/bir.2020.9.4.13<br>Horvatovic, M.A., Rankovic, M., Milic, D., Miljanovic, B. & Bajic, A. 2022, 'The first record of signal crayfish, Pacifastacus leniusculus (Dana, 1852) and its projected expansion in Serbia under global climate change', *Biologia Serbica* 44(2). https://doi.org/10.5281/zenodo.7451154<br>Kouba, A., Petrusek, A., Kozák, P. 2014. "Continental-wide distribution of crayfish species in Europe: update and maps", *Knowledge and Management of Aquatic Ecosystems* 413, 05. https://doi.org/10.1051/kmae/2014007<br>Ludanyi, M., Peeters, E.T.H.M., Kiss, B., Gaspar, A., Roessink, I., Magura, T. & Muller, Z., 2022, 'The current status of Pacifastacus leniusculus (Dana, 1852) and their effect on aquatic macroinvertebrate communities in Hungarian watercourses'. *Aquatic Invasions* 17 (4): 543-559. https://doi.org/10.3391/ai.2022.17.4.05<br>Madzivanzira, T.C., South, J., Wood, L.E., Nunes, A.L. & Weyl, O.L.F., 2020, 'A review of freshwater crayfish introductions in Africa', *Reviews in Fisheries Science & Aquaculture*, 1-21, https://doi.org/10.1080/23308249.2020.1802405<br>Nunes, A.L., Zengeya, T.A., Measey, G.J. & Weyl, O.L.F., 2017, 'Freshwater crayfish invasions in South Africa: past, present and potential future', *African Journal of Aquatic Science* 42, 309–323. https://doi.org/10.2989/16085914.2017.1405788<br>Schuster, G.A., Taylor, C.A. & Cordeiro, J., 2010, *Pacifastacus leniusculus. The IUCN Red List of Threatened Species 2010*: e.T153648A4526314. http://dx.doi.org/10.2305/IUCN.UK.2010-3.RLTS.T153648A4526314.en_[Accessed 31 January 2019]. |

| | | |
|---|---|---|
| **BAC9 Geographic scope = the *Area* under consideration** | | |
| *Area* of assessment: The geographic scope of this assessment is limited to South Africa. | | |
| Comments: | | |
| **BAC10 Is the *Taxon* present in the *Area*?** | | |
| Response: No | | Confidence: Low |
| Comments: There is anecdotal evidence that *P. leniusculus* might be present in the pet trade in South Africa but this needs to be confirmed (de Moor 2002; Faulkes 2015, Mandivanzira et al. 2020) | | |
| References:<br>de Moor, I., 2002, 'Potential impacts of alien freshwater crayfish in South Africa', *African Journal of Aquatic Science* 27, 125–139. https://doi.org/10.2989/16085914.2002.9626584.<br>Faulkes, Z., 2015, 'The global trade in crayfish as pets', *Crustacean Research* 44, 75–92. https://doi.org/10.18353/crustacea.44.0_75.<br>Madzivanzira, T.C., South, J., Wood, L.E., Nunes, A.L. & Weyl, O.L.F., 2020, 'A review of freshwater crayfish introductions in Africa', *Reviews in Fisheries Science & Aquaculture*, 1-21, https://doi.org/10.1080/23308249.2020.1802405. | | |
| **BAC11 Availability of physical specimen** | | |
| Response: No specimens available | | Confidence in ID: |
| Herbarium or museum accession number: | | |
| References: | | |
| **BAC12 Is the *Taxon* native to the *Area* or part of the *Area*?** | | |
| The *Taxon* is native to (part of) the *Area*. | No | Confidence: High |
| The *Taxon* is alien in (part of) the *Area*. | Yes | Confidence: High |
| Comments: No native freshwater crayfish occur in South Africa (de Moor 2002; Nunes et al. 2017, Mandivanzira et al. 2020) | | |

References:
de Moor, I., 2002, ‘Potential impacts of alien freshwater crayfish in South Africa’, *African Journal of Aquatic Science* 27, 125–139. https://doi.org/10.2989/16085914.2002.9626584.
Nunes, A.L., Zengeya, T.A., Measey, G.J. & Weyl, O.L.F., 2017, ‘Freshwater crayfish invasions in South Africa: past, present and potential future’, *African Journal of Aquatic Science* 42, 309–323. https://doi.org/10.2989/16085914.2017.1405788.
Madzivanzira, T.C., South, J., Wood, L.E., Nunes, A.L. & Weyl, O.L.F., 2020, ‘A review of freshwater crayfish introductions in Africa’, *Reviews in Fisheries Science & Aquaculture*, 1-21, https://doi.org/10.1080/23308249.2020.1802405.

**BAC13 What is the *Taxon*’s introduction status in the *Area*?**

| | | |
|---|---|---|
| The *Taxon* is in cultivation/containment. | Don’t know | Confidence: Low |
| The *Taxon* is present outside of cultivation/containment. | Don’t know | Confidence: Low |
| The *Taxon* has established/naturalised. | No | Confidence: Low |
| The *Taxon* is invasive. | No | Confidence: High |

Comments: There are only records for the introduction of the following freshwater crayfish species in South Africa: *Cherax destructor*, *C. cainii*, *C. tenuimanus*, *C. quadricarinatus*, and *Procambarus clarkii* (de Moor 2002; Nunes et al. 2017). There is however some anecdotal that *Pacifastacus leniusculus* might be present in the pet trade in South Africa but this needs to be verified (Nunes et al. 2017, Mandivanzira et al. 2020).

References:
de Moor, I., 2002, ‘Potential impacts of alien freshwater crayfish in South Africa’, *African Journal of Aquatic Science* 27, 125–139. https://doi.org/10.2989/16085914.2002.9626584.
Nunes, A.L., Zengeya, T.A., Measey, G.J. & Weyl, O.L.F., 2017, ‘Freshwater crayfish invasions in South Africa: past, present and potential future’, *African Journal of Aquatic Science* 42, 309–323. https://doi.org/10.2989/16085914.2017.1405788.
Madzivanzira, T.C., South, J., Wood, L.E., Nunes, A.L. & Weyl, O.L.F., 2020, ‘A review of freshwater crayfish introductions in Africa’, *Reviews in Fisheries Science & Aquaculture*, 1-21, https://doi.org/10.1080/23308249.2020.1802405.

**BAC14 Primary (introduction) pathways**

| | | |
|---|---|---|
| Release | Intentional releases | Confidence: Medium |
| Escape | aquaculture/mariculture | Confidence: Medium |
| | pet/aquarium/terrarium species (including live food for such species) | Confidence: Medium |
| Contaminant | none | Confidence: High |
| Stowaway | none | Confidence: High |
| Corridor | none | Confidence: High |
| Unaided | natural dispersal across borders of invasive alien organisms | Confidence: High |

Comments: *Pacifastacus leniusculus* has been introduced to regions outside of the native range via aquaculture (Holdich 1993; Holdich, Reynolds, & Sibley 2009), and is also traded in the pet trade industry (Chucholl 2013; Faulkes 2015). Furthermore, *P. leniusculus* is capable of moving overland (Holdich et al. 2009; Hudina et al. 2010)

| |
|---|
| References:<br>Chucholl, C., 2013, 'Invaders for sale: Trade and determinants of introduction of ornamental freshwater crayfish', *Biological Invasions* 15, 125–141. https://doi.org/10.1007/s10530-012-0273-2<br>Faulkes, Z, 2015, 'The global trade in crayfish as pets', *Crustacean Research* 44, 75–92. https://doi.org/10.18353/crustacea.44.0_75.<br>Holdich, D.M., 1993, 'A review of astaciculture: Freshwater crayfish farming', *Aquatic Living Resources* 6, 307–317. https://doi.org/10.1051/alr:1993032<br>Holdich, D.M., Reynolds, J.D., & Sibley, P.J., 2009, 'A review of the ever increasing threat to European crayfish from non-indigenous crayfish species', *Knowledge and Management of Aquatic Ecosystems* 11, 394–395. https://doi.org/10.1051/kmae/2009025.<br>Hudina, S., Faller, M., Lucić, A., Klobučar, G. & Maguire, I., 2009, 'Distribution and dispersal of two invasive crayfish species in the Drava River basin, Croatia', *Knowledge and Management of Aquatic Ecosystems* 9, 394–395. https://doi.org/10.1051/kmae/2009023. |

**2. Likelihood**

| **LIK1 Likelihood of entry via unaided primary pathways** | |
|---|---|
| Response: Very unlikely (p = 0.0027) | Confidence: Medium |
| Rationale: There are no known populations of *P. leniusculus* in South Africa's neighbouring countries, therefore, entry into the country through unaided pathways such as connected waterways is very unlikely (Madzivanzira et al. 2020; Nunes et al. 2017). | |
| References:<br>Madzivanzira, T.C., South, J., Wood, L.E., Nunes, A.L. & Weyl, O.L.F., 2020, 'A review of freshwater crayfish introductions in Africa', *Reviews in Fisheries Science & Aquaculture*, 1-21, https://doi.org/10.1080/23308249.2020.1802405.<br>Nunes, A.L., Zengeya, T.A., Measey, G.J. & Weyl, O.L.F., 2017, 'Freshwater crayfish invasions in South Africa: past, present and potential future', *African Journal of Aquatic Science* 42, 309–323, https://doi.org/10.2989/16085914.2017.1405788 | |

| **LIK2 Likelihood of entry via human aided primary pathways** | |
|---|---|
| Response: Fairly probable (p = 0.5) | Confidence: High |
| Rationale: *Pacifastacus leniusculus* has been widely introduced for aquaculture and for pet trade. It is also known to escape from aquaculture facilities and has been intentionally released into the wild by humans as unwanted pets (Chucholl 2013; Faulkes 2015). Therefore, it is fairly probable (p = 0.5) that *P. leniusculus* can enter the country via human aided pathways. | |
| References:<br>Chucholl, C., 2013, 'Invaders for sale: Trade and determinants of introduction of ornamental freshwater crayfish', *Biological Invasions* 15, 125–141. https://doi.org/10.1007/s10530-012-0273-2.<br>Faulkes, Z, 2015, 'The global trade in crayfish as pets', *Crustacean Research* 44, 75–92. https://doi.org/10.18353/crustacea.44.0_75. | |

| **LIK3 Habitat suitability** | |
|---|---|
| Response: Fairly probable (p = 0.5) | Confidence: Low |

| |
|---|
| Rationale: *Pacifastacus leniusculus* occurs in both lentic and lotic habitats (Soes & Koese 2010). It can potential occur in in several river systems in the Western and Eastern Cape provinces that were predicted as climatic suitable for its establishment (Zhang et al. 2019). |
| References:<br><br>Zhang, Z., Capinha, C., Usio, N., Weterings, R., Liu, X., Li, Y., Landeria, J.M., Zhou, Q. & Yokota, M. 2019, 'Impacts of climate change on the global potential distribution of two notorious invasive crayfishes', *Freshwater Biology* 65, 353-365. https://doi.org/10.1111/fwb.13429. |

| **LIK4 Climate suitability** | |
|---|---|
| Response: Fairly probable (p = 0.5) | Confidence: Low |
| Rationale: *Pacifastacus leniusculus* is a cool water species and the optimal temperature for growth is approximately 15-20°C (Ackefors 1999). Several areas in Western and Eastern Cape provinces of South Africa were predicted as climatical suitable for *Pacifastacus leniusculus* (Zhang et al. 2019). | |
| References:<br><br>Ackefors, H. 1999. 'The positive effects of established crayfish introductions in Europe'. In F. Gherardi, & D. M. Holdich (Eds.), Crayfish in Europe as alien species. How to make the best of a bad situation? (pp. 49–61). Rotterdam, The Netherlands: A.A. Balkema.<br>Zhang, Z., Capinha, C., Usio, N., Weterings, R., Liu, X., Li, Y., Landeria, J.M., Zhou, Q. & Yokota, M., 2019, 'Impacts of climate change on the global potential distribution of two notorious invasive crayfishes', *Freshwater Biology* 65, 353-365. https://doi.org/10.1111/fwb.13429 | |

| **LIK5 Unaided secondary (dispersal) pathways** | |
|---|---|
| Response: Fairly probable (p = 0.5) | Confidence: Low |
| Rationale: *Pacifastacus leniusculus* can migrate short distances overland to colonise new areas (Holdich et al. 2009; Hudina et al. 2010). There are no known populations of *P. leniusculus* in South Africa (Madzivanzira et al. 2020), however, if it was introduced, unaided secondary is likely to occur. Therefore, the response has been set to fair probable (p = 0.5). | |
| References:<br>Hudina, S., Faller, M., Lucić, A., Klobučar, G. & Maguire, I., 2009, 'Distribution and dispersal of two invasive crayfish species in the Drava River basin, Croatia', *Knowledge and Management of Aquatic Ecosystems* 9, 394–395. https://doi.org/10.1051/kmae/2009023<br>Holdich, D.M., Reynolds, J.D., Souty-Grosset, C. & Sibley, P.J., 2009, 'A review of the ever increasing threat to European crayfish from non-indigenous crayfish species', *Knowledge and Management of Aquatic Ecosystems* 11, 1–46. https://doi.org/10.1051/kmae/2009025<br>Madzivanzira, T.C., South, J., Wood, L.E., Nunes, A.L. & Weyl O.L.F., 2020, 'A review of freshwater crayfish introductions in Africa', *Reviews in Fisheries Science & Aquaculture* 1–24. https://doi.org/10.1080/23308249.2020.1802405. | |

| **LIK6 Human aided secondary (dispersal) pathways** | |
|---|---|
| Response: Fairly probable (p = 0.5) | Confidence: Low |
| Rationale: *Pacifastacus leniusculus* is a popular species in the global pet trade and has also been intentionally be released into the wild as unwanted pets (Chucholl 2013; Faulkes 2015; Holdich et al. 2009). If it was | |

| |
|---|
| introduced in South Africa for aquaculture or for the pet trade, then there is a strong chance it would be spread around the country for the same purposes. |
| References:<br>Chucholl, C., 2013, 'Invaders for sale: Trade and determinants of introduction of ornamental freshwater crayfish', *Biological Invasions* 15, 125–141. https://doi.org/10.1007/s10530-012-0273-2.<br>Faulkes, Z, 2015, 'The global trade in crayfish as pets', *Crustacean Research* 44, 75–92. https://doi.org/10.18353/crustacea.44.0_75.<br>Holdich, D.M., Reynolds, J.D. & Sibley, P.J., 2009, 'A review of the ever increasing threat to European crayfish from non-indigenous crayfish species', *Knowledge and Management of Aquatic Ecosystems* 11, 394–395. https://doi.org/10.1051/kmae/2009025. |

#### 3. Consequences

**CON1 Environmental impact**

**CON1a: Competition**

| Response: Major (MR) | Confidence: Low |
|---|---|

Rationale: *Pacifastacus leniusculus* contributes to the decline of several native species in introduced areas through competition for resources (Dana et al. 2010; Lodge et al. 2012; Weinlader & Furerfer 2002). Significant competition and reproductive interference often result in the displacement and extinction of native species where *P. leniusculus* out-competes native crayfish species (i.e., *Astacus astacus, Austropotamobius torrentium, Cambaroides japonicas* and *Pacifastacus nigrescens*) for shelter, making them more susceptible to predation (Huber & Schubart 2005; Pockl & Pekny 2002; Westman et al. 2002). There are no native crayfish species in South Africa and introduced crayfish have the potential to outcompete native potamonautid crabs (Mandivanzira et al. 2021).

References:

Dana, E.D., López-Santiago J., García-de-Lomas, J., García-Ocaña, D.M., Gámez, V. & Ortega, F., 2010, 'Long-term management of the invasive *Pacifastacus leniusculus* (Dana, 1852) in a small mountain stream', *Aquatic Invasions* 5, 317–322. https://doi.org/10.3391/ai.2010.5.3.10.

Huber, M.G. & Schubart, C.D., 2005, 'Distribution and reproductive biology of *Austropotamobius torrentium* in Bavaria and documentation of a contact zone with the alien crayfish *Pacifastacus leniusculus', Bulletin Français de la Pêche et de la Pisciculture* 376, 759–776. https://doi.org/10.1051/kmae:2005031.

Lodge, D.M., Deines, A., Gherardi, F., Yeo, D.C.J., Arcella, T. & Baldridge, A.K., et al., 2012, 'Global introductions of crayfishes: Evaluating the impact of species invasions on ecosystem services', *Annual Review of Ecology Evolution Systematics* 43, 449–472. , https://doi.org/10.1146/annurev-ecolsys-111511-103919

Madzivanzira, TC, South, J, Weyl, OLF. 2021, 'Invasive crayfish outperform Potamonautid crabs at higher temperatures', *Freshwater Biology* 66, 978–991. https://doi.org/10.1111/fwb.13691

Pöckl, M. & Pekny, R., 2002. 'Interaction between native and alien species of crayfish in Austria: case studies', *Bulletin Français de la Pêche et de la Pisciculture* 367, 763–776. https://doi.org/10.1051/kmae:2002064.

Weinländer, M. & Füreder, L., 2009, 'The continuing spread of *Pacifastacus leniusculus* in Carinthia (Austria)', *Knowledge and Management of Aquatic Ecosystems* 17, 394–395. . https://doi.org/10.1051/kmae/20010011.

Westman, K., Savolainen, R. & Julkunen, M., 2002, 'Replacement of the native crayfish *Astacus astacus* by the introduced species *Pacifastacus leniusculus* in a small, enclosed Finnish lake: A 30-year study', *Ecography* 25, 53–73, https://doi.org/10.1034/j.1600-0587.2002.250107.x.

**CON1b: Predation**

| Response: Moderate (MO) | Confidence: Medium |
|---|---|

Rationale: *Pacifastacus leniusculus* is a functional omnivore feeding on plant material, zoobenthos, detritus, fish, and sometimes other crayfish (Dana et al. 2010; Moorhouse 2018). *Pacifastacus leniusculus* tends to consume more animal than plant material and impacts invertebrate densities through direct predation (Crawford et al. 2006; Moorhouse 2018). In invaded areas, there is an overall decrease in richness and abundance of native species such as snails, leeches, caddisflies, newts, and fish (Crawford et al. 2006; Girdner et al. 2018, Galib et al. 2021).

References:

Crawford, L., Yeomans, W.E. & Adams, C.E., 2006, 'The impact of introduced signal crayfish *Pacifastacus leniusculus* on stream invertebrate communities', *Aquatic Conservation: Marine and Freshwater Ecosystems* 16, 611–621. https://doi.org/10.1002/aqc.761.

Dana, E.D., López-Santiago J., García-de-Lomas, J., García-Ocaña, D.M., Gámez, V. & Ortega, F., 2010, 'Long-term management of the invasive *Pacifastacus leniusculus* (Dana, 1852) in a small mountain stream', *Aquatic Invasions* 5, 317–322. , https://doi.org/10.3391/ai.2010.5.3.10.

Galib, S.M., Findlay, J.S., Lucas, M.C. 2021, 'Strong impacts of signal crayfish invasion on upland stream fish and invertebrate communities', *Freshwater Biology* 66, 223–240. https://doi.org/10.1111/fwb.13631

Girdner, S.F., Ray, A.M., Buktenica, M.W., Hering, D.K., Mack, J.A. & Umek, J.W., 2018, 'Replacement of a unique population of newts (*Taricha granulosa mazamae*) by introduced signal crayfish (*Pacifastacus leniusculus*) in Crater Lake, Oregon', *Biological Invasions* 20, 721–740. https://doi.org/10.1007/s10530-017-1570-6.

Moorhouse, T.P., Poole, A.E., Evans, L.C., Bradley, D.C. & Macdonald, D.W., 2014, 'Intensive removal of signal crayfish (*Pacifastacus leniusculus*) from rivers increases numbers and taxon richness of macroinvertebrate species', *Ecology and Evolution* 4, 494–504 https://doi.org/10.1002/ece3.903.

**CON1c: Hybridisation**

| Response: Data Deficient (DD) | Confidence: Low |
|---|---|

Rationale: No native freshwater crayfish in South Africa (Crandall & de Grave 2017).

References:

Crandall, K.A. & De Grave, S., 2017, 'An updated classification of the freshwater crayfishes (Decapoda: Astacidea) of the world, with a complete species list, *Journal of Crustacean Biology*, 37 (5), 615–653. https://doi.org/10.1093/jcbiol/rux070

**CON1d: Transmission of disease**

| Response: Major (MR) | Confidence: Medium |
|---|---|

Rationale: *Pacifastacus leniusculus* is a vector for crayfish plague, a disease caused by the parasitic oomycete *Aphanomyces astaci* (Lodge et al. 2012; Longshaw 2011). In Europe, the transmission of crayfish plague by *P. leniusculus* to native crayfish species has been linked to a decline in populations and in some case population collapses of several native species of crayfish such as *Astacus astacus*, *Austropotamobius pallipes*, and *Austropotamobius torrentium* (Dunn et al. 2009; Holdich & Reeve 1991; Holdich et al. 2009). There are no native crayfish species in South Africa but crayfish plague pathogen has been shown to infect other freshwater decapods such as native potamonautid crabs crabs (Svoboda et al. 2014)

References:

Dunn, J.C, McClymont, H.E., Christmas, M. & Dunn, A.M., 2009, Competition and parasitism in the native white clawed crayfish *Austropotamobius pallipes* and the invasive signal crayfish *Pacifastacus leniusculus* in the UK, *Biological Invasions* 11, 315–324. https://doi.org/10.1007/s10530-008-9249-7.

Holdich, D.M., Reynolds, J.D. & Sibley, P.J., 2009, 'A review of the ever increasing threat to European crayfish from non-indigenous crayfish species', *Knowledge and Management of Aquatic Ecosystems* 11, 394–395. https://doi.org/10.1051/kmae/2009025.

Holdich, D.M. & Reeve, I.D., 1991, 'Distribution of freshwater crayfish in the British Isles, with particular reference to crayfish plague, alien introductions and water quality', *Aquatic Conservation: Marine and Freshwater Ecosystems* 1, 139–158. https://doi.org/10.1002/aqc.3270010204.

Lodge, D.M., Deines, A., Gherardi, F., Yeo, D.C.J., Arcella. T, Baldridge, A.K. et al., 2012, 'Global introductions of crayfishes: Evaluating the impact of species invasions on ecosystem services', *Annual*

*Review of Ecology Evolution Systematics* 43, 449–472. https://doi.org/10.1146/annurev-ecolsys-111511-103919.
Longshaw, M. 2011, 'Diseases of crayfish: A review', *Journal of Invertebrate Pathology* 106, 54–70. https://doi.org/10.1016/j.jip.2010.09.013.
Svoboda, J., Strand, D.A., Vrålstad, T., Grandjean, F., Edsman, L., Kozák, P., Kouba, A., Fristad, R.F., Bahadir Koca, S. and Petrusek, A. 2014, 'The crayfish plague pathogen can infect freshwater-inhabiting crabs', *Freshwater Biology*, 59, 918-929. https://doi–org.uplib.idm.oclc.org/10.1111/fwb.12315

**CON1e: Parasitism**

Response: Data Deficient (DD) | Confidence: Low

Rationale:

References:

**CON1f: Poisoning/toxicity**

Response: Data Deficient (DD) | Confidence: Low

Rationale:

References:

**CON1g: Bio-fouling or other direct physical disturbance**

Response: Data Deficient (DD) | Confidence: Low

Rationale:

References:

**CON1h: Grazing/herbivory/browsing**

Response: Data Deficient (DD) | Confidence: Low

Rationale: Although there is no information available to assess the level of impact, *P. leniusculus* is a functional omnivore feeding on plant material (e.g., Guan & Wiles 1998, Carvalho et al. 2022).

References:

Guan, R. & Wiles, P.R., 1998, 'Feeding ecology of the signal crayfish *Pacifastacus leniusculus* in a British lowland river', *Aquaculture* 168, 177–193. https://doi.org/10.1016/S0044-8486(98)00377-9.
Carvalho, F., Sousa, R., Cássio, F., Pascoal C. 2022, 'Temperature and interspecific competition alter the impacts of two invasive crayfish species on a key ecosystem process. *Biological Invasions* 24, 3757–3768 https://doi.org/10.1007/s10530-022-02875-7

**CON1i: Chemical, physical or structural impact on ecosystem**

Response: Moderate (MO) | Confidence: Medium

Rationale: Burrowing activities of *P. leniusculus* can cause structural damage to river banks and increase bank erosion (Holdich et al. 2009). Although it is considered to be a non-burrowing species, in its invaded range, it constructs burrows under rocks and river banks (Dana et al. 2010). In Europe, the burrows can reach high densities, and can have a severe impact on river bank geomorphology, causing them to collapse (Holdich et al. 2009). On River Lark in the UK, burrowing by *P. leniusculus* has been reported to cause erosion at the rate of one meter per year (Guan 2010).

| |
|---|
| *Pacifastacus leniusculus* were found to have higher diversity of epibiont algae (species that live on the surface of other organisms) than sympatric native species (Falasco et al. 2018). This indicates that they can provide alternative substrate for agal species to grow on and this can this can lead to changes in community structure and function. |
| References:<br>Dana, E.D., López-Santiago J., García-de-Lomas, J., García-Ocaña, D.M., Gámez, V. & Ortega, F., 2010, 'Long-term management of the invasive *Pacifastacus leniusculus* (Dana, 1852) in a small mountain stream', *Aquatic Invasions* 5, 317–322. https://doi.org/10.3391/ai.2010.5.3.10.<br>Guan, R.Z., 2010, 'Burrowing behaviour of signal crayfish, '*Pacifastacus leniusculus* (Dana) in the River Great Ouse, England', *Freshwater Forum* 4,155–168. http://hdl.handle.net/1834/22157<br>Holdich, D.M., Reynolds, J.D. & Sibley, P.J., 2009, 'A review of the ever increasing threat to European crayfish from non-indigenous crayfish species', *Knowledge and Management of Aquatic Ecosystems* 11, 394–395. https://doi.org/10.1051/kmae/2009025.<br>Falasco, E., Bo, T., Ghia, D., Gruppuso, L., Bona, F., Fenoglio, S. 2018 "Diatoms prefer strangers: non-indigenous crayfish hosts completely different epizoic algal diatom communities from sympatric native species." . Biological Invasions 20: 2767-2776. https://doi.org/10.1007/s10530-018-1728-x |

| **CON1k: Indirect impacts through interactions with other species** | |
|---|---|
| Response: Data Deficient (DD) | Confidence: Low |
| Rationale: | |
| References: | |

| **CON1 Maximum environmental impact** | |
|---|---|
| Response: Major (MR) | Confidence: Low |
| Rationale: Direct predation and competition for food and shelter by *P. leniusculus* have led to local extinctions and a decrease in the abundance of invertebrates, native newts, crayfish and fish species in Japan, the USA, and several countries in Europe (Crawford et al. 2006; Lodge et al. 2012, Holdich et al. 2009, Lodge et al. 2012, Galib et al. 2021). *Pacifastacus leniusculus* is a vector for crayfish plague that was responsible for the decline of the European crayfish populations (Holdich et al. 2009; Lodge et al. 2012). | |
| References:<br><br>Crawford, L., Yeomans, W.E. & Adams, C.E., 2006, 'The impact of introduced signal crayfish *Pacifastacus leniusculus* on stream invertebrate communities', *Aquatic Conservation: Marine and Freshwater Ecosystems* 16, 611–621. https://doi.org/10.1002/aqc.761.<br>Galib, S.M., Findlay, J.S., Lucas, M.C. 2021, 'Strong impacts of signal crayfish invasion on upland stream fish and invertebrate communities', *Freshwater Biology* 66, 223–240. https://doi.org/10.1111/fwb.13631<br>Holdich, D.M., Reynolds, J.D. & Sibley, P.J., 2009, 'A review of the ever increasing threat to European crayfish from non-indigenous crayfish species', *Knowledge and Management of Aquatic Ecosystems* 11, 394–395. https://doi.org/10.1051/kmae/2009025.<br>Lodge, D.M., Deines, A., Gherardi, F., Yeo, D.C.J., Arcella. T, Baldridge, A.K. et al., 2012, 'Global introductions of crayfishes: Evaluating the impact of species invasions on ecosystem services', *Annual Review of Ecology Evolution Systematics* 43, 449–472. https://doi.org/10.1146/annurev-ecolsys-111511-103919. | |

| **CON2 Socio-economic impact** |
|---|
| **CON2a: Safety** |

| Response: Data Deficient (DD) | Confidence: Low |
|---|---|
| Rationale: | |
| References: | |
| **CON2b: Material and immaterial assets** | |
| Response: Minor (MN) | Confidence: Low |
| Rationale: *Pacifastacus leniusculus* introduced in Europe to alleviate the pressure on native crayfish such as *Astacus astacus* whose population had declined due to crayfish plague (Holdich et al. 2009). The introduction of *P. leniusculus* has however contributed to further decline in populations of native crayfish because *P. leniusculus* are carriers of the crayfish plague and have competitively displaced native crayfish in areas of introduction (e.g., Dana et al. 2010). *Astacus astacus* is considered to be more valuable than *P. leniusculus* and but the economic losses as a result of the introduction of *P. leniusculus* is unknown (Johnsen & Taugbøl 2010, Kouba et al. 2022). In addition, the burrowing activities of *P. leniusculus* can cause structural damage to river banks and increase bank erosion (Holdich et al. 2009, Kouba et al. 2022). | |
| References:<br>Dana, E.D., López-Santiago J., García-de-Lomas, J., García-Ocaña, D.M., Gámez, V. & Ortega, F., 2010, 'Long-term management of the invasive *Pacifastacus leniusculus* (Dana, 1852) in a small mountain stream', *Aquatic Invasions* 5, 317–322. https://doi.org/10.3391/ai.2010.5.3.10.<br>Holdich, D.M., Reynolds, J.D. & Sibley, P.J., 2009, 'A review of the ever increasing threat to European crayfish from non-indigenous crayfish species', *Knowledge and Management of Aquatic Ecosystems* 11, 394–395. https://doi.org/10.1051/kmae/2009025.<br>Johnsen SI, Taugbøl T. (2010): NOBANIS – Invasive Alien Species Fact Sheet – *Pacifastacus leniusculus.* Online Database of the European Network on Invasive Alien Species – NOBANIS www.nobanis.org. [Date of access 13/11/2019].<br>Kouba, A., Oficialdegui, F.J., Cuthbert, R.N., Kourantidou, M., South, J., Tricarico, E., Gozlan, R.E., Courchamp, F. and Haubrock, P.J., 2022. 'Identifying economic costs and knowledge gaps of invasive aquatic crustaceans', *Science of the Total Environment*, *813*, 152325 https://doi.org/10.1016/j.scitotenv.2021.152325 | |
| **CON2c: Health** | |
| Response: Data Deficient (DD) | Confidence: Low |
| Rationale: | |
| References: | |
| **CON2d: Social, spiritual and cultural relations** | |
| Response: Minor (MN) | Confidence: Low |
| Rationale: *Pacifastacus leniusculus* introduced in Europe to alleviate the pressure on native crayfish such as *Astacus astacus* whose population had declined due to crayfish plague (Holdich et al. 2009). This introduction has however contributed to further decline in native crayfish populations because *P. leniusculus* are carriers of the crayfish plague and have competitively displaced native crayfish in areas of introduction (e.g., Dana et al. 2010). The native crayfish species have a high cultural value and in places such as Scandinavia, the loss of *A. astacus* is considered to be serious cultural blow (Gherardi 2011). | |

References:

Dana, E.D., López-Santiago J., García-de-Lomas, J., García-Ocaña, D.M., Gámez, V. & Ortega, F., 2010, ‘Long-term management of the invasive *Pacifastacus leniusculus* (Dana, 1852) in a small mountain stream’, *Aquatic Invasions* 5, 317–322. https://doi.org/10.3391/ai.2010.5.3.10.

Holdich, D.M., Reynolds, J.D. & Sibley, P.J., 2009, ‘A review of the ever increasing threat to European crayfish from non-indigenous crayfish species’, Knowledge and Management of Aquatic Ecosystems 11, 394–395. https://doi.org/10.1051/kmae/2009025.

Gherardi F. 2011b. Towards a sustainable human use of crayfish (Crustacea, Decapoda, Astacidea). *Knowledge and Management of Aquatic Ecosystems*. 401, 02 https://doi.org/10.1051/kmae/2011038

**CON2 Maximum socio-economic impact**

| Response: Minor (MN) | Confidence: Low |
|---|---|

Rationale: *Pacifastacus leniusculus* introduced in Europe to alleviate the pressure on native crayfish such as *Astacus astacus* whose population had declined due to crayfish plague (Holdich et al. 2009). This introduction has however contributed to further decline in native crayfish populations because *P. leniusculus* are carriers of the crayfish plague and have competitively displaced native crayfish in areas of introduction (e.g., Dana et al. 2010). The native crayfish species have a high cultural and economic values and the introduction of *P. leniusculus* has lead to a decline in the benefits derived from the native crayfish (Gherardi 2011, Lodge et al. 2012; Kouba et al. 2022).

References:

Dana, E.D., López-Santiago J., García-de-Lomas, J., García-Ocaña, D.M., Gámez, V. & Ortega, F., 2010, ‘Long-term management of the invasive *Pacifastacus leniusculus* (Dana, 1852) in a small mountain stream’, *Aquatic Invasions* 5, 317–322. https://doi.org/10.3391/ai.2010.5.3.10.

Gherardi F. 2011b. Towards a sustainable human use of crayfish (Crustacea, Decapoda, Astacidea). *Knowledge and Management of Aquatic Ecosystems*. 401, 02 https://doi.org/10.1051/kmae/2011038

Holdich, D.M., Reynolds, J.D. & Sibley, P.J., 2009, ‘A review of the ever increasing threat to European crayfish from non-indigenous crayfish species’, Knowledge and Management of Aquatic Ecosystems 11, 394–395. https://doi.org/10.1051/kmae/2009025.

Johnsen SI, Taugbøl T. (2010): NOBANIS – Invasive Alien Species Fact Sheet – *Pacifastacus leniusculus.* Online Database of the European Network on Invasive Alien Species – NOBANIS www.nobanis.org. [Date of access 13/11/2019].

Kouba, A., Oficialdegui, F.J., Cuthbert, R.N., Kourantidou, M., South, J., Tricarico, E., Gozlan, R.E., Courchamp, F. and Haubrock, P.J., 2022’, ‘Identifying economic costs and knowledge gaps of invasive aquatic crustaceans’, *Science of the Total Environment*, 813, 152325 https://doi.org/10.1016/j.scitotenv.2021.152325

Lodge, D.M., Deines, A., Gherardi, F., Yeo, D.C.J., Arcella. T, Baldridge, A.K. et al., 2012, ‘Global introductions of crayfishes: Evaluating the impact of species invasions on ecosystem services’, *Annual Review of Ecology Evolution Systematics* 43, 449–472. https://doi.org/10.1146/annurev-ecolsys-111511-103919

**CON3 Closely related species’ environmental impact**

| Response: Not scored as CON1 is scored | Confidence: High |
|---|---|

Rationale: There is information to score CON1, so CON3 does not need to be scored as per the framework.

References:

**CON4 Closely related species’ socio-economic impact**

| Response: Not scored as CON2 is scored | Confidence: High |
|---|---|
| Rationale: There is information to score CON2, so CON4 does not need to be scored as per the framework. | |
| References: | |

| **CON5 Potential impact** | |
|---|---|
| **Response:** Major (MR) | **Confidence:** Low |
| Rationale: *Pacifastacus leniusculus* is a significant competitor and has displaced several species of native freshwater crayfish and other fauna in introduced areas (Dunn et al. 2009; Huber & Schubart 2005; Galib et al. 2021). While South Africa may not have native freshwater crayfish, closely related decapods such as crabs, may have a resource overlap with *P. leniusculus* which may result in competition (Mandivanzira et al. 2021). *Pacifastacus leniusculus* is a vector for crayfish plague that caused population crashes of native crayfish species in Europe (Dana et al. 2010; Holdich & Reeves 1991). Crayfish plague can affect other native freshwater decapods such as native potamonautid crabs (Svoboda et al. 2014). Furthermore, *P. leniusculus* is a facultative omnivore, and may cause negative impacts on macroinvertebrates and macrophyte communities (Dunn et al. 2009; Galib et al. 2021; Westman et al. 2002). In areas outside of its native range, *P. leniusculus* constructs burrows which can weaken riverbanks and dam walls, changing their bank geomorphology, leading to erosion (Guan 1994). | |
| References:<br><br>Dana, E.D., López-Santiago J., García-de-Lomas, J., García-Ocaña, D.M., Gámez, V. & Ortega, F., 2010, 'Long-term management of the invasive *Pacifastacus leniusculus* (Dana, 1852) in a small mountain stream', *Aquatic Invasions* 5 317–322. https://doi.org/10.3391/ai.2010.5.3.10.<br>Dunn, J.C, McClymont, H.E., Christmas, M. & Dunn, A.M., 2009, Competition and parasitism in the native white clawed crayfish *Austropotamobius pallipes* and the invasive signal crayfish *Pacifastacus leniusculus* in the UK, *Biological Invasions* 11, 315–324. https://doi.org/10.1007/s10530-008-9249-7.<br>Galib, S.M., Findlay, J.S., Lucas, M.C. 2021, 'Strong impacts of signal crayfish invasion on upland stream fish and invertebrate communities', *Freshwater Biology* 66, 223–240. https://doi.org/10.1111/fwb.13631<br>Guan, R.Z., 1994, 'Burrowing behaviour of signal crayfish, *Pacifastacus leniusculus* (Dana) in the River Great Ouse, England', *Freshwater Forum* 4,155–168.<br>Holdich, D.M. & Reeve, I.D., 1991, 'Distribution of freshwater crayfish in the British Isles, with particular reference to crayfish plague, alien introductions and water quality', *Aquatic Conservation: Marine and Freshwater Ecosystems* 1, 139–158. https://doi.org/10.1002/aqc.3270010204.<br>Huber, M.G. & Schubart, C.D., 2005, 'Distribution and reproductive biology of *Austropotamobius torrentium* in Bavaria and documentation of a contact zone with the alien crayfish *Pacifastacus leniusculus*', *Bulletin Français de la Pêche et de la Pisciculture* 376,759–776. https://doi.org/10.1051/kmae:2005031<br>Madzivanzira, TC, South, J, Weyl, OLF. 2021, 'Invasive crayfish outperform Potamonautid crabs at higher temperatures', *Freshwater Biology* 66, 978–991. https://doi.org/10.1111/fwb.13691<br>Svoboda, J., Strand, D.A., Vrålstad, T., Grandjean, F., Edsman, L., Kozák, P., Kouba, A., Fristad, R.F., Bahadir Koca, S. and Petrusek, A. 2014, 'The crayfish plague pathogen can infect freshwater-inhabiting crabs', *Freshwater Biology*, 59, 918-929. https://doi.org/10.1111/fwb.12315<br>Westman, K., Savolainen R. & Julkunen, M., 2002, 'Replacement of the native crayfish *Astacus astacus* by the introduced species *Pacifastacus leniusculus* in a small, enclosed Finnish lake: A 30–year study', *Ecography* 25, 53–73. https://doi.org/10.1034/j.1600-0587.2002.250107.x | |

**4. Management**

| MAN1 What is the feasibility to stop future immigration? | |
|---|---|
| Response: High | Confidence: Low |

Rationale: *Pacifastacus leniusculus* is a common species in the global pet trade industry (Faulkes 2015). It is suspected that *P. leniusculus* is present in the pet trade in South Africa but this needs to be confirmed (Madzivanzira et al. 2021, Nunes et al. 2017). The species is not in neighbouring so unaided migration is unlikely. It should be possible to regulate international introductions by listing the species on the prohibited list and having an effective border surveillance and monitoring strategy.

References:

Faulkes, Z., 2015, 'The global trade in crayfish as pets', *Crustacean Research* 44**,** 75–92. https://doi.org/10.18353/crustacea.44.0_75.

Madzivanzira, T.C., South, J. & Weyl, O.L.F., 2021, 'Invasive crayfish outperform Potamonautid crabs at higher temperatures', *Freshwater Biology* 66, 978–991. https://doi.org/10.1111/fwb.13691

Nunes, A.L., Zengeya, T.A., Measey, G.J. & Weyl, O.L.F., 2017, 'Freshwater crayfish invasions in South Africa: past, present and potential future', *African Journal of Aquatic Science* 42, 309–323, https://doi.org/10.2989/16085914.2017.1405788.

**MAN2 Benefits of the Taxon**

| MAN2a Socio-economic benefits of the *Taxon* | |
|---|---|
| Response: Low | Confidence: Low |

Rationale: *Pacifastacus leniusculus* is introduced mainly through aquaculture (Holdich 1993; Holdich et al. 2009), and the pet trade industry (Chucholl 2013; Faulkes 2015). The presence of *P. leniusculus* in South Africa still needs to be confirmed (Nines et al. 2017), and if it is here, it is likely to be in low numbers, therefore the socio-economic benefits are also likely to be low (Madzivanzira et al. 2021).

References:

Chucholl, C., 2013, 'Invaders for sale: Trade and determinants of introduction of ornamental freshwater crayfish', *Biological Invasions* 15, 125–141. https://doi.org/10.1007/s10530-012-0273-2

Faulkes, Z, 2015, 'The global trade in crayfish as pets', *Crustacean Research* 44, 75–92. https://doi.org/10.18353/crustacea.44.0_75.

Holdich, D.M., 1993, 'A review of astaciculture: Freshwater crayfish farming', *Aquatic Living Resources* 6, 307–317. https://doi.org/10.1051/alr:1993032

Holdich, D.M., Reynolds, J.D., & Sibley, P.J., 2009, 'A review of the ever increasing threat to European crayfish from non-indigenous crayfish species', *Knowledge and Management of Aquatic Ecosystems* 11, 394–395. https://doi.org/10.1051/kmae/2009025.

Madzivanzira, T.C,. South, J. & Weyl, O.L.F., 2021, 'Invasive crayfish outperform Potamonautid crabs at higher temperatures', *Freshwater Biology* 66, 978–991. https://doi.org/10.1111/fwb.13691

Nunes, A.L., Zengeya, T.A., Measey, G.J. & Weyl, O.L.F., 2017, 'Freshwater crayfish invasions in South Africa: past, present and potential future', *African Journal of Aquatic Science* 42, 309–323, https://doi.org/10.2989/16085914.2017.1405788.

| MAN2b Environmental benefits of the *Taxon* | |
|---|---|
| Response: Low | Confidence: High |

Rationale: There are no recorded environmental benefits of *P. leniusculus*.

References:

**MAN3 Ease of management**

**MAN3a How accessible are populations?**

| Response: 2 (don’t know) | Confidence: Low |
| --- | --- |

Rationale: There are no locality records for *P. leniusculus* in South Africa (Madzivanzira et al. 2021, Nunes et al. 2017).

References:

Madzivanzira, T.C, South, J. & Weyl, O.L.F., 2021, ‘Invasive crayfish outperform Potamonautid crabs at higher temperatures’, *Freshwater Biology* 66, 978–991. https://doi.org/10.1111/fwb.13691

Nunes, A.L., Zengeya, T.A., Measey, G.J. & Weyl, O.L.F., 2017, ‘Freshwater crayfish invasions in South Africa: past, present and potential future’, *African Journal of Aquatic Science* 42, 309–323, https://doi.org/10.2989/16085914.2017.1405788.

**MAN3b Is detectability critically time-dependent?**

| Response: Difficult (2) | Confidence: High |
| --- | --- |

Rationale: *Pacifastacus leniusculus* is a fairly small, largely nocturnal, burrowing freshwater species that is difficult to detect (Dana et al. 2010; Guan 2010).

References:

Dana, E.D., López-Santiago J., García-de-Lomas, J., García-Ocaña, D.M., Gámez, V. & Ortega, F., 2010, ‘Long-term management of the invasive *Pacifastacus leniusculus* (Dana, 1852) in a small mountain stream’, *Aquatic Invasions* 5, 317–322. https://doi.org/10.3391/ai.2010.5.3.10.

Guan, R.Z., 2010, ‘Burrowing behaviour of signal crayfish, ‘*Pacifastacus leniusculus* (Dana) in the River Great Ouse, England’, *Freshwater Forum* 4,155–168.

**MAN3c Time to reproduction**

| Response:1 (1–3 years) | Confidence: High |
| --- | --- |

Rationale: *Pacifastacus leniusculus* reach sexual maturity in two-three years (or once individuals reach six to nine cm in length (head to tail)) however maturity can also occur as early as a year (Guan and Wiles 1999).

References:

Guan, R. Z. & Wiles, P. R., 1999, ‘Growth and reproduction of the introduced crayfish *Pacifastacus leniusculus* in a British lowland river’. *Fisheries Research*, 42, 245-259. https://doi.org/10.1016/S0165-7836(99)00044-2

**MAN3d Propagule persistence**

| Response:1 (1–5 years) | Confidence: High |
| --- | --- |

Rationale: *Pacifastacus leniusculus* are highly fecund, on average a female produces 200-400 eggs (sometimes 500 eggs), and hatchlings hang under the female's tail for many weeks until they can feed on their own (Guan and Wiles 1999).

References:

Guan, R. Z., & Wiles, P. R. 1999, ‘Growth and reproduction of the introduced crayfish *Pacifastacus leniusculus* in a British lowland river’. *Fisheries Research*, 42, 245-259.

| **MAN3 Ease of management** | |
|---|---|
| Response: Difficult | Confidence: High |

Rationale:

*Pacifastacus leniusculus* is a fairly small, largely nocturnal, burrowing freshwater species that is difficult to detect. There are several methods that have been used to control alien populations of *P. leniusculus* these include mechanical removal using traps and electrofishing, biocides and sterile male release (Peay et al. 2019, Ballentyne et al. 2019). However, there are several issues that make management of crayfish invasions challenging (Gherardi et al. 2011). For example, the use of biocides is often effective for localized populations in dams and ponds (Ballantyne et al. 2019; Peay et al. 2019) but it is not effective for widespread species because of the large doses of treatment required and higher potential of adverse effects on non-targeted species (Gherardi et al. 2011). Mechanical controls are also effective in small discrete water bodies but are expensive, require commitment to follow on to complete eradication and can lead unwanted community compensation mechanisms such as enhanced recruitment (Peay et al. 2014, Larson et al. 2022). *Pacifastacus leniusculus* also constructs burrows that could impede some control methods (Peay & Dunn 2006).

*Pacifastacus leniusculus* is a popular species in the global pet trade industry and they are unconfirmed records that it might be present South Africa. The pet trade industry is generally difficult to regulate because trade restrictions imposed on species of conservation concern that are albeit popular and in high demand, often lead to illegal trading that is compounded by a lack of reliable record-keeping of the type and number of organisms presently in the pet trade (Smith et al., 2008; Chang et al., 2009; Gippet & Bertelsmeier, 2021).

References:

Ballantyne, L., Baum, D., Bean, C.W., Long, J. & Whitaker, S., 2019, 'Successful eradication of signal crayfish (*Pacifastacus leniusculus*) using a non-specific biocide in a small isolated water body in Scotland'. In: Veitch, C.R., Clout, M.N., Martin, A.R., J.C. Russell, J.C. & West, C.J. (eds.). Island invasives: scaling up to meet the challenge, pp. 443–446. Occasional Paper SSC no. 62. Gland, Switzerland: IUCN.

Chang, A.L., Grossman, J.D., Spezio, T.S., Weiskel, H.W., Blum, J.C., Burt, J.W., Muir, A.A., Piovia-Scott, J., Veblen, K.E. & Grosholz, E.D. 2009. 'Tackling aquatic invasions: risks and opportunities for the aquarium fish industry', *Biological Invasions* 11, 773–785. https://doi.org/10.1007/s10530-008-9292-4

Dana, E.D., López-Santiago J., García-de-Lomas, J., García-Ocaña, D.M., Gámez, V. & Ortega, F., 2010, 'Long-term management of the invasive *Pacifastacus leniusculus* (Dana, 1852) in a small mountain stream', *Aquatic Invasions* 5, 317 322. https://doi.org/10.3391/ai.2010.5.3.10.

Gherardi, F, Aquiloni, L., Diéguez-Uribeondo, J. & Tricarico E., 2011, 'Managing invasive crayfish: Is there a hope?', *Aquatic Sciences*, 73, 185–200. https:// doi.org/ 10.1007/s00027-011-0181-z

Gippet, J.M.& Bertelsmeier, C. 2021, 'Invasiveness is linked to greater commercial success in the global pet trade', *Proceedings of the National Academy of Sciences* 118, e2016337118. https://doi.org/10.1073/pnas.2016337118

Green, N., Andreou, D., Bentley, M., Stebbing, P., Hart, A. & Britton, J.R. 2022, 'Mechanical male sterilisation in invasive signal crayfish *Pacifastacus leniusculus*: persistence and functionality in captive and wild conditions. *Knowledge & Management of Aquatic Ecosystems* 423, 20 https://doi.org/10.1051/kmae/2022014

Larson, C.E., Bo, T., Candiotto, A., Fenoglio, S. & Doretto, A. 2022. 'Predicting invasive signal crayfish (*Pacifastacus leniusculus*) spread using a traditional survey and river network simulation', *River Research and Applications* 38, 1424-1435, https://doi.org/ 10.1002/rra.3986

Peay, S & Dunn, A.M. 2014, 'The behavioural response of the invasive signal crayfish *Pacifastacus leniusculus* to experimental dewatering of burrows and its implications for eradication treatment and management of ponds with crayfish', *Ethology Ecology & Evolution*, 26, 277- 298, https://doi.org/10.1080/03949370.2014.903379

Peay, S., Johnsen, S.I., Bean, C.W., Dunn, A.M., Sandodden, R. & Edsman, L. 2019, 'Biocide treatment of invasive signal crayfish: successes, failures and lessons learned. *Diversity, 11,* 29. https://doi.org/10.3390/d11030029

Peay, S., Dunn, A. M., Kunin, W. E., McKimm, R & Harrod, C. 2014, 'A method test of the use of electric shock treatment to control invasive signal crayfish in streams', *Aquatic Conservation: Marine. and Freshwater. Ecosystems.*, 25,: 874–880. https://doi.org/10.1002/aqc.2541.

Smith, K.F., Behrens, M.D., Max, L.M. & Daszak, P. 2008. 'U.S. drowning in unidentified fishes: scope, implications, and regulation of live fish import', *Conservation Letters* 1, 103–109. https://doi.org/10.1111/j.1755-263X.2008.00014.x

**MAN4 Has the feasibility of eradication been evaluated?**

| Response: No | Confidence: Low |
|---|---|

Rationale: Although feasibility of eradication has not been evaluated for SA, experience in other parts of the world suggest that once *P. leniusculus* has become established and widespread, eradication is infeasible (Gherardi et al. 2011). Eradication using biocides is effective for localised populations such as in small dams/ponds but biocides often have non-target impacts on native fauna (Ballantyne et al. 2019).

References:

Ballantyne, L., Baum, D., Bean, C.W, Long, J. & Whitaker S., 2019, 'Successful eradication of signal crayfish (*Pacifastacus leniusculus*) using a non-specific biocide in a small isolated water body in Scotland', *Island invasives: scaling up to meet the challenge* 62, 443–446.

Gherardi, F., Aquiloni, L., Diéguez-Uribeondo, J. & Tricarico, E., 2011, 'Managing invasive crayfish: Is there a hope?', *Aquatic Sciences* 73, 185–200. https://doi.org/10.1007/s00027-011-0181-z.

**MAN5 Control options and monitoring approaches available for the *Taxon***

Response: Biocides can be considered when *P. leniusculus* is confined to small dams. However, it has been noted that since *P. leniusculus* burrows, dewatering the areas sufficiently prior to biocides application is vital (Peay & Dunn 2014). This is to ensure that all individuals will be targeted to prevent alien crayfish to establish populations again. This method is recommended as a rapid response and not continuous eradication method because it is not target specific and once an area is treated it requires time to recover (Peay & Dunn 2014). Trapping has been used historically; however, some studies caution against using trapping as the only control method because results indicate that this method may be body size-selective, removing adults only and consequently, juveniles grow rapidly due to lack of competition, creating positive feedback loops (Manfrin et al. 2019; Chadwick et al. 2020)

References:

Chadwick, D.D.A., Pritchard, E.G., Bradley, P., Sayer, C.D., Chadwick, M.A., Eagle, L.J.B. & Axmacher, J.C., 2020. A novel triple down method highlights deficiencies in invasive alien crayfish survey and control techniques. *Journal of Applied Ecology* 00: 1–11. https://doi.org/10.1111/1365-2664.13758

Manfrin, C., Souty-Grosset, C., Anastacio, P.M., Reynolds, J. & Giulianini, P.G., 2019, 'Detection and Control of Invasive Freshwater Crayfish: From Traditional to Innovative Methods', *Diversity* 11, 1–16. https://doi.org/10.3390/d11010005

Peay, S & Dunn A.M., 2014, 'The behavioural response of the invasive signal crayfish *Pacifastacus leniusculus* to experimental dewatering of burrows and its implications for eradication treatment and management of ponds with crayfish', *Ethology Ecology & Evolution*, 26, 277–298, https://doi.org/10.1080/03949370.2014.903379

**MAN6 Any other management considerations to highlight?**

| | |
|---|---|
| Response | No |

**5. Calculations**

**Likelihood = Fairly probable**

| Parameter | Likelihood | Stages | Final assessment |
|---|---|---|---|
| LIK1 | 0.0027 | P(entry) = 0.5 | P (invasion) = 0.125 |
| LIK2 | 0.5 | | |
| LIK3 | 0.5 | P(establishment) = 0.5 | |
| LIK4 | 0.5 | | |
| LIK5 | 0.5 | P (spread) = 0.5 | |
| LIK6 | 0.5 | | |

**Consequence = Major (MR)**

| Parameter | Mechanism/sector | Response |
|---|---|---|
| CON1a | Competition | MR |
| CON1b | Predation | MO |
| CON1c | Hybridisation | DD |
| CON1d | Disease transmission | MR |
| CON1e | Parasitism | DD |
| CON1f | Poisoning/toxicity | DD |
| CON1g | Bio-fouling or other direct physical disturbance | DD |
| CON1h | Grazing/herbivory/browsing | DD |
| CON1i | Chemical, physical, structural impact | MN |
| CON1k | Indirect impacts through interactions with other species | DD |
| CON1 | Maximum environmental impact | MR |
| CON2a | Safety | DD |
| CON2b | Material and immaterial assets | MN |
| CON2c | Health | DD |
| CON2d | Social, spiritual and cultural relations | MN |

| | | |
|---|---|---|
| CON2 | Maximum socio-economic impact | MN |
| CON3 | Environmental impact of closely related taxa | Not scored as CON1 scored |
| CON4 | Socio-economic impact of closely related taxa | Not scored as CON2 scored |
| CON5 | Potential impact based on traits, experiments, or models | MR |

**Risk score = High**

| | | Consequences | | | | |
|---|---|---|---|---|---|---|
| | | MC | MN | MO | MR | MV |
| Likelihood | Extremely unlikely | low | low | low | medium | medium |
| | Very unlikely | low | low | low | medium | high |
| | Unlikely | low | low | medium | high | high |
| | Fairly probable | medium | medium | high | high | high |
| | Probable | medium | high | high | high | high |

**Ease of management = Difficult**

| Parameter | Question | Response |
|---|---|---|
| MAN3a | How accessible are populations? | 2 |
| MAN3b | Is detectability critically time-dependent? | 2 |
| MAN3c | Time to reproduction | 1 |
| MAN3d | Propagule persistence | 1 |
| **MAN3** | **SUM** | 6 |

**Appendix BAC7**: The global native range of *Pacifastacus leniusculus* (Dana, 1852) indicated in orange (The IUCN Red List of Threatened Species: *Pacifastacus leniusculus* – published in 2010. http://dx.doi.org/10.2305/IUCN.UK.2010-3.RLTS.T153648A4526314.en).

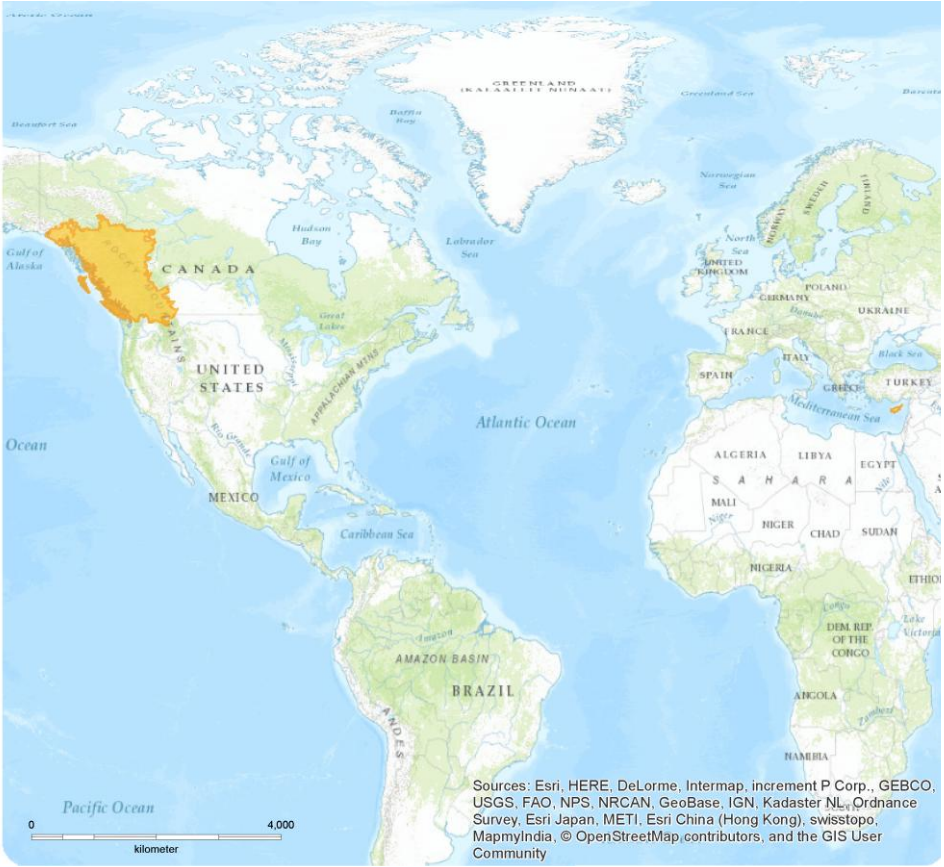

**Appendix BAC8(a)**: The global alien range of *Pacifastacus leniusculus* (Dana, 1852) indicated in pink (CABI 2021, from https://www.cabidigitallibrary.org/doi/10.1079/cabicompendium.70581) .

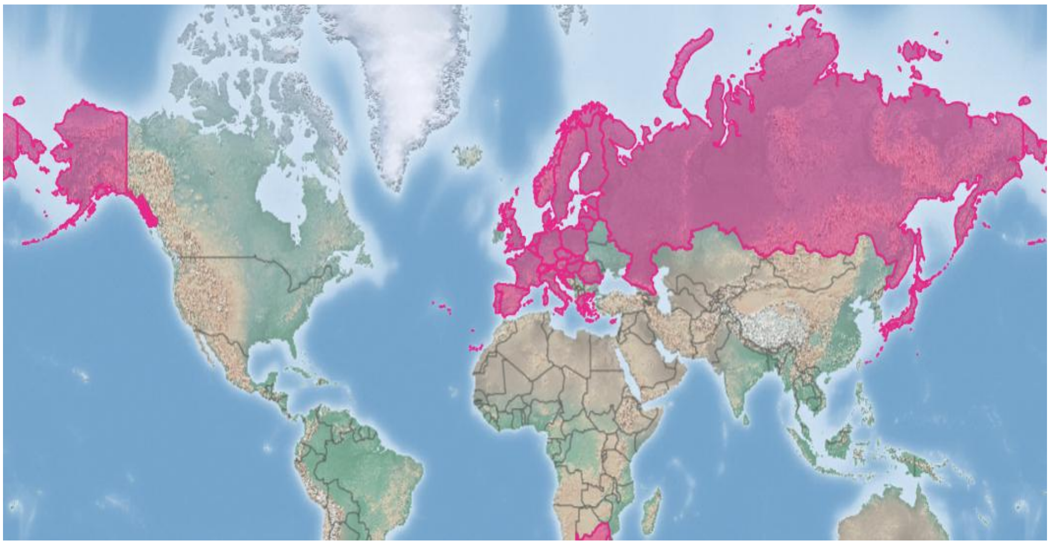

#### 1.11 Risk analysis report for Danube crayfish (*Pontastacus leptodactylus*)

| | |
|---|---|
| ***Taxon:*** *Pontastacus leptodactylus* | ***Area:*** South Africa |
| **Picture of *Taxon***<br>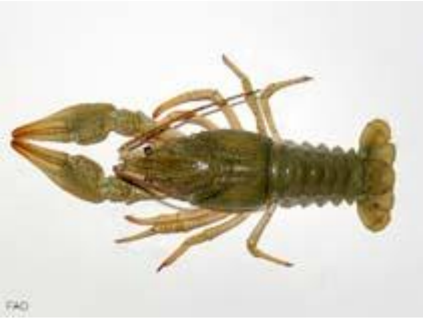<br>Photo: FAO | **Alien distribution map**<br>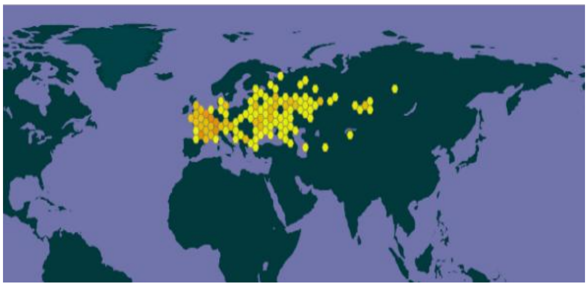<br>Reference: GBIF, 2023 |
| **Risk Assessment summary:**<br>Narrow-clawed crayfish (*Pontastacus leptodactylus*) are native to eastern Europe but have been introduced to several countries in western Europe. There are no know populations of *P. leptodactylus* in South Africa and in neighbouring countries, therefore via unaided pathways is unlikely. The illegal pet trade industry remains a concern because *P. leptodactylus* may be moved around by humans and there is a high risk of release of unwanted pets. There are no documented impacts from *P. leptodactylus*, and potential impacts were inferred from *Pacifastacus leniusculus*. In the introduced range, *P. leniusculus* has displaced native crayfish species through the transmission of crayfish plague and outcompeting them for food and shelter. *Pacifastacus leniusculus* has also caused negative impacts on macroinvertebrate communities through predation. Although there are no native crayfish species in South Africa, the crayfish plague has been shown to infect other freshwater decapods and introduced crayfish have the potential to outcompete native decapods such as potamonautid crabs. | **Risk score:**<br>High |
| **Management options summary:**<br>*Pontastacus leptodactylus* is a small, largely nocturnal, burrowing freshwater species that is difficult to detect. Several methods have been used to control alien populations of *P. leptodactylus* and these include mechanical removal using traps and electrofishing, biocides, and biological control. However, there are several issues that make the use of these methods challenging. The benefits of *P. leptodactylus* were scored as low. | **Ease of management:**<br><br>Medium |
| **Recommendations:** | **Listing under NEM:BA A&IS lists of 2020:** |

| | |
|---|---|
| *Pontastacus leptodactylus* poses a high risk to South Africa. If *P. leptodactylus* were to establish outside of captivity, such populations would likely be very difficult to extirpate. As such it is recommended that *P. leptodactylus* remains listed as category 1a, active efforts made to establish whether the species is preset in the country, and that any individuals found are controlled. This would likely require a dedicated assessment of the ornamental trade. A reassessment in five years is thus required and, if absent from the country, *P. leptodactylus s*hould be added to a future prohibited list. | 1a[1]<br><br>**Recommended listing category:**<br><br>1a[2] |

[1] The current listing is "*Astacus leptodactylus* (Eschscholtz, 1823)" the recommended listing is "*Pontastacus leptodactylus* (Eschscholtz, 1823)", i.e., it is proposed that the regulatory name be updated to the accepted scientific name of the taxon under assessment.

[2] The presence of *P. leptodactylus* in South Africa should be confirmed and the taxon should be reassessed in five years (i.e., by 2029).

#### 1. Background

| **BAC1 Name of assessor(s)** | | |
|---|---|---|
| Name of lead assessor | Lee-Anne Botha | |
| Additional assessor (1) | Tsungai Zengeya | |
| Additional assessor (2) | Christian Chimimba | |
| **BAC2 Contact details of assessor (s)** | | |
| Lead assessor | Organisational affiliation: Department of Zoology and Entomology, University of Pretoria and The South African National Biodiversity Institute (SANBI). | |
| | | |
| Additional assessor (1) | Organisational affiliation: Department of Zoology and Entomology, University of Pretoria and The South African National Biodiversity Institute (SANBI). | |
| | | |
| Additional assessor (2) | Organisational affiliation: Department of Zoology and Entomology, University of Pretoria | |
| | | |
| **BAC3 Name(s) and contact details of expert(s) consulted** | | |
| Expert (1) | Name: | |
| | email: | |
| | Phone: | |
| Expert (2) | Name: | |
| | email: | |
| | Phone: | |
| Comments: none | | |
| **BAC4 Scientific name of *Taxon* under assessment** | | |
| *Taxon* name: *Pontastacus leptodactylus* | | Authority: (Eschscholtz, 1823) |
| Comments: The scientific name of the taxon under assessment is accepted (GBIF 2023). | | |
| References:<br>*Pontastacus leptodactylus* (Eschscholtz, 1823) in GBIF Secretariat (2023). GBIF Backbone Taxonomy. Checklist dataset https://doi.org/10.15468/39omei accessed via GBIF.org on 2024-07-15. | | |
| **BAC5 Synonym(s) considered** | | |
| Synonyms: *Astacus leptodactylus* ssp. *kessleri* \| *Astacus leptodactylus* ssp. *eichwaldi* \| *Astacus leptodactylus* | | |
| Comments: The above listed synonyms of *Pontastacus leptodactylus* were used for the literature search for this risk assessment and obtained from GBIF (2024) and Gherardi & Souty-Grousset (2017). | | |
| References:<br>Gherardi, F.& Souty-Grosset, C., 2017, *Pontastacus leptodactylus. The IUCN Red List of Threatened Species 2017*: e.T153745A120103207. http://dx.doi.org/10.2305/IUCN.UK.2017-3.RLTS.T153745A120103207.en.<br>*Pontastacus leptodactylus* (Eschscholtz, 1823) in GBIF Secretariat (2023). GBIF Backbone Taxonomy. Checklist dataset https://doi.org/10.15468/39omei accessed via GBIF.org on 2024-07-15. | | |
| **BAC6 Common name(s) considered** | | |
| Common names: Danube crayfish\| Galican Crayfish \| Long-clawed Crayfish \| Narrow-clawed Crayfish \| Pond Crayfish \| Slender-clawed Crayfish \| Swamp Crayfish \| Turkish Crayfish | | |
| Comments: There are several English common names known for *Pontastacus leptodactylus* (Gherardi & Souty-Grousset 2017) and the above listed names were used in the literature search for this risk analysis | | |
| References:<br>Gherardi, F. & Souty-Grosset, C., 2017, *Pontastacus leptodactylus. The IUCN Red List of Threatened Species 2017*: e.T153745A120103207. | | |
| **BAC7 What is the native range of the *Taxon*?** | | |

| | |
|---|---|
| Response: Austria \| Azerbaijan \| Belarus \| Bosnia and Herzegovina\| Bulgaria \| Croatia \| Georgia \| Greece \| Hungary \| Iran \| Israel \| Kazakhstan \| Kyrgyzstan \| Moldova \| Romania \| Russian Federation \| Serbia \| Slovakia \| Turkey \| Turkmenistan \| Ukraine | Confidence: Medium |

Comments: Native to several eastern European countries (Kouba et al. 2014; Gherardi & Souty-Grousset 2017)

References:

Gherardi, F. & Souty-Grosset C. 2017. *Pontastacus leptodactylus. The IUCN Red List of Threatened Species 2017*: e.T153745A120103207.

Kouba, A., Petrusek, A. & Kozák, P., 2014, 'Continental-wide distribution of crayfish species in Europe: update and maps', *Knowledge Management of Aquatic Ecosystems*, 413, 5, https://doi.org/10.1051/kmae/2014007

**BAC8 What is the global alien range of the *Taxon*?**

| | |
|---|---|
| Response: Armenia, Belgium, Czech Republic, Denmark, Finland, France, Germany, Italy, Latvia, Lithuania, Luxembourg, Netherlands, Poland, Switzerland, United Kingdom (Great Britain), Uzbekistan | Confidence: Low |

Comments: *Pontastacus leptodactylus* has been introduced into several western European countries (Kouba et al. 2014; Gherardi & Souty-Grousset 2017)

References:

Gherardi, F. & Souty-Grosset C. 2017. *Pontastacus leptodactylus. The IUCN Red List of Threatened Species 2017*: e.T153745A120103207.

Kouba, A., Petrusek, A. & Kozák, P., 2014, 'Continental-wide distribution of crayfish species in Europe: update and maps', *Knowledge Management of Aquatic Ecosystems*, 413, 5, https://doi.org/10.1051/kmae/2014007

**BAC9 Geographic scope = the *Area* under consideration**

*Area* of assessment: South Africa

Comments: The geographic scope of this assessment is limited to South Africa.

**BAC10 Is the *Taxon* present in the *Area*?**

| | |
|---|---|
| Response: No | Confidence: Medium |

Comments: There are no records of *Pontastacus leptodactylus* being present in South Africa (Nunes et al. 2017).

References:

Nunes, A.L., Zengeya, T.A., Measey, G.J. & Weyl, O.L.F., 2017b, 'Freshwater crayfish invasions in South Africa: past, present and potential future', *African Journal of Aquatic Science* 42, 309–323, https://doi.org/10.2989/16085914.2017.1405788

**BAC11 Availability of physical specimen**

| | |
|---|---|
| Response: No | Confidence in ID: |

Herbarium or museum accession number: *Pontastacus leptodactylus is* not present in South Africa (Nunes et al. 2017)

References:

Nunes, A.L., Zengeya, T.A., Measey, G.J. & Weyl, O.L.F., 2017b, 'Freshwater crayfish invasions in South Africa: past, present and potential future', *African Journal of Aquatic Science* 42, 309–323, https://doi.org/10.2989/16085914.2017.1405788

**BAC12 Is the *Taxon* native to the *Area* or part of the *Area*?**

| | | |
|---|---|---|
| The *Taxon* is native to (part of) the *Area*. | No | Confidence: High |
| The *Taxon* is alien in (part of) the *Area*. | Yes | Confidence: High |

Comments: There are no native crayfish species in South Africa (de Moor 2002, Nunes et al. 2017)

| |
|---|
| References:<br>de Moor, I., 2002, 'Potential impacts of alien freshwater crayfish in South Africa', *African Journal of Aquatic Science* 27, 125–139, , https://doi.org/10.2989/16085914.2002.9626584<br>Nunes, A.L., Zengeya, T.A., Measey, G.J. & Weyl, O.L.F., 2017b, 'Freshwater crayfish invasions in South Africa: past, present and potential future', *African Journal of Aquatic Science* 42, 309–323, https://doi.org/10.2989/16085914.2017.1405788 |

| **BAC13 What is the *Taxon*'s introduction status in the *Area*?** | | |
|---|---|---|
| The *Taxon* is in cultivation/containment. | Don't know | Confidence: Low |
| The *Taxon* is present outside of cultivation/containment. | No | Confidence: Low |
| The *Taxon* has established/naturalised. | No | Confidence: Low |
| The *Taxon* is invasive. | No | Confidence: Low |
| Comments: *Pontastacus leptodactylus* has been introduced into several countries in Europe for aquaculture (Kouba et al. 2014) but there are no verified records of its presence in South Africa (Nunes et al. 2017). | | |
| References:<br>Kouba A, Petrusek A, Kozák P. 2014. Continental-wide distribution of crayfish species in Europe: update and maps. *Knowledge Management of Aquatic Ecosystems* 413:5, https://doi.org/10.1051/kmae/2014007<br>Nunes, A.L., Zengeya, T.A., Measey, G.J. & Weyl, O.L.F., 2017b, 'Freshwater crayfish invasions in South Africa: past, present and potential future', *African Journal of Aquatic Science* 42, 309–323, https://doi.org/10.2989/16085914.2017.1405788 | | |

| **BAC14 Primary (introduction) pathways** | | |
|---|---|---|
| Release | None | Confidence: |
| Escape | Aquaculture \| Pet trade | Confidence: High |
| Contaminant | None | Confidence: |
| Stowaway | None | Confidence: |
| Corridor | None | Confidence: |
| Unaided | None | Confidence: |
| Comments: *Pontastacus leptodactylus* are stocked deliberately for the consumption trade and are also available in the pet trade in Europe (Harlioğlu and Harlioğlu 2004; Chucholl 2013; Faulkes 2015). | | |
| References:<br>Chucholl, C., 2013, 'Invaders for sale: trade and determinants of introduction of ornamental freshwater crayfish', *Biological Invasions* 15, 125–141. https://doi.org/10.1007/s10530-012-0273-2.<br>Faulkes, Z., 2015, 'The global trade in crayfish as pets', *Crustacean Research*, 44, 75–92.<br>Harlioğlu, M.M. & Harlioğlu, A.G., 2004, 'The harvest of freshwater crayfish, *Astacus leptodactylus* (Eschscholtz, 1823) in Turkey', *Reviews in Fish Biology and Fisheries*, 14, 415–419, https://doi.org/10.1007/s11160-005-0812-3 | | |

#### 2. Likelihood

| **LIK1 Likelihood of entry via unaided primary pathways** | |
|---|---|
| Response: Very unlikely (p = 0.0027) | Confidence: Medium |
| Rationale: There is no evidence that *P. leptodactylus* is present in South Africa or has established in the wild in neighbouring countries (Nunes et al. 2017; Madzivanzira et al. 2020) therefore the probability of *P. leptodactylus* entering South Africa through unaided pathways (connected waterways) is very unlikely (p = 0.0027). | |
| References: | |

Madzivanzira, T.C., South, J., Wood, L.E., Nunes, A.L. & Weyl, O.L.F., 2020, 'A review of freshwater crayfish introductions in Africa', *Reviews in Fisheries Science & Aquaculture*, 1-21. https://doi.org/10.1080/23308249.2020.1802405.

Nunes, A.L., Zengeya, T.A., Measey, G.J. & Weyl, O.L.F., 2017, 'Freshwater crayfish invasions in South Africa: past, present and potential future', African Journal of Aquatic Science 42, 309–323, https://doi.org/10.2989/16085914.2017.1405788

| **LIK2 Likelihood of entry via human aided primary pathways** | |
|---|---|
| Response: Fairly probable (p = 0.5) | Confidence: Low |

Rationale: *Pontastacus leptodactylus* are stocked deliberately for the consumption trade and are also available in the pet trade in Europe (Chucholl 2013; Kouba et al. 2014; Faulkes 2015). However, there is no evidence to show that *P. leptodactylus* is currently traded in the South Africa pet trade industry, but this needs to be verified through dedicated surveys.

References:

Chucholl, C., 2013, 'Invaders for sale: trade and determinants of introduction of ornamental freshwater crayfish', *Biological Invasions* 15, 125–141. https://doi.org/10.1007/s10530-012-0273-2.

Faulkes, Z., 2015, 'The global trade in crayfish as pets', *Crustacean Research*, 44, 75–92. https://doi.org/10.18353/crustacea.44.0_75.

Kouba, A., Petrusek, A. & Kozák, P., 2014, 'Continental-wide distribution of crayfish species in Europe: update and maps', *Knowledge Management of Aquatic Ecosystems*, 413, 5, https://doi.org/10.1051/kmae/2014007

| **LIK3 Habitat suitability** | |
|---|---|
| Response: Fairly probable (p = 0.5) | Confidence: Low |

Rationale: *Pontastacus leptodactylus* is a habitat generalist that inhabits both fresh and brackish waters, and in several habitats such as lakes, canals, rivers and estuaries (Gherardi and Souty-Grosset 2017). It can tolerate a wide range of temperature (4-32°C) and salinity fluctuations (4-14 ppt) as well as temporary decreases in oxygen content (down to 2 mg/L) (CABI 2024)

References:

CABI, 2024. *Pontastacus leptodactylus* (Danube crayfish). Invasive Species Compendium, CABI Publishing, Wallingford, UK. https://www.cabidigitallibrary.org/doi/full/10.1079/cabicompendium.72034 (accessed 10 July 2024)

Gherardi, F. & Souty-Grosset, C. 2017, *Pontastacus leptodactylus. The IUCN Red List of Threatened Species 2017*: e.T153745A120103207. http://dx.doi.org/10.2305/IUCN.UK.2017-3.RLTS.T153745A120103207.en

| **LIK4 Climate suitability** | |
|---|---|
| Response: Fairly probable (p = 0.5) | Confidence: Low |

Rationale: *Pontastacus leptodactylus* can tolerate a wide range of temperature (4-32°C) and it is known to occur in several climate types that include temperate/mesothermal (average temperature of coldest month (> 0°C and < 18°C and mean warmest month >10 °C) and continental/microthermal climate (average temperature of coldest month (> 0°C and mean warmest month >10 °C ) (CABI 2024). These climates conditions are available in South Africa.

| References: |
| --- |
| CABI, 2024. *Pontastacus leptodactylus* (Danube crayfish). Invasive Species Compendium, CABI Publishing, Wallingford, UK. https://www.cabidigitallibrary.org/doi/full/10.1079/cabicompendium.72034 (accessed 10 July 2024). |

| **LIK5 Unaided secondary (dispersal) pathways** | |
| --- | --- |
| Response: Very unlikely (p = 0.0027) | Confidence: Low |
| Rationale: *Pontastacus leptodactylus* can disperse naturally in aquatic environments but has limited ability to disperse overland (de Vaate et al. 2002). There are no known populations of *P. leptodactylus* present in countries neighbouring South Africa that could lead to secondary dispersal events through connected waterways (Nunes et al. 2017; Madzivanzira et al. 2020). | |
| References:<br>Madzivanzira, T.C., South, J., Wood, L.E., Nunes, A.L. & Weyl, O.L.F., 2020, 'A review of freshwater crayfish introductions in Africa', *Reviews in Fisheries Science & Aquaculture* 1-21. https://doi.org/10.1080/23308249.2020.1802405<br>de Vaate, A.B., Jazdzewski, K., Ketelaars, H.A.M., Gollasch, S. & Van der Velde, G. 2002, 'Geographical patterns in range extension of Ponto-Caspian macroinvertebrate species in Europe', *Canadian Journal of Fisheries and Aquatic Sciences,* 59(7), 1159-1174.<br>Nunes, A.L., Zengeya, T.A., Measey, G.J. & Weyl, O.L.F., 2017, 'Freshwater crayfish invasions in South Africa: past, present and potential future', *African Journal of Aquatic Science* 42, 309–323, https://doi.org/10.2989/16085914.2017.1405788 | |

| **LIK6 Human aided secondary (dispersal) pathways** | |
| --- | --- |
| Response: Fairly probable (p = 0.5) | Confidence: Low |
| Rationale: Intentional release into the wild through the pet/ aquarium and aquaculture remains a concern. *Pontastacus leptodactylus* are stocked deliberately for the consumption trade and are also available in the pet trade in Europe (Chucholl 2013; Faulkes 2015). | |
| References:<br>Chucholl, C., 2013, 'Invaders for sale: trade and determinants of introduction of ornamental freshwater crayfish', *Biological Invasions* 15, 125–141. https://doi.org/10.1007/s10530-012-0273-2.<br>Faulkes, Z., 2015, 'The global trade in crayfish as pets', *Crustacean Research*, 44, 75–92. https://doi.org/10.18353/crustacea.44.0_75. | |

**3. Consequences**

| **CON1 Environmental impact** | |
| --- | --- |
| **CON1a: Competition** | |
| Response: Data Deficient (DD) | Confidence: |
| Rationale: No documented information available to assess the level of impact. | |
| References: | |

| **CON1b: Predation** | |
|---|---|
| Response: Data Deficient (DD) | Confidence: |
| Rationale: No documented information available to assess the level of impact. | |
| References: | |
| **CON1c: Hybridisation** | |
| Response: Data Deficient (DD) | Confidence: |
| Rationale: No documented information available to assess the level of impact. | |
| References: | |
| **CON1d: Transmission of disease** | |
| Response: Data Deficient (DD) | Confidence: |
| Rationale: No documented information available to assess the level of impact. | |
| References: | |
| **CON1e: Parasitism** | |
| Response: Data Deficient (DD) | Confidence: |
| Rationale: No documented information available to assess the level of impact. | |
| References: | |
| **CON1f: Poisoning/toxicity** | |
| Response: Data Deficient (DD) | Confidence: |
| Rationale: No documented information available to assess the level of impact. | |
| References: | |
| **CON1g: Bio-fouling or other direct physical disturbance** | |
| Response: Data Deficient (DD) | Confidence: |
| Rationale: No documented information available to assess the level of impact. | |
| References: | |
| **CON1h: Grazing/herbivory/browsing** | |
| Response: Data Deficient (DD) | Confidence: |
| Rationale: No documented information available to assess the level of impact. | |
| References: | |
| **CON1i: Chemical, physical or structural impact on ecosystem** | |

| | |
|---|---|
| Response: Data Deficient (DD) | Confidence: |
| Rationale: No documented information available to assess the level of impact. | |
| References: | |
| **CON1k: Indirect impacts through interactions with other species** | |
| Response: Data Deficient (DD) | Confidence: |
| Rationale: No documented information available to assess the level of impact. | |
| References: | |
| **CON1 Maximum environmental impact** | |
| Response: Data Deficient (DD) | Confidence: |
| Rationale: No documented information available to assess the level of impact. | |
| References: | |

| | |
|---|---|
| **CON2 Socio-economic impact** | |
| **CON2a: Safety** | |
| Response: Data Deficient (DD) | Confidence: |
| Rationale: No documented information available to assess the level of impact. | |
| References: | |
| **CON2b: Material and immaterial assets** | |
| Response: Data Deficient (DD) | Confidence: |
| Rationale: No documented information available to assess the level of impact. | |
| References: | |
| **CON2c: Health** | |
| Response: Data Deficient (DD) | Confidence: |
| Rationale: No documented information available to assess the level of impact. | |
| References: | |
| **CON2d: Social, spiritual and cultural relations** | |
| Response: Data Deficient (DD) | Confidence: |
| Rationale: No documented information available to assess the level of impact. | |
| References: | |

| CON2 Maximum socio-economic impact | |
|---|---|
| Response: Data Deficient (DD) | Confidence: |
| Rationale: No documented information available to assess the level of impact. | |
| References: | |

| CON3 Closely related species' environmental impact | |
|---|---|
| Response: Major (MR) | Confidence: Low |

Rationale: There are no recorded environmental impacts for *P. leptodactylus*, therefore *Pacifastacus leniusculus* was selected as a closely related taxon to estimate the environmental impacts that may arise from introductions of *P. leptodactylus*.

**Competition**

*Pacifastacus leniusculus* contributes to the decline of several native species in introduced areas through competition for resources (Dana et al. 2010; Lodge et al. 2012; Weinlader & Furerfer 2002). Significant competition and reproductive interference often result in the displacement and extinction of native species where *P. leniusculus* out-competes native crayfish species (i.e., *Astacus astacus, Austropotamobius torrentium, Cambaroides japonicas*, and *Pacifastacus nigrescens*) for shelter, making them more susceptible to predation (Huber & Schubart 2005; Pockl & Pekny 2002; Westman, Savolainen & Julkunen 2002). There are no native crayfish species in South Africa but introduced crayfish have the potential to outcompete native potamonautid crabs (Madzivanzira et al. 2021).

**Predation**

*Pacifastacus leniusculus* is a functional omnivore feeding on plant material, zoobenthos, detritus, fish, and sometimes other crayfish (Dana et al. 2010; Moorhouse 2018). *Pacifastacus leniusculus* tends to consume more animal than plant material and impacts invertebrate densities through direct predation (Crawford, Yeomans & Adams 2006; Moorhouse 2018). In invaded areas, there is an overall decrease in richness and abundance of native species such as snails, leeches, caddisflies, newts, and fish (Crawford et al. 2006; Galib, Findlay & Lucas 2021; Girdner et al. 2018).

**Transmission of disease**

*Pacifastacus leniusculus* is a vector for crayfish plague, a disease caused by the parasitic oomycete *Aphanomyces astaci* (Lodge et al. 2012; Longshaw 2011). In Europe, the transmission of crayfish plague by *P. leniusculus* to native crayfish species has been linked to a decline in populations and in some case population collapses of several native species of crayfish such as *Astacus astacus*, *Austropotamobius pallipes*, and *A. torrentium* (Dunn et al. 2009; Holdich & Reeve 1991; Holdich et al. 2009). There are no native crayfish species in South Africa, but crayfish plague pathogen has been shown to infect other freshwater decapods such as native potamonautid crabs (Svoboda et al. 2014).

**Chemical, physical or structural impact on ecosystem**

Burrowing activities of *P. leniusculus* have caused structural damage to river banks and increase bank erosion (Holdich et al. 2009). *Pacifastacus leniusculus* is considered to be a non-burrowing species in the native range, but in the invasive range, *P. leniusculus* constructs burrows under rocks and river banks (Dana et al. 2010). In Europe, the burrows can reach high densities, and can have a severe impact on river bank geomorphology, causing them to collapse (Holdich et al. 2009). On the River Lark in the UK, burrowing by *P. leniusculus* has been reported to cause erosion at the rate of one meter per year (Guan 2010). *Pacifastacus leniusculus* were found to have higher diversity of epibiont algae (species that live on the surface of other organisms) than sympatric native species (Falasco et al. 2018). This indicates that they can provide alternative substrate for agal species to grow on and this can lead to changes in community structure and function.

References:

Crawford, L., Yeomans, W.E. & Adams, C.E., 2006, 'The impact of introduced signal crayfish *Pacifastacus leniusculus* on stream invertebrate communities', *Aquatic Conservation: Marine and Freshwater Ecosystems* 16, 611–621. https://doi.org/10.1002/aqc.761.

Dana, E.D., López–Santiago, J., García–de–Lomas, J., García–Ocaña, D.M., Gámez, V. & Ortega, F., 2010, 'Long–term management of the invasive *Pacifastacus leniusculus* (Dana, 1852) in a small mountain stream', *Aquatic Invasions* 5, 317–322, https://doi.org/10.3391/ai.2010.5.3.10.

Dunn, J.C., McClymont, H.E., Christmas, M. & Dunn, A.M., 2009, 'Competition and parasitism in the native white clawed crayfish *Austropotamobius pallipes* and the invasive signal crayfish *Pacifastacus leniusculus* in the UK', *Biological Invasions* 11, 315–324. https://doi.org/10.1007/s10530-008-9249-7.

Galib, S.M., Findlay, J.S. & Lucas, M.C., 2021, 'Strong impacts of signal crayfish invasion on upland stream fish and invertebrate communities', *Freshwater Biology* 66, 223–240. https://doi.org/10.1111/fwb.13631.

Girdner, S.F., Ray, A.M., Buktenica, M.W., Hering, D.K., Mack, J.A. & Umek, J.W., 2018, 'Replacement of a unique population of newts (*Taricha granulosa mazamae*) by introduced signal crayfish (*Pacifastacus leniusculus*) in Crater Lake, Oregon', *Biological Invasions* 20, 721–740. https://doi.org/10.1007/s10530-017-1570-6.

Holdich, D.M. & Reeve, I.D., 1991, 'Distribution of freshwater crayfish in the British Isles, with particular reference to crayfish plague, alien introductions and water quality', *Aquatic Conservation: Marine and Freshwater Ecosystems* 1, 139–158. https://doi.org/10.1002/aqc.3270010204.

Holdich, D.M., Reynolds, J.D. & Sibley, P.J., 2009, 'A review of the ever increasing threat to European crayfish from non–indigenous crayfish species', *Knowledge and Management of Aquatic Ecosystems* 11, 394–395. https://doi.org/10.1051/kmae/2009025.

Huber, M.G. & Schubart, C.D., 2005, 'Distribution and reproductive biology of *Austropotamobius torrentium* in Bavaria and documentation of a contact zone with the alien crayfish *Pacifastacus leniusculus', Bulletin Français de la Pêche et de la Pisciculture* 376, 759–776. https://doi.org/10.1051/kmae:2005031

Lodge, D.M., Deines, A., Gherardi, F., Yeo, D.C., Arcella, T., Baldridge, A.K., et al., 2012, 'Global introductions of crayfishes: Evaluating the impact of species invasions on ecosystem services', *Annual Review of Ecology Evolution Systematics* 43, 449–472. https://doi.org/10.1146/annurev-ecolsys-111511-103919

Longshaw, M., 2011, 'Diseases of crayfish: A review', *Journal of Invertebrate Pathology* 106, 54–70. https://doi.org/10.1016/j.jip.2010.09.013.

Madzivanzira, T.C., South, J. & Weyl, O.L.F., 2021, 'Invasive crayfish outperform Potamonautid crabs at higher temperatures', *Freshwater Biology* 66, 978–991. https://doi.org/10.1111/fwb.13691.

Moorhouse, T.P., Poole, A.E., Evans, L.C., Bradley, D.C. & Macdonald, D.W., 2014, 'Intensive removal of signal crayfish (*Pacifastacus leniusculus*) from rivers increases numbers and taxon richness of macroinvertebrate species', *Ecology and Evolution* 4, 494–504. https://doi.org/10.1002/ece3.903

Pöckl, M. & Pekny, R., 2002. 'Interaction between native and alien species of crayfish in Austria: case studies', *Bulletin Français de la Pêche et de la Pisciculture* 367, 763–776. https://doi.org/10.1051/kmae:2002064.

Svoboda, J., Strand, D.A., Vrålstad, T., Grandjean, F., Edsman, L., Kozák, P., et al., 2014, 'The crayfish plague pathogen can infect freshwater–inhabiting crabs', *Freshwater Biology* 59, 918–929. https://doi–org.uplib.idm.oclc.org/10.1111/fwb.12315.

Weinländer, M. & Füreder, L., 2009, 'The continuing spread of *Pacifastacus leniusculus* in Carinthia (Austria)', *Knowledge and Management of Aquatic Ecosystems* 17, 394–395. https://doi.org/10.1051/kmae/20010011.

Westman, K., Savolainen, R. & Julkunen, M., 2002, 'Replacement of the native crayfish *Astacus astacus* by the introduced species *Pacifastacus leniusculus* in a small, enclosed Finnish lake: A 30–year study', *Ecography* 25, 53–73. https://doi.org/10.1034/j.1600-0587.2002.250107.x.

| CON4 Closely related species' socio-economic impact | |
|---|---|
| Response: Minor (MN) | Confidence: Low |

Rationale: There are no recorded socio-economic impacts for *P. leptodactylus*, therefore *Pacifastacus leniusculus* was selected as a closely related taxon to estimate the socio-economic impacts that may arise from introductions of *P. leptodactylus*.

**Material and immaterial assets**

*Pacifastacus leniusculus* was introduced to Europe to alleviate the pressure on native crayfish such as *Astacus astacus* whose populations had declined due to crayfish plague (Holdich et al. 2009). The introduction of *P. leniusculus* has however contributed to further decline in populations of native crayfish because *P. leniusculus* are carriers of the crayfish plague and have competitively displaced native crayfish in areas of introduction (Dana et al. 2010). *Astacus astacus* is considered to be a more valuable crayfish species than *P. leniusculus,* although the economic losses as a result of the introduction of *P. leniusculus* is unknown (Johnsen & Taugbøl 2010; Kouba et al. 2022). In addition, the burrowing activities of *P. leniusculus* can cause structural damage to river banks and increase bank erosion (Holdich et al. 2009; Kouba et al. 2022).

**Social, spiritual and cultural relations**

*Pacifastacus leniusculus* was introduced to Europe to alleviate the pressure on native crayfish such as *Astacus astacus* whose populations had declined due to crayfish plague (Holdich et al. 2009). This introduction has however contributed to further decline in native crayfish populations because *P. leniusculus* are carriers of the crayfish plague and have competitively displaced native crayfish in areas of introduction (e.g., Dana et al. 2010). The native crayfish species have a high cultural value and in places such as Scandinavia, the loss of *A. astacus* is serious cultural blow (Gherardi 2011).

References:

Dana, E.D., López-Santiago, J., García-de-Lomas, J., García-Ocaña, D.M., Gámez, V. & Ortega, F., 2010, 'Long–term management of the invasive *Pacifastacus leniusculus* (Dana, 1852) in a small mountain stream', *Aquatic Invasions* 5, 317–322. , https://doi.org/10.3391/ai.2010.5.3.10.

Gherardi, F., 2011, 'Towards a sustainable human use of crayfish (Crustacea, Decapoda, Astacidea)', *Knowledge and Management of Aquatic Ecosystems*, 401, 02 https://doi.org/10.1051/kmae/2011038.

Holdich, D.M., Reynolds, J.D. & Sibley, P.J., 2009, 'A review of the ever increasing threat to European crayfish from non–indigenous crayfish species', *Knowledge and Management of Aquatic Ecosystems* 11, 394–395. https://doi.org/10.1051/kmae/2009025.

Johnsen, S.I. & Taugbøl, T., 2010, 'NOBANIS – Invasive Alien Species Fact Sheet – *Pacifastacus leniusculus*', Online Database of the European Network on Invasive Alien Species – NOBANIS www.nobanis.org. [Date of access 13/11/2019].

Kouba, A., Oficialdegui, F.J., Cuthbert, R.N., Kourantidou, M., South, J., Tricarico, E., et al, 2022. 'Identifying economic costs and knowledge gaps of invasive aquatic crustaceans', *Science of the Total Environment*, 813, 152325 https://doi.org/10.1016/j.scitotenv.2021.152325.

| CON5 Potential impact | |
|---|---|
| Response: MR | Confidence: High |

Rationale: Based on the information gathered from the impact assessment for *P. leniusculus.*

*Pacifastacus leniusculus* is a significant competitor and has displaced several species of native freshwater crayfish and other fauna in introduced areas (Dunn et al. 2009; Galib et al. 2021; Huber & Schubart 2005). While South Africa may not have native freshwater crayfish, closely related decapods such as crabs, may have a resource overlap with *P. leniusculus* which may result in competition (Madzivanzira et al. 2021). *Pacifastacus*

| |
|---|
| *leniusculus* is a vector for crayfish plague that caused population crashes of native crayfish species in Europe (Dana et al. 2010; Holdich & Reeves 1991). Crayfish plague can affect other native freshwater decapods such as native potamonautid crabs (Svoboda et al. 2014). Furthermore, *P. leniusculus* is a facultative omnivore, and may cause negative impacts on macroinvertebrates and macrophyte communities (Dunn et al. 2009; Galib et al. 2021; Westman et al. 2002). In areas outside of its native range, *P. leniusculus* constructs burrows which can weaken riverbanks and dam walls, changing their bank geomorphology, leading to erosion (Guan 1994). |
| References:<br>Dana ED, López–Santiago J, García–de–Lomas J, García–Ocaña DM, Gámez V & Ortega F, 2010, 'Long–term management of the invasive *Pacifastacus leniusculus* (Dana, 1852) in a small mountain stream', *Aquatic Invasions* 5 317–322. https://doi.org/10.3391/ai.2010.5.3.10.<br>Dunn, J.C., McClymont, H.E., Christmas, M. & Dunn, A.M., 2009, 'Competition and parasitism in the native white clawed crayfish *Austropotamobius pallipes* and the invasive signal crayfish *Pacifastacus leniusculus* in the UK', *Biological Invasions* 11, 315–324. https://doi.org/10.1007/s10530-008-9249-7.<br>Galib, S.M., Findlay, J.S. & Lucas, M.C., 2021, 'Strong impacts of signal crayfish invasion on upland stream fish and invertebrate communities', *Freshwater Biology* 66, 223–240. https://doi.org/10.1111/fwb.13631.<br>Guan, R.Z., 1994, 'Burrowing behaviour of signal crayfish, *Pacifastacus leniusculus* (Dana) in the River Great Ouse, England', *Freshwater Forum* 4,155–168.<br>Holdich, D.M. & Reeve, I.D., 1991, 'Distribution of freshwater crayfish in the British Isles, with particular reference to crayfish plague, alien introductions and water quality', *Aquatic Conservation: Marine and Freshwater Ecosystems* 1, 139–158. https://doi.org/10.1002/aqc.3270010204.<br>Huber, M.G. & Schubart, C.D., 2005, 'Distribution and reproductive biology of *Austropotamobius torrentium* in Bavaria and documentation of a contact zone with the alien crayfish *Pacifastacus leniusculus*', *Bulletin Français de la Pêche et de la Pisciculture* 376, 759–776. https://doi.org/10.1051/kmae:2005031.<br>Madzivanzira, T.C., South, J. & Weyl, O.L.F., 2021, 'Invasive crayfish outperform Potamonautid crabs at higher temperatures', *Freshwater Biology* 66, 978–991. https://doi.org/10.1111/fwb.13691.<br>Svoboda, J., Strand, D.A., Vrålstad, T., Grandjean, F., Edsman, L., Kozák, P., et al., 2014, 'The crayfish plague pathogen can infect freshwater–inhabiting crabs', *Freshwater Biology* 59, 918–929. https://doi.org/10.1111/fwb.12315<br>Westman, K., Savolainen, R. & Julkunen, M., 2002, 'Replacement of the native crayfish *Astacus astacus* by the introduced species *Pacifastacus leniusculus* in a small, enclosed Finnish lake: A 30–year study', *Ecography* 25, 53–73. . https://doi.org/10.1034/j.1600-0587.2002.250107.x. |

**4. Management**

| **MAN1 What is the feasibility to stop future immigration?** | |
|---|---|
| Response: Medium | Confidence: Low |
| Rationale: *Pontastacus leptodactylus* is a common species in the global pet trade and aquaculture industry (Faulkes 2015). It is suspected that *P. leptodactylus* is present in the pet trade in South Africa, but this needs to be confirmed (Madzivanzira et al. 2021; Nunes et al. 2017). The species is absent in neighbouring countries, so unaided immigration is unlikely. Moreover, it should be possible to regulate intentional introductions legislative regulations and having an effective border surveillance and monitoring strategy (Zengeya & Wilson 2020; Wilson & Kumschick 2024). | |
| References:<br>Faulkes, Z., 2015, 'The global trade in crayfish as pets', *Crustacean Research* 44**,** 75–92. https://doi.org/10.18353/crustacea.44.0_75. | |

Madzivanzira, T.C., South, J., Wood, L.E., Nunes, A.L. & Weyl, O.L.F., 2020, 'A review of freshwater crayfish introductions in Africa', *Reviews in Fisheries Science & Aquaculture* 1-21. https://doi.org/10.1080/23308249.2020.1802405

Nunes, A.L., Zengeya, T.A., Measey, G.J. & Weyl, O.L.F., 2017, 'Freshwater crayfish invasions in South Africa: past, present and potential future', *African Journal of Aquatic Science* 42, 309–323, https://doi.org/10.2989/16085914.2017.1405788

Wilson, J. R., & Kumschick, S., 2024, 'The regulation of alien species in South Africa', *South African Journal of Science*, 120(5/6).

Zengeya, T.A. & Wilson, J.R., (eds.) 2020, *The status of biological invasions and their management in South Africa in 2019*, pp.71, South African National Biodiversity Institute, Kirstenbosch and DSI-NRF Centre of Excellence for Invasion Biology, Stellenbosch. http://dx.doi.org/10.5281/zenodo.3947613.

**MAN2 Benefits of the Taxon**

**MAN2a Socio-economic benefits of the *Taxon***

| Response: Low | Confidence: Low |
|---|---|

Rationale: Globally *P. leptodactylus* is introduced mainly through aquaculture (Kouba et al. 2014), and the pet trade industry (Chucholl 2013; Faulkes 2015). No socio-economic benefits have been recorded to date in South Africa and even if *P. leptodactylus* is present the benefits are likely to be low (Madzivanzira et al. 2020).

References:

Kouba, A., Petrusek, A. & Kozák, P., 2014, 'Continental-wide distribution of crayfish species in Europe: update and maps', *Knowledge Management of Aquatic Ecosystems*, 413, 5. https://doi.org/10.1051/kmae/2014007

Chucholl, C., 2013, 'Invaders for sale: Trade and determinants of introduction of ornamental freshwater crayfish', *Biological Invasions* 15, 125–141. https://doi.org/10.1007/s10530-012-0273-2.

Faulkes, Z., 2015, 'The global trade in crayfish as pets', *Crustacean Research* 44, 75–92. https://doi.org/10.18353/crustacea.44.0_75.

Madzivanzira, T.C., South, J., Wood, L.E., Nunes, A.L. & Weyl, O.L.F., 2020, 'A review of freshwater crayfish introductions in Africa', *Reviews in Fisheries Science & Aquaculture* 1-21. https://doi.org/10.1080/23308249.2020.1802405

**MAN2b Environmental benefits of the *Taxon***

| Response: Low | Confidence: High |
|---|---|

Rationale: There are no recorded environmental benefits of *P. leptodactylus*

References:

**MAN3 Ease of management**

**MAN3a How accessible are populations?**

| Response: 2 (difficult to access) | Confidence: Low |
|---|---|

Rationale: There are no locality records for *P. leptodactylus* in South Africa (Madzivanzira et al. 2020; Nunes et al. 2017). However, given that it is a small burrowing freshwater, accessibility is likely to be difficult (Peeters et al. 2023; CABI 2024).

References:

| | |
|---|---|
| CABI, 2024. *Pontastacus leptodactylus* (Danube crayfish). Invasive Species Compendium, CABI Publishing, Wallingford, UK. https://www.cabidigitallibrary.org/doi/full/10.1079/cabicompendium.72034 (accessed 10 July 2024).<br>Madzivanzira, T.C., South, J., Wood, L.E., Nunes, A.L. & Weyl, O.L.F., 2020, 'A review of freshwater crayfish introductions in Africa', *Reviews in Fisheries Science & Aquaculture* 1-21. https://doi.org/10.1080/23308249.2020.1802405<br>Nunes, A.L., Zengeya, T.A., Measey, G.J. & Weyl, O.L.F., 2017, 'Freshwater crayfish invasions in South Africa: past, present and potential future', *African Journal of Aquatic Science* 42, 309–323, https://doi.org/10.2989/16085914.2017.1405788<br>Peeters, E. T., de Vries, R., Elzinga, J., Ludányi, M., van Himbeeck, R., & Roessink, I. (2023). Triggers affecting crayfish burrowing behaviour. *Aquatic Ecology*, 1-16. https://doi.org/10.1007/s10452-023-10057-3 | |
| **MAN3b Is detectability critically time-dependent?** | |
| Response: 2 (yes) | Confidence: High |
| Rationale: *Pontastacus leptodactylus* is a small burrowing freshwater species that is difficult to detect (Peeters et al. 2023; CABI 2024). | |
| References:<br><br>CABI, 2024. *Pontastacus leptodactylus* (Danube crayfish). Invasive Species Compendium, CABI Publishing, Wallingford, UK. https://www.cabidigitallibrary.org/doi/full/10.1079/cabicompendium.72034 (accessed 10 July 2024).<br>Peeters, E. T., de Vries, R., Elzinga, J., Ludányi, M., van Himbeeck, R., & Roessink, I. (2023). Triggers affecting crayfish burrowing behaviour. *Aquatic Ecology*, 1-16. https://doi.org/10.1007/s10452-023-10057-3 | |
| **MAN3c Time to reproduction** | |
| Response:1 (1–3 years) | Confidence: Medium |
| Rationale: *Pontastacus leptodactylus* reach sexual maturity in three years in length (Berber & Mazlum 2009; CABI 2024). | |
| References:<br><br>CABI, 2024. *Pontastacus leptodactylus* (Danube crayfish). Invasive Species Compendium, CABI Publishing, Wallingford, UK. https://www.cabidigitallibrary.org/doi/full/10.1079/cabicompendium.72034 (accessed 10 July 2024).<br>Berber, S., & Mazlum, Y., 2009, 'Reproductive efficiency of the narrow-clawed crayfish, *Astacus leptodactylus*, in several populations in Turkey', *Crustaceana*, 531-542. https://www.jstor.org/stable/27743310 | |
| **MAN3d Propagule persistence** | |
| Response:0 (<1 year) | Confidence: Medium |
| Rationale: *Pontastacus leptodactylus* are highly fecund, on average a female produces 200–400 eggs (sometimes 700-800 eggs), with parental care for both eggs and hatchlings (Baber & Mazlum 2009; CABI 2024) | |
| References:<br><br>CABI, 2024. *Pontastacus leptodactylus* (Danube crayfish). Invasive Species Compendium, CABI Publishing, Wallingford, UK. https://www.cabidigitallibrary.org/doi/full/10.1079/cabicompendium.72034 (accessed 10 July 2024). | |

Berber, S., & Mazlum, Y., 2009, 'Reproductive efficiency of the narrow-clawed crayfish, *Astacus leptodactylus*, in several populations in Turkey', *Crustaceana*, 531-542. https://www.jstor.org/stable/27743310

| **MAN3 Ease of management** | |
|---|---|
| Response: Medium | Confidence: Medium |

Rationale: *Pontastacus leptodactylus* is a small burrowing freshwater species that is difficult to detect, it is highly fecund (200–400 eggs) and parental cares increases the survival probability of hatchlings (Baber & Mazlum 2009; CABI 2024).

References:

Berber, S., & Mazlum, Y., 2009, 'Reproductive efficiency of the narrow-clawed crayfish, *Astacus leptodactylus*, in several populations in Turkey', *Crustaceana*, 531-542. https://www.jstor.org/stable/27743310

CABI, 2024. *Pontastacus leptodactylus* (Danube crayfish). Invasive Species Compendium, CABI Publishing, Wallingford, UK. https://www.cabidigitallibrary.org/doi/full/10.1079/cabicompendium.72034 (accessed 10 July 2024)

Peeters, E. T., de Vries, R., Elzinga, J., Ludányi, M., van Himbeeck, R., & Roessink, I., 2023, 'Triggers affecting crayfish burrowing behaviour', *Aquatic Ecology*, 1-16. https://doi.org/10.1007/s10452-023-10057-3.

| **MAN4 Has the feasibility of eradication been evaluated?** | |
|---|---|
| Response: No | Confidence: Low |

Rationale: There are no known population of *Pontastacus leptodactylus* in South Africa, therefore eradication feasibility has not been no evaluated (Nunes et al. 2017) but see MAN5 for summary of control efforts used to manage invasive crayfish populations.

References:

Nunes, A.L., Zengeya, T.A., Measey, G.J. & Weyl, O.L.F., 2017, 'Freshwater crayfish invasions in South Africa: past, present and potential future', *African Journal of Aquatic Science* 42, 309–323, https://doi.org/10.2989/16085914.2017.1405788.

**MAN5 Control options and monitoring approaches available for the *Taxon***

Response: Several methods can be used to monitor and control invasive crayfish populations including mechanical trapping (e.g., Barkhuizen et al. 2022; Madzivanzira et al. 2021), electrofishing (Nunes et al. 2017), use of biocides, and biological control (Aquiloni et al 2010; Stebbing et al. 2014). However, there are several issues that make management of crayfish invasions challenging (Gherardi et al. 2011). For example, the use of biocides is often effective for localised populations in dams and ponds (Peay et al. 2019) but they are non-specific and need to be applied with caution (Gherardi et al. 2011). Mechanical control methods are also effective in small discrete water bodies but are expensive and require commitment to follow on to complete eradication (Loureiro et al. 2018). In addition, alien crayfish species are regarded as potential conflict-generating species in South Africa and potential management interventions should consider the concerns of stakeholders to avoid conflict (Zengeya et al. 2017).

References:

Aquiloni, L., Brusconi, S., Cecchinelli, E., Tricarico, E., Mazza, G., Paglianti, A., et al., 2010, 'Biological control of invasive populations of crayfish: the European eel (*Anguilla anguilla*) as a predator of *Procambarus clarkii*', *Biological Invasions*, 12, 3817-3824. https://doi.org/10.1007/s10530-010-9774-z

| |
|---|
| Barkhuizen, L.M., Madzivanzira, T.C. & South, J. 2022, 'Population ecology of a wild population of red swamp crayfish *Procambarus clarkii* (Girard, 1852) in the Free State Province, South Africa and implications for eradication efforts', *BioInvasions Records* 11, 181–191. https://doi.org/10.3391/bir.2022.11.1.18.<br>Gherardi, F., Aquiloni, L., Diéguez-Uribeondo, J. & Tricarico E., 2011, 'Managing invasive crayfish: is there a hope?', *Aquatic Sciences* 73, 185–200. https://doi.org/10.1007/s00027-011-0181-z.<br>Loureiro, T.G., Anastácio, P.M., de Siqueira Bueno, S.L. & Araujo, P.B., 2018, 'Management of invasive populations of the freshwater crayfish *Procambarus clarkii* (Decapoda, Cambaridae): test of a population-control method and proposal of a standard monitoring approach', *Environmental Monitoring Assessment* 190, 559 https://doi.org/10.1007/s10661-018-6942-6.<br>Madzivanzira, T.C, South, J., Nhiwatiwa, T. & Weyl, O.L.F., 2021, 'Standardisation of Australian redclaw crayfish *Cherax quadricarinatus* sampling gear in southern Africa', *Water SA* 47, 380–384. https://hdl.handle.net/10520/ejc-waters-v47-n3-a10.<br>Nunes, A.L., Zengeya, T.A., Hoffman, A.C., Measey, G.J. & Weyl, O.L., 2017, 'Distribution and establishment of the alien Australian redclaw crayfish, *Cherax quadricarinatus*, in South Africa and Swaziland', *PeerJ* 5, 1-21. https://doi.org/10.7717/peerj.3135.<br>Peay, S., Johnsen, S.I, Bean, S.W., Dunn, A.M., Sandodden, R. & Edsman, L, 2019, 'Biocide treatment of invasive signal crayfish: Successes, failures and lessons learned', *Diversity* 11, 29. https://doi.org/10.3390/d11030029.<br>Stebbing, P., Longshaw, M. & Scott, A., 2014, 'Review of methods for the management of nonindigenous crayfish, with particular reference to Great Britain', *Ethology Ecology & Evolution* 26, 204-231. https://doi.org/10.1080/03949370.2014.908326.<br>Woodford, D.J., Ivey, P., Novoa, A., Shackleton, R., Richardson, D., Weyl, O., Van Wilgen, B. and Zengeya, T., 2017, 'Managing conflict-generating invasive species in South Africa: Challenges and trade-offs. *Bothalia* 47: a2160. https://hdl.handle.net/10520/EJC-727738f3d |

| **MAN6 Any other management considerations to highlight?** | |
|---|---|
| Response | No |

**5. Calculations**

**Likelihood = Very unlikely**

| Parameter | Likelihood | Stages | Final assessment |
|---|---|---|---|
| LIK1 | 0.0027 | P(entry) = 0.5 | P (invasion) = 0.125 |
| LIK2 | 0.5 | | |
| LIK3 | 0.5 | P(establishment) = 0.5 | |
| LIK4 | 0.5 | | |
| LIK5 | 0.0027 | P (spread) = 0.5 | |
| LIK6 | 0.5 | | |

**Consequence = Major (MR)**

| Parameter | Mechanism/sector | Response |
|---|---|---|
| CON1 | Maximum environmental impact | DD |

| | | |
|---|---|---|
| CON2 | Maximum socio-economic impact | DD |
| CON3a | Competition | MR |
| CON3b | Predation | MO |
| CON3c | Hybridisation | DD |
| CON3d | Disease transmission | MR |
| CON3e | Parasitism | DD |
| CON3f | Poisoning/toxicity | DD |
| CON3g | Bio-fouling or other direct physical disturbance | DD |
| CON3h | Grazing/herbivory/browsing | DD |
| CON3i | Chemical, physical, structural impact | MO |
| CON3k | Indirect impacts through interactions with other species | DD |
| CON3 | Maximum environmental impact | MR |
| CON4a | Safety | DD |
| CON4b | Material and immaterial assets | MN |
| CON4c | Health | DD |
| CON4d | Social, spiritual and cultural relations | MN |
| CON4 | Maximum socio-economic impact | MN |
| CON5 | Potential impact based on traits, experiments, or models | MR |

**Risk score = High**

| | | Consequences | | | | |
|---|---|---|---|---|---|---|
| | | MC | MN | MO | MR | MV |
| Likelihood | Extremely unlikely | low | low | low | medium | medium |
| | Very unlikely | low | low | low | medium | high |
| | Unlikely | low | low | medium | high | high |
| | Fairly probable | medium | medium | high | high | high |
| | Probable | medium | high | high | high | high |

**Ease of management = Medium**

| Parameter | Question | Response |
|---|---|---|
| MAN3a | How accessible are populations? | 2 |
| MAN3b | Is detectability critically time-dependent? | 2 |
| MAN3c | Time to reproduction | 1 |
| MAN3d | Propagule persistence | 0 |
| **MAN3** | **SUM** | 5 |

**Appendix BAC7**:

**Appendix BAC8(a)**: Global alien range of *Pontastacus leptodactylus. From GBIF: Pontastacus leptodactylus* (Eschscholtz, 1823) in GBIF Secretariat. GBIF Backbone Taxonomy. Checklist *dataset https://doi.org/10.15468/39omei [*accessed via GBIF.org on 2023-10-27]

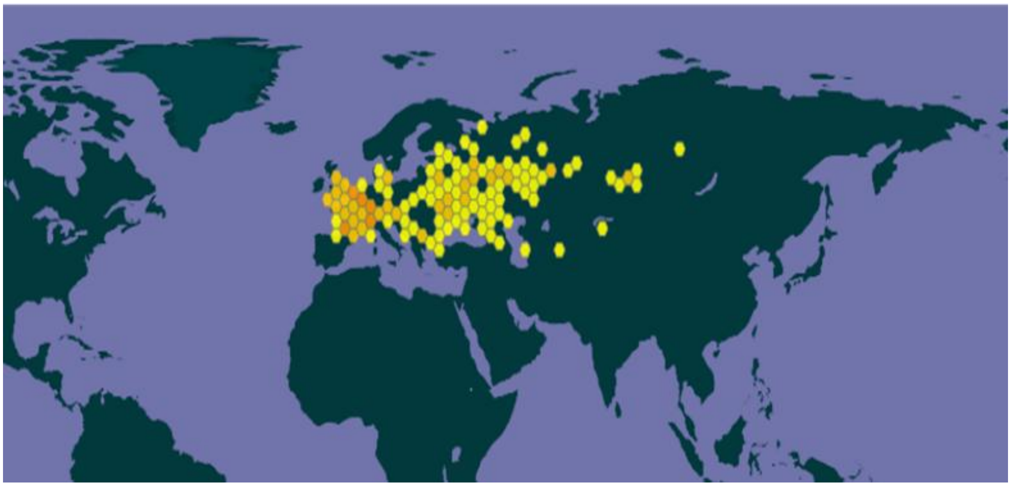

**Appendix BAC8(b)**:

### 1.12 Risk analysis report for White River crayfish (*Procambarus acutus*)

| ***Taxon***: *Procambarus acutus* (Girard, 1852) | ***Area***: South Africa |
|---|---|
| **Picture of *Taxon***<br>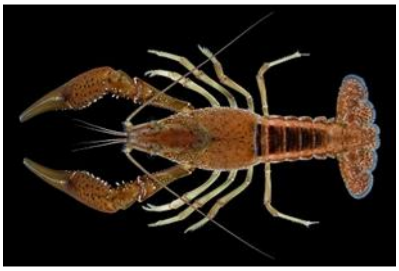<br>https://nas.er.usgs.gov/queries/factsheet.aspx?SpeciesID=216 | **Alien distribution map**<br>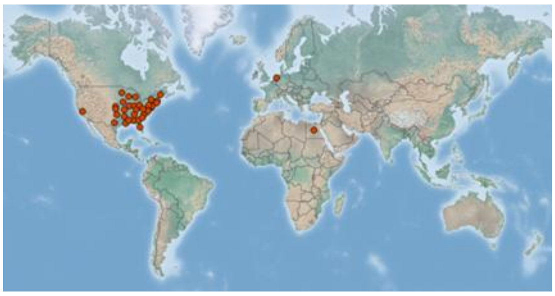<br>https://www.cabi.org/isc/datasheet/67841 |

| | |
|---|---|
| **Risk Assessment summary:**<br><br>The White River crayfish (*Procambarus acutus*) is native to central and eastern USA and has been introduced to several areas outside its natural range in the USA. It has also been introduced to Egypt and the Netherlands. There are no know populations of *P. acutus* in South Africa and in neighbouring countries, therefore via unaided pathways is unlikely. The illegal pet trade industry remains a concern because *P. acutus* may be moved around by humans and there is a high risk of release of unwanted pets. There are no documented impacts from *P. acutus*, and potential impacts were inferred from *Procambarus clarkii*, a closely-related species. *Procambarus clarkii* is a facultative omnivore and an aggressive competitor that often displaces native species in introduced areas through predation, competitive exclusions, and transmission of diseases; has caused habitat alteration through excessive grazing of macrophytes, burrowing activities that increase rates of soil erosion; and has caused physical damage to agricultural infrastructure such as irrigation canals. *Procambarus clarkii* serves as vector for several parasites and diseases, some of which are zoonotic. | **Risk score:**<br><br>Medium |
| **Management options summary:**<br><br>*Procambarus acutus* is a small, largely nocturnal, burrowing freshwater species that is difficult to detect. Several methods can be used to control alien populations of *P. acutus* and these include mechanical removal using traps and electrofishing, biocides, and biological control. However, there are several issues that make the use of these methods challenging. The benefits of *P. acutus* were scored as low. | **Ease of management:**<br><br>Medium |
| **Recommendations:**<br><br>*Procambarus acutus* poses a medium risk to South Africa. If *P. acutus* were to establish outside of captivity, such populations would likely be difficult to extirpate. | **Listing under NEM:BA A&IS lists of 2020:**<br><br>Not listed |

| | |
|---|---|
| *Procambarus acutus* is recommended to be added to a future prohibited species list, active efforts made to confirm that the species is absent in the country, and that any individuals found are controlled. This would likely require a dedicated assessment of the ornamental trade. A reassessment in five years is thus required and, if present in the country, *Procambarus acutus* should be listed under category 1a. | **Recommended listing category:**<br><br>Prohibited list[1] |

[1] The absence of *P. acutus* in South Africa should be confirmed and the taxon should be reassessed in five years (i.e., by 2029).

#### 1. Background

| **BAC1 Name of assessor(s)** | |
|---|---|
| Name of lead assessor | Lee-Anne Botha |
| Additional assessor (1) | Tsungai Zengeya |
| Additional assessor (2) | Christian Chimimba |
| **BAC2 Contact details of assessor (s)** | |
| Lead assessor | Organisational affiliation: Department of Zoology and Entomology, University of Pretoria and The South African National Biodiversity Institute (SANBI). |
| | |
| Additional assessor (1) | Organisational affiliation: Department of Zoology and Entomology, University of Pretoria and The South African National Biodiversity Institute (SANBI). |
| | |
| Additional assessor (2) | Organisational affiliation: Department of Zoology and Entomology, University of Pretoria |
| | |
| **BAC3 Name(s) and contact details of expert(s) consulted** | |
| Expert (1) | Name: |
| | email: |
| | Phone: |
| Expert (2) | Name: |
| | email: |
| | Phone: |
| Comments: none | |

| **BAC4 Scientific name of *Taxon* under assessment** | |
|---|---|
| *Taxon* name: *Procambarus acutus* | Authority: (Girard, 1852) |
| Comments: The name of the taxon under assessment is valid (Crandall & De Grave 2017; GBIF 2023) | |
| References:<br>Crandall, K.A. & De Grave, S., 2017, 'An updated classification of the freshwater crayfishes (Decapoda: Astacidea) of the world, with a complete species list', *Journal of Crustacean Biology*, 27, 615–653. https://doi.org/10.1093/jcbiol/rux070.<br>*Procambarus acutus* (Girard, 1852) in GBIF Secretariat (2023). GBIF Backbone Taxonomy. Checklist dataset https://doi.org/10.15468/39omei accessed via GBIF.org on 2024-07-16. | |
| **BAC5 Synonym(s) considered** | |
| Synonyms: *Cambarus a*cutus Girard, 1852 | |
| Comments: There are several synonyms listed for *P. acutus* (e.g., Crandall & de Grave 2017; CABI 2024) but only *Cambarus acutus* was considered for the literature search for this risk analysis. | |
| References:<br>Crandall, K.A. & De Grave, S., 2017, 'An updated classification of the freshwater crayfishes (Decapoda: Astacidea) of the world, with a complete species list', *Journal of Crustacean Biology*, 27, 615–653. https://doi.org/10.1093/jcbiol/rux070.<br>CABI, 2024. *Procambarus acutus acutus*. Invasive Species Compendium, CABI Publishing, Wallingford, UK. https://www.cabidigitallibrary.org/doi/10.1079/cabicompendium.67841 (accessed 10 July 2024) | |
| **BAC6 Common name(s) considered** | |
| Common names: White River crayfish \| eastern white river crayfish | |
| Comments: The above listed common English names for *P. acutus* were considered for the literature search for this risk analysis (CABI, 2024). | |

References:

CABI, 2024. *Procambarus acutus acutus*. Invasive Species Compendium, CABI Publishing, Wallingford, UK. https://www.cabidigitallibrary.org/doi/10.1079/cabicompendium.67841 (accessed 10 July 2024)

**BAC7 What is the native range of the *Taxon*?** (add map in Appendix BAC7)

| | |
|---|---|
| Response: Coastal plain from Maine to Georgia, and from the Florida panhandle to Texas, and Minnesota to Ohio. | Confidence: High |

Comments: Native range encompasses most of central and eastern USA (Loughman 2007; Crandall 2010)

References:

Crandall, K.A., 2010, *Procambarus* acutus. The IUCN Red List of Threatened Species 2010: e.T154022A4577805. http://dx.doi.org/10.2305/IUCN.UK.2010-3.RLTS.T154022A4577805.en

Loughman, Z., 2007, 'First Record of *Procambarus* (*Ortmannicus*) *acutus* (White River Crayfish) in West Viriginia, with notes on its natural history', *Northeastern Naturalist* 14(3), 495-500. https://doi.org/10.1656/1092-6194(2007)14[495:FROPOA]2.0.CO;2.

**BAC8 What is the global alien range of the *Taxon*?** (add map in Appendix BAC8)

| | |
|---|---|
| Response: Egypt, Netherlands, and USA (California, Connecticut, Kentucky, Maine, Massachusetts, and Rhode Island). | Confidence: Medium |

Comments: Introduced to several areas outside its natural range in the USA and was also introduced to Egypt and the Netherlands (Ibrahim et al. 1997; Taylor et al. 2007; Holdich et al. 2009; Soes & Koese 2010, CABI 2024).

References:

Holdich, D.M., Reynolds, J.D., & Sibley, P.J., 2009, 'A review of the ever increasing threat to European crayfish from non-indigenous crayfish species', *Knowledge and Management of Aquatic Ecosystems* 11, 394–395. https://doi.org/10.1051/kmae/2009025

Soes, D.M. & Koese, B., 2010, Invasive crayfish in the Netherlands: a preliminary risk analysis. *Interim report, Bureau Waardenburg bv, Stichting EIS-Nederland, Invasive Alien Species Team, Waardenburg*.

Ibrahim, A.M., Magdy, T.K. & Murbarak, M.F., 1997, 'Ecological studies on the exotic crayfishes, *Procambarus clarkii* and *Procambarus zonangulus* in the River Nile, Egypt. *International Journal of Ecology and Environmental Sciences*, 23, 217-228.

CABI, 2024. *Procambarus acutus acutus*. Invasive Species Compendium, CABI Publishing, Wallingford, UK. https://www.cabidigitallibrary.org/doi/10.1079/cabicompendium.67841 (accessed 10 July 2024)

**BAC9 Geographic scope = the *Area* under consideration**

*Area* of assessment: South Africa

Comments: The geographic scope of this assessment is limited to South Africa

**BAC10 Is the *Taxon* present in the *Area*?**

| | |
|---|---|
| Response: No | Confidence: High |

Comments: There are no records of *P. acutus* in South Africa (Nunes et al. 2017).

References:

Nunes, A.L., Zengeya, T.A., Measey, G.J. & Weyl, O.L.F., 2017b, 'Freshwater crayfish invasions in South Africa: past, present and potential future', *African Journal of Aquatic Science* 42, 309–323. https://doi.org/10.2989/16085914.2017.1405788

**BAC11 Availability of physical specimen**

| | |
|---|---|
| Response: No | Confidence in ID: |

Herbarium or museum accession number: There are no records of *P. acutus* in South Africa (Nunes et al. 2017).

References:

Nunes, A.L., Zengeya, T.A., Measey, G.J. & Weyl, O.L.F., 2017b, 'Freshwater crayfish invasions in South Africa: past, present and potential future', *African Journal of Aquatic Science* 42, 309–323. https://doi.org/10.2989/16085914.2017.1405788

**BAC12 Is the *Taxon* native to the *Area* or part of the *Area*?**

| | | |
|---|---|---|
| The *Taxon* is native to (part of) the *Area*. | No | Confidence: High |

| | | |
|---|---|---|
| The *Taxon* is alien in (part of) the *Area*. | Yes | Confidence: High |

Comments: There are no native crayfish species in South Africa (de Moor 2002, Nunes et al. 2017).

References:

de Moor I., 2002., ‘Potential impacts of alien freshwater crayfish in South Africa’, *African Journal of Aquatic Science*, 27, 125–139. https://doi.org/10.2989/16085914.2002.9626584.

Nunes, A.L., Zengeya, T.A., Measey, G.J. & Weyl, O.L.F., 2017, ‘Freshwater crayfish invasions in South Africa: past, present and potential future’, *African Journal of Aquatic Science*, 42, 309–323. https://doi.org/10.2989/16085914.2017.1405788.

**BAC13 What is the *Taxon*’s introduction status in the *Area*?**

| | | |
|---|---|---|
| The *Taxon* is in cultivation/containment. | Don’t known | Confidence: Low |
| The *Taxon* is present in the wild. | None | Confidence: High |
| The *Taxon* has established/naturalised. | None | Confidence: High |
| The *Taxon* is invasive. | None | Confidence: High |

Comments: *Procambarus acutus* has been introduced into areas in USA for aquaculture and as live bait (CABI 2024) but there are no verified records of its presence in South Africa (Nunes et al. 2017).

References:

CABI, 2024. *Procambarus acutus acutus*. Invasive Species Compendium, CABI Publishing, Wallingford, UK. https://www.cabidigitallibrary.org/doi/10.1079/cabicompendium.67841 (accessed 10 July 2024)

Nunes, A.L., Zengeya, T.A., Measey, G.J. & Weyl, O.L.F., 2017b, ‘Freshwater crayfish invasions in South Africa: past, present and potential future’, *African Journal of Aquatic Science* 42, 309–323. https://doi.org/10.2989/16085914.2017.1405788.

**BAC14 Primary (introduction) pathways**

| | | |
|---|---|---|
| Release | None | Confidence: |
| Escape | Pet trade \| Aquaculture \| Live food and bait | Confidence: Medium |
| Contaminant | Stock augmentation | Confidence: Low |
| Stowaway | None | Confidence: |
| Corridor | None | Confidence: |
| Unaided | None | Confidence: |

Comments:

*Procambarus acutus* has been introduced intentionally into several areas in the USA (e.g., California and New England) for aquaculture purposes and used as live bait (Holdich et al. 2009; Soes & Koese 2010; Faulkes 2015). The possibility remains that it has been moved unintentionally as a contaminant of *P. clarkii*’s stocks (Soes & Koese 2010).

References:

Faulkes, Z., 2015, ‘The global trade in crayfish as pets’, *Crustacean Research*, 44, 75–92.

Holdich, D.M., Reynolds, J.D., & Sibley, P.J., 2009, ‘A review of the ever increasing threat to European crayfish from non-indigenous crayfish species’, *Knowledge and Management of Aquatic Ecosystems* 11, 394–395. https://doi.org/10.1051/kmae/2009025.

Soes, D.M. & Koese, B., 2010, ‘Invasive crayfish in the Netherlands: A preliminary risk analysis. *Interim report, Bureau Waardenburg bv, Stichting EIS-Nederland’*, Invasive Alien Species Team, Waardenburghttps://nas.er.usgs.gov/queries/factsheet.aspx?SpeciesID=216

#### 2. Likelihood

| **LIK1 Likelihood of entry via unaided primary pathways** | |
|---|---|
| Response: Very unlikely (p = 0.0027) | Confidence: Medium |
| Rationale: There is no evidence that *P. acutus* is present in South Africa or has established in the wild in neighbouring countries (Nunes et al. 2017; Madzivanzira et al. 2020) therefore the probability of *F. rusticus* entering South Africa through unaided pathways (connected waterways) is very unlikely (p = 0.0027). | |
| References:<br><br>Madzivanzira, T.C., South, J., Wood, L.E., Nunes, A.L. & Weyl, O.L.F., 2020, 'A review of freshwater crayfish introductions in Africa', *Reviews in Fisheries Science & Aquaculture*, 1-21. https://doi.org/10.1080/23308249.2020.1802405.<br>Nunes, A.L., Zengeya, T.A., Measey, G.J. & Weyl, O.L.F., 2017, 'Freshwater crayfish invasions in South Africa: past, present and potential future', African Journal of Aquatic Science 42, 309–323. https://doi.org/10.2989/16085914.2017.1405788 | |

| **LIK2 Likelihood of entry via human aided primary pathways** | |
|---|---|
| Response: Unlikely (p = 0.027) | Confidence: Low |
| Rationale: *Procambarus acutus* has been introduced into areas outside its native range in USA for aquaculture and as live bait (CABI 2024). It is not very popular in the aquarium pet trade (Faulkes 2015) and there is no evidence to show that *P. acutus* is currently traded in the South Africa pet trade industry, but this needs to be verified through dedicated surveys (Nunes et al. 2017) | |
| References:<br><br>CABI, 2024. *Procambarus acutus acutus*. Invasive Species Compendium, CABI Publishing, Wallingford, UK. https://www.cabidigitallibrary.org/doi/10.1079/cabicompendium.67841 (accessed 10 July 2024).<br>Faulkes, Z., 2015, 'The global trade in crayfish as pets', *Crustacean Research*, 44, 75–92. https://doi.org/10.18353/crustacea.44.0_75.<br>Nunes, A.L., Zengeya, T.A., Measey, G.J. & Weyl, O.L.F., 2017b, 'Freshwater crayfish invasions in South Africa: past, present and potential future', *African Journal of Aquatic Science* 42, 309–323. https://doi.org/10.2989/16085914.2017.1405788. | |

| **LIK3 Habitat suitability** | |
|---|---|
| Response: Fairly probable (p = 0.5) | Confidence: High |
| Rationale: *Procambarus acutus* is a habitat generalist and is widely tolerant in most riverine habitats and it prefers still or slow flowing waters with mud, sand or gravel substrates (Crandall 2010, Soes and Kose 2010). It constructs simple shallow burrows to survive temporary harsh environmental conditions (Crandall 2010; Soes & Koese 2010). These habitats are available in South Africa. | |
| References:<br><br>Crandall, K.A., 2010, *Procambarus acutus*. The IUCN Red List of Threatened Species 2010: e.T154022A4577805. http://dx.doi.org/10.2305/IUCN.UK.2010-3.RLTS.T154022A4577805.en<br>Soes, D.M., Koese, B., 2010, Invasive crayfish in the Netherlands: a preliminary risk analysis. *Interim report, Bureau Waardenburg bv, Stichting EIS-Nederland, Invasive Alien Species Team, Waardenburg*. | |

| **LIK4 Climate suitability** | |
|---|---|
| Response: Fairly probable (p = 0.5) | Confidence: Low |
| Rationale: *Procambarus acutus* is known to prefer temperate/mesothermal climate (average temperature of coldest month (> 0°C and < 18°C and mean warmest month >10 °C) (CABI 2024). These climate conditions are available in South Africa. | |
| References:<br><br>CABI, 2024. *Procambarus acutus acutus*. Invasive Species Compendium, CABI Publishing, Wallingford, UK. https://www.cabidigitallibrary.org/doi/10.1079/cabicompendium.67841 (accessed 10 July 2024). | |

| **LIK5 Unaided secondary (dispersal) pathways** | |
|---|---|
| Response: Very unlikely (p = 0.0027) | Confidence: High |
| Rationale: There are no known populations of *P. acuctus* in countries neighbouring South Africa that could lead to secondary dispersal events through connected waterways (Nunes et al. 2017; Madzivanzira et al. 2020). | |
| References:<br><br>Madzivanzira, T.C., South, J., Wood, L.E., Nunes, A.L. & Weyl, O.L.F., 2020, 'A review of freshwater crayfish introductions in Africa', *Reviews in Fisheries Science & Aquaculture* 1-21. https://doi.org/10.1080/23308249.2020.1802405<br>Nunes, A.L., Zengeya, T.A., Measey, G.J. & Weyl, O.L.F., 2017b, 'Freshwater crayfish invasions in South Africa: past, present and potential future', *African Journal of Aquatic Science* 42, 309–323. https://doi.org/10.2989/16085914.2017.1405788. | |

| **LIK6 Human aided secondary (dispersal) pathways** | |
|---|---|
| Response: Fairly probable (p = 0.5) | Confidence: Low |
| Rationale: Intentional release into the wild through the pet trade, live bait and aquaculture remains a concern. *Procambarus acutus* has been introduced into areas outside its natural range through aquaculture, live bait and aquarium pet trade (Faulkes 2015, CABI 2024). | |
| References:<br>CABI, 2024. *Procambarus acutus acutus*. Invasive Species Compendium, CABI Publishing, Wallingford, UK. https://www.cabidigitallibrary.org/doi/10.1079/cabicompendium.67841 (accessed 10 July 2024).<br>Faulkes, Z., 2015, 'The global trade in crayfish as pets', *Crustacean Research*, 44, 75–92. https://doi.org/10.18353/crustacea.44.0_75. | |

**3. Consequences**

| CON1 Environmental impact | |
|---|---|
| **CON1a: Competition** | |
| Response: Data Deficient (DD) | Confidence: |
| Rationale: No documented information available to assess the level of impact. | |
| References: | |
| **CON1b: Predation** | |
| Response: Data Deficient (DD) | Confidence: |
| Rationale: No documented information available to assess the level of impact. | |
| References: | |
| **CON1c: Hybridisation** | |
| Response: Data Deficient (DD) | Confidence: |
| Rationale: No documented information available to assess the level of impact. | |
| References: | |
| **CON1d: Transmission of disease** | |
| Response: Data Deficient (DD) | Confidence: |
| Rationale: No documented information available to assess the level of impact. | |
| References: | |
| **CON1e: Parasitism** | |
| Response: Data Deficient (DD) | Confidence: |
| Rationale: No documented information available to assess the level of impact. | |
| References: | |
| **CON1f: Poisoning/toxicity** | |
| Response: Data Deficient (DD) | Confidence: |
| Rationale: No documented information available to assess the level of impact. | |
| References: | |
| **CON1g: Bio-fouling or other direct physical disturbance** | |
| Response: Data Deficient (DD) | Confidence: |
| Rationale: No documented information available to assess the level of impact. | |

| |
|---|
| References: |

| | |
|---|---|
| **CON1h: Grazing/herbivory/browsing** | |
| Response: Data Deficient (DD) | Confidence: |
| Rationale: No documented information available to assess the level of impact. | |
| References: | |
| **CON1i: Chemical, physical or structural impact on ecosystem** | |
| Response: Data Deficient (DD) | Confidence: |
| Rationale: No documented information available to assess the level of impact. | |
| References: | |
| **CON1k: Indirect impacts through interactions with other species** | |
| Response: Data Deficient (DD) | Confidence: |
| Rationale: No documented information available to assess the level of impact. | |
| References: | |
| **CON1 Maximum environmental impact** | |
| Response: Data Deficient (DD) | Confidence: |
| Rationale: No documented information available to assess the level of impact. | |
| References: | |

| | |
|---|---|
| **CON2 Socio-economic impact** | |
| **CON2a: Safety** | |
| Response: | Confidence: Low |
| Rationale: No documented information available to assess the level of impact. | |
| References: | |
| **CON2b: Material and immaterial assets** | |
| Response: | Confidence: Low |
| Rationale: No documented information available to assess the level of impact. | |
| References: | |
| **CON2c: Health** | |

| | |
|---|---|
| Response: | Confidence: Low |
| Rationale: No documented information available to assess the level of impact. | |
| References: | |
| **CON2d: Social, spiritual and cultural relations** | |
| Response: | Confidence: Low |
| Rationale: No documented information available to assess the level of impact. | |
| References: | |
| **CON2 Maximum socio-economic impact** (Figure S3) | |
| Response: | Confidence: Low |
| Rationale: No documented information available to assess the level of impact. | |
| References: | |

| **CON3 Closely related species' environmental impact** | |
|---|---|
| Response: Major (MR) | Confidence: High |
| Rationale: There are no recorded environmental impacts for *P. acutus*, therefore *Procambarus clarkii* was selected as a closely related taxon to estimate the environmental impacts that may arise from introductions of *P. acutus*.<br><br>*Procambarus clarkii* is an aggressive species and is known to out-competes native species for shelter and spawning sites leading to reproductive interference (Cruz et al. 2006; Lodge et al. 2012). For example, in areas were *P. clarkii* has been introduced, some amphibian species (e.g., *Bufo bufo*, *B. calamita*, *Rana* spp., *Taricha torosa* and *Triturus vulgaris*) have been excluded or displaced from their natural habitats, resulting in local extinctions through reproductive failure (Cruz et al. 2006; Lodge et al. 2012). In Europe and Japan, decreases in distributional ranges and abundance of native crayfish populations (*Astacus astacus*, *Austropotamobius pallipes* and *A. torrentium*) as a direct result of competition with *P. clarkii* were observed (Cruz et al. 2006; Lodge et al. 2012). Furthermore, direct competition for food has caused dietary niche constriction and declines in populations of native crabs in some areas invaded by *P. clarkii* (Jackson et al. 2016).<br><br>Predation by *P. clarkii* in the alien range can result in local or population extinction of at least one native species (Lodge et al. 2012; Souty-Grosset et al. 2016). For example, predation by *P. clarkii* has resulted in population declines of several species of fish and amphibians by reducing their breeding success through predation on eggs and larval amphibians (Cruz et al. 2006; Ficetola et al. 2011). Indirect effects of predation results in cause trophic cascades that lead to changes in ecosystem structure and function. For example, predation on invertebrates can release algae from grazing pressure and lead to changes in the abundance and dominance of species in algal communities (Gherardi & Barbaresi 2008).<br><br>*Procambarus clarkii* is a host of various pathogens, parasites, and diseases (Longshaw 2011; Lodge et al. 2012). Transmission of diseases and parasites by *P. clarkii* has led to local or population extinctions of at least one native species, leading to changes in community composition (Lodge et al. 2012). For example, *P. clarkii* is a vector of crayfish plague, a disease caused by the parasitic oomycete, *Aphanomyces astaci* (Aquiloni et al, | |

2011; Longshaw 2011; Souty-Grosset et al. 2016). In Europe, transmission of crayfish plague by *P. clarkii* has been linked to a decline in populations of several native crayfish species of as *Astacus astacus*, *Austropotamobius pallipes* and *A. torrentium* (Holdich et al. 2009; Longshaw 2011). *Procambarus clarkii* is also host to white spot syndrome disease – a viral infection of crustaceans, and fungal pathogens such as *Batrachochytrium dendrobatids* that causes chytridiomycosis – a lethal skin infection in amphibians (Longshaw 2011; McMahon et al. 2013).

Grazing by *P. clarkii* leads to habitat loss and modification through the removal of macrophytes, in turn leading to population declines of species that feed on macrophytes, use macrophyte stands as nesting sites and refugia from predation (Rosenthal et al. 2005). *Procambarus clarkii* leads community composition changes through trophic cascades (Gherardi & Aquistapace 2007; Souty-Grosset et al. 2016). For example, in Lake Chozas (northwest Spain), grazing by *P. clarkii* caused a reduction in macrophyte communities leading to cascades up the food chain with declines in invertebrates, amphibians, and waterfowl (Rodriguez et al 2003). Grazing can also cause changes to community composition and structure, such as changing ecosystems from macrophyte-dominated areas with clear water, to turbid phytoplankton-dominated areas (Matsizaki et al. 2009). Excessive grazing can also lead to accelerated rates of important processes such as litter breakdown and decomposition (Gherardi & Aquistapace 2007; Souty-Grosset et al. 2016).

*Procambarus clarkii* is considered an ecosystem engineer due to its ability to change ecosystems through its burrowing activities (Souty-Grosset et al. 2016). For example, its burrowing activities can cause a decrease in the water quality through bioturbation leading to increased turbidity and influx release of nutrients from sediments, often leading to algal blooms (Angeler et al. 2001; Yamamoto 2010). The impaired water quality also affects the quality of the habitats for other aquatic fauna (Angeler et al. 2001; Rodriguez et al. 2003). For example, increased turbidity can impede foraging and respiratory processes of fish (Rodriguez et al. 2003). Burrowing activities can cause structural damage to river banks and increase bank erosion, and also cause damage to water retention infrastructure such as dam walls and dykes (Souty-Grosset et al. 2016).

References:

Angeler, D.G., Sánchez-Carrillo, S., García, G. & Alvarez-Cobelas, M., 2001, 'The influence of *Procambarus clarkii* (Cambaridae, Decapoda) on water quality and sediment characteristics in a Spanish floodplain wetland', *Hydrobiologia* 464, 89–98. https://doi.org/10.1023/A:1013950129616.

Aquiloni, L., Martín, M.P., Gherardi, F. & Diéguez-Uribeondo, J., 2011, 'The North American crayfish *Procambarus clarkii* is the carrier of the oomycete *Aphanomyces astaci* in Italy', *Biological Invasions* 13, 35–367. https://doi.org/10.1007/s10530-010-9828-2.

Cruz, M.J., Rebelo, R. & Crespo, E.G., 2006, 'Effects of an introduced crayfish, *Procambarus clarkii*, on the distribution of south-western Iberian amphibians in their breeding habitats', *Ecography* 29, 329–338. https://doi.org/10.1111/j.2006.0906-7590.04333.x.

Jackson, M.C., Grey, J., Miller, K., Britton, J.R. & Donohue, I., 2016, 'Dietary niche constriction when invaders meet natives: Evidence from freshwater decapods', *Journal of Animal Ecology* 85, 1098–1107. https://doi.org/10.1051/kmae:2005031.

Ficetola, G.F., Siesa, M.E., Manenti, R., Bottoni, L., De Bernardi, F. & Padoa-Schioppa, E., 2011, 'Early assessment of the impact of alien species: Differential consequences of an invasive crayfish on adult and larval amphibians', *Diversity and Distributions* 17, 1141–1151. https://doi.org/10.1111/j.1472-4642.2011.00797.x

Gherardi, F. & Acquistapace, P., 2007, 'Invasive crayfish in Europe: The impact of *Procambarus clarkii* on the littoral community of a Mediterranean lake', *Freshwater Biology* 52, 1249–59. https://doi.org/10.1111/j.1365-2427.2007.01760.x.

Gherardi, F. & Barbaresi, S., 2008, 'Feeding opportunism of the red swamp crayfish *Procambarus clarkia*, an invasive species', *Freshwater crayfish* 16, 77–85. https://doi.org/10.1080/1023624021000014761

Holdich, D.M., Reynolds, J.D. & Sibley, P.J., 2009, 'A review of the ever increasing threat to European crayfish from non-indigenous crayfish species', *Knowledge and Management of Aquatic Ecosystems* 11, 394–395. https://doi.org/10.1051/kmae/2009025.

Lodge, D.M., Deines, A., Gherardi, F., Yeo, D.C., Arcella, T., Baldridge, A.K., et al., 2012, 'Global introductions of crayfishes: evaluating the impact of species invasions on ecosystem services', *Annual Review of Ecology, Evolution, and Systematics* 43**,** 449–472. https://doi.org/10.1146/annurev-ecolsys-111511-103919

Longshaw, M. 2011, 'Diseases of crayfish: A review', *Journal of Invertebrate Pathology* 106, 54–70. https://doi.org/10.1016/j.jip.2010.09.013.

Matsuzaki, S.S., Usio, N., Takamura, N. & Washitani, I., 2009, 'Contrasting impacts of invasive engineers on freshwater ecosystems: An experiment and meta-analysis', *Oecologia* 158, 673–686. https://doi.org/10.1007/s00442-008-1180-1.

McMahon, T.A., Brannelly, L.A., Chatfield, M.W., Johnson, P.T., Joseph, M.B., McKenzie, V.J., et al., 2013, 'Chytrid fungus *Batrachochytrium dendrobatidis* has non-amphibian hosts and releases chemicals that cause pathology in the absence of infection', *Proceedings of the National Academy of Sciences of the United States of America* 110, 210–215, https://doi.org/10.1073/pnas.1200592110.

Rodríguez, C.F., Bécares, E. & Fernández-Aláez, M., 2003, 'Shift from clear to turbid phase in Lake Chozas (NW Spain) due to the introduction of American red swamp crayfish (*Procambarus clarkii*)', *Hydrobiologia* 506, 421–26. https://doi.org/10.1023/B:HYDR.0000008626.07042.87.

Rosenthal, S.K., Lodge, D.M., Mavuti, K.M., Muohi, W., Ochieng, P., Mungai, B.N., et al., 2005, 'Comparing macrophyte herbivory by introduced Louisiana crayfish (*Procambarus clarkii*) (Crustacea: Cambaridea) and native Dytiscid beetles (*Cybister tripunctatus*) (Coleoptera: Dytiscidae), in Kenya', *African Journal of Aquatic Science* 30, 157–62 https://doi.org/10.2989/16085910509503850.

Souty-Grosset, C., Anastácio, P.M., Aquiloni, L., Banha, F., Choquer, J., Chucholl, C., et al. 2016. 'The red swamp crayfish *Procambarus clarkii* in Europe: Impacts on aquatic ecosystems and human well-being', *Limnologica* 58, 78–93. https://doi.org/10.1016/j.limno.2016.03.003.

Yamamoto, Y., 2010, 'Contribution of bioturbation by the red swamp crayfish *Procambarus clarkii* to the recruitment of bloom-forming cyanobacteria from sediment', *Journal of Limnology* 69, 102- 111. https://doi.org/10.4081/jlimnol.2010.102.

| CON4 Closely related species' socio-economic impact | |
|---|---|
| Response: Moderate (MO) | Confidence: Medium |

Rationale: There are no recorded socio-economic impacts for *P. acutus*, therefore *Procambarus clarkii* was selected as a closely related taxon to estimate the environmental impacts that may arise from introductions of *P. acutus*.

*Procambarus clarkii* often inhabits agricultural fields in which burrowing activities cause damage to infrastructure (i.e., irrigation canals and dam walls) (Arce & Diéguez-Uribeondo 2015; Lodge et al. 2012), and alters soil hydrology leading to water loss. Grazing causes crop damage and reduces crop yield (Anastácio et al. 2005; Arce & Diéguez-Uribeondo 2015). For example in Europe, *P. clarkii* affects rice production through water loss, damage to rice field banks and ditches, direct consumption of rice seed and plants, and clogging of pipes (Anastácio et al. 2005; Souty-Grosset et al. 2016). *Procambarus clarkii* also damages gill nets and spoils fish caught in nets (Gherardi et al. 2011).

*Procambarus clarkii* bio-accumulates toxins and metals from the environment (Alcorlo  et al. 2016; Gherardi et al. 2002; Souty-Grosset et al. 2016), which are often harmful. These toxins and metals can be transferred to higher food web levels through the consumption of affected crayfish by humans and predators such as otters, birds, and fish (Anda et al. 2001; Gherardi et al. 2011; Lodge et al. 2012). *Procambarus clarkii* is a vector for several parasites and diseases including the parasitic fluke flatworms (*Paragonimus* spp.) that causes lung fluke disease in humans, tularemia-causing bacterium *Francisella tularensis*, rat lungworm *Angiostrongylus cantonensis* that causes meningitis, and the nematode *Gnathostoma spinigerum* that causes human gnathostomiasis (Edgerton et al. 2002; Lane et al 2009; Souty-Grosset et al. 2016).

References:

Anastácio, P.M., Correia, A.M. & Menino, J.P., 2005, 'Processes and patterns of plant destruction by crayfish: effects of crayfish size and developmental stages of rice', *Archiv für Hydrobiologie* 162, 37-51. h https://doi.org/10.1127/0003-9136/2005/0162-0037.

Alcorlo, P., Otero, M., Crehuet, M., Baltanás, A. & Montes, C., 2006, 'The use of the red swamp crayfish (*Procambarus clarkii* Girard) as indicator of the bioavailability of heavy metals in environmental monitoring in the River Guadiamar (SW Spain)', *Science of the Total Environment* 366, 380-–390. https://doi.org/10.1016/j.scitotenv.2006.02.023

Anda, P., Segura del Pozo, J., Díaz García, J.M., Escudero, R., García Peña, F.J., López Velasco, M.C., et al., 2001, 'Waterborne outbreak of tularemia associated with crayfish fishing', *Emerging Infectious Diseases* 7, 575–82. https://doi.org/10.3201/eid0707.010740.

Arce, J.A. & Diéguez-Uribeondo, J., 2015, 'Structural damage caused by the invasive crayfish *Procambarus clarkii* (Girard, 1852) in rice fields of the Iberian Peninsula: A study case', *Fundamental and Applied Limnology* 186, 259-269. https://doi.org/10.1127/fal/2015/0715.

Edgerton, B.F., Evans, L.H., Stephens, F.J. & Overstreet, R.M., 2002, 'Synopsis of freshwater crayfish diseases and commensal organisms', *Aquaculture* 206, 57–135. https://doi.org/10.1016/S0044-8486(01)00865-1.

Gherardi, F., Barbaresi, S., Vaselli, O. & Bencini, A., 2002, 'A comparison of trace metal accumulation in indigenous and alien freshwater macro-decapods', *Marine and Freshwater Behaviour and Physiology* 35, 179–88. https://doi.org/10.1080/1023624021000014761.

Gherardi, F., Robert, F.J., Kenneth, B.B., Mavuti, M., Pacini, N., Grey, J., et al., 2011, 'A review of allodiversity in Lake Naivasha, Kenya: Developing conservation actions to protect East African lakes from the negative impacts of alien species', *Biological Conservation* 144, 2585–2596. https://doi.org/10.1016/j.biocon.2011.07.020.

Lane, M.A., Barsanti, M.C., Santos, C.A., Yeung, M., Lubner, S.J. & Weil, G.J., 2009, 'Human paragonimiasis in North America following ingestion of raw crayfish', *Clinical and Infectious Diseases* 49, 55–61. https://doi.org/10.1086/605534.

Lodge, D.M., Deines, A., Gherardi, F., Yeo, D.C., Arcella, T., Baldridge, A.K., et al., 2012, 'Global introductions of crayfishes: evaluating the impact of species invasions on ecosystem services', *Annual Review of Ecology, Evolution, and Systematics* 43**,** 449–472. https://doi.org/10.1146/annurev-ecolsys-111511-103919

Souty-Grosset, C., Anastácio, P.M., Aquiloni, L., Banha, F., Choquer, J., Chucholl, C., et al. 2016. 'The red swamp crayfish *Procambarus clarkii* in Europe: Impacts on aquatic ecosystems and human well-being', *Limnologica* 58, 78–93. , https://doi.org/10.1016/j.limno.2016.03.003.

| **CON5 Potential impact** | |
|---|---|
| Response: Major (MR) | Confidence: Medium |

Rationale: *Procambarus acutus* and *Procambarus clarkii* share ecological traits (Soes & Koese 2010). This species is an omnivore with high tolerance to environmental conditions, its aggressive nature proved that it could outcompete native species and replace them within an ecosystem. Its intensive grazing and stalk cutting is also a threat to macrophyte communities (Gherardi 2011). *Procambarus acutus* have similar life strategy, therefore it may have the same detrimental impacts in areas of introduction as its invasive congener. Being a functional omnivore, this species may also have detrimental impacts on macroinvertebrates and macrophyte communities in areas where invaded (Gherardi et al. 2011).

References:

Gherardi, F., Britton, J.R., Mavuti, K.M., Pacini, N., Grey, J., Tricarico, E. & Harper, D.M., 2011, 'A review of allodiversity in Lake Naivasha, Kenya: Developing conservation actions to protect East African lakes from the negative impacts of alien species', *Biological Conservation*, 144, 2585–2596. https://doi.org/10.1016/j.biocon.2011.07.020

Nunes, A.L., Zengeya, T.A., Measey, G.J. & Weyl, O.L.F., 2017b, 'Freshwater crayfish invasions in South Africa: past, present and potential future', *African Journal of Aquatic Science* 42, 309–323, https://doi.org/10.2989/16085914.2017.1405788

Soes, D.M. & Koese B. 2010. Invasive crayfish in the Netherlands: a preliminary risk analysis.*Interim report, Bureau Waardenburg bv, Stichting EIS-Nederland, Invasive Alien Species Team, Waardenburg*.

**4. Management**

| MAN1 What is the feasibility to stop future immigration? | |
|---|---|
| Response: High | Confidence: Low |
| Rationale: *Procambarus acutus* is not a common species in the global pet trade and aquaculture industry (Faulkes 2015). The species is absent in neighbouring countries (Madzivanzira et al. 2021), so unaided immigration is unlikely. Moreover, it should be possible to regulate intentional introductions legislative regulations and having an effective border surveillance and monitoring strategy (Zengeya & Wilson 2020). | |
| References:<br><br>Faulkes, Z., 2015., 'The global trade in crayfish as pets', *Crustacean Research*, 44, 75–92. https://doi.org/10.18353/crustacea.44.0_75<br>Madzivanzira, T.C., South, J., Weyl, O.L.F., 2021, 'Invasive crayfish outperform potamonautid crabs at higher temperatures', Freshwater Biology 66, 978–991. https://doi.org/10.1111/fwb.13691<br>Nunes, A.L., Zengeya, T.A., Measey, G.J. & Weyl, O.L.F., 2017, 'Freshwater crayfish invasions in South Africa: past, present and potential future', African Journal of Aquatic Science 42, 309–323, https://doi.org/10.2989/16085914.2017.1405788. | |

| MAN2 Benefits of the Taxon | |
|---|---|
| **MAN2a Socio-economic benefits of the *Taxon*** | |
| Response: Low | Confidence: Low |
| Rationale: *Procambarus acutus* has been introduced into areas outside its native range mainly through aquaculture and as live bait (CABI 2024). No socio-economic benefits have been recorded to date in South Africa are even if *P. acutus* is present the benefits are likely to be low (Madzivanzira et al. 2020). | |
| References:<br>CABI, 2024. *Procambarus acutus acutus*. Invasive Species Compendium, CABI Publishing, Wallingford, UK. https://www.cabidigitallibrary.org/doi/10.1079/cabicompendium.67841 (accessed 10 July 2024).<br>Madzivanzira, T.C., South, J., Wood, L.E., Nunes, A.L. & Weyl, O.L.F., 2020, 'A review of freshwater crayfish introductions in Africa', *Reviews in Fisheries Science & Aquaculture* 1-21. https://doi.org/10.1080/23308249.2020.1802405. | |
| **MAN2b Environmental benefits of the *Taxon*** | |
| Response: None | Confidence: |
| Rationale: There are no known environmental benefits from this taxon | |
| References: | |

| MAN3 Ease of management | |
|---|---|
| **MAN3a How accessible are populations?** | |
| Response: 2 (difficult to access) | Confidence: Low |
| Rationale: There are no locality records for *P.acutus* in South Africa (Madzivanzira et al. 2020; Nunes et al. 2017). However, given that it is an aquatic species, accessibility is likely to be difficult. | |

| |
|---|
| References:<br>Madzivanzira, T.C., South, J., Wood, L.E., Nunes, A.L. & Weyl, O.L.F., 2020, 'A review of freshwater crayfish introductions in Africa', *Reviews in Fisheries Science & Aquaculture* 1-21. https://doi.org/10.1080/23308249.2020.1802405.<br>Nunes, A.L., Zengeya, T.A., Measey, G.J. & Weyl, O.L.F., 2017, 'Freshwater crayfish invasions in South Africa: past, present and potential future', *African Journal of Aquatic Science* 42, 309–323, https://doi.org/10.2989/16085914.2017.1405788 |

| **MAN3b Is detectability critically time-dependent?** | |
|---|---|
| Response: 2 (yes) | Confidence: High |
| Rationale: *Procambarus acutus* is a small burrowing freshwater species that is difficult to detect (Peeters et al. 2023; CABI 2024). | |
| References:<br><br>CABI, 2024. *Procambarus acutus acutus*. Invasive Species Compendium, CABI Publishing, Wallingford, UK. https://www.cabidigitallibrary.org/doi/10.1079/cabicompendium.67841 (accessed 10 July 2024).<br>Peeters, E. T., de Vries, R., Elzinga, J., Ludányi, M., van Himbeeck, R., & Roessink, I., 2023, 'Triggers affecting crayfish burrowing behaviour', *Aquatic Ecology*, 1-16. https://doi.org/10.1007/s10452-023-10057-3 | |

| **MAN3c Time to reproduction** | |
|---|---|
| Response:1 (1–3 years) | Confidence: Medium |
| Rationale: *Procambarus acutus* reach sexual maturity in three years in length (Mazlum 2005; CABI 2024). | |
| References:<br><br>CABI, 2024. *Procambarus acutus acutus*. Invasive Species Compendium, CABI Publishing, Wallingford, UK. https://www.cabidigitallibrary.org/doi/10.1079/cabicompendium.67841 (accessed 10 July 2024).<br>Mazlum, Y., 2005, 'Reproduction of eastern white river crayfish, *Procambarus acutus acutus* (Girard, 1852), in two different habitats', *Journal of Animal and Veterinary Advances*, 4(11), 933-936. | |

| **MAN3d Propagule persistence** | |
|---|---|
| Response:0 (<1 year) | Confidence: Medium |
| Rationale: *Procambarus acutus* is a seasonal spawner that is highly fecund, on average a female produces 32–556 with parental care for both eggs and hatchlings (Mazlum 2005; CABI 2024) | |
| References:<br><br>CABI, 2024. *Procambarus acutus acutus*. Invasive Species Compendium, CABI Publishing, Wallingford, UK. https://www.cabidigitallibrary.org/doi/10.1079/cabicompendium.67841 (accessed 10 July 2024).<br><br>Mazlum, Y., 2005, 'Reproduction of eastern white river crayfish, *Procambarus acutus acutus* (Girard, 1852), in two different habitats', *Journal of Animal and Veterinary Advances*, 4(11), 933-936. | |

| **MAN3 Ease of management** (SUM from Table S4) | |
|---|---|
| Response: Medium | Confidence: Medium |
| Rationale: *Procambarus acutus* is a small burrowing freshwater species that is difficult to detect, it is highly fecund (32–556 eggs) and parental cares increases the survival probability of hatchlings (Mazlum 2005; Peeters et al. 2023; CABI 2024). | |

References:

CABI, 2024. *Procambarus acutus acutus*. Invasive Species Compendium, CABI Publishing, Wallingford, UK. https://www.cabidigitallibrary.org/doi/10.1079/cabicompendium.67841 (accessed 10 July 2024).

Mazlum, Y., 2005, 'Reproduction of eastern white river crayfish, *Procambarus acutus acutus* (Girard, 1852), in two different habitats', *Journal of Animal and Veterinary Advances*, 4(11), 933-936.

Peeters, E. T., de Vries, R., Elzinga, J., Ludányi, M., van Himbeeck, R., & Roessink, I., 2023, 'Triggers affecting crayfish burrowing behaviour', *Aquatic Ecology*, 1-16. https://doi.org/10.1007/s10452-023-10057-3

| **MAN4 Has the feasibility of eradication been evaluated?** | |
|---|---|
| Response: No | Confidence: Low |

Rationale: There are no known population of *Procambarus acutus* in South Africa, therefore eradication feasibility has not been no evaluated (Nunes et al. 2017) but see MAN5 for summary of control efforts used to manage invasive crayfish populations.

References:

Nunes, A.L., Zengeya, T.A., Measey, G.J. & Weyl, O.L.F., 2017, 'Freshwater crayfish invasions in South Africa: past, present and potential future', *African Journal of Aquatic Science* 42, 309–323, https://doi.org/10.2989/16085914.2017.1405788.

**MAN5 Control options and monitoring approaches available for the *Taxon***

Response: Several methods can be used to monitor and control invasive crayfish populations including mechanical trapping (e.g., Barkhuizen et al. 2022; Madzivanzira et al. 2021), electrofishing (Nunes et al. 2017), use of biocides, and biological control (Aquiloni et al 2010; Stebbing et al. 2014). However, there are several issues that make management of crayfish invasions challenging (Gherardi et al. 2011). For example, the use of biocides is often effective for localised populations in dams and ponds (Peay et al. 2019) but they are non-specific and need to be applied with caution (Gherardi et al. 2011). Mechanical control methods are also effective in small discrete water bodies but are expensive and require commitment to follow on to complete eradication (Loureiro et al. 2018). In addition, alien crayfish species are regarded as potential conflict-generating species in South Africa and potential management interventions should consider the concerns of stakeholders to avoid conflict (Zengeya et al. 2017).

References:

Aquiloni, L., Brusconi, S., Cecchinelli, E., Tricarico, E., Mazza, G., Paglianti, A. & Gherardi, F. 2010, 'Biological control of invasive populations of crayfish: the European eel (*Anguilla anguilla*) as a predator of *Procambarus clarkii*', *Biological Invasions* 12 3817-3824. https://doi.org/10.1007/s10530-007-9094-0.

Barkhuizen, L.M, Madzivanzira, T.C., South, J., 2022, 'Population ecology of a wild population of red swamp crayfish *Procambarus clarkii* (Girard, 1852) in the Free State Province, South Africa and implications for eradication efforts', *BioInvasions Records* 11, 181–191. https://doi.org/10.3391/bir.2022.

Gherardi, F., Aquiloni, L., Diéguez-Uribeondo, J. & Tricarico E., 2011, 'Managing invasive crayfish: is there a hope? *Aquatic Sciences* 73, 185–200. https://doi.org/10.1007/s00027-011-0181-z.

Loureiro, T.G, Anastácio, P.M, de Siqueira Bueno, S.L. & Araujo, P.B., 2018, 'Management of invasive populations of the freshwater crayfish *Procambarus clarkii* (Decapoda, Cambaridae): test of a population-control method and proposal of a standard monitoring approach', *Environmental Monitoring Assessment* 190, 559, https://doi.org/10.1007/s10661-018-6942-6.

Madzivanzira, T.C., South, J., Nhiwatiwa, T. & Weyl, O.L.F., 2021, 'Standardisation of Australian redclaw crayfish *Cherax quadricarinatus* sampling gear in southern Africa', *Water SA* 47, 380–384. https://hdl.handle.net/10520/ejc-waters-v47-n3-a10.

Nunes, A.L., Zengeya, T.A., Hoffman, A.C., Measey, G.J. & Weyl, O.L., 2017, 'Distribution and establishment of the alien Australian redclaw crayfish, *Cherax quadricarinatus*, in South Africa and Swaziland', *PeerJ* 5, 1-21. https://doi.org/10.7717/peerj.3135.

Peay, S., Johnsen, S.I, Bean, S.W., Dunn, A.M., Sandodden, R. & Edsman, L, 2019, 'Biocide treatment of invasive signal crayfish: Successes, failures and lessons learned', *Diversity* 11, 29. https://doi.org/10.3390/d11030029

Stebbing, P., Longshaw, M., Scott, A., 2014, 'Review of methods for the management of nonindigenous crayfish, with particular reference to Great Britain', *Ethology Ecology & Evolution* 26, 204-231. https://doi.org/10.1080/03949370.2014.908326.

Woodford, D.J., Ivey, P., Novoa, A., Shackleton, R., Richardson, D., Weyl, O., Van Wilgen, B. and Zengeya, T., 2017, 'Managing conflict-generating invasive species in South Africa: Challenges and trade-offs', *Bothalia* 47, a2160. https://doi. org/10.4102/abc.v47i2.2160.

| **MAN6 Any other management considerations to highlight?** (if yes, fill in Appendix MAN6) | |
|---|---|
| Response | No |

**5. Calculations**

**Likelihood = Very Unlikely**

| Parameter | Likelihood | Stages | Final assessment |
|---|---|---|---|
| LIK1 | 0.0027 | P(entry) = 0.027 | P (invasion) = 0.007 |
| LIK2 | 0.027 | | |
| LIK3 | 0.5 | P(establishment) = 0.5 | |
| LIK4 | 0.5 | | |
| LIK5 | 0.0027 | P (spread) = 0.5 | |
| LIK6 | 0.5 | | |

**Consequence = Major (MR)**

| Parameter | Mechanism/sector | Response |
|---|---|---|
| CON1 | Maximum environmental impact | DD |
| CON2 | Maximum socio-economic impact | DD |
| CON1a | Competition | DD |
| CON1b | Predation | DD |
| CON1c | Hybridisation | DD |

| | | |
|---|---|---|
| CON1d | Disease transmission | DD |
| CON1e | Parasitism | DD |
| CON1f | Poisoning/toxicity | DD |
| CON1g | Bio-fouling or other direct physical disturbance | DD |
| CON1h | Grazing/herbivory/browsing | DD |
| CON1i | Chemical, physical, structural impact | DD |
| CON1k | Indirect impacts through interactions with other species | DD |
| CON2a | Safety | DD |
| CON2b | Material and immaterial assets | DD |
| CON2c | Health | DD |
| CON2d | Social, spiritual and cultural relations | DD |
| CON3 | Maximum environmental impact (closely related taxa) | MR |
| CON4 | Maximum socio-economic impact (closely related taxa) | MO |
| CON5 | Potential impact based on traits, experiments, or models | MR |

**Risk = Medium**

| | | **Consequences** | | | | |
|---|---|---|---|---|---|---|
| | | MC | MN | MO | MR | MV |
| **Likelihood** | Extremely unlikely | low | low | low | medium | medium |
| | Very unlikely | low | low | low | medium | high |
| | Unlikely | low | low | medium | high | high |
| | Fairly probable | medium | medium | high | high | high |
| | Probable | medium | high | high | high | high |

**Easy of management: Medium**

| Parameter | Question | Response |
|---|---|---|
| MAN3a | How accessible are populations? | 2 |
| MAN3b | Is detectability critically time-dependent? | 2 |
| MAN3c | Time to reproduction | 1 |
| MAN3d | Propagule persistence | 0 |
| **MAN3** | **SUM** | 5 |

**Appendix**

**BAC7**: Global alien range of *Procambarus acutus acutus.* Map from CABI: https://www.cabi.org/isc/datasheet/67841

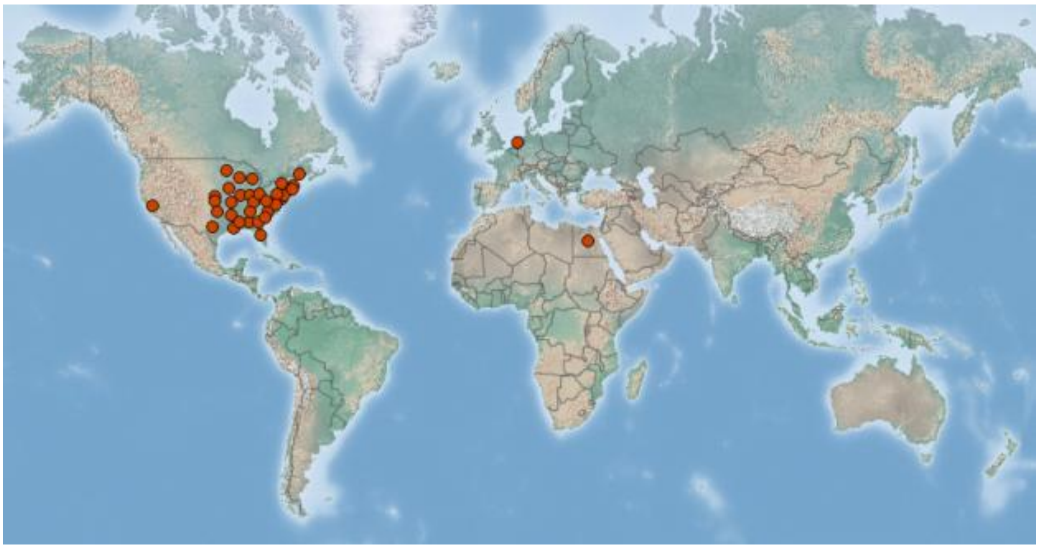

### 1.13 Risk analysis report for Red swamp crayfish (*Procambarus clarkii*)

| **_Taxon:_** *Procambarus clarkii* (Girard, 1852) | **_Area:_** South Africa |
| --- | --- |
| **Picture of _Taxon_**<br>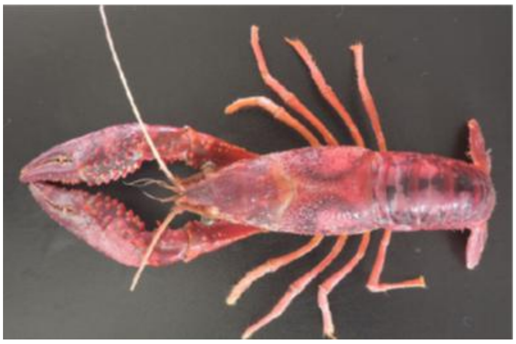<br>Photo: M Antunes (2020), http://dx.doi.org/10.1590/2358-2936e2020037 | **Alien distribution map**<br>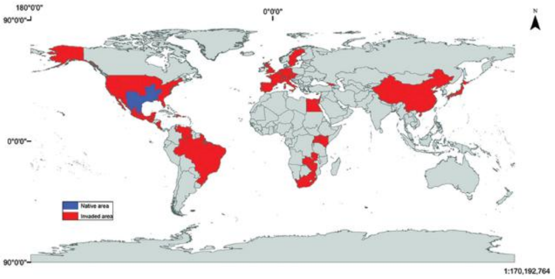<br><br>Reference: Loureiro T.G., Anastácio, P.M.S.G., Araujo, P.B., Souty-Grosset, C. & Almerão, M.P., 2015, 'Red swamp crayfish: biology, ecology and invasion–an overview', *Nauplius* 23(1), 1–19. https://doi.org/10.1590/S0104-64972014002214. |

| **Risk Assessment summary:** | **Risk score:** |
| --- | --- |
| The red swamp crayfish, *Procambarus clarkii* (Girard 1852) is native to northeastern Mexico and south-central USA and has been introduced worldwide for aquaculture and the ornamental pet trade. In South Africa, *P. clarkii* is known to have established outside of captivity in Mpumalanga (Driehoek farm), the Free State (Mimosa Dam) and the Western Cape (Vredendal: lower Olifants and agricultural irrigation systems). The potential for further spread is high because *P. clarkii* is likely to be moved by humans in the aquarium trade industry and *P. clarkii* can disperse over land between water bodies. *Procambarus clarkii* is a facultative omnivore and an aggressive competitor that often displaces native species in introduced areas through predation, competitive exclusions, and transmission of diseases; has caused habitat alteration through excessive grazing of macrophytes, burrowing activities that increase rates of soil erosion; and has caused physical damage to agricultural infrastructure such as irrigation canals. *Procambarus clarkii* serves as vector for several parasites and diseases, some of which are zoonotic. | High |
| **Management summary:**<br><br>As an aquatic species, it is difficult to detect and access *P. clarkii* individuals and populations. Moreover, the burrowing behaviour of *P. clarkii* complicates management efforts as does *P. clarkii*'s rapid and prolific breeding. Management efforts should focus on preventing spread to new areas; and the trade and movement of *P. clarkii* through the pet trade should be assessed. The current distribution of *P. clarkii* means that eradication is likely to be infeasible. Even though *P. clarkii* is considered an important aquacultural species in other countries, this is not the case in South Africa. | **Ease of management:**<br><br>Difficult |
| **Recommendations:**<br><br>*Procambarus clarkii* poses a high risk to South Africa, management is likely to be difficult, and eradication is infeasible. Therefore, *P. clarkii* is recommended to be listed as Category 1b. Public engagement to prevent further spread is recommended. | **Listing under NEM:BA A&IS lists of 2020:**<br><br>Not listed |

| | |
|---|---|
| | **Recommended listing category:**<br><br>1b |

#### 1. Background

| BAC1 Name of assessor(s) | |
|---|---|
| Name of lead assessor | Lee-Anne Botha |
| Additional assessor (1) | Tsungai Zengeya |
| Additional assessor (2) | Christian Chimimba |
| **BAC2 Contact details of assessor (s)** | |
| Lead assessor | Organisational affiliation: South African National Biodiversity Institute |
| | |
| Additional assessor (1) | Organisational affiliation: Department of Zoology and Entomology, University of Pretoria/South African National Biodiversity Institute (SANBI) |
| | |
| Additional assessor (2) | Organisational affiliation: Department of Zoology and Entomology, University of Pretoria |
| | |
| **BAC3 Name(s) and contact details of expert(s) consulted** | |
| Expert (1) | Name: |
| | email: |
| | Phone: |
| Expert (2) | Name: |
| | email: |
| | Phone: |
| Comments: none | |
| **BAC4 Scientific name of *Taxon* under assessment** | |
| *Taxon* name: *Procambarus clarkii* | Authority: (Girard, 1852) |
| Comments: The name of the taxon under assessment is valid (Crandall & De Grave 2017; Integrated Taxonomic Information System 2023). | |
| References:<br>Integrated Taxonomic Information System (ITIS), 2023, '*Procambarus clarkii* (Girard, 1852)', accessed 09 June 2023, from https://www.itis.gov/servlet/SingleRpt/SingleRpt.<br>Crandall, K.A. & De Grave, S., 2017, 'An updated classification of the freshwater crayfishes (Decapoda: Astacidea) of the world, with a complete species list', *Journal of Crustacean Biology* 27, 615–653. https://doi.org/10.1093/jcbiol/rux070. | |
| **BAC5 Synonym(s) considered** | |
| Synonyms: None | |
| Comments: No synonyms were considered. All the literature refers to the species as *P. clarkii* (red swamp crayfish) | |
| References: | |
| **BAC6 Common name(s) considered** | |
| Common names: red swamp crayfish \| Louisiana crayfish | |
| Comments: *Procambarus clarkii* is commonly known as the red swamp crayfish and the Louisiana crayfish (Gherardi 2006). These common names were considered in this assessment. | |
| References:<br>Gherardi, F., 2006, 'Crayfish invading Europe: The case study of *Procambarus clarkii*', *Hydrobiologia* 595, 295–301. https://doi.org/10.1080/10236240600869702. | |
| **BAC7 What is the native range of the *Taxon*?** | |
| Response: North America | Confidence: High |
| Comments: *Procambarus clarkii* is native to north-eastern Mexico and south-central USA (Gherardi 2006; Hobbs 1989). | |

References:
Gherardi, F., 2006, 'Crayfish invading Europe: The case study of *Procambarus clarkii*', *Hydrobiologia* 595, 295–301. https://doi.org/10.1080/10236240600869702.
Hobbs, H., 1989, 'A review of global crayfish introductions with particular emphasis on two North American species (Decapoda, Cambaridae)', *Crustaceana* 56, 299–316. https://doi.org/10.1163/156854089X00275.

**BAC8 What is the global alien range of the *Taxon*?**

| Response: All continents except Antarctica | Confidence: Medium |
|---|---|

Comments: Established populations of *P. clarkii* occurs in the following countries: Belgium, Belize, Brazil; Chile, China, Colombia, Costa Rica, Cyprus, Dominican Republic, Ecuador, Egypt; France, Georgia, Germany, Italy, Japan, Kenya, Mexico, Netherlands, Philippines, Portugal, South Africa, Sudan, Spain, Switzerland, Taiwan, Uganda, United Kingdom, United States of America (Arizona, Arkansas, California, Hawaii, Idaho, Indiana, Maryland, Nevada, New Mexico – Native, North Carolina, Ohio, Oklahoma – Native, Oregon, South Carolina, Utah, Virginia, West Virginia – Present – Origin uncertain), Venezuela (Bolivarian Republic) and Zambia (Lodge et al. 2012; Loureiro et al. 2015). See Appendix BAC8.

References:
Lodge, D.M., Deines, A., Gherardi, F., Yeo, D.C., Arcella, T., Baldridge, A.K., et al., 2012, 'Global introductions of crayfishes: Evaluating the impact of species invasions on ecosystem services', *Annual Review of Ecology Evolution Systematics* 43, 449–472. https://doi.org/10.1146/annurev-ecolsys-111511-103919
Loureiro, T.G., Anastácio, P.M.S.G., Araujo, P.B., Souty-Grosset, C. & Almerão, M.P., 2015, 'Red swamp crayfish: biology, ecology and invasion–an overview', *Nauplius* 23(1), 1–19. . https://doi.org/10.1590/S0104-64972014002214.

**BAC9 Geographic scope = the *Area* under consideration**

*Area* of assessment: South Africa

Comments: The geographic scope of this assessment is limited to South Africa.

**BAC10 Is the *Taxon* present in the *Area*?**

| Response: Yes | Confidence: High |
|---|---|

Comments: The first record of *P. clarkii* was in 1988 at the Driehoek Farm *ca*. 10 km from the town of Dullstroom, Mpumalanga Province (25°28′24.50″S, 30°07′23.61″E) (de Moor 2002; Nunes et al. 2017a, 2017b). *Procambarus clarkii* has since been found in Mimosa Dam (27°58′56.28″S, 26°44′16.79″E) in the Free State Province and Vredendal (lower Olifants and irrigation systems) in the Western Cape (Barkhuizen, Madzivanzira, & South 2022; Weyl et al. 2020). See Appendix BAC10.

References:
Barkhuizen, L.M., Madzivanzira, T.C. & South, J., 2022, 'Population ecology of a wild population of red swamp crayfish *Procambarus clarkii* (Girard, 1852) in the Free State Province, South Africa and implications for eradication efforts', *BioInvasions Records* 11, 181-191. . https://doi.org/10.3391/bir.2022.
de Moor, I., 2002, 'Potential impacts of alien freshwater crayfish in South Africa', *African Journal of Aquatic Science* 27, 125–139. https://doi.org/10.2989/16085914.2002.9626584.
Nunes, A.L., Hoffman, C.A., Zengeya, T.A., Measey, G.J. & Weyl, O.L.F, 2017a, 'Red swamp crayfish, *Procambarus clarkii*, found in South Africa 22 years after attempted eradication', *Aquatic Conservation: Marine & Freshwater Ecosystem* 27, 1334–1340. https://doi.org/10.1002/aqc.2741.
Nunes, A.L., Zengeya, T.A., Measey, G.J. & Weyl, O.L.F., 2017b, 'Freshwater crayfish invasions in South Africa: past, present and potential future', *African Journal of Aquatic Science* 42, 309–323. https://doi.org/10.2989/16085914.2017.1405788.
Weyl, O.L.F., Ellender, B., Wassermann, R.J., Truter, M., Dalu, T., Zengeya, T.A. et al., 2020, 'Alien freshwater fauna in South Africa', In: van Wilgen B.W., Measey, J., Richardson, D.M., Wilson, J.R., Zengeya, T.A. (eds), *Biological invasions in South Africa*, pp 151–182, Springer, Berlin.

**BAC11 Availability of physical specimen**

| Response: Yes | Confidence in ID: Medium |
|---|---|

Herbarium or museum accession number: A sample was collected at one reservoir on the Driehoek Farm. The specimen will be sent to the Albany Museum. As the specimen has not been formally deposited yet, the confidence is scored as medium.

References:

**BAC12 Is the *Taxon* native to the *Area* or part of the *Area*?**

| | | |
|---|---|---|
| The *Taxon* is native to (part of) the *Area*. | No | Confidence: High |
| The *Taxon* is alien in (part of) the *Area*. | Yes | Confidence: High |

Comments: There are no freshwater crayfish species native to South Africa (de Moor 2002; Nunes et al. 2017a, 2017b)

References:

de Moor, I., 2002, ‘Potential impacts of alien freshwater crayfish in South Africa’, *African Journal of Aquatic Science* 27, 125–139. https://doi.org/10.2989/16085914.2002.9626584.

Nunes, A.L., Hoffman, C.A., Zengeya, T.A., Measey, G.J. & Weyl, O.L.F, 2017a, ‘Red swamp crayfish, *Procambarus clarkii*, found in South Africa 22 years after attempted eradication’, *Aquatic Conservation: Marine & Freshwater Ecosystems* 27, 1334–1340. https://doi.org/10.1002/aqc.2741.

Nunes, A.L., Zengeya, T.A., Measey, G.J. & Weyl, O.L.F., 2017b, ‘Freshwater crayfish invasions in South Africa: past, present and potential future’, *African Journal of Aquatic Science* 42, 309–323. https://doi.org/10.2989/16085914.2017.1405788.

**BAC13 What is the *Taxon*’s introduction status in the *Area*?**

| | | |
|---|---|---|
| The *Taxon* is in cultivation/containment. | Unknown | Confidence: Low |
| The *Taxon* is present outside of cultivation/containment. | Yes | Confidence: High |
| The *Taxon* has established/naturalised. | Yes | Confidence: High |
| The *Taxon* is invasive. | Yes | Confidence: Low |

Comments: During a survey at Driehoek Farm in 2016 a specimen of *P. clarkii* was found (22 years after an eradication attempt of the population at the farm) (Nunes et al. 2017a). The specimen was found to be reproductively active, indicating that a small but viable population may still be present. In 2018, a population of *P. clarkii* was found in Mimosa Dam in the Free State province and indications are that the population is established and breeding (Barkhuizen et al. 2022; Madzivanzira et al. 2021). In 2022, *P. clarkii* was also found in the Olifants tributary and agricultural irrigations systems in Vredendal, Western Cape. *Procambarus clarkii* may also be present in the pet trade but this has not been verified (Nunes et al. 2017a, 2017b).

References:

Barkhuizen, L.M., Madzivanzira, T.C. & South, J., 2022, ‘Population ecology of a wild population of red swamp crayfish *Procambarus clarkii* (Girard, 1852) in the Free State Province, South Africa and implications for eradication efforts’, *BioInvasions Records* 11, 181-191. https://doi.org/10.3391/bir.2022.

Madzivanzira, T.C., South, J., Wood, L.E., Nunes, A.L. & Weyl, O.L.F., 2021, ‘A review of freshwater crayfish introductions in Africa’, *Reviews in Fisheries Science & Aquaculture* 29(2), 218-214. https://doi.org/10.1080/23308249.2020.1802405.

Nunes, A.L., Hoffman, C.A., Zengeya, T.A., Measey, G.J. & Weyl, O.L.F, 2017a, ‘Red swamp crayfish, *Procambarus clarkii*, found in South Africa 22 years after attempted eradication’, *Aquatic Conservation: Marine & Freshwater Ecosystem* 27, 1334–1340. https://doi.org/10.1002/aqc.2741.

Nunes, A.L., Zengeya, T.A., Measey, G.J. & Weyl, O.L.F., 2017b, ‘Freshwater crayfish invasions in South Africa: past, present and potential future’, *African Journal of Aquatic Science* 42, 309–323. https://doi.org/10.2989/16085914.2017.1405788.

**BAC14 Primary (introduction) pathways**

| | | |
|---|---|---|
| Release | none | Confidence: Low |
| Escape | aquaculture/mariculture | Confidence: High |
| | pet/aquarium/terrarium species (including live food for fish species) | Confidence: High |
| | live food and live bait | Confidence: High |

| | | |
|---|---|---|
| Contaminant | none | Confidence: High |
| Stowaway | none | Confidence: High |
| Corridor | none | Confidence: High |
| Unaided | none | Confidence: High |
| Comments: The primary pathway of introduction is escape from confinement through aquaculture (Holdich et al. 2009), through the pet trade industry (Faulkes 2015; Patoka et al. 2018), and intentional release as bait for fish and laboratory specimens that are also released facilitating escape (Nunes et al. 2017b) | | |
| References: Faulkes, Z., 2015, 'The global trade in crayfish as pets', *Crustacean Research* 44, 75–92. https://doi.org/10.18353/crustacea.44.0_75. Holdich, D.M., Reynolds, J.D. & Sibley, P.J., 2009, 'A review of the ever increasing threat to European crayfish from non-indigenous crayfish species', *Knowledge and Management of Aquatic Ecosystems* 11, 394–395, https://doi.org/10.1051/kmae/2009025 Nunes, A.L., Zengeya, T.A., Measey, G.J. & Weyl, O.L.F., 2017b, 'Freshwater crayfish invasions in South Africa: past, present and potential future', *African Journal of Aquatic Science* 42, 309–323. https://doi.org/10.2989/16085914.2017.1405788. Patoka, J., Magalhães, A.L.B., Kouba, A., Faulkes, Z., Jerikho, R. & Vitule, J.R.S., 2018, 'Invasive aquatic pets: failed policies increase risks of harmful invasions', *Biodiversity and Conservation* 27, 3037–3046. https://doi.org/10.1007/s10531-018-1581-3 . | | |

**2. Likelihood**

| **LIK1 Likelihood of entry via unaided primary pathways** | |
|---|---|
| Response: Present in the country (p = 1) | Confidence: High |
| Rationale: *Procambarus clarkii* is present in South Africa and neighbouring countries, for example, Zimbabwe (Loureiro et al. 2015; Nunes et al. 2017a; Barkhuizen et al. 2022). In terms of the likelihood of future introductions, *P. clarkii* is known to disperse over long distances on land in order to find suitable habitats (Gherardi et al. 2000). Therefore, it is probable (p = 1) that *P. clarkii* will enter the country via unaided primary pathways such as connected waterways in future. | |
| References: Gherardi, F., Barbaresi, S. & Salvi, G., 2000, 'Spatial and temporal patterns in the movement of *Procambarus clarkii,* an invasive crayfish', *Aquatic Sciences* 62, 179–193. https://doi.org/10.1007/PL00001330. Barkhuizen, L.M., Madzivanzira, T.C. & South, J., 2022, 'Population ecology of a wild population of red swamp crayfish *Procambarus clarkii* (Girard, 1852) in the Free State Province, South Africa and implications for eradication efforts', *BioInvasions Records* 11(1), 181–191. https://doi.org/10.3391/bir.2022. Loureiro, T.G., Anastácio, P.M.S.G., Araujo, P.B., Souty-Grosset, C. & Almerão, M.P., 2015, 'Red swamp crayfish: biology, ecology and invasion–an overview', *Nauplius 23*, 1–19. https://doi.org/10.1590/S0104-64972014002214. Nunes, A.L., Hoffman, C.A., Zengeya, T.A., Measey, G.J. & Weyl, O.L.F, 2017a, 'Red swamp crayfish, *Procambarus clarkii*, found in South Africa 22 years after attempted eradication', *Aquatic Conservation: Marine Freshwater Ecosystem* 27, 1334–1340. https://doi.org/10.1002/aqc.2741. Nunes, A.L., Zengeya, T.A., Measey, G.J. & Weyl, O.L.F., 2017b, 'Freshwater crayfish invasions in South Africa: past, present and potential future', *African Journal of Aquatic Science* 42, 309–323. https://doi.org/10.2989/16085914.2017.1405788. | |

| **LIK2 Likelihood of entry via human aided primary pathways** | |
|---|---|
| Response: Present in the country (p = 1) | Confidence: High |

| |
|---|
| Rationale: *Procambarus clarkii* is present in the country and neighbouring countries (Loureiro et al. 2015; Nunes et al. 2017a). In terms of the likelihood of future introductions, the primary pathway of introduction includes escapees from aquaculture facilities, pet trade industry, and accidental introduction as bait for fish (Faulkes 2015; Nunes et al. 2017b). |
| References:<br>Faulkes, Z., 2015, 'The global trade in crayfish as pets', *Crustacean Research* 44, 75–92. https://doi.org/10.18353/crustacea.44.0_75.<br>Loureiro, T.G., Anastácio, P.M.S.G., Araujo, P.B., Souty-Grosset, C. & Almerão, M.P., 2015, 'Red swamp crayfish: biology, ecology and invasion–an overview', *Nauplius* 23(1), 1–19. https://doi.org/10.1590/S0104-64972014002214.<br>Nunes, A.L., Zengeya, T.A., Measey, G.J. & Weyl, O.L.F., 2017b, 'Freshwater crayfish invasions in South Africa: past, present and potential future', *African Journal of Aquatic Science* 42, 309–323. https://doi.org/10.2989/16085914.2017.1405788. |

| **LIK3 Habitat suitability** | |
|---|---|
| Response: Probable (p = 1) | Confidence: Medium |
| Rationale: *Procambarus clarkii* is tolerant to a wide range of environmental conditions ranging from environments with low salinity, oxygen levels, to those with extreme low and high temperatures (de Moor 2002). *Procambarus clarkii* can occur in a wide range of habitats and can therefore colonise and establish anywhere in South Africa in areas with rivers, lakes, ponds, streams, canals, and seasonally flooded swamps, marshes, and estuaries (de Moor 2002; Souty-Grosset et al. 2016). In Portugal and Italy, *P. clarkii* is known to occur in caves (Souty-Grosset et al. 2016). In South Africa, *P. clarkii* has established in man-made impoundments (de Moor 2002; Madzivanzira et al. 2021). | |
| References:<br>de Moor, I., 2002, 'Potential impacts of alien freshwater crayfish in South Africa', *African Journal of Aquatic Science* 27, 125–139, https://doi.org/10.2989/16085914.2002.9626584.<br>Madzivanzira, T.C., South, J., Wood, L.E., Nunes, A.L. & Weyl, O.L.F., 2021, 'A review of freshwater crayfish introductions in Africa', *Reviews in Fisheries Science & Aquaculture*, 1–21. https://doi.org/10.1080/23308249.2020.1802405<br>Souty-Grosset, C., Anastacio, P.M., Aquiloni, L., Banha, F., Choquer, J., Chucholl, C., et al. 2016, 'The red swamp crayfish *Procambarus clarkii* in Europe: Impacts on aquatic ecosystems and human well-being', *Limnologica* 58, 78–93, https://doi.org/10.1016/j.limno.2016.03.003. | |

| **LIK4 Climate suitability** | |
|---|---|
| Response: Probable (p = 1) | Confidence: High |
| Rationale: *Procambarus clarkii* occurs in environments with optimal conditions between 20–25˚C, but has been seen in South Africa to withstand water temperatures exceeding 25 ˚C but not 35 ˚C (de Moor 2002). Sub-optimal temperatures (<12 ˚C) inhibit growth but *P. clarkii* can survive lower temperatures (>0˚ C) by hibernating in burrows (Peruzza et al. 2015). Several catchment areas in South Africa were predicted to be climatically suitable for *P. clarkii*, include the Greater Berg, Bree, Gourits, Kromme, Swartkops, Bushmans, Keiskamma, Great Kei, Mzimvubu, uMngeni and Phongolo Rivers (Nunes et al. 2017b). | |
| References:<br>de Moor, I., 2002, 'Potential impacts of alien freshwater crayfish in South Africa', *African Journal of Aquatic Science* 27, 125–139. https://doi.org/10.2989/16085914.2002.9626584. . | |

Nunes, A.L., Zengeya, T.A., Measey, G.J. & Weyl, O.L.F., 2017b, 'Freshwater crayfish invasions in South Africa: past, present and potential future', *African Journal of Aquatic Science* 42, 309–323. https://doi.org/10.2989/16085914.2017.1405788.

Peruzza, L., Piazza, F., Manfrin, C., Bonzi, L.C., Battistella, S. & Giulianini, P.G., 2015, 'Reproductive plasticity of a *Procambarus clarkii* population living 10°C below its thermal optimum', *Aquatic Invasions* 10(2), 199–208. https://doi.org/10.3391/ai.2015.10.2.08.

| **LIK5 Unaided secondary (dispersal) pathways** | |
|---|---|
| Response: Probable (p = 1) | Confidence: Medium |
| Rationale: *Procambarus clarkii* can travel long distances overland to find water resources, therefore it is probable (p = 1) that *P. clarkii* would be able to move into new areas. *Procambarus clarkii* has stationary and nomadic phases (Gherardi 2006). During the nomadic phase, breeding males have been reported to travel up to 17 km in four days, and can survive up to 10 hours outside water (Gherardi 2006; Souty-Grosset et al. 2016). Juveniles can also be translocated to new areas by water birds (Souty-Grosset et al. 2016). | |
| References: Gherardi, F., 2006, 'Crayfish invading Europe: The case study of *Procambarus clarkii*', *Hydrobiologia* 595, 295–301. https://doi.org/10.1080/10236240600869702. Souty-Grosset, C., Anastacio, P.M., Aquiloni, L., Banha, F., Choquer, J., Chucholl, C. & Tricarico, E., 2016, 'The red swamp crayfish *Procambarus clarkii* in Europe: Impacts on aquatic ecosystems and human well-being', *Limnologica* 58, 78–93. https://doi.org/10.1016/j.limno.2016.03.003. | |

| **LIK6 Human aided secondary (dispersal) pathways** | |
|---|---|
| Response: Probable (p = 1) | Confidence: Medium |
| Rationale: *Procambarus clarkii* could be introduced into new areas through aquaculture and the pet trade (Faulkes 2015; Holdich et al. 2009; Madzivanzira et al. 2021). *Procambarus clarkii* is also likely to be translocated intentionally as bait for fishing (Nunes et al. 2017a). In addition, areas with known naturalised populations can act as a source for spreading into other regions in South Africa. *Procambarus clarkii* is not currently regulated under the NEM:BA Alien and Invasive Species regulations and is available for purchase online (Nunes et al. 2017a; Madzivanzira et al. 2021). This could promote farming operations for *P. clarkii* resulting in further dispersal of the species (Nunes et al. 2017a; Madzivanzira et al. 2021). | |
| References: Faulkes, Z., 2015, 'The global trade in crayfish as pets', *Crustacean Research* 44, 75–92. https://doi.org/10.18353/crustacea.44.0_75 Holdich, D.M., Reynolds, J.D. & Sibley, P.J., 2009, 'A review of the ever increasing threat to European crayfish from non-indigenous crayfish species', *Knowledge and Management of Aquatic Ecosystems* 11, 394–395. https://doi.org/10.1051/kmae/2009025. Madzivanzira, T.C., South, J., Wood, L.E., Nunes, A.L. & Weyl, O.L.F., 2021, 'A review of freshwater crayfish introductions in Africa', *Reviews in Fisheries Science & Aquaculture*,1–21. https://doi.org/10.1080/23308249.2020.1802405. Nunes, A.L., Hoffman, C.A., Zengeya, T.A., Measey, G.J. & Weyl, O.L.F, 2017a, 'Red swamp crayfish, *Procambarus clarkii*, found in South Africa 22 years after attempted eradication', *Aquatic Conservation: Marine & Freshwater Ecosystems* 27, 1334–1340. https://doi.org/10.1002/aqc.2741. | |

**3. Consequences**

| **CON1 Environmental impact** |
|---|

| **CON1a: Competition** | |
|---|---|
| Response: Major (MR) | Confidence: High |
| Rationale: *Procambarus clarkii* is very aggressive and is known to out-compete native species for shelter and spawning sites, which often leads to reproductive interference (Cruz et al. 2006). In introduced areas, some amphibian species (e.g., *Bufo bufo*, *Bufo calamita*, *Rana* spp., *Taricha torosa* and *Triturus vulgaris*) have been excluded or displaced from their natural habitats, resulting in local extinctions through either larval predation or amphibians spawning in areas that do not offer sufficient protection to avoid interaction with *P. clarkii,* resulting in low recruitment (Cruz et al. 2006; Lodge et al. 2012). In addition, as a result of direct competition, there has been a decrease in the distributional ranges and abundance of native crayfish populations (*Astacus astacus*, *Austropotamobius pallipes* and *Austropotamobius torrentium*) in some areas in Europe and Japan (Cruz et al. 2006; Lodge et al. 2012). Furthermore, direct competition for food has caused dietary niche constriction and declines in populations of native crabs (*Potamonautes* sp.) in some areas invaded by *P. clarkii* (Gherardi & Daniels 2004; Jackson et al. 2016). | |
| References:<br>Cruz, M.J., Rebelo, R. & Crespo, E.G., 2006, 'Effects of an introduced crayfish, *Procambarus clarkii*, on the distribution of south-western Iberian amphibians in their breeding habitats', *Ecography* 29. 329–338. https://doi.org/10.1111/j.2006.0906-7590.04333.x.<br>Gherardi, F. & Daniels, W.H., 2004, 'Agonism and shelter competition between invasive and indigenous crayfish species' *Canadian Journal of Zoology* 82, 1923–1932. https://doi.org/10.1139/z04-185<br>Jackson, M.C., Grey, J., Miller, K., Britton, J.R. & Donohue, I., 2016, 'Dietary niche constriction when invaders meet natives: Evidence from freshwater decapods', *Journal of Animal Ecology* 85, 1098–1107https://doi.org/10.1051/kmae:2005031.<br>Lodge, D.M., Deines, A., Gherardi, F., Yeo, D.C., Arcella, T., Baldridge, A.K., et al., 2012, 'Global introductions of crayfishes: Evaluating the impact of species invasions on ecosystem services', *Annual Review of Ecology Evolution Systematics* 43, 449–472. https://doi.org/10.1146/annurev-ecolsys-111511-103919. | |
| **CON1b: Predation** | |
| Response: Major (MR) | Confidence: High |
| Rationale: In the alien range, there is evidence that predation by *P. clarkii* has resulted in the loss of populations of at least one native species (Lodge et al. 2012; Souty-Grosset et al. 2016). *Procambarus clarkii* has been implicated in population declines of several species of fish and amphibians by reducing their breeding success through predation on eggs and larvae respectively (Gamradt et al. 1996; Cruz et al. 2006; Mueller et al. 2006; Ficetola et al. 2011). | |
| References:<br>Cruz, M.J., Rebelo, R. & Crespo, E.G., 2006, 'Effects of an introduced crayfish, *Procambarus clarkii*, on the distribution of south-western Iberian amphibians in their breeding habitats', *Ecography* 29, 329–338, https://doi.org/10.1111/j.2006.0906–7590.04333.x.<br>Ficetola, G.F., Siesa, M.E., Manenti, R., Bottoni, L., De Bernardi, F. & Padoa-Schioppa, E., 2011, 'Early assessment of the impact of alien species: Differential consequences of an invasive crayfish on adult and larval amphibians', *Diversity and Distributions* 17, 1141–1151. https://doi.org/10.1111/j.1472-4642.2011.00797.x<br>Gamradt, S.C. & Kats, L.B., 1996, 'Effect of introduced crayfish and mosquitofish on California newts', *Conservation Biology* 10(4), 1155–1162. https://doi.org/10.1046/j.1523-1739.1996.10041155.x<br>Lodge, D.M., Deines, A., Gherardi, F., Yeo, D.C., Arcella, T., Baldridge, A.K., et al., 2012, 'Global introductions of crayfishes: Evaluating the impact of species invasions on ecosystem services', *Annual Review of Ecology Evolution Systematics* 43, 449–472, https://doi.org/10.1146/annurev-ecolsys-111511-103919. | |

Mueller, G.A., Carpenter, J. & Thornbrugh, D., 2006, ‘Bullfrog tadpole (*Rana catesbeiana*) and red swamp crayfish (*Procambarus clarkii*) predation on early life stages of endangered razorback sucker (*Xyrauchen texanus*)’, *The Southwestern Naturalist* 51(2), 258–261. https://doi.org/10.1894/0038-4909(2006)51[258:BTRCAR]2.0.CO;2

Souty-Grosset, C., Anastacio, P.M., Aquiloni, L., Banha, F., Choquer, J., Chucholl, C., et al., 2016, ‘The red swamp crayfish *Procambarus clarkii* in Europe: Impacts on aquatic ecosystems and human well-being’, *Limnologica* 58, 78–93. https://doi.org/10.1016/j.limno.2016.03.003.

**CON1c: Hybridisation**

| Response: Data Deficient (DD) | Confidence: Low |
|---|---|

Rationale: Hybridisation is unlikely in South Africa because there are no native freshwater crayfish species (de Moor 2002; Nunes et al. 2017b). The other alien crayfish species that are currently in South Africa are both in the *Cherax* genus native to Australia of which one is confined to aquaculture facilities (de Moor 2002; Nunes et al. 2017b).

References:

de Moor, I., 2002, ‘Potential impacts of alien freshwater crayfish in South Africa’, *African Journal of Aquatic Science* 27, 125–139. , https://doi.org/10.2989/16085914.2002.9626584.

Nunes, A.L., Zengeya, T.A., Measey, G.J. & Weyl, O.L.F., 2017b, ‘Freshwater crayfish invasions in South Africa: past, present and potential future’, *African Journal of Aquatic Science* 42, 309–323. https://doi.org/10.2989/16085914.2017.1405788.

**CON1d: Transmission of disease**

| Response: Major (MR) | Confidence: High |
|---|---|

Rationale: *Procambarus clarkii* can harbour many pathogens, parasites, and diseases that can be transmitted to other congeneric species (Longshaw 2011; Mastitsky et al. 2010). There is evidence that transmission of diseases and parasites by *P. clarkii* to native species has caused the loss of populations of at least one native species, leading to changes in community composition (Lodge et al. 2012). For example, *P. clarkii* is a vector of crayfish plague, a disease caused by the parasitic oomycete *Aphanomyces astaci* (Longshaw 2011; Souty-Grosset et al. 2016). In Europe, transmission of crayfish plague by *P. clarkii* to native crayfish species has been linked to a decline in populations of several native crayfish species such as *Astacus astacus*, *Austropotamobius pallipes* and *A. torrentium* (Holdich et al. 2009; Longshaw 2011). *Procambarus clarkii* can also harbour white spot syndrome disease (a viral infection of crustaceans), and fungal pathogens such as *Batrachochytrium dendrobatids* that causes chytridiomycosis (a lethal skin infection of amphibians) (Longshaw 2011; McMahon et al. 2013).

References:

Mastitsky, S.E., Karatayev, A.Y., Burlakova, L.E. and Molloy, D.P., 2010. ‘BIODIVERSITY RESEARCH: Parasites of exotic species in invaded areas: does lower diversity mean lower epizootic impact?’, Diversity *and Distributions*, *16,* 798-803. https://doi.org/10.1111/j.1472-4642.2010.00693.x

Holdich, D.M., Reynolds, J.D. & Sibley, P.J., 2009, ‘A review of the ever increasing threat to European crayfish from non-indigenous crayfish species’, *Knowledge and Management of Aquatic Ecosystems* 394–395, 11. https://doi.org/10.1051/kmae/2009025.

Lodge, D.M., Deines, A., Gherardi, F., Yeo, D.C., Arcella, T., Baldridge, A.K., et al., 2012, ‘Global introductions of crayfishes: Evaluating the impact of species invasions on ecosystem services’, *Annual Review of Ecology Evolution Systematics* 43, 449–472. https://doi.org/10.1146/annurev-ecolsys-111511-103919

Longshaw, M. 2011, ‘Diseases of crayfish: A review’, *Journal of Invertebrate Pathology* 106, 54–70. https://doi.org/10.1016/j.jip.2010.09.013

| |
|---|
| Karatayev, L.E. Burlakova, & Molloy, D.P., 2010, 'Parasites of exotic species in invaded areas: does lower diversity mean lower epizootic impact?' *Diversity and Distributions* 16, 798–803. https://doi.org/10.1007/s00442-008-1180-1<br>McMahon, T.A., Brannelly, L.A., Chatfield, M.W., Johnson, P.T., Joseph, M.B., McKenzie, V.J., et al., 2013, 'Chytrid fungus *Batrachochytrium dendrobatidis* has non-amphibian hosts and releases chemicals that cause pathology in the absence of infection', *Proceedings of the National Academy of Sciences of the United States of America* 110, 210–215. https://doi.org/10.1073/pnas.1200592110.<br>Souty-Grosset, C., Anastacio, P.M., Aquiloni, L., Banha, F., Choquer, J., Chucholl, C., et al., 2016, 'The red swamp crayfish *Procambarus clarkii* in Europe: Impacts on aquatic ecosystems and human well-being', *Limnologica* 58, 78–93. https://doi.org/10.1016/j.limno.2016.03.003. |

| **CON1e: Parasitism** | |
|---|---|
| Response: Data Deficient (DD) | Confidence: High |
| Rationale: *Procambarus clarkii* is not a parasitic taxon. | |
| References: | |

| **CON1f: Poisoning/toxicity** | |
|---|---|
| Response: Minimal Concern (MC) | Confidence: Low |
| Rationale: *Procambarus clarkii* can bio-accumulate toxins and metals from the environment (Gherardi et al. 2011; Souty-Grosset et al. 2016). These pollutants are often harmful and are transferred to higher food web levels through the consumption of affected crayfish by predators such as otters, birds, and fish (Gherardi et al. 2011; Lodge et al. 2012). | |
| References:<br>Gherardi, F., Britton, J.R., Mavuti, K.M., Pacini, N., Grey, J., Tricarico, E., et al., 2011, 'A review of allodiversity in Lake Naivasha, Kenya: Developing conservation actions to protect East African lakes from the negative impacts of alien species', *Biological Conservation* 144, 2585–2596. https://doi.org/10.1016/j.biocon.2011.07.020.<br>Lodge, D.M., Deines, A., Gherardi, F., Yeo, D.C., Arcella, T., Baldridge, A.K., et al., 'Global introductions of crayfishes: Evaluating the impact of species invasions on ecosystem services', *Annual Review of Ecology Evolution Systematics* 43, 449–472. https://doi.org/10.1146/annurev-ecolsys-111511-103919.<br>Souty-Grosset, C., Anastacio, P.M., Aquiloni, L., Banha, F., Choquer, J., Chucholl, C., et al., 2016, 'The red swamp crayfish *Procambarus clarkii* in Europe: Impacts on aquatic ecosystems and human well-being', *Limnologica* 58, 78–93. https://doi.org/10.1016/j.limno.2016.03.003. | |

| **CON1g: Bio-fouling or other direct physical disturbance** | |
|---|---|
| Response: Data Deficient (DD) | Confidence: Low |
| Rationale: | |
| References: | |

| **CON1h: Grazing/herbivory/browsing** | |
|---|---|
| Response: Moderate (MO) | Confidence: Medium |
| Rationale: Grazing by *P. clarkii* leads to habitat loss and modification through the removal of macrophytes. Habitat loss can lead to a decline in populations of species that utilise the macrophyte stands as a food source, nesting sites, and as refugia from predation (Rosenthal et al. 2005). *Procambarus clarkii* leads to changes to | |

| |
|---|
| community composition through trophic cascades (Gherardi & Aquistapace 2007; Souty-Grosset et al. 2016). For example, in Lake Chozas (northwest Spain), grazing by *P. clarkii* caused a reduction in macrophyte communities and this impact cascaded up the food chain with declines in invertebrates, amphibians, and waterfowl (Rodriguez et al. 2003). Similar results were also observed in Lake Naivasha in Kenya (Smart et al. 2002). |
| References:<br>Gherardi, F. & Acquistapace P., 2007, 'Invasive crayfish in Europe: The impact of *Procambarus clarkii* on the littoral community of a Mediterranean lake', *Freshwater Biology* 52, 1249–59. https://doi.org/10.1111/j.1365-2427.2007.01760.x.<br>Rodríguez, C.F., Bécares, E. & Fernández-Aláez, M., 2003, 'Shift from clear to turbid phase in Lake Chozas (NW Spain) due to the introduction of American red swamp crayfish (*Procambarus clarkii*)', *Hydrobiologia* 506, 421–26, https://doi.org/10.1023/B:HYDR.0000008626.07042.87.<br>Rosenthal, S.K., Lodge, D.M., Mavuti, K.M., Muohi, W., Ochieng, P., Mungai, B.N. & Mkoji, G.M., 2005, 'Comparing macrophyte herbivory by introduced Louisiana crayfish (*Procambarus clarkii*) (Crustacea: Cambaridea) and native Dytiscid beetles (*Cybister tripunctatus*) in Kenya, *African Journal of Aquatic Science*, 30, 157–162, https://doi.org/10.2989/16085910509503850.<br>Smart, A.C., Harper, D.M., Malaisse, F., Schmitz, S., Coley, S. & de Beauregard, A-C.G., 2002, 'Feeding of the exotic Louisiana red swamp crayfish, *Procambarus clarkii* (Crustacea, Decapoda), in an African tropical lake: Lake Naivasha, Kenya', *Hydrobiologia* 488, 129–142. https://doi.org/10.1023/A:1023326530914.<br>Souty-Grosset, C., Anastacio, P.M., Aquiloni, L., Banha, F., Choquer, J., Chucholl, C., et al., 2016, 'The red swamp crayfish *Procambarus clarkii* in Europe: Impacts on aquatic ecosystems and human well-being', *Limnologica* 58, 78–93. , https://doi.org/10.1016/j.limno.2016.03.003. |

| **CON1i: Chemical, physical or structural impact on ecosystem** | |
|---|---|
| Response: MN (Minor) | Confidence: Medium |
| Rationale:<br><br>**Water quality**: *Procambarus clarkii* is often considered an ecosystem engineer due to its ability to change ecosystems through its burrowing activity. For example, its burrowing activity can cause a decrease in the water quality through bioturbation leading to increased turbidity and influx release of nutrients from sediment (Rodriguez et al 2003; Yamamoto 2010). This often leads to algal blooms. The impaired water quality also affects the quality of habitat for other aquatic fauna. For example, increased turbidity can impede foraging and respiratory processes of fish (Souty-Grosset et al. 2016).<br><br>**Erosion**: Burrowing activities can cause structural damage to river banks and increase bank erosion (Angeler et al. 2001). | |
| References:<br>Angeler, D.G., Sánchez-Carrillo, S., García, G.& Alvarez-Cobelas, M., 2001, 'The influence of *Procambarus clarkii* (Cambaridae, Decapoda) on water quality and sediment characteristics in a Spanish floodplain wetland', *Hydrobiologia* 464, 89–98. https://doi.org/10.1023/A:1013950129616.<br>Rodríguez, C.F., Bécares, E. & Fernández-Aláez, M., 2003, 'Shift from clear to turbid phase in Lake Chozas (NW Spain) due to the introduction of American red swamp crayfish (*Procambarus clarkii*)', *Hydrobiologia* 506, 421–26. https://doi.org/10.1023/B:HYDR.0000008626.07042.87.<br>Souty-Grosset, C., Anastacio, P.M., Aquiloni, L., Banha, F., Choquer, J., Chucholl, C., et al., 2016, 'The red swamp crayfish *Procambarus clarkii* in Europe: Impacts on aquatic ecosystems and human well-being', *Limnologica* 58, 78–93. https://doi.org/10.1016/j.limno.2016.03.003. | |

Yamamoto, Y., 2010, 'Contribution of bioturbation by the red swamp crayfish *Procambarus clarkii* to the recruitment of bloom-forming cyanobacteria from sediment', *Journal of limnology* 69, 102–111. https://doi.org/10.4081/jlimnol.2010.102.

**CON1k: Indirect impacts through interactions with other species**

| Response: Minimal Concern (MC) | Confidence: Medium |
|---|---|

Rationale: *Procambarus clarkii* often occurs in sympatry with other species in its alien range that actively feeds on it for example birds that can lead to dispersal into new areas (Lovas-Kiss et al. 2018; Southy-Grosset et al. 2016).

References:

Lovas-Kiss, A., Sanchez, M.I., Molnar, A.V., Valls, L., Armengol, X., Mesquita-Joanes, F., et al., 2018, 'Crayfish invasion facilitates dispersal of plants and invertebrates by gulls', *Freshwater Biology* 63(4), 392–404. https://doi.org/10.1111/fwb.13080.

Souty-Grosset, C., Anastacio, P.M., Aquiloni, L., Banha, F., Choquer, J., Chucholl, C., et al., 2016, 'The red swamp crayfish *Procambarus clarkii* in Europe: Impacts on aquatic ecosystems and human well-being', *Limnologica* 58, 78–93. https://doi.org/10.1016/j.limno.2016.03.003.

**CON1: Maximum environmental impact**

| Response: Major (MR) | Confidence: High |
|---|---|

Rationale Direct predation and competition for food, shelter, and spawning sites have led to the loss of populations and a decrease in the abundance of native amphibians and crayfish species (Cruz et al. 2006; Gherardi & Acquistapace 2007; Jackson et al. 2016). Impacts have been recorded in the Iberian Peninsula, Sweden, Italy (Gherardi & Acquistapace 2007), Japan, and U.S.A. (California) (Holdich et al. 2009; Lodge et al. 2012). Alien crayfish expose native fauna to diseases and parasites that could affect their overall health (Edgerton et al. 2002; Longshaw 2011). For example, *P. clarkii* has been implicated in the transmission of crayfish plague that caused devastating impacts in Europe resulting in the collapse of native freshwater crayfish communities (Holdich & Reeve 1991; Souty-Grosset et al. 2016).

References:

Cruz, M.J., Rebelo, R. & Crespo, E.G., 2006, 'Effects of an introduced crayfish, *Procambarus clarkii*, on the distribution of south-western Iberian amphibians in their breeding habitats', *Ecography* 29, 329–338. https://doi.org/10.1111/j.2006.0906-7590.04333.x.

Edgerton, B.F., Evans, L.H., Stephens, F.J. & Overstreet, R.M., 2002, 'Synopsis of freshwater crayfish diseases and commensal organisms', *Aquaculture* 206, 57–135. https://doi.org/10.1016/S0044-8486(01)00865-1.

Gherardi, F. & Acquistapace P., 2007, 'Invasive crayfish in Europe: The impact of *Procambarus clarkii* on the littoral community of a Mediterranean lake', *Freshwater Biology* 52, 1249–59. https://doi.org/10.1111/j.1365-2427.2007.01760.x.

Holdich, D.M. & Reeve, I.D., 1991, 'Distribution of freshwater crayfish in the British Isles, with particular reference to crayfish plague, alien introductions and water quality', *Aquatic Conservation: Marine & Freshwater Ecosystems* 1, 139–158. https://doi.org/10.1002/aqc.3270010204.

Holdich, D.M., Reynolds, J.D. & Sibley, P.J., 2009, 'A review of the ever increasing threat to European crayfish from non-indigenous crayfish species', *Knowledge and Management of Aquatic Ecosystems* 11, 394–395. https://doi.org/10.1051/kmae/2009025.

Jackson, M.C., Grey, J., Miller, K., Britton, J.R. & Donohue, I., 2016, 'Dietary niche constriction when invaders meet natives: Evidence from freshwater decapods', *Journal of Animal Ecology* 85, 1098–1107. https://doi.org/10.1051/kmae:2005031.

Lodge, D.M., Deines, A., Gherardi, F., Yeo, D.C., Arcella, T., Baldridge, A.K., et al., 2012, 'Global introductions of crayfishes: Evaluating the impact of species invasions on ecosystem services', *Annual*

*Review of Ecology Evolution Systematics* 43, 449–472. https://doi.org/10.1146/annurev-ecolsys-111511-103919..

Longshaw, M. 2011, 'Diseases of crayfish: A review', *Journal of Invertebrate Pathology* 106, 54–70. https://doi.org/10.1016/j.jip.2010.09.013.

Souty-Grosset, C., Anastacio, P.M., Aquiloni, L., Banha, F., Choquer, J., Chucholl, C., et al., 2016, 'The red swamp crayfish *Procambarus clarkii* in Europe: Impacts on aquatic ecosystems and human well-being', *Limnologica* 58, 78–93. https://doi.org/10.1016/j.limno.2016.03.003.

| **CON2 Socio-economic impact** | |
|---|---|
| **CON2a: Safety** | |
| Response: Data Deficient (DD) | Confidence: Low |
| Rationale: | |
| References: | |
| **CON2b: Material and immaterial assets** | |
| Response: Moderate (MO) | Confidence: Medium |

Rationale: *Procambarus clarkii* often inhabits agricultural fields and their burrowing activities can cause damage to infrastructure such as irrigation canals and dam walls (Arce & Diéguez-Uribeondo 2015; Lodge et al. 2012). Burrowing activities can also alter soil hydrology leading to water loss. Grazing causes crop damage and reduces yield (Anastácio et al. 2005). In Europe and North Africa, *P. clarkii* affected rice production through field water loss, damage to rice field banks and ditches, direct consumption of rice seed and plants, and clogging of pipes (Abdel-Kader 2016; Sara & El Moutaouaki 2019; Souty-Grosset et al. 2016). *Procambarus clarkii* affects the fishing industry by damaging gill nets and spoiling the fish caught in the nets (Gherardi et al. 2011).

References:

Abdel-Kader, S.M., 2016, 'Some ecological and toxicological studies on crayfish, *Procambarus clarkii* at Sharkia governorate,' *Journal of Basic and Applied Zoology* 74, 62–67. https://doi.org/10.1016/j.jobaz.2016.08.001.

Anastácio, P.M., Correia, A.M. & Menino, J.P., 2005, 'Processes and patterns of plant destruction by crayfish: effects of crayfish size and developmental stages of rice', *Archiv für Hydrobiologie* 162, 37–51, https://doi.org/10.1127/0003-9136/2005/0162-0037.

Arce, J.A. & Diéguez-Uribeondo, J., 2015, 'Structural damage caused by the invasive crayfish *Procambarus clarkii* (Girard, 1852) in rice fields of the Iberian Peninsula: A study case', *Fundamental and Applied Limnology* 186, 259–269. https://doi.org/10.1127/fal/2015/0715.

Gherardi, F., Britton, J.R., Mavuti, K.M., Pacini, N., Grey, J., Tricarico, E., et al., 2011, 'A review of allodiversity in Lake Naivasha, Kenya: Developing conservation actions to protect East African lakes from the negative impacts of alien species', *Biological* Conservation 144, 2585–2596. , https://doi.org/10.1016/j.biocon.2011.07.020.

Lodge, D.M., Deines, A., Gherardi, F., Yeo, D.C., Arcella, T., Baldridge, A.K., et al., 2012, 'Global introductions of crayfishes: Evaluating the impact of species invasions on ecosystem services', *Annual Review of Ecology Evolution Systematics* 43, 449–472. https://doi.org/10.1146/annurev-ecolsys-111511-103919.

Sara, S. & El Moutaouakil, M.E., 2019, 'Study on the Spread of *Procambarus clarkii* at Gharb (Morocco) and its impact on rice growing', *Journal of Agriculture, Science and Technology*, 9, 81–92. https://doi.org/10.17265/2161-6256/2019.02.002.

Souty-Grosset, C., Anastacio, P.M., Aquiloni, L., Banha, F., Choquer, J., Chucholl, C., et al., 2016, 'The red swamp crayfish *Procambarus clarkii* in Europe: Impacts on aquatic ecosystems and human well-being', *Limnologica* 58, 78–93. https://doi.org/10.1016/j.limno.2016.03.003.

**CON2c: Health**

| Response: Minor (MN) | Confidence: Medium |
|---|---|

Rationale: *Procambarus clarkii* can bio-accumulate toxins and metals from the environment (Gherardi et al. 2011; Souty-Grosset et al. 2016). These pollutants are often harmful and can be transferred to humans via consumption (Gherardi et al. 2011; Lodge et al. 2012).

*Procambarus clarkii* serves as a vector for several parasites and diseases some of which are zoonotic, for example the parasitic fluke flatworms (*Paragonimus* spp.) that causes lung fluke disease in humans, tularemia–causing bacterium *Francisella tularensis*, rat lungworm *Angiostrongylus cantonensis* that causes meningitis, and the nematode *Gnathostoma spinigerum* that causes human gnathostomiasis (de Moor 2002; Putra et al. 2017; Souty-Grosset et al. 2016).

References:

de Moor, I., 2002, 'Potential impacts of alien freshwater crayfish in South Africa', *African Journal of Aquatic Science* 27, 125–139. https://doi.org/10.2989/16085914.2002.9626584.

Gherardi, F., Britton, J.R., Mavuti, K.M., Pacini, N., Grey, J., Tricarico, E. & Harper, D.M., 2011, 'A review of allodiversity in Lake Naivasha, Kenya: Developing conservation actions to protect East African lakes from the negative impacts of alien species', *Biological* Conservation 144, 2585–2596. https://doi.org/10.1016/j.biocon.2011.07.020.

Putra, M.D, Bláha, M., Wardiatno, Y., Krisanti, M., Bystřický, P.K, Kouba, A., et al., 2017, '*Procambarus clarkii* (Girard, 1852) and crayfish plague as new threats for biodiversity in Indonesia', *Aquatic Conservation: Marine and Freshwater Ecosystems* 28, 1434–1440. https://doi.org/10.1002/aqc.2970.

Souty-Grosset, C., Anastacio, P.M., Aquiloni, L., Banha, F., Choquer, J., Chucholl, C. & Tricarico, E., 2016, 'The red swamp crayfish *Procambarus clarkii* in Europe: Impacts on aquatic ecosystems and human well-being', *Limnologica* 58, 78–93. https://doi.org/10.1016/j.limno.2016.03.003.

**CON2d: Social, spiritual and cultural relations**

| Response: Data Deficient (DD) | Confidence: Medium |
|---|---|

Rationale:

References:

**CON2 Maximum socio-economic impact**

| Response: Moderate (MO) | Confidence: Medium |
|---|---|

Rationale: In the alien range, *P. clarkii* has caused harmful impacts in agricultural fields (Anastácio et al. 2005) and has disrupted some recreational activities leading to economic loss (Gherardi et al. 2011). Additionally, *P. clarkii* is a host of various parasites and diseases (*Paragonimus* spp., *Francisella tularensis*, *Angiostrongylus cantonensis*, and *Gnathostoma spinigerum*) (de Moor 2002; Putra et al. 2017; Souty-Grosset et al. 2016). *Procambarus clarkii* also bio-accumulates toxins and metals that are harmful to humans if consumed (Gherardi et al. 2011; Lodge et al. 2012; Souty-Grosset et al. 2016).

References:

de Moor, I., 2002, 'Potential impacts of alien freshwater crayfish in South Africa', *African Journal of Aquatic Science* 27, 125–139. https://doi.org/10.2989/16085914.2002.9626584.
Anastácio, P.M., Correia, A.M. & Menino, J.P., 2005, 'Processes and patterns of plant destruction by crayfish: effects of crayfish size and developmental stages of rice', *Archiv für Hydrobiologie* 162, 37–51 https://doi.org/10.1127/0003-9136/2005/0162-0037.
Gherardi, F., Britton, J.R., Mavuti, K.M., Pacini, N., Grey, J., Tricarico, E., et al., 2011, 'A review of allodiversity in Lake Naivasha, Kenya: Developing conservation actions to protect East African lakes from the negative impacts of alien species', *Biological* Conservation 144, 2585–2596. https://doi.org/10.1016/j.biocon.2011.07.020.
Lodge, D.M., Deines, A., Gherardi, F., Yeo, D.C., Arcella, T., Baldridge, A.K., et al., 2012, 'Global introductions of crayfishes: Evaluating the impact of species invasions on ecosystem services', *Annual Review of Ecology Evolution Systematics* 43, 449–472. https://doi.org/10.1146/annurev-ecolsys-111511-103919.
Putra, M.D, Bláha, M., Wardiatno, Y., Krisanti, M., Bystřický, P.K, Kouba, A., et al., 2017, '*Procambarus clarkii* (Girard, 1852) and crayfish plague as new threats for biodiversity in Indonesia', *Aquatic Conservation: Marine and Freshwater Ecosystems* 28, 1434–1440. https://doi.org/10.1002/aqc.2970.
Souty-Grosset, C., Anastacio, P.M., Aquiloni, L., Banha, F., Choquer, J., Chucholl, C. & Tricarico, E., 2016, 'The red swamp crayfish *Procambarus clarkii* in Europe: Impacts on aquatic ecosystems and human well-being', *Limnologica* 58, 78–93. https://doi.org/10.1016/j.limno.2016.03.003.

| **CON3 Closely related species' environmental impact** | |
|---|---|
| Response: Not scored as CON1 scored | Confidence: |
| Rationale: There is information to score CON1, so CON3 does not need to be scored as per the framework. | |
| References: | |

| **CON4 Closely related species' socio-economic impact** | |
|---|---|
| Response: Not scored as CON2 scored | Confidence: |
| Rationale: There is information to score CON2, so CON4 does not need to be scored as per the framework. | |
| References: | |

| **CON5 Potential impact** | |
|---|---|
| Response: Major (MR) | Confidence: High |
| Rationale: *Procambarus clarkii* can tolerate a wide range of environmental conditions (Nunes et al. 2017b), and is an aggressive competitor that can outcompete and displace native species in invaded ecosystems (Gherardi et al. 2011; Jackson et al. 2016). *Procambarus clarkii* can also cause habitat alteration through intensive grazing of macrophytes and burrowing activities can destabilise river banks and water infrastructure such as dams and canals (Souty-Grosset et al. 2016). Habitat loss can lead to a decline in populations of species that utilise the macrophyte stands as a food source, nesting sites, and as refugia from predation (Rosenthal et al 2005). Grazing and browsing activities of *P. clarkii* can causes economic losses in agricultural fields where these activities can damage infrastructure (e.g., to irrigation canals and dam walls) (Lodge et al. 2012; Arce & Diéguez-Uribeondo 2015) and can reduce yields by damaging crops (Anastácio et al. 2005; Souty-Grosset et al. 2016). *Procambarus clarkii* can also have economic repercussions on the fishing industry by damaging gill nets and spoiling the fish caught in the nets (Gherardi et al. 2011). | |

*Procambarus clarkii* has resulted in the decline of several species of fish and amphibians through predation on eggs and larval amphibians (Cruz et al. 2006; Francesco et al. 2011; Lodge et al. 2012; Souty-Grosset et al. 2016). Indirect effects of predation can also cause trophic cascades that can lead to changes in ecosystem structure and function. For example, predation on invertebrates can release algae from grazing pressure and lead to changes in the abundance and dominance of species in algal communities (Gherardi & Barbaresi 2008).

*Procambarus clarkii* is host of various pathogens, parasites, and diseases (i.e., *Aphanomyces astaci*; *Batrachochytrium dendrobatids* and white spot syndrome disease) that can be transmitted to congeneric species (Holdich et al. 2009; Longshaw 2011; Lodge et al. 2012; McMahon et al. 2013; Souty-Grosset et al. 2016). The transmission of pathogens, parasites and diseases to other crayfish taxa can result in population extinctions leading to changes in community composition (Lodge et al. 2012). Furthermore, *P. clarkii* is a host and vector of several parasites and diseases (*Paragonimus* spp. can cause lung fluke disease in humans, *Francisella tularensis*, and *Angiostrongylus cantonensis* (rat lungworm) can cause meningitis, and *Gnathostoma spinigerum* can causes human gnathostomiasis) (de Moor 2002; Souty-Grosset et al. 2016; Putra et al. 2017). Additionally, *P. clarkii* can accumulate toxins and metals from the environment (Gherardi et al. 2011; Souty-Grosset et al. 2016) which can be transferred to other predators and humans who consume crayfish (Gherardi et al. 2011; Lodge et al. 2012). Recreational activities such as angling are also affected because *P. clarkii* damage nets and prey on popular fish species resulting in some economic loss.

References:

Anastácio, P.M., Correia, A.M. & Menino, J.P., 2005, 'Processes and patterns of plant destruction by crayfish: effects of crayfish size and developmental stages of rice', *Archiv für Hydrobiologie* 162, 37–51, https://doi.org/10.1127/0003-9136/2005/0162-0037.

Arce, J.A. & Diéguez-Uribeondo, J., 2015, 'Structural damage caused by the invasive crayfish *Procambarus clarkii* (Girard, 1852) in rice fields of the Iberian Peninsula: A study case', *Fundamental and Applied Limnology* 186, 259–269, https://doi.org/10.1127/fal/2015/0715.

Cruz, M.J., Rebelo, R. & Crespo, E.G., 2006, 'Effects of an introduced crayfish, *Procambarus clarkii*, on the distribution of south-western Iberian amphibians in their breeding habitats', *Ecography* 29, 329–338. https://doi.org/10.1111/j.2006.0906-7590.04333.x

de Moor, I., 2002, 'Potential impacts of alien freshwater crayfish in South Africa', *African Journal of Aquatic Science* 27, 125–139. https://doi.org/10.2989/16085914.2002.9626584.

Ficetola, G.F., Siesa, M.E., Manenti, R., Bottoni, L., De Bernardi, F. & Padoa-Schioppa, E., 2011, 'Early assessment of the impact of alien species: Differential consequences of an invasive crayfish on adult and larval amphibians', *Diversity and Distributions* 17, 1141–1151. https://doi.org/10.1111/j.1472-4642.2011.00797.x.

Gherardi, F. & Barbaresi, S., 2008, 'Feeding opportunism of the red swamp crayfish *Procambarus clarkii*, an invasive species, *Freshwater crayfish* 16, 77–85. https://doi.org/10.1080/1023624021000014761

Gherardi, F., Britton, J.R., Mavuti, K.M., Pacini, N., Grey, J., Tricarico, E. & Harper, D.M., 2011, 'A review of allodiversity in Lake Naivasha, Kenya: Developing conservation actions to protect East African lakes from the negative impacts of alien species', *Biological* Conservation 144, 2585–2596. https://doi.org/10.1016/j.biocon.2011.07.020.

Jackson, M.C., Grey, J., Miller, K., Britton, J.R. & Donohue, I., 2016, 'Dietary niche constriction when invaders meet natives: Evidence from freshwater decapods', *Journal of Animal Ecology* 85, 1098–1107. https://doi.org/10.1051/kmae:2005031.

Lodge, D.M., Deines, A., Gherardi, F., Yeo, D.C., Arcella, T., Baldridge, A.K., et al., 2012, 'Global introductions of crayfishes: Evaluating the impact of species invasions on ecosystem services', *Annual Review of Ecology Evolution Systematics* 43, 449–472. https://doi.org/10.1146/annurev-ecolsys-111511-103919

Longshaw, M. 2011, ‘Diseases of crayfish: A review’, *Journal of Invertebrate Pathology* 106, 54–70. https://doi.org/10.1016/j.jip.2010.09.013.

McMahon, T.A., Brannelly, L.A., Chatfield, M.W., Johnson, P.T., Joseph, M.B., McKenzie, V.J., et al., 2013, ‘Chytrid fungus *Batrachochytrium dendrobatidis* has non-amphibian hosts and releases chemicals that cause pathology in the absence of infection’, *Proceedings of the National Academy of Sciences of the United States of America* 110, 210–215. https://doi.org/10.1073/pnas.1200592110.

Nunes, A.L., Hoffman, C.A., Zengeya, T.A., Measey, G.J. & Weyl, O.L.F, 2017a, ‘Red swamp crayfish, *Procambarus clarkii*, found in South Africa 22 years after attempted eradication’, *Aquatic Conservation: Marine & Freshwater Ecosystem* 27, 1334–1340. https://doi.org/10.1002/aqc.2741.

Nunes, A.L., Zengeya, T.A., Measey, G.J. & Weyl, O.L.F., 2017b, ‘Freshwater crayfish invasions in South Africa: past, present and potential future’, *African Journal of Aquatic Science* 42, 309–323. https://doi.org/10.2989/16085914.2017.1405788.

Putra, M.D, Bláha, M., Wardiatno, Y., Krisanti, M., Bystřický, P.K, Kouba, A., et al., 2017, ‘*Procambarus clarkii* (Girard, 1852) and crayfish plague as new threats for biodiversity in Indonesia’, *Aquatic Conservation: Marine & Freshwater Ecosystems* 28, 1434–1440. https://doi.org/10.1002/aqc.2970.

Rosenthal, S.K., Lodge, D.M., Mavuti, K.M., Muohi, W., Ochieng, P., Mungai, B.N. & Mkoji, G.M., 2005, ‘Comparing macrophyte herbivory by introduced Louisiana crayfish (*Procambarus clarkii*) (Crustacea: Cambaridea) and native Dytiscid beetles (*Cybister tripunctatus*) in Kenya, *African Journal of Aquatic Science*, 30, 157–162, https://doi.org/10.2989/16085910509503850.

Souty-Grosset, C., Anastacio, P.M., Aquiloni, L., Banha, F., Choquer, J., Chucholl, C., et al. 2016, ‘The red swamp crayfish *Procambarus clarkii* in Europe: Impacts on aquatic ecosystems and human well-being’, *Limnologica* 58, 78–93. https://doi.org/10.1016/j.limno.2016.03.003.

#### 4. Management

| **MAN1 What is the feasibility to stop future immigration?** | |
|---|---|
| Response: Low | Confidence: Low |
| Rationale: Preventing introductions via intentional introductions may be feasible through regulation of imports (Zengeya & Wilson 2020), however preventing unintentional introductions (e.g., through natural dispersal) will be more difficult. *Procambarus clarkii* have established populations in Zimbabwe, the extent of invasion still needs to be determined (Loureiro et al. 2015), however this is cause for concern because connected waterways can facilitate dispersal. | |
| References:<br><br>Loureiro, T.G., Anastácio, P.M.S.G., Araujo, P.B., Souty-Grosset, C. & Almerão, M.P., 2015, ‘Red swamp crayfish: biology, ecology and invasion–an overview’, *Nauplius* 23(1), 1–19. https://doi.org/10.1590/S0104-64972014002214.<br>Zengeya, T.A. & Wilson, J.R., 2020, *The status of biological invasions and their management in South Africa in 2019*, South African National Biodiversity Institute, Kirstenbosch and DSI-NRF Centre of Excellence for Invasion Biology, Stellenbosch. http://dx.doi.org/10.5281/zenodo.3947613. | |

| **MAN2 Benefits of the Taxon** | |
|---|---|
| **MAN2a Socio-economic benefits of the *Taxon*** | |
| Response: Low | Confidence: Medium |
| Rationale: *Procambarus clarkii* is widely used in aquaculture in several countries, with the highest production in China and the US. In 2005, China produced over 80 000 tonnes (estimated value US$ 303 million) and the US produced roughly 33 000 tonnes (estimated value US$ 48.6 million). Some fisheries exist in other countries | |

(Costa Rica Kenya, Mexico Portugal, Spain and Zambia) but there are discrepancies between different sources. It is, therefore, difficult to estimate the monetary value generated.

There is no information on the production of P. *clarkii* in South Africa, but other crayfish species (*C. temuimanus*) has been farmed on very small scales (Burgess 2007). Therefore, the response has been scored as low (Nunes et al. 2017b; Madzivanzira et al. 2021).

*Procambarus clarkii* is considered to be popular in the pet trade industry elsewhere in the world (Faulkes 2015) but the benefits derived from the pet trade industry in South Africa have yet to be evaluated.

References:

Faulkes, Z., 2015, 'The global trade in crayfish as pets', *Crustacean Research* 44, 75–92. https://doi.org/10.18353/crustacea.44.0_75.

Madzivanzira, T.C., South, J., Wood, L.E., Nunes, A.L. & Weyl, O.L.F., 2021, 'A review of freshwater crayfish introductions in Africa', *Reviews in Fisheries Science & Aquaculture*,1–21. https://doi.org/10.1080/23308249.2020.1802405.

Nunes, A.L., Zengeya, T.A., Measey, G.J. & Weyl, O.L.F., 2017b, 'Freshwater crayfish invasions in South Africa: past, present and potential future', *African Journal of Aquatic Science* 42, 309–323. https://doi.org/10.2989/16085914.2017.1405788

Souty-Grosset, C., Anastácio, P.M., Aquiloni, L., Banha, F., Choquer, J., Chucholl, C., et al., 2016, 'The red swamp crayfish *Procambarus clarkii* in Europe: Impacts on aquatic ecosystems and human well-being', *Limnologica* 58, 78–93https://doi.org/10.1016/j.limno.2016.03.003.

**MAN2b Environmental benefits of the *Taxon***

| Response: Low | Confidence: Low |
|---|---|

Rationale: There are no recorded environmental benefits of *P. clarkii*.

References:

**MAN3 Ease of management**

**MAN3a How accessible are populations?**

| Response: 2 (difficult to access) | Confidence: Medium |
|---|---|

Rationale: *Procambarus clarkii* is known to occur at several sites in Mpumalanga, Free State and Western Cape Provinces. Two populations are confined to dams (Barkhuizen et al. 2022; Nunes et al. 2017a) and the one in the Western Cape occurs in the tributary of the lower Olifants River and irrigation systems around Vredendal. One dam however, is on private property, and thus would require access to be granted by the landowner (Nunes et al. 2017a, 2017b). Being a aquatic species, accessibility is difficult.

References:

Barkhuizen, L.M., Madzivanzira, T.C. & South, J., 2022, 'Population ecology of a wild population of red swamp crayfish *Procambarus clarkii* (Girard, 1852) in the Free State Province, South Africa and implications for eradication efforts', *BioInvasions Records* 11, 181-191. https://doi.org/10.3391/bir.2022.

Nunes, A.L., Hoffman, C.A., Zengeya, T.A., Measey, G.J. & Weyl, O.L.F, 2017a, 'Red swamp crayfish, *Procambarus clarkii*, found in South Africa 22 years after attempted eradication', *Aquatic Conservation: Marine & Freshwater Ecosystems* 27, 1334–1340. https://doi.org/10.1002/aqc.2741.

Nunes, A.L., Zengeya, T.A., Measey, G.J. & Weyl, O.L.F., 2017b, 'Freshwater crayfish invasions in South Africa: past, present and potential future', *African Journal of Aquatic Science* 42, 309–323. https://doi.org/10.2989/16085914.2017.1405788.

**MAN3b Is detectability critically time-dependent?**

| Response: 2 (yes) | Confidence: High |
|---|---|

Rationale: *Procambarus clarkii* can be detected throughout the year (de Moor 2002; Nunes et al. 2017a, 2017b). However, freshwater species in general can go undetected for years and the burrowing activities of *P. clarkii* could further impede detection (Barkhuizen et al. 2022; Nunes et al. 2017a).

References:
Barkhuizen, L.M., Madzivanzira, T.C. & South, J., 2022, 'Population ecology of a wild population of red swamp crayfish *Procambarus clarkii* (Girard, 1852) in the Free State Province, South Africa and implications for eradication efforts', *BioInvasions Records* 11, 181-191. https://doi.org/10.3391/bir.2022.
de Moor, I., 2002, 'Potential impacts of alien freshwater crayfish in South Africa', *African Journal of Aquatic Science* 27, 125–139https://doi.org/10.2989/16085914.2002.9626584.
Nunes, A.L., Hoffman, C.A., Zengeya, T.A., Measey, G.J. & Weyl, O.L.F, 2017a, 'Red swamp crayfish, *Procambarus clarkii*, found in South Africa 22 years after attempted eradication', *Aquatic Conservation: Marine & Freshwater Ecosystems* 27, 1334–1340. https://doi.org/10.1002/aqc.2741.
Nunes, A.L., Zengeya, T.A., Measey, G.J. & Weyl, O.L.F., 2017b, 'Freshwater crayfish invasions in South Africa: past, present and potential future', *African Journal of Aquatic Science* 42, 309–323. https://doi.org/10.2989/16085914.2017.1405788.

**MAN3c Time to reproduction**

| Response: 2 (<1 year) | Confidence: High |
|---|---|

Rationale: *Procambarus clarkii* is a prolific breeder, reaching sexual maturity in three months (or once individuals reach six to 13 cm in length) and it can spawn twice a year (Ackefors 1999; Barbaresi & Gherardi 2000). Females display parental care with newly born individuals staying with the mother in burrows up to two moult cycles (Barbaresi & Gherardi 2000).

References:

Ackefors, H., 1999, 'The positive effects of established crayfish introductions in Europe', In" Gherardi, F. & Holdich, D.M. (eds.), *Crustacean Issues 11: 'Crayfish in Europe as Alien Species (How to make the best of a bad situation?)',* pp. 49–62, A.A. Balkema, Rotterdam, Netherlands.
Barbaresi, S & Gherardi, F., 2000, 'The invasion of the alien crayfish *Procambarus clarkii* in Europe, with particular reference to Italy', *Biological Invasions* 2, 259–264.

**MAN3d Propagule persistence**

| Response: 0 (<1 year) | Confidence: High |
|---|---|

Rationale: *Procambarus clarkii* is solely an r-strategist with a short lifespan with multiple spawning events (Huner & Barr 1991; Loureiro et al. 2015). The number of eggs laid is usually determined by the size of female. A female can produce up to 500 eggs and hatchlings hang under the female's tail for many weeks until they can feed on their own (Loureiro et al. 2015).

References:
Huner, J.V. & Barr, J.E., 1991, *Red swamp crayfish, biology, and exploitation (3rd ed)*, Louisiana State University, Baton Rouge, Louisiana, USA

Loureiro, T.G., Anastácio, P.M.S.G., Araujo, P.B., Souty-Grosset, C. & Almerão, M.P., 2015, 'Red swamp crayfish: biology, ecology and invasion–an overview', *Nauplius* 23(1), 1–19. https://doi.org/10.1590/S0104-64972014002214.

**MAN3 Ease of management**

| Response: Difficult | Confidence: High |
|---|---|

Rationale: It is difficult to detect and access *P. clarkia* individuals and populations. Moreover, the burrowing behaviour of *P. clarkii* complicates management. *Procambarus clarkii* also has a short life span of one to two years with multiple spawning events (Huner & Barr 1991; Loureiro et al. 2015; Vogt 2012). Females can lay up to 500 eggs that remain under the females' tail for weeks until they are able to feed on their own (Loureiro et al. 2015).

References:

Huner, J.V. & Barr, J.E., 1991, *Red swamp crayfish, biology, and exploitation (3rd ed)*, Louisiana State University, Baton Rouge, Louisiana, USA

Loureiro, T.G., Anastácio, P.M.S.G., Araujo, P.B., Souty-Grosset, C. & Almerão, M.P., 2015, 'Red swamp crayfish: biology, ecology and invasion–an overview', *Nauplius* 23(1), 1–19. https://doi.org/10.1590/S0104-64972014002214.

Vogt, G., 2012, 'Ageing and longevity in the Decapoda (Crustacea): A review, Zoologischer Anzeiger', *A Journal of Comparative Zoology* 251, 1–25 https://doi.org/10.1016/j.jcz.2011.05.003.

**MAN4 Has the feasibility of eradication been evaluated?**

| Response: No | Confidence: High |
|---|---|

Rationale: No formal eradication study has been conducted for *P. clarkii* in South Africa. However, an attempt at eradication was conducted 22 years ago at the Driehoek Farm—this was not successful and the method used remains unknown (Nunes et al. 2017a).

References:

Nunes, A.L., Hoffman, C.A., Zengeya, T.A., Measey, G.J. & Weyl, O.L.F, 2017a, 'Red swamp crayfish, *Procambarus clarkii*, found in South Africa 22 years after attempted eradication', *Aquatic Conservation: Marine & Freshwater Ecosystems* 27, 1334–1340. https://doi.org/10.1002/aqc.2741.

**MAN5 Control options and monitoring approaches available for the *Taxon***

Response: Eradication is not easy once *P. clarkii* has established because of its hardy nature, and conventional control methods such as biocides are neither practical nor desirable as a result of the large quantities of toxins required to treat an area (Nunes et al. 2017a). Furthermore, biocides are not target-specific, and often also adversely affect native fauna (Nunes et al. 2017a).

Mechanical methods would not be effective in the removal of individuals or treatment of burrows (Nunes et al. 2017a, 2017b). However, mechanical methods (intensive trapping, electrofishing) together with de-watering the ponds (physical removal) should be considered (Nunes et al. 2017a). Some control techniques (e.g., complete dewatering waterbodies and the use of biocides) might be undesirable to land owners because of the financial and social implications (Nunes et al. 2017a, 2017b).

| |
|---|
| Whichever control option is chosen, implementation must occur over the period of one year at a minimum, and repeated monitoring efforts must be conducted (Nunes et al. 2017a). |
| References:<br><br>Nunes, A.L., Hoffman, C.A., Zengeya, T.A., Measey, G.J. & Weyl, O.L.F, 2017a, 'Red swamp crayfish, *Procambarus clarkii*, found in South Africa 22 years after attempted eradication', *Aquatic Conservation: Marine & Freshwater Ecosystem* 27, 1334–1340. https://doi.org/10.1002/aqc.2741.<br>Nunes, A.L., Zengeya, T.A., Measey, G.J. & Weyl, O.L.F., 2017b, 'Freshwater crayfish invasions in South Africa: past, present and potential future', *African Journal of Aquatic Science* 42, 309–323. https://doi.org/10.2989/16085914.2017.1405788. |

| **MAN6 Any other management considerations to highlight?** | |
|---|---|
| Response | No |

**5. Calculations**

**Likelihood = Probable**

| Parameter | Likelihood | Stages | Final assessment |
|---|---|---|---|
| LIK1 | 1 | P(entry) = 1 | P (invasion) = 1 |
| LIK2 | 1 | | |
| LIK3 | 1 | P(establishment) = 1 | |
| LIK4 | 1 | | |
| LIK5 | 1 | P (spread) =1 | |
| LIK6 | 1 | | |

**Consequence = Major (MR)**

| Parameter | Mechanism/sector | Response |
|---|---|---|
| CON1a | Competition | MR |
| CON1b | Predation | MR |
| CON1c | Hybridisation | DD |
| CON1d | Disease transmission | MR |
| CON1e | Parasitism | DD |
| CON1f | Poisoning/toxicity | MC |
| CON1g | Bio-fouling or other direct physical disturbance | DD |
| CON1h | Grazing/herbivory/browsing | MO |

| | | |
|---|---|---|
| CON1i | Chemical, physical, structural impact | MN |
| CON1k | Indirect impacts through interactions with other species | MC |
| CON1 | Maximum environmental impact | MR |
| CON2a | Safety | DD |
| CON2b | Material and immaterial assets | MO |
| CON2c | Health | MN |
| CON2d | Social, spiritual and cultural relations | DD |
| CON2 | Maximum socio-economic impact | MO |
| CON3 | Environmental impact of closely related taxa | Not scored as CON1 scored |
| CON4 | Socio-economic impact of closely related taxa | Not scored as CON2 scored |
| CON5 | Potential impact based on traits, experiments, or models | MR |

**Risk score = High**

| | | Consequences | | | | |
|---|---|---|---|---|---|---|
| | | MC | MN | MO | MR | MV |
| Likelihood | Extremely unlikely | low | low | low | medium | medium |
| | Very unlikely | low | low | low | medium | high |
| | Unlikely | low | low | medium | high | high |
| | Fairly probable | medium | medium | high | high | high |
| | Probable | medium | high | high | high | high |

**Ease of management = Difficult**

| Parameter | Question | Response |
|---|---|---|
| MAN3a | How accessible are populations? | 2 |
| MAN3b | Is detectability critically time-dependent? | 2 |
| MAN3c | Time to reproduction | 2 |
| MAN3d | Propagule persistence | 0 |
| **MAN3** | **SUM** | 6 |

**Appendix BAC7**: Map of the native range of *Procambarus clarkii* in North America 2023. Orange indicates the native range and brown indicates non-native populations. Map accessed from: https://nas.er.usgs.gov/queries/factsheet.aspx?SpeciesID=217#:~:text=Native%20Range%3A%20Gulf%20coastal%20plain,2007).&text=Nonindigenous%20Occurrences%3A,Table%201

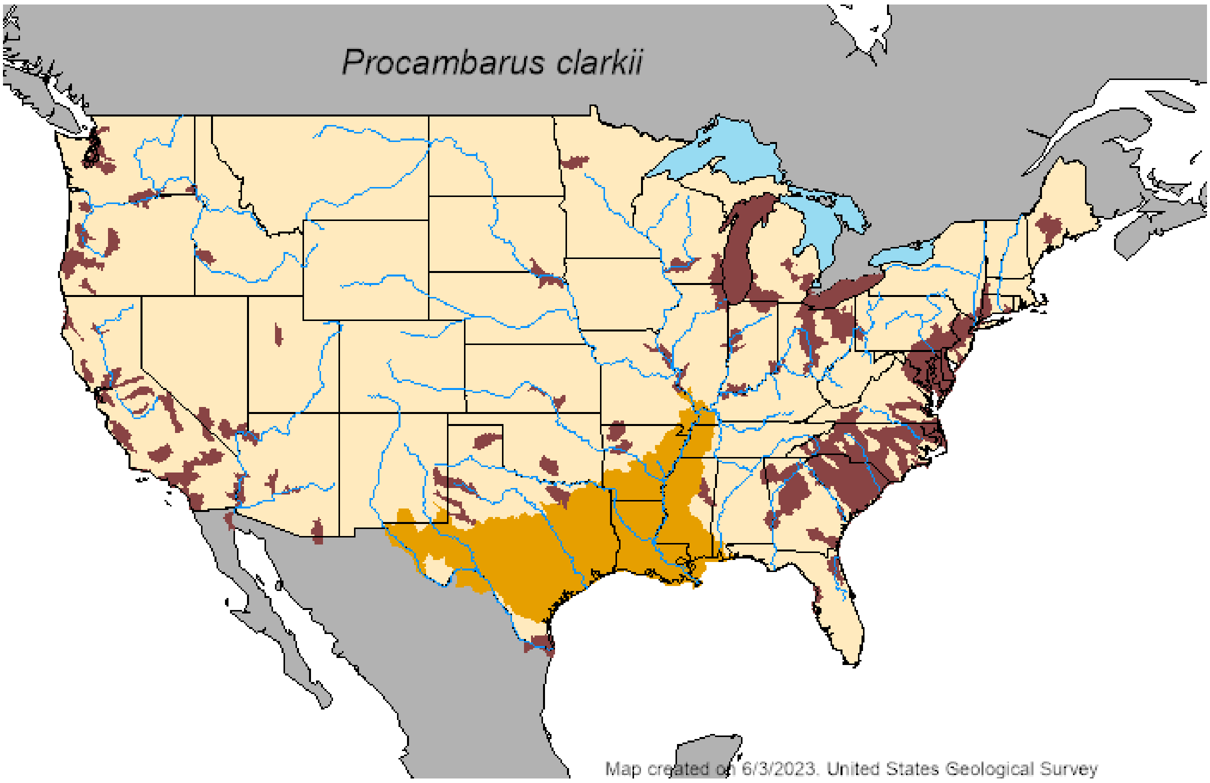

**Appendix BAC8(a)**: Map of the global alien range of *Procambarus clarkii* (indicated in red). Map obtained from Loureiro et al. (2015).

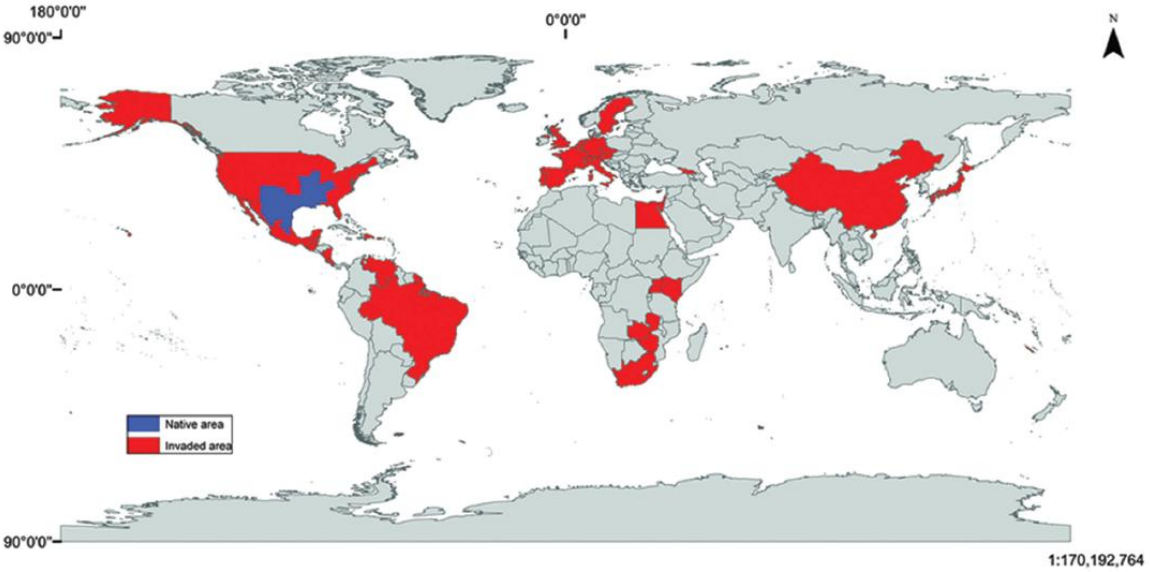

**Appendix BAC10**: The South African distribution of *Procambarus clarkii*. Locality data was taken from Barkhuisen et al. (2022) and Nunes et al. (2017a).

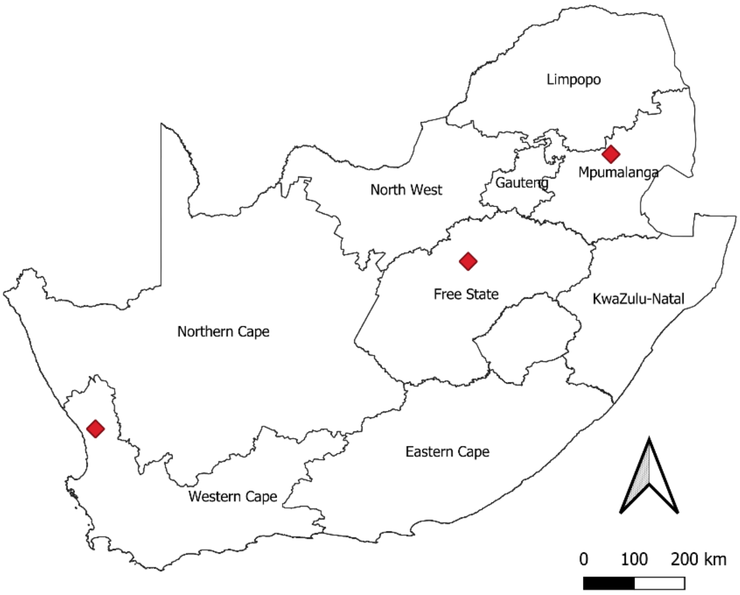

References:

Barkhuizen, L.M., Madzivanzira, T.C. & South, J. 2022, 'Population ecology of a wild population of red swamp crayfish *Procambarus clarkii* (Girard, 1852) in the Free State Province, South Africa and implications for eradication efforts', *BioInvasions Records* 11, 181–191. https://doi.org/10.3391/bir.2022.

Loureiro T.G., Anastácio, P.M.S.G., Araujo, P.B., Souty-Grosset, C. & Almerão, M.P., 2015, 'Red swamp crayfish: biology, ecology and invasion–an overview', *Nauplius* 23(1), 1–19. https://doi.org/10.1590/S0104-64972014002214.

Nunes, A.L., Hoffman, C.A., Zengeya, T.A., Measey, G.J. & Weyl, O.L.F., 2017a, 'Red swamp crayfish, *Procambarus clarkii*, found in South Africa 22 years after attempted eradication', *Aquatic Conservation: Marine & Freshwater Ecosystem* 27, 1334-1340. https://doi.org/10.1002/aqc.2741.

#### 1.14 Risk analysis report for Marmorkrebs (*Procambarus virginalis*)

***Taxon***: *Procambarus virginalis* (Lyko, 2017)

***Area***: South Africa

**Picture of *Taxon***

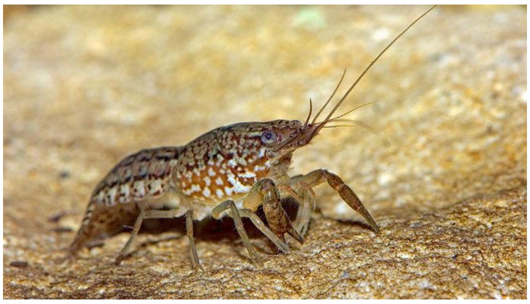

http://www.sciencemag.org/news/2018/02/aquarium-accident-may-have-given-crayfish-dna-take-over-world

**Alien distribution map**

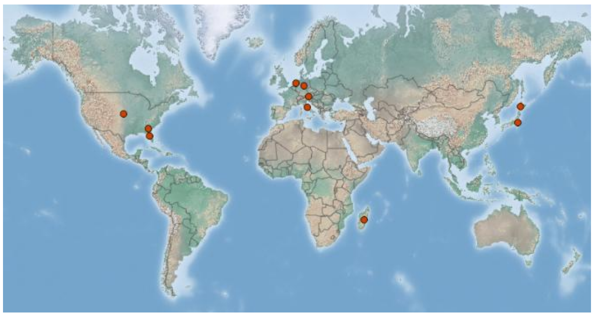

**https://www.gbif.org/species/2227309**

**Risk Assessment summary:**

Marmorkrebs (*Procambarus virginalis*), is a parthenogenetic crayfish whose native origin has been deffisulty to decern but likely to be Florida, USA. It has become widely established in several countries in Europe and Madagascar, however, there are no known populations in South Africa or any neighboring countries. *Procambarus virginalis* is a highly sought after aquarium pet species and is available to buy online in several countries and is likely to be moved by humans through the pet trade. There are no documented impacts from *P. virginalis* and potential impacts were inferred from *Procambarus clarkii*, a closely-related species. *Procambarus clarkii* is a facultative omnivore and an aggressive competitor that often displaces native species in areas of introduction through predation, competitive exclusions, and transmission of diseases. It has also been implicated in causing habitat alteration through excessive grazing of macrophytes, and its burrowing activities increase rates of soil erosion and cause physical damage to agricultural infrastructure such as irrigation canals. *Procambarus clarkii* serves as vector for several parasites and diseases some of which are zoonotic. *Procambarus virginalis* is likely to have similar impacts in recipient areas of introduction.

**Risk score:**

High

**Management options summary:**

There are no records of *Procambarus virginalis* being in present in South Africa. When evaluating suitable methods however, it is important to take into account the burrowing behaviour of the species. The species is popular in the aquarium trade, and as a result, the potential of accidental or intentional release of the species into new areas by humans is high.

**Ease of management:**

Difficult

**Recommendations:**

*Procambarus virginalis* is not currently listed under the NEM:BA Alien and Invasive Species (A&IS) Regulations.

**Listing under NEM:BA A&IS lists of 2014 as amended 2020**

| | |
|---|---|
| *Procambarus virginalis* is recommended to be added to a future prohibited species list. The illegal pet trade industry still poses a significant risk for intentional and accidental release of the species into the wild. There is, therefore, a need to assess the trade of and movement of the taxon through the aquarium trade in order to implement appropriate management strategies. | Not listed<br>**Recommended listing category:**<br>Prohibited list [1] |

[1] The absence of *Procambarus virginalis* in South Africa should be confirmed and the taxon should be reassessed in five years (i.e., by 2029).

#### 1. Background

| **BAC1 Name of assessor(s)** | |
|---|---|
| Name of lead assessor | Lee-Anne Botha |
| Additional assessor (1) | Tsungai Zengeya |
| Additional assessor (2) | Christian Chimimba |
| **BAC2 Contact details of assessor (s)** | |
| Lead assessor | Organisational affiliation: Department of Zoology and Entomology, University of Pretoria and The South African National Biodiversity Institute (SANBI). |
| | |
| Additional assessor (1) | Organisational affiliation: Department of Zoology and Entomology, University of Pretoria and The South African National Biodiversity Institute (SANBI). |
| | |
| Additional assessor (2) | Organisational affiliation: Department of Zoology and Entomology, University of Pretoria |
| | |
| **BAC3 Name(s) and contact details of expert(s) consulted** | |
| Expert (1) | Name: |
| | email: |
| | Phone: |
| Expert (2) | Name: |
| | email: |
| | Phone: |
| Comments: none | |
| **BAC4 Scientific name of *Taxon* under assessment** | |
| *Taxon* name: *Procambarus virginalis* | Authority: Lyko, 2017 |
| Comments: *Procambarus virginalis* is listed as the accepted name for marbled crayfish (Lyko 2017; GBIF 2023) | |
| References:<br>*Procambarus virginalis* Lyko, 2017 in Döring M (2022). English Wikipedia - Species Pages. Wikimedia Foundation. Checklist dataset https://doi.org/10.15468/c3kkgh accessed via GBIF.org on 2025-02-14.<br>Lyko, F., 2017. The marbled crayfish (Decapoda: Cambaridae) represents an independent new species. Zootaxa, 4363(4), pp.544-552, https://doi.org/10.11646/zootaxa.4363.4.6. | |
| **BAC5 Synonym(s) considered** | |
| Synonyms: *Procambarus fallax* f. *virginalis* | |
| Comments: *Procambarus fallax* f. *virginalis* is listed as a synonym of *Procambarus virginalis* (GBIF 2023) and it was used in the listreature search for this risk analyis | |
| References:<br>*Procambarus fallax f. virginalis* Martin, Dorn, Kawai, van der Heiden & Scholtz, 2010 in GBIF Secretariat (2023). GBIF Backbone Taxonomy. Checklist dataset https://doi.org/10.15468/39omei accessed via GBIF.org on 2024-07-17. | |
| **BAC6 Common name(s) considered** | |
| Common names: Marbled crayfish \| Marmorkrebs | |
| Comments: There are two English common names for *Procambarus virginalis* (Marbled crayfish and Marmorkrebs) (GBIF 2023; CABI, 2024) and these were used in the listreature search for this risk analyis | |

References:
CABI, 2024. *Procambarus fallax* f. *virginalis* (Marmorkrebs). Invasive Species Compendium, CABI Publishing, Wallingford, UK. https://www.cabidigitallibrary.org/doi/10.1079/cabicompendium.110477 (accessed 10 July 2024)
*Procambarus fallax f. virginalis* Martin, Dorn, Kawai, van der Heiden & Scholtz, 2010 in GBIF Secretariat (2023). GBIF Backbone Taxonomy. Checklist dataset https://doi.org/10.15468/39omei accessed via GBIF.org on 2024-07-17.

| **BAC7 What is the native range of the *Taxon*?** (add map in Appendix BAC7) | |
|---|---|
| Response: Florida, USA | Confidence: Medium |

Comments: It was first discovered in a pet shop in Germany in the mid-1990s (Lukhap 2001) and has become a very popular pet species since then (Feria and Faulkes 2011). It native range is unknown due to uncertanity over its taxonomic status - "Most recently, Martin et al. (2010a) suggested that the Marmorkrebs is the parthenogenetic form of *Procambarus fallax* (Hagen, 1870) and proposed the tentative scientific name *Procambarus fallax* f. *virginalis*. *Procambarus fallax* occurs in southern Georgia and Florida, and it is therefore reasonable to assume that the Marmorkrebs originates also from the southeastern United States, although an indigenous population has never been reported" (Chucholl and Pfeiffer, 2010). A phylogeographic reconstruction of the marbel crayfish origin indicated that it might have originated form Florida in the USA (Gutekunst et al. 2021).

References:
Chucholl, C. & Pfeiffer, M. 2010, 'First evidence for an established Marmorkrebs (Decapoda, Astacida, Cambaridae) population in Southwestern Germany, in syntopic occurrence with *Orconectes limosus* (Rafinesque, 1817)', *Aquatic invasions* 5, 405-412. https://doi.org/10.3391/ai.2010.5.4.10
Feria, T.P. & Faulkes, Z., 2011, 'Forecasting the distribution of Marmorkrebs, a parthenogenetic crayfish with high invasive potential, in Madagascar, Europe, and North America', *Aquatic Invasions* 6, 55–67. https://doi.org/10.3391/ai.2011.6.1.07.
Gutekunst, J., Maiakovska, O., Hanna, K., Provataris, P., Horn, H., Wolf, S., Skelton, C.E., Dorn, N.J. and Lyko, F., 2021, 'Phylogeographic reconstruction of the marbled crayfish origin', *Communications Biology*, *4*(1), p.1096. https://doi.org/10.1038/s42003-021-02609-w
Lukhaup, C. (2001). Procambarus sp.–der Marmorkrebs. *Aquaristik aktuell*, *7*(8), 48-51.
Martin P, Dorn N, Kawai T, Heiden Cvan der, Scholtz G, 2010, 'The enigmatic Marmorkrebs (marbled crayfish) is the parthenogenetic form of *Procambarus fallax* (Hagen, 1870). *Contributions to Zoology*, 79(3), 107-118. https://doi.org/10.1163/18759866-07903003

| **BAC8 What is the global alien range of the *Taxon*?** (add map in Appendix BAC8) | |
|---|---|
| Response: Austria, Croatia, Czech Republic, Germany, Hungary, Ireland, Italy, Japan, Kazakhstan, Madagascar, Netherlands, Romania, Russia, Slovakia, Sweden, Ukraine, United Kingdom, USA | Confidence: Medium |

Comments: Introductions have occureed in several countires in Europe, USA and Madagascar (Jones et al. 2009; Chucholl and Pfeiffer, 2010; Faulkes 2010; Soes & Koese 2010; Feria & Faulkes 2011; Chucholl et al. 2012; Faulkes et al. 2012; Bohman et al. 2013; Faulkes 2014; Kouba et al. 2014; Patoka et al. 2014; Maguire et al. 2018; Lőkkös et al. 2016;; Novitsky & Son 2016; Lipták et al. 2017; Pârvulescu et al. 2017; Uderbayev et al. 2017)

References:

Bohman, P., Edsman, L., Martin, P., & Scholtz, G. 2013, 'The first Marmorkrebs (Decapoda: Astacida: Cambaridae) in Scandinavia', *BioInvasions Records*, 2(3), 227-232 https://doi.org/10.3391/bir.2013.2.3.09 .

Chucholl, C. & Pfeiffer, M. 2010, 'First evidence for an established Marmorkrebs (Decapoda, Astacida, Cambaridae) population in Southwestern Germany, in syntopic occurrence with Orconectes limosus (Rafinesque, 1817)', *Aquatic invasions* 5, 405-412. https://doi.org/10.3391/ai.2010.5.4.10.

Chucholl, C., Morawetz, K., & Groß, H. 2012, 'The clones are coming-strong increase in Marmorkrebs [*Procambarus fallax* (Hagen, 1870) f. *virginalis*] records from Europe'. *Aquatic Invasions*, 7(4), 511-519. https://doi.org/10.3391/ai.2012.7.4.008.

Faulkes, Z. 2010, 'The spread of the parthenogenetic marbled crayfish, Marmorkrebs (*Procambarus* sp.), in the North American pet trade', *Aquatic Invasions*, 5(4), 447-450. https://doi.org/10.3391/ai.2010.5.4.16

Faulkes, Z. 2014, 'A bomb set to drop: parthenogenetic Marmorkrebs for sale in Ireland, a European location without non-indigenous crayfish. *Management of Biological Invasions*, 6(1), 111-114. https://doi.org/10.3391/mbi.2015.6.1.09.

Faulkes, Z., Feria, T. P., & Muñoz, J. 2012, 'Do Marmorkrebs, *Procambarus fallax* f. *virginalis*, threaten freshwater Japanese ecosystems? *Aquatic Biosystems*, 8(1), 13. https://doi.org/10.1186/2046-9063-8-13

Feria, T.P. & Faulkes, Z., 2011, 'Forecasting the distribution of Marmorkrebs, a parthenogenetic crayfish with high invasive potential, in Madagascar, Europe, and North America', *Aquatic Invasions* 6, 55-67. https://doi.org/10.3391/ai.2011.6.1.07.

Jones, J.P.G., Rasamy, J.R., Harvey, A., Toon, A., Birgit Oidtmann, B., Randrianarison, M.H., Raminosoa, N. & Ravoahangimalala, O.R., 2009, 'The perfect invader: a parthenogenetic crayfish poses a new threat to Madagascar's freshwater biodiversity', *Biological Invasions*, 11, 1475–1482, https://doi.org/10.1007/s10530-008-9334-y.

Kouba, A., Petrusek, A. & Kozák, P. 2014, 'Continental-wide distribution of crayfish species in Europe: update and maps', *Knowledge and Management of Aquatic Ecosystems*, (413), 05. https://doi.org/10.1051/kmae/2014007.

Lipták, B., Mojžišová, M., Gruľa, D., Christophoryová, J., Jablonski, D., Bláha, M., Petrusek, A. & Kouba, A. 2017, 'Slovak section of the Danube has its well-established breeding ground of marbled crayfish *Procambarus fallax* f. *virginalis*. *Knowledge & Management of Aquatic Ecosystems*, (418), 40. https://doi.org/10.1051/kmae/2017029.

Lőkkös, A., Müller, T., Kovács, K., Várkonyi, L., Specziár, A., & Martin, P. 2016, 'The alien, parthenogenetic marbled crayfish (Decapoda: Cambaridae) is entering Kis-Balaton (Hungary), one of Europe's most important wetland biotopes', *Knowledge and Management of Aquatic Ecosystems*, (417), 16. https://doi.org/10.1051/kmae/2016003.

Maguire, I., Klobučar, G., Žganec, K., Jelić, M., Lucić, A., & Hudina, S. 2018. 'Recent changes in distribution pattern of freshwater crayfish in Croatia− threats and perspectives', *Knowledge & Management of Aquatic Ecosystems*, (419), 2. https://doi.org/10.1051/kmae/2017053

Novitsky, R. A., & Son, M. O. 2016, 'The first records of Marmorkrebs [*Procambarus fallax* (Hagen, 1870) f. *virginalis*](Crustacea, Decapoda, Cambaridae) in Ukraine. *Ecologica Montenegrina*, 5, 44-46. https://doi.org/10.37828/em.2016.5.8.

Pârvulescu, L., Togor, A., Lele, S. F., Scheu, S., Şinca, D., & Panteleit, J. 2017, 'First established population of marbled crayfish *Procambarus fallax* (Hagen, 1870) f. *virginalis* (Decapoda, Cambaridae) in Romania. BioInvasions Record, 6(4). https://doi.org/10.3391/bir.2017.6.4.09.

Patoka, J., Petrtýl, M., & Kalous, L. 2014, 'Garden ponds as potential introduction pathway of ornamental crayfish. *Knowledge and Management of Aquatic Ecosystems*, (414), 13. https://doi.org/10.1051/kmae/2014019.

Soes, M. & Koese, B. 2010, 'Invasive freshwater crayfish in the Netherlands: a preliminary risk analysis. Invasive freshwater crayfish in the Netherlands: a preliminary risk analysis'.

Uderbayev, T., Patoka, J., Beisembayev, R., Petrtýl, M., Bláha, M., & Kouba, A. 2017, 'Risk assessment of pet-traded decapod crustaceans in the Republic of Kazakhstan, the leading country in Central Asia. *Knowledge & Management of Aquatic Ecosystems*, (418), 30. https://doi.org/10.1051/kmae/2017018

**BAC9 Geographic scope = the *Area* under consideration**

*Area* of assessment: South Africa

Comments: The geograhic scope of the assessment is limited to South Africa

**BAC10 Is the *Taxon* present in the *Area*?**

| | |
|---|---|
| Response: No | Confidence: High |

Comments: There are no records of *Procambarus virginalis* in South Africa (Nunes et al. 2017).

References:

Nunes, A.L., Zengeya, T.A., Measey, G.J. & Weyl, O.L.F., 2017, 'Freshwater crayfish invasions in South Africa: past, present and potential future', *African Journal of Aquatic Science* 42, 309–323, https://doi.org/10.2989/16085914.2017.1405788

**BAC11 Availability of physical specimen**

| | |
|---|---|
| Response: No | Confidence in ID: |

Herbarium or museum accession number: There are no records of *Procambarus virginalis* in South Africa (Nunes et al. 2017).

References:

Nunes, A.L., Zengeya, T.A., Measey, G.J. & Weyl, O.L.F., 2017, 'Freshwater crayfish invasions in South Africa: past, present and potential future', *African Journal of Aquatic Science* 42, 309–323, https://doi.org/10.2989/16085914.2017.1405788

**BAC12 Is the *Taxon* native to the *Area* or part of the *Area*?**

| | | |
|---|---|---|
| The *Taxon* is native to (part of) the *Area*. | No | Confidence: High |
| The *Taxon* is alien in (part of) the *Area*. | Yes | Confidence: High |

Comments: The origin of *Procambarus virginalis* is presumed to be Florida in the USA and there are no native crayfish species in South Africa (de Moor 2002; Martin et al. 2010; Nunes et al. 2017; Gutekunst et al. 2021).

References:

de Moor, I., 2002, 'Potential impacts of alien freshwater crayfish in South Africa', *African Journal of Aquatic Science*, 27, 125-139, https://doi.org/10.2989/16085914.2002.9626584

Gutekunst, J., Maiakovska, O., Hanna, K., Provataris, P., Horn, H., Wolf, S., Skelton, C.E., Dorn, N.J. and Lyko, F., 2021, 'Phylogeographic reconstruction of the marbled crayfish origin', *Communications Biology*, 4(1), p.1096. https://doi.org/10.1038/s42003-021-02609-w

Martin P, Dorn N, Kawai T, Heiden Cvan der, Scholtz G, 2010a. The enigmatic Marmorkrebs (marbled crayfish) is the parthenogenetic form of *Procambarus fallax* (Hagen, 1870). *Contributions to Zoology*, 79(3), 107-118. https://doi.org/10.1163/18759866-07903003

Nunes, A.L., Zengeya, T.A., Measey, G.J. & Weyl, O.L.F., 2017, 'Freshwater crayfish invasions in South Africa: past, present and potential future', *African Journal of Aquatic Science*, 42, 309-323, https://doi.org/10.2989/16085914.2017.1405788

**BAC13 What is the *Taxon*'s introduction status in the *Area*?**

| | | |
|---|---|---|
| The *Taxon* is in cultivation/containment. | Don't know | Confidence: Low |
| The *Taxon* is present in the wild. | None | Confidence: |
| The *Taxon* has established/naturalised. | None | Confidence: |
| The *Taxon* is invasive. | None | Confidence: |

Comments: There are no verified records of *Procambarus virginalis* presence in South Africa (Nunes et al. 2017).

References:

Nunes, A.L., Zengeya, T.A., Measey, G.J. & Weyl, O.L.F., 2017b, 'Freshwater crayfish invasions in South Africa: past, present and potential future', *African Journal of Aquatic Science* 42, 309–323. https://doi.org/10.2989/16085914.2017.1405788.

**BAC14 Primary (introduction) pathways**

| | | |
|---|---|---|
| Release | NA | Confidence: |

| | | |
|---|---|---|
| Escape | Aquarium trade \| Aquaculture | Confidence: High |
| Contaminant | NA | Confidence: |
| Stowaway | NA | Confidence: |
| Corridor | NA | Confidence: |
| Unaided | NA | Confidence: |
| Comments: *Procambarus virginalis* has been introduced to several countries globally mainly through the pet trade (e.g., Souty-Grosset et al., 2006; Feria et al. 2011; Faulkes et al. 2012; Chucholl and Pfeiffer, 2010; Olden & Carvalho 2024). Secondary introductions for human consumption have also occurred in Madagascar (Jones et al. 2009). | | |

References:

Chucholl, C. & Pfeiffer, M. 2010, 'First evidence for an established Marmorkrebs (Decapoda, Astacida, Cambaridae) population in Southwestern Germany, in syntopic occurrence with *Orconectes limosus* (Rafinesque, 1817)', *Aquatic invasions* 5, 405-412. https://doi.org/10.3391/ai.2010.5.4.10

Faulkes Z. Feria TP, Muñoz J. 2012. Do Marmorkrebs, *Procambarus fallax f. virginalis*, threaten freshwater Japanese ecosystems?. *Aquatic Biosystems* 8: 13, https://doi.org/10.1186/2046-9063-8-13

Faulkes, Z, 2015, 'The global trade in crayfish as pets', *Crustacean Research* 44, 75–92. https://doi.org/10.18353/crustacea.44.0_75

Feria, T.P. & Faulkes, Z., 2011, 'Forecasting the distribution of Marmorkrebs, a parthenogenetic crayfish with high invasive potential, in Madagascar, Europe, and North America', *Aquatic Invasions* 6, 55–67. https://doi.org/10.3391/ai.2011.6.1.07.

Jones, J.P.G., Rasamy, J.R., Harvey, A., Toon, A., Birgit Oidtmann, B., Randrianarison, M.H., Raminosoa, N. & Ravoahangimalala, O.R., 2009, 'The perfect invader: a parthenogenetic crayfish poses a new threat to Madagascar's freshwater biodiversity', Biological Invasions, 11, 1475–1482, https://doi.org/10.1007/s10530-008-9334-y

Souty-Grosset C, Holdich D.M., Noel P.Y., Reynolds J.D., Haffner, P, 2006. Atlas of crayfish in Europe. Paris, France: Muséum national d´Histoire naturelle, 187 pp.

Olden, J.D., Carvalho, F.A. C., 2024, 'Global invasion and biosecurity risk from the online trade in ornamental crayfish, *Conservation Biology*, 38, e14359. https://doi.org/10.1111/cobi.14359

**2. Likelihood**

| **LIK1 Likelihood of entry via unaided primary pathways** | |
|---|---|
| Response: Very unlikely (p = 0.0027) | Confidence: Medium |
| Rationale: There is no evidence that *Procambarus virginalis* is present in South Africa or has established in the wild in neighbouring countries (Nunes et al. 2017; Madzivanzira et al. 2020) therefore the probability o it entering South Africa through unaided pathways (connected waterways) is very unlikely (p = 0.0027). | |

References:

Madzivanzira, T.C., South, J., Wood, L.E., Nunes, A.L. & Weyl, O.L.F., 2020, 'A review of freshwater crayfish introductions in Africa', *Reviews in Fisheries Science & Aquaculture*, 1-21. https://doi.org/10.1080/23308249.2020.1802405.

Nunes, A.L., Zengeya, T.A., Measey, G.J. & Weyl, O.L.F., 2017, 'Freshwater crayfish invasions in South Africa: past, present and potential future', *African Journal of Aquatic Science* 42, 309–323, https://doi.org/10.2989/16085914.2017.1405788.

| **LIK2 Likelihood of entry via human aided primary pathways** | |
|---|---|
| Response: Probable (p = 1) | Confidence: Medium |

| |
|---|
| Rationale: *Procambarus virginalis* is a very popular species in the global pet trade due to its size and because of its pathogenesis abilities that makes it very easy to rear (Faulkes 2015; Olden &Carvalho 2024). It is present in Madagascar, and it is probable that it could be introduced into South Africa through the pet trade. There is no evidence to show that *P. virginalis* is currently traded in the South Africa pet trade industry, but this needs to be verified through dedicated surveys (Nunes et al. 2017) |
| References:<br><br>Faulkes, Z, 2015, 'The global trade in crayfish as pets', *Crustacean Research* 44, 75–92. https://doi.org/10.18353/crustacea.44.0_75.<br><br>Nunes, A.L., Zengeya, T.A., Measey, G.J. & Weyl, O.L.F., 2017, 'Freshwater crayfish invasions in South Africa: past, present and potential future', African Journal of Aquatic Science 42, 309–323, https://doi.org/10.2989/16085914.2017.1405788.<br><br>Olden, J.D., Carvalho, F.A. C., 2024, 'Global invasion and biosecurity risk from the online trade in ornamental crayfish, *Conservation Biology*, 38, e14359. https://doi.org/10.1111/cobi.14359 |

| **LIK3 Habitat suitability** | |
|---|---|
| Response: Fairly probable (p = 0.5) | Confidence: Low |
| Rationale: Due to the uncertainty with regards to the native range of *Procambarus virginalis* and no record of indigenous populations, nothing is known about its natural habitat (CABI 2024). Ocupies a variety of habitats from rivers, lakes, swamps, and drainage ditches and fishponds (Dorn and Volin 2009; Chucholl and Pfeiffer 2010; Feria and Faulkes 2011). *Procambarus virginalis* is closely related to *Procambarus fallax* and some habitat preferences can also be inferred from *P. fallax* – occurs in slow flowing water habitats such as wetlands and marshes (Hendrix & Loftus 2000; Martin et al. 2010). Mainly aquatic and only burrows under extreme conditions (Dorn & Volin 2009). | |
| References:<br><br>CABI, 2024. *Procambarus fallax* f. *virginalis* (Marmorkrebs). Invasive Species Compendium, CABI Publishing, Wallingford, UK. https://www.cabidigitallibrary.org/doi/10.1079/cabicompendium.110477 (accessed 10 July 2024)<br>Chucholl C, Pfeiffer M. 2010. First evidence for an established Marmorkrebs (Decapoda, Astacida, Cambaridae) population in Southwestern Germany, in syntopic occurrence with *Orconectes limosus* (Rafinesque, 1817). *Aquatic invasions* 5: 405-412. https://doi.org/10.3391/ai.2010.5.4.10<br>Dorn NJ, Volin JC, 2009. Resistance of crayfish (*Procambarus* spp.) populations to wetland drying depends on species and substrate. *Journal of the North American Benthological Society*, 28(4):766-777. https://doi.org/10.1899/08-151.1<br>Feria, T.P. & Faulkes, Z., 2011, 'Forecasting the distribution of Marmorkrebs, a parthenogenetic crayfish with high invasive potential, in Madagascar, Europe, and North America*', Aquatic Invasions* 6, 55-67. https://doi.org/10.3391/ai.2011.6.1.07.<br>Hendrix AN, Loftus WF, 2000. Distribution and relative abundance of the crayfishes *Procambarus alleni* (Faxon) and P. fallax (Hagen) in southern Florida. *Wetlands*, 20:194-199. https://doi.org/10.1672/0277-5212(2000)020[0194:DARAOT]2.0.CO;2 | |

| **LIK4 Climate suitability** | |
|---|---|
| Response: Fairly probable (p = 0.5) | Confidence: Low |

| |
|---|
| Rationale: *Procambarus virginalis* seem to be tolerant of a wide range of environmental conditions, including low oxygenation and temporary exposure to temperatures < 8°C and > 30°C (Seitz et al. 2005; Feria and Faulkes 2011; CABI 2023). These climatic conditions are available in South Africa. |
| References:<br><br>CABI, 2024. *Procambarus fallax* f. *virginalis* (Marmorkrebs). Invasive Species Compendium, CABI Publishing, Wallingford, UK. https://www.cabidigitallibrary.org/doi/10.1079/cabicompendium.110477 (accessed 10 July 2024)<br>Feria, TP, Faulkes Z. 2011. Forecasting the distribution of Marmorkrebs, a parthenogenetic crayfish with high invasive potential, in Madagascar, Europe, and North America. *Aquatic Invasions* 6: 55-67. https://doi.org/10.3391/ai.2011.6.1.07<br>Seitz, R., Vilpoux, K., Hopp, U., Harzsch, S. & Maier, G., 2005, 'Ontogeny of the Marmorkrebs (marbled crayfish): a parthenogenetic crayfish with unknown origin and phylogenetic position', *Journal of Experimental Zoology Part A: Comparative Experimental Biology*, 303, 393-405. https://doi.org/10.1002/jez.a.143 |

| **LIK5 Unaided secondary (dispersal) pathways** | |
|---|---|
| Response: Very unlikely (p = 0.0027) | Confidence: Medium |
| Rationale: There is no evidence that *Procambarus virginalis* is present in South Africa or has established in the wild in neighbouring countries (Nunes et al. 2017; Madzivanzira et al. 2020) therefore the probability of *Procambarus virginalis* entering South Africa through unaided pathways (connected waterways) is very unlikely (p = 0.0027). | |
| References:<br><br>Madzivanzira, T.C., South, J., Wood, L.E., Nunes, A.L. & Weyl, O.L.F., 2020, 'A review of freshwater crayfish introductions in Africa', *Reviews in Fisheries Science & Aquaculture*, 1-21. https://doi.org/10.1080/23308249.2020.1802405.<br>Nunes, A.L., Zengeya, T.A., Measey, G.J. & Weyl, O.L.F., 2017, 'Freshwater crayfish invasions in South Africa: past, present and potential future', African Journal of Aquatic Science 42, 309–323, https://doi.org/10.2989/16085914.2017.1405788. | |

| **LIK6 Human aided secondary (dispersal) pathways** | |
|---|---|
| Response: Fairly probable (p = 0.5) | Confidence: Medium |
| Rationale: Intentional release into the wild through the pet trade and aquaculture remains a concern (Feria & Faulkes 2011; Faulkes 2015). For example, secondary dispersal has occurred in Madagascar for human consumption (Jones et al. 2009). | |
| References:<br>Faulkes, Z, 2015, 'The global trade in crayfish as pets', *Crustacean Research* 44, 75-92. https://doi.org/10.18353/crustacea.44.0_75.<br>Feria, T.P. & Faulkes, Z., 2011, 'Forecasting the distribution of Marmorkrebs, a parthenogenetic crayfish with high invasive potential, in Madagascar, Europe, and North America', *Aquatic Invasions* 6, 55–67. https://doi.org/10.3391/ai.2011.6.1.07.<br>Jones, J.P.G., Rasamy, J.R., Harvey, A., Toon, A., Birgit Oidtmann, B., Randrianarison, M.H., Raminosoa, N. & Ravoahangimalala, O.R., 2009, 'The perfect invader: a parthenogenetic crayfish poses a new threat to Madagascar's freshwater biodiversity', Biological Invasions, 11, 1475–1482, https://doi.org/10.1007/s10530-008-9334-y | |

**3. Consequences**

| **CON1 Environmental impact** | |
|---|---|
| **CON1a: Competition** | |
| Response: Data Deficient (DD) | Confidence: |
| Rationale: | |
| References: | |
| **CON1b: Predation** | |
| Response: Data Deficient (DD) | Confidence: |
| Rationale: | |
| References: | |
| **CON1c: Hybridisation** | |
| Response: Data Deficient (DD) | Confidence: |
| Rationale: | |
| References: | |
| **CON1d: Transmission of disease** | |
| Response: Data Deficient (DD) | Confidence: |
| Rationale: | |
| References: | |
| **CON1e: Parasitism** | |
| Response: Data Deficient (DD) | Confidence: |
| Rationale: | |
| References: | |
| **CON1f: Poisoning/toxicity** | |
| Response: Data Deficient (DD) | Confidence: |
| Rationale: | |
| References: | |
| **CON1g: Bio-fouling or other direct physical disturbance** | |
| Response: Data Deficient (DD) | Confidence: |
| Rationale: | |

| | |
|---|---|
| References: | |
| **CON1h: Grazing/herbivory/browsing** | |
| Response: Data Deficient (DD) | Confidence: |
| Rationale: | |
| References: | |
| **CON1i: Chemical, physical or structural impact on ecosystem** | |
| Response: Data Deficient (DD) | Confidence: |
| Rationale: | |
| References: | |
| **CON1k: Indirect impacts through interactions with other species** | |
| Response: Data Deficient (DD) | Confidence: |
| Rationale: | |
| References: | |
| **CON1 Maximum environmental impact** | |
| Response: Data Deficient (DD) | Confidence: |
| Rationale: | |
| References: | |

| | |
|---|---|
| **CON2 Socio-economic impact** | |
| **CON2a: Safety** | |
| Response: Data Deficient (DD) | Confidence: |
| Rationale: | |
| References: | |
| **CON2b: Material and immaterial assets** | |
| Response: Data Deficient (DD) | Confidence: |
| Rationale: | |
| References: | |
| **CON2c: Health** | |

| | |
|---|---|
| Response: Data Deficient (DD) | Confidence: |
| Rationale: | |
| References: | |
| **CON2d: Social, spiritual and cultural relations** | |
| Response: Data Deficient (DD) | Confidence: |
| Rationale: | |
| References: | |
| **CON2 Maximum socio-economic impact** | |
| Response: Data Deficient (DD) | Confidence: |
| Rationale: | |
| References: | |

| | |
|---|---|
| **CON3 Closely related species' environmental impact** | |
| Response: Major (MR) | Confidence: High |
| Rationale: There are no recorded environmental impacts for *P. virginalis*, therefore *P. clarkii* was selected as a closely related taxon to estimate the environmental impacts that may arise from introductions of *P. virginalis*.<br><br>*Procambarus clarkii* is an aggressive species and is known to out-competes native species for shelter and spawning sites leading to reproductive interference (Cruz et al. 2006; Lodge et al. 2012). For example, in areas were *P. clarkii* has been introduced, some amphibian species (e.g., *Bufo bufo*, *B. calamita*, *Rana* spp., *Taricha torosa* and *Triturus vulgaris*) have been excluded or displaced from their natural habitats, resulting in local extinctions through reproductive failure (Cruz et al. 2006; Lodge et al. 2012). In Europe and Japan, decreases in distributional ranges and abundance of native crayfish populations (*Astacus astacus*, *Austropotamobius pallipes* and *A. torrentium*) as a direct result of competition with *P. clarkii* were observed (Cruz et al. 2006; Lodge et al. 2012). Furthermore, direct competition for food has caused dietary niche constriction and declines in populations of native crabs in some areas invaded by *P. clarkii* (Jackson et al. 2016).<br><br>Predation by *P. clarkii* in the alien range can result in local or population extinction of at least one native species (Lodge et al. 2012; Souty-Grosset et al. 2016). For example, predation by *P. clarkii* has resulted in population declines of several species of fish and amphibians by reducing their breeding success through predation on eggs and larval amphibians (Cruz et al. 2006; Ficetola et al. 2011). Indirect effects of predation results in cause trophic cascades that lead to changes in ecosystem structure and function. For example, predation on invertebrates can release algae from grazing pressure and lead to changes in the abundance and dominance of species in algal communities (Gherardi & Barbaresi 2008). | |

*Procambarus clarkii* is a host of various pathogens, parasites, and diseases (Longshaw 2011; Lodge et al. 2012). Transmission of diseases and parasites by *P. clarkii* has led to local or population extinctions of at least one native species, leading to changes in community composition (Lodge et al. 2012). For example, *P. clarkii* is a vector of crayfish plague, a disease caused by the parasitic oomycete, *Aphanomyces astaci* (Aquiloni et al, 2011; Longshaw 2011; Souty-Grosset et al. 2016). In Europe, transmission of crayfish plague by *P. clarkii* has been linked to a decline in populations of several native crayfish species of as *Astacus astacus*, *Austropotamobius pallipes* and *A. torrentium* (Holdich et al. 2009; Longshaw 2011). *Procambarus clarkii* is also host to white spot syndrome disease – a viral infection of crustaceans, and fungal pathogens such as *Batrachochytrium dendrobatids* that causes chytridiomycosis – a lethal skin infection in amphibians (Longshaw 2011; McMahon et al. 2013).

Grazing by *P. clarkii* leads to habitat loss and modification through the removal of macrophytes, in turn leading to population declines of species that feed on macrophytes, use macrophyte stands as nesting sites and refugia from predation (Rosenthal et al. 2005). *Procambarus clarkii* leads community composition changes through trophic cascades (Gherardi & Aquistapace 2007; Souty-Grosset et al. 2016). For example, in Lake Chozas (northwest Spain), grazing by *P. clarkii* caused a reduction in macrophyte communities leading to cascades up the food chain with declines in invertebrates, amphibians, and waterfowl (Rodriguez et al 2003). Grazing can also cause changes to community composition and structure, such as changing ecosystems from macrophyte-dominated areas with clear water, to turbid phytoplankton-dominated areas (Matsizaki et al. 2009). Excessive grazing can also lead to accelerated rates of important processes such as litter breakdown and decomposition (Gherardi & Aquistapace 2007; Souty-Grosset et al. 2016).

*Procambarus clarkii* is considered an ecosystem engineer due to its ability to change ecosystems through its burrowing activities (Souty-Grosset et al. 2016). For example, its burrowing activities can cause a decrease in the water quality through bioturbation leading to increased turbidity and influx release of nutrients from sediments, often leading to algal blooms (Angeler et al. 2001; Yamamoto 2010). The impaired water quality also affects the quality of the habitats for other aquatic fauna (Angeler et al. 2001; Rodriguez et al. 2003). For example, increased turbidity can impede foraging and respiratory processes of fish (Rodriguez et al. 2003). Burrowing activities can cause structural damage to river banks and increase bank erosion, and also cause damage to water retention infrastructure such as dam walls and dykes (Souty-Grosset et al. 2016).

References:

Angeler, D.G., Sánchez-Carrillo, S., García, G. & Alvarez-Cobelas, M., 2001, 'The influence of *Procambarus clarkii* (Cambaridae, Decapoda) on water quality and sediment characteristics in a Spanish floodplain wetland', *Hydrobiologia* 464, 89-98. https://doi.org/10.1023/A:1013950129616

Aquiloni, L., Martín, M.P., Gherardi, F. & Diéguez-Uribeondo, J., 2011, 'The North American crayfish *Procambarus clarkii* is the carrier of the oomycete *Aphanomyces astaci* in Italy', *Biological Invasions* 13, 35-367. https://doi.org/10.1007/s10530-010-9828-2.

Cruz, M.J., Rebelo, R. & Crespo, E.G., 2006, 'Effects of an introduced crayfish, *Procambarus clarkii*, on the distribution of south-western Iberian amphibians in their breeding habitats', *Ecography* 29, 329-338. https://doi.org/10.1111/j.2006.0906-7590.04333.x.

Jackson, M.C., Grey, J., Miller, K., Britton, J.R. & Donohue, I., 2016, 'Dietary niche constriction when invaders meet natives: Evidence from freshwater decapods', *Journal of Animal Ecology* 85, 1098–1107. https://doi.org/10.1111/1365-2656.12533.

Ficetola, G.F., Siesa, M.E., Manenti, R., Bottoni, L., De Bernardi, F. & Padoa-Schioppa, E., 2011, 'Early assessment of the impact of alien species: Differential consequences of an invasive crayfish on adult and larval amphibians', *Diversity and Distributions* 17, 1141-1151. https://doi.org/10.1111/j.1472-4642.2011.00797.x.

Gherardi, F. & Acquistapace, P., 2007, 'Invasive crayfish in Europe: The impact of *Procambarus clarkii* on the littoral community of a Mediterranean lake', *Freshwater Biology* 52, 1249-59. https://doi.org/10.1111/j.1365-2427.2007.01760.x.

Gherardi, F. & Barbaresi, S., 2008, 'Feeding opportunism of the red swamp crayfish *Procambarus clarkia*, an invasive species', *Freshwater crayfish* 16, 77–85.

Holdich, D.M., Reynolds, J.D. & Sibley, P.J., 2009, 'A review of the ever increasing threat to European crayfish from non-indigenous crayfish species', *Knowledge and Management of Aquatic Ecosystems* 11, 394-395. https://doi.org/10.1051/kmae/2009025.

Lodge, D.M., Deines, A., Gherardi, F., Yeo, D.C., Arcella, T., Baldridge, A.K., et al., 2012, 'Global introductions of crayfishes: evaluating the impact of species invasions on ecosystem services', *Annual Review of Ecology, Evolution, and Systematics* 43, 449-472. https://doi.org/10.1146/annurev-ecolsys-111511-103919.

Longshaw, M. 2011, 'Diseases of crayfish: A review', *Journal of Invertebrate Pathology* 106, 54-70. https://doi.org/10.1016/j.jip.2010.09.013.

Matsuzaki, S.S., Usio, N., Takamura, N. & Washitani, I., 2009, 'Contrasting impacts of invasive engineers on freshwater ecosystems: An experiment and meta-analysis', *Oecologia* 158, 673-686. https://doi.org/10.1007/s00442-008-1180-1.

McMahon, T.A., Brannelly, L.A., Chatfield, M.W., Johnson, P.T., Joseph, M.B., McKenzie, V.J., et al., 2013, 'Chytrid fungus *Batrachochytrium dendrobatidis* has non-amphibian hosts and releases chemicals that cause pathology in the absence of infection', *Proceedings of the National Academy of Sciences of the United States of America* 110, 210–215. https://doi.org/10.1073/pnas.1200592110.

Rodríguez, C.F., Bécares, E. & Fernández-Aláez, M., 2003, 'Shift from clear to turbid phase in Lake Chozas (NW Spain) due to the introduction of American red swamp crayfish (*Procambarus clarkii*)', *Hydrobiologia* 506, 421-26. https://doi.org/10.1023/B:HYDR.0000008626.07042.87.

Rosenthal, S.K., Lodge, D.M., Mavuti, K.M., Muohi, W., Ochieng, P., Mungai, B.N., et al., 2005, 'Comparing macrophyte herbivory by introduced Louisiana crayfish (*Procambarus clarkii*) (Crustacea: Cambaridea) and native Dytiscid beetles (*Cybister tripunctatus*) (Coleoptera: Dytiscidae), in Kenya', *African Journal of Aquatic Science* 30, 157-62. https://doi.org/10.2989/16085910509503850.

Souty-Grosset, C., Anastácio, P.M., Aquiloni, L., Banha, F., Choquer, J., Chucholl, C., et al. 2016. 'The red swamp crayfish *Procambarus clarkii* in Europe: Impacts on aquatic ecosystems and human well-being', *Limnologica* 58, 78-93. https://doi.org/10.1016/j.limno.2016.03.003.

Yamamoto, Y., 2010, 'Contribution of bioturbation by the red swamp crayfish *Procambarus clarkii* to the recruitment of bloom-forming cyanobacteria from sediment', *Journal of Limnology* 69, 102- 111. https://doi.org/10.4081/jlimnol.2010.102

| **CON4 Closely related species' socio-economic impact** | |
|---|---|
| Response: Moderate (MO) | Confidence: Medium |

Rationale: There are no recorded socio-economic impacts for *P. virginalis*, therefore *Procambarus clarkii* was selected as a closely related taxon to estimate the environmental impacts that may arise from introductions of *P. virginalis*.

*Procambarus clarkii* often inhabits agricultural fields in which burrowing activities cause damage to infrastructure (i.e., irrigation canals and dam walls) (Arce & Diéguez-Uribeondo 2015; Lodge et al. 2012), and alters soil hydrology leading to water loss. Grazing causes crop damage and reduces crop yield (Anastácio et al. 2005; Arce & Diéguez-Uribeondo 2015). For example in Europe, *P. clarkii* affects rice production through water loss, damage to rice field banks and ditches, direct consumption of rice seed and plants, and clogging of pipes (Anastácio et al. 2005; Souty-Grosset et al. 2016). *Procambarus clarkii* also damages gill nets and spoils fish caught in nets (Gherardi et al. 2011).

*Procambarus clarkii* bio-accumulates toxins and metals from the environment (Alcorlo et al. 2016; Gherardi et al. 2002; Souty-Grosset et al. 2016), which are often harmful. These toxins and metals can be transferred to higher food web levels through the consumption of affected crayfish by humans and predators such as otters, birds, and fish (Anda et al. 2001; Gherardi et al. 2011; Lodge et al. 2012). *Procambarus clarkii* is a vector for several parasites and diseases including the parasitic fluke flatworms (*Paragonimus* spp.) that causes lung fluke disease in humans, tularemia-causing bacterium *Francisella tularensis*, rat lungworm *Angiostrongylus cantonensis* that causes meningitis, and the nematode *Gnathostoma spinigerum* that causes human gnathostomiasis (Edgerton et al. 2002; Lane et al 2009; Souty-Grosset et al. 2016).

References:

Anastácio, P.M., Correia, A.M. & Menino, J.P., 2005, 'Processes and patterns of plant destruction by crayfish: effects of crayfish size and developmental stages of rice', *Archiv für Hydrobiologie* 162, 37-51. https://doi.org/10.1127/0003-9136/2005/0162-0037.

Alcorlo, P., Otero, M., Crehuet, M., Baltanás, A. & Montes, C., 2006, 'The use of the red swamp crayfish (*Procambarus clarkii* Girard) as indicator of the bioavailability of heavy metals in environmental monitoring in the River Guadiamar (SW Spain)', *Science of the Total Environment* 366, 380-390. https://doi.org/10.1016/j.scitotenv.2006.02.023

Anda, P., Segura del Pozo, J., Díaz García, J.M., Escudero, R., García Peña, F.J., López Velasco, M.C., et al., 2001, 'Waterborne outbreak of tularemia associated with crayfish fishing', *Emerging Infectious Diseases* 7, 575-82. https://doi.org/10.3201/eid0707.010740.

Arce, J.A. & Diéguez-Uribeondo, J., 2015, 'Structural damage caused by the invasive crayfish *Procambarus clarkii* (Girard, 1852) in rice fields of the Iberian Peninsula: A study case', *Fundamental and Applied Limnology* 186, 259-269. https://doi.org/10.1127/fal/2015/0715.

Edgerton, B.F., Evans, L.H., Stephens, F.J. & Overstreet, R.M., 2002, 'Synopsis of freshwater crayfish diseases and commensal organisms', *Aquaculture* 206, 57–135. https://doi.org/10.1016/S0044-8486(01)00865-1 .

Gherardi, F., Barbaresi, S., Vaselli, O. & Bencini, A., 2002, 'A comparison of trace metal accumulation in indigenous and alien freshwater macro-decapods', *Marine and Freshwater Behaviour and Physiology* 35, 179-88. https://doi.org/10.1080/1023624021000014761.

Gherardi, F., Robert, F.J., Kenneth, B.B., Mavuti, M., Pacini, N., Grey, J., et al., 2011, 'A review of allodiversity in Lake Naivasha, Kenya: Developing conservation actions to protect East African lakes from the negative impacts of alien species', *Biological Conservation* 144, 2585–2596. https://doi.org/10.1016/j.biocon.2011.07.020.

Lane, M.A., Barsanti, M.C., Santos, C.A., Yeung, M., Lubner, S.J. & Weil, G.J., 2009, 'Human paragonimiasis in North America following ingestion of raw crayfish', *Clinical and Infectious Diseases* 49, 55-61. https://doi.org/10.1086/605534.

Lodge, D.M., Deines, A., Gherardi, F., Yeo, D.C., Arcella, T., Baldridge, A.K., et al., 2012, 'Global introductions of crayfishes: evaluating the impact of species invasions on ecosystem services', *Annual Review of Ecology, Evolution, and Systematics* 43, 449-472. https://doi.org/10.1146/annurev-ecolsys-111511-103919.

Souty-Grosset, C., Anastácio, P.M., Aquiloni, L., Banha, F., Choquer, J., Chucholl, C., et al. 2016. 'The red swamp crayfish *Procambarus clarkii* in Europe: Impacts on aquatic ecosystems and human well-being', *Limnologica* 58, 78-93. https://doi.org/10.1016/j.limno.2016.03.003.

**CON5 Potential impact**

| Response: Major (MR) | Confidence: High |
|---|---|

Rationale:

*Procambarus virginalis*, is a highly sought after aquarium pet species, due to its small size reproduce via parthogenisis making it easy to rear. *Procambarus virginalis* is in the same genus as *Procambarus clarkii*, it is therefore highly likely for it to have detrimental impacts in areas where it manages to establish populations. It is clear from the assessment that *P. clarkii* can becomes invasive in areas of introduction. It can tolerate a wide range of environmental conditions (Nunes et al. 2017b), and it is an aggressive competitor that can out-compete and displace native species in invaded ecosystems (Gherardi et al. 2011, Jackson et al. 2016). It is thereore highly likely that *Procambarus virginalis* can have similar impacts in its invaded range.

References:

Chucholl C, Pfeiffer M. 2010. First evidence for an established Marmorkrebs (Decapoda, Astacida, Cambaridae) population in Southwestern Germany, in syntopic occurrence with *Orconectes limosus* (Rafinesque, 1817). *Aquatic invasions* 5, 405-412. https://doi.org/10.3391/ai.2010.5.4.10

Feria, T.P. & Faulkes, Z., 2011, 'Forecasting the distribution of Marmorkrebs, a parthenogenetic crayfish with high invasive potential, in Madagascar, Europe, and North America*', Aquatic Invasions* 6, 55–67. https://doi.org/10.3391/ai.2011.6.1.07.

Gherardi, F., Britton, J.R., Mavuti, K.M., Pacini, N., Grey, J., Tricarico, E. & Harper, D.M., 2011, 'A review of allodiversity in Lake Naivasha, Kenya: Developing conservation actions to protect East African lakes from the negative impacts of alien species', *Biological Conservation,* 144, 2585-2596. https://doi.org/10.1016/j.biocon.2011.07.020.

Jackson, M.C., Grey, J., Miller, K., Britton, J.R., Donohue, I., 2016, 'Dietary niche constriction when invaders meet natives: Evidence from freshwater decapods', *Journal of Animal Ecology*, 85, 1098–1107, https://doi.org/10.1111/1365-2656.12533 .

Nunes AL, Zengeya TA, Measey GJ, Weyl OLF. 2017. Freshwater crayfish invasions in South Africa: Past, present and potential future. *African Journal of Aquatic Science* 42, 309-323. https://doi.org/10.2989/16085914.2017.1405788.

Soes M, Koese, B. 2010. Invasive freshwater crayfish in the Netherlands: a preliminary risk analysis. Invasive freshwater crayfish in the Netherlands: a preliminary risk analysis

#### 4. Management

**MAN1 What is the feasibility to stop future immigration?**

| Response: High | Confidence: Low |
|---|---|

Rationale: *Procambarus virginalis* is a common species in the global pet trade and aquaculture industry (Faulkes 2015; Olden & Carvalho 2024). The species is absent in neighbouring countries (Madzivanzira et al. 2020), so

| |
|---|
| unaided immigration is unlikely. Moreover, it should be possible to regulate intentional introductions legislative regulations and having an effective border surveillance and monitoring strategy (Zengeya & Wilson 2020; Wilson & Kumschick 2024). |
| References:<br>Faulkes Z. 2015. The global trade in crayfish as pets. *Crustacean Research* 44: 75-92. https://doi.org/10.18353/crustacea.44.0_75.<br>Madzivanzira, T.C., South, J., Wood, L.E., Nunes, A.L. & Weyl, O.L.F., 2020, 'A review of freshwater crayfish introductions in Africa', Reviews in Fisheries Science & Aquaculture, 1-21. https://doi.org/10.1080/23308249.2020.1802405.<br>Nunes, A.L., Zengeya, T.A., Measey, G.J. & Weyl, O.L.F., 2017, 'Freshwater crayfish invasions in South Africa: past, present and potential future', African Journal of Aquatic Science 42, 309–323, https://doi.org/10.2989/16085914.2017.1405788.<br>Olden, J.D., Carvalho, F.A. C., 2024, 'Global invasion and biosecurity risk from the online trade in ornamental crayfish, *Conservation Biology*, 38, e14359. https://doi.org/10.1111/cobi.14359<br>Wilson, J. R., & Kumschick, S., 2024, 'The regulation of alien species in South Africa', *South African Journal of Science*, 120(5/6), https://doi.org/10.17159/sajs.2024/17002.<br>Zengeya, T.A. & Wilson, J.R. (eds.), 2020, 'The status of biological invasions and their management in South Africa in 2019', South African National Biodiversity Institute, Kirstenbosch and DSI-NRF Centre of Excellence for Invasion Biology, Stellenbosch, http://dx.doi.org/10.5281/zenodo.3947613. |

| **MAN2 Benefits of the Taxon** | |
|---|---|
| **MAN2a Socio-economic benefits of the *Taxon*** | |
| Response: Low | Confidence: Low |
| Rationale: *Procambarus virginalis* has been introduced into areas outside its native range mainly for pet trade and aquaculture (Faulkes 2015; Olden & Carvalho 2024). No socio-economic benefits have been recorded to date in South Africa are even if *Procambarus virginalis* is present the benefits are likely to be low (Nunes et al. 2017; Madzivanzira et al. 2021). | |
| References:<br><br>Faulkes Z. 2015. The global trade in crayfish as pets. *Crustacean Research* 44: 75-92. https://doi.org/10.18353/crustacea.44.0_75.<br>Madzivanzira, T.C., South, J., Wood, L.E., Nunes, A.L. & Weyl, O.L.F., 2020, 'A review of freshwater crayfish introductions in Africa', Reviews in Fisheries Science & Aquaculture, 1-21. https://doi.org/10.1080/23308249.2020.1802405.<br>Nunes, A.L., Zengeya, T.A., Measey, G.J. & Weyl, O.L.F., 2017, 'Freshwater crayfish invasions in South Africa: past, present and potential future', African Journal of Aquatic Science 42, 309–323, https://doi.org/10.2989/16085914.2017.1405788.<br>Olden, J.D., Carvalho, F.A. C., 2024, 'Global invasion and biosecurity risk from the online trade in ornamental crayfish, *Conservation Biology*, 38, e14359. https://doi.org/10.1111/cobi.14359. | |
| **MAN2b Environmental benefits of the *Taxon*** | |
| Response: None | Confidence: |
| Rationale: There are no known environmental benefits from this taxon | |
| References: | |

| **MAN3 Ease of management** | |
|---|---|
| **MAN3a How accessible are populations?** | |
| Response: 2 (difficult to access) | Confidence: Low |

| |
|---|
| Rationale: There are no locality records for *Procambarus virginalis* in South Africa (Madzivanzira et al. 2020; Nunes et al. 2017). However, given that it is an aquatic species, accessibility is likely to be difficult. |
| References:<br><br>Madzivanzira, T.C., South, J., Wood, L.E., Nunes, A.L. & Weyl, O.L.F., 2020, 'A review of freshwater crayfish introductions in Africa', *Reviews in Fisheries Science & Aquaculture*, 1-21. https://doi.org/10.1080/23308249.2020.1802405.<br>Nunes, A.L., Zengeya, T.A., Measey, G.J. & Weyl, O.L.F., 2017, 'Freshwater crayfish invasions in South Africa: past, present and potential future', *African Journal of Aquatic Science* 42, 309–323, https://doi.org/10.2989/16085914.2017.1405788 |
| **MAN3b Is detectability critically time-dependent?** |

| | |
|---|---|
| Response: 2 (difficult to access) | Confidence: Medium |

| |
|---|
| Rationale: *Procambarus virginalis* is a small burrowing freshwater species that is difficult to detect (Dorn & Volin 2009; Martin et al. 2010). |
| References:<br><br>Martin, P., Dorn, N., Kawai, T., Heiden, C. van der. & Scholtz G, 2010, 'The enigmatic Marmorkrebs (marbled crayfish) is the parthenogenetic form of *Procambarus fallax* (Hagen, 1870)', *Contributions to Zoology*, 79(3), 107-118. https://doi.org/10.1163/18759866-07903003.<br><br>Dorn, N.J. & Volin, J.C. 2009, 'Resistance of crayfish (*Procambarus* spp.) populations to wetland drying depends on species and substrate', *Journal of the North American Benthological Society*, 28(4), 766-777. https://doi.org/10.1899/08-151.1. |
| **MAN3c Time to reproduction** |

| | |
|---|---|
| Response:1 (1–3 years) | Confidence: Medium |

| |
|---|
| Rationale: *Procambarus virginalis* reach sexual maturity within a year (Seitz et al. 2005). |
| References:<br><br>Seitz, R., Vilpoux, K., Hopp, U., Harzsch, S. & Maier, G. 2005, 'Ontogeny of the Marmorkrebs (marbled crayfish): a parthenogenetic crayfish with unknown origin and phylogenetic position'. *Journal of Experimental Zoology* A, 303(5), 393-405. https://doi.org/10.1002/jez.a.143 |
| **MAN3d Propagule persistence** |

| | |
|---|---|
| Response:0 (<1 year) | Confidence: Medium |

| |
|---|
| Rationale: *Procambarus virginalis* reproduces by apomictic parthenogenesis – females produce offspring from unfertilized eggs and no males are known to exist (Scholtz et al., 2003; Martin et al., 2007). No males are known to exist (Seitz et al., 2005; Jones et al., 2009). One female is enough to start a population. Fecundity varies between 45 - 724 eggs (Sietz et al. 2005; Jones et al. 2009, Chucholl & Pfeiffer 2010). |
| References:<br><br>Chucholl, C. & Pfeiffer, M. 2010, 'First evidence for an established Marmorkrebs (Decapoda, Astacida, Cambaridae) population in Southwestern Germany, in syntopic occurrence with *Orconectes limosus* (Rafinesque, 1817)', *Aquatic Invasions*, 5(4), 405-412. https://doi.org/10.3391/ai.2010.5.4.10 |

Jones, J.P.G., Rasamy, J.R., Harvey, A., Toon, A., Birgit Oidtmann, B., Randrianarison, M.H., Raminosoa, N. & Ravoahangimalala, O.R., 2009, 'The perfect invader: a parthenogenetic crayfish poses a new threat to Madagascar's freshwater biodiversity', *Biological Invasions*, 11, 1475–1482, https://doi.org/10.1007/s10530-008-9334-y

Martin P, Kohlmann K, Scholtz G, 2007, 'The parthenogenetic Marmorkrebs (marbled crayfish) produces genetically uniform offspring', *Naturwissenschaften*, 94(10), 843-846. https://doi.org/10.1007/s00114-007-0260-0

Scholtz, G., Braband, A., Tolley, L., Reimann, A., Mittmann, B., Lukhaup, C., Steuerwald, F. & Vogt, G, 2003, 'Parthenogenesis in an outsider crayfish', *Nature*, 421(6925), 806-806. https://doi.org/10.1038/421806a

Seitz R, Vilpoux K, Hopp U, Harzsch S, Maier G, 2005. Ontogeny of the Marmorkrebs (marbled crayfish): a parthenogenetic crayfish with unknown origin and phylogenetic position. *Journal of Experimental Zoology* A, 303(5):393-405. https://doi.org/10.1002/jez.a.143

| **MAN3 Ease of management** (SUM from Table S4) | |
|---|---|
| Response: Difficult | Confidence: Medium |

Rationale: *Procambarus virginalis* is a small burrowing freshwater species that is difficult to detect, reproduces parthenogenetically, and is highly fecund (45 – 724 eggs) (Scholtz et al., 2003; Sietz et al. 2005; Martin et al., 2007; Jones et al. 2009, Chucholl & Pfeiffer 2010). It does not need to maintain a viable population as one female can start a new population.

References:

Chucholl, C. & Pfeiffer, M. 2010, 'First evidence for an established Marmorkrebs (Decapoda, Astacida, Cambaridae) population in Southwestern Germany, in syntopic occurrence with *Orconectes limosus* (Rafinesque, 1817)', *Aquatic Invasions*, 5(4), 405-412. https://doi.org/10.3391/ai.2010.5.4.10

Jones, J.P.G., Rasamy, JR, Harvey A, Toon A, Oidtmann B, Randrianarison MH, Raminosoa N. & Ravoahangimalala, O.R. 2009, 'The perfect invader: a parthenogenic crayfish poses a new threat to Madagascar's freshwater biodiversity', *Biological Invasions*, 11(6), 1475-1482. https://doi.org/10.1007/s10530-008-9334-y

Martin, P., Kohlmann, K. & Scholtz, G. 2007, 'The parthenogenetic Marmorkrebs (marbled crayfish) produces genetically uniform offspring. *Naturwissenschaften*, 94(10), 843-846. https://doi.org/10.1007/s00114-007-0260-0

Scholtz, G., Braband, A., Tolley, L., Reimann, A., Mittmann, B., Lukhaup, C., Steuerwald, F. & Vogt, G. 2003, 'Parthenogenesis in an outsider crayfish', *Nature*, 421(6925), 806-806. https://doi.org/10.1038/421806a

Seitz, R., Vilpoux, K., Hopp, U., Harzsch, S. & Maier, G. 2005, 'Ontogeny of the Marmorkrebs (marbled crayfish): a parthenogenetic crayfish with unknown origin and phylogenetic position', *Journal of Experimental Zoology A*, 303(5), 393-405. https://doi.org/10.1002/jez.a.143

| **MAN4 Has the feasibility of eradication been evaluated?** | |
|---|---|
| Response: No | Confidence: Low |

Rationale: There are no known population of *Procambarus virginalis* in South Africa, therefore eradication feasibility has not been no evaluated (Nunes et al. 2017) but see MAN5 for summary of control efforts used to manage invasive crayfish populations.

References:

Nunes, A.L., Zengeya, T.A., Measey, G.J. & Weyl, O.L.F., 2017, ‘Freshwater crayfish invasions in South Africa: past, present and potential future’, *African Journal of Aquatic Science* 42, 309–323, https://doi.org/10.2989/16085914.2017.1405788

**MAN5 Control options and monitoring approaches available for the *Taxon***

Response: Several methods can be used to monitor and control invasive crayfish populations including mechanical trapping (e.g., Barkhuizen et al. 2022; Madzivanzira et al. 2021), electrofishing (Nunes et al. 2017), use of biocides, and biological control (Aquiloni et al 2010; Stebbing et al. 2014). However, there are several issues that make management of crayfish invasions challenging (Gherardi et al. 2011). For example, the use of biocides is often effective for localised populations in dams and ponds (Peay et al. 2019) but they are non-specific and need to be applied with caution (Gherardi et al. 2011). Mechanical control methods are also effective in small discrete water bodies but are expensive and require commitment to follow on to complete eradication (Loureiro et al. 2018). In addition, alien crayfish species are regarded as potential conflict-generating species in South Africa and potential management interventions should consider the concerns of stakeholders to avoid conflict (Zengeya et al. 2017).

References:

Aquiloni, L., Brusconi, S., Cecchinelli, E., Tricarico, E., Mazza, G., Paglianti, A. & Gherardi, F. 2010, ‘Biological control of invasive populations of crayfish: the European eel (*Anguilla anguilla*) as a predator of *Procambarus clarkii*’, *Biological Invasions* 12 3817-3824. https://doi.org/10.1007/s10530-007-9094-0.

Barkhuizen, L.M, Madzivanzira, T.C., South, J., 2022, ‘Population ecology of a wild population of red swamp crayfish *Procambarus clarkii* (Girard, 1852) in the Free State Province, South Africa and implications for eradication efforts’, *BioInvasions Records* 11, 181–191. https://doi.org/10.3391/bir.2022.

Gherardi, F., Aquiloni, L., Diéguez-Uribeondo, J. & Tricarico E., 2011, ‘Managing invasive crayfish: is there a hope? *Aquatic Sciences* 73, 185–200. https://doi.org/10.1007/s00027-011-0181-z.

Loureiro, T.G, Anastácio, P.M, de Siqueira Bueno, S.L. & Araujo, P.B., 2018, ‘Management of invasive populations of the freshwater crayfish *Procambarus clarkii* (Decapoda, Cambaridae): test of a population-control method and proposal of a standard monitoring approach’, *Environmental Monitoring Assessment* 190, 559, https://doi.org/10.1007/s10661-018-6942-6.

Madzivanzira, T.C., South, J., Nhiwatiwa, T. & Weyl, O.L.F., 2021, ‘Standardisation of Australian redclaw crayfish *Cherax quadricarinatus* sampling gear in southern Africa’, *Water SA* 47, 380–384. https://hdl.handle.net/10520/ejc-waters-v47-n3-a10.

Nunes, A.L., Zengeya, T.A., Hoffman, A.C., Measey, G.J. & Weyl, O.L., 2017, ‘Distribution and establishment of the alien Australian redclaw crayfish, *Cherax quadricarinatus*, in South Africa and Swaziland’, *PeerJ* 5, 1-21. https://doi.org/10.7717/peerj.3135.

Peay, S., Johnsen, S.I, Bean, S.W., Dunn, A.M., Sandodden, R. & Edsman, L, 2019, ‘Biocide treatment of invasive signal crayfish: Successes, failures and lessons learned’, *Diversity* 11, 29. https://doi.org/10.3390/d11030029

Stebbing, P., Longshaw, M., Scott, A., 2014, ‘Review of methods for the management of nonindigenous crayfish, with particular reference to Great Britain’, *Ethology Ecology & Evolution* 26, 204-231. https://doi.org/10.1080/03949370.2014.908326.

Woodford, D.J., Ivey, P., Novoa, A., Shackleton, R., Richardson, D., Weyl, O., Van Wilgen, B. and Zengeya, T., 2017, ‘Managing conflict-generating invasive species in South Africa: Challenges and trade-offs’, *Bothalia* 47, a2160. https://doi. org/10.4102/abc.v47i2.2160.

**MAN6 Any other management considerations to highlight?** (if yes, fill in Appendix MAN6)

| | |
|---|---|
| Response | Yes / No |

**5. Calculations**

**Likelihood = Fairly probable**

| Parameter | Likelihood | Stages | Final assessment |
|---|---|---|---|
| LIK1 | 0.0027 | P(entry) = 1 | P (invasion) = 0.25 |
| LIK2 | 1 | | |
| LIK3 | 0.5 | P(establishment) = 0.5 | |
| LIK4 | 0.5 | | |
| LIK5 | 0.0027 | P (spread) = 0.5 | |
| LIK6 | 0.5 | | |

**Consequence = Major (MR)**

| Parameter | Mechanism/sector | Response |
|---|---|---|
| CON1 | Maximum environmental impact | DD |
| CON2 | Maximum socio-economic impact | DD |
| CON1a | Competition | DD |
| CON1b | Predation | DD |
| CON1c | Hybridisation | DD |
| CON1d | Disease transmission | DD |
| CON1e | Parasitism | DD |
| CON1f | Poisoning/toxicity | DD |
| CON1g | Bio-fouling or other direct physical disturbance | DD |
| CON1h | Grazing/herbivory/browsing | DD |
| CON1i | Chemical, physical, structural impact | DD |
| CON1k | Indirect impacts through interactions with other species | DD |
| CON3 | Maximum environmental impact (closely related taxa) | MR |
| CON1a | Safety | DD |
| CON1b | Material and immaterial assets | DD |
| CON1c | Health | DD |

| | | |
|---|---|---|
| CON1d | Social, spiritual and cultural relations | DD |
| CON4 | Maximum socio-economic impact (closely related taxa) | MO |
| CON5 | Potential impact based on traits, experiments, or models | MR |

**Risk score: High**

| | | Consequences | | | | |
|---|---|---|---|---|---|---|
| | | MC | MN | MO | MR | MV |
| Likelihood | Extremely unlikely | low | low | low | medium | medium |
| | Very unlikely | low | low | low | medium | high |
| | Unlikely | low | low | medium | high | high |
| | Fairly probable | medium | medium | high | high | high |
| | Probable | medium | high | high | high | high |

**Ease of management: difficult**

| Parameter | Question | Response |
|---|---|---|
| MAN3a | How accessible are populations? | 2 |
| MAN3b | Is detectability critically time-dependent? | 2 |
| MAN3c | Time to reproduction | 1 |
| MAN3d | Propagule persistence | 1 |
| **MAN3** | **SUM** | 6 |

**Appendix BAC7**: Global alien range of *Procambarus virginalis.* Map from CABI: **https://www.gbif.org/species/2227309**

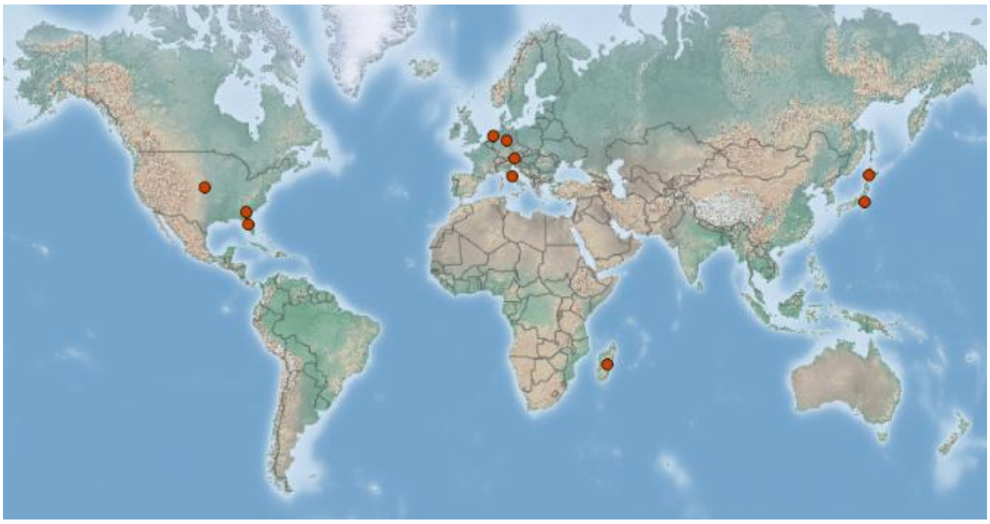
