## Supplement 2 for "Applying standardised methods to assess the potential risks of alien freshwater crayfish introductions to South Africa"

2.1 A summary of the impact assessment of alien crayfish species done using the Environmental Impact Classification of Alien Taxa (EICAT) and the relevant literature cited. Impact scores, from highest to lowest are: 1) Massive (MV); 2) Major (MR); 3) Moderate (MO), 4) Minor (MN); 5) and Minimal Concern (MC).

| Species | Impact mechanisms | Impact | Impact score | References | Region | Confidence score |
|---|---|---|---|---|---|---|
| *Cherax quadricarinatus* | Competition | Outcompete and displace native species | MC | Haubrouck et al. 2021; Madzivanzira et al. 2021; Madzivanzira et al. 2022; Zengeya et al. 2022 | Australia \| Africa | Medium |
| | Predation | Predation on invertebrates and fish | MC | Haubrouck et al. 2021; Madzivanzira et al. 2021; Madzivanzira et al. 2022; Zengeya et al. 2022 | Australia \| Africa | Medium |
| | Transmission of diseases | Transmit diseases to native species | MR | Haubrock et al. 2021 | Asia | Medium |
| | Poisoning/toxicity | Bioaccumulate toxins that can be transmitted up the food webs | MC | Haubrouck et al. 2021 | Australia | High |
| | Grazing | Reduced macrophyte abundance and diversity | MC | Haubrouck et al. 2021, Madzivanzira et al. 2021; Madzivanzira et al. 2022, Zengeya et al. 2022 | Australia \| Africa | |
| | Structural changes | Changes to ecosystem structure and functioning | MC | Haubrock et al. 2021; Marufu et al. 2018; Robinet 2010; Todd and D`Andrea 2003; Zengeya et al. 2022 | Africa \| North America | Medium |
| *Faxonius limosus* | Competition | Outcompete and displace native species | MR | Holdich et al. 2006, Holdich and Black 2007; Kozák et al. 2007; Musil et al. 2010 | Europe | Medium |
| | Predation | Decline in native species population of invertebrates | MO | Vojkovská et al. 2014; Šidagytė et al. 2017 | Europe | Low |
| | Transmission of diseases | Transmit diseases to native species | MR | Holdich et al. 2006; Holdich and Black 2007; Kozák et al. 2007; Musil et al. 2010; Pârvulescu et al. 2012 | Europe | Medium |
| | Grazing | Decrease in macrophyte cover due to excessive grazing | MN | Vojkovská et al. 2014; Šidagytė et al. 2017 | Europe | Low |

| Species | Impact mechanisms | Impact | Impact score | References | Region | Confidence score |
|---|---|---|---|---|---|---|
| | Structural changes | Burrowing activity causes physical damage to riverbanks and impacts water quality through bioturbation | MN | Starzner et al. 2000; Aldridge 2011 | Europe | Low |
| *Faxonius immunis* | Competition | Outcompete native species | MO | Jansen et al. 2009; EU 2022 | Europe \| North America | Low |
| | Predation | Decline in native species population of fish and invertebrates | MR | Chucholl 2012; EU 2022 | Europe | Medium |
| | Transmission of diseases | Transmit diseases to native species | MN | Schrimpf et al. 2013 | Europe | Low |
| | Physical disturbance | Burrowing activity causes physical damage to riverbanks and impacts water quality through increased erosion | MN | Souty-Grosset et al. 2016 | Europe | Low |
| | Grazing | Reduction in macrophyte cover, macroinvertebrates abundance and diversity | MN | EU 2022 | Europe | |
| *Faxonius rusticus* | Competition | Outcompete native crayfish – decline in abundance | MO | Bobeldyk and Lamberti 2008 | North America | Medium |
| | | Competitive exclusion | MO | Hill and Lodge 1994 | North America | Medium |
| | | Outcompete native species | MO | Garvey and Stein 1993 | North America | Medium |
| | | Displacing native congeners | MO | Taylor and Redmer 1996 | North America | Medium |
| | | Competitive exclusion | MO | Bergman and Moore 2003 | North America | Medium |
| | Predation | Decline in snail diversity | MR | Kreps et al. 2012 | North America | High |
| | | Decline in abundance of native fish species | MO | Wilson et al. 2004, Jonas et al. 2005 | North America | High |
| | | Decrease in invertebrate diversity and abundance | MO | Klocker & Strayer 2004; Kreps et al. 2012; Lodge et al. 2005; Wilson et al. 2004 | North America | High |
| | | Predate on eggs- reproductive interference | MN | Baldrige and Lodge 2013 | North America | Medium |
| | Hybridisation | Hybridise with native species | MN | Alcella et al. 2014; Perry et al. 2002 | North America | Medium |
| | Transmission of disease | Transmit disease to native species | MN | Panteleit et al. 2019; Butler et al. 2020, Stratton et al. 2023 | North America | |
| | Grazing | Reduced macrophyte abundance and diversity | MR | Wilson et al. 2004 | North America | High |
| | | Reduced macrophyte abundance | MO | Roth et al. 2006 | North America | Medium |

| Species | Impact mechanisms | Impact | Impact score | References | Region | Confidence score |
|---|---|---|---|---|---|---|
| | Indirect impacts through interactions with other species | Interactions with other these species facilitate negative impacts native fauna | MO | Johnson et al. 2009 | North America | Medium |
| *Faxonius virilis* | Competition | Competition - decline, displacement and extirpation of native populations of invertebrates, crayfish, frogs and fish | MR | Hanson et al. 1990; Rogowski and Stockwell 2006; Philips et al. 2009; Loughman and Welsh 2010; Shaw et al. 2021 | North America | Medium |
| | Predation | Predation - decline, displacement and extirpation of native populations of invertebrates, crayfish, frogs and fish | MR | Hanson et al. 1990; Rogowski and Stockwell 2006; Philips et al. 2009; Loughman and Welsh 2010; Shaw et al. 2021 | North America | Medium |
| | Transmission of disease | Transmit disease to native species | MN | Ahern et al. 2008, Longshaw 2011 | Europe | Medium |
| | Grazing | Decrease in macrophyte cover due to excessive grazing | MO | Chambers et al. 1990; Roessink et al. 2017 | North America \| Europe | Low |
| | Structural changes | Habitat alteration due to decrease in macrophyte cover due to excessive grazing | MN | Chambers et al. 1990; Roessink et al. 2017 | North America \| Europe | Low |
| *Pacifastacus leniusculus* | Competition | Displacing native crayfish | MR | Almeida et al. 2014 | Europe | High |
| | | Competitive exclusion | MR | Westman et al. 2002; Dana et al. 2010, Huber and Schubart 2005; Pockl and Pekny 2002 | Europe | High |
| | | Displacing native crayfish species, competitive exclusion | MO | Dunn et al. 2009 | Europe | Medium |
| | Predation | Decline in abundance of invertebrates | MR | Crawford et al. 2006; Mathers et al. 2016 | Europe | High |
| | | Decline in native species | MR | Dana et al. 2010 | Europe | Low |
| | | Decline in newt numbers | MO | Girdner et al. 2018 | North America | High |
| | | Decline in abundance of molluscs | MO | Meira et al. 2019 | Europe | High |
| | | Decline in mussel numbers | MO | Sousa et al. 2019 | Europe | High |
| | | Decline in invertebrate richness and abundance | MO | Galib et al. 2021 | Europe | High |

| Species | Impact mechanisms | Impact | Impact score | References | Region | Confidence score |
|---|---|---|---|---|---|---|
| *Pacifastacus leniusculus* | | Affect salmonid recruitment | MN | Peay et al. 2009 | Europe | Medium |
| | Transmission of diseases | Transmission of crayfish plague led to decline in numbers and local extinction of native crayfish | MR | Chucholl and Schrimpf 2016 | Europe | High |
| | | Local disappearance of native crayfish | MR | Almeida et al. 2014 | Europe | Medium |
| | | Transmitted diseases to native species | MR | Weinlader and Furer 2009 | Europe | Medium |
| | | Transmitted diseases to native crayfish and crabs | MN | Svaboda et al. 2014 | Europe | Low |
| | Structural changes | Burrowing has led to the collapse of riverbanks | MO | Guan et al. 1994 | Europe | Medium |
| | | Burrowing activities damage riverbanks and increase erosion | MO | Holdrich et al. 2009 | Europe | Medium |
| *Procambarus clarkii* | Competition | Decrease in abundance and distribution range of native species | MR | Cruz et al. 2006 | Europe | Medium |
| | | Niche constriction and decrease in abundance of native crabs | MR | Jackson et al. 2016 | Africa | Medium |
| | Predation | Reproduction interference by excluding amphibians from breeding sites | MR | Cruz and Rebelo 2005 | Europe | High |
| | | Displacement and extirpation of native species | MR | Souty-Grosset et al. 2016 | Europe | High |
| | | Led to collapse of amphibian populations in invaded areas | MR | Cruz et al. 2008 | Europe | High |
| | | Displaced newts in areas of introduction | MR | Gamradt and Katz 1996 | North America | High |
| | | Reduced the abundance of amphibian populations in invaded areas | MO | Cruz et al. 2006 | Europe | High |
| | | Reduced the abundance of amphibian populations in invaded areas | MO | Ficetola et al. 2011 | Europe | Medium |
| | | Reduced the abundance of invertebrates in invaded areas | MN | Bucciarelli et al. 2019; Meira et al. 2020 | Europe | High |
| | | Predates on native amphibians | MN | Banci et al. 2013 | South America | High |
| | Transmission of diseases | Transmission of disease to native species leading population declines | MR | Gherardi 2010 ; Souty-Grosset et al. 2016 | Europe | Medium |

| Species | Impact mechanisms | Impact | Impact score | References | Region | Confidence score |
|---|---|---|---|---|---|---|
| *Procambarus clarkii* | Poisoning/toxicity | Bioaccumulate toxins that can be amplifies across the food web | MC | Gherardi et al. 2011; Souty-Grosset et al. 2016 | Africa \| Europe | Low |
| | Grazing | Habitat loss and modification through the removal of macrophytes leading decline in native species populations | MO | Smart et al. 2002; Rodríguez et al. 2003 ; Gherardi and Aquistapace 2007; Souty-Grosset et al. 2016 | Africa \| Europe | medium |
| | Structural impact on ecosystem | Change water from clear to turbid | MN | Rodriguez et al. 2003 | Europe | High |
| | | Burrowing may reduce levee stability | MO | Acre and Diéguez Uribeondo 2015 | Europe | Medium |
| | | Burrowing damage dam walls and irrigation structures | MN | Correia and Fereira 2005 | Europe | Medium |
| | | Burrowing may reduce levee stability | MN | Haubrock et al. 2019 | Europe | Low |
| | Indirect impacts through interactions with other species | Facilitated dispersal by other alien species into new areas | MC | Southy-Grosset et al. 2016 ; Lovas-Kiss et al. 2018 | Europe | Medium |

### Literature cited for EICAT assessment of freshwater crayfish

Acre, J.A. & Diéguez-Uribeondo, J., 2015, ′Structural damage caused by the invasive crayfish *Procambarus clarkii* (Girard, 1852) in rice fields of the Iberian Peninsula: a study case′ *Fundamental of Applied Limnology,* 186, 259–269, https://doi.org/10.1127/fal/2015/0715.

Ahern, D., England, J. & Ellis, A., 2008, 'The virile crayfish, *Orconectes virilis* (Hagen, 1870) (Crustacea: Decapoda: Cambaridae), identified in the UK. Invasive Species in Inland Waters of Europe and North America: Distribution and Impacts', *Aquatic Invasions*, 3(1), 102-104. https://doi.org/10.3391/ai.2008.3.1.18.

Aldridge, D., 2011. Spinycheek Crayfish, *Orconectes limosus*. Sand Hutton, UK: GB Non-native Species Secretariat. http://www.nonnativespecies.org/factsheet/factsheet.cfm?speciesId=2441 .

Almeida, D., Ellis, A., England, J. & Copp, G.H., 2014, "Time series analysis of native and non-native crayfish dynamics in the Thames River Basin (south-eastern England)", *Aquatic Conservation: Marine and Freshwater Ecosystems* 24, 192–202, https://doi.org/10.1002/aqc.2366.

Arcella, T.E., Perry, W.L., Lodge, D.M. & Feder, J.L., 2014, 'The role of hybridization in a species invasion and extirpation of resident fauna: hybrid vigor and breakdown in the rusty crayfish, *Orconectes rusticus', Journal of Crustacean Biology* 34, 157–164, https://doi.org/10.1163/1937240X-00002204.

Baldridge, A.K. & Lodge, D.M., 2013, 'Intraguild predation between spawning smallmouth bass (*Micropterus dolomieu*) and nest-raiding crayfish (*Orconectes rusticus*): implications for bass nesting success' *Freshwater Biology*, 58, 2355–2365, https://doi.org/10.1111/fwb.12215.

Banci, K.R.S., Viera, N.F.T., Marinho, P.S., Calixto, P.O. & Marques, O.A.V., 2013, 'Predation of *Rhinella ornata* (Anura, Bufonidae) by the alien crayish (Crustacea, Astacidae) *Procambarus clarkii* (Girard, 1852) in São Paulo, Brazil', *Herpetology Notes* 6, 339–341.

Bergman, D.A. & Moore, P.A., 2003, 'Field Observations of Intraspecific Agonistic Behavior of Two Crayfish Species, Orconectes rusticus and Orconectes virilis, in Different Habitats', *Biology Bulletin*, 205: 26–35.

Bobeldyk, A.M. & Lamberti, G.A., 2008. 'A decade after invasion: Evaluating the continuing effects of rusty crayfish on a Michigan River. *Journal of Great Lakes Research*, 34, 265-275, https://doi.org/10.3394/0380-1330(2008)34[265:ADAIET]2.0.CO;2.

Bucciarelli, G.M., Suh, D., Lamb, AD, Roberts D, Sharpton D, Shaffer HB, Fisher, R.N. and Kats, L.B., 2019. Assessing effects of non-native crayfish on mosquito survival. *Conservation Biology 33*:122–131, https://doi.org/10.1111/cobi.13198.

Butler, E., P. Crigler, G. Robbins, and J. E. Blair. 2020. Preliminary survey of *Aphanomyces* sp. associated with native and invasive crayfish in the lower Susquehanna watershed of south central Pennsylvania. *Journal of Freshwater Ecology* 35(1), 223-233, https://doi.org/10.1080/02705060.2020.1779141.

Chambers, P.A., Hanson, J.M. & Burke, J.M., 1990, 'The impact of the crayfish *Orconectes virilis* on aquatic macrophytes', *Freshwater Biology* 24, 81–91, https://doi.org/10.1111/j.1365-2427.1990.tb00309.x .

Chucholl, C. & Schimpf, A., 2016, 'The decline of endangered stone crayfish (*Austropotamobius torrentium*) in southern Germany is related to the spread of invasive alien species and land use change', *Aquatic Conservation: Marine and Freshwater Ecosystems*, 26, 44–56, https://doi.org/10.1002/aqc.2568.

Chucholl, C., 2012, 'Understanding invasion success: life-history traits and feeding habits of the alien crayfish Orconectes immunis (Decapoda, Astacida, Cambaridae)', *Knowledge and Management of Aquatic Ecosystems* 404, 04, https://doi.org/10.1051/kmae/2011082.

Correia, A.M. & Ferreira, O., 2005, 'Burrowing behavior of the introduced red swamp crayfish procambarus clarkii (decapoda: cambaridae) in Portugal', *Journal of Crustacean Biology* 15, 248–257, https://doi.org/10.1163/193724095X00262.

Crawford, L., Yeomans, W.E. & Adams, C.E., 2006, 'The impact of introduced signal crayfish *Pacifastacus leniusculus* on stream invertebrate communities. *Aquatic Conservation: Marine and Freshwater Ecosystems*, 16, 611–621, https://doi.org/10.1002/aqc.761.

Cruz MJ, Rebelo R. 2005. Vulnerability of Southwest Iberian amphibians to an introduced crayfish, *Procambarus clarkia*. *Amphibia-Reptilia* 26:293–30.

Cruz, M.J., Rebelo, R. & Crespo, E.G., 2006, 'Effects of an introduced crayfish, *Procambarus clarkii*, on the distribution of south-western Iberian amphibians in their breeding habitats. *Ecography*, 29, 329–338, https://doi.org/10.1111/j.2006.0906-7590.04333.x.

Cruz, M.J., Segurado, P., Sousa, P.M. & Rebelo, R., 2008, 'Collapse of the amphibian community of the Paul do Boquilobo Natural Reserve (central Portugal) after the arrival of the exotic American crayfish *Procambarus clarkii', Herpetological Journal* 18, 197–204.

Dana, E.D., López-Santiago J., García-de-Lomas, J., García-Ocaña, D.M., Gámez, V. & Ortega, F., 2010, 'Long-term management of the invasive *Pacifastacus leniusculus* (Dana, 1852) in a small mountain stream', *Aquatic Invasions* 5, 317–322, https://doi.org/10.3391/ai.2010.5.3.10.

Dunn, J.C., McClymont, H.E., Christmas, M. and Dunn, A.M., 2009, 'Competition and parasitism in the native White Clawed Crayfish Austropotamobius pallipes and the invasive Signal Crayfish Pacifastacus leniusculus in the UK', *Biological Invasions*, *11*, 315–324, https://doi.org/10.1007/s10530-008-9249-7.

European Commission 2022, Directorate-General for Environment, Study on invasive alien species – Development of risk assessments to tackle priority species and enhance prevention – Final report (and annexes), Publications Office of the European Union, 2022, https://data.europa.eu/doi/10.2779/302048.

Ficetola, G.F., Siesa, M.E., Manenti, R., Bottoni, L., De Bernardi, F. & Padoa-Schioppa, E., 2011, 'Early assessment of the impact of alien species: Differential consequences of an invasive crayfish on adult and larval amphibians', *Diversity and Distributions* 17,1141–1151, https://doi.org/10.1111/j.1472-4642.2011.00797.x.

Galib, S.M., Findlay, J.S., Lucas, M.C. 2021, 'Strong impacts of signal crayfish invasion on upland stream fish and invertebrate communities', *Freshwater Biology* 66, 223–240, https://doi.org/10.1111/fwb.13631.

Gamradt, S.C. & Kats, L.B., 1996, 'Effect of introduced crayfish and mosquitofish on California newts', *Conservation Biology* 10(4), 1155–1162, https://doi.org/10.1046/j.1523-1739.1996.10041155.x.

Garvey, J.E. & Stein, R.A, 1993, 'Evaluating how chela size influences the invasion potential of an introduced crayfish (*Orconectes rusticus*)', *American Midland Naturalist* 129, 172–181, https://doi.org/10.1046/j.1523-1739.1996.10041155.x.

Gherardi, F. & Acquistapace P., 2007, 'Invasive crayfish in Europe: The impact of *Procambarus clarkii* on the littoral community of a Mediterranean lake', *Freshwater Biology* 52, 1249–59, https://doi.org/10.1111/j.1365-2427.2007.01760.x.

Gherardi, F., 2010, 'Invasive crayfish and freshwater fishes of the world', *Scientific and Technical Review of the Office International des Epizooties,* 29, 241–254, https://doi.org/10.20506/rst.29.2.1973.

Gherardi, F., Britton, J.R., Mavuti, K.M., Pacini, N., Grey, J., Tricarico, E., et al., 2011, 'A review of allodiversity in Lake Naivasha, Kenya: Developing conservation actions to protect East African lakes from the negative impacts of alien species', *Biological Conservation* 144, 2585–2596, https://doi.org/10.1016/j.biocon.2011.07.020.

Girdner, S.F., Ray, A.M., Buktenica, M.W., Hering, D.K., Mack, J.A. & Umek, J.W., 2018, 'Replacement of a unique population of newts (*Taricha granulosa mazamae*) by introduced signal crayfish (*Pacifastacus leniusculus*) in Crater Lake, Oregon', *Biological Invasions* 20, 721–740, https://doi.org/10.1007/s10530-017-1570-6.

Guan, R.Z., 1994, 'Burrowing behaviour of signal crayfish, *Pacifastacus leniusculus* (Dana) in the River Great Ouse, England', In *Freshwater Forum* 4, 155–168.

Hanson, J.M., Chambers, P.A. & Prepas, E.E., 1990, 'Selective foraging by the crayfish *Orconectes virilis* and its impact on macroinvertebrates', *Freshwater Biology*, 24, 69–80, https://doi.org/10.1111/j.1365-2427.1990.tb00308.x.

Haubrock, P.J., Inghilesi, A.F., Mazza, G., Bendoni, M., Solari, L. & Ttricarico, E., 2019, 'Burrowing activity of *Procambarus clarkii* on levees: analysing behaviour and burrow structure', *Wetlands Ecology Management*, 27, 497–511, https://doi.org/10.1007/s11273-019-09674-3.

Haubrock, P.J., Oficialdegui, F.J., Zeng, Y., Patoka, J., Yeo, D.C.J. & Kouba, A. 2021, 'The redclaw crayfish: A prominent aquaculture species with invasive potential in tropical and subtropical biodiversity hotspots', *Reviews in Aquaculture* 13, 1488–1530, https://doi.org/10.1111/raq.12531.

Hill, A.M. & Lodge, D.M., 1994, 'Diel changes in resource demand: competition and predation in species replacement among crayfishes', *Ecology*, 75, 2118–2126, https://doi.org/10.2307/1941615.

Holdich D.M., Haffner, P. & Noël, P., 2006, 'Species files', In: Souty-Grosset, C., Holdich, D.M., Noël, P.Y., Reynolds, J.D. & Haffner, P. (eds.), *Atlas of Crayfish in Europe*, Muséum national d'Histoire naturelle, Paris, Patrimoines naturels, 64, 50–129.

Holdich, D. & Black, J., 2007, 'The spiny-cheek crayfish, *Orconectes limosus* (Rafinesque, 1817) [Crustacea: Decapoda: Cambaridae], digs into the UK', *Aquatic Invasions*, 2(1), 1-16. http://dx.doi.org/10.3391/ai.2007.2.1.1.

Holdich, D.M., Haffner, P. & Noël, P., 2006, 'Species files', In: Souty-Grosset, C., Holdich, D.M., Noël, P.Y., Reynolds, J.D. & Haffner, P. (eds.), *Atlas of Crayfish in Europe*, Muséum national d'Histoire naturelle, Paris, Patrimoines naturels, 64, 50–129.

Holdich, D.M., Reynolds, J.D. & Sibley, P.J., 2009, 'A review of the ever increasing threat to European crayfish from non-indigenous crayfish species', *Knowledge and Management of Aquatic Ecosystems* 11, 394–395, https://doi.org/10.1051/kmae/2009025.

Huber, M.G. & Schubart, C.D., 2005, 'Distribution and reproductive biology of *Austropotamobius torrentium* in Bavaria and documentation of a contact zone with the alien crayfish *Pacifastacus leniusculus', Bulletin Français de la Pêche et de la Pisciculture* 376, 759–776, https://doi.org/10.1051/kmae:2005031.

Jackson, M.C., Grey, J., Miller, K., Britton, J.R. & Donohue, I., 2016, 'Dietary niche constriction when invaders meet natives: Evidence from freshwater decapods', *Journal of Animal Ecology*, 85, 1098–1107, https://doi.org/10.1111/1365-2656.12533.

Jansen, W., Geard, N., Mosindy, T., Olson, G. & Turner, M., 2009, "Relative abundance and habitat association of three crayfish (*Orconectes virilis*, *O. rusticus*, and *O. immunis*) near an invasion front of *O. rusticus*, and long-term changes in their distribution in Lake of the Woods, Canada", *Aquatic Invasions*, 4, 627–649, https://doi.org/10.1111/1365-2656.12533.

Johnson, P.T., Olden, J.D., Solomon, C.T. & Vander Zanden, M.J., 2009, 'Interactions among invaders: community and ecosystem effects of multiple invasive species in an experimental aquatic system', *Oecologia* 159, 161–170, https://doi.org/10.1007/s00442-008-1176-x.

Jonas, J.L., Claramunt, R.M., Fitzsimons, J.D., Marsden, J.E. & Ellrott, B.J., 2005, 'Estimates of egg deposition and effects of lake trout (*Salvelinus namaycush*) egg predators in three regions of the Great Lakes. *Canadian Journal of Fisheries and Aquatic Sciences* 62: 2254–2264, https://doi.org/10.1139/f05-141.

Klocker, C.A. & Strayer, D.L., 2004, 'Interactions among an invasive crayfish (*Orconectes rusticus*), a native crayfish (*Orconectes limosus*), and native bivalves (Sphaeriidae and Unionidae)', *Northeastern Naturalist*, 11, 167–178, https://doi.org/10.1656/1092-6194(2004)011[0167:IAAICO]2.0.CO;2.

Kozák, P., Buřič, M., Policar, T., Hamáčková, J., Lepičová, A., 2007, 'The effect of inter- and intra-specific competition on survival and growth rate of native juvenile noble crayfish *Astacus astacus* and alien spiny-cheek crayfish *Orconectes limosus*', *Hydrobiologia*, 59085-94, https://doi.org/10.1007/s10750-007-0760-0.

Kreps, T.A., Baldridge, A.K. & Lodge, D.M., 2012, 'The impact of an invasive predator (*Orconectes rusticus*) on freshwater snail communities: Insights on habitat-specific effects from a multilake long-term study', *Canadian Journal of Fisheries and Aquatic Sciences*, 69, 1164–1173, https://doi.org/10.1139/f2012-052.

Lodge, D.M., Kershner, M.W. & Aloi, J.E., 1995, 'Effects of an omnivorous crayfish (*Orconectes rusticus*) on a freshwater littoral food web', *Ecology* 75, 1265–1281, https://doi.org/10.2307/1937452.

Longshaw, M., 2011, 'Diseases of crayfish: A review', *Journal of Invertebrate Pathology* 106, 54–70, https://doi.org/10.1016/j.jip.2010.09.013.

Loughman, Z. J. & Welsh, S.A., 2010, 'Distribution and conservation standing of West Virginia crayfishes', *Southeastern Naturalist*, 9(3), 63–78, https://doi.org/10.1656/058.009.s304.

Lovas-Kiss, A., Sanchez, M.I., Molnar, A.V., Valls, L., Armengol, X., Mesquita-Joanes, F., et al., 2018, 'Crayfish invasion facilitates dispersal of plants and invertebrates by gulls', *Freshwater Biology* 63(4), 392–404, https://doi.org/10.1111/fwb.13080.

Madzivanzira, T.C., Weyl, O.L.F. & South, J.. 2022, 'Ecological and potential socioeconomic impacts of two globally invasive crayfish', *NeoBiota* 72, 25-43, https://doi.org/10.3897/neobiota.72.71868.

Madzivanzira, T.C., South, J., Weyl, O.L.F., 2021, 'Invasive crayfish outperform potamonautid crabs at higher temperatures', *Freshwater Biology* 66, 978-991, https://doi.org/10.1111/fwb.13691.

Marufu, L., Dalu, T., Barson, M., Simango, R., Utete, B. & Nhiwatiwa, T., 2018, 'The diet of an invasive crayfish, *Cherax quadricarinatus* (Von Martens, 1868), in Lake Kariba, inferred using stomach content and stable isotope analyses', *BioInvasions Records*, 7, 12–132, https://doi.org/10.3391/bir.2018.7.2.03.

Mathers, K.L., Chadd, R.P., Dunbar, M.J., Extence, C.A., Reeds, J., Rice, S.P. & Wood, P.J., 2016, 'The long-term effects of invasive signal crayfish (Pacifastacus leniusculus) on instream macroinvertebrate communities', *Science of the Total Environment*, *556*, 207–218, https://doi.org/10.1016/j.scitotenv.2016.01.215.

Meira, A., Lopes-Lima, M., Varandas, S., Teixeira, A., Arenas, F. & Sousa, R., 2019 'Invasive crayfishes as a threat to freshwater bivalves: Interspecific differences and conservation implications', *Science of the Total Environment*, *649*, 938–948, https://doi.org/10.1016/j.scitotenv.2018.08.341.

Musil, M., Buřič, M., Policar, T., Kouba, A. & Kozák, P., 2010, 'Comparison of diurnal and nocturnal activity between noble crayfish (*Astacus astacus*) and spinycheek crayfish (*Orconectes limosus*)', *Freshwater Crayfish*, 17189–17193.

Panteleit, J., T. Horvath, J. Jussila, J. Makkonen, W. Perry, R. Schulz, K. Theissinger, and A. Schrimpf. 2019. Invasive rusty crayfish (*Faxonius rusticus*) populations in North America are infected with the crayfish plague disease agent (*Aphanomyces astaci*). *Freshwater Science*, 38(2), 425–433, https://doi.org/10.1086/703417

Pârvulescu, L., Schrimpf, A., Kozubíková, E., Resino, S. C., Vrålstad, T., Petrusek, A. & Schulz, R., 2012, 'Invasive crayfish and crayfish plague on the move: first detection of the plague agent *Aphanomyces astaci* in the Romanian Danube', *Diseases of Aquatic Organisms*, 98(1), 85–94, https://doi.org/10.3354/dao02432.

Peay, S., Guthrie, N., Spees, J., Nilsson, E. and Bradley, P., 2009, 'The impact of signal crayfish (Pacifastacus leniusculus) on the recruitment of salmonid fish in a headwater stream in Yorkshire, England', *Knowledge and Management of Aquatic Ecosystems*, 394–395, 12, https://doi.org/10.1051/kmae/2010003.

Perry, W.L., Lodge, D.M. and Feder, J.L., 2002, 'Importance of hybridization between indigenous and nonindigenous freshwater species: an overlooked threat to North American biodiversity', *Systematic Biology*, 51, 255–275, https://doi.org/10.1080/10635150252899761.

Phillips, I.D., Vinebrooke, R.D. & Turner, M.A., 2009, 'Ecosystem consequences of potential range expansions of *Orconectes virilis* and *Orvconectes rusticus* crayfish in Canada- a review. *Environmental Review*, 17, 235–248, https://doi.org/10.1139/A09-011.

Pöckl, M. & Pekny, R., 2002. 'Interaction between native and alien species of crayfish in Austria: case studies', *Bulletin Français de la Pêche et de la Pisciculture* 367, 763–776, https://doi.org/10.1051/kmae:2002064.

Robinet, O., 2010, 'Stratégie de lutte contre les espèces invasives à la Réunion. Parc national Ile de La Réunion., France

Rodríguez, C.F., Bécares, E. & Fernández-Aláez, M., 2003, 'Shift from clear to turbid phase in Lake Chozas (NW Spain) due to the introduction of American red swamp crayfish (*Procambarus clarkii*)', *Hydrobiologia* 506, 421–26, https://doi.org/10.1023/B:HYDR.0000008626.07042.87.

Roessink, I., Gylstra, R., Heuts, P. G. M., Specken, B. & Ottburg, F., 2017, 'Impact of invasive crayfish on water quality and aquatic macrophytes in the Netherlands', *Aquatic Invasions* 12(3), 397–404, https://doi.org/10.3391/ai.2017.12.3.12.

Rogowski, D.L. & Stockwell, C.A., 2006, 'Assessment of potential impacts of exotic species on populations of a threatened species, white sands pupfish, *Cyprinodon Tularosa*', *Biological Invasions*, 8(1), 79–87, https://doi.org/10.1007/s10530-005-0238-9.

Roth, B.M., Hein, C.L. & Vander Zanden, M.J., 2006, 'Using bioenergetics and stable isotopes to assess the trophic role of rusty crayfish (*Orconectes rusticus*) in lake littoral zones', *Canadian Journal of Fisheries and Aquatic Sciences*, 63, 335–344, https://doi.org/10.1139/f05-217.

Schrimpf, A., Chucholl, C., Schmidt, T., & Schulz, R., 2013, 'Crayfish plague agent detected in populations of the invasive North American crayfish *Orconectes immunis* (Hagen, 1870) in the Rhine River, Germany', *Aquatic Invasions*, 8(1), http://dx.doi.org/10.3391/ai.2013.8.1.12.

Shaw, P.C., Larson, E.R. & Billman, E.J., 2021, 'Invasion of Virile Crayfish *Faxonius virilis* (Hagen 1870) in the Lower Henrys Fork Drainage, Idaho', *Northwest Science* 95(1), 106–113, https://doi.org/10.3955/046.095.0107.

Šidagytė, E., Razlutskij, V., Alekhnovich, A., Rybakovas, A., Moroz, M., Šniaukštaitė, V., Gintaitas, V. & Arbačiauskas, K., 2017, 'Predatory diet and potential effects of *Orconectes limosus* on river macroinvertebrate assemblages of the southeastern Baltic Sea basin: implications for ecological assessment', *Aquatic Invasions*, 12(4), https://doi.org/10.3391/ai.2017.12.4.09.

Smart, A.C., Harper, D.M., Malaisse, F., Schmitz, S., Coley, S. & de Beauregard, A-C.G., 2002, 'Feeding of the exotic Louisiana red swamp crayfish, *Procambarus clarkii* (Crustacea, Decapoda), in an African tropical lake: Lake Naivasha, Kenya', *Hydrobiologia* 488, 129–142, https://doi.org/10.1023/A:1023326530914.

Sousa, R., Nogueira, J.G., Ferreira, A., Carvalho, F., Lopes-Lima, M., Varandas, S. & Teixeira, A. A., 2019, 'Tale of shells and claws: The signal crayfish as a threat to the pearl mussel Margaritifera margaritifera in Europe', *Science of the Total Environment*, 665, 329–337, https://doi.org/10.1016/j.scitotenv.2019.02.094.

Souty-Grosset, C, Manuel, P., Aquiloni, L., Banha, F., Choquer, J., Chucholl, C., Tricarico, E., 2016, 'The red swamp crayfish *Procambarus clarkii* in Europe: Impacts on aquatic ecosystems and human well-being', *Limnologica* 58, 78–93, https://doi.org/10.1016/j.limno.2016.03.003.

Statzner, B., Fièvet, E., Champagne, J. Y., Morel, R. & Herouin, E., 2000, 'Crayfish as geomorphic agents and ecosystem engineers: biological behavior affects sand and gravel erosion in experimental streams', *Limnology and Oceanography*, 45(5), 1030-1040. https://doi.org/10.4319/lo.2000.45.5.1030.

Stratton, C. E., B. A. Kabalan, S. A. Bolds, L. S. Reisinger, D. C. Behringer, and J. Bojko. 2023. *Cambaraspora faxoni* n. sp. (Microsporidia: Glugeida) from native and invasive crayfish in the USA and a novel host of *Cambaraspora floridanus*. *Journal of Invertebrate Pathology* 199:107949. https://doi.org/10.1016/j.jip.2023.107949.

Svoboda, J., Strand, D.A., Vralstad, T., Grandjean, F., Lennart, L., Kozak, P., Kouba, A., Fristad, R.F., Koca, S.B. & Petrusek, A., 2014, 'The crayfish plague pathogen can infect freshwater inhabiting crabs', *Freshwater Biology* 59: 918–929, https://doi.org/10.1111/fwb.12315.

Taylor, C.A. & Redmer, M., 1996, 'Dispersal of the crayfish orconectes rusticus in Illinois, with notes on species displacement and habitat preference', *Journal of Crustacean Biology*, 16: 547–551, https://doi.org/10.1163/193724096X00577.

Todd, S. & D'Andrea, M., 2003, 'Alien crayfish invasion of Jamaican rivers', *Crayfish News* 25, 17–18.

Vojkovská, R., Horká, I. & Ďuriš, Z., 2014, 'The diet of the spiny-cheek crayfish *Orconectes limosus* in the Czech Republic', *Central European Journal of Biology*, 9(1), 58–69, https://doi.org/10.2478/s11535-013-0189-y.

Weinländer, M. & Füreder, L., 2009, 'The continuing spread of Pacifastacus leniusculus in Carinthia (Austria)', *Knowledge and Management of Aquatic Ecosystems*, 394–395, 17, https://doi.org/10.1051/kmae/20010011.

Westman, K., Savolainen, R. & Julkunen, M., 2002, 'Replacement of the native crayfish *Astacus astacus* by the introduced species *Pacifastacus leniusculus* in a small, enclosed Finnish lake: A 30-year study', *Ecography* 25, 53–73, https://doi.org/10.1034/j.1600-0587.2002.250107.x.

Wilson, K.A., Magnuson, J.J., Lodge, D.M., Hill, A.M., Kratz, T.K., Perry, W.L. & Willis, T.V., 2004, 'A long-term rusty crayfish (Orconectes rusticus) invasion: dispersal patterns and community change in a north temperate lake', *Canadian Journal of Fisheries and Aquatic Sciences*, 61, 2255– 2266, https://doi.org/10.1139/f04-170.

Zengeya, T.A., Lombard, R.J.H., Nelwamondo, V.E., Nunes, A.L., Measey, J.G., Weyl, O.L.F., 2022, 'Trophic niche of an invasive generalist consumer: Australian redclaw crayfish, *Cherax quadricarinatus*, in the Inkomati River Basin, South Africa', *Austral Ecology*, 47, 1480–1494, https://doi.org/10.1111/aec.13230.

**2.2** A summary of the impact assessment using the Socio-Economic Impact Classification for Alien Taxa (SEICAT) and the relevant literature cited. Impact scores, from highest to lowest are: 1) Massive (MV); 2) Major (MR); 3) Moderate (MO); 4) Minor (MN); and 5) Minimal Concern (MC).

| Species | Constituent of human well-being | Activity | Impact score | Reference | Region | Confidence score |
|---|---|---|---|---|---|---|
| *Cherax quadricarinatus* | Material and immaterial assets | Supplanted native crayfish species through competition and predation | MO | Vega-Villasante et al. 2015; Haubrock et al. 2021 | North America | Medium |
| | | Damaged fishing gear | MO | Douthwaite et al. 2018 ; Haubrock et al. 2021; Madzivanzira et al. 2022; Chakandinakira et al. 2023; Madzivanzira et al 2023 | Africa | Medium |
| *Faxonius immunis* | Material and immaterial assets | Considered an agricultural pest due to burrowing and grazing activity | MN | Souty-Grosset et al. 2016 | Europe | Low |
| *Faxonius limosus* | Material and immaterial assets | Led to the collapse of crayfish fisheries and loss of livelihood for local communities | MO | Laurent 1998 | Europe | Medium |
| | | Burrowing activity increase bank erosion and damage to nearby infrastructure | MO | Aldridge 2011 | Europe | Medium |
| | | Predation on native fish disrupts angling activities | MO | Souty-Grosset et al. 2006 | Europe | Medium |
| | Social, spiritual and cultural relations | Led to the collapse of crayfish fisheries and loss of livelihood for local communities | MO | Laurent 1998; Aldridge 2011; Souty-Grosset et al. 2016 | Europe | Medium |
| *Faxonius rusticus* | Material and immaterial assets | Reduced population of native fish that are a target of sport fishing | MO | Keller et al. 2008 | North America | Low |
| *Faxonius virilis* | Material and immaterial assets | Burrowing activity increase bank erosion and damage to nearby infrastructure | MN | GISD 2023; Nemesis 2023 | North America | Low |
| *Pacifastacus leniusculus* | Material and immaterial assets | Supplanted native species that are used for aquaculture | MO | Holdich et al. 2009, Kouba et al. 2022 | Europe | Low |
| | Social, spiritual and cultural activities | Reduced population of native fish that are a target of sport fishing | MN | Holdich et al. 2009; Gherardi 2011 | Europe | Low |
| *Procambarus clarkii* | Material and immaterial | Damage to rice fields | MO | Gherardi et al. 2011 | Europe | Medium |

| Species | Constituent of human well-being | Activity | Impact score | Reference | Region | Confidence score |
|---|---|---|---|---|---|---|
| | | Burrowing and grazing activities decreased rice production and clog pipes | MO | Gherardi et al. 2011 | Europe | Medium |
| | | Decreased rice production | MN | Anastácio et al. 2005 | Europe | Medium |
| | | Affected fishing industry by damaging gill nets and spoiling the fish caught in the nets | MO | Gherardi et al. 2011 | Africa | Medium |
| | | Burrowing and grazing activities decreased rice production | MO | Sara and Moutaouakil 2019 | Africa | Medium |
| | Health | Transmission of disease to native species | MO | Lane et al. 2009 | North America | High |
| | | Transmission of disease to native species | MO | Anda et al. 2011 | Europe | High |
| | | *Procambarus clarkii* serves as vector for several parasites and diseases | MN | Putra et al. 2018 | Asia | Low |
| | | It accumulates cyanobacteria toxins and heavy metals that can be transferred to its consumers, above all birds but also humans included | MN | Gherardi et al. 2011 | Europe | Low |
| | | Transmission of diseases to native species | MO | Souty-Grosset et al. 2016 | Europe | Low |
| | | Pinching caused injuries to locals in invaded sites | MN | Sara and El Moutaouakil 2019 | Africa | High |
| | Social, spiritual and cultural activities | Disrupt recreational activities such as angling | MO | Gherardi et al. 2011 | Europe | Low |

**Literature cited for SEICAT assessment of freshwater crayfish**


Aldridge, D., 2011. Spinycheek Crayfish, *Orconectes limosus*. Sand Hutton, UK: GB Non-native Species Secretariat, https://doi.org/10.1079/cabicompendium.72033.

Anastácio, P.M., Correia, A.M. & Menino, J.P., 2005, 'Processes and patterns of plant destruction by crayfish: effects of crayfish size and developmental stages of rice', *Archiv für Hydrobiologie* 162, 37–51, https://doi.org/10.1127/0003-9136/2005/0162-0037.

Anda, P., del Pozo, J.S., García, J.M.D., Escudero, R., Peña, F.J.G., Velasco, M.C.L., Sellek, R.E., Chillarón, M.R.J., Serrano, L.P.S. & Navarro J.F.M., 2001, 'Waterborne Outbreak of Tularemia Associated with Crayfish Fishing', *Emerging Infectious Diseases*, 7, 575–82¸ https://doi.org/10.3201/eid0707.010740.

Douthwaite, R., Jones, E., Tyser, A. & Vrdoljak, S., 2018, 'The introduction, spread and ecology of redclaw crayfish *Cherax quadricarinatus* in the Zambezi catchment', *African Journal of Aquatic Science* 43, 353–366, https://doi.org/10.2989/16085914.2018.1517080.

Gherardi, F., Britton, J.R., Mavuti, K.M., Pacini, N., Grey, J., Tricarico, E. & Harper, D.M., 2011b, 'A review of allodiversity in Lake Naivasha, Kenya: Developing conservation actions to protect East African lakes from the negative impacts of alien species', *Biological Conservation*, 144, 2585–2596, https://doi.org/10.1016/j.biocon.2011.07.020.

GISD, 2023. Species profile: *Orconectes virilis*. Global Invasive Species Database, http://www.iucngisd.org/gisd/species.php?sc=218.

Haubrock, P.J., Oficialdegui, F.J., Zeng, Y., Patoka, J., Yeo, D.C.J. & Kouba, A. 2021, 'The redclaw crayfish: A prominent aquaculture species with invasive potential in tropical and subtropical biodiversity hotspots', *Reviews in Aquaculture* 13, 1488–1530, https://doi.org/10.1111/raq.12531.

Holdich, D.M., Reynolds, J.D., Souty-Grosset, C. & Sibley, P.J., 2009, 'A review of the ever-increasing threat to European crayfish from non-indigenous crayfish species', Knowledge and Management of Aquatic Ecosystems, 11, 1–46, https://doi.org/10.1051/kmae/2009025.

Keller, R.P., Frang, K. & Lodge, D.M., 2008, 'Preventing the spread of invasive species: Economic benefits of intervention guided by ecological predictions', *Conservation Biology*, 22, 80–88, https://doi.org/10.1111/j.1523-1739.2007.00811.

Kouba, A., Oficialdegui, F.J., Cuthbert, R.N., Kourantidou, M., South, J., Tricarico, E., Gozlan, R.E., Courchamp, F. and Haubrock, P.J., 2022, 'Identifying economic costs and knowledge gaps of invasive aquatic crustaceans', *Science of the Total Environment*, *813*, p.152325, https://doi.org/10.1016/j.scitotenv.2021.152325.

Lane, M.A., Barsanti, M.C., Santos, C.A., Yeung, M., Lubner, S.J. & Weil, G.J., 2009, 'Human paragonimiasis in North America following ingestion of raw crayfish', *Clinical Infectious Diseases*, 49, 55–61, https://doi.org/10.1086/605534.

Laurent, P.J., 1988, '*Austropotamobius pallipes* and *A. torrentium*, with observations on their interactions with other species in Europe. In: *Biology of freshwater crayfish: biology, management and exploitation*, [ed. by Holdich, D.M., Lowery, R.S]. London, UK: Croom Helm. 341–364.

Madzivanzira, T.C., Chakandinakira, A.T., Mungenge, C.P., O'Brien, G., Dalu, T. & South, J., 2023, 'Get it before it gets to my catch: misdirection traps to mitigate against socioeconomic impacts associated with crayfish invasion', *Management of Biological Invasions* 14(2), 335–346, https://doi.org/10.3391/mbi.2023.14.2.10.

Madzivanzira, T.C., Weyl, O.L.F. & South, J., 2022, 'Ecological and potential socioeconomic impacts of two globally invasive crayfish', *NeoBiota* 72, 25–43, https://doi.org/10.3897/neobiota.72.71868.

Nemesis, 2023. *Faxonius virilis*, Crustaceans-Crayfish: virile crayfish. Edgewater, MD: National Estuarine and Marine Exotic Species Information System (NEMESIS), Smithsonian Environmental Research Center. https://invasions.si.edu/nemesis/species_summary/97425.

Peay, S., Guthrie, N., Spees, J., Nilsson, E. & Bradley, P., 2009, 'The impact of signal crayfish (*Pacifastacus leniusculus*) on the recruitment of salmonoidfish in a headwater stream in Yorkshire, England', *Knowledge and Management of Aquatic Ecosystems*, 394–395, https://doi.org/10.1051/kmae/2010003.

Putra, M.D., Bláha, M., Wardiatno, Y., Krisanti, M., Yonvitner, Jerikho, R., Kamal, M.M., Mojžišová, M., Bystřický, P.K., Kouba, A. & Kalous, L., 2018, '*Procambarus clarkii* (Girard, 1852) and crayfish plague as new threats for biodiversity in Indonesia', *Aquatic Conservation: Marine and Freshwater Ecosystems*, *28*, 1434–1440, https://doi.org/10.1002/aqc.2970.

Sara, S. & El Moutaouakil, M.E., 2019, 'Study on the Spread of *Procambarus clarkii* at Gharb (Morocco) and its impact on rice growing', *Journal of Agriculture, Science and Technology*, 9, 81–92, https://doi.org/10.17265/2161-6256/2019.02.002.

Souty-Grosset, C, Manuel P, Aquiloni L, Banha F, Choquer J, Chucholl C, Tricarico E. 2016. The red swamp crayfish *Procambarus clarkii* in Europe : Impacts on aquatic ecosystems and human well-being *Limnologica* 58: 78–93, https://doi.org/10.1016/j.limno.2016.03.003.

Vega-Villasante F, Ávalos-Aguilar J, Nolasco-Soria H, Vargas-Ceballos M, Bortolini-Rosales J, Chong-Carrillo O, Ruiz-Núñez M, Morales-Hernández J. Wild populations of the invasive Australian red claw crayfish Cherax quadricarinatus (Crustacea, Decapoda) near the northern coast of Jalisco, Mexico: a new fishing and profitable resource. *Lat. Am. J. Aquat. Res.*. 2017;43(4): 781-785. https://doi.org/10.3856/vol43-issue4-fulltext-17.
